# Environmental features or interspecific relations: What determines distribution of Syrian and Great Spotted woodpeckers hybrids?

**DOI:** 10.64898/2026.09.17.752445

**Authors:** Antoni Bakai, Emilia Grzędzicka, Bartłomiej Kusal, Zuzanna Pochłopień, Tomasz Figarski, Gerard Gorman, Łukasz Kajtoch

**Affiliations:** Institute of Systematics and Evolution of Animals Polish Academy of Sciences, Sławkowska 17 31-016, Kraków, Poland; Department of Forest Sciences University of Lodz Branch in Tomaszów Mazowiecki Konstytucji 3 Maja 65/67, 97−200 Tomaszów Mazowiecki, Poland; Hungarian Woodpecker Group, Hungarian Ornithological and Nature Conservation Society, Budapest, Hungary

**Keywords:** Dendrocopos major, Dendrocopos syriacus, hybrids, habitat selection, urban greenery, orchards, Sturnus vulgaris

## Abstract

Urban and rural woodlands have become important habitats for wildlife, providing opportunities for ecological and ethological adaptation to changing environmental conditions. Due to limited tree resources in anthropogenic woodlands, different species are often forced to interact more closely, and one possible outcome of such interactions is hybridisation. Recent studies have shown that hybrids between the uncommon Syrian Woodpecker (*Dendrocopos major*) and the often common Great Spotted Woodpecker (*Dendrocopos syriacus*) are relatively widespread, yet rare, in sympatric populations (Bakai et al. 2026). To understand the factors determining their occurrence in rural and urban landscapes, we conducted an intensive survey across Europe. We examined the effects of environmental factors, particularly woodland characteristics, as well as the co-occurrence of hybrids with the parental species and European Starlings (*Sturnus vulgaris*) (hereafter Starling). Spatial analyses revealed that the parental species generally have allotopic distributions, with Syrian Woodpeckers (hereafter SW) being more common in scattered woodlands in both rural and urban landscapes, while Great Spotted Woodpeckers (hereafter GW) are more commonly associated with denser woodlands and forests. Although in some areas the two species can co-occur, which provides opportunities for interspecific hybridisation, hybrid offspring showed no clear preference for any particular woodland type. However, they tended to occur in proximity to one of the parental species, usually the species that was locally rarer. Starlings were not identified as a factor influencing hybrid distribution. These results suggest that hybrids have a relatively broad ecological tolerance, potentially facilitated by heterosis. These findings contribute to our understanding of the factors influencing interspecific hybridisation and may have implications for the management of urban and rural woodlands to preserve habitats for the SW, an uncommon and strictly protected species in the west of the European Union.

## Introduction

Wooded areas in cities and agricultural landscapes have become important habitats for animals (Gentili et al. 2024, Feber et al. 2025). Despite their often limited extent, they can have mature and structurally diverse trees that provide refugia for arboreal taxa and support species-rich bird communities (Fröhlich et al. 2022). The distribution and persistence of species in urban and rural green spaces can present challenges, but these environments may also provide opportunities for ecological and behavioural adaptation to changing environmental conditions (Minias & Janiszewski 2016). Due to the usually limited availability and spatial concentration of tree resources in anthropogenic woodlands, some species can interact more often than they do in natural habitats. Although such interactions are predominantly competitive, interactions between closely related species may also facilitate interspecific mating and hybridisation.

Among arboreal bird species, woodpeckers (Picidae) are noteworthy in the context of interspecific hybridisation as this phenomenon is relatively widespread amongst them (Ottenburghs & Nicolaï 2024). However, in most of the known hybridising species pairs, the occurrence of hybrids is poorly known owing to data being restricted to occasional observations (Bakai et al. 2026a). The exceptions are the New World *Sphyrapicus* supsuckers (Billerman et al. 2019, Natola et al. 2021), *Colaptes* flickers (Aguillon et al. 2022) and *Melanerpes* woodpeckers (Barrowclough et al. 2017) and the Sri Lankan *Dinopium* flamebacks (Ranasinghe et al. 2024). The only pair of picid species that regularly hybridise in Europe are GW and SW. This pair of species is particularly interesting in regard to the study of related impacts in urban and rural woodlands, as SW is strongly synanthropic (Gorman 1997).

Hybridisation between SW and the GW has been known since the SW began to expand its range from the Middle East into Europe, where it came into contact with its congener (Reiser 1894, Glutz von Blotzheim & Bauer 1980, Dudzik & Polakowski 2011, Michalczuk 2014). This phenomenon has been mainly studied in Poland (e.g. Michalczuk et al. 2014, Figarski & Kajtoch 2018a, Gurgul et al.2019), with very little information available from other regions, even those where hybrids have also been observed (Kroneisl-Ruckner 1957, Winkler 1971). A recent review of the literature on *Dendrocopos* hybridisation (Bakai et al. 2026a,b) and data collected in citizen science projects, documented hybrid frequency in areas where the two species co-occur (Bakai et al. 2025). Moreover, nearly all previous studies have focused on interspecific ecological and ethological differences between pure SWs and GWs, although these aspects have not yet been investigated in their hybrids.

Similarly to other actively hybridising woodpecker species, SW × GW hybrids are able to successfully breed with their parental species (Dudzik & Polakowski 2011, authors’ own data), as well as together (Kajtoch & Kusal 2022). This results in multiple generations of “backcrossed” birds with gene introgression in parental species (Grugul et al. 2019). According to a previous study (Bakai et al. 2025), hybridisation between SW and GW woodpeckers is categorised by a hybrid swarm, where hybrids reproduce spontaneously across the whole sympatric range, rather than in distinct hybrid zones. This might indicate that hybrid occurrence is more likely explained by local environmentalal factors or interspecific interactions than geographical gradients.

The respective biology of GW and SW are well known (Gorman 1997). GW tends to occupy tree stands of a larger area and with dense vegetation, while SW prefers fragmented woodlands with fruit and softwood trees. In anthropogenic environments such as rural and urban greenery, both species can, however, occupy similar habitats (Michalczuk & Michalczuk 2016a, Figarski & Kajtoch 2018b) or a mosaic of habitats which allows for frequent contact and hence interspecific occasional interbreeding. Nevertheless, it should be emphasised that most of the relevant study on this originates from Poland with few data from other populations. Despite a rather large body of knowledge on the biology of the two parental species and also comparative ecological studies existing (Michalczuk & Michalczuk 2016b, Kajtoch & Figarski 2017, Figarski & Kajtoch 2018b, Fröhlich & Ciach 2013), information on their hybrids is relatively scarce, and they are often omitted from faunistical and ecological studies (Bakai et al. 2026b). Therefore which environmental factors (for example, which woodland types, if any, are favoured) may promote hybridisation between these woodpeckers is unknown Apart from environmental characteristics and the habitat preferences of the two species, which could be intermediate in hybrids, interspecific interactions may also play a role in the formation and occurrence of hybrids. When considering such interactions, the most obvious assumption is that the parental species themselves influence the occurrence of their hybrids. However, other taxa are also likely to contribute to this process, for instance, through competition for nesting sites and/or food resources. One of the well-known competitors for woodpecker cavities is the Starling, a bird which will occupy those of both SW and GW (Winkler 1973; Mazgajski 2000; Smith 2005).

In this study we aim to examine the habitat requirements of hybrids in relation to their parental species and other taxa in synanthropic woodlands in order to try to understand what determines the occurrence of hybrids in woodpecker populations.

Subsequently, we tested the following hypotheses:

1. The habitat preferences of GW and SW in anthropogenic landscapes documented in Polish populations, appear to be consistent across Europe. Specifically, GW tends to occupy larger and denser woodlands, whereas SW is more frequently associated with scattered woodlands and secondary habitats.
2. Hybrids occupy landscapes that are intermediate between, or combine characteristics of, those inhabited by the parental species. Specifically, hybrids are most frequently found in transitional zones between dense and scattered woodlands, where SW and GW may locally nest in close proximity to one another. This forces hybrids to breed in close proximity to their parental species, typically near the territories of the locally less abundant species.
3. Urban and rural woodlands contribute more to the occurrence of hybrids than the co-occurrence of their parental species and other taxa.

## Methods

### 2.1 Study area

The study was conducted on two levels - European and Lesser Poland (part of the SE Poland). The European level included data from Poland, Slovakia, Hungary, Austria, Czech, Germany, Ukraine, Romania, Bulgaria and Greece. Two urban and two rural transects were established in 55 cities across Central and SE Europe (within the range of SW and in adjacent areas) (see Bakai et al. 2025). Distances between the cities (urban and rural transects) were around 80 km, forming a uniform grid across the studied geographical range (Fig S1).

The Lesser Poland level included Krakow and 16 surrounding villages and small towns up to 50 km from this city. Here, counts were conducted on 16 urban, 16 suburban, and 16 rural transects.

### 2.2 Field data collection

The records of woodpeckers were obtained by counting along transects. Transects were planned along presumed suitable habitat for both species and their hybrids (based on data from previous surveys and records of SW and hybrids from online databases like eBird, iNaturalist and ornitho.pl/ornitho.at), namely urban and rural green areas excluding dense continuous forests where SW is seldom present, or treeless areas where no woodpeckers at all could be found. Transects were 1.5 km in length, with the exception of transects from Lesser Poland that were 2.4 km long. Survey points were situated 300m apart along the transect, making five survey points for shorter and eight for longer transects.

At each survey point playback of calls and drums was used, following the inventory method of SW in Poland (Michalczuk & Michalczuk 2006), as both species are known to actively respond to each others’ vocalisations and drums (Figarski 2017). Any woodpeckers that appeared were observed with binoculars for at least five minutes and, when possible, photographs were taken. Woodpeckers were identified based on their appearance and vocalisations (Gorman 1997, Gorman & Kajtoch 2026). Additionally, any Starlings present, and whether they were singing and/or nesting, noted.

Counts were conducted in 2023-2025 during the breeding season (mid-March to mid-April), when woodpeckers were occupying territories and their activity and response to playback was highest. Two surveys were done at each site, with at least a two-week interval between them. Additionally, on transects where hybrids or mixed pairs were found, a third count was done, in mid-May to mid-June, to search for cavities and nestlings.

### 2.3 Habitat features

In the European level analyses, a study plot formed by an area of a 150 m radius around each count point was established. Inside each plot areas of both tree canopy and infrastructure (buildings, railroads, highways) were calculated. Additionally, the presence of other particular habitat features were noted (Tab S1, S2, S3).

For the analysis on a regional level, Lesser Poland was chosen owing to the studies on SW and hybrids that had already been carried out there in recent years (Kajtoch & Figarski 2017). In this region, apart from features used on the European level, the canopy area was additionally divided into three categories: managed woodland in residential areas (managed park areas, cemeteries, residential greenery), orchards and gardens and wild or semi-natural vegetation (abandoned tree stands, urban or rural forests) (Tab S1, S3). Additionally, distances between three types of woodland were calculated in a buffer zone of 300 m around these plots. These distances were used to assess if hybrid occurrence is associated with the proximity of different woodland types, which are suitable for their parental species (e.g. orchards for SW, forests for GW).

All measurements were conducted on 2024-2025 Google satellite maps (google.com/maps) in QGIS v 3.34. The infrastructure area was measured based on QGIS quick Open Street Maps layers (https://quickosm.github.io/QuickOSM/user-guide/map-preset/), requesting “buildings” objects, with additional manually made polygons where the objects did not reflect local infrastructure correctly (mainly in S Europe rural areas). As there were no suitable layers for green areas, all polygons for area computation were made manually based on satellite images and field observations.

Apart from environmental factors and co-occurrence with parental species, hybrid distribution could also be shaped by competitive species like other woodpeckers and secondary cavity nesters. Apart from parental species (SW and GW), the Starling can be highly competitive with hybrids, too. This species often usurps nest cavities excavated by GW and SW, even these that are occupied (Winkler 1973, Mazgajski 2000, Smith 2005).

### 2.4 Statistical analyses

All statistical analyses were prepared using R (v 4.5.1; R Core Team 2025) and R-studio (Posit team 2024).

To test the ecological relationships between woodpeckers and their habitats, data was treated on two landscape levels. Considering the whole dataset collected in Europe, 55 pairs of urban and 55 pairs of rural transects were examined. Data from 12 points from Kraków and nearby villages where hybrids were recorded was added to this analysis. For examining the regional population in Lesser Poland, we considered 16 rural, 16 suburban, and 16 urban transects. For statistical tests with multiple environmental predictors (Tab S1), we conducted a constrained ordination analysis in vegan package to investigate the trends in habitat selection for woodpeckers (Oksanen et al. 2026). After confirming the linearity of data with DCA (DC1 < 3), tbRDA on Hellinger-transformed count data was made to test GW, SW and also hybrid responses to predictors. On the European level analysis, the following habitat features were used as a predictors: canopy cover area, infrastructure area; presence/abscense of dense canopy, loose canopy, coniferous trees, forest patches, urban parks, graveyards, wooded lanes, residential greenery, orchards, riparian woodlands, small buildings with sporadic distribution, small buildings in dense aggregations, tall buildings with sporadic distribution, tall buildings in dense aggregations and other infrastructure (Tab S1). On the Lesser Poland level, instead of total canopy cover area and the presence/absence of orchards, forest patches, urban parks, wooded lanes, graveyards and residential greenery, the area of orchards, managed greenery area and unmanaged woodland areas were used (Tab S1). As woodpecker relationships with resources and habitat features may be essentially different in rural and urban environments, we conducted unconstrained linear ordination analysis (PCA) in factoMineR package (Lê et al. 2008), to use the first two principal components as landscape type characteristics. To increase the explained variance, the habitat features whose contributions to the main axes were less than 10% were removed and new principal components generated.

Because bird count data (GW, SW, hybrids) had poisson distribution, for European level data, Generalised Linear Mixed Models (GLMMs) were generated using the package glmmTMB (Brooks et al. 2017). On the Lesser Poland level, Generalised Linear Models were generated without random factors, both having a log link function to fit the model. Unidentified woodpecker records were removed from the analysis to avoid type I errors. To test habitat preferences for GW, SW and their hybrids, together with possible interspecific interactions effects, models including PC1 and PC2 together with presence of other woodpeckers (GW, SW, H) and Starlings as predictors, were generated. In order to examine more specifically the particular habitat features that influenced the woodpeckers, models with all the discrete habitat features present on a corresponding level as predictors were generated. Due to the large geographical scope of the study on the European level, we used transects as a random factor, to account for possible geographical variability. Models were then placed in the MumMIn package (Bartoń 2026). These models were later tested for autocorrelation with a Moran I test, in the DHARMa package (Hartig 2026), and if the autocorrelation test was positive, a Matern function in spaMM package was used (Rousset & Ferdy 2014).

To examine woodpeckers distribution patterns SADIE analyses were performed in epiphy package on the Central European and Lesser Poland levels. We focused on transects only from Central Europe, as transects from Greece, Bulgaria, Romania and Eastern Ukraine were situated along narrow patches of habitat, which meant that it was not possible to extrapolate geographic data into the whole Balkans or Ukraine. Kernel density graphs were made with statstat and sf packages for GW, SW and H on both levels to visualise distribution patterns. To examine the association between GW, SW and H, the SADIEs of each were compared, and association indexes between each pair of taxa calculated (1 = positive association, 1 = avoidance, close to 0 = no effect).

To test whether differences in distances between different tree stand types in study plots have an effect on hybrid occurrence, GLMs were generated, testing if the length of the shortest distance between two of three different tree stand types had a significant effect on hybrid presence. Kernel Density graphs depicting the frequency of hybrid presence in distance between two different tree stand types (Managed-Orchard, Managed-Wild, Orchard-Wild) were made, with distances reaching outside the buffer range (300 m) marked as 500 m. Graphs were also made for GW and SW, to check for possible bias.

For data visualisation, the ggplot2 (Wickham 2016), sjPlot (Lüdecke 2025), factoextra (Kassambara & Mundt 2026), epiphy (Gigot 2023), spatstat (Baddeley et al. 2015) and sf (Pebesma & Bivand 2023) packages were used.

## Results

### 3.1 Bird counts

A total of 220 transects containing 1100 counting points with birds count and environmental data were conducted in ten countries (Tab S2, S3). The number of transects in each country were as follows: Poland 72, Slovakia 28, Czech 24, Bulgaria 20, Romania 20, Hungary 20, Ukraine 16, Austria 12, Germany 8, Greece 4.

In addition, in Lesser Poland a dense sampling was carried out over 16 urban, 16 suburban and 16 rural transects (384 point surveys in total).

Hybrids were recorded at 60 points. In summary, we obtained records of 947 GW, 310 SW, and 68 hybrids.

### 3.2 Environment characteristics

The PCA conducted on all studied environmental features on the European level showed a low variance (PC1=19.8%, PC2=12.2%): 7 features contributing less than 10% to the axes were removed (Fig S2). The PCA on the remaining features comprised 50.5% of the total variance and showed two distinct gradients of woodpecker occupied environments (Fig 1). PC1 corresponded to the gradient between wild unmanaged green areas with larger areas of dense vegetation and little infrastructure to built-up areas with a managed loose canopy. PC2 corresponded to the gradient between sporadic rural greenery with orchards and small buildings to dense greenery in urban parks and residential greenery with tall buildings. Both principal components indicated a transition from rural to urban landscape.

**Figure 1.**
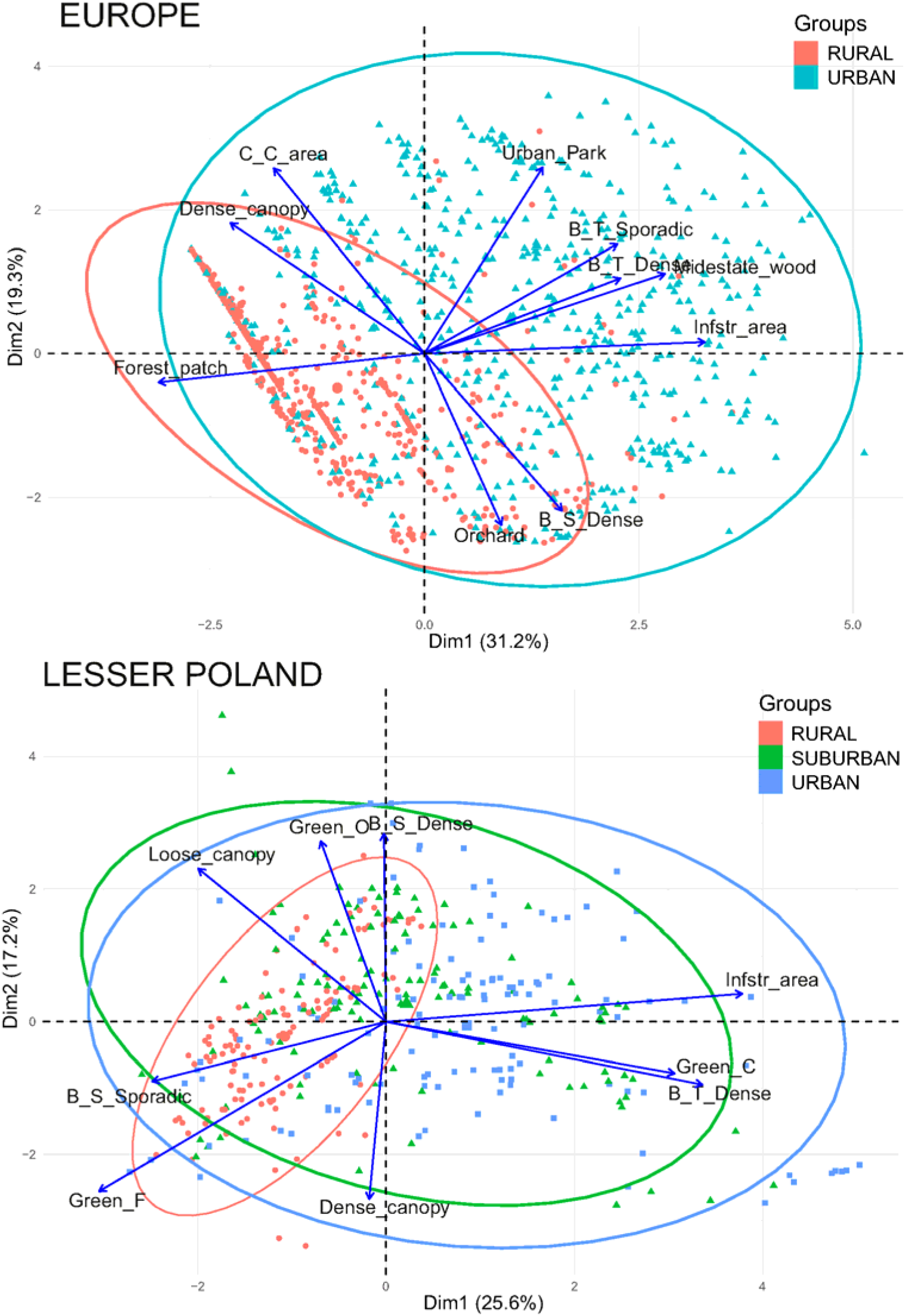
Principal Component Analysis biplot showing the gradient of environmental variables present within the proximity of 150 m of woodpecker count points along the first two principal components at a European and SE Poland level. The count points are grouped depending on whether their landscape type was rural (R), suburban (S) or urban (U). The photos represent typical rural and urban type habitats described by lower-left and upper-right quadrants.

Following the same procedures, PCA on Local level represented 42.8 % of total variance (Fig 1). The first axis PC1 corresponded to typical urban and rural greenery in Lesser Poland, while the second PC2 corresponded to the gradient between dense unmanaged vegetation and residential greenery with gardens and sporadic trees scattered along roads.

### 3.3 Woodpecker habitat use and interspecies relationships

The tbRDA on Hellinger-transformed count data indicates less than 15% of variance on both levels (Fig S3, Tab S4), therefore interpretation requires caution. Nevertheless, it shows distinct trends for both parental species, and a lack of any pattern for hybrids on both levels (Fig 2). There are clear positive effects of canopy cover area and density of canopy on GW presence, while SW is attracted to orchards and the like with dense rural housing and other elements of open rural landscapes. In the meantime, both species seemed to benefit from riparian vegetation and patches of unmanaged greenery, though avoiding large build-up areas. Taking in account the PCA results on habitat structure, there reasons to believe that both species occupy different habitats even in the same environment are evident. Hybrids, on the other hand, do not seem to have any particular habitat preference according to this redundancy analysis.

**Figure 2.**
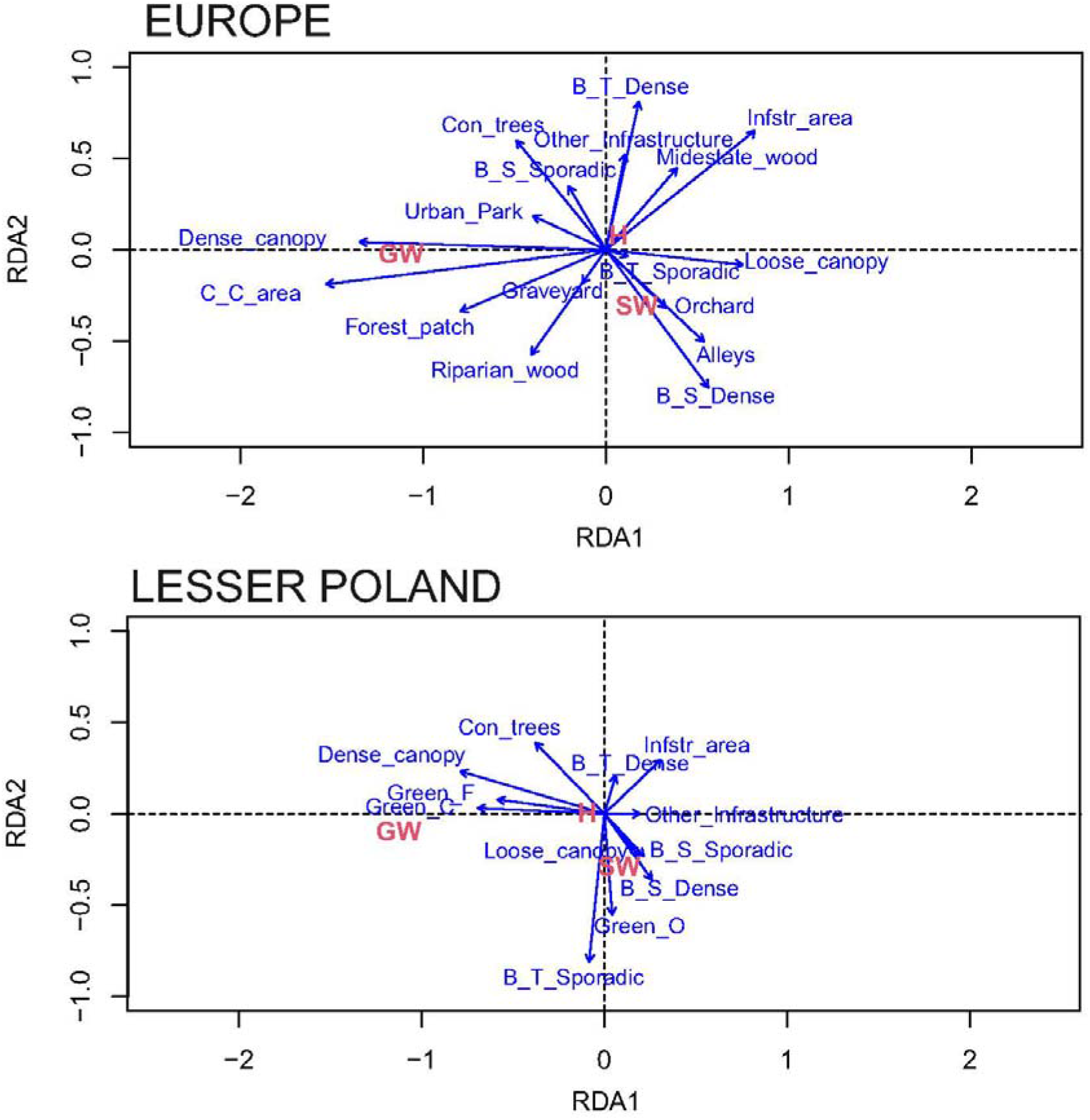
Transformation-based Redundancy Analysis (tbRDA) ordination plots depicting Great Spotted Woodpecker, Syrian Woodpecker and their hybrids between two habitat preferences over two first constrained axes: RDA1 and RDA2 at a European and Local level. The relationship of particular habitat elements to ordination axes are shown with blue arrows.

On the European level, the GLMMs testing relation of woodpeckers with other species in the environment represented by principal components (Tab 1) showed a negative association for GW with both PC1 (tendency towards larger patches of dense semi-natural vegetation instead of anthropogenic environment with managed greenery and buildings) and PC2 (tendency towards landscape with dense residential greenery with orchards and gardens instead of urban landscapes with managed parks and tall infrastructure). At the same time GW has a positive association with hybrids and Starlings. Similar models did not show significant effects of the environment or Starling presence neither for SW nor hybrids. However, both SW and hybrid models showed significant positive associations between these birds. A significant negative autocorrelation was detected in the SW model (p = 0.04), thus the model was adjusted, but significance of the predictors was not changed.

The similar GLMMs on the local level showed a significant negative association between GW and PC2 (preference towards more forest-like habitats while avoiding residential greenery), while positive association with Starlings (Tab 2). The influence of both local level principal components on SW occurrence was not significant. The models testing hybrid occurrence showed a significant negative association of the latter with PC1 (tendency towards semi-natural tree stands among arable fields and villages with scattered single houses instead of managed urban greenery and dense urban infrastructure). Both SW and hybrid models indicated a significant positive association between these birds, while the presence of GW and Starlings has no significant effect on them. The significant positive autocorrelation was indicated in the GW model (p = 0.008), and after adjusting general effects was the same.

The European level GLMMs with the discrete habitat elements as predictors also showed a significant positive influence of some urban greenery and large dense tree stands on GW presence and abundance. In the case of SW, the model confirmed only significant positive effects of dense rural buildings aggregations and negative effects of forest and other unmanaged greenery patches. A Moran I test detected a significant autocorrelation in GW (p = 0.035) and SW (p = 0.031) models, and after fitting with geographical coordinates, the models showed a significant positive influence of coniferous trees on GW and significant negative effect of coniferous trees on SW. The model with hybrids as a dependent variable revealed a significant positive effect of dense canopy presence at nesting sites. The summaries of the adjusted optimal models are present in Tab 3. The summaries of initial models with all predictors are in the supplemental Tab S4.

The Lesser Poland level GLMMs showed a significant positive effect of managed and unmanaged trees canopy area on GW presence and abundance. SW, on the other hand, showed a preference for landscapes with sporadic rural houses and an avoidance of coniferous trees. Hybrids had a positive relationship with dense vegetation. No significant autocorrelation was detected in local level models. The summaries of the optimal models are present in Tab 4. The summaries of initial models with all predictors are in the supplemental Tab S5.

The GLMMs analysis on hybrid presence (data type: binomial, link function: logit) and a relationship with the shortest distances between different types of tree stands (Tab S1) on the Lesser Poland level, did not show any significant effects (Estimate <0.001, p = 0.941). The density plots showed a peak of hybrid appearance in the study plots where the closest distance between two different tree stand types was around 50 m, but also a second peak in plots where no other tree stand type was present (Fig S4). A similar picture was observed on density plots where both GW and SW are present (Fig S4).

### 3.4 Woodpecker spatial distribution and interspecies relationships

The European level SADIE analyses for GW, SW and hybrid spatial distribution showed a significant clustering of both species but not of hybrids. GWs were mainly aggregated in the west and north-west, while SW clusters were concentrated in the south-east (Tab 5, Fig 3). On the Lesser Poland level, hybrids and SW showed a pattern of significant clustering, while GWs were distributed randomly (Tab 5, Fig 4). No differences were observed between rural and urban transects on the European level (Fig S5), however, on the local level more clear species distribution patterns were seen (Fig 4).

**Figure 3.**
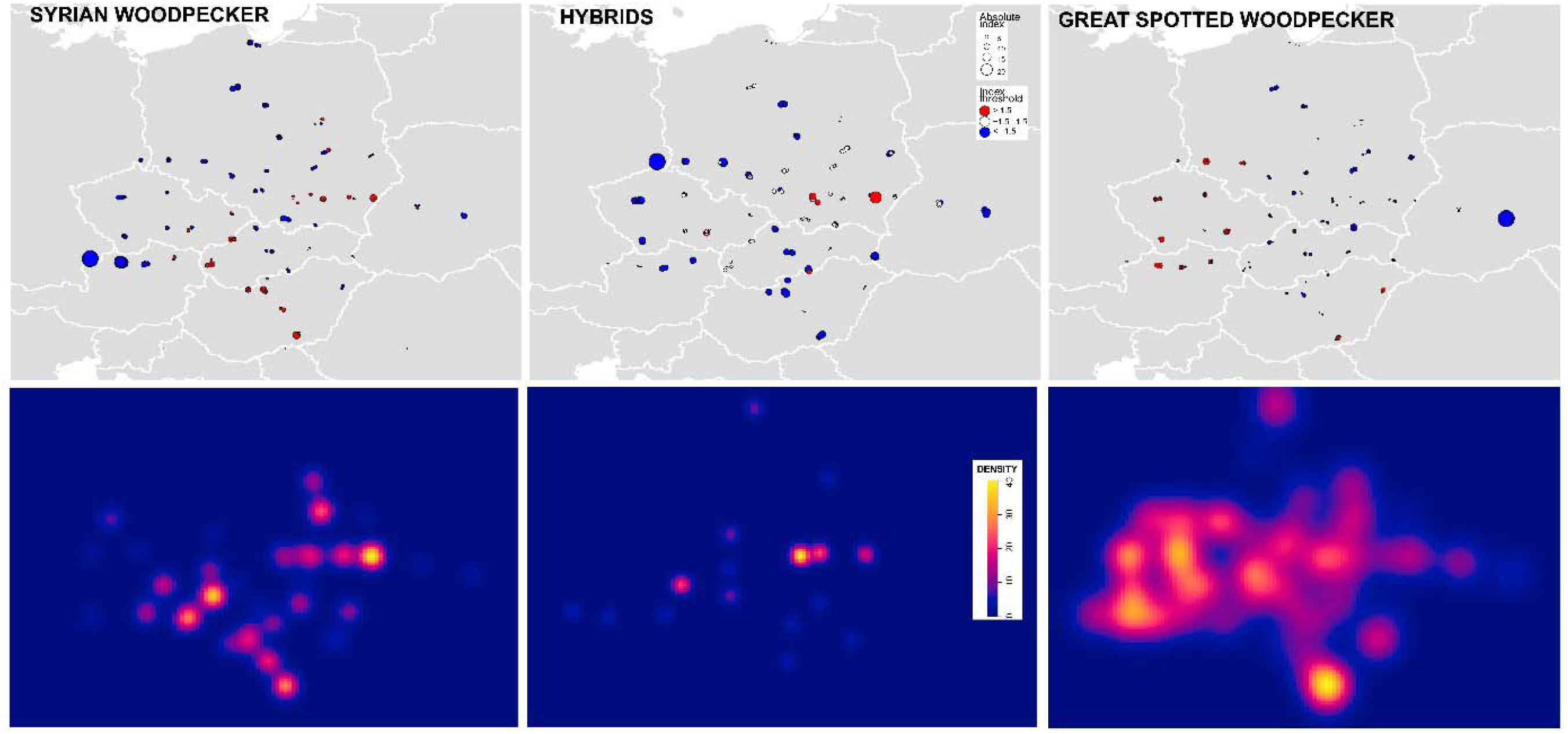
Graphical visualisation of Great Spotted Woodpecker, Syrian Woodpecker and their hybrids distribution in Central Europe. SADIE red & blue plots (top) indicate the number of woodpeckers on the counting points above (red), below (blue) and average (white). Kernel Density Estimation pixel plots (bottom) are based on each single bird record.

**Figure 4.**
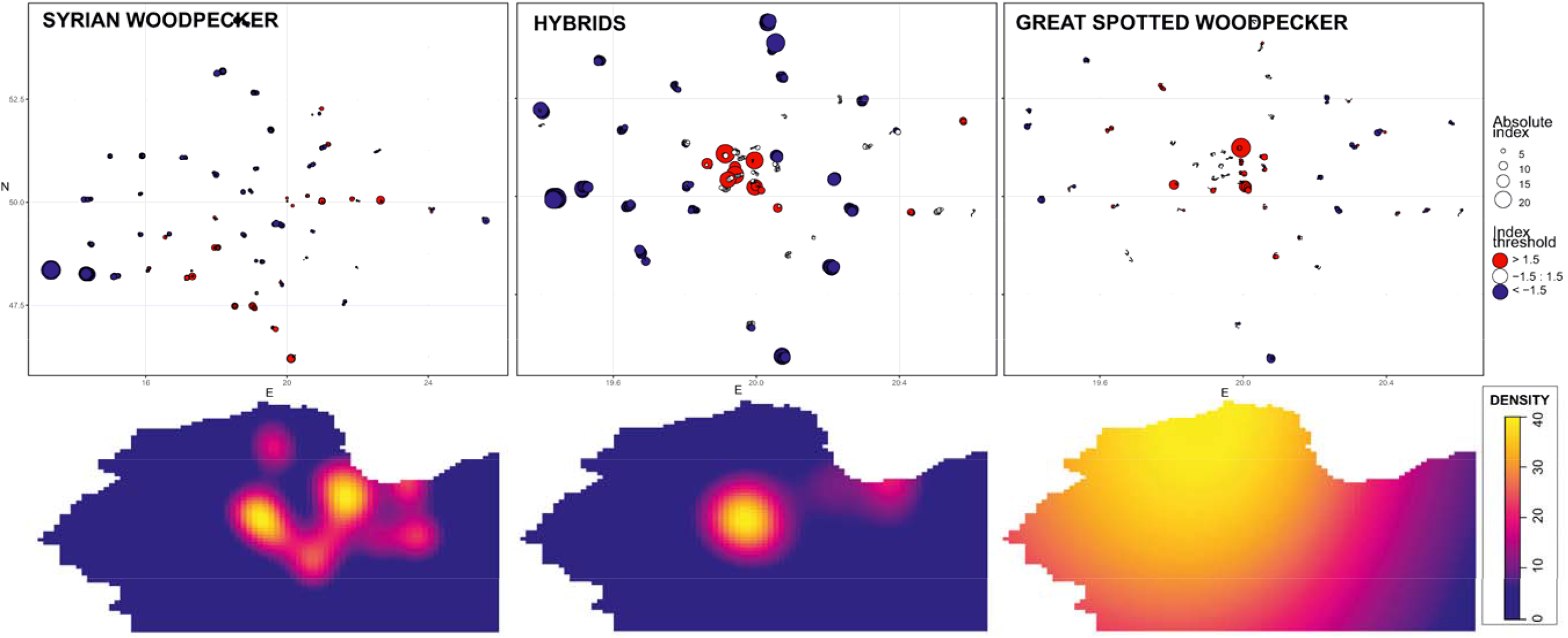
Graphical visualisation of Great Spotted Woodpecker, Syrian Woodpecker and their hybrids distribution in SE Poland. SADIE red & blue plots (top) indicate the number of woodpeckers on the counting points above (red), below (blue) and average (white). Kernel Density Estimation pixel plots (bottom) based on each single bird record are grouped according to landscape type (Rural, Suburban, Urban).

The correlation analyses on the both levels did not show significant negative or positive association indexes (Tab 6).

## Discussion

The major finding of this study was that the widespread distribution of SW x GW hybrids is accompanied by a broad ecological tolerance. Therefore, the overall scarcity of hybrids is unlikely to be caused by environmental factors, but rather by limited interspecific breeding, which possibly results from differences in the behaviour of the two species (Figarski 2017), rather than from differences in their reproductive biology (Kajtoch & Kusal 2022).

### SW and GW habitat preferences

Landscape factors shaping the distribution of SW and GW are consistent with the current knowledge of their ecology (Gorman 1997) and with the niche segregation previously reported for Polish populations (Figarski & Kajtoch 2018a). However, our analyses have revealed a difference between European and regional level preferences. On the European level GW showed preferences towards rural landscapes with dense semi-natural tree stands and the presence of coniferous trees, while in Lesser Poland GW tended to be indifferent to whether any given tree stands was managed or not. Yet, on both levels GW consistently showed a preference towards larger wooded areas and an avoidance of dense residential greenery with gardens. On the contrary, SWs showed a preference for agricultural landscapes and avoided coniferous tree stands on the European level. Yet, on the Lesser Poland level this species prefers areas of urban housing and a positive association with gardens but avoids built-up areas. The contrast between the preferences of GW and SW on the European and regional levels could be caused by the different landscape structure and elements in Lesser Poland compared to the rest of the European range. Indeed, the contrast to the Pannonian Basin and the Balkans was particularly marked. Our results are in accordance with previous studies on GW in Poland, which described this species as a habitat generalist capable of breeding in virtually any wooded habitat, including both forests, and urban greenery (Mazgajski 1998; Hebda et al. 2017). Although it shows a preference for large, continuous forest complexes, GW readily occupies smaller woodlots and urban woodlands and has expanded considerably across Europe in recent decades (Keller et al. 2020). In more southern parts of Europe, however, GW is still rarely observed in cities, where anthropogenic greenery is dominated by SW (Bakai et al. 2026). The later species is described in the literature as primarily occupying scattered woodlands embedded within urban and agricultural landscapes, showing a preference for tree-lined avenues, orchards and other semi-open wooded habitats (Michalczuk & Michalczuk 2016a, 2020). This habitat preference reflects its evolutionary association with forest-steppe environments in its native range in the Middle East and western Asia (Aghanajafizadeh et al. 2011). The absence of residential areas has had a positive effect on SW in Lesser Poland, which is likely to be a result of some recent changes in landscape use, particularly around villages, where important elements for this species, such as old fruit trees, have been removed (Michalczuk & Michalczuk 2015, Kajtoch & Kusal 2023).

### SW × GW hybrid habitat preferences

According to our analysis, hybrids, unlike parental species, did not appear to be associated with any particular woodland type. Hybrid individuals were recorded in a wide variety of wooded habitats in both urban and rural landscapes, suggesting a broader ecological tolerance than that of both parental species. The only significant habitat variable that predicted hybrid occurrence was the presence of dense tree stand patches at nesting sites. Why hybrids would prefer this type of vegetation is unclear, although dense vegetation might offer better foraging opportunities than scattered trees. A preference for managed tree stands in Lesser Poland may be a result of available habitat niche in that region, however, it may be due to a higher proportion of GW genes in hybrid genomes in this area (Gurgul et al. 2019, ongoing study). Indeed, most of the hybrid woodpeckers in the north of Europe show a GW-like phenotype (Bakai et al. 2025). At the same time, when pairing with SW, hybrids easily enter typical Lesser Poland agriculture landscape with scattered unmanaged vegetation and a few, scattered infrastructure.

### Ecotone habitats and hybrid genesis

Our analysis did not confirm the hypothesis that hybrid tend to occur where the contrasting habitats of their parental species are found, for example where urban parks adjoin allotmenta and gardens or where forests border orchards and farmland woodlots. The pattern shown by the Kernel density graphs for both hybrids and parental species reflects overall habitat structure rather than any ecological preferences of the studied birds. Even if hybrid origin is associated with such ecotones, the high plasticity and dispersability of these birds make them unlikely to remain in the places where they hatched. Thus, the location where an adult hybrid is observed does not necessarily correspond to its natal site, as dispersing individuals may subsequently establish territories in a variety of other habitats. Consequently, the observed habitat associations of adult hybrids does not reflect the environments in which hybridisation originally occurred. To study the origins of hybrid woodpeckers, the characteristics of mixed pair nesting habitat, and of those pairs raising young which originated from extrapair interspecific copulations, should be examined. The mechanism of hybrid bird origin has been considered the key source of Green Woodpecker (*Picus viridis*) × Grey-headed Woodpecker (*Picus canus*) hybrids (Friedmann et al. 2011, Ławicki et al. 2015). Consequently, hybrid origins cannot be interpreted solely on the basis of mixed breeding pair distribution. Quantifying the relative contribution of social pairing and extra-pair fertilisation to hybrid formation remains an important challenge for future behavioural and genomic studies. Unfortunately, to date, there are too few records of mixed pairs and none of any extra-pair mixed offspring to conduct such an analysis (Bakai et al. 2025, 2026b). Different study designs would be needed to collect suitable data.

### Woodpecker spatial distribution and interspecies relationships

The contrasting habitat preferences result in largely allotopic distributions, although local overlap of territories creates opportunities for interspecific pair formation and extra-pair copulations (Figarski & Kajtoch 2018b). Still, our spatial analyses neither indicated positive nor negative associations between these woodpeckers. On the vast geographical level, GW were more aggregated in the westernmost and northernmost parts of the European level study area, whereas their distribution became increasingly dispersed towards the south and east. The opposite pattern was observed for SW, which formed clusters primarily in south-eastern regions and were almost absent from western Europe. This pattern closely mirrors the broader European distributions and regional population densities of both species (Keller et al. 2020, Bakai et al. 2025). Although GWs could be found throughout Lesser Poland, SWs were restricted to a few clusters in Krakow and some villages and small towns to the east of the city. Hybrids were mostly found in the same areas where SW was present, although some hybrid individuals were found west of Krakow, where pure SWs were absent. This could again be explained by the high plasticity and dispersability of hybrids. Indeed, some records were far beyond the known SW range in Europe (Bakai et al. 2025). The same pattern was observed on the European level. The positive association of GW with hybrids might arise from the tendency of hybrids to form pairs with GW.

The significance of Starlings as a negative factor for woodpeckers (Winkler 1973; Mazgajski 2000; Smith 2005) was not confirmed in this study. The Generalised linear model analysis even showed a significant positive association between GWs and Starlings, however, but this might have been due to the opposite effect, as Starlings are attracted to habitats with woodpeckers. Despite previous reports in the literature concerning negative interactions between SWs and Starlings, our GLM analyses did not reveal any significant associations between these species nor between Starlings and hybrids, but this might be due to the methodology used in this study. Here, we examined only the presence of birds (woodpeckers and Starlings), while interactions between these species, including competition for nesting sites, might be more apparent when the availability and occupancy of nesting cavities are examined (Winkler 1973; Mazgajski 2000; Smith 2005).

### Questions, conclusions and possible implications of the study

The broad habitat spectrum occupied by GW X SW hybrids raises interesting questions regarding their respective ecological characteristics. Hybrids were recorded across diverse woodland types spanning both urban and rural landscapes, suggesting greater ecological flexibility than observed in either parental species. Whether this apparent flexibility reflects hybrid vigour (heterosis), complementary combinations of parental traits, increased phenotypic plasticity, or simply the cumulative effects of dispersal and habitat availability, is unknown. Resolving these alternative explanations will require genomic analyses combined with detailed ecological studies of hybrid fitness, habitat selection and reproductive success.

What do these findings imply for the future of the parental species? Given its abundance, hybridisation is unlikely to pose a threat to the GW at the continental level. The European population of GW is estimated at 17.2–27.3 million adults (BirdLife International 2021), the vast majority of which breed entirely allopatrically with SW. The total European population of SW is estimated at approximately 0.3–0.8 million adults (BirdLife International 2021), thus it, too, is sufficiently large that hybridisation is unlikely to threaten the species.

The situation may differ, however, at the periphery of the SW’s distribution in north-central and north-eastern Europe. In those areas populations are relatively small, fragmented and have shown recent declines following decades of rapid expansion during the twentieth century (Michalczuk 2014; Michalczuk & Michalczuk 2015; Keller et al. 2020; Kajtoch & Kusal 2023). The causes of this reversal remain poorly understood. Habitat loss undoubtedly contributes, particularly through the removal of traditional orchards, roadside lines of tree and mature urban trees (Kajtoch 2023). Increasing competition with the expanding GW has also been proposed as a contributing factor (Bakai et al. 2026b). Our findings suggest that hybridisation should now be considered an additional process potentially influencing peripheral populations.

From a conservation perspective, these findings emphasise the importance of the long-term monitoring of hybridisation, particularly within peripheral populations of the SW. Hybridisation itself is a natural evolutionary phenomenon and should not automatically be regarded as detrimental. Instead, effective conservation requires understanding its frequency, spatial dynamics and long-term demographic consequences. Such knowledge can only be achieved by combining field observations, citizen science data (Bakai et al. 2025), long-term monitoring programmes (being developed) and high-resolution genomic analyses (in progress).

Furthermore, this study demonstrates that parks, orchards, wooded lanes and unmanaged tree stands are important for woodpecker ecology. Eco-evolutionary processes can operate in such anthropogenic environments and may even be facilitated by the high structural complexity and heterogeneity of urban and rural greenery, which can promote the co-occurrence of species. Hybridisation in urban and rural woodlands is an interesting phenomenon that deserves further investigation across a broad range of taxa, including plants, fungi and mammals.

## Acknowledgements

This paper was prepared as a part of a grant funded by the National Science Centre, Poland (project number UMO-2022/47/O/NZ9/02044, granted to Ł. Kajtoch). We are grateful to the following for their contributions to field surveys and data sharing: Maciej Aleksandrowicz, Krzysztof Basista, Johanka Bláhová, Marcin Borowik, Stanisław Broński, Jan Čapek, Martin Černý, Tomasz Chodkiewicz, Peter Chrašč, Balázs Csibrány, Krzysztof Czajowski, Krzysztof Czarnocki, Adam Dmoch, Ryszard Dworak, Alena Fišerová, Mariusz Godlewski, Marta Gołek, Paweł Grochowski, Fatima Hayatli, Lucie Hornátová, Tomasz Janiszewski, Aleksandra Janiszewska, Joanna Kawka, Katarzyna Kusal, Mikołaj Krzyżanowski, Sławomir Kuczmarski, Piotr Kłonowski, Władysław Lasoń, Peter Lešo, Maria Madej, Konrad Malec, Marcin Matysek, Szymon Mazgaj, Iryna Miedviedieva, Magdalena Naber, Mateusz Niedziółka, Samuel Pačenovský, Peter Puchala, Patryk Rowiński, András Schmidt, Jerzy Smykla, Tadeusz Sobus, Bartłomiej Stankiewicz, Jan Špička, Daniela Svojanovská, Zbyszek Swiacki, Marta Świtala, Krzysztof Tabernacki, Karolina Tchoń, Matěj Tvarůžka, Lukas Vana, Petr Veselý, Ottó Veszelinov, Łukasz Wardecki, Kristýna Wehrichová, Marta Wołoszyn, Jakub Wyka, Vadym Zhulenko, Piotr Zielinski, Tomasz Ziółkowski, Tomasz Ziółkowski, Antoni Życki, Paweł Żarkiewicz, Karolina Żukowska.

---

**Table 1.** Generalised Linear Mixed Models demonstrating the relationships of Great Spotted Woodpecker, Syrian Woodpecker and their hybrids abundance according to habitat characteristic elements compressed to two main principal components PC1 and PC2 and the coincidence of other woodpeckers and European Starlings (St). PC1 corresponds to the natural environment (dense canopy, canopy area, unmanaged greenery, etc.) versus human transformed environments (buildings, infrastructure area, managed greenery, etc.). PC2 corresponds to semi open rural habitat versus urban habitat and city forests. The SW model was adapted for autocorrelation. The significant P values (<0.05) are highlighted in bold. The analysis was made using European data.

| Predictors | Estimate | SE | Z value | p |
| --- | --- | --- | --- | --- |
| Syrian Woodpecker |  |  |  |  |
| PC1 | -0.009 | 0.049 | -0.192 | 0.847 |
| PC2 | 0.092 | 0.060 | 1.536 | 0.124 |
| GW | -0.091 | 0.097 | -0.934 | 0.350 |
| H | 0.683 | 0.225 | 3.041 | **0.002** |
| St | 0.340 | 0.175 | 1.940 | 0.052 |
| Hybrid |  |  |  |  |
| PC1 | 0.119 | 0.099 | 1.203 | 0.229 |
| PC2 | -0.064 | 0.120 | -0.536 | 0.592 |
| GW | 0.271 | 0.156 | 1.732 | 0.083 |
| SW | 0.502 | 0.184 | 2.724 | **0.006** |
| St | 0.012 | 0.327 | 0.036 | 0.971 |
| Great Spotted Woodpecker |  |  |  |  |
| PC1 | -0.202 | 0.031 | -6.629 | <<**0.001** |
| PC2 | -0.223 | 0.036 | -6.194 | <<**0.001** |
| SW | -0.103 | 0.082 | -1.252 | 0.211 |
| H | 0.315 | 0.149 | 2.120 | **0.034** |
| St | 0.445 | 0.095 | 4.711 | <<**0.001** |

**Table 2.** Generalised Linear Models demonstrating the relationships of Great Spotted Woodpecker, Syrian Woodpecker and their hybrids abundance to habitat characteristic elements compressed to two main principal components PC1 and PC2 with and the coincidence of other woodpeckers and Starlings (St). PC1 corresponds to the natural environment (dense canopy, canopy area, unmanaged greenery, etc.) versus human transformed environment (buildings, infrastructure area, managed greenery). PC2 corresponds to semi open rural habitat versus urban habitat and city forests. The GW model was adapted for autocorrelation. The significant P values (<0.05) are highlighted in bold. The analysis was made using SE Poland data.

| Predictors | Estimate | SE | Z value | p |
| --- | --- | --- | --- | --- |
| Syrian Woodpecker |  |  |  |  |
| PC1 | 0.101 | 0.121 | 0.834 | 0.404 |
| PC2 | 0.194 | 0.123 | 1.570 | 0.116 |
| GW | -0.356 | 0.213 | -1.671 | 0.095 |
| H | 0.758 | 0.245 | 03.paż | **0.002** |
| St | 0.436 | 0.316 | 1.381 | 0.167 |
| Hybrid |  |  |  |  |
| PC1 | -0.223 | 0.106 | -2.105 | **0.035** |
| PC2 | -0.033 | 0.141 | -0.237 | 0.812 |
| GW | 0.339 | 0.183 | 1.851 | 0.064 |
| SW | 0.764 | 0.261 | 2.924 | **0.003** |
| St | 0.580 | 0.378 | 1.533 | 0.125 |
| Great Spotted Woodpecker |  |  |  |  |
| PC1 | 0.006 | 0.0394 | 0.141 | 0.888 |
| PC2 | -0.247 | 0.051 | -4.813 | **<<0.001** |
| SW | -0.209 | 0.188 | -1.111 | 0.266 |
| H | 0.251 | 0.147 | 1.705 | 0.088 |
| St | 0.633 | 0.131 | 4.833 | **<<0.001** |

**Table 3.**
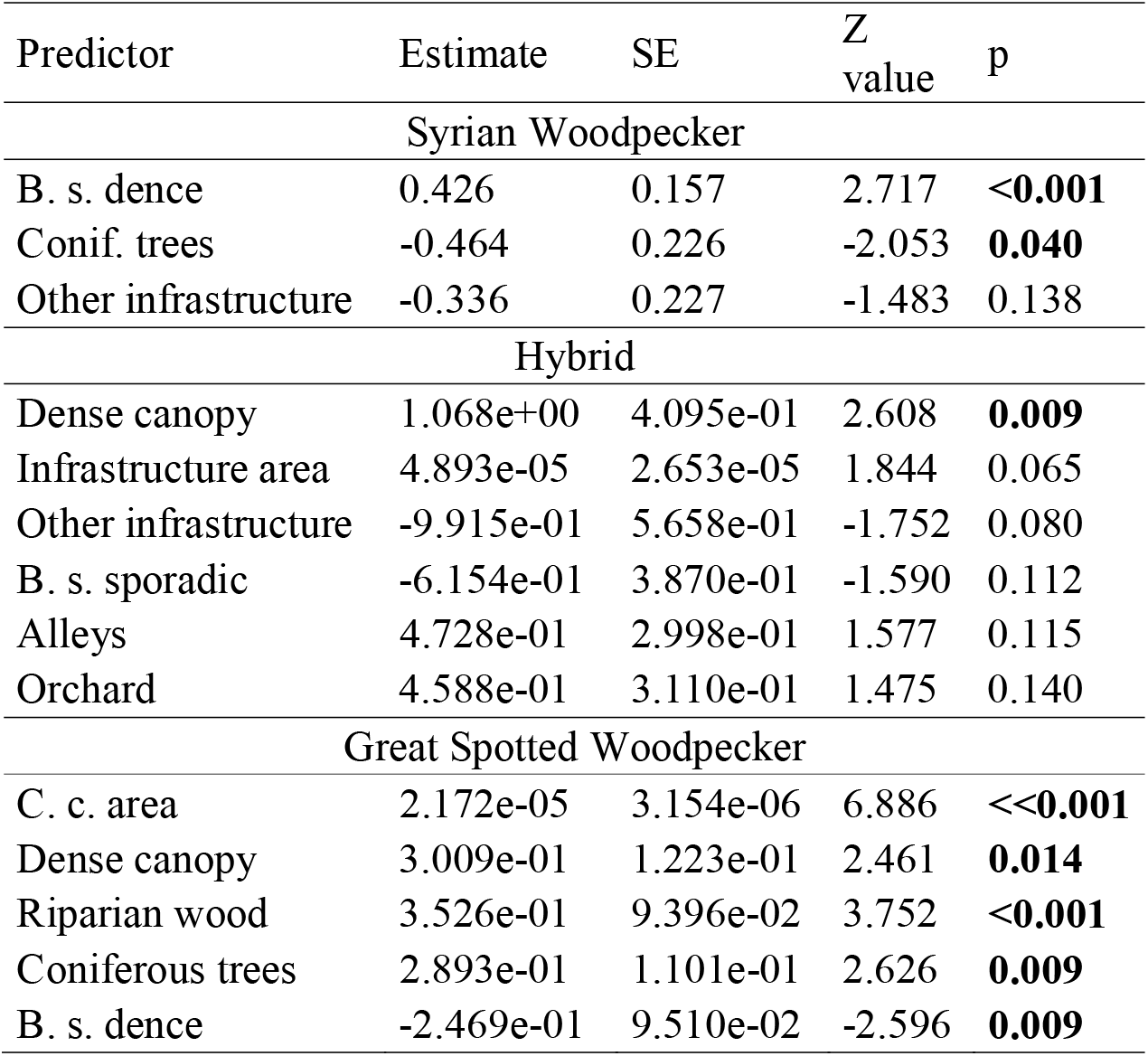
Generalised Linear Mixed Models demonstrating the relationship of Great Spotted Woodpecker, Syrian Woodpecker and their hybrids abundance to mix of particular habitat characteristic elements: Canopy cover area (C. c. area), Infrastructure area, Dense canopy, Forest patch, Riparian wood, Coniferous trees, Alleys, small buildings in single or sporadic placement (B. s. sporadic), small buildings in dense placement (B. s. dense), and other infrastructure. The significant P values (<0.05) are highlighted in bold. The analysis was made using European data.

| Predictor | Estimate | SE | Z<br>value | p |
| --- | --- | --- | --- | --- |
| Syrian Woodpecker |  |  |  |  |
| B. s. dence | 0.426 | 0.157 | 2.717 | **<0.001** |
| Conif. trees | -0.464 | 0.226 | -2.053 | **0.040** |
| Other infrastructure | -0.336 | 0.227 | -1.483 | 0.138 |
| Hybrid |  |  |  |  |
| Dense canopy | 1.068e+00 | 4.095e-01 | 2.608 | **0.009** |
| Infrastructure area | 4.893e-05 | 2.653e-05 | 1.844 | 0.065 |
| Other infrastructure | -9.915e-01 | 5.658e-01 | -1.752 | 0.080 |
| B. s. sporadic | -6.154e-01 | 3.870e-01 | -1.590 | 0.112 |
| Alleys | 4.728e-01 | 2.998e-01 | 1.577 | 0.115 |
| Orchard | 4.588e-01 | 3.110e-01 | 1.475 | 0.140 |
| Great Spotted Woodpecker |  |  |  |  |
| C. c. area | 2.172e-05 | 3.154e-06 | 6.886 | **<<0.001** |
| Dense canopy | 3.009e-01 | 1.223e-01 | 2.461 | **0.014** |
| Riparian wood | 3.526e-01 | 9.396e-02 | 3.752 | **<0.001** |
| Coniferous trees | 2.893e-01 | 1.101e-01 | 2.626 | **0.009** |
| B. s. dence | -2.469e-01 | 9.510e-02 | -2.596 | **0.009** |

**Table 4.** Generalised Linear Models demonstrating the relationships of Great Spotted Woodpecker, Syrian Woodpecker and their hybrids abundance to a mix of particular habitat characteristic elements: Managed canopy area, Orchard canopy area, Wild canopy area, Infrastructure area, Dense canopy, Riparian wood, tall buildings in single or sporadic placement (B. t. sporadic). The significant P values (<0.05) highlighted in bold. The analysis was made using SE Poland data.

| Predictor | Estimate | SE | Z<br>value | p |
| --- | --- | --- | --- | --- |
| Syrian Woodpecker |  |  |  |  |
| B. t. sporadic | 1.132e+00 | 3.464e-01 | 3.267 | **0.001** |
| Orchard green area | 4.792e-05 | 2.119e-5 | 2.262 | **0.024** |
| Infrastructure area | -6.055e-05 | 2.703e-05 | -2.240 | **0.025** |
| Hybrid |  |  |  |  |
| Managed canopy area | 2.711e-05 | 9.771e-06 | 2.744 | **0.005** |
| Dense canopy | 9.837e-01 | 4.915e-01 | 2.002 | **0.045** |
| B. t. sporadic | 6.636e-01 | 3.404e-01 | 1.949 | 0.051 |
| Great Spotted Woodpecker |  |  |  |  |
| Managed canopy area | 3.443e-05 | 4.495e-06 | 7.660 | **<<0.001** |
| Wild canopy area | 3.159e-05 | 5.578e-06 | 5.663 | **<<0.001** |
| Orchard canopy area | 3.142e-05 | 1.632e-05 | 1.925 | 0.054 |
| Riparian wood | 2.604e-01 | 1.369e-01 | 1.902 | 0.057 |
| Dense canopy | 2.745e-01 | 1.681e-01 | 1.633 | 0.102 |

**Table 4.**
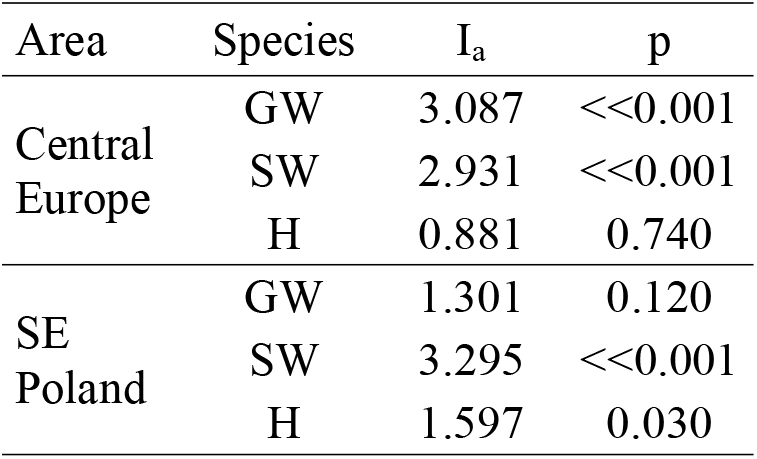
SADIE for Great Spotted Woodpecker, Syrian Woodpecker and their hybrids in Central Europe and SE Poland. Aggregation index (I_a_) values significantly bigger than 1 indicate a cluster of observations, while index values near 1 indicate a random distribution across the study area.

| Area | Species | $I_a$ | p |
| --- | --- | --- | --- |
| Central Europe | GW | 3.087 | <<0.001 |
|  | SW | 2.931 | <<0.001 |
|  | H | 0.881 | 0.740 |
| SE Poland | GW | 1.301 | 0.120 |
|  | SW | 3.295 | <<0.001 |
|  | H | 1.597 | 0.030 |

**Table 5.** SADIE for Great Spotted Woodpecker, Syrian Woodpecker and their hybrids, in Central Europe and SE Poland. Aggregation index (I_a_) values significantly bigger than 1 indicate a cluster of observations, while index values near 1 indicate a random distribution across the study area.

| Area | Species | $I_a$ | p |
| --- | --- | --- | --- |
| Central Europe | GW | 3.043 | **<<0.001** |
|  | SW | 2.728 | **<<0.001** |
|  | H | 1.198 | 0.200 |
| Lesser Poland | GW | 1.301 | 0.120 |
|  | SW | 3.295 | **<<0.001** |
|  | H | 1.468 | **0.040** |

**Table 6.** The association indexes between populations of Great Spotted Woodpecker, Syrian Woodpecker, their hybrids, *Picus* woodpeckers (GrW) and Starlings (St) at European and local (SE Poland) levelss. Association index values range from 1 (strong association) to -1 (strong dissociation).

| Species pair | GWxSW | GWxH | SWxH |
| --- | --- | --- | --- |
| European level |  |  |  |
| Association Index | -0.072 | 0.014 | 0.074 |
| P | 0.801 | 0.314 | 0.213 |
| Local level |  |  |  |
| Association Index | -0.050 | 0.233 | 0.045 |
| P | 0.364 | 0.091 | 1.00 |

## References

Aghanajafizadeh, S., Heydari, F., Naderi, G. and Hemami, MR. (2011) Nesting hole site selection by the Syrian Woodpecker, Dendrocopos syriacus, in Yazd province, Iran (Aves: Picidae). Zoology in the Middle East 53:3–6. 10.1080/09397140.2011.10638494

Aguillon, S.M. and Rohwer, V.G. (2022), Revisiting a classic hybrid zone: Movement of the northern flicker hybrid zone in contemporary times. Evolution, 76: 1082–1090. 10.1111/evo.14474

Baddeley A, Rubak E, Turner R (2015). Spatial Point Patterns: Methodology and Applications with R. Chapman and Hall/CRC Press, London. ISBN 9781482210200. https://www.routledge.com/Spatial-Point-Patterns-Methodology-and-Applications-with-R/Baddeley-Rubak-Turner/p/book/9781482210200/

Bakai, A., Figarski, T., Grzędzicka, E., Kusal, B., Gorman, G., Černý, M., Fišerová, A., Veselý, P., Zielinski, P., Smykla, J., Pochłopień, Z., Lešo, P., Czajowski, K., Aleksandrowicz, M., Borowik, M., Broński, S., Chodkiewicz, T., Janiszewska, A., Kawka, J., Kłonowski, P., Krzyżanowski, M., Kuczmarski, S., Lasoń, W., Loudová, K., Matysek, M., Mazgaj, S., Miedviedieva, I., Murzyn, B., Niedziółka, M., Puchala, P., Schmidt, A., Sobus, T., Tchoń, K., Veszelinov, O., Wehrichová, K., Zhulenko, V., Ziółkowski, T., Życki, A. (2025). Citizen science and detailed searches reveal widespread occurrence of hybrid-like Syrian x Great-spotted woodpeckers. bioRxiv. 10.64898/2025.12.22.695873

Bakai, A., Fuchs J., Gorman, G., Sajdak, D., Kajtoch, Ł. (2026a). A systematic review of interspecific breeding in woodpeckers. IBIS 168: 415:430. 10.1111/ibi.70014

Bakai, A., Gorman, G. and Kajtoch, Ł. (2026b) A review of co-occurrence and hybridization as neglected factors in studies of Syrian Woodpecker (Dendrocopos syriacus) and Great Spotted Woodpecker (Dendrocopos major). Journal of Ornithology 167: 1–15. 10.1007/s10336-025-02298-w

Barrowclough, G. F., Groth, J. G., Bramlett, E. K., Lai, J. E. and Mauck, W. M. (2017). Phylogeography and geographic variation in the Red-bellied Woodpecker (Melanerpes carolinus): characterization of mtDNA and plumage hybrid zones. Wilson J. Ornithol. 130: 671–683. 10.1676/17-070.1

Bartoń K (2026). MuMIn: Multi-Model Inference. R package version 1.48.19, https://CRAN.R-project.org/package=MuMIn.

Billerman, S.M., Cicero, C., Bowie, R.C.K. and Carling, M.D. (2019). Phenotypic and genetic introgression across a moving woodpecker hybrid zone. Mol. Ecol. 28: 1692–1708.

Brooks ME, Kristensen K, van Benthem KJ, Magnusson A, Berg CW, Nielsen A, Skaug HJ, Maechler M, Bolker BM (2017). “glmmTMB Balances Speed and Flexibility Among Packages for Zero-inflated Generalized Linear Mixed Modeling.” The R Journal, 9(2), 378–400. doi:10.32614/RJ-2017-066.

Dudzik, K. and Polakowski, M. (2011). The cases of mixed broods and identification of Syrian Woodpecker Dendrocopos syriacus and Great Spotted Woodpecker Dendrocopos major hybrids in Poland. Chrońmy Przyrodę Ojczystą 67:254–260.

Feber, R. E., Johnson, P. J., and Bourn, N. A. D. (2025). Quantifying the value of trees outside woods for promoting biodiversity on farmland. Ecological Solutions and Evidence, 6, e70042. 10.1002/2688-8319.70042

Figarski, T. (2017). Contrasting seasonal reactions of two sibling woodpeckers to playback stimulation in urban areas — implications for inventory and monitoring of the Syrian woodpecker. Behaviour, 154: 981–996. https://www.jstor.org/stable/26488532.

Figarski, T. and Kajtoch, Ł. (2018). Hybrids and mixed pairs of Syrian and great-spotted woodpeckers in urban populations. J. Ornithol. 159: 311–314.

Figarski T., Kajtoch Ł. (2018). Differences in habitat requirements between two sister Dendrocopos woodpeckers in urban environments: implication for the conservation of Syrian Woodpecker. Acta Ornithologica. 53: 23–36

Fox, J. and Weisberg, S. (2019). An R Companion to Applied Regression, 3rd Edition. Thousand Oaks, CA <https://www.john-fox.ca/Companion/index.html>

Friedmann V.S. 2011. A riddle of the Green Woodpecker: how appear hybrids with the Grey Woodpecker? Berkut. 20 (1-2).

Fröhlich, A. and Ciach, M. (2013). Distribution and abundance of the Syrian Woodpecker Dendrocopos syriacus in Kraków. Ornis Polonica, 54: 237–246.

Fröhlich, A., Hawryło, P., and Ciach, M. (2022). Urbanization filters woodpecker assemblages: Habitat specialization limits population abundance of dead wood dependent organisms in the urban landscape. Global Ecology and Conservation, 38, e02220. 10.1016/j.gecco.2022.e02220.

Gentili, R., Quaglini, L.A., Galasso, G. et al. (2024) Urban refugia sheltering biodiversity across world cities. Urban Ecosyst 27, 219–230. 10.1007/s11252-023-01432-x

Gigot, C. (2023). epiphy: Analysis of Plant Disease Epidemics. doi:10.32614/CRAN.package.epiphy. <https://doi.org/10.32614/CRAN.package.epiphy>, R package version 0.5.0, <https://CRAN.R-project.org/package=epiphy>.

Glutz von Blotzheim U. N., Bauer K. (1980). Handbuch der Vögel Mitteleuropas. 9. Akademi sche Verlag, Wiesbaden.

Gorman, G. 1997. Hybridisation by Syrian woodpeckers. Br. Birds 90: 578.

Gorman G., Kajtoch Ł. 2026. Identification of Great Spotted Woodpecker x Syrian Woodpecker hybrids. British Birds 119: 166–171.

Gurgul, A., Miksza-Cybulska, A., Szmatoła, T., Semik-Gurgul, E., Jasielczuk, I., Bugno-Poniewierska, M., Figarski, T. and Kajtoch, Ł. (2019). Evaluation of genotyping by sequencing for population genetics of sibling and hybridising birds: An example using Syrian and great spotted woodpeckers. J. Ornithol. 160: 287–294.

Hartig F (2026). _DHARMa: Residual Diagnostics for Hierarchical (Multi-Level / Mixed) Regression. Models. doi:10.32614/CRAN.package.DHARMa <https://doi.org/10.32614/CRAN.package.DHARMa>, R package version 0.5.0, <https://CRAN.R-project.org/package=DHARMa>.

Hebda, G., Wesołowski, T. and Rowiński, P. (2017). Nest Sites of a Strong Excavator, the Great Spotted Woodpecker Dendrocopos major, in a Primeval Forest. Ardea, 105:61–71. 10.5253/arde.v105i1.a8

Kajtoch, Ł. and Figarski, T. (2017). Comparative distribution of Syrian and Great spotted woodpeckers in different landscapes of Poland. Folia Zoologica. 66: 29–36.

Kajtoch, Ł. and Kusal, B. 2022. The first case of a successful brood from a double hybrid mixed pair (Dendrocopos syriacus 9 Dendrocopos major (Picidae)). Ibis 164: 1273–1277.

Kajtoch, Ł. and Kusal, B. (2023) Decline in the population of the Syrian Woodpecker Dendrocopos syriacus in the Krakow agglomeration. Ornis Polonica, 64: 119–128. 10.12657/ornis.2023.2.3

Keller, V., Herrando, S., Voříšek, P., Franch, M., Kipson, M., Milanesi, P., Martí, D., Anton, M., Klvaňová, A., Kalyakin, M.V., Bauer, H.-G. and Foppen, R.P.B. (2020). European Breeding Bird Atlas 2: Distribution, Abundance and Change. European Bird Census Council and Lynx Edicions, Barcelona.

Kassambara A, Mundt F (2026). factoextra: Extract and Visualize the Results of Multivariate Data Analyses. R package version 2.2.0. With contributions from Laszlo Erdey (Faculty of Economics and Business, University of Debrecen, Hungary), https://CRAN.R-project.org/package=factoextra.

Kroneisl-Rucner, R. (1957). Bird-banding in 1956. 1. Results of the bird-banding carried out by the Ornithological Institute, Department of the Museum at Zagreb, 11th report; 2. Foreign recoveries made in Yugoslavia, 7th report. Larus, 11: 5–22.

Lê S, Josse J, Husson F (2008). “FactoMineR: A Package for Multivariate Analysis.” Journal of Statistical Software, 25(1), 1–18. doi:10.18637/jss.v025.i01.

Ławicki, Ł., Cofta, T., Beuch, S., Dmoch, A., Sikora. A., Aftyka, S., Czechowski, P., Bocheński, M., Sieczak, K. and Mazgaj, S. (2015). Identification and occurrence of hybrids Grey-headed x European Green Woodpecker in Poland. Dutch Birding 37: 215–228.

Mazgajski, T. (1998) Nest-site characteristic of Great Spotted Woodpecker Dendrocopos major in Central Poland. Polish Journal of Ecology 46(1):33–41

Mazgajski, T. D. (2000). Competition for nest sites between the Starling Sturnus vulgaris and other cavity nesters—study in forest park. Acta Ornithologica, 35(1), 103–107.

Michalczuk, J. (2014) Expansion of the Syrian Woodpecker Dendrocopos syriacus in Europe and Western Asia. Ornis Polonica, 55: 149–161.

Michalczuk, J. and Michalczuk, M. (2006). Reaction to Playback and Density Estimations of Syrian Woodpeckers Dendrocopos syriacus in Agricultural Areas of South-Eastern Poland. Acta Ornithologica, 41: 33–39. DOI:10.3161/068.041.0109

Michalczuk, J. and Michalczuk, M. (2015) Decline of the Syrian Woodpecker Dendrocopos syriacus population in rural landscape in Lesser Poland in 2004–2012. Ornis Polonica 56: 67–75. 10.12657/ornis.2015.2.1

Michalczuk, J. and Michalczuk, M. (2016a). Habitat preferences of Picidae woodpeckers in the agricultural landscape of Lesser Poland: is the Syrian woodpecker Dendrocopos syriacus colonizing a vacant ecological niche? North-West. J. Zool. 12: 14–21.

Michalczuk, J. and Michalczuk, M. (2016b). Coexistence of Syrian Woodpecker Dendrocopos syriacus and Great Spotted Woodpecker Dendrocopos major in nonforest tree stands of the agricultural landscape in Lesser Poland. Turkish Journal of Zoology 40:743–748. DOI:10.3906/zoo-1601-13.

Michalczuk, J., McDevitt, A.D., Mazgajski, T.D., Figarski, T., Ilieva, M., Bujoczek, M. and Kajtoch, Ł. (2014). Tests of multiple molecular markers for the identification of great spotted and Syrian woodpeckers and their hybrids. J. Ornithol. 155: 591–600.

Minias, P. and Janiszewski, T. (2016), Territory selection in the city: can birds reliably judge territory quality in a novel urban environment?. J Zool, 300: 120–126. 10.1111/jzo.12362

Natola, L., Curtis, A., Hudon, J. and Burg, T.M. (2021). Introgression between Sphyrapicus nuchalis and S. varius sapsuckers in a hybrid zone in west-central Alberta. J. Avian Biol. 52: e02717.

Oksanen J, Simpson G, Blanchet F, Kindt R, Legendre P, Minchin P, O’Hara R, Solymos P, Stevens M, Szoecs E, Wagner H, Bedward M, Bolker B, Borcard D, Carvalho G, De Caceres M, Durand S, Evangelista H, Hannigan G, Hill M, Lahti L, Martino C, Ouellette M, Ribeiro Cunha E, Smith T, Stier A, Ter Braak C and, Weedon J (2026). vegan: Community Ecology Package. R package version 2.8–0, https://vegandevs.github.io/vegan/.

Ottenburghs, J. and Nicolaï, MP. (2024). Hybridization constrains the evolution of mimicry complexes in woodpeckers, Journal of Avian Biology 2024.

Pebesma E, Bivand R (2023). Spatial Data Science: With applications in R. Chapman and Hall/CRC. doi:10.1201/9780429459016. https://r-spatial.org/book/.

Pons, J.-M., Masson, C., Olioso, G. and Fuchs, J. (2019). Gene flow and genetic admixture across a secondary contact zone between two divergent lineages of the Eurasian green woodpecker Picus viridis. J. Ornithol. 160: 935–945.

R Core Team 2023. R: A language and environment for statistical computing. Vienna: R Foundation for Statistical Computing. https://www.R-project.org/

Ranasinghe, R.W., Seneviratne, S.S. and Irwin, D. 2024. Cryptic hybridisation dynamics in a three-way hybrid zone of Dinopium flamebacks on a tropical Island. Ecol. Evol. 14: e70716.

Rousset, F. and Ferdy, J. (2014). Testing environmental and genetic effects in the presence of spatial autocorrelation. Ecography, 37: 781–790. <10.1111/ecog.00566>.

Reiser, O. Materialien zu einer Ornis Balcanica. II. Wien: Gerold 1894.

Rüge, K. (1969). Beobachtungen am Blutspecht Dendrocopos syriacus im Burgenland. Vogelwelt, 90, 201–223.

Seehausen O. 2004. Hybridization and adaptive radiation. Trends in Ecology & Evolution, 19, 198–207

Smith, K. W. (2005). Has the reduction in nest-site competition from Starlings Sturnus vulgaris been a factor in the recent increase of Great Spotted Woodpecker Dendrocopos major numbers in Britain? Bird Study, 52(3), 307–313. 10.1080/00063650509461404

H. Wickham. ggplot2: Elegant Graphics for Data Analysis. Springer-Verlag New York, 2016.

Winkler, H. (1971). Die artliche Isolation des Blutspechts Picoides (Dendrocopos) syriacus. Egretta 14:1–20.

Winkler, H. (1973). Food acquisition and competition in two Holarctic woodpeckers. Oecologia, 12(3), 193–208. 10.1007/BF00345517

