## Supplementary file for "Environmental features or interspecific relations: What determines distribution of Syrian and Great Spotted woodpeckers hybrids?"

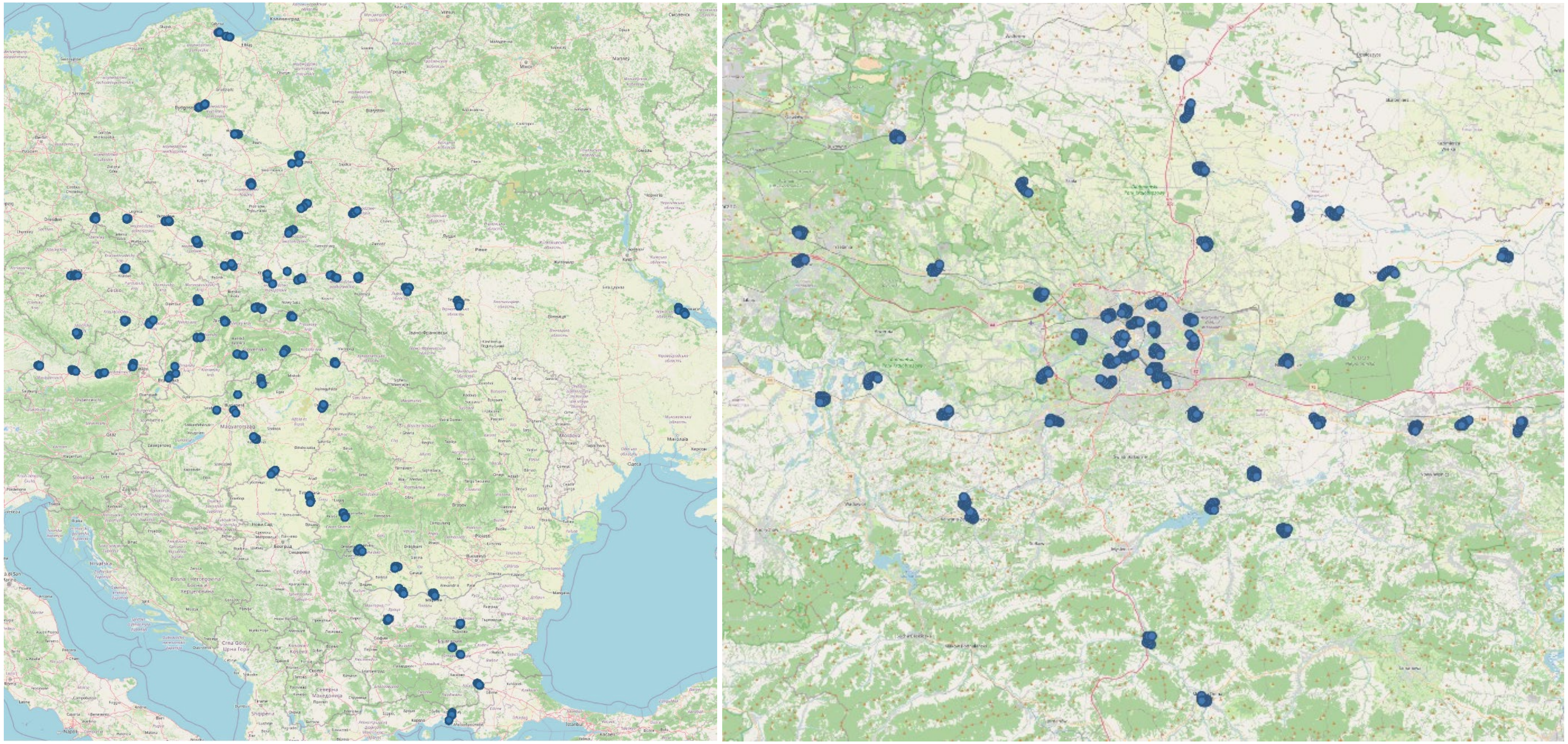

**Figure S1.** Study area of Syrian Woodpecker × Great Spotted Woodpecker hybridisation, in European (A) and regional Lesser Poland (SE Poland)(B) areas. The maps contain 1.5 (European) and 2.4 km (Local) transects where woodpecker hybrids were recorded.

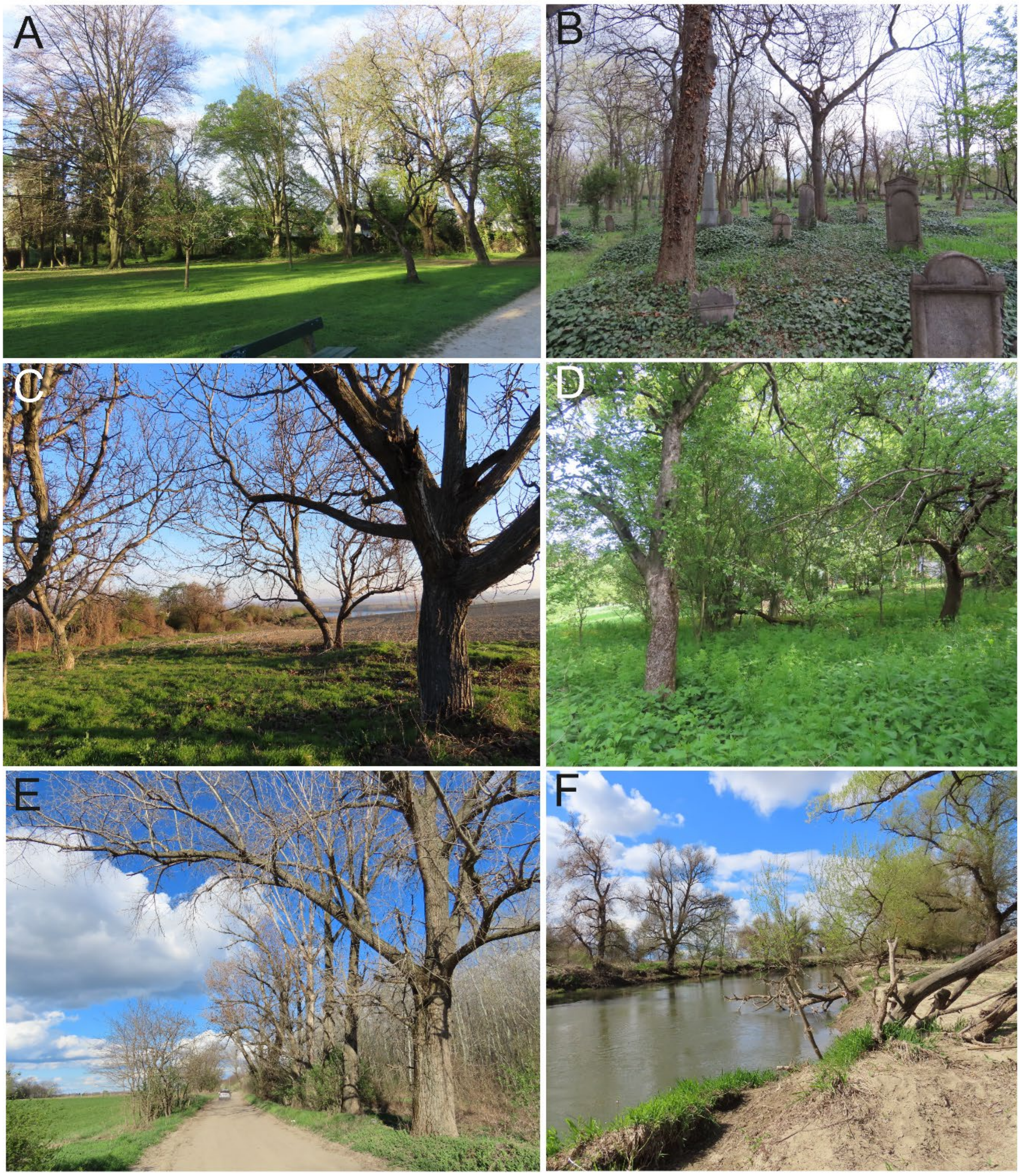


**Figure S2.** Examples of woodland types occupied by Syrian × Great Spotted Woodpecker hybrids in European (A) urban park, (B) cemetery, (C) rural orchard, (D) suburban allotment, (E) rural lanes, (F) riparian woodland.

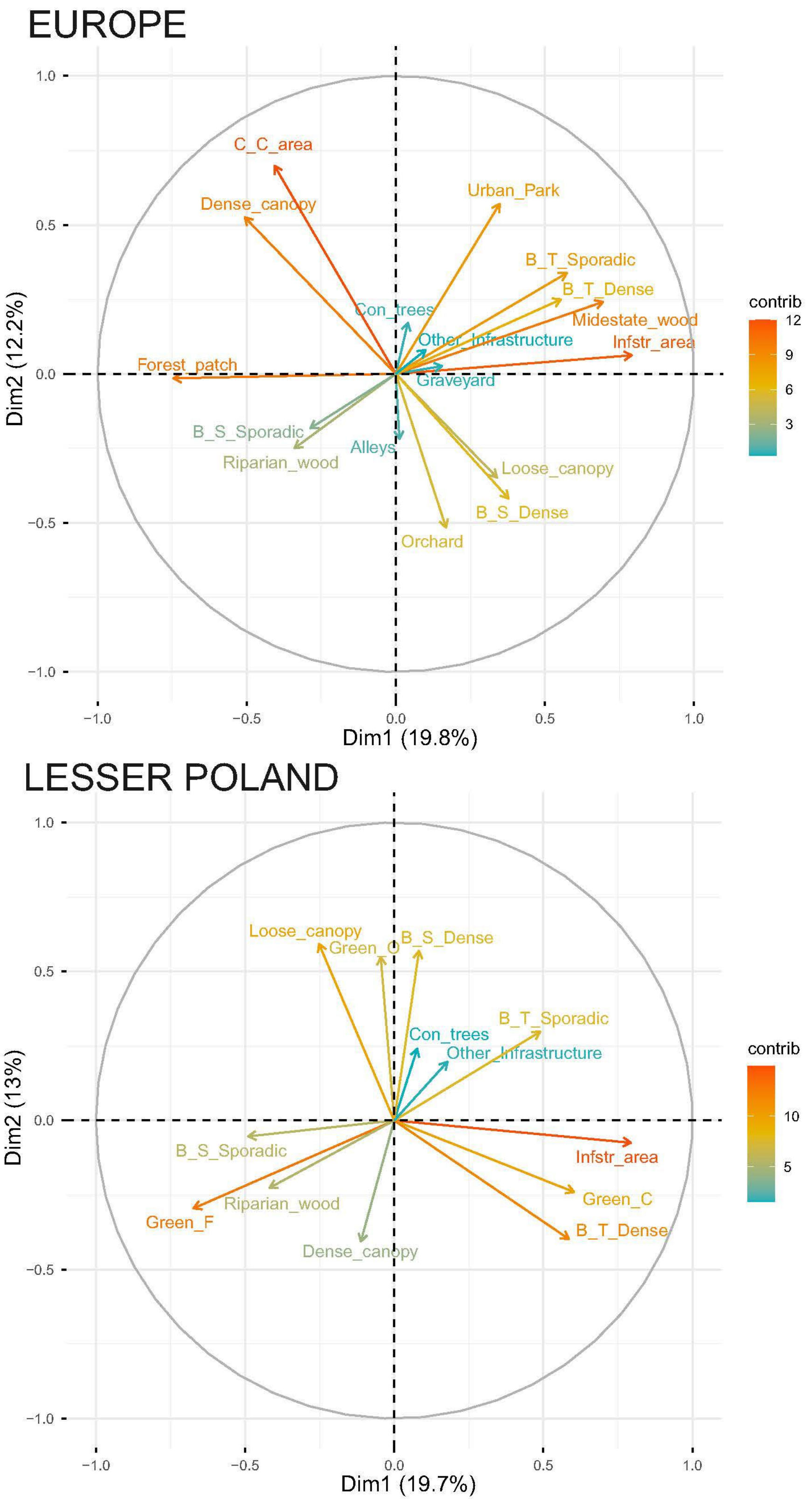


**Figure S3**. The Principal Component Analysis of environmental variables present in an area of 150 m around count point records of Great Spotted Woodpeckers, Syrian Woodpeckers and their hybrids. The variable trends shown along the first two PCs illustrate the highest proportions of variation and the contribution of each variable is shown by arrow length and colour.

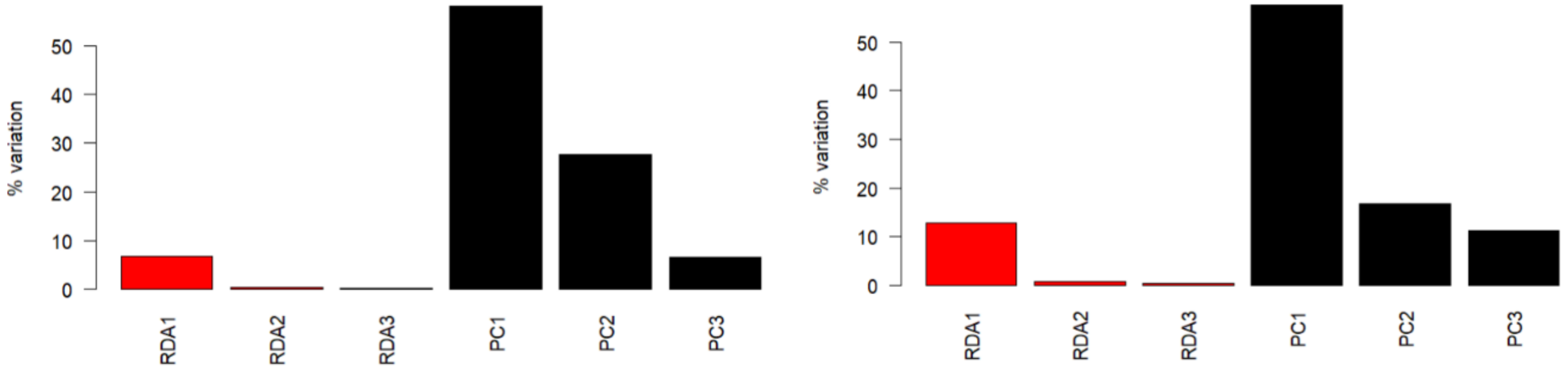


**Figure S4.** Transformation-based Redundancy analysis bar-plots for Great Spotted Woodpecker, Syrian Woodpecker and their hybrids in relation to environmental factors. A) Based on European scale predictors: Canopy cover area, Infrastructure area, Dense canopy, Loose canopy, Coniferous trees, Forest patch, Urban park, Cemetery, Alleys, Mid-estate wood, Orchard, Riparian wood, Buildings small sporadic, Buildings small dense, Buildings tall sporadic, Buildings tall dense and Other infrastructure; B) SE Poland scale predictors: Managed trees area, Orchard-area, Unmanaged trees area, Infrastructure area, Dense canopy, Loose canopy, Coniferous trees, Buildings small sporadic, Buildings small dense, Buildings tall sporadic, Buildings tall dense and Other infrastructure; Bars corresponding to the explained variation share (%) of three first constrained (red) and unconstrained (black) axes.

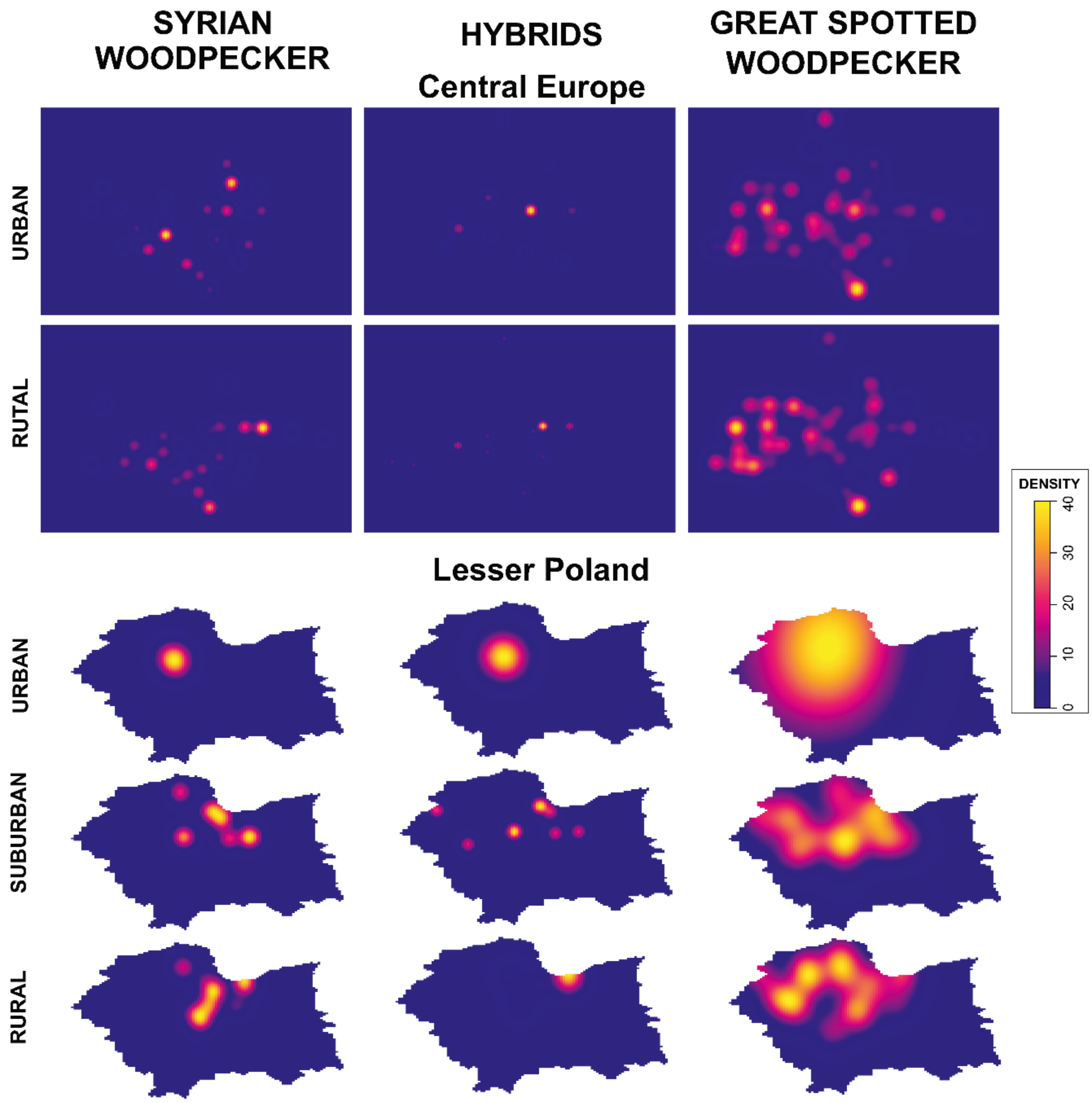


**Figure S5.** Graphical visualisation of Syrian Woodpecker, Great Spotted Woodpecker, and the distribution of hybrids in Central Europe and SE Poland. Kernel Density Estimation pixel plots based on each single bird record and grouped according to landscape type (Rural, Urban, Suburban).

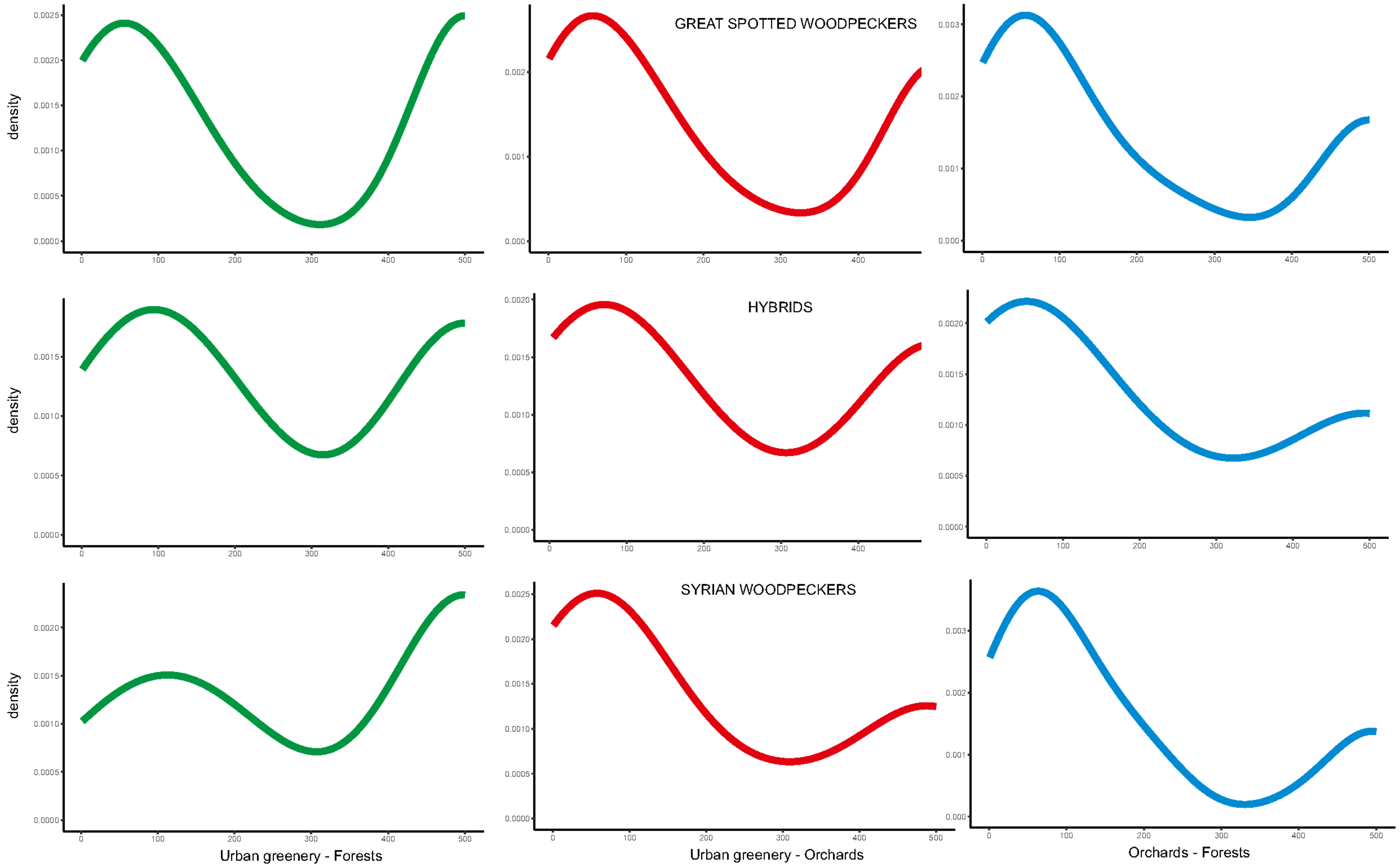


**Figure S6'.** Kernel Density plots depicting the Syrian Woodpecker, Great Spotted Woodpecker, and the presence of hybrids depending on the distance between one of the tree-stand types (Managed, Orchard, Wild) present in the study plot (R = 150 m) to closest different tree-stand type in the area within this plot and an additional 150 m around it. The . If the closest tree-stand type was outside the study plot area and 150m buffer (total R=300), the distance was marked as 500 m.

**Table S1.** Description of environmental predictors used to model Great Spotted Woodpecker, Syrian Woodpecker and hybrid habitat preferences.

| Variable name | data type | units | description |
|---|---|---|---|
| NS | numeric | degrees | longitude |
| WE | numeric | degrees | latitude |
| T_name | factor | - | name of the transect |
| Point_ID | factor | - | name of the specific point on transects |
| GW | numeric | number | number of Great Spotted Woodpeckers within a 150m radius of a point |
| SW | numeric | number | number of Syrian Woodpeckers within a 150m radius of a point |
| H | numeric | number | number of hybrids or pure (GWxSW) mixed pairs within a 150m radius of a point |
| C_C_area | numeric | m2 | canopy cover - area of all trees (crowns) within a 150m radius of a point |
| Infstr_area | numeric | m2 | "infrastructure" - area of all buildings, railways and highways within a 150m radius of a point |
| Dense_canopy | binary | 0/1 | presence or absence of dense (and at least more than 50m2) patch of trees within a 150m radius of a point |
| Loose_canopy | binary | 0/1 | presence or absence of sporadic trees or loose patches of trees within a 150m radius of a point |
| Con_trees | binary | 0/1 | presence or absence of coniferous trees (pine, spruce) within a 150m radius of a point |
| Forest_patch | binary | 0/1 | presence or absence of unmanaged wood in 150m radius of a point |
| Urban_Park | binary | 0/1 | presence or absence of managed urban park within a 150m radius of a point |
| Graveyard | binary | 0/1 | presence or absence of graveyard within a 150m radius of a point |
| Alleys | binary | 0/1 | presence or absence of long narrow tree belts within a 150m radius of a point |
| Mid-estate_wood | binary | 0/1 | presence or absence of mid-estate wood within a 150m radius of a point |
| Orchard | binary | 0/1 | presence or absence of orchard within a 150m radius of a point |
| Riparian_wood | binary | 0/1 | presence or absence of riparian wood (soft-wood trees around water bodies) within a 150m radius of a point |
| B_S_Sporadic | binary | 0/1 | presence or absence of sporadic small buildings within a 150m radius of a point |

| | | | |
|---|---|---|---|
| B_S_Dense | binary | 0/1 | presence or absence of dense aggregations of small buildings within a 150m radius of a point |
| B_T_Sporadic | binary | 0/1 | presence or absence of sporadic tall buildings within a 150m radius of a point |
| B_T_Dense | binary | 0/1 | presence or absence of dense aggregations of tall buildings (as in old-towns) within a 150m radius of a point |
| Other_Infrastructure | binary | 0/1 | presence or absence of railways, highways, marketplaces, industrial infrastructure and others, within a 150m radius of a point |
| Green_C | numeric | m2 | Area of managed woodland in residential areas (parks, mid-estate wood) |
| Green_O | numeric | m2 | Area of orchards, gardens and trees on private property with fruit trees. |
| Green_F | numeric | m2 | Area of wild or quasi-natural vegetation with understorey (abandoned tree stands, urban or rural forests, trees on arable land, etc.) |
| CO | numeric | m | Distance between managed wood patches within a 150 m radius of the count point and closest vegetation with fruit trees within a 300 m radius |
| CF | numeric | m | Distance between managed wood patches within a 150 m radius around a count point and closest quasi-natural vegetation with a 300 m radius |
| OF | numeric | m | Distance between vegetation with fruit trees within a 150 m radius of a count point and closest quasi-natural vegetation within a 300 m radius |

**Table S4.** Transformation-based Redundancy analysis for Great Spotted Woodpecker, Syrian Woodpecker and their hybrids in relation to environmental factors. A) Based on European scale predictors: Canopy cover area, Infrastructure area, Dense canopy, Loose canopy, Coniferous trees, Forest patch, Urban park, Graveyard, Alleys, Mid-estate wood, Orchard, Riparian wood, Buildings small sporadic, Buildings small dense, Buildings tall sporadic, Buildings tall dense and Other infrastructure; B) SE Poland scale predictors: Managed trees area, Orchard-area, Unmanaged trees area, Infrastructure area, Dense canopy, Loose canopy, Coniferous trees, Buildings small sporadic, Buildings small dense, Buildings tall sporadic, Buildings tall dense and Other infrastructure.

| Scale | European | | Lesser Poland | |
|---|---|---|---|---|
| | Inertia | Proportion | Inertia | Proportion |
| Total | 0.361 | 1.000 | 0.333 | 1.000 |
| Constrained | 0.027 | 0.076 | 0.047 | 0.142 |
| Unconstrained | 0.334 | 0.924 | 0.286 | 0.858 |
| Eigenvalues for | RDA1 | RDA2 | RDA1 | RDA2 |
| constrained axes | 0.025 | 0.002 | 0.043 | 0.003 |
| Eigenvalues for | PC1 | PC2 | PC1 | PC2 |
| unconstrained axes | 0.210 | 0.100 | 0.192 | 0.056 |

Table S2. Database of examined transects with records of woodpeckers and environmental variables collected across Europe.

| NS | EW | T_name | Point_ID | Country | Type | GW | GWpr | SW | SWpr | H | Hpr | St | Altitude | Distance_SP | Distance_DE | C_C_area | Infstr_area | Dense_canopy | Loose_canopy | Con_trees | Forest_patch | Urban_Park | Graveyard | Alleys | Midestate_wood | Orchard | Riparian_wood | B_S_Sporadic | B_S_Dense | B_T_Sporadic | B_T_Dense | Other_Infrastructure | GrW |
|---|---|---|---|---|---|---|---|---|---|---|---|---|---|---|---|---|---|---|---|---|---|---|---|---|---|---|---|---|---|---|---|---|---|
| 48.35922 | 13.29235 | BAD_R_1 | badr1_1 | Germany | R | 3 | 1 | 0 | 0 | 0 | 0 | 1 | 330,808 | 1481,73382 | 107 | 30599,428 | 0 | 1 | 1 | 0 | 1 | 0 | 0 | 0 | 0 | 1 | 1 | 0 | 1 | 0 | 0 | 0 | 0 |
| 48.36037 | 13.29532 | BAD_R_1 | badr1_2 | Germany | R | 1 | 1 | 0 | 0 | 0 | 0 | 0 | 330 | 1481,654753 | 107 | 32281,835 | 0 | 1 | 0 | 0 | 1 | 0 | 0 | 0 | 0 | 0 | 1 | 1 | 0 | 0 | 0 | 0 | 0 |
| 48.36197 | 13.29858 | BAD_R_1 | badr1_3 | Germany | R | 0 | 0 | 0 | 0 | 0 | 0 | 0 | 326,908 | 1481,657867 | 107 | 15630,529 | 0 | 1 | 0 | 0 | 1 | 0 | 0 | 0 | 0 | 0 | 1 | 0 | 0 | 0 | 0 | 0 | 0 |
| 48.3633 | 13.30305 | BAD_R_1 | badr1_4 | Germany | R | 1 | 1 | 0 | 0 | 0 | 0 | 1 | 324 | 1481,700923 | 107 | 14572,364 | 271,472197 | 1 | 0 | 0 | 1 | 0 | 0 | 0 | 0 | 0 | 1 | 0 | 0 | 0 | 0 | 0 | 0 |
| 48.36486 | 13.30712 | BAD_R_1 | badr1_5 | Germany | R | 3 | 1 | 0 | 0 | 1 | 1 | 1 | 323 | 1481,649057 | 107 | 21208,283 | 3200,17051 | 1 | 0 | 0 | 1 | 0 | 0 | 0 | 0 | 0 | 1 | 0 | 0 | 0 | 0 | 0 | 1 |
| 48.33988 | 13.34132 | BAD_R_2 | badr2_1 | Germany | R | 0 | 0 | 0 | 0 | 0 | 0 | 0 | 316,325 | 1485,946925 | 108 | 57583,516 | 373,4277344 | 1 | 0 | 0 | 1 | 0 | 0 | 1 | 0 | 0 | 0 | 1 | 0 | 0 | 0 | 0 | 0 |
| 48.3423 | 13.34441 | BAD_R_2 | badr2_2 | Germany | R | 1 | 1 | 0 | 0 | 0 | 0 | 0 | 316,596 | 1485,779401 | 108 | 33386,268 | 0 | 1 | 1 | 0 | 1 | 0 | 0 | 1 | 0 | 0 | 1 | 0 | 0 | 0 | 0 | 0 | 0 |
| 48.34515 | 13.34653 | BAD_R_2 | badr2_3 | Germany | R | 1 | 1 | 0 | 0 | 0 | 0 | 1 | 314,016 | 1485,629689 | 108 | 60582,142 | 0 | 1 | 0 | 0 | 1 | 0 | 0 | 0 | 0 | 0 | 0 | 0 | 0 | 0 | 0 | 0 | 0 |
| 48.34826 | 13.34829 | BAD_R_2 | badr2_4 | Germany | R | 1 | 1 | 0 | 0 | 0 | 0 | 0 | 316 | 1485,430457 | 108 | 2975,781 | 0 | 1 | 1 | 0 | 1 | 0 | 0 | 1 | 0 | 0 | 0 | 1 | 0 | 0 | 0 | 0 | 1 |
| 48.35039 | 13.35078 | BAD_R_2 | badr2_5 | Germany | R | 0 | 0 | 0 | 0 | 0 | 0 | 1 | 313,808 | 1485,257661 | 108 | 3173,565 | 1360,666956 | 0 | 1 | 0 | 1 | 0 | 0 | 1 | 0 | 0 | 0 | 1 | 0 | 0 | 0 | 0 | 0 |
| 48.35426 | 13.34232 | BAD_U_1 | badu1_1 | Germany | U | 0 | 0 | 0 | 0 | 0 | 0 | 0 | 320,766 | 1482,399836 | 106 | 608,376 | 7227,044234 | 0 | 1 | 0 | 0 | 0 | 0 | 1 | 0 | 0 | 0 | 0 | 1 | 0 | 0 | 0 | 0 |
| 48.35611 | 13.34614 | BAD_U_1 | badu1_2 | Germany | U | 0 | 0 | 0 | 0 | 0 | 0 | 1 | 320 | 1482,23975 | 106 | 5230,257 | 12860,90666 | 0 | 1 | 0 | 0 | 0 | 0 | 1 | 1 | 1 | 0 | 0 | 1 | 0 | 0 | 0 | 0 |
| 48.35655 | 13.35034 | BAD_U_1 | badu1_3 | Germany | U | 0 | 0 | 0 | 0 | 0 | 0 | 0 | 319,094 | 1481,996952 | 106 | 2912,671 | 12326,84947 | 0 | 1 | 0 | 0 | 0 | 0 | 1 | 0 | 0 | 0 | 0 | 1 | 0 | 0 | 0 | 0 |
| 48.35486 | 13.35223 | BAD_U_1 | badu1_4 | Germany | U | 0 | 0 | 0 | 0 | 0 | 0 | 0 | 320 | 1481,772812 | 106 | 2453,632 | 9965,030512 | 0 | 1 | 0 | 0 | 0 | 0 | 0 | 1 | 0 | 0 | 0 | 1 | 0 | 0 | 0 | 0 |
| 48.35328 | 13.35024 | BAD_U_1 | badu1_5 | Germany | U | 0 | 0 | 0 | 0 | 0 | 0 | 1 | 319,834 | 1481,826125 | 106 | 4686,645 | 14782,63906 | 0 | 1 | 1 | 0 | 0 | 0 | 0 | 0 | 1 | 0 | 0 | 1 | 0 | 0 | 0 | 0 |
| 48.35002 | 13.31294 | BAD_U_2 | badu2_1 | Germany | U | 1 | 1 | 0 | 0 | 0 | 0 | 1 | 325 | 1483,721023 | 108 | 29062,19 | 10184,69724 | 1 | 1 | 0 | 0 | 1 | 0 | 0 | 0 | 0 | 0 | 0 | 1 | 1 | 0 | 0 | 0 |
| 48.34802 | 13.31612 | BAD_U_2 | badu2_2 | Germany | U | 0 | 0 | 0 | 0 | 0 | 0 | 0 | 322 | 1483,473818 | 108 | 25842,5 | 3221,695435 | 0 | 1 | 0 | 0 | 1 | 0 | 1 | 1 | 1 | 0 | 1 | 0 | 1 | 0 | 0 | 0 |
| 48.34993 | 13.32073 | BAD_U_2 | badu2_3 | Germany | U | 2 | 1 | 0 | 0 | 0 | 0 | 1 | 322,158 | 1483,606131 | 108 | 15903,495 | 8928,442978 | 1 | 1 | 0 | 1 | 0 | 0 | 1 | 1 | 0 | 0 | 0 | 0 | 1 | 1 | 0 | 0 |
| 48.35181 | 13.32397 | BAD_U_2 | badu2_4 | Germany | U | 0 | 0 | 0 | 0 | 0 | 0 | 1 | 321 | 1483,812491 | 108 | 47388,251 | 1683,079729 | 1 | 1 | 0 | 1 | 0 | 0 | 0 | 1 | 0 | 0 | 0 | 0 | 1 | 0 | 0 | 0 |
| 48.35349 | 13.32171 | BAD_U_2 | badu2_5 | Germany | U | 1 | 1 | 0 | 0 | 0 | 0 | 1 | 322 | 1484,117108 | 108 | 49599,141 | 248,2682544 | 1 | 1 | 0 | 1 | 1 | 0 | 0 | 0 | 0 | 0 | 1 | 0 | 0 | 0 | 0 | 1 |
| 48.17211 | 17.26891 | BRA_R_1 | brar1_1 | Slovakia | R | 1 | 1 | 0 | 0 | 0 | 0 | 0 | 131 | 1222,598055 | 83 | 51286,722 | 975 | 1 | 1 | 0 | 1 | 0 | 0 | 1 | 0 | 1 | 0 | 0 | 1 | 0 | 0 | 0 | 0 |
| 48.17417 | 17.27234 | BRA_R_1 | brar1_2 | Slovakia | R | 0 | 0 | 0 | 0 | 0 | 0 | 0 | 131 | 1222,524495 | 83 | 50312,243 | 7717 | 1 | 1 | 0 | 1 | 0 | 0 | 0 | 0 | 1 | 0 | 0 | 1 | 0 | 0 | 0 | 0 |
| 48.17735 | 17.27334 | BRA_R_1 | brar1_3 | Slovakia | R | 0 | 0 | 0 | 0 | 0 | 0 | 0 | 130,989 | 1222,669914 | 83 | 53186,804 | 2032 | 1 | 1 | 0 | 1 | 0 | 0 | 1 | 0 | 1 | 0 | 1 | 0 | 1 | 0 | 0 | 0 |
| 48.17924 | 17.2763 | BRA_R_1 | brar1_4 | Slovakia | R | 0 | 0 | 0 | 0 | 0 | 0 | 0 | 130,736 | 1222,61331 | 83 | 39278,096 | 0 | 1 | 0 | 0 | 1 | 0 | 0 | 0 | 0 | 0 | 0 | 0 | 0 | 0 | 0 | 0 | 0 |
| 48.18102 | 17.27939 | BRA_R_1 | brar1_5 | Slovakia | R | 0 | 0 | 0 | 0 | 0 | 0 | 0 | 130 | 1222,542333 | 83 | 22271,637 | 0 | 1 | 1 | 0 | 1 | 0 | 0 | 0 | 0 | 0 | 0 | 0 | 0 | 0 | 0 | 0 | 0 |
| 48.19575 | 17.31447 | BRA_R_2 | brar2_1 | Slovakia | R | 1 | 1 | 1 | 1 | 0 | 0 | 0 | 129 | 1221,382555 | 87 | 26900,96 | 10108 | 1 | 1 | 0 | 1 | 0 | 0 | 0 | 0 | 1 | 1 | 0 | 1 | 0 | 0 | 0 | 0 |
| 48.19539 | 17.31869 | BRA_R_2 | brar2_2 | Slovakia | R | 0 | 0 | 0 | 0 | 0 | 0 | 0 | 127,885 | 1221,105149 | 87 | 14793,245 | 12684 | 0 | 1 | 0 | 0 | 0 | 0 | 1 | 0 | 0 | 0 | 0 | 1 | 0 | 0 | 0 | 0 |
| 48.19685 | 17.3216 | BRA_R_2 | brar2_3 | Slovakia | R | 1 | 1 | 0 | 0 | 0 | 0 | 0 | 128 | 1221,02487 | 87 | 29440,013 | 10056 | 0 | 1 | 0 | 0 | 0 | 0 | 1 | 0 | 0 | 0 | 0 | 1 | 0 | 0 | 0 | 0 |
| 48.1983 | 17.3247 | BRA_R_2 | brar2_4 | Slovakia | R | 0 | 0 | 1 | 1 | 0 | 0 | 0 | 128,07 | 1220,932848 | 87 | 15386,133 | 12239 | 0 | 1 | 0 | 0 | 0 | 0 | 1 | 0 | 0 | 0 | 0 | 1 | 0 | 0 | 0 | 0 |
| 48.20036 | 17.32499 | BRA_R_2 | brar2_5 | Slovakia | R | 0 | 0 | 0 | 0 | 0 | 0 | 0 | 128,296 | 1221,049653 | 87 | 20392,002 | 9362 | 0 | 1 | 0 | 0 | 0 | 0 | 1 | 0 | 0 | 0 | 0 | 1 | 0 | 0 | 1 | 0 |
| 48.22558 | 17.38937 | BRA_R_3 | brar3_1 | Slovak | R | 0 | 0 | 0 | 0 | 0 | 0 | 0 | 140,088 | 1218,832215 | 92 | 5040,507 | 4033 | 0 | 1 | 0 | 1 | 0 | 0 | 0 | 0 | 1 | 0 | 1 | 0 | 0 | 0 | 0 | 0 |
| 48.22312 | 17.391 | BRA_R_3 | brar3_2 | Slovak | R | 0 | 0 | 1 | 1 | 0 | 0 | 0 | 139,768 | 1218,572562 | 92 | 5194,756 | 5796 | 0 | 1 | 0 | 1 | 0 | 0 | 0 | 0 | 1 | 0 | 0 | 1 | 0 | 0 | 0 | 0 |
| 48.22073 | 17.39281 | BRA_R_3 | brar3_3 | Slovak | R | 0 | 0 | 0 | 0 | 0 | 0 | 0 | 135,256 | 1218,306484 | 92 | 4252,811 | 13267 | 0 | 1 | 0 | 0 | 0 | 1 | 0 | 1 | 1 | 0 | 0 | 1 | 1 | 0 | 0 | 0 |
| 48.2184 | 17.39294 | BRA_R_3 | brar3_4 | Slovak | R | 0 | 0 | 0 | 0 | 0 | 0 | 0 | 124,9 | 1218,141347 | 92 | 7534,676 | 10754 | 0 | 1 | 0 | 0 | 0 | 1 | 0 | 1 | 1 | 0 | 0 | 1 | 1 | 0 | 0 | 0 |
| 48.2158 | 17.39247 | BRA_R_3 | brar3_5 | Slovak | R | 0 | 0 | 0 | 0 | 0 | 0 | 0 | 125 | 1217,999664 | 92 | 6378,674 | 15449 | 0 | 1 | 0 | 0 | 0 | 0 | 0 | 0 | 1 | 0 | 0 | 1 | 0 | 0 | 0 | 0 |
| 48.22066 | 17.41841 | BRA_R_4 | brar4_1 | Slovak | R | 0 | 0 | 1 | 1 | 0 | 0 | 0 | 126 | 1216,762664 | 93 | 6661,337 | 8850 | 0 | 1 | 0 | 0 | 0 | 0 | 1 | 1 | 1 | 0 | 0 | 1 | 0 | 1 | 0 | 0 |
| 48.21931 | 17.41478 | BRA_R_4 | brar4_2 | Slovak | R | 0 | 0 | 0 | 0 | 0 | 0 | 0 | 123,347 | 1216,887348 | 93 | 1265,239 | 12644 | 0 | 1 | 0 | 0 | 0 | 0 | 0 | 1 | 1 | 0 | 0 | 1 | 1 | 0 | 0 | 0 |
| 48.21777 | 17.41139 | BRA_R_4 | brar4_3 | Slovak | R | 0 | 0 | 0 | 0 | 0 | 0 | 0 | 125 | 1216,994579 | 93 | 3243,554 | 11001 | 0 | 1 | 0 | 0 | 0 | 0 | 0 | 0 | 1 | 0 | 0 | 1 | 0 | 0 | 1 | 0 |
| 48.21652 | 17.40786 | BRA_R_4 | brar4_4 | Slovak | R | 0 | 0 | 0 | 0 | 0 | 0 | 0 | 124,548 | 1217,137266 | 93 | 7962,323 | 12987 | 0 | 1 | 0 | 0 | 1 | 0 | 1 | 0 | 1 | 0 | 0 | 1 | 0 | 0 | 0 | 0 |
| 48.21439 | 17.40998 | BRA_R_4 | brar4_5 | Slovak | R | 0 | 0 | 1 | 1 | 0 | 0 | 0 | 123 | 1216,859286 | 93 | 7420,801 | 8888 | 0 | 1 | 0 | 0 | 1 | 0 | 0 | 1 | 1 | 1 | 0 | 1 | 1 | 0 | 1 | 0 |
| 48.33884 | 17.31263 | BRA_R_5 | brar5_1 | Slovak | R | 0 | 0 | 1 | 1 | 0 | 0 | 1 | 200,614 | 1230,865207 | 88 | 27838,652 | 6571 | 0 | 1 | 0 | 1 | 0 | 0 | 0 | 0 | 1 | 0 | 1 | 1 | 0 | 1 | 0 | 0 |
| 48.3366 | 17.31298 | BRA_R_5 | brar5_2 | Slovak | R | 0 | 0 | 2 | 1 | 0 | 0 | 1 | 180,073 | 1230,706321 | 88 | 20162,175 | 12690 | 0 | 1 | 0 | 1 | 0 | 0 | 0 | 0 | 1 | 0 | 0 | 0 | 0 | 1 | 0 | 0 |
| 48.33391 | 17.31329 | BRA_R_5 | brar5_3 | Slovak | R | 0 | 0 | 1 | 1 | 0 | 0 | 0 | 169,076 | 1230,520555 | 88 | 11562,833 | 18168 | 0 | 1 | 0 | 1 | 0 | 0 | 0 | 1 | 1 | 0 | 0 | 0 | 1 | 1 | 0 | 0 |
| 48.3313 | 17.31423 | BRA_R_5 | brar5_4 | Slovak | R | 0 | 0 | 0 | 0 | 0 | 0 | 1 | 163,525 | 1230,280615 | 88 | 7134,498 | 8771 | 0 | 1 | 0 | 1 | 0 | 0 | 1 | 0 | 1 | 0 | 1 | 0 | 0 | 1 | 0 | 0 |
| 48.33017 | 17.31676 | BRA_R_5 | brar5_5 | Slovak | R | 0 | 0 | 0 | 0 | 0 | 0 | 1 | 159,612 | 1230,063266 | 88 | 35660,376 | 1094 | 1 | 1 | 0 | 1 | 0 | 0 | 0 | 0 | 0 | 0 | 1 | 0 | 0 | 0 | 0 | 0 |
| 48.33377 | 17.29424 | BRA_R_6 | brar6_1 | Slovak | R | 1 | 1 | 0 | 0 | 0 | 0 | 0 | 208,794 | 1231,643969 | 87 | 21708,984 | 333 | 1 | 1 | 0 | 1 | 0 | 0 | 1 | 0 | 0 | 0 | 1 | 0 | 0 | 0 | 0 | 1 |
| 48.33298 | 17.2981 | BRA_R_6 | brar6_2 | Slovak | R | 0 | 0 | 0 | 0 | 0 | 0 | 0 | 194,568 | 1231,360235 | 87 | 4589,47 | 798 | 0 | 1 | 0 | 1 | 0 | 0 | 1 | 0 | 1 | 0 | 1 | 0 | 0 | 0 | 0 | 0 |
| 48.33507 | 17.29885 | BRA_R_6 | brar6_3 | Slovak | R | 0 | 0 | 2 | 1 | 0 | 0 | 0 | 197,343 | 1231,448933 | 87 | 5433,044 | 1895 | 0 | 1 | 0 | 1 | 0 | 0 | 0 | 0 | 1 | 0 | 1 | 1 | 0 | 0 | 0 | 0 |
| 48.33763 | 17.29919 | BRA_R_6 | brar6_4 | Slovak | R | 0 | 0 | 3 | 1 | 0 | 0 | 0 | 195,468 | 1231,593241 | 87 | 10796,982 | 6216 | 1 | 1 | 0 | 1 | 0 | 0 | 0 | 0 | 1 | 0 | 1 | 1 | 0 | 0 | 0 | 0 |
| 48.33924 | 17.29604 | BRA_R_6 | brar6_5 | Slovak | R | 0 | 0 | 0 | 0 | 0 | 0 | 1 | 210,154 | 1231,900465 | 87 | 18703,949 | 685 | 1 | 1 | 0 | 1 | 0 | 0 | 0 | 0 | 1 | 0 | 1 | 0 | 0 | 0 | 0 | 1 |
| 48.16188 | 17.16879 | BRA_U_1 | brau1_1 | Slovakia | U | 0 | 0 | 1 | 1 | 0 | 0 | 0 | 134 | 1228,004878 | 75 | 28156,792 | 8794 | 0 | 1 | 0 | 0 | 1 | 0 | 1 | 1 | 0 | 0 | 0 | 0 | 1 | 0 | 0 | 0 |
| 48.16427 | 17.16713 | BRA_U_1 | brau1_2 | Slovakia | U | 1 | 1 | 1 | 1 | 0 | 0 | 0 | 135 | 1228,259324 | 75 | 28043,772 | 14121 | 0 | 1 | 0 | 0 | 1 | 0 | 1 | 1 | 0 | 0 | 0 | 1 | 1 | 0 | 1 | 0 |
| 48.16472 | 17.16328 | BRA_U_1 | brau1_3 | Slovakia | U | 1 | 1 | 1 | 1 | 0 | 0 | 0 | 135 | 1228,52144 | 75 | 35552,216 | 10923 | 1 | 1 | 0 | 0 | 1 | 1 | 1 | 1 | 1 | 0 | 0 | 1 | 1 | 0 | 1 | 0 |
| 48.16274 | 17.16124 | BRA_U_1 | brau1_4 | Slovakia | U | 0 | 0 | 0 | 0 | 0 | 0 | 0 | 135 | 1228,518205 | 75 | 53657,765 | 5080 | 1 | 1 | 0 | 0 | 0 | 1 | 0 | 1 | 0 | 0 | 0 | 0 | 1 | 0 | 1 | 0 |
| 48.16013 | 17.16125 | BRA_U_1 | brau1_5 | Slovakia | U | 0 | 0 | 1 | 1 | 0 | 0 | 0 | 133,753 | 1228,350435 | 75 | 35021,372 | 10416 | 1 | 1 | 0 | 0 | 0 | 1 | 0 | 1 | 0 | 0 | 0 | 0 | 1 | 0 | 0 | 0 |
| 48.14719 | 17.19214 | BRA_U_2 | brau2_1 | Slovakia | U | 0 | 0 | 1 | 1 | 0 | 0 | 0 | 132 | 1225,642808 | 77 | 51286,722 | 6082 | 1 | 1 | 0 | 1 | 0 | 0 | 0 | 0 | 1 | 0 | 0 | 1 | 0 | 0 | 0 | 0 |
| 48.14646 | 17.19538 | BRA_U_2 | brau2_2 | Slovakia | U | 0 | 0 | 0 | 0 | 0 | 0 | 0 | 132 | 1225,398908 | 77 | 50312,243 | 3005 | 1 | 1 | 0 | 0 | 1 | 1 | 0 | 0 | 1 | 0 | 1 | 1 | 0 | 0 | 0 | 0 |
| 48.1458 | 17.19832 | BRA_U_2 | brau2_3 | Slovakia | U | 1 | 1 | 1 | 1 | 0 | 0 | 1 | 131 | 1225,178005 | 77 | 53186,804 | 2135 | 1 | 1 | 1 | 0 | 1 | 1 | 0 | 1 | 1 | 1 | 0 | 1 | 0 | 0 | 0 | 0 |
| 48.14596 | 17.2021 | BRA_U_2 | brau2_4 | Slovakia | U | 0 | 0 | 0 | 0 | 0 | 0 | 0 | 131,799 | 1224,958918 | 77 | 39278,096 | 4195 | 1 | 1 | 1 | 0 | 1 | 0 | 1 | 0 | 1 | 1 | 0 | 1 | 0 | 0 | 0 | 0 |
| 48.14798 | 17.20483 | BRA_U_2 | brau2_5 | Slovakia | U | 0 | 0 | 1 | 1 | 0 | 0 | 0 | 132 | 1224,922984 | 77 | 49172,65 | 6802 | 1 | 1 | 0 | 0 | 1 | 0 | 1 | 0 | 1 | 1 | 0 | 1 | 0 | 0 | 0 | 0 |
| 48.16014 | 17.08958 | BRA_U_3 | brau3_1 | Slovak | U | 2 | 1 | 0 | 0 | 0 | 0 | 0 | 194,746 | 1232,701316 | 69 | 59534,449 | 7683 | 1 | 0 | 0 | 0 | 1 | 0 | 0 | 0 | 0 | 0 | 0 | 0 | 1 | 0 | 0 | 0 |
| 48.15866 | 17.0929 | BRA_U_3 | brau3_2 | Slovak | U | 0 | 0 | 0 | 0 | 0 | 0 | 0 | 206,89 | 1232,418442 | 69 | 60654,808 | 2539 | 1 | 1 | 0 | 0 | 1 | 0 | 0 | 1 | 0 | 0 | 1 | 0 | 0 | 0 | 0 | 0 |
| 48.15732 | 17.09522 | BRA_U_3 | brau3_3 | Slovak | U | 1 | 1 | 0 | 0 | 0 | 0 | 0 | 222,381 | 1232,185474 | 69 | 54614,52 | 6606 | 1 | 1 | 1 | 0 | 1 | 0 | 0 | 1 | 0 | 0 | 1 | 1 | 0 | 0 | 0 | 0 |
| 48.15806 | 17.09869 | BRA_U_3 | brau3_4 | Slovak | U | 0 | 0 | 0 | 0 | 0 | 0 | 0 | 230,273 | 1232,024335 | 69 | 32569,622 | 12336 | 1 | 1 | 0 | 0 | 0 | 0 | 0 | 1 | 1 | 0 | 0 | 1 | 1 | 0 | 0 | 0 |
| 48.15945 | 17.10056 | BRA_U_3 | brau3_5 | Slovak | U | 0 | 0 | 0 | 0 | 0 | 0 | 0 | 235,876 | 1231,995347 | 69 | 39930,065 | 8904 | 1 | 1 | 1 | 0 | 1 | 0 | 0 | 1 | 1 | 0 | 0 | 1 | 0 | 0 | 0 | 0 |
| 48.156 | 17.05954 | BRA_U_4 | brau4_1 | Slovak | U | 0 | 0 | 0 | 0 | 0 | 0 | 0 | 157,282 | 1234,273869 | 66 | 26703,723 | 9739 | 1 | 1 | 0 | 0 | 0 | 0 | 0 | 1 | 1 | 0 | 0 | 1 | 1 | 0 | 0 | 0 |
| 48.1582 | 17.05724 | BRA_U_4 | brau4_2 | Slovak | U | 2 | 1 | 0 | 0 | 0 | 0 | 0 | 163,693 | 1234,553122 | 66 | 30386,556 | 11062 | 1 | 1 | 0 | 0 | 0 | 0 | 0 | 1 | 1 | 0 | 0 | 1 | 1 | 0 | 0 | 1 |
| 48.16038 | 17.05499 | BRA_U_4 | brau4_3 | Slovak | U | 0 | 0 | 1 | 1 | 0 | 0 | 0 | 166,668 | 1234,830483 | 66 | 32400,716 | 11830 | 1 | 1 | 1 | 1 | 0 | 0 | 0 | 1 | 1 | 0 | 0 | 1 | 1 | 0 | 0 | 1 |
| 48.16281 | 17.05394 | BRA_U_4 | brau4_4 | Slovak | U | 0 | 0 | 0 | 0 | 0 | 0 | 0 | 171,48 | 1235,052468 | 66 | 63091,969 | 3905 | 1 | 0 | 0 | 1 | 0 | 0 | 0 | 0 | 1 | 0 | 1 | 0 | 1 | 0 | 0 | 0 |
| 48.16516 | 17.05193 | BRA_U_4 | brau4_5 | Slovak | U | 0 | 0 | 0 | 0 | 0 | 0 | 0 | 174,65 | 1235,323277 | 66 | 61741,395 | 2392 | 1 | 1 | 0 | 1 | 0 | 0 | 0 | 1 | 0 | 0 | 1 | 0 | 1 | 0 | 0 | 0 |
| 48.13583 | 17.11857 | BRA_U_5 | brau5_1 | Slovak | U | 0 | 0 | 0 | 0 | 0 | 0 | 0 | 137,988 | 1229,391118 | 71 | 39343,225 | 1527 | 1 | 1 | 0 | 0 | 1 | 0 | 0 | 1 | 0 | 0 | 1 | 0 | 0 | 0 | 0 | 1 |
| 48.13523 | 17.11513 | BRA_U_5 | brau5_2 | Slovak | U | 1 | 1 | 0 | 0 | 0 | 0 | 0 | 138,28 | 1229,561587 | 71 | 29000,242 | 3539 | 0 | 1 | 0 | 0 | 1 | 0 | 0 | 1 | 0 | 0 | 1 | 0 | 0 | 0 | 0 | 0 |
| 48.13351 | 17.1137 | BRA_U_5 | brau5_3 | Slovak | U | 0 | 0 | 0 | 0 | 0 | 0 | 0 | 135,432 | 1229,535814 | 71 | 53691,207 | 758 | 1 | 1 | 0 | 0 | 1 | 0 | 0 | 1 | 0 | 0 | 0 | 0 | 1 | 0 | 0 | 1 |
| 48.13339 | 17.11017 | BRA_U_5 | brau5_4 | Slovak | U | 0 | 0 | 0 | 0 | 0 | 0 | 0 | 135,309 | 1229,749039 | 71 | 44018,158 | 16331 | 1 | 1 | 1 | 0 | 1 | 0 | 0 | 0 | 0 | 0 | 0 | 0 | 0 | 1 | 1 | 0 |
| 48.13513 | 17.10711 | BRA_U_5 | brau5_5 | Slovak | U | 0 | 0 | 0 | 0 | 0 | 0 | 0 | 135,721 | 1230,042376 | 71 | 53628,042 | 3239 | 0 | 1 | 1 | 0 | 1 | 0 | 0 | 0 | 0 | 0 | 0 | 0 | 1 | 1 | 0 | 1 |
| 48.11598 | 17.1193 | BRA_U_6 | brau6_1 | Slovak | U | 1 | 1 | 0 | 0 | 0 | 0 | 0 | 134,205 | 1228,077625 | 71 | 26126,834 | 13773 | 1 | 1 | 0 | 0 | 0 | 0 | 0 | 1 | 0 | 0 | 0 | 0 | 1 | 0 | 0 | 1 |
| 48.11476 | 17.11578 | BRA_U_6 | brau6_2 | Slovak | U | 1 | 1 | 1 | 1 | 0 | 0 | 0 | 134 | 1228,219309 | 71 | 40018,663 | 7099 | 1 | 1 | 0 | 0 | 0 | 0 | 0 | 1 | 0 | 0 | 0 | 0 | 1 | 0 | 0 | 0 |
| 48.11326 | 17.1136 | BRA_U_6 | brau6_3 | Slovak | U | 3 | 1 | 0 | 0 | 0 | 0 | 0 | 133 | 1228,257294 | 71 | 35677,301 | 9293 | 1 | 1 | 0 | 0 | 0 | 0 | 0 | 1 | 0 | 0 | 0 | 0 | 1 | 0 | 0 | 0 |
| 48.11116 | 17.11229 | BRA_U_6 | brau6_4 | Slovak | U | 0 | 0 | 0 | 0 | 0 | 0 | 0 | 135 | 1228,199659 | 71 | 17937,544 | 18682 | 1 | 1 | 0 | 0 | 0 | 0 | 0 | 1 | 0 | 0 | 0 | 0 | 1 | 0 | 1 | 0 |
| 48.10967 | 17.11548 | BRA_U_6 | brau6_5 | Slovak | U | 1 | 1 | 0 | 0 | 0 | 0 | 0 | 134,409 | 1227,909625 | 71 | 44187,94 | 5217 | 1 | 1 | 0 | 0 | 0 | 0 | 0 | 1 | 0 | 0 | 0 | 0 | 1 | 0 | 0 | 0 |
| 48.21958 | 17.39642 | BRA_U_9 | brau9_1 | Slovakia | U | 0 | 0 | 1 | 1 | 0 | 0 | 0 | 128 | 1218,010392 | 94 | 7623,673 | 19197 | 0 | 1 | 0 | 0 | 0 | 0 | 1 | 1 | 0 | 0 | 0 | 0 | 1 | 1 | 0 | 0 |
| 48.22329 | 17.4038 | BRA_U_9 | brau9_2 | Slovakia | U | 0 | 0 | 0 | 0 | 0 | 0 | 0 | 125,05 | 1217,813656 | 94 | 1257,661 | 16791 | 0 | 1 | 0 | 0 | 0 | 0 | 0 | 0 | 0 | 0 | 1 | 1 | 0 | 1 | 0 | 0 |
| 48.22422 | 17.41017 | BRA_U_9 | brau9_3 | Slovakia | U | 0 | 0 | 1 | 1 | 0 | 0 | 0 | 125 | 1217,493933 | 94 | 824,939 | 13678 | 0 | 1 | 0 | 0 | 0 | 1 | 0 | 0 | 1 | 0 | 0 | 1 | 0 | 0 | 1 | 0 |
| 48.2197 | 17.41142 | BRA_U_9 | brau9_4 | Slovakia | U | 0 | 0 | 1 | 1 | 0 | 0 | 1 | 124,183 | 1217,121426 | 94 | 728,411 | 15829 | 0 | 1 | 0 | 0 | 0 | 0 | 0 | 0 | 1 | 0 | 0 | 1 | 0 | 0 | 0 | 0 |
| 48.22048 | 17.41693 | BRA_U_9 | brau9_5 | Slovakia | U | 0 | 0 | 0 | 0 | 0 | 0 | 0 | 125,272 | 1216,841979 | 94 | 2152,857 | 14681 | 0 | 1 | 0 | 0 | 0 | 0 | 1 | 0 | 0 | 0 | 0 | 1 | 0 | 0 | 1 | 0 |
| 49.23133 | 16.65725 | BRN_R_1 | brnr1_1 | Czech | R | 1 | 1 | 0 | 0 | 0 | 0 | 0 | 261,718 | 1329,335362 | 100 | 32781,219 | 3534 | 1 | 1 | 0 | 1 | 0 | 0 | 0 | 0 | 1 | 0 | 0 | 1 | 0 | 0 | 0 | 0 |
| 49.22843 | 16.65882 | BRN_R_1 | brnr1_2 | Czech | R | 0 | 0 | 0 | 0 | 0 | 0 | 0 | 231,235 | 1329,04061 | 100 | 48500,247 | 3411 | 1 | 1 | 0 | 1 | 0 | 0 | 1 | 0 | 1 | 1 | 0 | 1 | 0 | 0 | 0 | 0 |
| 49.22593 | 16.66186 | BRN_R_1 | brnr1_3 | Czech | R | 2 | 1 | 0 | 0 | 0 | 0 | 0 | 216,088 | 1328,698597 | 100 | 45577,592 | 405 | 1 | 1 | 0 | 1 | 0 | 0 | 0 | 0 | 1 | 1 | 1 | 0 | 0 | 0 | 1 | 0 |
| 49.22693 | 16.6656 | BRN_R_1 | brnr1_4 | Czech | R | 1 | 1 | 0 | 0 | 0 | 0 | 0 | 223,16 | 1328,553775 | 100 | 58699,672 | 0 | 1 | 1 | 0 | 1 | 0 | 0 | 1 | 0 | 1 | 0 | 1 | 0 | 0 | 0 | 1 | 0 |
| 49.22939 | 16.66659 | BRN_R_1 | brnr1_5 | Czech | R | 0 | 0 | 0 | 0 | 0 | 0 | 0 | 223,993 | 1328,664779 | 100 | 62059,683 | 553 | 1 | 0 | 0 | 1 | 0 | 0 | 1 | 0 | 0 | 0 | 0 | 0 | 0 | 0 | 1 | 0 |
| 49.22817 | 16.53683 | BRN_R_2 | brnr2_1 | Czech | R | 0 | 0 | 0 | 0 | 0 | 0 | 0 | 217,412 | 1336,028359 | 90 | 14802,289 | 7513 | 0 | 1 | 0 | 1 | 0 | 0 | 0 | 1 | 1 | 0 | 0 | 1 | 0 | 0 | 1 | 1 |
| 49.2289 | 16.54164 | BRN_R_2 | brnr2_2 | Czech | R | 2 | 1 | 0 | 0 | 0 | 0 | 0 | 222,048 | 1335,803215 | 90 | 20788,252 | 7352 | 0 | 1 | 0 | 1 | 0 | 0 | 1 | 0 | 1 | 0 | 1 | 1 | 0 | 0 | 1 | 0 |
| 49.2311 | 16.54027 | BRN_R_2 | brnr2_3 | Czech | R | 0 | 0 | 0 | 0 | 0 | 0 | 0 | 239,049 | 1336,018544 | 90 | 27741,553 | 710 | 1 | 1 | 1 | 1 | 0 | 0 | 0 | 0 | 1 | 0 | 1 | 0 | 0 | 0 | 0 | 0 |
| 49.23305 | 16.53835 | BRN_R_2 | brnr2_4 | Czech | R | 1 | 1 | 0 | 0 | 0 | 0 | 0 | 262,956 | 1336,277652 | 90 | 22741,681 | 203 | 1 | 1 | 0 | 1 | 0 | 0 | 0 | 0 | 1 | 0 | 1 | 0 | 0 | 0 | 0 | 1 |

| | | | | | | | | | | | | | | | | | | | | | | | | | | | | | | | | | |
|---|---|---|---|---|---|---|---|---|---|---|---|---|---|---|---|---|---|---|---|---|---|---|---|---|---|---|---|---|---|---|---|---|---|
| 49.23583 | 16.53832 | BRN_R_2 | brnr2_5 | Czech | R | 0 | 0 | 0 | 0 | 0 | 0 | 0 | 283,108 | 1336,468229 | 90 | 23616,96 | 1079 | 1 | 1 | 0 | 1 | 0 | 0 | 1 | 0 | 1 | 0 | 1 | 0 | 0 | 0 | 0 | 0 |
| 49.23892 | 16.64435 | BRN_R_3 | brnr3_1 | Czech | R | 0 | 0 | 0 | 0 | 0 | 0 | 0 | 245,674 | 1330,594075 | 100 | 24103,937 | 5439 | 1 | 1 | 0 | 1 | 0 | 0 | 0 | 1 | 1 | 0 | 1 | 0 | 1 | 0 | 0 | 0 |
| 49.23774 | 16.64834 | BRN_R_3 | brnr3_2 | Czech | R | 0 | 0 | 0 | 0 | 0 | 0 | 0 | 249,955 | 1330,28985 | 100 | 24759,999 | 4315 | 1 | 1 | 0 | 1 | 0 | 0 | 1 | 0 | 0 | 0 | 0 | 1 | 1 | 0 | 0 | 0 |
| 49.23817 | 16.6521 | BRN_R_3 | brnr3_3 | Czech | R | 0 | 0 | 0 | 0 | 0 | 0 | 0 | 269,081 | 1330,097771 | 100 | 20191,27 | 341 | 0 | 1 | 0 | 1 | 0 | 0 | 1 | 0 | 1 | 0 | 1 | 0 | 0 | 0 | 0 | 0 |
| 49.23946 | 16.6542 | BRN_R_3 | brnr3_4 | Czech | R | 0 | 0 | 0 | 0 | 0 | 0 | 0 | 278,424 | 1330,065755 | 100 | 18137,897 | 0 | 1 | 1 | 0 | 1 | 0 | 0 | 1 | 0 | 0 | 0 | 0 | 0 | 0 | 0 | 0 | 0 |
| 49.24187 | 16.65482 | BRN_R_3 | brnr3_5 | Czech | R | 0 | 0 | 0 | 0 | 0 | 0 | 0 | 272,038 | 1330,198259 | 100 | 23506,801 | 1473 | 1 | 1 | 0 | 1 | 0 | 0 | 1 | 1 | 0 | 0 | 1 | 0 | 0 | 0 | 0 | 0 |
| 49.24053 | 16.54177 | BRN_R_4 | brnr4_1 | Czech | R | 0 | 0 | 0 | 0 | 0 | 0 | 1 | 294,864 | 1336,591932 | 90 | 24434,379 | 396 | 1 | 1 | 0 | 1 | 0 | 0 | 0 | 0 | 1 | 0 | 1 | 0 | 0 | 0 | 0 | 0 |
| 49.2414 | 16.53769 | BRN_R_4 | brnr4_2 | Czech | R | 1 | 1 | 0 | 0 | 0 | 0 | 0 | 275,74 | 1336,879664 | 90 | 68325,767 | 122 | 1 | 1 | 0 | 1 | 0 | 0 | 0 | 0 | 1 | 0 | 0 | 0 | 0 | 0 | 0 | 0 |
| 49.24352 | 16.53409 | BRN_R_4 | brnr4_3 | Czech | R | 3 | 1 | 0 | 0 | 0 | 0 | 0 | 246,156 | 1337,189178 | 90 | 57011,194 | 0 | 1 | 0 | 0 | 1 | 0 | 0 | 0 | 0 | 0 | 0 | 0 | 0 | 0 | 0 | 0 | 0 |
| 49.2454 | 16.53137 | BRN_R_4 | brnr4_4 | Czech | R | 1 | 1 | 0 | 0 | 0 | 0 | 0 | 253,002 | 1337,502411 | 90 | 49506,546 | 0 | 1 | 0 | 0 | 1 | 0 | 0 | 0 | 0 | 0 | 0 | 0 | 0 | 0 | 0 | 0 | 0 |
| 49.24779 | 16.53149 | BRN_R_4 | brnr4_5 | Czech | R | 1 | 1 | 0 | 0 | 0 | 0 | 0 | 259,584 | 1337,677435 | 90 | 60818,752 | 0 | 1 | 0 | 0 | 1 | 0 | 0 | 0 | 0 | 0 | 0 | 0 | 0 | 0 | 0 | 0 | 0 |
| 49.14618 | 16.55357 | BRN_R_5 | brnr5_1 | Czech | R | 1 | 1 | 0 | 0 | 0 | 0 | 1 | 313,677 | 1329,501463 | 93 | 14583,213 | 0 | 1 | 0 | 0 | 1 | 0 | 0 | 0 | 0 | 0 | 0 | 0 | 0 | 0 | 0 | 0 | 0 |
| 49.14831 | 16.54976 | BRN_R_5 | brnr5_2 | Czech | R | 1 | 1 | 2 | 1 | 1 | 1 | 0 | 308,884 | 1329,84587 | 93 | 9667,162 | 0 | 1 | 0 | 0 | 1 | 0 | 0 | 1 | 0 | 0 | 0 | 0 | 0 | 0 | 0 | 0 | 0 |
| 49.15015 | 16.5469 | BRN_R_5 | brnr5_3 | Czech | R | 3 | 1 | 2 | 1 | 1 | 1 | 1 | 310,614 | 1330,155119 | 93 | 10651,976 | 605 | 0 | 1 | 0 | 1 | 0 | 0 | 1 | 0 | 1 | 0 | 1 | 0 | 0 | 0 | 0 | 0 |
| 49.15307 | 16.54474 | BRN_R_5 | brnr5_4 | Czech | R | 1 | 1 | 0 | 0 | 0 | 0 | 1 | 305,997 | 1330,471545 | 93 | 44197,846 | 623 | 1 | 1 | 0 | 1 | 0 | 0 | 0 | 0 | 1 | 0 | 1 | 0 | 0 | 0 | 0 | 0 |
| 49.15554 | 16.54306 | BRN_R_5 | brnr5_5 | Czech | R | 1 | 1 | 0 | 0 | 0 | 0 | 0 | 301,305 | 1330,744426 | 93 | 21477,026 | 308 | 1 | 1 | 1 | 1 | 0 | 0 | 0 | 0 | 1 | 0 | 1 | 0 | 0 | 0 | 0 | 0 |
| 49.12822 | 16.58415 | BRN_R_6 | brnr6_1 | Czech | R | 0 | 0 | 0 | 0 | 0 | 0 | 1 | 307,036 | 1326,506977 | 97 | 17014,92 | 0 | 0 | 1 | 0 | 0 | 0 | 0 | 0 | 0 | 1 | 0 | 0 | 0 | 0 | 0 | 0 | 0 |
| 49.12656 | 16.5878 | BRN_R_6 | brnr6_2 | Czech | R | 0 | 0 | 2 | 1 | 0 | 0 | 1 | 293,536 | 1326,184655 | 97 | 29897,057 | 0 | 1 | 1 | 0 | 1 | 0 | 0 | 0 | 0 | 1 | 0 | 0 | 0 | 0 | 0 | 0 | 0 |
| 49.12658 | 16.59254 | BRN_R_6 | brnr6_3 | Czech | R | 0 | 0 | 0 | 0 | 0 | 0 | 1 | 270,48 | 1325,90868 | 97 | 22019,661 | 16 | 1 | 1 | 0 | 1 | 0 | 0 | 0 | 0 | 1 | 0 | 0 | 0 | 0 | 0 | 0 | 0 |
| 49.12718 | 16.59689 | BRN_R_6 | brnr6_4 | Czech | R | 0 | 0 | 0 | 0 | 0 | 0 | 1 | 248,196 | 1325,699455 | 97 | 21332,836 | 17 | 0 | 1 | 0 | 0 | 0 | 0 | 0 | 0 | 1 | 0 | 1 | 0 | 0 | 0 | 0 | 1 |
| 49.13063 | 16.59754 | BRN_R_6 | brnr6_5 | Czech | R | 0 | 0 | 1 | 1 | 0 | 0 | 0 | 230,091 | 1325,896931 | 97 | 11208,696 | 1057 | 0 | 1 | 0 | 0 | 0 | 0 | 0 | 0 | 1 | 0 | 1 | 0 | 1 | 0 | 0 | 0 |
| 49.11811 | 16.57786 | BRN_R_7 | brnr7_1 | Czech | R | 0 | 0 | 0 | 0 | 0 | 0 | 0 | 225,026 | 1326,331821 | 96 | 18334,672 | 7744 | 1 | 1 | 0 | 1 | 0 | 0 | 1 | 1 | 1 | 0 | 1 | 1 | 1 | 0 | 0 | 0 |
| 49.12082 | 16.57853 | BRN_R_7 | brnr7_2 | Czech | R | 0 | 0 | 2 | 1 | 0 | 0 | 0 | 243,57 | 1326,1852 | 96 | 13049,634 | 0 | 0 | 1 | 0 | 1 | 0 | 0 | 1 | 0 | 1 | 0 | 1 | 0 | 0 | 0 | 0 | 0 |
| 49.11964 | 16.58247 | BRN_R_7 | brnr7_3 | Czech | R | 0 | 0 | 0 | 0 | 0 | 0 | 0 | 229,22 | 1326,03624 | 96 | 16148,338 | 3310 | 0 | 1 | 0 | 0 | 0 | 0 | 0 | 1 | 1 | 0 | 1 | 1 | 1 | 0 | 0 | 0 |
| 49.11728 | 16.58144 | BRN_R_7 | brnr7_4 | Czech | R | 0 | 0 | 0 | 0 | 0 | 0 | 0 | 208,624 | 1325,924167 | 96 | 6439,955 | 14566 | 0 | 1 | 0 | 0 | 0 | 0 | 0 | 0 | 1 | 0 | 0 | 1 | 0 | 1 | 0 | 1 |
| 49.11512 | 16.58125 | BRN_R_7 | brnr7_5 | Czech | R | 0 | 0 | 0 | 0 | 0 | 0 | 0 | 201,568 | 1325,803003 | 96 | 12746,506 | 6389 | 0 | 1 | 0 | 0 | 0 | 0 | 0 | 1 | 1 | 1 | 0 | 1 | 0 | 0 | 0 | 0 |
| 49.20585 | 16.60677 | BRN_U_1 | brnu1_1 | Czech | U | 0 | 0 | 0 | 0 | 0 | 0 | 0 | 214,613 | 1330,485919 | 96 | 43501,994 | 6137 | 1 | 1 | 0 | 0 | 1 | 0 | 0 | 0 | 0 | 0 | 0 | 0 | 0 | 1 | 0 | 0 |
| 49.20648 | 16.61067 | BRN_U_1 | brnu1_2 | Czech | U | 2 | 1 | 2 | 1 | 0 | 0 | 0 | 206 | 1330,304753 | 96 | 37114,464 | 4174 | 1 | 1 | 0 | 0 | 1 | 0 | 0 | 0 | 0 | 0 | 0 | 0 | 0 | 1 | 0 | 0 |
| 49.20735 | 16.61431 | BRN_U_1 | brnu1_3 | Czech | U | 0 | 0 | 0 | 0 | 0 | 0 | 0 | 226,617 | 1330,156785 | 96 | 24888,557 | 10962 | 0 | 1 | 1 | 0 | 1 | 0 | 0 | 1 | 1 | 0 | 0 | 0 | 1 | 1 | 0 | 0 |
| 49.20957 | 16.61327 | BRN_U_1 | brnu1_4 | Czech | U | 0 | 0 | 0 | 0 | 0 | 0 | 0 | 232,573 | 1330,366734 | 96 | 11416,395 | 20438 | 0 | 1 | 0 | 0 | 0 | 0 | 1 | 1 | 1 | 0 | 0 | 0 | 1 | 1 | 0 | 0 |
| 49.21146 | 16.61287 | BRN_U_1 | brnu1_5 | Czech | U | 1 | 1 | 2 | 1 | 0 | 0 | 0 | 219,655 | 1330,517298 | 96 | 20172,29 | 11280 | 0 | 1 | 0 | 0 | 1 | 0 | 1 | 1 | 1 | 0 | 0 | 0 | 1 | 1 | 1 | 0 |
| 49.19187 | 16.56446 | BRN_U_2 | brnu2_1 | Czech | U | 0 | 0 | 0 | 0 | 0 | 0 | 0 | 214,608 | 1331,96854 | 93 | 45161,059 | 4740 | 1 | 1 | 0 | 1 | 1 | 0 | 0 | 1 | 0 | 0 | 1 | 0 | 1 | 0 | 0 | 0 |
| 49.18898 | 16.56444 | BRN_U_2 | brnu2_2 | Czech | U | 2 | 1 | 0 | 0 | 2 | 1 | 1 | 215,704 | 1331,769113 | 93 | 49849,889 | 4486 | 1 | 1 | 0 | 1 | 1 | 0 | 0 | 1 | 0 | 0 | 1 | 0 | 0 | 0 | 0 | 0 |
| 49.18717 | 16.56769 | BRN_U_2 | brnu2_3 | Czech | U | 2 | 1 | 0 | 0 | 0 | 0 | 1 | 216,12 | 1331,461375 | 93 | 56665,781 | 1365 | 1 | 1 | 0 | 1 | 0 | 0 | 0 | 1 | 0 | 1 | 1 | 0 | 0 | 0 | 1 | 0 |
| 49.18409 | 16.56778 | BRN_U_2 | brnu2_4 | Czech | U | 1 | 1 | 0 | 0 | 0 | 0 | 0 | 228,86 | 1331,251302 | 93 | 58877,653 | 0 | 1 | 1 | 0 | 1 | 0 | 0 | 0 | 0 | 1 | 0 | 1 | 0 | 0 | 0 | 1 | 0 |
| 49.18144 | 16.56605 | BRN_U_2 | brnu2_5 | Czech | U | 2 | 1 | 0 | 0 | 0 | 0 | 0 | 257,036 | 1331,163811 | 93 | 55841,358 | 0 | 1 | 1 | 0 | 1 | 0 | 0 | 0 | 0 | 1 | 0 | 1 | 0 | 0 | 0 | 1 | 1 |
| 49.1724 | 16.59507 | BRN_U_3 | brnu3_1 | Czech | U | 2 | 1 | 0 | 0 | 0 | 0 | 0 | 226,748 | 1328,878123 | 96 | 22609,564 | 4416 | 0 | 1 | 1 | 0 | 0 | 1 | 1 | 1 | 0 | 0 | 0 | 1 | 1 | 0 | 0 | 0 |
| 49.17134 | 16.59147 | BRN_U_3 | brnu3_2 | Czech | U | 1 | 1 | 0 | 0 | 0 | 0 | 0 | 245,957 | 1329,015326 | 96 | 40246,397 | 393 | 0 | 1 | 1 | 0 | 0 | 1 | 0 | 0 | 0 | 0 | 1 | 0 | 0 | 0 | 0 | 0 |
| 49.16885 | 16.5916 | BRN_U_3 | brnu3_3 | Czech | U | 0 | 0 | 0 | 0 | 0 | 0 | 0 | 245,24 | 1328,835323 | 96 | 27413,065 | 0 | 0 | 1 | 1 | 0 | 0 | 1 | 0 | 0 | 0 | 0 | 0 | 0 | 0 | 0 | 0 | 0 |
| 49.16736 | 16.59441 | BRN_U_3 | brnu3_4 | Czech | U | 0 | 0 | 0 | 0 | 0 | 0 | 0 | 234,248 | 1328,565941 | 96 | 36721,379 | 0 | 0 | 1 | 1 | 0 | 0 | 1 | 0 | 0 | 0 | 0 | 0 | 0 | 0 | 0 | 0 | 0 |
| 49.16531 | 16.59655 | BRN_U_3 | brnu3_5 | Czech | U | 0 | 0 | 0 | 0 | 0 | 0 | 0 | 229,304 | 1328,306652 | 96 | 27134,562 | 6353 | 0 | 1 | 1 | 0 | 0 | 1 | 1 | 1 | 0 | 0 | 0 | 0 | 1 | 0 | 0 | 0 |
| 49.17694 | 16.58037 | BRN_U_4 | brnu4_1 | Czech | U | 0 | 0 | 1 | 1 | 0 | 0 | 1 | 287,3 | 1330,03487 | 93 | 31436,866 | 3464 | 1 | 1 | 0 | 0 | 0 | 0 | 0 | 0 | 1 | 0 | 1 | 0 | 1 | 0 | 0 | 0 |
| 49.1788 | 16.58065 | BRN_U_4 | brnu4_2 | Czech | U | 0 | 0 | 0 | 0 | 0 | 0 | 1 | 295,149 | 1329,735583 | 93 | 18733,138 | 3612 | 0 | 1 | 1 | 0 | 0 | 0 | 0 | 0 | 1 | 0 | 1 | 1 | 1 | 0 | 0 | 0 |
| 49.1801 | 16.5841 | BRN_U_4 | brnu4_3 | Czech | U | 1 | 1 | 0 | 0 | 0 | 0 | 1 | 278,12 | 1330,144281 | 93 | 31219,649 | 2136 | 0 | 1 | 1 | 0 | 0 | 0 | 1 | 0 | 1 | 0 | 1 | 1 | 0 | 0 | 0 | 0 |
| 49.17853 | 16.58727 | BRN_U_4 | brnu4_4 | Czech | U | 3 | 1 | 1 | 1 | 1 | 1 | 1 | 245,878 | 1330,034534 | 93 | 39629,371 | 3406 | 1 | 1 | 1 | 1 | 0 | 0 | 0 | 1 | 1 | 0 | 1 | 0 | 0 | 0 | 0 | 1 |
| 49.17568 | 16.5861 | BRN_U_4 | brnu4_5 | Czech | U | 1 | 1 | 0 | 0 | 0 | 0 | 0 | 273,496 | 1329,624387 | 93 | 50067,239 | 0 | 1 | 1 | 0 | 1 | 1 | 0 | 0 | 0 | 1 | 0 | 1 | 0 | 0 | 0 | 0 | 0 |
| 49.19966 | 16.66017 | BRN_U_5 | brnu5_1 | Czech | U | 0 | 0 | 2 | 1 | 0 | 0 | 0 | 281,164 | 1326,991035 | 100 | 38106,796 | 11081 | 1 | 1 | 0 | 1 | 1 | 0 | 0 | 1 | 1 | 0 | 1 | 0 | 1 | 0 | 0 | 0 |
| 49.19704 | 16.65865 | BRN_U_5 | brnu5_2 | Czech | U | 0 | 0 | 0 | 0 | 0 | 0 | 0 | 261,56 | 1326,903237 | 100 | 28470,599 | 4251 | 1 | 1 | 0 | 1 | 1 | 0 | 1 | 1 | 1 | 0 | 0 | 0 | 1 | 0 | 1 | 0 |
| 49.19506 | 16.6618 | BRN_U_5 | brnu5_3 | Czech | U | 2 | 1 | 0 | 0 | 0 | 0 | 0 | 295,616 | 1326,582801 | 100 | 23085,286 | 720 | 1 | 1 | 0 | 1 | 0 | 0 | 0 | 1 | 1 | 0 | 0 | 0 | 1 | 0 | 0 | 0 |
| 49.19358 | 16.6591 | BRN_U_5 | brnu5_4 | Czech | U | 0 | 0 | 0 | 0 | 0 | 0 | 0 | 283,693 | 1326,630828 | 100 | 41210,02 | 3337 | 1 | 1 | 0 | 1 | 1 | 0 | 0 | 1 | 1 | 0 | 1 | 0 | 1 | 0 | 0 | 0 |
| 49.1948 | 16.65609 | BRN_U_5 | brnu5_5 | Czech | U | 0 | 0 | 1 | 1 | 0 | 0 | 0 | 240,315 | 1326,896525 | 100 | 56411,168 | 6746 | 1 | 1 | 0 | 1 | 1 | 0 | 1 | 0 | 1 | 0 | 1 | 0 | 0 | 0 | 0 | 0 |
| 49.20139 | 16.64584 | BRN_U_6 | brnu6_1 | Czech | U | 0 | 0 | 0 | 0 | 0 | 0 | 0 | 216,024 | 1327,936631 | 99 | 26561,839 | 12596 | 1 | 0 | 0 | 1 | 0 | 0 | 0 | 0 | 1 | 0 | 1 | 0 | 1 | 0 | 0 | 0 |
| 49.20397 | 16.64556 | BRN_U_6 | brnu6_2 | Czech | U | 1 | 1 | 0 | 0 | 0 | 0 | 0 | 224,145 | 1328,129863 | 99 | 27207,473 | 8513 | 1 | 1 | 0 | 1 | 0 | 0 | 0 | 0 | 1 | 0 | 1 | 0 | 0 | 0 | 0 | 0 |
| 49.20602 | 16.64801 | BRN_U_6 | brnu6_3 | Czech | U | 2 | 1 | 0 | 0 | 0 | 0 | 0 | 229,126 | 1328,128089 | 99 | 35482,309 | 3996 | 1 | 1 | 1 | 1 | 0 | 1 | 0 | 1 | 1 | 0 | 1 | 1 | 1 | 0 | 0 | 1 |
| 49.20759 | 16.65076 | BRN_U_6 | brnu6_4 | Czech | U | 0 | 0 | 0 | 0 | 0 | 0 | 0 | 273,882 | 1328,074695 | 99 | 59730,072 | 457 | 0 | 1 | 1 | 0 | 0 | 1 | 0 | 1 | 1 | 0 | 1 | 0 | 1 | 1 | 0 | 0 |
| 49.20751 | 16.65415 | BRN_U_6 | brnu6_5 | Czech | U | 1 | 1 | 0 | 0 | 0 | 0 | 0 | 288,022 | 1327,877063 | 99 | 53167,477 | 2370 | 0 | 1 | 0 | 0 | 0 | 0 | 0 | 1 | 1 | 0 | 1 | 1 | 1 | 1 | 0 | 0 |
| 47.78658 | 19.13291 | BUD_R_1 | budr1_1 | Hungary | R | 0 | 0 | 0 | 0 | 0 | 0 | 0 | 115,788 | 1086,123277 | 235 | 9717,751 | 18166 | 1 | 1 | 0 | 1 | 0 | 0 | 1 | 1 | 0 | 0 | 0 | 1 | 0 | 1 | 1 | 1 |
| 47.78908 | 19.1336 | BUD_R_1 | budr1_2 | Hungary | R | 1 | 1 | 0 | 0 | 0 | 0 | 0 | 122,036 | 1086,261482 | 235 | 31097,743 | 11345 | 1 | 1 | 1 | 1 | 0 | 0 | 0 | 1 | 0 | 0 | 1 | 0 | 1 | 0 | 0 | 0 |
| 47.79053 | 19.13669 | BUD_R_1 | budr1_3 | Hungary | R | 0 | 0 | 1 | 1 | 0 | 0 | 1 | 132,016 | 1086,183159 | 235 | 26402,223 | 16990 | 0 | 1 | 0 | 0 | 0 | 0 | 1 | 1 | 1 | 0 | 0 | 1 | 1 | 0 | 0 | 0 |
| 47.78852 | 19.13894 | BUD_R_1 | budr1_4 | Hungary | R | 0 | 0 | 0 | 0 | 0 | 0 | 1 | 131,816 | 1085,910782 | 235 | 9757,429 | 21515 | 0 | 1 | 0 | 0 | 0 | 0 | 0 | 1 | 1 | 0 | 0 | 1 | 0 | 0 | 0 | 0 |
| 47.78644 | 19.13984 | BUD_R_1 | budr1_5 | Hungary | R | 1 | 1 | 3 | 1 | 0 | 0 | 0 | 127,078 | 1085,713845 | 235 | 14240,104 | 18013 | 0 | 1 | 1 | 0 | 0 | 0 | 0 | 1 | 1 | 0 | 0 | 1 | 0 | 0 | 0 | 0 |
| 47.47397 | 18.52279 | BUD_R_2 | budr2_1 | Hungary | R | 0 | 0 | 0 | 0 | 0 | 0 | 0 | 187,677 | 1101,029619 | 168 | 6172,936 | 9192 | 0 | 1 | 0 | 0 | 0 | 0 | 1 | 0 | 1 | 0 | 0 | 1 | 0 | 0 | 0 | 0 |
| 47.47616 | 18.51987 | BUD_R_2 | budr2_2 | Hungary | R | 0 | 0 | 1 | 1 | 0 | 0 | 0 | 195,55 | 1101,350917 | 168 | 9502,75 | 10970 | 0 | 1 | 0 | 0 | 0 | 0 | 1 | 0 | 1 | 0 | 0 | 1 | 1 | 0 | 0 | 0 |
| 47.47912 | 18.52055 | BUD_R_2 | budr2_3 | Hungary | R | 0 | 0 | 1 | 1 | 0 | 0 | 0 | 175,003 | 1101,503804 | 168 | 7999,257 | 4086 | 0 | 1 | 0 | 1 | 0 | 0 | 0 | 1 | 1 | 0 | 1 | 1 | 0 | 0 | 0 | 0 |
| 47.48213 | 18.52111 | BUD_R_2 | budr2_4 | Hungary | R | 0 | 0 | 1 | 1 | 0 | 0 | 0 | 180,665 | 1101,667307 | 168 | 5761,691 | 7499 | 0 | 1 | 0 | 0 | 0 | 0 | 0 | 0 | 1 | 0 | 0 | 1 | 0 | 0 | 0 | 0 |
| 47.48449 | 18.51941 | BUD_R_2 | budr2_5 | Hungary | R | 1 | 1 | 0 | 0 | 0 | 0 | 0 | 186,752 | 1101,925695 | 168 | 16958,444 | 1949 | 1 | 1 | 0 | 1 | 0 | 0 | 1 | 0 | 1 | 0 | 1 | 1 | 0 | 0 | 0 | 0 |
| 47.48524 | 19.01717 | BUD_U_1 | budu1_1 | Hungary | U | 1 | 1 | 0 | 0 | 0 | 0 | 1 | 184,808 | 1072,003951 | 221 | 18146,682 | 13005 | 0 | 1 | 0 | 0 | 0 | 0 | 1 | 1 | 0 | 0 | 0 | 1 | 1 | 0 | 0 | 0 |
| 47.4837 | 19.01944 | BUD_U_1 | budu1_2 | Hungary | U | 0 | 0 | 1 | 1 | 0 | 0 | 0 | 199,744 | 1071,765013 | 221 | 27594,398 | 12759 | 1 | 1 | 0 | 1 | 0 | 0 | 0 | 1 | 0 | 0 | 0 | 0 | 1 | 0 | 0 | 0 |
| 47.48355 | 19.01526 | BUD_U_1 | budu1_3 | Hungary | U | 0 | 0 | 1 | 1 | 0 | 0 | 0 | 222,766 | 1072,003795 | 221 | 24048,568 | 12257 | 1 | 1 | 1 | 1 | 0 | 0 | 0 | 1 | 1 | 0 | 1 | 0 | 1 | 0 | 0 | 0 |
| 47.48338 | 19.01189 | BUD_U_1 | budu1_4 | Hungary | U | 0 | 0 | 0 | 0 | 0 | 0 | 0 | 233,094 | 1072,193169 | 221 | 11789,422 | 10483 | 0 | 1 | 1 | 0 | 0 | 0 | 0 | 1 | 1 | 0 | 0 | 1 | 1 | 0 | 0 | 0 |
| 47.48508 | 19.01281 | BUD_U_1 | budu1_5 | Hungary | U | 2 | 1 | 0 | 0 | 0 | 0 | 1 | 211,02 | 1072,254015 | 221 | 10992,212 | 12181 | 0 | 1 | 1 | 0 | 0 | 0 | 0 | 1 | 1 | 0 | 0 | 1 | 1 | 0 | 0 | 1 |
| 47.42465 | 19.07929 | BUD_U_2 | budu2_1 | Hungary | U | 0 | 0 | 0 | 0 | 0 | 0 | 1 | 106,527 | 1064,17972 | 225 | 13436,899 | 11718 | 0 | 1 | 1 | 0 | 0 | 0 | 1 | 1 | 1 | 0 | 0 | 1 | 1 | 0 | 0 | 0 |
| 47.42453 | 19.08319 | BUD_U_2 | budu2_2 | Hungary | U | 0 | 0 | 1 | 1 | 0 | 0 | 1 | 102,159 | 1063,937206 | 225 | 4722,904 | 15662 | 0 | 1 | 1 | 0 | 0 | 0 | 0 | 0 | 1 | 0 | 0 | 1 | 0 | 0 | 0 | 0 |
| 47.42258 | 19.08246 | BUD_U_2 | budu2_3 | Hungary | U | 0 | 0 | 1 | 1 | 0 | 0 | 0 | 104,041 | 1063,848863 | 225 | 5609,917 | 16365 | 0 | 1 | 0 | 0 | 0 | 0 | 1 | 0 | 1 | 0 | 0 | 1 | 0 | 0 | 0 | 0 |
| 47.42264 | 19.07793 | BUD_U_2 | budu2_4 | Hungary | U | 0 | 0 | 1 | 1 | 0 | 0 | 0 | 106,728 | 1064,120849 | 225 | 7279,031 | 13006 | 0 | 1 | 0 | 0 | 0 | 0 | 1 | 0 | 1 | 0 | 0 | 1 | 1 | 0 | 0 | 0 |
| 47.42401 | 19.07495 | BUD_U_2 | budu2_5 | Hungary | U | 1 | 1 | 0 | 0 | 0 | 0 | 1 | 102,541 | 1064,394453 | 225 | 5865,405 | 15060 | 0 | 1 | 1 | 0 | 0 | 0 | 1 | 0 | 1 | 0 | 0 | 1 | 1 | 0 | 0 | 0 |
| 53.15577 | 18.14314 | BYD_R_1 | bydr1_1 | Poland | R | 0 | 0 | 0 | 0 | 0 | 0 | 1 | 59,315 | 1573,704794 | -100 | 18440,064 | 861 | 1 | 1 | 0 | 1 | 0 | 0 | 0 | 0 | 0 | 0 | 0 | 0 | 0 | 0 | 0 | 1 |
| 53.15823 | 18.14049 | BYD_R_1 | bydr1_2 | Poland | R | 0 | 0 | 0 | 0 | 0 | 0 | 1 | 79,949 | 1574,030462 | -100 | 12392,047 | 0 | 1 | 1 | 0 | 1 | 0 | 0 | 0 | 0 | 0 | 0 | 0 | 0 | 0 | 0 | 0 | 0 |
| 53.16077 | 18.1427 | BYD_R_1 | bydr1_3 | Poland | R | 0 | 0 | 0 | 0 | 0 | 0 | 0 | 94,28 | 1574,174473 | -100 | 27033,897 | 0 | 0 | 1 | 0 | 1 | 0 | 0 | 0 | 0 | 0 | 0 | 0 | 0 | 0 | 0 | 0 | 0 |
| 53.1619 | 18.14681 | BYD_R_1 | bydr1_4 | Poland | R | 0 | 0 | 0 | 0 | 0 | 0 | 1 | 77,124 | 1574,114233 | -100 | 24532,832 | 1416 | 1 | 1 | 0 | 1 | 0 | 0 | 0 | 0 | 0 | 0 | 1 | 0 | 0 | 0 | 0 | 0 |
| 53.16153 | 18.1513 | BYD_R_1 | bydr1_5 | Poland | R | 0 | 0 | 0 | 0 | 0 | 0 | 1 | 53,656 | 1573,9038 | -100 | 16792,651 | 2998 | 1 | 1 | 0 | 1 | 0 | 0 | 0 | 0 | 1 | 0 | 1 | 1 | 0 | 0 | 0 | 0 |
| 53.17784 | 18.17526 | BYD_R_2 | bydr2_1 | Poland | R | 0 | 0 | 0 | 0 | 0 | 0 | 1 | 31,79 | 1574,434647 | -100 | 9795,98 | 3217 | 1 | 1 | 0 | 1 | 0 | 0 | 0 | 0 | 0 | 0 | 1 | 0 | 0 | 0 | 0 | 0 |
| 53.17672 | 18.18028 | BYD_R_2 | bydr2_2 | Poland | R | 0 | 0 | 0 | 0 | 0 | 0 | 1 | 31,992 | 1574,137811 | -100 | 2240,571 | 190 | 0 | 1 | 0 | 0 | 0 | 0 | 1 | 0 | 1 | 0 | 1 | 0 | 0 | 0 | 0 | 0 |
| 53.17717 | 18.18438 | BYD_R_2 | bydr2_3 | Poland | R | 0 | 0 | 0 | 0 | 0 | 0 | 1 | 29,623 | 1574,01558 | -100 | 20124,909 | 1445 | 0 | 1 | 0 | 1 | 0 | 0 | 0 | 0 | 0 | 1 | 0 | 0 | 1 | 0 | 1 | 0 |
| 53.17982 | 18.18622 | BYD_R_2 | bydr2_4 | Poland | R | 0 | 0 | 0 | 0 | 0 | 0 | 1 | 28,924 | 1574,173151 | -100 | 13405,757 | 0 | 0 | 1 | 0 | 1 | 0 | 0 | 1 | 0 | 0 | 1 | 0 | 0 | 0 | 0 | 0 | 0 |
| 53.18198 | 18.18781 | BYD_R_2 | bydr2_5 | Poland | R | 0 | 0 | 0 | 0 | 0 | 0 | 1 | 28,887 | 1574,314804 | -100 | 6955,932 | 0 | 0 | 1 | 0 | 1 | 0 | 0 | 0 | 0 | 0 | 1 | 0 | 0 | 0 | 0 | 0 | 0 |
| 53.11583 | 17.9835 | BYD_U_1 | bydu1_1 | Poland | U | 1 | 1 | 0 | 0 | 0 | 0 | 0 | 66,624 | 1576,443379 | -97 | 17943,642 | 13682 | 0 | 1 | 0 | 0 | 0 | 0 | 0 | 1 | 1 | 0 | 0 | 0 | 1 | 0 | 0 | 0 |
| 53.11802 | 17.98347 | BYD_U_1 | bydu1_2 | Poland | U | 1 | 1 | 0 | 0 | 0 | 0 | 1 | 63,986 | 1576,643309 | -97 | 25104,739 | 11834 | 0 | 1 | 0 | 0 | 0 | 0 | 0 | 1 | 0 | 0 | 0 | 0 | 1 | 1 | 0 | 0 |
| 53.11968 | 17.987 | BYD_U_1 | bydu1_3 | Poland | U | 1 | 1 | 0 | 0 | 0 | 0 | 1 | 67,522 | 1576,620967 | -97 | 32842,592 | 8863 | 1 | 1 | 0 | 0 | 1 | 0 | 0 | 1 | 0 | 0 | 0 | 0 | 1 | 0 | 0 | 0 |
| 53.11949 | 17.99206 | BYD_U_1 | bydu1_4 | Poland | U | 0 | 0 | 0 | 0 | 0 | 0 | 1 | 70,18 | 1576,426995 | -97 | 23732,468 | 19185 | 1 | 1 | 0 | 0 | 1 | 0 | 0 | 1 | 1 | 0 | 0 | 1 | 1 | 1 | 0 | 0 |
| 53.11718 | 17.99514 | BYD_U_1 | bydu1_5 | Poland | U | 0 | 0 | 0 | 0 | 0 | 0 | 1 | 70,848 | 1576,098864 | -97 | 10407,364 | 17291 | 0 | 1 | 0 | 0 | 0 | 0 | 0 | 1 | 0 | 0 | 0 | 0 | 1 | 0 | 0 | 0 |
| 53.12449 | 18.01258 | BYD_U_2 | bydu2_1 | Poland | U | 0 | 0 | 0 | 0 | 0 | 0 | 1 | 37,164 | 1576,059178 | -98 | 35643,95 | 10529 | 0 | 1 | 0 | 0 | 1 | 0 | 0 | 1 | 0 | 0 | 0 | 0 | 1 | 1 | 0 | 0 |
| 53.12584 | 18.0083 | BYD_U_2 | bydu2_2 | Poland | U | 0 | 0 | 0 | 0 | 0 | 0 | 0 | 39,024 | 1576,350556 | -98 | 9795,028 | 24406 | 0 | 1 | 0 | 0 | 0 | 0 | 0 | 1 | 0 | 0 | 0 | 0 | 1 | 1 | 0 | 0 |
| 53.12824 | 18.00869 | BYD_U_2 | bydu2_3 | Poland | U | 0 | 0 | 0 | 0 | 0 | 0 | 0 | 44 | 1576,54792 | -98 | 8575,488 | 29122 | 0 | 1 | 0 | 0 | 0 | 0 | 0 | 1 | 1 | 0 | 0 | 0 | 0 | 1 | 0 | 0 |
| 53.13017 | 18.01193 | BYD_U_2 | bydu2_4 | Poland | U | 0 | 0 | 0 | 0 | 0 | 0 | 1 | 46,032 | 1576,589631 | -98 | 26009,886 | 14444 | 0 | 1 | 0 | 0 | 1 | 0 | 0 | 1 | 1 | 0 | 0 | 0 | 0 | 1 | 0 | 0 |
| 53.13069 | 18.0162 | BYD_U_2 | bydu2_5 | Poland | U | 0 | 0 | 0 | 0 | 0 | 0 | 1 | 46,649 | 1576,474834 | -98 | 24753,232 | 16327 | 1 | 1 | 0 | 0 | 1 | 0 | 0 | 1 | 0 | 0 | 0 | 0 | 0 | 1 | 0 | 0 |
| 45.3274 | 22.2637 | CAR_R_1 | carr1_1 | Romania | R | 0 | 0 | 1 | 1 | 0 | 0 | 0 | 241,32 | 727,7379853 | 341 | 10101,448 | 0 | 0 | 1 | 0 | 1 | 0 | 0 | 0 | 1 | 0 | 0 | 0 | 0 | 0 | 0 | 0 | 0 |
| 45.32477 | 22.26595 | CAR_R_1 | carr1_2 | Romania | R | 0 | 0 | 0 | 0 | 0 | 0 | 0 | 244 | 727,4221419 | 341 | 17119,225 | 0 | 0 | 1 | 0 | 1 | 0 | 0 | 0 | 0 | 0 | 1 | 0 | 0 | 0 | 0 | 0 | 0 |
| 45.32228 | 22.26864 | CAR_R_1 | carr1_3 | Romania | R | 0 | 0 | 0 | 0 | 0 | 0 | 0 | 245,814 | 727,0830082 | 341 | 23205,237 | 0 | 0 | 1 | 0 | 1 | 0 | 0 | 1 | 0 | 0 | 1 | 0 | 0 | 0 | 0 | 0 | 0 |
| 45.31944 | 22.26889 | CAR_R_1 | carr1_4 | Romania | R | 1 | 1 | 0 | 0 | 0 | 0 | 0 | 249 | 726,8731518 | 341 | 16855,317 | 0 | 0 | 1 | 0 | 1 | 0 | 0 | 1 | 0 | 0 | 1 | 0 | 0 | 0 | 0 | 0 | 0 |
| 45.31721 | 22.2702 | CAR_R_1 | carr1_5 | Romania | R | 0 | 0 | 0 | 0 | 0 | 0 | 0 | 250 | 726,643348 | 341 | 6045,229 | 0 | 0 | 1 | 0 | 0 | 0 | 0 | 0 | 0 | 0 | 1 | 0 | 0 | 0 | 0 | 0 | 0 |
| 45.34864 | 22.27454 | CAR_R_2 | carr2_1 | Romania | R | 1 | 1 | 0 | 0 | 0 | 0 | 0 | 253,552 | 728,5178957 | 341 | 24626,355 | 0 | 1 | 1 | 0 | 1 | 0 | 0 | 1 | 0 | 1 | 0 | 0 | 0 | 0 | 0 | 0 | 0 |
| 45.34863 | 22.27829 | CAR_R_2 | carr2_2 | Romania | R | 0 | 0 | 0 | 0 | 0 | 0 | 0 | 261,059 | 728,2843659 | 341 | 18417,508 | 0 | 1 | 1 | 0 | 1 | 0 | 0 | 1 | 0 | 0 | 0 | 0 | 0 | 0 | 0 | 0 | 0 |

| | | | | | | | | | | | | | | | | | | | | | | | | | | | | | | | | | |
|---|---|---|---|---|---|---|---|---|---|---|---|---|---|---|---|---|---|---|---|---|---|---|---|---|---|---|---|---|---|---|---|---|---|
| 45.34893 | 22.28254 | CAR_R_2 | carr2_3 | Romania | R | 0 | 0 | 0 | 0 | 0 | 0 | 1 | 268,419 | 728,0433866 | 341 | 15954,876 | 0 | 1 | 1 | 0 | 1 | 0 | 0 | 1 | 0 | 0 | 0 | 0 | 0 | 0 | 0 | 0 | 0 |
| 45.34896 | 22.28702 | CAR_R_2 | carr2_4 | Romania | R | 0 | 0 | 0 | 0 | 0 | 0 | 0 | 269 | 727,7690835 | 341 | 15214,684 | 0 | 1 | 1 | 0 | 1 | 0 | 0 | 1 | 0 | 0 | 0 | 0 | 0 | 0 | 0 | 0 | 0 |
| 45.34883 | 22.29125 | CAR_R_2 | carr2_5 | Romania | R | 0 | 0 | 0 | 0 | 0 | 0 | 1 | 270,394 | 727,498941 | 341 | 3331,218 | 0 | 0 | 1 | 0 | 0 | 0 | 0 | 0 | 1 | 0 | 0 | 0 | 0 | 0 | 0 | 0 | 0 |
| 45.40859 | 22.21342 | CAR_U_1 | caru1_1 | Romania | U | 0 | 0 | 2 | 1 | 0 | 0 | 1 | 201,312 | 736,3934567 | 347 | 20249,18 | 11580 | 1 | 1 | 0 | 1 | 0 | 0 | 1 | 0 | 0 | 1 | 0 | 1 | 0 | 0 | 0 | 0 |
| 45.40915 | 22.20998 | CAR_U_1 | caru1_2 | Romania | U | 0 | 0 | 0 | 0 | 0 | 0 | 1 | 201 | 736,6440278 | 347 | 16496,075 | 8324 | 0 | 1 | 0 | 0 | 0 | 0 | 1 | 1 | 0 | 1 | 0 | 1 | 0 | 0 | 0 | 0 |
| 45.40894 | 22.20614 | CAR_U_1 | caru1_3 | Romania | U | 0 | 0 | 0 | 0 | 0 | 0 | 1 | 200 | 736,8646993 | 347 | 32009,163 | 124 | 0 | 1 | 0 | 0 | 0 | 0 | 1 | 1 | 0 | 1 | 1 | 0 | 0 | 0 | 0 | 0 |
| 45.41022 | 22.20378 | CAR_U_1 | caru1_4 | Romania | U | 0 | 0 | 0 | 0 | 0 | 0 | 1 | 200 | 737,0991317 | 347 | 44135,063 | 923 | 1 | 1 | 0 | 1 | 0 | 0 | 1 | 0 | 1 | 1 | 1 | 0 | 0 | 0 | 0 | 0 |
| 45.41287 | 22.20298 | CAR_U_1 | caru1_5 | Romania | U | 0 | 0 | 1 | 1 | 0 | 0 | 1 | 199 | 737,3305481 | 347 | 33089,352 | 1680 | 1 | 1 | 1 | 1 | 0 | 0 | 1 | 0 | 0 | 1 | 1 | 0 | 0 | 0 | 0 | 0 |
| 45.41219 | 22.22648 | CAR_U_2 | caru2_1 | Romania | U | 0 | 0 | 1 | 1 | 0 | 0 | 1 | 215 | 735,8372829 | 347 | 14073,65 | 13139 | 0 | 1 | 0 | 0 | 0 | 0 | 1 | 1 | 1 | 0 | 0 | 1 | 1 | 0 | 0 | 0 |
| 45.41275 | 22.23024 | CAR_U_2 | caru2_2 | Romania | U | 0 | 0 | 0 | 0 | 0 | 0 | 0 | 215 | 735,6441366 | 347 | 8526,815 | 9719 | 0 | 1 | 0 | 0 | 0 | 0 | 0 | 1 | 1 | 0 | 0 | 1 | 0 | 1 | 0 | 0 |
| 45.41138 | 22.23272 | CAR_U_2 | caru2_3 | Romania | U | 0 | 0 | 0 | 0 | 0 | 0 | 1 | 214 | 735,397722 | 347 | 3700,835 | 7424 | 0 | 1 | 0 | 0 | 0 | 0 | 1 | 0 | 1 | 0 | 1 | 1 | 0 | 0 | 0 | 0 |
| 45.41025 | 22.23414 | CAR_U_2 | caru2_4 | Romania | U | 0 | 0 | 0 | 0 | 0 | 0 | 0 | 218,804 | 735,2327321 | 347 | 6216,685 | 11155 | 1 | 1 | 0 | 0 | 0 | 0 | 1 | 0 | 1 | 0 | 0 | 1 | 0 | 0 | 1 | 0 |
| 45.40911 | 22.23055 | CAR_U_2 | caru2_5 | Romania | U | 0 | 0 | 0 | 0 | 0 | 0 | 0 | 213,016 | 735,3750287 | 347 | 13112,125 | 5533 | 0 | 1 | 0 | 0 | 0 | 0 | 1 | 1 | 0 | 0 | 0 | 1 | 0 | 0 | 0 | 0 |
| 49.383691 | 32.184460 | CHE_R_1 | cher1_1 | Ukraine | R | 1 | 1 | 1 | 1 | 0 | 0 | 1 | 85,344 | 951,5669096 | 385 | 29644,867 | 2319 | 1 | 1 | 0 | 1 | 0 | 0 | 0 | 0 | 1 | 1 | 1 | 1 | 0 | 0 | 0 | 0 |
| 49.382772 | 32.181298 | CHE_R_1 | cher1_2 | Ukraine | R | 0 | 0 | 0 | 0 | 0 | 0 | 1 | 85,604 | 951,4325141 | 385 | 12988,876 | 3926 | 0 | 1 | 0 | 1 | 0 | 0 | 0 | 0 | 1 | 0 | 1 | 0 | 0 | 0 | 0 | 0 |
| 49.382801 | 32.176029 | CHE_R_1 | cher1_3 | Ukraine | R | 0 | 0 | 0 | 0 | 0 | 0 | 1 | 79 | 951,292225 | 385 | 10737,739 | 1118 | 0 | 1 | 0 | 1 | 0 | 0 | 0 | 0 | 1 | 1 | 1 | 1 | 0 | 0 | 0 | 0 |
| 49.381472 | 32.170957 | CHE_R_1 | cher1_4 | Ukraine | R | 2 | 1 | 0 | 0 | 0 | 0 | 1 | 81 | 951,0716663 | 385 | 11088,675 | 1508 | 0 | 1 | 0 | 1 | 0 | 0 | 0 | 0 | 0 | 1 | 1 | 0 | 0 | 0 | 0 | 0 |
| 49.378556 | 32.167478 | CHE_R_1 | cher1_5 | Ukraine | R | 0 | 0 | 0 | 0 | 0 | 0 | 1 | 80,682 | 950,6931915 | 385 | 5768,245 | 0 | 0 | 1 | 0 | 1 | 0 | 0 | 0 | 0 | 0 | 0 | 0 | 0 | 0 | 0 | 0 | 0 |
| 49.336937 | 32.211714 | CHE_R_2 | cher2_1 | Ukraine | R | 0 | 0 | 2 | 1 | 0 | 0 | 1 | 106 | 947,1159889 | 391 | 16674,112 | 3554 | 0 | 1 | 0 | 0 | 0 | 0 | 0 | 0 | 1 | 0 | 0 | 1 | 0 | 0 | 0 | 0 |
| 49.338831 | 32.214987 | CHE_R_2 | cher2_2 | Ukraine | R | 0 | 0 | 0 | 0 | 0 | 0 | 0 | 105,218 | 947,381575 | 391 | 11664,999 | 3474 | 1 | 1 | 0 | 1 | 0 | 0 | 0 | 0 | 1 | 0 | 0 | 1 | 0 | 0 | 0 | 0 |
| 49.342208 | 32.215802 | CHE_R_2 | cher2_3 | Ukraine | R | 0 | 0 | 0 | 0 | 0 | 0 | 0 | 83,99 | 947,7006258 | 391 | 12972,447 | 521 | 1 | 1 | 0 | 1 | 0 | 0 | 0 | 0 | 1 | 0 | 1 | 0 | 0 | 0 | 0 | 0 |
| 49.343200 | 32.219310 | CHE_R_2 | cher2_4 | Ukraine | R | 1 | 1 | 0 | 0 | 0 | 0 | 0 | 83,48 | 947,9171024 | 391 | 6328,343 | 1054 | 0 | 1 | 0 | 1 | 0 | 0 | 0 | 0 | 1 | 1 | 1 | 0 | 0 | 0 | 0 | 0 |
| 49.341921 | 32.222840 | CHE_R_2 | cher2_5 | Ukraine | R | 1 | 1 | 0 | 0 | 0 | 0 | 0 | 87,964 | 947,8733818 | 391 | 35050,943 | 1662 | 1 | 1 | 0 | 1 | 0 | 0 | 0 | 1 | 1 | 0 | 1 | 0 | 0 | 0 | 0 | 0 |
| 49.415682 | 32.023094 | CHE_U_1 | cheu1_1 | Ukraine | U | 0 | 0 | 2 | 1 | 0 | 0 | 0 | 112,469 | 958,0823693 | 373 | 61375,751 | 1874 | 1 | 0 | 0 | 1 | 0 | 0 | 0 | 0 | 0 | 0 | 0 | 1 | 0 | 0 | 0 | 0 |
| 49.417790 | 32.023356 | CHE_U_1 | cheu1_2 | Ukraine | U | 1 | 1 | 0 | 0 | 0 | 0 | 1 | 111,072 | 957,8893154 | 373 | 50602,468 | 479 | 1 | 1 | 0 | 1 | 0 | 0 | 0 | 0 | 0 | 1 | 0 | 0 | 0 | 0 | 0 | 0 |
| 49.419893 | 32.025098 | CHE_U_1 | cheu1_3 | Ukraine | U | 0 | 0 | 1 | 1 | 0 | 0 | 1 | 110,738 | 957,636627 | 373 | 48334,829 | 2959 | 1 | 0 | 0 | 1 | 0 | 0 | 0 | 0 | 0 | 1 | 0 | 1 | 0 | 0 | 0 | 0 |
| 49.420992 | 32.022807 | CHE_U_1 | cheu1_4 | Ukraine | U | 0 | 0 | 0 | 0 | 0 | 0 | 0 | 110,06 | 957,3135642 | 373 | 39164,482 | 0 | 1 | 1 | 0 | 0 | 1 | 0 | 0 | 1 | 0 | 0 | 1 | 0 | 0 | 0 | 0 | 0 |
| 49.25197 | 32.01139 | CHE_U_1 | cheu1_5 | Ukraine | U | 0 | 0 | 0 | 0 | 0 | 0 | 0 | 162 | 957,4396981 | 373 | 38405,069 | 1200 | 1 | 1 | 0 | 1 | 1 | 0 | 0 | 0 | 0 | 0 | 1 | 0 | 0 | 0 | 0 | 0 |
| 49.474258 | 32.016668 | CHE_U_2 | cheu2_1 | Ukraine | U | 1 | 1 | 0 | 0 | 0 | 0 | 0 | 117,007 | 951,7959448 | 380 | 55044,184 | 725 | 1 | 1 | 0 | 0 | 1 | 0 | 0 | 0 | 0 | 1 | 0 | 0 | 0 | 0 | 1 | 0 |
| 49.469321 | 32.021795 | CHE_U_2 | cheu2_2 | Ukraine | U | 2 | 1 | 0 | 0 | 0 | 0 | 1 | 124,726 | 951,8926208 | 380 | 51297,071 | 2326 | 1 | 1 | 0 | 0 | 1 | 0 | 0 | 0 | 0 | 0 | 0 | 1 | 0 | 0 | 0 | 0 |
| 49.466024 | 32.023132 | CHE_U_2 | cheu2_3 | Ukraine | U | 0 | 0 | 1 | 1 | 0 | 0 | 0 | 121,464 | 952,137184 | 380 | 33870,063 | 5438 | 1 | 1 | 0 | 0 | 1 | 0 | 0 | 1 | 0 | 0 | 0 | 0 | 1 | 0 | 1 | 0 |
| 49.466784 | 32.025107 | CHE_U_2 | cheu2_4 | Ukraine | U | 0 | 0 | 0 | 0 | 0 | 0 | 0 | 117,8 | 952,411005 | 380 | 33139,433 | 5742 | 0 | 1 | 0 | 0 | 1 | 0 | 0 | 1 | 0 | 0 | 0 | 0 | 1 | 0 | 1 | 0 |
| 49.466412 | 32.027749 | CHE_U_2 | cheu2_5 | Ukraine | U | 3 | 1 | 0 | 0 | 0 | 0 | 0 | 112,07 | 952,5327945 | 380 | 21318,115 | 9388 | 0 | 1 | 0 | 0 | 0 | 0 | 0 | 1 | 0 | 0 | 0 | 0 | 1 | 0 | 1 | 0 |
| 48.95325 | 14.46334 | CZB_R_1 | czbr1_1 | Czech | R | 3 | 1 | 0 | 0 | 0 | 0 | 1 | 391 | 1443,108607 | 1 | 28172,953 | 0 | 1 | 1 | 0 | 1 | 0 | 0 | 0 | 1 | 0 | 0 | 0 | 0 | 0 | 0 | 0 | 0 |
| 48.95152 | 14.46405 | CZB_R_1 | czbr1_2 | Czech | R | 2 | 1 | 0 | 0 | 0 | 0 | 0 | 391,388 | 1442,963078 | 1 | 35954,343 | 2777 | 1 | 1 | 0 | 1 | 0 | 0 | 1 | 0 | 1 | 1 | 0 | 1 | 0 | 0 | 0 | 1 |
| 48.94914 | 14.46209 | CZB_R_1 | czbr1_3 | Czech | R | 1 | 1 | 0 | 0 | 0 | 0 | 0 | 392 | 1442,946255 | 1 | 12153,773 | 1323 | 0 | 1 | 0 | 1 | 0 | 0 | 1 | 0 | 1 | 1 | 0 | 1 | 0 | 0 | 0 | 0 |
| 48.94677 | 14.45967 | CZB_R_1 | czbr1_4 | Czech | R | 0 | 0 | 0 | 0 | 0 | 0 | 0 | 392,118 | 1442,958889 | 1 | 6855,449 | 0 | 0 | 1 | 0 | 1 | 0 | 0 | 1 | 0 | 0 | 1 | 0 | 0 | 0 | 0 | 0 | 0 |
| 48.94435 | 14.4575 | CZB_R_1 | czbr1_5 | Czech | R | 0 | 0 | 0 | 0 | 0 | 0 | 0 | 394 | 1442,952622 | 1 | 10148,821 | 3631 | 0 | 1 | 0 | 0 | 0 | 0 | 0 | 0 | 1 | 0 | 1 | 0 | 0 | 0 | 0 | 0 |
| 48.89915 | 14.45012 | CZB_R_2 | czbr2_1 | Czech | R | 1 | 1 | 0 | 0 | 0 | 0 | 1 | 492,06 | 1440,781397 | 1 | 12075,288 | 3813 | 0 | 1 | 0 | 0 | 0 | 1 | 0 | 1 | 1 | 0 | 1 | 1 | 0 | 0 | 0 | 0 |
| 48.89641 | 14.45107 | CZB_R_2 | czbr2_2 | Czech | R | 0 | 0 | 0 | 0 | 0 | 0 | 1 | 496,011 | 1440,563006 | 1 | 5724,915 | 7866 | 0 | 1 | 0 | 0 | 0 | 0 | 0 | 1 | 1 | 0 | 0 | 1 | 1 | 0 | 0 | 1 |
| 48.89571 | 14.45465 | CZB_R_2 | czbr2_3 | Czech | R | 2 | 1 | 0 | 0 | 0 | 0 | 1 | 485,961 | 1440,298281 | 1 | 6564,115 | 6060 | 0 | 1 | 0 | 0 | 0 | 0 | 0 | 0 | 1 | 0 | 0 | 1 | 0 | 0 | 0 | 1 |
| 48.89556 | 14.45875 | CZB_R_2 | czbr2_4 | Czech | R | 3 | 1 | 0 | 0 | 0 | 0 | 1 | 474,532 | 1440,033286 | 1 | 8372,734 | 2406 | 0 | 1 | 0 | 0 | 0 | 0 | 1 | 0 | 1 | 0 | 1 | 0 | 0 | 0 | 0 | 1 |
| 48.89624 | 14.4621 | CZB_R_2 | czbr2_5 | Czech | R | 1 | 1 | 0 | 0 | 0 | 0 | 1 | 467,604 | 1439,863414 | 1 | 18451,096 | 1635 | 1 | 1 | 0 | 1 | 0 | 0 | 1 | 0 | 0 | 0 | 0 | 0 | 0 | 0 | 0 | 0 |
| 48.99263 | 14.43746 | CZB_R_3 | czbr3_1 | Czech | R | 3 | 1 | 0 | 0 | 0 | 0 | 0 | 387,933 | 1447,019832 | 3 | 39478,734 | 0 | 1 | 1 | 0 | 1 | 0 | 0 | 0 | 0 | 0 | 0 | 0 | 0 | 0 | 0 | 0 | 0 |
| 48.99497 | 14.43988 | CZB_R_3 | czbr3_2 | Czech | R | 2 | 1 | 0 | 0 | 0 | 0 | 1 | 386,615 | 1447,010802 | 3 | 52246,182 | 0 | 1 | 1 | 0 | 1 | 0 | 0 | 0 | 0 | 0 | 0 | 0 | 0 | 0 | 0 | 0 | 1 |
| 48.99726 | 14.4422 | CZB_R_3 | czbr3_3 | Czech | R | 1 | 1 | 0 | 0 | 0 | 0 | 0 | 384 | 1447,002779 | 3 | 53047,018 | 0 | 1 | 1 | 0 | 1 | 0 | 0 | 0 | 0 | 0 | 1 | 0 | 0 | 0 | 0 | 0 | 0 |
| 49.00002 | 14.44122 | CZB_R_3 | czbr3_4 | Czech | R | 1 | 1 | 0 | 0 | 0 | 0 | 0 | 382,564 | 1447,224301 | 3 | 34012,132 | 0 | 1 | 1 | 0 | 1 | 0 | 0 | 0 | 0 | 1 | 1 | 1 | 0 | 0 | 0 | 0 | 0 |
| 49.00248 | 14.43873 | CZB_R_3 | czbr3_5 | Czech | R | 0 | 0 | 0 | 0 | 0 | 0 | 1 | 381,959 | 1447,524268 | 3 | 5397,516 | 0 | 0 | 1 | 0 | 0 | 0 | 0 | 1 | 0 | 0 | 1 | 0 | 0 | 0 | 0 | 0 | 0 |
| 48.99002 | 14.41331 | CZB_R_4 | czbr4_1 | Czech | R | 1 | 1 | 0 | 0 | 0 | 0 | 1 | 391 | 1448,374322 | 4 | 5184,192 | 0 | 0 | 1 | 0 | 0 | 0 | 0 | 0 | 0 | 0 | 1 | 0 | 0 | 0 | 0 | 0 | 0 |
| 48.98815 | 14.40994 | CZB_R_4 | czbr4_2 | Czech | R | 0 | 0 | 0 | 0 | 0 | 0 | 1 | 393,784 | 1448,475896 | 4 | 5748,788 | 324 | 0 | 1 | 0 | 0 | 0 | 0 | 1 | 0 | 1 | 1 | 1 | 0 | 0 | 0 | 0 | 1 |
| 48.98556 | 14.41099 | CZB_R_4 | czbr4_3 | Czech | R | 2 | 1 | 0 | 0 | 0 | 0 | 1 | 396,564 | 1448,259147 | 4 | 5393,382 | 646 | 1 | 0 | 0 | 1 | 0 | 0 | 0 | 0 | 0 | 0 | 1 | 0 | 0 | 0 | 0 | 1 |
| 48.98479 | 14.41485 | CZB_R_4 | czbr4_4 | Czech | R | 3 | 1 | 0 | 0 | 0 | 0 | 0 | 403,244 | 1447,97194 | 4 | 35184,335 | 0 | 1 | 0 | 0 | 1 | 0 | 0 | 0 | 0 | 0 | 0 | 0 | 0 | 0 | 0 | 0 | 1 |
| 48.98452 | 14.41886 | CZB_R_4 | czbr4_5 | Czech | R | 0 | 0 | 0 | 0 | 0 | 0 | 1 | 405 | 1447,693309 | 4 | 34804,911 | 0 | 1 | 0 | 0 | 0 | 1 | 0 | 0 | 0 | 0 | 0 | 0 | 0 | 0 | 0 | 0 | 0 |
| 48.97326 | 14.46093 | CZB_U_1 | czbu1_1 | Czech | U | 2 | 1 | 0 | 0 | 0 | 0 | 1 | 385,736 | 1444,428618 | 1 | 20663,764 | 6572 | 0 | 1 | 0 | 0 | 1 | 0 | 1 | 0 | 1 | 0 | 1 | 0 | 0 | 0 | 1 | 1 |
| 48.97318 | 14.45553 | CZB_U_1 | czbu1_2 | Czech | U | 0 | 0 | 0 | 0 | 0 | 0 | 1 | 387,399 | 1444,75048 | 1 | 16848,364 | 12738 | 0 | 1 | 0 | 0 | 1 | 0 | 1 | 1 | 1 | 0 | 1 | 1 | 1 | 0 | 0 | 1 |
| 48.9725 | 14.45419 | CZB_U_1 | czbu1_3 | Czech | U | 1 | 1 | 0 | 0 | 0 | 0 | 1 | 387 | 1444,804488 | 1 | 23626,238 | 3281 | 1 | 1 | 0 | 0 | 1 | 0 | 1 | 0 | 1 | 0 | 0 | 1 | 0 | 0 | 0 | 1 |
| 48.9705 | 14.45217 | CZB_U_1 | czbu1_4 | Czech | U | 0 | 0 | 0 | 0 | 0 | 0 | 1 | 387 | 1444,81434 | 1 | 36173,234 | 0 | 1 | 1 | 0 | 0 | 1 | 0 | 0 | 0 | 0 | 1 | 0 | 0 | 0 | 0 | 0 | 1 |
| 48.96847 | 14.45415 | CZB_U_1 | czbu1_5 | Czech | U | 1 | 1 | 0 | 0 | 0 | 0 | 0 | 387 | 1444,570549 | 1 | 43025,273 | 0 | 1 | 1 | 0 | 0 | 1 | 0 | 0 | 0 | 0 | 1 | 0 | 0 | 0 | 0 | 0 | 0 |
| 48.97667 | 14.47723 | CZB_U_2 | czbu2_1 | Czech | U | 2 | 1 | 0 | 0 | 0 | 0 | 0 | 387,988 | 1443,615169 | 1 | 16684,938 | 29929 | 0 | 1 | 0 | 0 | 1 | 0 | 0 | 1 | 0 | 0 | 0 | 0 | 0 | 1 | 0 | 0 |
| 48.9782 | 14.47407 | CZB_U_2 | czbu2_2 | Czech | U | 2 | 1 | 0 | 0 | 0 | 0 | 0 | 386,833 | 1443,901612 | 1 | 13091,29 | 30529 | 0 | 1 | 0 | 0 | 1 | 0 | 0 | 1 | 0 | 0 | 0 | 0 | 0 | 1 | 0 | 0 |
| 48.98046 | 14.47205 | CZB_U_2 | czbu2_3 | Czech | U | 1 | 1 | 0 | 0 | 0 | 0 | 0 | 385 | 1444,15971 | 1 | 5403,796 | 27411 | 0 | 1 | 0 | 0 | 0 | 0 | 1 | 1 | 0 | 0 | 0 | 0 | 0 | 1 | 0 | 0 |
| 48.98186 | 14.46876 | CZB_U_2 | czbu2_4 | Czech | U | 1 | 1 | 0 | 0 | 0 | 0 | 0 | 385 | 1444,446293 | 1 | 17171,789 | 20260 | 0 | 1 | 0 | 0 | 0 | 0 | 1 | 1 | 0 | 0 | 0 | 0 | 1 | 1 | 0 | 0 |
| 48.98416 | 14.46789 | CZB_U_2 | czbu2_5 | Czech | U | 0 | 0 | 0 | 0 | 0 | 0 | 0 | 385,394 | 1444,635555 | 1 | 28831,166 | 11317 | 1 | 1 | 0 | 0 | 1 | 0 | 1 | 1 | 0 | 0 | 0 | 0 | 1 | 1 | 0 | 0 |
| 50.81273 | 19.15196 | CZE_R_1 | czer1_1 | Poland | R | 0 | 0 | 0 | 0 | 0 | 0 | 0 | 273,712 | 1324,77807 | 170 | 42650,897 | 0 | 1 | 1 | 0 | 1 | 0 | 0 | 0 | 0 | 0 | 0 | 0 | 0 | 0 | 0 | 0 | 0 |
| 50.81214 | 19.15508 | CZE_R_1 | czer1_2 | Poland | R | 0 | 0 | 0 | 0 | 0 | 0 | 0 | 280 | 1324,588121 | 170 | 27913,571 | 0 | 1 | 1 | 0 | 1 | 0 | 0 | 0 | 0 | 0 | 0 | 0 | 0 | 0 | 0 | 0 | 0 |
| 50.81348 | 19.15815 | CZE_R_1 | czer1_3 | Poland | R | 0 | 0 | 0 | 0 | 0 | 0 | 0 | 266,952 | 1324,566649 | 170 | 18502,975 | 0 | 1 | 1 | 0 | 1 | 0 | 0 | 0 | 0 | 0 | 0 | 0 | 0 | 0 | 0 | 0 | 0 |
| 50.81525 | 19.15988 | CZE_R_1 | czer1_4 | Poland | R | 0 | 0 | 0 | 0 | 0 | 0 | 0 | 257 | 1324,641729 | 170 | 9785,662 | 0 | 0 | 1 | 0 | 0 | 0 | 0 | 1 | 0 | 1 | 0 | 1 | 0 | 0 | 0 | 0 | 0 |
| 50.81646 | 19.15589 | CZE_R_1 | czer1_5 | Poland | R | 0 | 0 | 0 | 0 | 0 | 0 | 0 | 260,54 | 1324,923115 | 170 | 10923,163 | 0 | 0 | 1 | 0 | 0 | 0 | 0 | 1 | 0 | 1 | 0 | 1 | 0 | 0 | 0 | 0 | 0 |
| 50.8266 | 19.15667 | CZE_R_2 | czer2_1 | Poland | R | 3 | 1 | 0 | 0 | 0 | 0 | 0 | 241 | 1325,760791 | 169 | 37126,55 | 0 | 1 | 1 | 0 | 1 | 0 | 0 | 0 | 0 | 0 | 1 | 0 | 0 | 0 | 0 | 0 | 1 |
| 50.82834 | 19.15993 | CZE_R_2 | czer2_2 | Poland | R | 2 | 1 | 0 | 0 | 0 | 0 | 1 | 241 | 1325,767847 | 169 | 24963,14 | 0 | 1 | 1 | 0 | 1 | 0 | 0 | 0 | 0 | 0 | 1 | 0 | 0 | 0 | 0 | 0 | 0 |
| 50.82885 | 19.16403 | CZE_R_2 | czer2_3 | Poland | R | 1 | 1 | 0 | 0 | 0 | 0 | 1 | 240,86 | 1325,628057 | 169 | 18082,474 | 0 | 1 | 1 | 0 | 1 | 0 | 0 | 0 | 0 | 0 | 0 | 0 | 0 | 0 | 0 | 0 | 1 |
| 50.8303 | 19.16693 | CZE_R_2 | czer2_4 | Poland | R | 0 | 0 | 0 | 0 | 0 | 0 | 1 | 241 | 1325,623943 | 169 | 29459,359 | 0 | 1 | 1 | 0 | 1 | 0 | 0 | 0 | 0 | 0 | 0 | 0 | 0 | 0 | 0 | 0 | 0 |
| 50.83286 | 19.16587 | CZE_R_2 | czer2_5 | Poland | R | 0 | 0 | 0 | 0 | 0 | 0 | 0 | 243,039 | 1325,892696 | 169 | 24811,906 | 3265 | 1 | 1 | 0 | 1 | 0 | 0 | 0 | 0 | 1 | 0 | 1 | 0 | 0 | 0 | 0 | 1 |
| 50.81062 | 19.06964 | CZE_U_1 | czeu1_1 | Poland | U | 1 | 1 | 0 | 0 | 0 | 0 | 1 | 260,069 | 1328,276655 | 170 | 39047,29 | 1359 | 1 | 0 | 0 | 0 | 1 | 0 | 0 | 0 | 1 | 1 | 0 | 1 | 0 | 0 | 0 | 0 |
| 50.81218 | 19.07204 | CZE_U_1 | czeu1_2 | Poland | U | 0 | 0 | 0 | 0 | 0 | 0 | 1 | 259,948 | 1328,301596 | 170 | 51498,381 | 0 | 1 | 0 | 0 | 0 | 1 | 0 | 0 | 0 | 0 | 1 | 0 | 0 | 0 | 0 | 0 | 0 |
| 50.81199 | 19.07659 | CZE_U_1 | czeu1_3 | Poland | U | 0 | 0 | 0 | 0 | 0 | 0 | 0 | 257,881 | 1328,085587 | 170 | 56718,952 | 0 | 1 | 1 | 0 | 0 | 1 | 0 | 0 | 0 | 0 | 0 | 0 | 0 | 0 | 0 | 0 | 0 |
| 50.80992 | 19.07846 | CZE_U_1 | czeu1_4 | Poland | U | 0 | 0 | 0 | 0 | 0 | 0 | 0 | 254,712 | 1327,825081 | 170 | 39694,763 | 409 | 1 | 1 | 0 | 0 | 1 | 0 | 0 | 0 | 1 | 0 | 1 | 0 | 0 | 0 | 0 | 1 |
| 50.80837 | 19.08048 | CZE_U_1 | czeu1_5 | Poland | U | 1 | 1 | 0 | 0 | 0 | 0 | 1 | 254,964 | 1327,598215 | 170 | 25922,309 | 0 | 1 | 1 | 0 | 0 | 1 | 0 | 0 | 1 | 1 | 1 | 1 | 0 | 0 | 0 | 0 | 0 |
| 50.81414 | 19.09612 | CZE_U_2 | czeu2_1 | Poland | U | 1 | 1 | 0 | 0 | 0 | 0 | 1 | 279,317 | 1327,393854 | 170 | 33274,752 | 11313 | 1 | 1 | 0 | 0 | 1 | 0 | 0 | 1 | 0 | 0 | 0 | 0 | 0 | 1 | 0 | 0 |
| 50.81373 | 19.10046 | CZE_U_2 | czeu2_2 | Poland | U | 1 | 1 | 0 | 0 | 0 | 0 | 0 | 269,26 | 1327,171153 | 170 | 33842,287 | 8079 | 0 | 1 | 0 | 0 | 1 | 0 | 0 | 1 | 0 | 0 | 0 | 0 | 0 | 1 | 0 | 0 |
| 50.81149 | 19.10275 | CZE_U_2 | czeu2_3 | Poland | U | 1 | 1 | 0 | 0 | 0 | 0 | 0 | 267,7 | 1326,870782 | 170 | 48072,067 | 1216 | 1 | 1 | 0 | 0 | 1 | 0 | 0 | 1 | 0 | 0 | 0 | 0 | 1 | 1 | 0 | 0 |
| 50.80938 | 19.1017 | CZE_U_2 | czeu2_4 | Poland | U | 0 | 0 | 0 | 0 | 0 | 0 | 1 | 269,676 | 1326,765377 | 170 | 7024,878 | 13399 | 1 | 1 | 0 | 0 | 1 | 0 | 0 | 1 | 1 | 0 | 0 | 1 | 1 | 1 | 0 | 0 |
| 50.80816 | 19.0987 | CZE_U_2 | czeu2_5 | Poland | U | 0 | 0 | 0 | 0 | 0 | 0 | 0 | 269,496 | 1326,734798 | 170 | 16388,401 | 8704 | 1 | 1 | 0 | 0 | 1 | 0 | 0 | 1 | 1 | 0 | 0 | 1 | 1 | 0 | 0 | 0 |
| 47.59454 | 21.63678 | DEB_R_1 | debr1_1 | Hungary | R | 0 | 0 | 0 | 0 | 0 | 0 | 0 | 128,388 | 935,1223032 | 401 | 8211,806 | 200 | 1 | 1 | 0 | 1 | 0 | 0 | 0 | 1 | 0 | 0 | 0 | 0 | 0 | 0 | 0 | 0 |
| 47.59338 | 21.63979 | DEB_R_1 | debr1_2 | Hungary | R | 1 | 1 | 0 | 0 | 0 | 0 | 1 | 129,244 | 934,8746616 | 401 | 17673,27 | 3223 | 1 | 1 | 0 | 0 | 0 | 0 | 0 | 1 | 0 | 0 | 0 | 0 | 1 | 0 | 0 | 1 |
| 47.59158 | 21.64218 | DEB_R_1 | debr1_3 | Hungary | R | 2 | 1 | 0 | 0 | 0 | 0 | 1 | 127,688 | 934,6056685 | 401 | 14209,459 | 5629 | 1 | 1 | 0 | 0 | 0 | 0 | 0 | 1 | 1 | 0 | 1 | 1 | 0 | 0 | 0 | 0 |
| 47.58965 | 21.6409 | DEB_R_1 | debr1_4 | Hungary | R | 3 | 1 | 0 | 0 | 0 | 0 | 1 | 130,02 | 934,5115004 | 401 | 41083,965 | 0 | 1 | 0 | 0 | 1 | 0 | 0 | 0 | 0 | 0 | 1 | 0 | 0 | 0 | 0 | 0 | 1 |
| 47.5886 | 21.63754 | DEB_R_1 | debr1_5 | Hungary | R | 3 | 1 | 0 | 0 | 0 | 0 | 1 | 134,322 | 934,5950288 | 401 | 48950,53 | 0 | 1 | 0 | 0 | 1 | 0 | 0 | 0 | 0 | 0 | 0 | 0 | 0 | 0 | 0 | 0 | 0 |
| 47.58275 | 21.64667 | DEB_R_2 | debr2_1 | Hungary | R | 2 | 1 | 0 | 0 | 0 | 0 | 1 | 130,866 | 933,6521882 | 402 | 70760,526 | 0 | 1 | 0 | 0 | 1 | 0 | 0 | 0 | 0 | 0 | 0 | 0 | 0 | 0 | 0 | 0 | 0 |
| 47.58274 | 21.65149 | DEB_R_2 | debr2_2 | Hungary | R | 2 | 1 | 0 | 0 | 0 | 0 | 1 | 131,586 | 933,4071699 | 402 | 65864,274 | 0 | 1 | 1 | 0 | 1 | 0 | 0 | 0 | 0 | 0 | 0 | 0 | 0 | 0 | 0 | 0 | 1 |
| 47.58214 | 21.65604 | DEB_R_2 | debr2_3 | Hungary | R | 0 | 0 | 0 | 0 | 0 | 0 | 1 | 128,22 | 933,1278861 | 402 | 17683,84 | 7886 | 1 | 1 | 0 | 1 | 0 | 0 | 1 | 0 | 1 | 0 | 0 | 1 | 0 | 0 | 1 | 0 |
| 47.58162 | 21.66059 | DEB_R_2 | debr2_4 | Hungary | R | 0 | 0 | 1 | 1 | 0 | 0 | 1 | 128,708 | 932,8550536 | 402 | 12217,557 | 1668 | 0 | 1 | 0 | 0 | 0 | 0 | 0 | 0 | 1 | 0 | 1 | 1 | 0 | 0 | 0 | 0 |
| 47.5794 | 21.66033 | DEB_R_2 | debr2_5 | Hungary | R | 1 | 1 | 0 | 0 | 0 | 0 | 0 | 125,028 | 932,6853066 | 402 | 4179,406 | 13402 | 0 | 1 | 0 | 0 | 0 | 0 | 0 | 1 | 0 | 1 | 1 | 0 | 0 | 0 | 1 | 0 |
| 47.5526 | 21.6243 | DEB_U_1 | debu1_1 | Hungary | U | 3 | 1 | 0 | 0 | 0 | 0 | 1 | 130,36 | 932,3083169 | 402 | 54677,478 | 1109 | 1 | 1 | 1 | 0 | 1 | 0 | 1 | 1 | 0 | 0 | 0 | 0 | 0 | 0 | 0 | 0 |
| 47.55149 | 21.62825 | DEB_U_1 | debu1_2 | Hungary | U | 0 | 0 | 0 | 0 | 0 | 0 | 0 | 130,527 | 932,0161177 | 402 | 47060,202 | 10148 | 1 | 0 | 0 | 0 | 1 | 0 | 0 | 0 | 0 | 0 | 0 | 0 | 0 | 1 | 0 | 1 |
| 47.55059 | 21.63212 | DEB_U_1 | debu1_3 | Hungary | U | 0 | 0 | 0 | 0 | 0 | 0 | 0 | 126,678 | 931,7453001 | 402 | 50099,453 | 3838 | 1 | 1 | 0 | 0 | 1 | 0 | 0 | 0 | 0 | 0 | 0 | 0 | 1 | 0 | 0 | 1 |
| 47.54962 | 21.63561 | DEB_U_1 | debu1_4 | Hungary | U | 0 | 0 | 0 | 0 | 0 | 0 | 0 | 126,368 | 931,489352 | 402 | 45897,071 | 6220 | 1 | 1 | 0 | 0 | 1 | 0 | 1 | 1 | 0 | 0 | 0 | 0 | 1 | 0 | 0 | 0 |
| 47.5473 | 21.63836 | DEB_U_1 | debu1_5 | Hungary | U | 0 | 0 | 0 | 0 | 0 | 0 | 0 | 123,931 | 931,159243 | 402 | 15180,967 | 22005 | 0 | 1 | 0 | 0 | 0 | 0 | 1 | 1 | 0 | 0 | 1 | 1 | 0 | 1 | 1 | 0 |
| 47.51778 | 21.60288 | DEB_U_2 | debu2_1 | Hungary | U | 0 | 0 | 1 | 1 | 0 | 0 | 0 | 107,016 | 930,5501941 | 400 | 29142,48 | 6953 | 0 | 1 | 0 | 0 | 0 | 0 | 1 | 1 | 0 | 0 | 0 | 0 | 1 | 0 | 0 | 0 |
| 47.51563 | 21.60198 | DEB_U_2 | debu2_2 | Hungary | U | 0 | 0 | 0 | 0 | 0 | 0 | 0 | 115,676 | 930,4197274 | 400 | 23681,061 | 6167 | 0 | 1 | 0 | 0 | 0 | 0 | 1 | 1 | 0 | 0 | 0 | 0 | 1 | 0 | 1 | 0 |
| 47.51411 | 21.59889 | DEB_U_2 | debu2_3 | Hungary | U | 2 | 1 | 0 | 0 | 1 | 1 | 0 | 112,589 | 930,4538318 | 400 | 22071,81 | 3079 | 1 | 1 | 0 | 1 | 0 | 0 | 1 | 1 | 0 | 0 | 1 | 0 | 0 | 1 | 0 | 0 |
| 47.51453 | 21.59518 | DEB_U_2 | debu2_4 | Hungary | U | 1 | 1 | 0 | 0 | 0 | 0 | 0 | 107,692 | 930,6773085 | 400 | 8196,296 | 0 | 1 | 1 | 0 | 1 | 0 | 0 | 1 | 0 | 1 | 0 | 0 | 0 | 0 | 0 | 0 | 0 |
| 47.51621 | 21.59327 | DEB_U_2 | debu2_5 | Hungary | U | 0 | 0 | 0 | 0 | 0 | 0 | 0 | 108,081 | 930,912763 | 400 | 22926,481 | 0 | 1 | 1 | 0 | 1 | 0 | 0 | 0 | 0 | 0 | 0 | 0 | 0 | 0 | 0 | 0 | 1 |

| | | | | | | | | | | | | | | | | | | | | | | | | | | | | | | | | | |
|---|---|---|---|---|---|---|---|---|---|---|---|---|---|---|---|---|---|---|---|---|---|---|---|---|---|---|---|---|---|---|---|---|---|
| 44.67006 | 22.70094 | DRO_R_1 | dror1_1 | Romania | R | 0 | 0 | 1 | 1 | 0 | 0 | 0 | 60,984 | 656,6387364 | 336 | 3253,843 | 0 | 0 | 1 | 0 | 1 | 0 | 0 | 0 | 0 | 0 | 1 | 0 | 0 | 0 | 0 | 0 | 0 |
| 44.67258 | 22.69891 | DRO_R_1 | dror1_2 | Romania | R | 0 | 0 | 0 | 0 | 0 | 0 | 0 | 60,022 | 656,9307444 | 336 | 21421,443 | 0 | 1 | 1 | 0 | 1 | 0 | 0 | 0 | 0 | 0 | 1 | 0 | 0 | 0 | 0 | 0 | 0 |
| 44.67522 | 22.70133 | DRO_R_1 | dror1_3 | Romania | R | 0 | 0 | 0 | 0 | 0 | 0 | 0 | 72,996 | 656,9416634 | 336 | 11039,391 | 1381 | 0 | 1 | 0 | 0 | 0 | 0 | 0 | 1 | 1 | 1 | 1 | 0 | 0 | 0 | 0 | 0 |
| 44.67764 | 22.70077 | DRO_R_1 | dror1_4 | Romania | R | 0 | 0 | 0 | 0 | 0 | 0 | 0 | 71,389 | 657,1358772 | 336 | 17278,926 | 3131 | 0 | 1 | 0 | 1 | 0 | 0 | 0 | 1 | 1 | 1 | 1 | 0 | 0 | 0 | 0 | 1 |
| 44.67993 | 22.7005 | DRO_R_1 | dror1_5 | Romania | R | 0 | 0 | 0 | 0 | 0 | 0 | 0 | 76,95 | 657,3000657 | 336 | 9715,015 | 13678 | 1 | 1 | 0 | 1 | 0 | 0 | 0 | 1 | 1 | 0 | 0 | 1 | 0 | 0 | 0 | 0 |
| 44.60957 | 22.77011 | DRO_R_2 | dror2_1 | Romania | R | 0 | 0 | 0 | 0 | 0 | 0 | 0 | 110,452 | 648,297681 | 340 | 40938,34 | 337 | 1 | 1 | 0 | 1 | 0 | 0 | 1 | 0 | 1 | 0 | 1 | 0 | 0 | 0 | 0 | 0 |
| 44.61143 | 22.77254 | DRO_R_2 | dror2_2 | Romania | R | 0 | 0 | 1 | 1 | 0 | 0 | 0 | 119,459 | 648,259108 | 340 | 32440,07 | 1513 | 1 | 1 | 0 | 1 | 0 | 0 | 1 | 0 | 1 | 0 | 1 | 0 | 0 | 0 | 0 | 0 |
| 44.61374 | 22.77439 | DRO_R_2 | dror2_3 | Romania | R | 1 | 1 | 3 | 1 | 1 | 1 | 0 | 125,771 | 648,2854832 | 340 | 22211,357 | 2100 | 1 | 1 | 0 | 1 | 0 | 0 | 1 | 0 | 1 | 0 | 1 | 0 | 0 | 0 | 0 | 0 |
| 44.61686 | 22.77537 | DRO_R_2 | dror2_4 | Romania | R | 1 | 1 | 0 | 0 | 0 | 0 | 0 | 131,392 | 648,4163961 | 340 | 38945,658 | 3405 | 1 | 1 | 0 | 1 | 0 | 0 | 0 | 0 | 1 | 0 | 0 | 1 | 0 | 0 | 0 | 0 |
| 44.61935 | 22.77734 | DRO_R_2 | dror2_5 | Romania | R | 0 | 0 | 0 | 0 | 0 | 0 | 0 | 156,124 | 648,4473275 | 340 | 28001,265 | 6798 | 1 | 1 | 0 | 1 | 0 | 0 | 0 | 1 | 1 | 0 | 0 | 1 | 0 | 0 | 0 | 0 |
| 44.64109 | 22.65865 | DRO_U_1 | drou1_1 | Romania | U | 0 | 0 | 0 | 0 | 0 | 0 | 1 | 76,871 | 657,566231 | 333 | 20272,665 | 14069 | 0 | 1 | 0 | 0 | 0 | 0 | 1 | 0 | 0 | 0 | 0 | 1 | 1 | 0 | 0 | 0 |
| 44.64307 | 22.65659 | DRO_U_1 | drou1_2 | Romania | U | 0 | 0 | 0 | 0 | 0 | 0 | 0 | 76,052 | 657,8233005 | 333 | 16249,27 | 11040 | 0 | 1 | 0 | 0 | 0 | 0 | 0 | 1 | 1 | 1 | 0 | 1 | 1 | 0 | 0 | 0 |
| 44.64491 | 22.65384 | DRO_U_1 | drou1_3 | Romania | U | 0 | 0 | 0 | 0 | 0 | 0 | 0 | 78,233 | 658,1214231 | 333 | 8924,378 | 9654 | 0 | 1 | 0 | 0 | 0 | 0 | 0 | 1 | 1 | 0 | 0 | 1 | 0 | 1 | 1 | 0 |
| 44.64693 | 22.65111 | DRO_U_1 | drou1_4 | Romania | U | 0 | 0 | 0 | 0 | 0 | 0 | 0 | 81,892 | 658,426762 | 333 | 21703,828 | 7882 | 1 | 1 | 0 | 0 | 1 | 0 | 0 | 0 | 1 | 1 | 0 | 1 | 0 | 0 | 0 | 0 |
| 44.64665 | 22.64768 | DRO_U_1 | drou1_5 | Romania | U | 0 | 0 | 2 | 1 | 0 | 0 | 0 | 84,961 | 658,6343665 | 333 | 43187,573 | 2087 | 1 | 1 | 0 | 0 | 1 | 0 | 1 | 1 | 0 | 1 | 1 | 0 | 0 | 1 | 0 | 0 |
| 44.62415 | 22.64235 | DRO_U_2 | drou2_1 | Romania | U | 0 | 0 | 0 | 0 | 0 | 0 | 1 | 70,94 | 657,5698639 | 333 | 8263,999 | 18958 | 0 | 1 | 0 | 0 | 0 | 0 | 1 | 1 | 0 | 0 | 0 | 1 | 0 | 1 | 0 | 0 |
| 44.62303 | 22.64497 | DRO_U_2 | drou2_2 | Romania | U | 0 | 0 | 1 | 1 | 0 | 0 | 0 | 62,448 | 657,3280521 | 333 | 19640,14 | 12492 | 0 | 1 | 0 | 0 | 1 | 0 | 0 | 1 | 0 | 0 | 0 | 1 | 0 | 1 | 1 | 0 |
| 44.62245 | 22.64828 | DRO_U_2 | drou2_3 | Romania | U | 0 | 0 | 0 | 0 | 0 | 0 | 0 | 51,46 | 657,0741675 | 333 | 34016,893 | 9493 | 1 | 1 | 1 | 0 | 1 | 0 | 1 | 0 | 0 | 0 | 0 | 1 | 0 | 1 | 0 | 0 |
| 44.62287 | 22.65197 | DRO_U_2 | drou2_4 | Romania | U | 0 | 0 | 0 | 0 | 0 | 0 | 0 | 57,328 | 656,8594036 | 333 | 30893,599 | 7142 | 1 | 1 | 1 | 0 | 1 | 0 | 1 | 1 | 0 | 0 | 0 | 1 | 0 | 1 | 0 | 0 |
| 44.62339 | 22.65566 | DRO_U_2 | drou2_5 | Romania | U | 0 | 0 | 0 | 0 | 0 | 0 | 0 | 58,331 | 656,6509591 | 333 | 20635,945 | 17233 | 1 | 1 | 0 | 0 | 1 | 0 | 0 | 1 | 0 | 0 | 0 | 0 | 0 | 1 | 0 | 0 |
| 48.42517 | 21.9629 | ESK_R_1 | eskr1_1 | Slovak | R | 1 | 1 | 1 | 1 | 0 | 0 | 0 | 154,609 | 989,6486917 | 432 | 35054,398 | 0 | 1 | 1 | 0 | 1 | 0 | 0 | 0 | 0 | 0 | 0 | 0 | 0 | 0 | 0 | 0 | 0 |
| 48.42325 | 21.96419 | ESK_R_1 | eskr1_2 | Slovak | R | 0 | 0 | 0 | 0 | 0 | 0 | 0 | 161,527 | 989,3499528 | 432 | 11291,226 | 0 | 1 | 1 | 0 | 1 | 0 | 0 | 0 | 0 | 0 | 0 | 0 | 0 | 0 | 0 | 0 | 0 |
| 48.42096 | 21.96623 | ESK_R_1 | eskr1_3 | Slovak | R | 0 | 0 | 0 | 0 | 0 | 0 | 1 | 176,805 | 989,0503439 | 432 | 14746,028 | 0 | 1 | 1 | 0 | 1 | 0 | 0 | 1 | 0 | 0 | 0 | 0 | 0 | 0 | 0 | 0 | 0 |
| 48.41849 | 21.96677 | ESK_R_1 | eskr1_4 | Slovak | R | 0 | 0 | 0 | 0 | 0 | 0 | 0 | 172,902 | 988,7568379 | 432 | 508,117 | 0 | 0 | 1 | 0 | 1 | 0 | 0 | 0 | 0 | 0 | 0 | 0 | 0 | 0 | 0 | 0 | 1 |
| 48.41593 | 21.96694 | ESK_R_1 | eskr1_5 | Slovak | R | 0 | 0 | 0 | 0 | 0 | 0 | 0 | 165,043 | 988,469039 | 432 | 12879,937 | 3553 | 1 | 1 | 0 | 1 | 0 | 0 | 0 | 0 | 1 | 0 | 1 | 0 | 0 | 0 | 0 | 0 |
| 48.40843 | 21.94994 | ESK_R_2 | eskr2_1 | Slovak | R | 1 | 1 | 0 | 0 | 0 | 0 | 0 | 131,511 | 990,5537422 | 432 | 20082,784 | 176 | 1 | 1 | 0 | 1 | 0 | 0 | 1 | 0 | 1 | 0 | 1 | 0 | 0 | 0 | 0 | 0 |
| 48.40639 | 21.95259 | ESK_R_2 | eskr2_2 | Slovak | R | 0 | 0 | 0 | 0 | 0 | 0 | 0 | 106,356 | 990,3187888 | 432 | 16770,181 | 1083 | 1 | 1 | 0 | 1 | 0 | 0 | 0 | 0 | 1 | 0 | 1 | 0 | 0 | 0 | 0 | 0 |
| 48.40441 | 21.95531 | ESK_R_2 | eskr2_3 | Slovak | R | 0 | 0 | 0 | 0 | 0 | 0 | 0 | 98,986 | 990,0241989 | 432 | 16760,103 | 1227 | 1 | 1 | 0 | 1 | 0 | 0 | 0 | 0 | 1 | 0 | 1 | 0 | 0 | 0 | 0 | 0 |
| 48.40207 | 21.95732 | ESK_R_2 | eskr2_4 | Slovak | R | 0 | 0 | 0 | 0 | 0 | 0 | 1 | 99,841 | 989,7778693 | 432 | 37878,6 | 840 | 1 | 1 | 0 | 1 | 0 | 0 | 0 | 0 | 1 | 0 | 1 | 0 | 0 | 0 | 0 | 0 |
| 48.40015 | 21.95991 | ESK_R_2 | eskr2_5 | Slovak | R | 0 | 0 | 0 | 0 | 0 | 0 | 1 | 99,365 | 989,5439733 | 432 | 36723,208 | 766 | 1 | 1 | 0 | 1 | 0 | 0 | 0 | 0 | 1 | 0 | 1 | 0 | 0 | 0 | 0 | 1 |
| 48.39587 | 21.99798 | ESK_U_1 | esku1_1 | Slovak | U | 0 | 0 | 2 | 1 | 0 | 0 | 1 | 99 | 986,3698267 | 435 | 11385,408 | 2394 | 0 | 1 | 0 | 0 | 0 | 0 | 0 | 0 | 1 | 0 | 0 | 1 | 0 | 0 | 0 | 0 |
| 48.3936 | 22.00004 | ESK_U_1 | esku1_2 | Slovak | U | 0 | 0 | 0 | 0 | 0 | 0 | 1 | 102,144 | 986,0870228 | 435 | 9612,085 | 6022 | 0 | 1 | 0 | 0 | 0 | 0 | 0 | 0 | 1 | 0 | 0 | 1 | 0 | 0 | 0 | 0 |
| 48.39281 | 22.00385 | ESK_U_1 | esku1_3 | Slovak | U | 0 | 0 | 0 | 0 | 0 | 0 | 0 | 99,124 | 985,8451221 | 435 | 6781,035 | 5370 | 1 | 1 | 0 | 1 | 0 | 0 | 0 | 0 | 1 | 0 | 0 | 1 | 0 | 0 | 0 | 0 |
| 48.39018 | 22.00455 | ESK_U_1 | esku1_4 | Slovak | U | 0 | 0 | 1 | 1 | 0 | 0 | 0 | 100,648 | 985,5747872 | 435 | 15164,828 | 7171 | 1 | 1 | 0 | 1 | 0 | 0 | 0 | 0 | 1 | 0 | 0 | 1 | 0 | 0 | 0 | 0 |
| 48.38925 | 22.00781 | ESK_U_1 | esku1_5 | Slovak | U | 0 | 0 | 0 | 0 | 0 | 0 | 1 | 99 | 985,3446134 | 435 | 29270,438 | 1009 | 1 | 1 | 0 | 1 | 0 | 0 | 0 | 0 | 0 | 0 | 0 | 0 | 0 | 0 | 0 | 0 |
| 48.42998 | 21.974 | ESK_U_2 | esku2_1 | Slovak | U | 1 | 1 | 0 | 0 | 0 | 0 | 0 | 105,443 | 990,2274772 | 433 | 13043,953 | 7890 | 0 | 1 | 0 | 1 | 0 | 1 | 0 | 0 | 1 | 0 | 0 | 1 | 0 | 0 | 0 | 1 |
| 48.42729 | 21.97378 | ESK_U_2 | esku2_2 | Slovak | U | 0 | 0 | 0 | 0 | 0 | 0 | 0 | 112,148 | 990,4647682 | 433 | 18430,603 | 11067 | 1 | 1 | 0 | 1 | 0 | 0 | 0 | 0 | 1 | 0 | 0 | 1 | 0 | 1 | 0 | 0 |
| 48.42474 | 21.97596 | ESK_U_2 | esku2_3 | Slovak | U | 0 | 0 | 0 | 0 | 0 | 0 | 1 | 119,152 | 989,9223771 | 433 | 19269,438 | 12722 | 0 | 1 | 0 | 0 | 0 | 0 | 0 | 1 | 1 | 0 | 0 | 1 | 0 | 1 | 0 | 0 |
| 48.42289 | 21.97886 | ESK_U_2 | esku2_4 | Slovak | U | 0 | 0 | 0 | 0 | 0 | 0 | 0 | 112,596 | 989,6280834 | 433 | 8765,548 | 15995 | 0 | 1 | 0 | 0 | 0 | 0 | 0 | 1 | 1 | 0 | 0 | 1 | 1 | 1 | 0 | 0 |
| 48.42034 | 21.98033 | ESK_U_2 | esku2_5 | Slovak | U | 0 | 0 | 0 | 0 | 0 | 0 | 1 | 116,812 | 989,3341169 | 433 | 9361,132 | 14456 | 0 | 1 | 0 | 0 | 0 | 0 | 0 | 1 | 1 | 0 | 0 | 1 | 1 | 0 | 0 | 0 |
| 52.14510 | 21.17630 | FAL_R_1 | falr1_1 | Poland | R | 1 | 1 | 0 | 0 | 0 | 0 | 1 | 86 | 1368,13667 | 46 | 36879,891 | 500 | 1 | 1 | 1 | 1 | 0 | 0 | 0 | 0 | 0 | 0 | 1 | 0 | 0 | 0 | 0 | 0 |
| 52.14312 | 21.17640 | FAL_R_1 | falr1_2 | Poland | R | 1 | 1 | 0 | 0 | 0 | 0 | 1 | 86 | 1367,941625 | 46 | 40958,191 | 0 | 1 | 1 | 0 | 1 | 0 | 0 | 0 | 0 | 0 | 0 | 0 | 0 | 0 | 0 | 0 | 0 |
| 52.14147 | 21.17579 | FAL_R_1 | falr1_3 | Poland | R | 1 | 1 | 0 | 0 | 0 | 0 | 1 | 84,432 | 1367,800059 | 46 | 36321,563 | 0 | 1 | 1 | 0 | 1 | 0 | 0 | 0 | 0 | 0 | 1 | 0 | 0 | 0 | 0 | 0 | 0 |
| 52.13893 | 21.17849 | FAL_R_1 | falr1_4 | Poland | R | 1 | 1 | 0 | 0 | 0 | 0 | 1 | 87 | 1367,464138 | 46 | 42608,305 | 0 | 1 | 1 | 0 | 1 | 0 | 0 | 0 | 0 | 0 | 1 | 0 | 0 | 0 | 0 | 0 | 0 |
| 52.13714 | 21.18176 | FAL_R_1 | falr1_5 | Poland | R | 0 | 0 | 0 | 0 | 0 | 0 | 1 | 86,567 | 1367,181477 | 46 | 36743,918 | 22 | 1 | 1 | 0 | 1 | 0 | 0 | 0 | 0 | 0 | 1 | 0 | 0 | 0 | 0 | 0 | 0 |
| 54.3529 | 18.79458 | GDA_R_1 | gdar1_1 | Poland | R | 1 | 1 | 0 | 0 | 0 | 0 | 0 | 1,12 | 1660,499031 | -216 | 33300,647 | 0 | 1 | 0 | 1 | 1 | 0 | 0 | 0 | 0 | 0 | 0 | 0 | 0 | 0 | 0 | 0 | 0 |
| 54.3513 | 18.79089 | GDA_R_1 | gdar1_2 | Poland | R | 1 | 1 | 0 | 0 | 0 | 0 | 0 | 1,32 | 1660,473119 | -216 | 18928,535 | 1783 | 1 | 1 | 1 | 1 | 0 | 0 | 0 | 1 | 0 | 1 | 1 | 0 | 0 | 0 | 0 | 0 |
| 54.34993 | 18.79529 | GDA_R_1 | gdar1_3 | Poland | R | 1 | 1 | 0 | 0 | 1 | 1 | 0 | 3,781 | 1660,189664 | -216 | 33839,441 | 2767 | 1 | 0 | 1 | 1 | 0 | 0 | 0 | 0 | 0 | 0 | 0 | 1 | 0 | 0 | 0 | 0 |
| 54.34912 | 18.80048 | GDA_R_1 | gdar1_4 | Poland | R | 1 | 1 | 0 | 0 | 0 | 0 | 0 | 3 | 1659,937506 | -216 | 29064,646 | 3306 | 1 | 1 | 1 | 1 | 0 | 0 | 0 | 1 | 1 | 0 | 1 | 1 | 1 | 0 | 0 | 0 |
| 54.34883 | 18.80512 | GDA_R_1 | gdar1_5 | Poland | R | 0 | 0 | 0 | 0 | 0 | 0 | 0 | 3 | 1659,748358 | -216 | 24253,12 | 2166 | 1 | 1 | 1 | 1 | 0 | 0 | 1 | 1 | 1 | 0 | 0 | 1 | 0 | 0 | 0 | 0 |
| 54.33857 | 18.89419 | GDA_R_2 | gdar2_1 | Poland | R | 1 | 1 | 0 | 0 | 0 | 0 | 0 | 2 | 1655,764971 | -215 | 24536,608 | 2969 | 1 | 1 | 1 | 1 | 0 | 0 | 0 | 1 | 1 | 0 | 0 | 1 | 0 | 0 | 0 | 1 |
| 54.33661 | 18.89608 | GDA_R_2 | gdar2_2 | Poland | R | 1 | 1 | 0 | 0 | 0 | 0 | 0 | 3,977 | 1655,498643 | -215 | 38317,332 | 3327 | 1 | 1 | 1 | 1 | 0 | 0 | 1 | 1 | 1 | 0 | 1 | 1 | 1 | 0 | 0 | 0 |
| 54.33694 | 18.90026 | GDA_R_2 | gdar2_3 | Poland | R | 1 | 1 | 0 | 0 | 0 | 0 | 0 | 2,064 | 1655,386983 | -215 | 32084,366 | 873 | 1 | 1 | 1 | 1 | 0 | 0 | 0 | 0 | 1 | 0 | 1 | 0 | 0 | 0 | 0 | 1 |
| 54.33633 | 18.90566 | GDA_R_2 | gdar2_4 | Poland | R | 0 | 0 | 0 | 0 | 0 | 0 | 0 | 3,296 | 1655,145087 | -215 | 30499,473 | 3027 | 0 | 1 | 1 | 1 | 0 | 0 | 0 | 1 | 1 | 0 | 1 | 1 | 0 | 0 | 0 | 0 |
| 54.33714 | 18.90931 | GDA_R_2 | gdar2_5 | Poland | R | 0 | 0 | 0 | 0 | 0 | 0 | 0 | 3 | 1655,098215 | -215 | 11517,553 | 6177 | 0 | 1 | 0 | 0 | 0 | 0 | 1 | 1 | 1 | 0 | 1 | 1 | 1 | 0 | 0 | 0 |
| 54.41288 | 18.56046 | GDA_U_1 | gdau1_1 | Poland | U | 1 | 1 | 0 | 0 | 0 | 0 | 1 | 26 | 1674,207172 | -226 | 47228,391 | 1126 | 1 | 1 | 0 | 0 | 1 | 1 | 0 | 1 | 0 | 0 | 0 | 0 | 1 | 0 | 0 | 0 |
| 54.41395 | 18.56501 | GDA_U_1 | gdau1_2 | Poland | U | 0 | 0 | 0 | 0 | 0 | 0 | 1 | 23 | 1674,145532 | -226 | 14056,165 | 6837 | 0 | 1 | 0 | 0 | 0 | 0 | 1 | 1 | 0 | 0 | 0 | 0 | 1 | 0 | 1 | 0 |
| 54.41443 | 18.56945 | GDA_U_1 | gdau1_3 | Poland | U | 0 | 0 | 0 | 0 | 0 | 0 | 1 | 18,947 | 1674,041497 | -226 | 22844,808 | 7189 | 0 | 1 | 0 | 0 | 0 | 0 | 0 | 1 | 1 | 1 | 1 | 0 | 1 | 0 | 1 | 0 |
| 54.4154 | 18.57313 | GDA_U_1 | gdau1_4 | Poland | U | 0 | 0 | 0 | 0 | 0 | 0 | 1 | 16,56 | 1674,00796 | -226 | 13841,957 | 4831 | 0 | 1 | 0 | 0 | 0 | 0 | 1 | 1 | 1 | 1 | 0 | 1 | 0 | 0 | 0 | 0 |
| 54.41566 | 18.57694 | GDA_U_1 | gdau1_5 | Poland | U | 0 | 0 | 0 | 0 | 0 | 0 | 1 | 15,026 | 1673,896361 | -226 | 10912,96 | 6863 | 0 | 1 | 0 | 0 | 0 | 0 | 1 | 1 | 1 | 1 | 0 | 1 | 1 | 0 | 0 | 0 |
| 54.42454 | 18.59385 | GDA_U_2 | gdau2_1 | Poland | U | 1 | 1 | 0 | 0 | 0 | 0 | 1 | 2,564 | 1674,153266 | -226 | 29018,661 | 1875 | 0 | 1 | 1 | 0 | 1 | 0 | 0 | 0 | 0 | 1 | 0 | 1 | 0 | 0 | 0 | 0 |
| 54.42169 | 18.59492 | GDA_U_2 | gdau2_2 | Poland | U | 1 | 1 | 0 | 0 | 1 | 1 | 0 | 2,06 | 1673,849537 | -226 | 51562,708 | 1011 | 1 | 1 | 0 | 0 | 1 | 0 | 0 | 1 | 0 | 0 | 0 | 0 | 1 | 0 | 0 | 0 |
| 54.41982 | 18.59799 | GDA_U_2 | gdau2_3 | Poland | U | 2 | 1 | 0 | 0 | 0 | 0 | 1 | 2,153 | 1673,56308 | -226 | 54762,072 | 46 | 1 | 1 | 0 | 0 | 1 | 0 | 0 | 0 | 0 | 0 | 0 | 1 | 1 | 0 | 0 | 0 |
| 54.4178 | 18.60194 | GDA_U_2 | gdau2_4 | Poland | U | 4 | 1 | 0 | 0 | 0 | 0 | 1 | 2,921 | 1673,244888 | -226 | 44580,768 | 1866 | 1 | 1 | 1 | 0 | 1 | 0 | 0 | 0 | 0 | 1 | 0 | 0 | 0 | 1 | 0 | 0 |
| 54.41585 | 18.6049 | GDA_U_2 | gdau2_5 | Poland | U | 1 | 1 | 0 | 0 | 0 | 0 | 1 | 2,563 | 1672,956777 | -226 | 31474,948 | 0 | 1 | 1 | 1 | 1 | 1 | 0 | 0 | 1 | 0 | 1 | 0 | 0 | 0 | 0 | 0 | 0 |
| 52.19558 | 20.73707 | GOL_R_1 | golr1_1 | Poland | R | 0 | 0 | 0 | 0 | 0 | 0 | 1 | 91 | 1388,007933 | 36 | 5448 | 0 | 0 | 1 | 0 | 0 | 0 | 0 | 1 | 0 | 0 | 0 | 0 | 0 | 0 | 0 | 0 | 0 |
| 52.1981 | 20.7387 | GOL_R_1 | golr1_2 | Poland | R | 1 | 1 | 0 | 0 | 0 | 0 | 1 | 92 | 1388,183413 | 36 | 25264 | 0 | 1 | 1 | 0 | 0 | 0 | 0 | 1 | 0 | 1 | 0 | 0 | 0 | 0 | 0 | 0 | 0 |
| 52.19717 | 20.74241 | GOL_R_1 | golr1_3 | Poland | R | 0 | 0 | 0 | 0 | 0 | 0 | 0 | 93 | 1387,975002 | 36 | 10707 | 0 | 0 | 1 | 0 | 0 | 0 | 0 | 0 | 0 | 1 | 0 | 0 | 0 | 0 | 0 | 0 | 0 |
| 52.19549 | 20.7435 | GOL_R_1 | golr1_4 | Poland | R | 0 | 0 | 0 | 0 | 0 | 0 | 0 | 92,142 | 1387,775374 | 36 | 18353 | 31 | 0 | 1 | 0 | 0 | 0 | 0 | 0 | 1 | 0 | 0 | 1 | 0 | 0 | 0 | 1 | 0 |
| 52.19505 | 20.74788 | GOL_R_1 | golr1_5 | Poland | R | 0 | 0 | 0 | 0 | 0 | 0 | 0 | 93 | 1387,580996 | 36 | 13425 | 0 | 0 | 1 | 0 | 0 | 0 | 0 | 0 | 1 | 0 | 1 | 0 | 0 | 0 | 0 | 0 | 0 |
| 51.12673 | 14.94836 | GOR_R_1 | gorr1_1 | Germany | R | 0 | 0 | 1 | 1 | 0 | 0 | 0 | 239,676 | 1557,215066 | -96 | 6233 | 0 | 0 | 1 | 0 | 0 | 0 | 0 | 1 | 0 | 0 | 0 | 0 | 0 | 0 | 0 | 0 | 1 |
| 51.1265 | 14.9445 | GOR_R_1 | gorr1_2 | Germany | R | 0 | 0 | 0 | 0 | 0 | 0 | 0 | 245 | 1557,406066 | -96 | 650 | 0 | 0 | 0 | 0 | 0 | 0 | 0 | 1 | 0 | 0 | 0 | 0 | 0 | 0 | 0 | 0 | 0 |
| 51.12558 | 14.94017 | GOR_R_1 | gorr1_3 | Germany | R | 1 | 1 | 0 | 0 | 0 | 0 | 1 | 258,51 | 1557,571047 | -96 | 33663,971 | 0 | 1 | 1 | 0 | 1 | 0 | 0 | 1 | 1 | 0 | 0 | 0 | 0 | 0 | 0 | 0 | 0 |
| 51.12431 | 14.93582 | GOR_R_1 | gorr1_4 | Germany | R | 1 | 1 | 0 | 0 | 0 | 0 | 1 | 280,895 | 1557,712219 | -96 | 17373,008 | 0 | 1 | 1 | 0 | 1 | 0 | 0 | 1 | 0 | 0 | 0 | 0 | 0 | 0 | 0 | 0 | 1 |
| 51.12528 | 14.93116 | GOR_R_1 | gorr1_5 | Germany | R | 2 | 1 | 0 | 0 | 0 | 0 | 1 | 303,017 | 1558,030375 | -96 | 51138,264 | 0 | 1 | 1 | 0 | 1 | 0 | 0 | 0 | 0 | 0 | 0 | 0 | 0 | 0 | 0 | 0 | 0 |
| 51.11737 | 14.98668 | GOR_R_2 | gorr2_1 | Germany | R | 0 | 0 | 0 | 0 | 0 | 0 | 1 | 191,978 | 1554,495438 | -92 | 16114,728 | 5885,7 | 0 | 1 | 0 | 1 | 0 | 1 | 0 | 1 | 0 | 1 | 1 | 1 | 0 | 0 | 0 | 0 |
| 51.11894 | 14.99028 | GOR_R_2 | gorr2_2 | Germany | R | 2 | 1 | 0 | 0 | 0 | 0 | 0 | 186 | 1554,413448 | -92 | 8195,943 | 426 | 1 | 1 | 0 | 1 | 0 | 0 | 0 | 0 | 1 | 1 | 1 | 0 | 0 | 0 | 0 | 1 |
| 51.12131 | 14.99339 | GOR_R_2 | gorr2_3 | Germany | R | 2 | 1 | 0 | 0 | 0 | 0 | 0 | 185 | 1554,41754 | -92 | 6854,604 | 0 | 0 | 1 | 0 | 1 | 0 | 0 | 0 | 0 | 0 | 1 | 0 | 0 | 0 | 0 | 0 | 1 |
| 51.12361 | 14.99049 | GOR_R_2 | gorr2_4 | Germany | R | 2 | 1 | 0 | 0 | 0 | 0 | 1 | 184,001 | 1554,739765 | -92 | 14658,004 | 0 | 1 | 1 | 0 | 1 | 0 | 0 | 0 | 0 | 0 | 0 | 0 | 0 | 0 | 0 | 0 | 1 |
| 51.12487 | 14.98645 | GOR_R_2 | gorr2_5 | Germany | R | 0 | 0 | 1 | 1 | 0 | 0 | 1 | 187,794 | 1555,04819 | -92 | 9102,415 | 4961,7 | 0 | 1 | 0 | 1 | 0 | 0 | 0 | 0 | 1 | 0 | 1 | 0 | 0 | 0 | 0 | 0 |
| 51.16365 | 14.97461 | GOR_U_1 | goru1_1 | Germany | U | 1 | 1 | 0 | 0 | 0 | 0 | 1 | 211,253 | 1558,471727 | -96 | 29235 | 2054,9 | 1 | 0 | 0 | 1 | 1 | 0 | 0 | 0 | 0 | 0 | 0 | 0 | 0 | 0 | 0 | 0 |
| 51.16132 | 14.97683 | GOR_U_1 | goru1_2 | Germany | U | 1 | 1 | 0 | 0 | 0 | 0 | 1 | 204,596 | 1558,184074 | -96 | 35460 | 5072,6 | 1 | 1 | 0 | 0 | 1 | 0 | 1 | 1 | 0 | 0 | 1 | 1 | 1 | 0 | 0 | 0 |
| 51.1612 | 14.98076 | GOR_U_1 | goru1_3 | Germany | U | 0 | 0 | 0 | 0 | 0 | 0 | 1 | 201,612 | 1557,967773 | -96 | 33283,04 | 5831,5 | 0 | 1 | 0 | 0 | 1 | 0 | 1 | 1 | 1 | 0 | 1 | 0 | 0 | 0 | 1 | 1 |
| 51.16077 | 14.98486 | GOR_U_1 | goru1_4 | Germany | U | 1 | 1 | 0 | 0 | 0 | 0 | 1 | 212,08 | 1557,718808 | -96 | 33889 | 6830,2 | 1 | 1 | 1 | 0 | 0 | 1 | 0 | 1 | 1 | 0 | 1 | 0 | 0 | 1 | 0 | 1 |
| 51.1634 | 14.98533 | GOR_U_1 | goru1_5 | Germany | U | 0 | 0 | 0 | 0 | 0 | 0 | 1 | 220,143 | 1557,881112 | -96 | 46931 | 802 | 1 | 1 | 1 | 0 | 0 | 1 | 0 | 1 | 0 | 0 | 1 | 0 | 0 | 0 | 0 | 1 |
| 51.14285 | 14.98167 | GOR_U_2 | goru2_1 | Germany | U | 0 | 0 | 1 | 1 | 0 | 0 | 1 | 225,257 | 1556,596437 | -98 | 31588,71 | 13496,3 | 1 | 1 | 0 | 0 | 1 | 0 | 0 | 1 | 1 | 0 | 1 | 0 | 1 | 1 | 0 | 0 |
| 51.14216 | 14.98589 | GOR_U_2 | goru2_2 | Germany | U | 0 | 0 | 0 | 0 | 0 | 0 | 0 | 222,552 | 1556,321377 | -98 | 21070,98 | 14796,6 | 1 | 1 | 0 | 0 | 1 | 0 | 0 | 1 | 1 | 0 | 1 | 0 | 1 | 1 | 0 | 0 |
| 51.14147 | 14.98838 | GOR_U_2 | goru2_3 | Germany | U | 2 | 1 | 0 | 0 | 0 | 0 | 1 | 207,057 | 1556,137608 | -98 | 43929 | 9711,3 | 1 | 1 | 0 | 0 | 1 | 0 | 0 | 1 | 0 | 1 | 0 | 0 | 1 | 1 | 0 | 1 |
| 51.13881 | 14.98852 | GOR_U_2 | goru2_4 | Germany | U | 2 | 1 | 0 | 0 | 0 | 0 | 1 | 209,721 | 1555,937538 | -98 | 51705,421 | 10878,4 | 1 | 1 | 0 | 0 | 1 | 0 | 0 | 0 | 1 | 0 | 1 | 0 | 0 | 1 | 0 | 0 |
| 51.13751 | 14.985 | GOR_U_2 | goru2_5 | Germany | U | 2 | 1 | 0 | 0 | 0 | 0 | 1 | 227,036 | 1556,035057 | -98 | 37840,339 | 9020,2 | 1 | 1 | 0 | 0 | 1 | 0 | 0 | 1 | 1 | 1 | 1 | 0 | 0 | 0 | 1 | 1 |
| 50.2875 | 18.95301 | GSL_U_1 | gslu1_1 | Poland | U | 1 | 1 | 0 | 0 | 0 | 0 | 1 | 277 | 1289,410695 | 126 | 45658,891 | 3672 | 1 | 1 | 1 | 0 | 1 | 0 | 0 | 0 | 0 | 0 | 0 | 0 | 1 | 0 | 0 | 0 |
| 50.2883 | 18.94929 | GSL_U_1 | gslu1_2 | Poland | U | 0 | 0 | 0 | 0 | 0 | 0 | 0 | 279,489 | 1289,653248 | 126 | 7952,108 | 11276 | 0 | 1 | 0 | 0 | 0 | 1 | 1 | 1 | 0 | 0 | 0 | 0 | 1 | 1 | 0 | 0 |
| 50.29015 | 18.94989 | GSL_U_1 | gslu1_3 | Poland | U | 0 | 0 | 0 | 0 | 0 | 0 | 0 | 280 | 1289,778977 | 126 | 15343,043 | 11308 | 0 | 1 | 0 | 0 | 0 | 1 | 1 | 1 | 0 | 0 | 0 | 0 | 1 | 1 | 0 | 0 |
| 50.29228 | 18.95131 | GSL_U_1 | gslu1_4 | Poland | U | 0 | 0 | 0 | 0 | 0 | 0 | 1 | 280,641 | 1289,890494 | 126 | 20029,169 | 18109 | 0 | 1 | 0 | 0 | 0 | 1 | 1 | 1 | 0 | 0 | 0 | 0 | 1 | 1 | 0 | 0 |
| 50.29442 | 18.95262 | GSL_U_1 | gslu1_5 | Poland | U | 0 | 0 | 0 | 0 | 0 | 0 | 0 | 285,432 | 1290,006056 | 126 | 17172,239 | 26464 | 1 | 1 | 0 | 0 | 1 | 0 | 1 | 1 | 0 | 0 | 0 | 0 | 1 | 1 | 0 | 0 |
| 50.2957 | 18.95464 | GSL_U_1 | gslu1_6 | Poland | U | 0 | 0 | 0 | 0 | 0 | 0 | 1 | 287,48 | 1290,017091 | 126 | 25424,564 | 21281 | 1 | 1 | 0 | 0 | 1 | 0 | 0 | 1 | 0 | 0 | 0 | 0 | 1 | 1 | 0 | 0 |
| 50.29314 | 18.95566 | GSL_U_1 | gslu1_7 | Poland | U | 0 | 0 | 0 | 0 | 0 | 0 | 1 | 292,114 | 1289,755545 | 126 | 16557,371 | 19227 | 0 | 1 | 0 | 0 | 1 | 0 | 0 | 1 | 0 | 0 | 0 | 0 | 1 | 1 | 1 | 0 |
| 50.2905 | 18.9562 | GSL_U_1 | gslu1_8 | Poland | U | 0 | 0 | 0 | 0 | 0 | 0 | 1 | 284,248 | 1289,510112 | 126 | 13009,695 | 2771,111 | 0 | 1 | 0 | 0 | 1 | 0 | 0 | 1 | 0 | 0 | 0 | 0 | 1 | 0 | 1 | 0 |
| 50.20713 | 15.86187 | HRA_R_1 | hrar1_1 | Czech | R | 2 | 1 | 0 | 0 | 0 | 0 | 1 | 231,243 | 1442,074925 | 1 | 20995,521 | 3124 | 1 | 1 | 0 | 1 | 0 | 0 | 0 | 0 | 1 | 1 | 0 | 1 | 0 | 0 | 0 | 1 |
| 50.20885 | 15.8642 | HRA_R_1 | hrar1_2 | Czech | R | 4 | 1 | 0 | 0 | 0 | 0 | 1 | 230,897 | 1442,068302 | 1 | 47828,296 | 416 | 1 | 1 | 0 | 1 | 0 | 0 | 0 | 0 | 0 | 1 | 1 | 0 | 0 | 0 | 0 | 1 |
| 50.20996 | 15.86794 | HRA_R_1 | hrar1_3 | Czech | R | 4 | 1 | 0 | 0 | 0 | 0 | 1 | 231 | 1441,940912 | 1 | 30053,476 | 317 | 1 | 1 | 0 | 1 | 0 | 0 | 1 | 0 | 0 | 1 | 1 | 0 | 0 | 0 | 0 | 1 |
| 50.21179 | 15.87139 | HRA_R_1 | hrar1_4 | Czech | R | 0 | 0 | 0 | 0 | 0 | 0 | 1 | 231 | 1441,880629 | 1 | 31163,6 | 137 | 1 | 1 | 0 | 1 | 0 | 0 | 1 | 0 | 0 | 1 | 1 | 0 | 0 | 0 | 0 | 1 |
| 50.21058 | 15.87481 | HRA_R_1 | hrar1_5 | Czech | R | 0 | 0 | 0 | 0 | 0 | 0 | 1 | 231 | 1441,606567 | 1 | 12125,612 | 0 | 1 | 1 | 1 | 1 | 0 | 0 | 0 | 0 | 0 | 1 | 0 | 0 | 0 | 0 | 0 | 0 |

| | | | | | | | | | | | | | | | | | | | | | | | | | | | | | | | | | |
|---|---|---|---|---|---|---|---|---|---|---|---|---|---|---|---|---|---|---|---|---|---|---|---|---|---|---|---|---|---|---|---|---|---|
| 50.19453 | 15.82221 | HRA_R_2 | hrar2_1 | Czech | R | 0 | 0 | 0 | 0 | 0 | 0 | 1 | 228,986 | 1443,370815 | 1 | 15258,162 | 0 | 0 | 1 | 0 | 1 | 0 | 0 | 1 | 0 | 0 | 1 | 1 | 0 | 0 | 0 | 0 | 0 |
| 50.1937 | 15.81731 | HRA_R_2 | hrar2_2 | Czech | R | 0 | 0 | 0 | 0 | 0 | 0 | 0 | 228 | 1443,583788 | 1 | 13220,573 | 0 | 0 | 1 | 0 | 1 | 0 | 0 | 1 | 0 | 0 | 1 | 0 | 0 | 0 | 0 | 0 | 0 |
| 50.19207 | 15.8207 | HRA_R_2 | hrar2_3 | Czech | R | 1 | 1 | 0 | 0 | 0 | 0 | 0 | 226,452 | 1443,28164 | 1 | 31249,031 | 41 | 1 | 1 | 0 | 1 | 0 | 0 | 1 | 0 | 0 | 1 | 0 | 0 | 0 | 0 | 0 | 0 |
| 50.19027 | 15.82399 | HRA_R_2 | hrar2_4 | Czech | R | 0 | 0 | 0 | 0 | 0 | 0 | 1 | 227,354 | 1442,972702 | 1 | 9313,484 | 1174 | 1 | 1 | 0 | 1 | 0 | 0 | 1 | 0 | 1 | 1 | 0 | 1 | 0 | 0 | 0 | 0 |
| 50.18795 | 15.82384 | HRA_R_2 | hrar2_5 | Czech | R | 1 | 1 | 0 | 0 | 0 | 0 | 1 | 226,933 | 1442,817699 | 1 | 18151,584 | 1296 | 0 | 1 | 0 | 1 | 0 | 0 | 0 | 1 | 1 | 1 | 0 | 1 | 0 | 0 | 0 | 0 |
| 50.21147 | 15.83412 | HRA_U_1 | hrau1_1 | Czech | U | 2 | 1 | 0 | 0 | 0 | 0 | 1 | 232,827 | 1443,909863 | 1 | 21797,672 | 18196 | 0 | 1 | 0 | 0 | 1 | 0 | 0 | 1 | 0 | 0 | 0 | 0 | 0 | 1 | 0 | 0 |
| 50.21109 | 15.8393 | HRA_U_1 | hrau1_2 | Czech | U | 2 | 1 | 0 | 0 | 0 | 0 | 1 | 232,964 | 1443,598294 | 1 | 13212,762 | 13417 | 0 | 1 | 1 | 0 | 1 | 0 | 1 | 1 | 0 | 0 | 0 | 0 | 0 | 1 | 0 | 0 |
| 50.21298 | 15.84004 | HRA_U_1 | hrau1_3 | Czech | U | 1 | 1 | 0 | 0 | 0 | 0 | 1 | 233 | 1443,691401 | 1 | 19866,337 | 8101 | 0 | 1 | 1 | 0 | 1 | 0 | 1 | 1 | 0 | 0 | 0 | 0 | 1 | 0 | 0 | 0 |
| 50.2151 | 15.83825 | HRA_U_1 | hrau1_4 | Czech | U | 2 | 1 | 1 | 1 | 0 | 0 | 1 | 230,448 | 1443,93975 | 1 | 18010,219 | 1177 | 0 | 1 | 0 | 0 | 1 | 0 | 0 | 0 | 0 | 1 | 0 | 0 | 0 | 0 | 0 | 1 |
| 50.21608 | 15.83467 | HRA_U_1 | hrau1_5 | Czech | U | 1 | 1 | 0 | 0 | 0 | 0 | 1 | 231,797 | 1444,206288 | 1 | 25454,463 | 0 | 0 | 1 | 0 | 0 | 1 | 0 | 0 | 0 | 0 | 0 | 0 | 0 | 0 | 0 | 1 | 0 |
| 50.19982 | 15.8333 | HRA_U_2 | hrau2_1 | Czech | U | 0 | 0 | 0 | 0 | 0 | 0 | 0 | 229,69 | 1443,131435 | 1 | 21970,721 | 7900 | 1 | 1 | 0 | 1 | 0 | 0 | 0 | 1 | 0 | 0 | 0 | 0 | 1 | 0 | 0 | 0 |
| 50.19755 | 15.83454 | HRA_U_2 | hrau2_2 | Czech | U | 0 | 0 | 0 | 0 | 0 | 0 | 0 | 229,938 | 1442,903468 | 1 | 13711,584 | 11891 | 1 | 1 | 0 | 1 | 0 | 0 | 0 | 1 | 1 | 0 | 0 | 1 | 1 | 0 | 0 | 0 |
| 50.19497 | 15.83484 | HRA_U_2 | hrau2_3 | Czech | U | 1 | 1 | 0 | 0 | 0 | 0 | 0 | 228 | 1442,705593 | 1 | 23934,196 | 3599 | 1 | 1 | 0 | 1 | 1 | 0 | 0 | 0 | 1 | 1 | 1 | 1 | 0 | 0 | 0 | 0 |
| 50.19384 | 15.83134 | HRA_U_2 | hrau2_4 | Czech | U | 0 | 0 | 0 | 0 | 0 | 0 | 0 | 227,176 | 1442,818847 | 1 | 21088,657 | 2109 | 1 | 1 | 0 | 0 | 1 | 0 | 0 | 1 | 1 | 1 | 0 | 1 | 0 | 0 | 0 | 0 |
| 50.19169 | 15.82989 | HRA_U_2 | hrau2_5 | Czech | U | 1 | 1 | 0 | 0 | 0 | 0 | 0 | 229,241 | 1442,74723 | 1 | 20066,287 | 3299 | 1 | 1 | 0 | 1 | 0 | 0 | 1 | 0 | 1 | 0 | 0 | 1 | 0 | 0 | 0 | 0 |
| 50.05839 | 22.68272 | JAR_R_1 | jarr1_1 | Poland | R | 4 | 1 | 4 | 1 | 0 | 0 | 1 | 180 | 1114,017331 | 288 | 20665,918 | 4939 | 1 | 1 | 0 | 1 | 0 | 0 | 0 | 0 | 1 | 0 | 1 | 0 | 0 | 0 | 0 | 0 |
| 50.05663 | 22.67955 | JAR_R_1 | jarr1_2 | Poland | R | 2 | 1 | 0 | 0 | 0 | 0 | 0 | 178,224 | 1113,95634 | 288 | 14909,528 | 1703 | 0 | 1 | 0 | 1 | 0 | 0 | 1 | 0 | 1 | 1 | 1 | 1 | 0 | 0 | 0 | 0 |
| 50.05451 | 22.6771 | JAR_R_1 | jarr1_3 | Poland | R | 0 | 0 | 2 | 1 | 0 | 0 | 1 | 179,204 | 1113,83653 | 288 | 15599,441 | 1176 | 0 | 1 | 0 | 0 | 0 | 0 | 1 | 0 | 1 | 1 | 0 | 1 | 0 | 0 | 0 | 0 |
| 50.05204 | 22.67587 | JAR_R_1 | jarr1_4 | Poland | R | 0 | 0 | 2 | 1 | 0 | 0 | 0 | 179,205 | 1113,637903 | 288 | 25661,942 | 520 | 0 | 1 | 0 | 0 | 0 | 0 | 1 | 0 | 1 | 1 | 0 | 1 | 0 | 0 | 0 | 0 |
| 50.04993 | 22.67834 | JAR_R_1 | jarr1_5 | Poland | R | 0 | 0 | 2 | 1 | 0 | 0 | 0 | 177,234 | 1113,348048 | 288 | 16413,044 | 1719 | 0 | 1 | 0 | 0 | 0 | 0 | 1 | 0 | 1 | 1 | 1 | 1 | 0 | 0 | 0 | 0 |
| 50.0451 | 22.64292 | JAR_R_2 | jarr2_1 | Poland | R | 2 | 1 | 0 | 0 | 0 | 0 | 0 | 198,976 | 1114,107421 | 288 | 24452,704 | 749 | 1 | 1 | 0 | 1 | 0 | 1 | 1 | 1 | 0 | 0 | 1 | 0 | 0 | 0 | 0 | 0 |
| 50.04706 | 22.6439 | JAR_R_2 | jarr2_2 | Poland | R | 0 | 0 | 1 | 1 | 0 | 0 | 0 | 209,584 | 1114,273597 | 288 | 11200,122 | 0 | 0 | 1 | 0 | 0 | 0 | 0 | 1 | 0 | 0 | 0 | 0 | 0 | 0 | 1 | 0 | 0 |
| 50.04947 | 22.64507 | JAR_R_2 | jarr2_3 | Poland | R | 0 | 0 | 0 | 0 | 0 | 0 | 0 | 209,748 | 1114,466451 | 288 | 35879,018 | 0 | 1 | 1 | 0 | 1 | 0 | 0 | 1 | 0 | 0 | 0 | 0 | 0 | 0 | 0 | 0 | 0 |
| 50.05203 | 22.64748 | JAR_R_2 | jarr2_4 | Poland | R | 0 | 0 | 2 | 1 | 0 | 0 | 0 | 193,528 | 1114,630307 | 288 | 39849,836 | 3945 | 1 | 0 | 0 | 1 | 0 | 0 | 1 | 0 | 0 | 0 | 1 | 1 | 0 | 0 | 0 | 0 |
| 50.0513 | 22.65229 | JAR_R_2 | jarr2_5 | Poland | R | 0 | 0 | 0 | 0 | 0 | 0 | 1 | 180,32 | 1114,390794 | 288 | 11595,992 | 9000 | 1 | 1 | 1 | 1 | 0 | 0 | 1 | 0 | 0 | 1 | 0 | 1 | 0 | 0 | 1 | 0 |
| 50.02189 | 22.66826 | JAR_U_1 | jaru1_1 | Poland | U | 2 | 1 | 0 | 0 | 0 | 0 | 0 | 208,144 | 1110,976318 | 290 | 56852,049 | 2957 | 1 | 0 | 0 | 0 | 1 | 0 | 0 | 0 | 0 | 0 | 0 | 0 | 1 | 0 | 0 | 0 |
| 50.02329 | 22.67244 | JAR_U_1 | jaru1_2 | Poland | U | 2 | 1 | 0 | 0 | 0 | 0 | 0 | 206,06 | 1110,968629 | 290 | 43774,589 | 4120 | 1 | 0 | 0 | 0 | 1 | 0 | 0 | 0 | 0 | 0 | 0 | 1 | 0 | 0 | 0 | 0 |
| 50.02311 | 22.676 | JAR_U_1 | jaru1_3 | Poland | U | 0 | 0 | 0 | 0 | 0 | 0 | 0 | 200,274 | 1110,825431 | 290 | 14427,405 | 9400 | 0 | 1 | 1 | 0 | 0 | 0 | 0 | 1 | 1 | 0 | 0 | 1 | 0 | 1 | 0 | 0 |
| 50.02197 | 22.67964 | JAR_U_1 | jaru1_4 | Poland | U | 0 | 0 | 0 | 0 | 0 | 0 | 0 | 204 | 1110,587922 | 290 | 18543,516 | 11268 | 0 | 1 | 1 | 0 | 0 | 0 | 0 | 1 | 0 | 0 | 0 | 1 | 0 | 1 | 0 | 0 |
| 50.02117 | 22.68349 | JAR_U_1 | jaru1_5 | Poland | U | 0 | 0 | 1 | 1 | 0 | 0 | 0 | 183,92 | 1110,374942 | 290 | 13353,227 | 10303 | 0 | 1 | 0 | 0 | 0 | 0 | 0 | 1 | 1 | 0 | 0 | 1 | 0 | 1 | 0 | 0 |
| 50.0141 | 22.671 | JAR_U_2 | jaru2_1 | Poland | U | 1 | 1 | 0 | 0 | 0 | 0 | 0 | 217,4 | 1110,125794 | 292 | 20224,748 | 12217 | 1 | 1 | 1 | 1 | 0 | 0 | 0 | 1 | 0 | 0 | 1 | 0 | 0 | 0 | 1 | 0 |
| 50.01369 | 22.66634 | JAR_U_2 | jaru2_2 | Poland | U | 0 | 0 | 0 | 0 | 0 | 0 | 0 | 205,62 | 1110,249348 | 292 | 16041,898 | 7088 | 1 | 1 | 1 | 1 | 0 | 0 | 0 | 1 | 1 | 0 | 0 | 1 | 0 | 0 | 0 | 0 |
| 50.0118 | 22.66259 | JAR_U_2 | jaru2_3 | Poland | U | 1 | 1 | 0 | 0 | 0 | 0 | 0 | 198 | 1110,197206 | 292 | 11320,059 | 3876 | 0 | 1 | 1 | 0 | 0 | 0 | 0 | 1 | 1 | 0 | 0 | 1 | 0 | 0 | 0 | 0 |
| 50.00959 | 22.66029 | JAR_U_2 | jaru2_4 | Poland | U | 0 | 0 | 0 | 0 | 0 | 0 | 0 | 211,973 | 1110,06396 | 292 | 23738,034 | 3943 | 1 | 1 | 0 | 0 | 0 | 0 | 0 | 1 | 1 | 0 | 0 | 1 | 0 | 0 | 1 | 0 |
| 50.00756 | 22.65849 | JAR_U_2 | jaru2_5 | Poland | U | 1 | 1 | 2 | 1 | 0 | 0 | 0 | 216,432 | 1109,930132 | 292 | 35736,26 | 1242 | 1 | 1 | 0 | 0 | 0 | 1 | 0 | 1 | 0 | 0 | 0 | 1 | 1 | 1 | 0 | 1 |
| 49.63881 | 17.95359 | JIC_R_1 | jicr1_1 | Czech | R | 2 | 1 | 0 | 0 | 0 | 0 | 0 | 250,378 | 1287,560097 | 132 | 58335,049 | 1000 | 1 | 1 | 0 | 1 | 0 | 0 | 0 | 0 | 0 | 1 | 0 | 0 | 0 | 0 | 1 | 1 |
| 49.63634 | 17.95159 | JIC_R_1 | jicr1_2 | Czech | R | 2 | 1 | 0 | 0 | 0 | 0 | 0 | 250,773 | 1287,481445 | 132 | 42348,138 | 0 | 1 | 1 | 0 | 1 | 0 | 0 | 0 | 0 | 0 | 1 | 0 | 0 | 0 | 0 | 0 | 0 |
| 49.63388 | 17.94979 | JIC_R_1 | jicr1_3 | Czech | R | 0 | 0 | 0 | 0 | 0 | 0 | 0 | 251,732 | 1287,386271 | 132 | 7818,66 | 0 | 1 | 1 | 0 | 1 | 0 | 0 | 1 | 0 | 0 | 1 | 0 | 0 | 0 | 0 | 0 | 1 |
| 49.63142 | 17.94666 | JIC_R_1 | jicr1_4 | Czech | R | 2 | 1 | 1 | 1 | 0 | 0 | 0 | 251,133 | 1287,363565 | 132 | 12440,615 | 0 | 0 | 1 | 0 | 1 | 0 | 0 | 1 | 0 | 0 | 1 | 0 | 0 | 0 | 0 | 0 | 0 |
| 49.62838 | 17.94447 | JIC_R_1 | jicr1_5 | Czech | R | 1 | 1 | 1 | 1 | 1 | 1 | 0 | 252,985 | 1287,247564 | 132 | 24676,066 | 1234 | 1 | 1 | 0 | 1 | 0 | 0 | 1 | 0 | 0 | 1 | 1 | 0 | 0 | 0 | 0 | 0 |
| 49.6214 | 17.94318 | JIC_R_2 | jicr2_1 | Czech | R | 1 | 1 | 0 | 0 | 0 | 0 | 1 | 255 | 1286,785108 | 133 | 25921,215 | 2064 | 1 | 1 | 0 | 1 | 0 | 0 | 1 | 0 | 1 | 1 | 1 | 0 | 1 | 0 | 0 | 0 |
| 49.61943 | 17.9452 | JIC_R_2 | jicr2_2 | Czech | R | 0 | 0 | 0 | 0 | 0 | 0 | 1 | 257,72 | 1286,494242 | 133 | 13738,25 | 4939 | 0 | 1 | 0 | 0 | 0 | 0 | 0 | 1 | 1 | 1 | 1 | 1 | 0 | 0 | 0 | 0 |
| 49.61684 | 17.94571 | JIC_R_2 | jicr2_3 | Czech | R | 1 | 1 | 0 | 0 | 0 | 0 | 1 | 259,444 | 1286,266569 | 133 | 16973,623 | 3795 | 1 | 1 | 0 | 1 | 0 | 0 | 0 | 1 | 1 | 1 | 1 | 1 | 0 | 0 | 0 | 0 |
| 49.61406 | 17.94386 | JIC_R_2 | jicr2_4 | Czech | R | 2 | 1 | 1 | 1 | 0 | 0 | 0 | 261,064 | 1286,194374 | 133 | 15781,556 | 500 | 1 | 0 | 0 | 1 | 0 | 0 | 1 | 0 | 1 | 1 | 1 | 0 | 0 | 0 | 0 | 1 |
| 49.6123 | 17.93981 | JIC_R_2 | jicr2_5 | Czech | R | 2 | 1 | 0 | 0 | 0 | 0 | 0 | 265,244 | 1286,258976 | 133 | 12012,368 | 0 | 1 | 1 | 0 | 1 | 0 | 0 | 1 | 0 | 0 | 1 | 0 | 0 | 0 | 0 | 0 | 0 |
| 49.59124 | 18.012 | JIC_U_1 | jicu1_1 | Czech | U | 2 | 1 | 0 | 0 | 0 | 0 | 1 | 299,165 | 1280,864982 | 142 | 15181,158 | 14766 | 0 | 1 | 0 | 0 | 1 | 0 | 0 | 1 | 1 | 0 | 0 | 1 | 1 | 0 | 0 | 1 |
| 49.58901 | 18.00959 | JIC_U_1 | jicu1_2 | Czech | U | 2 | 1 | 0 | 0 | 0 | 0 | 1 | 313,564 | 1280,82016 | 142 | 26799,798 | 8574 | 0 | 1 | 0 | 0 | 1 | 0 | 0 | 0 | 1 | 0 | 0 | 1 | 1 | 0 | 0 | 0 |
| 49.58764 | 18.01326 | JIC_U_1 | jicu1_3 | Czech | U | 2 | 1 | 0 | 0 | 0 | 0 | 1 | 324,357 | 1280,52159 | 142 | 14541,807 | 10126 | 1 | 0 | 0 | 1 | 0 | 0 | 0 | 0 | 1 | 0 | 0 | 1 | 0 | 0 | 0 | 0 |
| 49.58591 | 18.0156 | JIC_U_1 | jicu1_4 | Czech | U | 2 | 1 | 0 | 0 | 0 | 0 | 0 | 352,184 | 1280,262665 | 142 | 41203,141 | 846 | 1 | 1 | 0 | 1 | 1 | 0 | 0 | 0 | 0 | 0 | 0 | 0 | 0 | 0 | 0 | 0 |
| 49.58642 | 18.01843 | JIC_U_1 | jicu1_5 | Czech | U | 2 | 1 | 0 | 0 | 0 | 0 | 1 | 321,703 | 1280,15599 | 142 | 34394,775 | 1311 | 1 | 1 | 0 | 1 | 1 | 0 | 0 | 0 | 1 | 0 | 1 | 0 | 0 | 0 | 0 | 0 |
| 49.58127 | 17.9667 | JIC_U_2 | jicu2_1 | Czech | U | 1 | 1 | 0 | 0 | 0 | 0 | 0 | 372,664 | 1282,492901 | 138 | 26970,137 | 306 | 1 | 1 | 0 | 1 | 0 | 0 | 1 | 0 | 1 | 0 | 1 | 0 | 0 | 0 | 0 | 1 |
| 49.57802 | 17.96529 | JIC_U_2 | jicu2_2 | Czech | U | 0 | 0 | 0 | 0 | 0 | 0 | 0 | 359,822 | 1282,351919 | 138 | 35211,933 | 3674 | 1 | 1 | 0 | 1 | 0 | 0 | 1 | 0 | 1 | 0 | 1 | 1 | 0 | 0 | 0 | 0 |
| 49.57789 | 17.96152 | JIC_U_2 | jicu2_3 | Czech | U | 0 | 0 | 0 | 0 | 0 | 0 | 0 | 388,661 | 1282,508036 | 138 | 24497,039 | 10002 | 1 | 0 | 0 | 1 | 0 | 1 | 0 | 0 | 1 | 0 | 1 | 1 | 0 | 0 | 0 | 0 |
| 49.57993 | 17.96003 | JIC_U_2 | jicu2_4 | Czech | U | 0 | 0 | 1 | 1 | 0 | 0 | 1 | 382,676 | 1282,741426 | 138 | 28798,702 | 3065 | 1 | 1 | 0 | 1 | 0 | 1 | 0 | 0 | 1 | 0 | 1 | 0 | 0 | 0 | 0 | 0 |
| 49.58237 | 17.96024 | JIC_U_2 | jicu2_5 | Czech | U | 1 | 1 | 0 | 0 | 0 | 0 | 0 | 399,06 | 1282,917608 | 138 | 35896,267 | 0 | 1 | 1 | 0 | 1 | 0 | 0 | 0 | 0 | 1 | 0 | 0 | 0 | 0 | 0 | 0 | 0 |
| 50.25259 | 18.74334 | KAT_R_1 | katr1_1 | Poland | R | 0 | 0 | 0 | 0 | 0 | 0 | 1 | 208,984 | 1296,504867 | 116 | 15910,734 | 1002 | 0 | 1 | 0 | 1 | 0 | 0 | 0 | 0 | 1 | 0 | 1 | 0 | 0 | 0 | 0 | 0 |
| 50.25537 | 18.74369 | KAT_R_1 | katr1_2 | Poland | R | 0 | 0 | 0 | 0 | 0 | 0 | 0 | 208,238 | 1296,715193 | 116 | 21397,213 | 0 | 1 | 1 | 0 | 1 | 0 | 0 | 0 | 0 | 0 | 0 | 0 | 0 | 0 | 0 | 0 | 0 |
| 50.25816 | 18.74403 | KAT_R_1 | katr1_3 | Poland | R | 0 | 0 | 0 | 0 | 0 | 0 | 0 | 209,366 | 1296,928629 | 116 | 15004,558 | 0 | 1 | 1 | 0 | 1 | 0 | 0 | 0 | 0 | 0 | 0 | 0 | 0 | 0 | 0 | 0 | 0 |
| 50.25823 | 18.74712 | KAT_R_1 | katr1_4 | Poland | R | 0 | 0 | 0 | 0 | 0 | 0 | 0 | 210,231 | 1296,787478 | 116 | 22135,006 | 0 | 1 | 1 | 0 | 1 | 0 | 0 | 0 | 0 | 0 | 1 | 0 | 0 | 0 | 0 | 0 | 0 |
| 50.25664 | 18.75064 | KAT_R_1 | katr1_5 | Poland | R | 0 | 0 | 0 | 0 | 0 | 0 | 1 | 211,304 | 1296,488234 | 116 | 4832,168 | 0 | 0 | 1 | 0 | 1 | 0 | 0 | 1 | 0 | 0 | 1 | 0 | 0 | 0 | 0 | 1 | 1 |
| 50.24499 | 18.76035 | KAT_R_2 | katr2_1 | Poland | R | 2 | 1 | 0 | 0 | 0 | 0 | 1 | 228 | 1295,063175 | 117 | 44081,558 | 2272 | 1 | 0 | 1 | 1 | 0 | 0 | 0 | 0 | 0 | 0 | 1 | 0 | 0 | 0 | 1 | 0 |
| 50.24818 | 18.75744 | KAT_R_2 | katr2_2 | Poland | R | 2 | 1 | 0 | 0 | 0 | 0 | 1 | 225,413 | 1295,465677 | 117 | 55829,411 | 0 | 1 | 1 | 1 | 1 | 0 | 0 | 0 | 0 | 0 | 0 | 0 | 0 | 0 | 0 | 0 | 0 |
| 50.25054 | 18.75643 | KAT_R_2 | katr2_3 | Poland | R | 1 | 1 | 0 | 0 | 0 | 0 | 1 | 221,365 | 1295,707989 | 117 | 24187,035 | 409 | 1 | 1 | 1 | 1 | 0 | 0 | 0 | 0 | 1 | 1 | 1 | 0 | 0 | 0 | 0 | 0 |
| 50.24885 | 18.75322 | KAT_R_2 | katr2_4 | Poland | R | 1 | 1 | 0 | 0 | 0 | 0 | 1 | 219,886 | 1295,721992 | 117 | 26874,522 | 0 | 1 | 1 | 1 | 1 | 0 | 0 | 0 | 0 | 0 | 1 | 0 | 0 | 0 | 0 | 0 | 0 |
| 50.24845 | 18.7499 | KAT_R_2 | katr2_5 | Poland | R | 2 | 1 | 0 | 0 | 0 | 0 | 0 | 219,534 | 1295,848442 | 117 | 21977,803 | 0 | 1 | 1 | 0 | 1 | 0 | 1 | 1 | 1 | 0 | 1 | 0 | 0 | 0 | 0 | 0 | 0 |
| 50.24652 | 19.00226 | KAT_U_2 | katu2_1 | Poland | U | 0 | 0 | 0 | 0 | 0 | 0 | 1 | 295,12 | 1283,67074 | 132 | 63904,884 | 492 | 1 | 0 | 0 | 0 | 1 | 0 | 0 | 0 | 0 | 0 | 0 | 0 | 0 | 0 | 0 | 1 |
| 50.24442 | 18.99959 | KAT_U_2 | katu2_2 | Poland | U | 1 | 1 | 0 | 0 | 0 | 0 | 1 | 298,088 | 1283,622861 | 132 | 61448,001 | 1058 | 1 | 0 | 0 | 0 | 1 | 0 | 0 | 0 | 1 | 0 | 0 | 1 | 0 | 0 | 0 | 1 |
| 50.24301 | 18.99979 | KAT_U_2 | katu2_3 | Poland | U | 1 | 1 | 0 | 0 | 0 | 0 | 1 | 303,164 | 1283,494453 | 132 | 50712,499 | 2186 | 1 | 1 | 1 | 0 | 1 | 0 | 0 | 1 | 1 | 0 | 0 | 1 | 1 | 0 | 0 | 0 |
| 50.2404 | 19.00067 | KAT_U_2 | katu2_4 | Poland | U | 3 | 1 | 0 | 0 | 0 | 0 | 0 | 304,852 | 1283,236214 | 132 | 52548,845 | 0 | 1 | 1 | 1 | 0 | 1 | 0 | 0 | 0 | 1 | 0 | 0 | 1 | 0 | 0 | 0 | 1 |
| 50.24131 | 19.00419 | KAT_U_2 | katu2_5 | Poland | U | 4 | 1 | 0 | 0 | 0 | 0 | 0 | 317,344 | 1283,145402 | 132 | 55260,195 | 993 | 1 | 1 | 1 | 0 | 1 | 1 | 0 | 1 | 0 | 0 | 0 | 0 | 1 | 0 | 0 | 0 |
| 46.91175 | 19.63966 | KEC_R_1 | kecr1_1 | Hungary | R | 1 | 1 | 2 | 1 | 0 | 0 | 0 | 115,224 | 995,8143564 | 253 | 33190,607 | 0 | 1 | 1 | 0 | 1 | 0 | 0 | 1 | 0 | 0 | 0 | 0 | 0 | 0 | 0 | 0 | 0 |
| 46.91385 | 19.64185 | KEC_R_1 | kecr1_2 | Hungary | R | 0 | 0 | 0 | 0 | 0 | 0 | 0 | 118,14 | 995,8225775 | 253 | 35944,264 | 1488 | 1 | 1 | 0 | 1 | 0 | 0 | 1 | 0 | 0 | 0 | 1 | 0 | 0 | 0 | 0 | 0 |
| 46.91333 | 19.64534 | KEC_R_1 | kecr1_3 | Hungary | R | 1 | 1 | 0 | 0 | 0 | 0 | 0 | 114,988 | 995,5755169 | 253 | 26048,254 | 524 | 1 | 0 | 0 | 1 | 0 | 0 | 0 | 0 | 0 | 0 | 1 | 0 | 0 | 0 | 0 | 0 |
| 46.91174 | 19.64876 | KEC_R_1 | kecr1_4 | Hungary | R | 0 | 0 | 0 | 0 | 0 | 0 | 0 | 115,272 | 995,2610098 | 253 | 30079,644 | 613 | 1 | 1 | 0 | 1 | 1 | 0 | 1 | 0 | 0 | 0 | 1 | 0 | 0 | 0 | 0 | 0 |
| 46.9098 | 19.65215 | KEC_R_1 | kecr1_5 | Hungary | R | 1 | 1 | 0 | 0 | 1 | 1 | 0 | 116,373 | 994,9248005 | 253 | 30499,881 | 0 | 1 | 1 | 0 | 1 | 0 | 0 | 1 | 1 | 0 | 1 | 0 | 0 | 0 | 0 | 0 | 1 |
| 46.95081 | 19.59254 | KEC_R_2 | kecr2_1 | Hungary | R | 0 | 0 | 1 | 1 | 0 | 0 | 0 | 121,928 | 1001,293507 | 251 | 36797,302 | 0 | 1 | 1 | 0 | 1 | 0 | 0 | 1 | 0 | 1 | 1 | 0 | 0 | 0 | 0 | 0 | 0 |
| 46.95363 | 19.59475 | KEC_R_2 | kecr2_2 | Hungary | R | 0 | 0 | 2 | 1 | 0 | 0 | 0 | 122,932 | 1001,35161 | 251 | 26109,36 | 824 | 1 | 1 | 0 | 1 | 0 | 0 | 1 | 0 | 1 | 0 | 1 | 0 | 0 | 0 | 0 | 0 |
| 46.95526 | 19.59766 | KEC_R_2 | kecr2_3 | Hungary | R | 1 | 1 | 0 | 0 | 0 | 0 | 0 | 121,488 | 1001,283981 | 251 | 43991,983 | 422 | 1 | 1 | 0 | 1 | 0 | 0 | 1 | 0 | 0 | 0 | 1 | 0 | 0 | 0 | 0 | 0 |
| 46.95674 | 19.60138 | KEC_R_2 | kecr2_4 | Hungary | R | 0 | 0 | 0 | 0 | 0 | 0 | 0 | 121,712 | 1001,158066 | 251 | 24967,247 | 1540 | 1 | 1 | 0 | 1 | 0 | 0 | 1 | 0 | 0 | 0 | 0 | 0 | 0 | 0 | 1 | 0 |
| 46.95702 | 19.60539 | KEC_R_2 | kecr2_5 | Hungary | R | 1 | 1 | 0 | 0 | 0 | 0 | 0 | 123,132 | 1000,931094 | 251 | 18789,601 | 620 | 1 | 1 | 0 | 1 | 0 | 0 | 1 | 0 | 0 | 0 | 1 | 0 | 0 | 0 | 0 | 0 |
| 46.90267 | 19.68131 | KEC_U_1 | kecu1_1 | Hungary | U | 0 | 0 | 0 | 0 | 0 | 0 | 0 | 116,498 | 992,6759539 | 255 | 14754,91 | 21493 | 0 | 1 | 0 | 0 | 1 | 0 | 0 | 1 | 0 | 0 | 0 | 1 | 0 | 1 | 0 | 0 |
| 46.90056 | 19.68373 | KEC_U_1 | kecu1_2 | Hungary | U | 2 | 1 | 0 | 0 | 0 | 0 | 0 | 119,01 | 992,3866574 | 255 | 37434,243 | 9975 | 1 | 1 | 0 | 1 | 1 | 0 | 0 | 1 | 0 | 0 | 0 | 1 | 1 | 0 | 0 | 0 |
| 46.89824 | 19.68153 | KEC_U_1 | kecu1_3 | Hungary | U | 0 | 0 | 1 | 1 | 0 | 0 | 0 | 115,991 | 992,3638529 | 255 | 38907,364 | 7440 | 1 | 1 | 0 | 1 | 0 | 0 | 0 | 1 | 1 | 0 | 0 | 1 | 1 | 0 | 0 | 1 |
| 46.89604 | 19.68372 | KEC_U_1 | kecu1_4 | Hungary | U | 1 | 1 | 0 | 0 | 0 | 0 | 0 | 117,916 | 992,0836181 | 255 | 29538,738 | 10344 | 1 | 1 | 0 | 1 | 0 | 0 | 0 | 1 | 0 | 0 | 0 | 1 | 0 | 0 | 1 | 0 |
| 46.89642 | 19.68634 | KEC_U_1 | kecu1_5 | Hungary | U | 0 | 0 | 0 | 0 | 0 | 0 | 0 | 114,224 | 991,9493796 | 255 | 34243,846 | 8177 | 1 | 1 | 0 | 1 | 0 | 0 | 0 | 1 | 0 | 0 | 0 | 0 | 1 | 0 | 0 | 0 |
| 46.91914 | 19.68437 | KEC_U_2 | kecu2_1 | Hungary | U | 0 | 0 | 1 | 1 | 0 | 0 | 0 | 118 | 993,5938795 | 256 | 32967,934 | 6592 | 1 | 1 | 1 | 1 | 1 | 1 | 1 | 0 | 0 | 0 | 0 | 1 | 0 | 0 | 0 | 0 |
| 46.91637 | 19.68671 | KEC_U_2 | kecu2_2 | Hungary | U | 1 | 1 | 0 | 0 | 0 | 0 | 0 | 114,989 | 993,2653311 | 256 | 28585,673 | 13012 | 1 | 1 | 0 | 0 | 1 | 0 | 1 | 1 | 0 | 0 | 0 | 1 | 0 | 1 | 0 | 0 |
| 46.91588 | 19.68283 | KEC_U_2 | kecu2_3 | Hungary | U | 0 | 0 | 0 | 0 | 0 | 0 | 0 | 118,644 | 993,4693725 | 256 | 58484,966 | 2409 | 1 | 1 | 0 | 0 | 1 | 1 | 0 | 1 | 0 | 0 | 1 | 0 | 0 | 0 | 0 | 0 |
| 46.91628 | 19.67961 | KEC_U_2 | kecu2_4 | Hungary | U | 2 | 1 | 0 | 0 | 0 | 0 | 1 | 119 | 993,6904617 | 256 | 31155,858 | 14627 | 1 | 1 | 0 | 0 | 0 | 1 | 1 | 1 | 0 | 0 | 0 | 0 | 1 | 0 | 1 | 0 |
| 46.91473 | 19.67654 | KEC_U_2 | kecu2_5 | Hungary | U | 0 | 0 | 1 | 1 | 0 | 0 | 1 | 116,443 | 993,7729917 | 256 | 30570,562 | 7417 | 1 | 1 | 0 | 0 | 1 | 0 | 1 | 1 | 0 | 0 | 0 | 0 | 1 | 0 | 0 | 0 |
| 50.92822 | 20.75315 | KIE_R_1 | kier1_1 | Poland | R | 0 | 0 | 0 | 0 | 0 | 0 | 1 | 305,677 | 1268,013669 | 177 | 6452,879 | 4800 | 0 | 1 | 0 | 0 | 0 | 0 | 0 | 1 | 1 | 0 | 0 | 1 | 0 | 0 | 0 | 0 |
| 50.92749 | 20.75758 | KIE_R_1 | kier1_2 | Poland | R | 1 | 1 | 0 | 0 | 0 | 0 | 0 | 300,722 | 1267,772535 | 177 | 5970,56 | 4746 | 0 | 1 | 0 | 0 | 0 | 0 | 0 | 1 | 1 | 1 | 0 | 1 | 0 | 0 | 0 | 0 |
| 50.92593 | 20.76071 | KIE_R_1 | kier1_3 | Poland | R | 1 | 1 | 0 | 0 | 0 | 0 | 1 | 300,748 | 1267,50554 | 177 | 3990,794 | 5680 | 0 | 1 | 0 | 0 | 0 | 0 | 1 | 1 | 0 | 0 | 0 | 1 | 0 | 0 | 0 | 0 |
| 50.92482 | 20.76444 | KIE_R_1 | kier1_4 | Poland | R | 0 | 0 | 0 | 0 | 0 | 0 | 1 | 298,658 | 1267,257063 | 177 | 12108,657 | 4275 | 1 | 1 | 0 | 1 | 0 | 0 | 0 | 1 | 0 | 1 | 0 | 1 | 0 | 0 | 0 | 1 |
| 50.92321 | 20.76784 | KIE_R_1 | kier1_5 | Poland | R | 0 | 0 | 0 | 0 | 0 | 0 | 1 | 301,789 | 1266,976435 | 177 | 12055,063 | 6762 | 1 | 1 | 0 | 1 | 0 | 0 | 1 | 1 | 1 | 0 | 0 | 1 | 1 | 0 | 0 | 0 |
| 50.91481 | 20.70931 | KIE_R_2 | kier2_1 | Poland | R | 1 | 1 | 0 | 0 | 0 | 0 | 0 | 340,168 | 1268,489796 | 178 | 26978,244 | 0 | 1 | 1 | 0 | 1 | 0 | 0 | 0 | 0 | 0 | 1 | 0 | 0 | 0 | 0 | 0 | 0 |
| 50.91351 | 20.71368 | KIE_R_2 | kier2_2 | Poland | R | 0 | 0 | 0 | 0 | 0 | 0 | 0 | 331,366 | 1268,201395 | 178 | 11008,643 | 0 | 0 | 1 | 0 | 1 | 0 | 0 | 0 | 0 | 0 | 1 | 0 | 0 | 0 | 0 | 0 | 1 |
| 50.91263 | 20.71769 | KIE_R_2 | kier2_3 | Poland | R | 1 | 1 | 0 | 0 | 0 | 0 | 1 | 322,016 | 1267,960803 | 178 | 19739,366 | 0 | 1 | 1 | 0 | 1 | 0 | 0 | 1 | 0 | 0 | 1 | 0 | 0 | 0 | 0 | 0 | 0 |
| 50.91076 | 20.72278 | KIE_R_2 | kier2_4 | Poland | R | 0 | 0 | 0 | 0 | 0 | 0 | 1 | 312,736 | 1267,589032 | 178 | 7788,499 | 5150 | 0 | 1 | 1 | 1 | 0 | 0 | 1 | 0 | 0 | 0 | 0 | 1 | 0 | 0 | 0 | 0 |
| 50.90911 | 20.72454 | KIE_R_2 | kier2_5 | Poland | R | 0 | 0 | 0 | 0 | 0 | 0 | 1 | 308,656 | 1267,366092 | 178 | 15864,872 | 3797 | 1 | 1 | 0 | 1 | 0 | 0 | 1 | 0 | 0 | 0 | 1 | 1 | 0 | 0 | 0 | 0 |
| 50.88051 | 20.62989 | KIE_U_1 | kieu1_1 | Poland | U | 0 | 0 | 0 | 0 | 0 | 0 | 1 | 260 | 1268,464839 | 179 | 18635,224 | 6169 | 0 | 1 | 0 | 0 | 1 | 0 | 0 | 0 | 1 | 1 | 0 | 1 | 1 | 0 | 0 | 1 |
| 50.88443 | 20.63046 | KIE_U_1 | kieu1_2 | Poland | U | 1 | 1 | 0 | 0 | 0 | 0 | 0 | 261,634 | 1268,80327 | 179 | 21134,218 | 12269 | 0 | 1 | 0 | 0 | 1 | 0 | 0 | 0 | 0 | 1 | 0 | 1 | 1 | 1 | 0 | 0 |
| 50.88652 | 20.63256 | KIE_U_1 | kieu1_3 | Poland | U | 1 | 1 | 0 | 0 | 0 | 0 | 0 | 268,948 | 1268,913367 | 179 | 15202,514 | 6315 | 0 | 1 | 0 | 0 | 1 | 0 | 1 | 1 | 0 | 1 | 0 | 1 | 0 | 0 | 0 | 1 |

| | | | | | | | | | | | | | | | | | | | | | | | | | | | | | | | | | |
|---|---|---|---|---|---|---|---|---|---|---|---|---|---|---|---|---|---|---|---|---|---|---|---|---|---|---|---|---|---|---|---|---|---|
| 50.88875 | 20.63494 | KIE_U_1 | kieu1_4 | Poland | U | 0 | 0 | 0 | 0 | 0 | 0 | 1 | 274,068 | 1269,021412 | 179 | 28100,127 | 4294 | 1 | 1 | 0 | 0 | 1 | 0 | 0 | 1 | 0 | 0 | 1 | 0 | 1 | 0 | 0 | 0 |
| 50.89144 | 20.63479 | KIE_U_1 | kieu1_5 | Poland | U | 0 | 0 | 0 | 0 | 0 | 0 | 1 | 269,259 | 1269,274622 | 179 | 18444,152 | 2430 | 1 | 1 | 0 | 1 | 1 | 0 | 0 | 0 | 0 | 0 | 0 | 0 | 0 | 0 | 0 | 1 |
| 50.85697 | 20.62100 | KIE_U_2 | kieu2_1 | Poland | U | 1 | 1 | 0 | 0 | 0 | 0 | 0 | 262,637 | 1266,654339 | 184 | 22486,294 | 823 | 1 | 1 | 0 | 0 | 0 | 1 | 0 | 0 | 1 | 0 | 0 | 1 | 0 | 0 | 0 | 0 |
| 50.85956 | 20.62353 | KIE_U_2 | kieu2_2 | Poland | U | 1 | 1 | 0 | 0 | 0 | 0 | 1 | 267,416 | 1266,791664 | 184 | 20103,698 | 2467 | 0 | 1 | 0 | 0 | 0 | 1 | 0 | 0 | 1 | 0 | 0 | 1 | 1 | 0 | 0 | 0 |
| 50.85938 | 20.62750 | KIE_U_2 | kieu2_3 | Poland | U | 1 | 1 | 0 | 0 | 0 | 0 | 1 | 279,464 | 1266,617049 | 184 | 12540,333 | 13219 | 0 | 1 | 0 | 0 | 0 | 0 | 1 | 0 | 0 | 1 | 0 | 0 | 1 | 1 | 1 | 0 | 0 |
| 50.85701 | 20.62887 | KIE_U_2 | kieu2_4 | Poland | U | 0 | 0 | 0 | 0 | 0 | 0 | 0 | 289 | 1266,345262 | 184 | 14989,61 | 8086 | 1 | 1 | 0 | 0 | 0 | 1 | 1 | 1 | 1 | 0 | 0 | 1 | 1 | 1 | 0 | 1 |
| 50.85651 | 20.62627 | KIE_U_2 | kieu2_5 | Poland | U | 0 | 0 | 0 | 0 | 0 | 0 | 1 | 282,008 | 1266,402902 | 184 | 13444,359 | 2638 | 1 | 1 | 0 | 0 | 0 | 1 | 1 | 1 | 0 | 0 | 0 | 1 | 0 | 0 | 0 | 1 |
| 41.1578 | 25.42333 | KOM_R_1 | komr1_1 | Grece | R | 0 | 0 | 0 | 0 | 0 | 0 | 0 | 163,387 | 309,7547283 | 366 | 64257,179 | 0 | 1 | 0 | 1 | 1 | 0 | 0 | 1 | 0 | 0 | 0 | 0 | 0 | 0 | 0 | 0 | 0 |
| 41.1554 | 25.42308 | KOM_R_1 | komr1_2 | Grece | R | 0 | 0 | 0 | 0 | 0 | 0 | 0 | 159,244 | 309,7744352 | 366 | 46077,343 | 0 | 1 | 1 | 1 | 1 | 0 | 0 | 0 | 0 | 1 | 0 | 0 | 0 | 0 | 0 | 0 | 0 |
| 41.1531 | 25.42294 | KOM_R_1 | komr1_3 | Grece | R | 0 | 0 | 0 | 0 | 0 | 0 | 0 | 165,155 | 309,7845784 | 366 | 38124,653 | 0 | 1 | 1 | 1 | 1 | 0 | 0 | 1 | 0 | 0 | 0 | 1 | 0 | 0 | 0 | 0 | 0 |
| 41.15134 | 25.42105 | KOM_R_1 | komr1_4 | Grece | R | 0 | 0 | 0 | 0 | 0 | 0 | 0 | 150,78 | 309,9462678 | 366 | 18467,285 | 0 | 1 | 1 | 1 | 1 | 0 | 0 | 1 | 0 | 0 | 0 | 0 | 0 | 0 | 0 | 0 | 0 |
| 41.14875 | 25.41981 | KOM_R_1 | komr1_5 | Grece | R | 0 | 0 | 2 | 1 | 0 | 0 | 0 | 128,684 | 310,049749 | 366 | 8862,612 | 0 | 0 | 1 | 0 | 0 | 0 | 0 | 1 | 0 | 0 | 0 | 0 | 0 | 0 | 0 | 0 | 0 |
| 41.17426 | 25.40751 | KOM_R_2 | komr2_1 | Grece | R | 0 | 0 | 2 | 1 | 0 | 0 | 0 | 91,748 | 311,0836666 | 366 | 28997,705 | 3314 | 1 | 1 | 1 | 1 | 0 | 0 | 1 | 1 | 1 | 1 | 1 | 1 | 0 | 0 | 0 | 0 |
| 41.17168 | 25.40783 | KOM_R_2 | komr2_2 | Grece | R | 0 | 0 | 2 | 1 | 0 | 0 | 0 | 101,14 | 311,0577981 | 366 | 21970,758 | 1147 | 0 | 1 | 0 | 1 | 0 | 0 | 1 | 1 | 1 | 0 | 1 | 0 | 0 | 0 | 0 | 0 |
| 41.16979 | 25.40493 | KOM_R_2 | komr2_3 | Grece | R | 0 | 0 | 0 | 0 | 0 | 0 | 0 | 92,748 | 311,3011667 | 366 | 20246,965 | 1425 | 0 | 1 | 0 | 1 | 0 | 0 | 1 | 0 | 1 | 1 | 1 | 0 | 0 | 0 | 0 | 0 |
| 41.16738 | 25.40341 | KOM_R_2 | komr2_4 | Grece | R | 0 | 0 | 2 | 1 | 0 | 0 | 0 | 82,573 | 311,4272734 | 366 | 27738,994 | 1102 | 0 | 1 | 0 | 1 | 0 | 0 | 1 | 0 | 1 | 1 | 1 | 0 | 0 | 0 | 1 | 0 |
| 41.16598 | 25.40075 | KOM_R_2 | komr2_5 | Grece | R | 0 | 0 | 2 | 1 | 0 | 0 | 0 | 70,528 | 311,6515742 | 366 | 23641,03 | 1830 | 1 | 1 | 0 | 0 | 0 | 0 | 1 | 0 | 1 | 1 | 1 | 0 | 0 | 0 | 1 | 0 |
| 40.99033 | 25.3064 | KOM_R_3 | komr3_1 | Grece | R | 0 | 0 | 0 | 0 | 0 | 0 | 0 | 4,241 | 320,1376384 | 352 | 9946,517 | 0 | 0 | 1 | 0 | 1 | 0 | 0 | 0 | 0 | 1 | 1 | 0 | 0 | 0 | 0 | 0 | 0 |
| 40.99304 | 25.30666 | KOM_R_3 | komr3_2 | Grece | R | 0 | 0 | 1 | 1 | 0 | 0 | 0 | 10,072 | 320,0953167 | 352 | 5022,488 | 0 | 0 | 1 | 0 | 1 | 0 | 0 | 0 | 0 | 0 | 1 | 0 | 0 | 0 | 0 | 0 | 0 |
| 40.99583 | 25.30853 | KOM_R_3 | komr3_3 | Grece | R | 0 | 0 | 0 | 0 | 0 | 0 | 0 | 9,512 | 319,9208211 | 352 | 3996,177 | 0 | 0 | 1 | 0 | 1 | 0 | 0 | 0 | 0 | 0 | 1 | 0 | 0 | 0 | 0 | 0 | 0 |
| 40.99853 | 25.31018 | KOM_R_3 | komr3_4 | Grece | R | 0 | 0 | 1 | 1 | 0 | 0 | 0 | 8,764 | 319,7642902 | 352 | 7927,041 | 0 | 0 | 1 | 0 | 1 | 0 | 0 | 0 | 0 | 0 | 1 | 0 | 0 | 0 | 0 | 0 | 0 |
| 41.00151 | 25.31123 | KOM_R_3 | komr3_5 | Grece | R | 0 | 0 | 1 | 1 | 0 | 0 | 0 | 8,821 | 319,6601996 | 352 | 8254,762 | 0 | 0 | 1 | 0 | 1 | 0 | 0 | 1 | 0 | 0 | 1 | 0 | 0 | 0 | 0 | 0 | 0 |
| 41.00746 | 25.10928 | KOM_R_4 | komr4_1 | Grece | R | 0 | 0 | 0 | 0 | 0 | 0 | 0 | 12,083 | 336,5869353 | 331 | 39383,186 | 0 | 1 | 1 | 1 | 1 | 0 | 0 | 1 | 0 | 0 | 0 | 0 | 0 | 0 | 0 | 0 | 0 |
| 41.00674 | 25.11266 | KOM_R_4 | komr4_2 | Grece | R | 0 | 0 | 2 | 1 | 0 | 0 | 0 | 4,736 | 336,3075061 | 331 | 13471,854 | 4060 | 1 | 1 | 1 | 1 | 0 | 0 | 0 | 1 | 0 | 1 | 0 | 0 | 0 | 0 | 0 | 0 |
| 41.00493 | 25.11539 | KOM_R_4 | komr4_3 | Grece | R | 0 | 0 | 0 | 0 | 0 | 0 | 0 | 3,792 | 336,0890386 | 331 | 4310,548 | 5479 | 0 | 1 | 0 | 0 | 0 | 0 | 0 | 1 | 1 | 0 | 0 | 1 | 0 | 0 | 0 | 0 |
| 41.00297 | 25.11781 | KOM_R_4 | komr4_4 | Grece | R | 0 | 0 | 0 | 0 | 0 | 0 | 0 | 5,196 | 335,8979462 | 331 | 31076,849 | 0 | 1 | 1 | 1 | 1 | 0 | 0 | 0 | 1 | 1 | 0 | 0 | 1 | 0 | 0 | 1 | 0 |
| 41.00246 | 25.12145 | KOM_R_4 | komr4_5 | Grece | R | 0 | 0 | 1 | 1 | 0 | 0 | 0 | 12,364 | 335,595445 | 331 | 54226,079 | 0 | 1 | 0 | 1 | 1 | 0 | 0 | 0 | 0 | 0 | 0 | 0 | 0 | 0 | 0 | 0 | 0 |
| 41.1121 | 25.39901 | KOM_U_1 | komu1_1 | Grece | U | 0 | 0 | 2 | 1 | 0 | 0 | 0 | 36 | 311,8383338 | 366 | 17437,39 | 10188 | 0 | 1 | 0 | 0 | 0 | 0 | 1 | 1 | 0 | 0 | 0 | 1 | 0 | 0 | 1 | 0 |
| 41.114 | 25.39751 | KOM_U_1 | komu1_2 | Grece | U | 0 | 0 | 1 | 1 | 0 | 0 | 0 | 37,764 | 311,9607767 | 366 | 10847,598 | 15380 | 0 | 1 | 0 | 0 | 0 | 0 | 1 | 1 | 0 | 0 | 0 | 0 | 0 | 1 | 1 | 0 |
| 41.11608 | 25.39983 | KOM_U_1 | komu1_3 | Grece | U | 0 | 0 | 0 | 0 | 0 | 0 | 0 | 43,224 | 311,762214 | 366 | 14008,559 | 16621 | 0 | 1 | 0 | 0 | 0 | 0 | 1 | 1 | 0 | 0 | 0 | 0 | 0 | 1 | 1 | 0 |
| 41.11713 | 25.40285 | KOM_U_1 | komu1_4 | Grece | U | 0 | 0 | 0 | 0 | 0 | 0 | 0 | 47,928 | 311,5074319 | 366 | 3729,987 | 21093 | 0 | 1 | 0 | 0 | 0 | 0 | 1 | 0 | 0 | 0 | 0 | 0 | 0 | 1 | 0 | 0 |
| 41.1171 | 25.40601 | KOM_U_1 | komu1_5 | Grece | U | 0 | 0 | 0 | 0 | 0 | 0 | 0 | 45,88 | 311,2421262 | 366 | 12776,858 | 15968 | 0 | 1 | 0 | 0 | 1 | 0 | 1 | 1 | 0 | 0 | 0 | 0 | 0 | 1 | 0 | 0 |
| 41.1239 | 25.38143 | KOM_U_2 | komu2_1 | Grece | U | 0 | 0 | 0 | 0 | 0 | 0 | 0 | 32,778 | 313,295528 | 366 | 25087,601 | 0 | 0 | 1 | 0 | 1 | 0 | 0 | 0 | 1 | 1 | 1 | 1 | 0 | 0 | 0 | 0 | 0 |
| 41.12352 | 25.38543 | KOM_U_2 | komu2_2 | Grece | U | 0 | 0 | 0 | 0 | 0 | 0 | 0 | 33,124 | 312,9599759 | 366 | 6808,767 | 233 | 0 | 1 | 0 | 0 | 0 | 0 | 1 | 0 | 0 | 0 | 1 | 0 | 0 | 0 | 1 | 0 |
| 41.12244 | 25.38896 | KOM_U_2 | komu2_3 | Grece | U | 0 | 0 | 0 | 0 | 0 | 0 | 0 | 32,639 | 312,6657772 | 366 | 8680,067 | 16034 | 0 | 1 | 0 | 0 | 0 | 0 | 1 | 1 | 0 | 0 | 0 | 1 | 1 | 0 | 0 | 0 |
| 41.12318 | 25.39179 | KOM_U_2 | komu2_4 | Grece | U | 0 | 0 | 0 | 0 | 0 | 0 | 0 | 36,199 | 312,4263854 | 366 | 20273,892 | 13023 | 0 | 1 | 0 | 0 | 1 | 0 | 1 | 1 | 0 | 0 | 0 | 0 | 1 | 1 | 0 | 0 |
| 41.12506 | 25.39159 | KOM_U_2 | komu2_5 | Grece | U | 0 | 0 | 0 | 0 | 0 | 0 | 0 | 33,568 | 312,4402752 | 366 | 11839,487 | 10127 | 0 | 1 | 0 | 0 | 0 | 0 | 0 | 1 | 0 | 0 | 1 | 0 | 1 | 0 | 1 | 0 |
| 44.30053 | 23.74315 | KRA_R_1 | krar1_1 | Romania | R | 0 | 0 | 2 | 1 | 0 | 0 | 0 | 72 | 565,43189 | 394 | 46619,215 | 0 | 1 | 1 | 0 | 1 | 0 | 0 | 0 | 0 | 0 | 1 | 0 | 0 | 0 | 0 | 0 | 1 |
| 44.30291 | 23.74179 | KRA_R_1 | krar1_2 | Romania | R | 0 | 0 | 0 | 0 | 0 | 0 | 0 | 71 | 565,6763322 | 394 | 52787,564 | 0 | 1 | 1 | 0 | 1 | 0 | 0 | 0 | 0 | 0 | 1 | 0 | 0 | 0 | 0 | 0 | 0 |
| 44.30488 | 23.73915 | KRA_R_1 | krar1_3 | Romania | R | 0 | 0 | 1 | 1 | 0 | 0 | 0 | 71,466 | 565,9761846 | 394 | 64461,431 | 0 | 1 | 0 | 0 | 1 | 0 | 0 | 0 | 0 | 0 | 1 | 0 | 0 | 0 | 0 | 0 | 0 |
| 44.30689 | 23.73529 | KRA_R_1 | krar1_4 | Romania | R | 1 | 1 | 0 | 0 | 0 | 0 | 0 | 73 | 566,3593954 | 394 | 70634,379 | 0 | 1 | 0 | 0 | 1 | 0 | 0 | 0 | 0 | 0 | 0 | 0 | 0 | 0 | 0 | 0 | 0 |
| 44.30571 | 23.73188 | KRA_R_1 | krar1_5 | Romania | R | 0 | 0 | 0 | 0 | 0 | 0 | 0 | 104,721 | 566,4965177 | 394 | 68123,182 | 0 | 1 | 0 | 0 | 1 | 0 | 0 | 0 | 0 | 0 | 0 | 0 | 0 | 0 | 0 | 0 | 0 |
| 44.26849 | 23.71798 | KRA_R_2 | krar2_1 | Romania | R | 0 | 0 | 0 | 0 | 0 | 0 | 0 | 114,22 | 564,9016674 | 394 | 16705,04 | 6826 | 1 | 1 | 0 | 1 | 0 | 0 | 0 | 0 | 1 | 1 | 0 | 1 | 0 | 0 | 0 | 0 |
| 44.26762 | 23.72075 | KRA_R_2 | krar2_2 | Romania | R | 0 | 0 | 0 | 0 | 0 | 0 | 0 | 106,13 | 564,6669713 | 394 | 15526,873 | 9888 | 0 | 1 | 0 | 0 | 0 | 0 | 1 | 0 | 1 | 1 | 0 | 1 | 0 | 0 | 0 | 0 |
| 44.2675 | 23.7246 | KRA_R_2 | krar2_3 | Romania | R | 0 | 0 | 0 | 0 | 0 | 0 | 0 | 103,44 | 564,4105838 | 394 | 15335,191 | 8190 | 0 | 1 | 0 | 0 | 0 | 0 | 1 | 0 | 1 | 1 | 1 | 1 | 0 | 0 | 0 | 0 |
| 44.26681 | 23.72856 | KRA_R_2 | krar2_4 | Romania | R | 0 | 0 | 0 | 0 | 0 | 0 | 0 | 107,238 | 564,1131355 | 394 | 8987,026 | 3313 | 0 | 1 | 0 | 0 | 0 | 0 | 1 | 0 | 1 | 1 | 1 | 1 | 0 | 0 | 0 | 0 |
| 44.26668 | 23.73187 | KRA_R_2 | krar2_5 | Romania | R | 0 | 0 | 0 | 0 | 0 | 0 | 0 | 119,419 | 563,8901628 | 394 | 6905,643 | 256 | 1 | 1 | 0 | 1 | 0 | 0 | 1 | 0 | 0 | 1 | 1 | 0 | 0 | 0 | 0 | 0 |
| 44.29638 | 23.82657 | KRA_U_1 | krau1_1 | Romania | U | 0 | 0 | 3 | 1 | 0 | 0 | 1 | 94,663 | 559,8583619 | 405 | 19805,57 | 2096 | 1 | 1 | 0 | 1 | 0 | 0 | 0 | 0 | 1 | 1 | 0 | 1 | 0 | 0 | 0 | 0 |
| 44.297 | 23.82262 | KRA_U_1 | krau1_2 | Romania | U | 0 | 0 | 3 | 1 | 0 | 0 | 0 | 90,286 | 560,5008999 | 405 | 48396,435 | 2405 | 1 | 1 | 0 | 1 | 0 | 0 | 1 | 0 | 0 | 1 | 0 | 0 | 1 | 0 | 0 | 0 |
| 44.29809 | 23.81823 | KRA_U_1 | krau1_3 | Romania | U | 0 | 0 | 2 | 1 | 0 | 0 | 0 | 91,946 | 560,665009 | 405 | 42982,909 | 1008 | 1 | 1 | 0 | 1 | 1 | 0 | 1 | 0 | 0 | 1 | 0 | 0 | 1 | 0 | 0 | 0 |
| 44.29708 | 23.81458 | KRA_U_1 | krau1_4 | Romania | U | 0 | 0 | 3 | 1 | 1 | 1 | 1 | 93,536 | 560,5624582 | 405 | 42217,95 | 0 | 1 | 1 | 0 | 1 | 1 | 0 | 1 | 0 | 0 | 1 | 0 | 0 | 0 | 0 | 0 | 0 |
| 44.29462 | 23.81353 | KRA_U_1 | krau1_5 | Romania | U | 0 | 0 | 2 | 1 | 0 | 0 | 0 | 96,708 | 560,1497986 | 405 | 24070,438 | 3921 | 0 | 1 | 0 | 1 | 0 | 0 | 1 | 1 | 0 | 0 | 0 | 1 | 1 | 0 | 0 | 0 |
| 44.30221 | 23.80139 | KRA_U_2 | krau2_1 | Romania | U | 0 | 0 | 2 | 1 | 0 | 0 | 0 | 81,956 | 561,8469314 | 405 | 43629,961 | 7193 | 1 | 1 | 0 | 0 | 1 | 0 | 0 | 1 | 0 | 0 | 0 | 1 | 1 | 0 | 0 | 0 |
| 44.30019 | 23.80067 | KRA_U_2 | krau2_2 | Romania | U | 0 | 0 | 2 | 1 | 0 | 0 | 0 | 76,728 | 561,7559881 | 405 | 56028,45 | 2000 | 1 | 1 | 0 | 0 | 1 | 0 | 0 | 0 | 0 | 1 | 0 | 0 | 0 | 0 | 0 | 0 |
| 44.29857 | 23.80362 | KRA_U_2 | krau2_3 | Romania | U | 0 | 0 | 2 | 1 | 0 | 0 | 0 | 79 | 561,4597188 | 405 | 53396,616 | 2963 | 1 | 1 | 0 | 0 | 1 | 0 | 0 | 0 | 0 | 1 | 0 | 0 | 1 | 0 | 0 | 0 |
| 44.29904 | 23.80686 | KRA_U_2 | krau2_4 | Romania | U | 0 | 0 | 2 | 1 | 0 | 0 | 0 | 84,696 | 561,2864306 | 405 | 53439,944 | 2463 | 1 | 1 | 0 | 0 | 1 | 0 | 0 | 0 | 0 | 1 | 0 | 0 | 0 | 0 | 0 | 0 |
| 44.29979 | 23.81018 | KRA_U_2 | krau2_5 | Romania | U | 0 | 0 | 0 | 0 | 0 | 0 | 0 | 85 | 561,1266174 | 405 | 54792,446 | 124 | 1 | 1 | 0 | 0 | 1 | 0 | 0 | 0 | 0 | 1 | 0 | 0 | 0 | 0 | 0 | 0 |
| 51.2133 | 22.48915 | LBN_R_1 | lbnr1_1 | Poland | R | 0 | 0 | 0 | 0 | 0 | 0 | 1 | 220 | 1233,909526 | 158 | 12095,33 | 4493,390486 | 0 | 1 | 1 | 0 | 0 | 0 | 0 | 0 | 1 | 0 | 1 | 0 | 0 | 0 | 0 | 0 |
| 51.21315 | 22.49343 | LBN_R_1 | lbnr1_2 | Poland | R | 0 | 0 | 0 | 0 | 0 | 0 | 1 | 225 | 1233,762998 | 158 | 12819,184 | 6346,605861 | 0 | 1 | 1 | 1 | 0 | 0 | 0 | 0 | 1 | 0 | 1 | 1 | 0 | 0 | 0 | 0 |
| 51.21373 | 22.49763 | LBN_R_1 | lbnr1_3 | Poland | R | 0 | 0 | 0 | 0 | 0 | 0 | 1 | 224 | 1233,665088 | 158 | 13229,769 | 4968,89873 | 0 | 1 | 1 | 1 | 0 | 0 | 0 | 0 | 1 | 0 | 1 | 0 | 0 | 0 | 0 | 0 |
| 51.21506 | 22.50143 | LBN_R_1 | lbnr1_4 | Poland | R | 1 | 1 | 0 | 0 | 0 | 0 | 0 | 212 | 1233,691315 | 158 | 49478,643 | 571,3398974 | 1 | 1 | 0 | 1 | 0 | 0 | 0 | 0 | 0 | 0 | 1 | 0 | 0 | 0 | 0 | 0 |
| 51.21757 | 22.50158 | LBN_R_1 | lbnr1_5 | Poland | R | 2 | 1 | 0 | 0 | 0 | 0 | 0 | 215 | 1233,921485 | 158 | 32148,533 | 3435,833333 | 1 | 1 | 0 | 1 | 0 | 0 | 1 | 1 | 0 | 0 | 1 | 0 | 0 | 0 | 1 | 0 |
| 51.27425 | 22.62816 | LBN_R_2 | lbnr2_1 | Poland | R | 0 | 0 | 0 | 0 | 0 | 0 | 0 | 180 | 1235,601586 | 153 | 26622,899 | 4582 | 1 | 1 | 1 | 1 | 0 | 0 | 1 | 0 | 0 | 0 | 1 | 0 | 0 | 0 | 0 | 0 |
| 51.27251 | 22.63145 | LBN_R_2 | lbnr2_2 | Poland | R | 0 | 0 | 0 | 0 | 0 | 0 | 0 | 167 | 1235,363768 | 153 | 13463,246 | 0 | 1 | 1 | 0 | 1 | 0 | 0 | 0 | 0 | 0 | 1 | 0 | 0 | 0 | 0 | 0 | 0 |
| 51.27039 | 22.63109 | LBN_R_2 | lbnr2_3 | Poland | R | 0 | 0 | 0 | 0 | 0 | 0 | 0 | 166 | 1235,156181 | 153 | 20458,313 | 473,7350642 | 1 | 1 | 0 | 1 | 0 | 0 | 0 | 0 | 1 | 1 | 1 | 0 | 0 | 0 | 0 | 0 |
| 51.26899 | 22.62738 | LBN_R_2 | lbnr2_4 | Poland | R | 0 | 0 | 0 | 0 | 0 | 0 | 0 | 165 | 1235,119716 | 153 | 15537,319 | 0 | 1 | 1 | 0 | 1 | 0 | 0 | 0 | 0 | 0 | 1 | 0 | 0 | 0 | 0 | 0 | 0 |
| 51.26996 | 22.62437 | LBN_R_2 | lbnr2_5 | Poland | R | 0 | 0 | 0 | 0 | 0 | 0 | 0 | 175 | 1235,307308 | 153 | 26215,643 | 1679 | 1 | 1 | 0 | 1 | 0 | 0 | 0 | 1 | 1 | 0 | 1 | 0 | 0 | 0 | 0 | 0 |
| 51.27349 | 22.55371 | LBN_U_1 | lbnu1_1 | Poland | U | 0 | 0 | 0 | 0 | 0 | 0 | 0 | 229 | 1238,128486 | 153 | 32748,223 | 11276 | 0 | 1 | 0 | 0 | 0 | 0 | 0 | 1 | 0 | 0 | 0 | 1 | 0 | 0 | 0 | 0 |
| 51.27537 | 22.55097 | LBN_U_1 | lbnu1_2 | Poland | U | 0 | 0 | 0 | 0 | 0 | 0 | 0 | 215 | 1237,839247 | 153 | 24383,943 | 12213 | 0 | 1 | 1 | 0 | 0 | 0 | 1 | 1 | 0 | 0 | 0 | 1 | 1 | 0 | 0 | 0 |
| 51.27103 | 22.55491 | LBN_U_1 | lbnu1_3 | Poland | U | 0 | 0 | 0 | 0 | 0 | 0 | 0 | 216 | 1237,544616 | 153 | 28541,236 | 12052 | 0 | 1 | 1 | 0 | 0 | 0 | 1 | 1 | 0 | 0 | 0 | 1 | 0 | 0 | 0 | 0 |
| 51.26945 | 22.55249 | LBN_U_1 | lbnu1_4 | Poland | U | 0 | 0 | 0 | 0 | 0 | 0 | 0 | 221 | 1237,518974 | 153 | 25045,077 | 8320 | 0 | 1 | 1 | 0 | 0 | 0 | 0 | 1 | 1 | 0 | 0 | 1 | 0 | 0 | 0 | 0 |
| 51.26935 | 22.5482 | LBN_U_1 | lbnu1_5 | Poland | U | 0 | 0 | 0 | 0 | 0 | 0 | 0 | 214 | 1237,58379 | 153 | 21084,957 | 10751 | 0 | 1 | 0 | 0 | 0 | 0 | 0 | 1 | 0 | 0 | 0 | 1 | 0 | 0 | 0 | 0 |
| 51.21559 | 22.57122 | LBN_U_2 | lbnu2_1 | Poland | U | 0 | 0 | 0 | 0 | 0 | 0 | 0 | 190 | 1231,545537 | 159 | 17069,592 | 11892 | 0 | 1 | 0 | 0 | 0 | 0 | 0 | 1 | 1 | 0 | 0 | 1 | 1 | 1 | 0 | 0 |
| 51.2151 | 22.56774 | LBN_U_2 | lbnu2_2 | Poland | U | 0 | 0 | 0 | 0 | 0 | 0 | 0 | 205 | 1231,615764 | 159 | 24929,636 | 11560 | 0 | 1 | 0 | 0 | 0 | 0 | 1 | 1 | 0 | 0 | 0 | 1 | 0 | 0 | 0 | 0 |
| 51.21414 | 22.56624 | LBN_U_2 | lbnu2_3 | Poland | U | 0 | 0 | 0 | 0 | 0 | 0 | 0 | 196 | 1231,565353 | 159 | 25945,548 | 7231 | 1 | 1 | 0 | 0 | 0 | 0 | 0 | 1 | 1 | 0 | 0 | 1 | 1 | 0 | 0 | 1 |
| 51.21215 | 22.56928 | LBN_U_2 | lbnu2_4 | Poland | U | 0 | 0 | 0 | 0 | 0 | 0 | 0 | 194 | 1231,282004 | 159 | 28777,694 | 5992 | 1 | 1 | 0 | 0 | 0 | 0 | 0 | 1 | 1 | 0 | 0 | 0 | 1 | 0 | 0 | 0 |
| 51.21033 | 22.57241 | LBN_U_2 | lbnu2_5 | Poland | U | 0 | 0 | 0 | 0 | 0 | 0 | 0 | 188 | 1231,013994 | 159 | 31638,171 | 6138 | 1 | 1 | 0 | 0 | 0 | 0 | 0 | 1 | 1 | 0 | 0 | 1 | 1 | 0 | 0 | 0 |
| 51.24408 | 22.53617 | LBN_U_3 | lbnu3_1 | Poland | U | 0 | 0 | 1 | 1 | 0 | 0 | 1 | 204 | 1235,493745 | 155 | 44836,154 | 3964 | 1 | 1 | 1 | 0 | 1 | 0 | 0 | 1 | 0 | 0 | 0 | 0 | 1 | 0 | 1 | 0 |
| 51.24504 | 22.53951 | LBN_U_3 | lbnu3_2 | Poland | U | 0 | 0 | 0 | 0 | 0 | 0 | 1 | 207 | 1235,452 | 155 | 25808,308 | 15722 | 1 | 1 | 0 | 0 | 1 | 0 | 0 | 1 | 0 | 0 | 0 | 0 | 1 | 0 | 0 | 0 |
| 51.24642 | 22.54225 | LBN_U_3 | lbnu3_3 | Poland | U | 0 | 0 | 0 | 0 | 0 | 0 | 0 | 205 | 1235,612133 | 155 | 19945,969 | 16027 | 0 | 1 | 0 | 0 | 0 | 0 | 0 | 1 | 1 | 0 | 0 | 0 | 1 | 0 | 0 | 0 |
| 51.24867 | 22.54368 | LBN_U_3 | lbnu3_4 | Poland | U | 0 | 0 | 0 | 0 | 0 | 0 | 1 | 207 | 1235,689972 | 155 | 36085,246 | 11715 | 1 | 1 | 1 | 0 | 1 | 0 | 1 | 1 | 0 | 0 | 0 | 0 | 1 | 1 | 1 | 0 |
| 51.25009 | 22.54571 | LBN_U_3 | lbnu3_5 | Poland | U | 0 | 0 | 0 | 0 | 0 | 0 | 0 | 216 | 1235,777705 | 155 | 42629,083 | 4013 | 1 | 1 | 1 | 0 | 1 | 0 | 0 | 1 | 0 | 0 | 1 | 0 | 1 | 0 | 0 | 0 |
| 51.23429 | 22.55476 | LBN_U_4 | lbnu4_1 | Poland | U | 0 | 0 | 0 | 0 | 0 | 0 | 0 | 172 | 1233,943084 | 156 | 18563,42 | 852 | 1 | 1 | 0 | 0 | 0 | 0 | 1 | 1 | 1 | 0 | 1 | 0 | 1 | 0 | 0 | 0 |
| 51.23583 | 22.55805 | LBN_U_4 | lbnu4_2 | Poland | U | 0 | 0 | 0 | 0 | 0 | 0 | 0 | 171 | 1233,950312 | 156 | 31054,199 | 2200 | 1 | 1 | 0 | 0 | 1 | 0 | 1 | 0 | 0 | 0 | 0 | 0 | 0 | 0 | 1 | 0 |
| 51.23572 | 22.56213 | LBN_U_4 | lbnu4_3 | Poland | U | 0 | 0 | 0 | 0 | 0 | 0 | 0 | 171 | 1233,829821 | 156 | 31127,199 | 7165 | 1 | 1 | 0 | 0 | 1 | 0 | 0 | 1 | 0 | 0 | 0 | 0 | 0 | 0 | 1 | 0 |
| 51.23811 | 22.56183 | LBN_U_4 | lbnu4_4 | Poland | U | 0 | 0 | 0 | 0 | 0 | 0 | 0 | 171 | 1234,081823 | 156 | 35094,362 | 2825 | 0 | 1 | 0 | 0 | 1 | 0 | 1 | 1 | 0 | 0 | 0 | 0 | 0 | 1 | 0 | 1 |
| 51.23972 | 22.5596 | LBN_U_4 | lbnu4_5 | Poland | U | 0 | 0 | 0 | 0 | 0 | 0 | 1 | 174 | 1234,310845 | 156 | 23318,348 | 2100 | 1 | 1 | 0 | 0 | 1 | 0 | 1 | 1 | 1 | 1 | 1 | 0 | 0 | 0 | 1 | 1 |
| 48.2408 | 14.40395 | LIN_R_1 | linr1_1 | Austria | R | 1 | 1 | 1 | 1 | 1 | 1 | 0 | 246,926 | 1406,846766 | -42 | 7873,414 | 2907 | 0 | 1 | 0 | 0 | 0 | 0 | 0 | 0 | 1 | 0 | 0 | 1 | 0 | 0 | 0 | 0 |
| 48.24223 | 14.39988 | LIN_R_1 | linr1_2 | Austria | R | 0 | 0 | 0 | 0 | 0 | 0 | 0 | 247,972 | 1407,26025 | -42 | 5238,778 | 585 | 0 | 1 | 0 | 1 | 0 | 0 | 0 | 0 | 1 | 0 | 1 | 0 | 0 | 0 | 0 | 0 |
| 48.24311 | 14.39567 | LIN_R_1 | linr1_3 | Austria | R | 2 | 1 | 0 | 0 | 0 | 0 | 0 | 246,527 | 1407,577833 | -42 | 32653,66 | 0 | 1 | 1 | 0 | 1 | 0 | 0 | 0 | 0 | 0 | 0 | 0 | 0 | 0 | 0 | 0 | 0 |
| 48.2428 | 14.39107 | LIN_R_1 | linr1_4 | Austria | R | 1 | 1 | 0 | 0 | 0 | 0 | 0 | 245,08 | 1407,860371 | -42 | 16455,504 | 0 | 1 | 1 | 0 | 1 | 0 | 0 | 0 | 0 | 0 | 0 | 0 | 0 | 0 | 0 | 0 | 0 |
| 48.24443 | 14.38838 | LIN_R_1 | linr1_5 | Austria | R | 1 | 1 | 0 | 0 | 0 | 0 | 0 | 246,159 | 1408,123327 | -42 | 4264,512 | 0 | 0 | 1 | 0 | 1 | 0 | 0 | 0 | 0 | 1 | 0 | 0 | 0 | 0 | 0 | 0 | 0 |
| 48.26141 | 14.358 | LIN_R_2 | linr2_1 | Austria | R | 3 | 1 | 0 | 0 | 0 | 0 | 1 | 250,739 | 1411,009566 | -43 | 30169,182 | 4286 | 1 | 1 | 1 | 1 | 1 | 0 | 1 | 1 | 0 | 1 | 0 | 0 | 1 | 0 | 0 | 0 |
| 48.26348 | 14.36257 | LIN_R_2 | linr2_2 | Austria | R | 1 | 1 | 0 | 0 | 0 | 0 | 1 | 250,014 | 1410,824098 | -43 | 29032,165 | 0 | 1 | 1 | 1 | 1 | 0 | 0 | 1 | 1 | 0 | 1 | 0 | 0 | 0 | 0 | 0 | 0 |
| 48.26332 | 14.36675 | LIN_R_2 | linr2_3 | Austria | R | 1 | 1 | 0 | 0 | 0 | 0 | 1 | 249,952 | 1410,543599 | -43 | 20583,655 | 2039 | 1 | 1 | 1 | 1 | 0 | 0 | 1 | 1 | 1 | 1 | 0 | 1 | 0 | 0 | 0 | 1 |
| 48.26243 | 14.37069 | LIN_R_2 | linr2_4 | Austria | R | 1 | 1 | 0 | 0 | 0 | 0 | 0 | 249,386 | 1410,240266 | -43 | 20214,174 | 133 | 1 | 0 | 1 | 1 | 0 | 0 | 0 | 0 | 1 | 1 | 0 | 1 | 0 | 0 | 0 | 0 |
| 48.26037 | 14.37364 | LIN_R_2 | linr2_5 | Austria | R | 2 | 1 | 0 | 0 | 0 | 0 | 1 | 245,069 | 1409,936597 | -43 | 28731,447 | 0 | 1 | 0 | 1 | 1 | 0 | 0 | 1 | 0 | 0 | 1 | 0 | 0 | 0 | 0 | 0 | 1 |
| 48.27845 | 14.28659 | LIN_U_1 | linu1_1 | Austria | U | 0 | 0 | 0 | 0 | 0 | 0 | 0 | 269,392 | 1416,567697 | -48 | 9272,53 | 13344 | 0 | 1 | 0 | 0 | 0 | 0 | 0 | 0 | 1 | 0 | 1 | 1 | 0 | 0 | 0 | 0 |
| 48.27861 | 14.29045 | LIN_U_1 | linu1_2 | Austria | U | 2 | 1 | 0 | 0 | 0 | 0 | 0 | 275,998 | 1416,326263 | -48 | 30855,359 | 7246 | 1 | 1 | 0 | 0 | 1 | 0 | 0 | 1 | 1 | 0 | 0 | 1 | 1 | 0 | 0 | 0 |
| 48.27705 | 14.29348 | LIN_U_1 | linu1_3 | Austria | U | 1 | 1 | 0 | 0 | 0 | 0 | 0 | 280 | 1416,043433 | -48 | 42013,682 | 3283 | 1 | 1 | 0 | 0 | 1 | 0 | 0 | 1 | 0 | 0 | 1 | 0 | 0 | 0 | 1 | 1 |
| 48.27516 | 14.29252 | LIN_U_1 | linu1_4 | Austria | U | 2 | 1 | 0 | 0 | 0 | 0 | 0 | 280 | 1416,006589 | -48 | 32286,737 | 12796 | 1 | 1 | 0 | 0 | 1 | 0 | 0 | 1 | 0 | 0 | 1 | 0 | 1 | 1 | 0 | 0 |
| 48.27566 | 14.28883 | LIN_U_1 | linu1_5 | Austria | U | 1 | 1 | 0 | 0 | 0 | 0 | 0 | 279,624 | 1416,273143 | -48 | 13746,334 | 16979 | 0 | 1 | 0 | 0 | 0 | 0 | 0 | 1 | 1 | 0 | 0 | 1 | 1 | 0 | 0 | 0 |
| 48.26275 | 14.30302 | LIN_U_2 | linu2_1 | Austria | U | 1 | 1 | 0 | 0 | 0 | 0 | 0 | 262 | 1414,659248 | -48 | 19358,84 | 3905 | 1 | 1 | 0 | 1 | 0 | 0 | 0 | 0 | 1 | 0 | 1 | 0 | 1 | 0 | 1 | 0 |

| | | | | | | | | | | | | | | | | | | | | | | | | | | | | | | | | | |
|---|---|---|---|---|---|---|---|---|---|---|---|---|---|---|---|---|---|---|---|---|---|---|---|---|---|---|---|---|---|---|---|---|---|
| 48.26189 | 14.30541 | LIN_U_2 | linu2_2 | Austria | U | 0 | 0 | 0 | 0 | 0 | 0 | 0 | 262 | 1414,44357 | -48 | 36260,226 | 2556 | 1 | 1 | 0 | 1 | 0 | 0 | 0 | 0 | 0 | 0 | 1 | 0 | 1 | 0 | 0 | 0 |
| 48.25966 | 14.30778 | LIN_U_2 | linu2_3 | Austria | U | 3 | 1 | 0 | 0 | 0 | 0 | 0 | 262 | 1414,173004 | -48 | 47166,21 | 2079 | 1 | 1 | 1 | 1 | 0 | 0 | 0 | 0 | 1 | 0 | 0 | 1 | 0 | 0 | 0 | 0 |
| 48.25741 | 14.30928 | LIN_U_2 | linu2_4 | Austria | U | 2 | 1 | 0 | 0 | 0 | 0 | 0 | 260 | 1414,003723 | -48 | 38114,645 | 438 | 1 | 1 | 0 | 1 | 1 | 0 | 0 | 0 | 0 | 0 | 0 | 0 | 0 | 0 | 0 | 0 |
| 48.25744 | 14.30441 | LIN_U_2 | linu2_5 | Austria | U | 1 | 1 | 0 | 0 | 0 | 0 | 0 | 261 | 1414,28155 | -48 | 41337,401 | 462 | 1 | 1 | 0 | 1 | 0 | 0 | 0 | 0 | 0 | 0 | 0 | 0 | 0 | 0 | 0 | 0 |
| 51.77361 | 19.48565 | LOD_U_1 | lodu1_1 | Poland | U | 2 | 1 | 0 | 0 | 0 | 0 | 1 | 228,34 | 1395,586629 | 74 | 30479,908 | 9980 | 1 | 1 | 0 | 0 | 1 | 0 | 1 | 1 | 0 | 0 | 0 | 1 | 1 | 1 | 0 | 0 |
| 51.77107 | 19.48589 | LOD_U_1 | lodu1_2 | Poland | U | 0 | 0 | 0 | 0 | 0 | 0 | 1 | 226,204 | 1395,347649 | 74 | 31368,88 | 10864 | 1 | 1 | 1 | 0 | 0 | 0 | 0 | 1 | 1 | 0 | 0 | 1 | 1 | 0 | 0 | 0 |
| 51.76991 | 19.48794 | LOD_U_1 | lodu1_3 | Poland | U | 2 | 1 | 0 | 0 | 0 | 0 | 1 | 229,648 | 1395,16055 | 74 | 44962,536 | 5381 | 1 | 1 | 1 | 0 | 1 | 0 | 1 | 1 | 1 | 0 | 0 | 1 | 1 | 0 | 0 | 0 |
| 51.77026 | 19.49222 | LOD_U_1 | lodu1_4 | Poland | U | 1 | 1 | 0 | 0 | 0 | 0 | 1 | 231 | 1395,020196 | 74 | 43989,981 | 7199 | 1 | 1 | 1 | 0 | 1 | 0 | 1 | 1 | 1 | 0 | 0 | 1 | 1 | 0 | 0 | 0 |
| 51.77057 | 19.49648 | LOD_U_1 | lodu1_5 | Poland | U | 1 | 1 | 0 | 0 | 0 | 0 | 1 | 234,052 | 1394,877732 | 74 | 41376,439 | 4991 | 1 | 1 | 1 | 0 | 1 | 0 | 1 | 1 | 1 | 0 | 0 | 1 | 1 | 0 | 0 | 0 |
| 51.77099 | 19.50071 | LOD_U_1 | lodu1_6 | Poland | U | 1 | 1 | 0 | 0 | 0 | 0 | 1 | 234,564 | 1394,746655 | 74 | 42467,858 | 1100 | 1 | 1 | 1 | 1 | 1 | 0 | 0 | 1 | 0 | 0 | 1 | 0 | 0 | 0 | 0 | 0 |
| 51.77261 | 19.50235 | LOD_U_1 | lodu1_7 | Poland | U | 1 | 1 | 0 | 0 | 0 | 0 | 0 | 236,508 | 1394,827302 | 74 | 60489,997 | 610 | 1 | 0 | 0 | 1 | 0 | 0 | 0 | 0 | 0 | 0 | 0 | 0 | 0 | 0 | 1 | 0 |
| 51.77515 | 19.50148 | LOD_U_1 | lodu1_8 | Poland | U | 2 | 1 | 0 | 0 | 0 | 0 | 0 | 242,026 | 1395,093315 | 74 | 38234,916 | 700 | 1 | 1 | 0 | 1 | 0 | 0 | 0 | 1 | 0 | 0 | 0 | 0 | 0 | 0 | 1 | 0 |
| 49.2838 | 20.75209 | LUB_R_1 | lubr1_1 | Slovakia | R | 0 | 0 | 0 | 0 | 0 | 0 | 0 | 505,832 | 1120,518051 | 290 | 24181,081 | 377 | 1 | 1 | 1 | 1 | 0 | 0 | 1 | 0 | 0 | 0 | 1 | 0 | 0 | 0 | 0 | 0 |
| 49.28385 | 20.74819 | LUB_R_1 | lubr1_2 | Slovakia | R | 0 | 0 | 0 | 0 | 0 | 0 | 0 | 506,276 | 1120,698177 | 290 | 46585,557 | 1383 | 1 | 1 | 0 | 1 | 0 | 0 | 1 | 0 | 0 | 1 | 0 | 0 | 0 | 0 | 1 | 0 |
| 49.28458 | 20.74355 | LUB_R_1 | lubr1_3 | Slovakia | R | 0 | 0 | 0 | 0 | 0 | 0 | 0 | 508,078 | 1120,974247 | 290 | 18657,841 | 952 | 1 | 1 | 0 | 1 | 0 | 0 | 1 | 0 | 0 | 1 | 0 | 0 | 0 | 0 | 1 | 0 |
| 49.28617 | 20.74016 | LUB_R_1 | lubr1_4 | Slovakia | R | 0 | 0 | 0 | 0 | 0 | 0 | 0 | 508,244 | 1121,267905 | 290 | 18690,122 | 807 | 0 | 1 | 0 | 1 | 0 | 0 | 1 | 0 | 0 | 1 | 0 | 0 | 0 | 0 | 1 | 0 |
| 49.28721 | 20.7433 | LUB_R_1 | lubr1_5 | Slovakia | R | 1 | 1 | 0 | 0 | 0 | 0 | 0 | 528,42 | 1121,2138 | 290 | 36567,333 | 3806 | 1 | 1 | 0 | 1 | 0 | 0 | 0 | 0 | 1 | 0 | 0 | 0 | 0 | 0 | 1 | 1 |
| 49.31373 | 20.70367 | LUB_R_2 | lubr2_1 | Slovakia | R | 1 | 1 | 0 | 0 | 0 | 0 | 0 | 563,197 | 1125,319714 | 295 | 48562,639 | 592 | 1 | 0 | 1 | 1 | 0 | 0 | 0 | 0 | 1 | 0 | 1 | 0 | 0 | 0 | 0 | 0 |
| 49.31563 | 20.7011 | LUB_R_2 | lubr2_2 | Slovakia | R | 2 | 1 | 0 | 0 | 0 | 0 | 0 | 620,752 | 1125,600926 | 295 | 29434,111 | 5637 | 1 | 1 | 1 | 1 | 0 | 0 | 1 | 0 | 0 | 0 | 0 | 0 | 1 | 0 | 0 | 0 |
| 49.31642 | 20.69779 | LUB_R_2 | lubr2_3 | Slovakia | R | 2 | 1 | 0 | 0 | 0 | 0 | 0 | 628,356 | 1125,819912 | 295 | 41487,287 | 3727 | 1 | 1 | 0 | 1 | 0 | 0 | 1 | 0 | 0 | 0 | 1 | 0 | 1 | 0 | 0 | 0 |
| 49.31362 | 20.69614 | LUB_R_2 | lubr2_4 | Slovakia | R | 0 | 0 | 1 | 1 | 0 | 0 | 0 | 630,783 | 1125,654599 | 295 | 19261,438 | 2938 | 1 | 1 | 1 | 1 | 0 | 0 | 1 | 1 | 0 | 0 | 0 | 1 | 0 | 0 | 0 | 0 |
| 49.31421 | 20.69118 | LUB_R_2 | lubr2_5 | Slovakia | R | 0 | 0 | 0 | 0 | 0 | 0 | 0 | 598,511 | 1125,931673 | 295 | 1236,052 | 0 | 0 | 1 | 0 | 1 | 0 | 0 | 1 | 0 | 0 | 0 | 0 | 0 | 0 | 0 | 0 | 0 |
| 49.30211 | 20.68166 | LUB_U_1 | lubu1_1 | Slovakia | U | 0 | 0 | 0 | 0 | 0 | 0 | 0 | 527,482 | 1125,321858 | 290 | 14781,011 | 5304 | 1 | 1 | 1 | 1 | 0 | 0 | 1 | 0 | 0 | 0 | 1 | 1 | 0 | 0 | 0 | 0 |
| 49.29989 | 20.68378 | LUB_U_1 | lubu1_2 | Slovakia | U | 0 | 0 | 0 | 0 | 0 | 0 | 0 | 538,371 | 1125,033367 | 290 | 13746,486 | 7957 | 0 | 1 | 1 | 0 | 0 | 1 | 1 | 0 | 0 | 0 | 0 | 1 | 1 | 0 | 0 | 0 |
| 49.29881 | 20.68706 | LUB_U_1 | lubu1_3 | Slovakia | U | 0 | 0 | 0 | 0 | 0 | 0 | 0 | 551 | 1124,789831 | 290 | 2205,301 | 14780 | 0 | 1 | 0 | 0 | 0 | 1 | 0 | 1 | 0 | 0 | 0 | 1 | 0 | 1 | 0 | 0 |
| 49.30005 | 20.69009 | LUB_U_1 | lubu1_4 | Slovakia | U | 0 | 0 | 0 | 0 | 0 | 0 | 0 | 547,554 | 1124,758261 | 290 | 14052,332 | 13964 | 1 | 1 | 1 | 0 | 0 | 0 | 1 | 1 | 1 | 1 | 0 | 1 | 1 | 1 | 0 | 0 |
| 49.30208 | 20.6905 | LUB_U_1 | lubu1_5 | Slovakia | U | 0 | 0 | 0 | 0 | 0 | 0 | 0 | 545,707 | 1124,91485 | 290 | 8163,156 | 25884 | 0 | 1 | 0 | 0 | 0 | 0 | 0 | 1 | 1 | 0 | 0 | 0 | 0 | 1 | 0 | 0 |
| 49.29618 | 20.68927 | LUB_U_2 | lubu2_1 | Slovakia | U | 0 | 0 | 0 | 0 | 0 | 0 | 0 | 552,628 | 1124,462929 | 290 | 9442,261 | 12215 | 0 | 1 | 0 | 0 | 0 | 0 | 1 | 1 | 0 | 0 | 0 | 1 | 1 | 0 | 1 | 0 |
| 49.29361 | 20.68795 | LUB_U_2 | lubu2_2 | Slovakia | U | 0 | 0 | 0 | 0 | 0 | 0 | 0 | 553,266 | 1124,299407 | 290 | 18510,879 | 12542 | 1 | 1 | 1 | 0 | 1 | 0 | 0 | 1 | 1 | 0 | 0 | 1 | 1 | 0 | 0 | 0 |
| 49.29089 | 20.68674 | LUB_U_2 | lubu2_3 | Slovakia | U | 0 | 0 | 0 | 0 | 0 | 0 | 0 | 548,144 | 1124,121053 | 290 | 8090,747 | 7076 | 1 | 1 | 1 | 0 | 1 | 0 | 0 | 1 | 1 | 0 | 0 | 1 | 0 | 0 | 0 | 0 |
| 49.29245 | 20.68391 | LUB_U_2 | lubu2_4 | Slovakia | U | 0 | 0 | 0 | 0 | 0 | 0 | 0 | 560,398 | 1124,386434 | 290 | 2081,073 | 6702 | 0 | 1 | 0 | 0 | 0 | 0 | 0 | 1 | 1 | 0 | 0 | 1 | 0 | 0 | 1 | 0 |
| 49.29447 | 20.68052 | LUB_U_2 | lubu2_5 | Slovakia | U | 0 | 0 | 0 | 0 | 0 | 0 | 0 | 573,024 | 1124,715353 | 290 | 7200,768 | 15160 | 0 | 1 | 0 | 0 | 0 | 0 | 0 | 1 | 1 | 0 | 0 | 1 | 1 | 0 | 1 | 0 |
| 49.93256 | 23.71391 | LVI_R_1 | lvir1_1 | Ukraine | R | 0 | 0 | 0 | 0 | 0 | 0 | 1 | 293,616 | 1067,803402 | 308 | 14508,492 | 2300 | 1 | 1 | 0 | 1 | 0 | 0 | 0 | 0 | 1 | 0 | 1 | 0 | 0 | 0 | 0 | 0 |
| 49.93402 | 23.71035 | LVI_R_1 | lvir1_2 | Ukraine | R | 0 | 0 | 0 | 0 | 0 | 0 | 0 | 308,469 | 1068,051965 | 308 | 29313,229 | 1875 | 1 | 1 | 0 | 1 | 0 | 0 | 0 | 0 | 1 | 1 | 1 | 0 | 0 | 0 | 0 | 0 |
| 49.93529 | 23.70663 | LVI_R_1 | lvir1_3 | Ukraine | R | 0 | 0 | 0 | 0 | 0 | 0 | 0 | 295 | 1068,299878 | 308 | 30567,071 | 2500 | 1 | 1 | 0 | 1 | 0 | 0 | 0 | 0 | 1 | 1 | 1 | 0 | 0 | 0 | 0 | 0 |
| 49.93658 | 23.70291 | LVI_R_1 | lvir1_4 | Ukraine | R | 0 | 0 | 0 | 0 | 0 | 0 | 0 | 289,672 | 1068,542649 | 308 | 37468,267 | 1040 | 1 | 1 | 0 | 1 | 0 | 0 | 0 | 0 | 0 | 1 | 1 | 0 | 0 | 0 | 0 | 0 |
| 49.9382 | 23.69939 | LVI_R_1 | lvir1_5 | Ukraine | R | 0 | 0 | 0 | 0 | 0 | 0 | 0 | 293,81 | 1068,815742 | 308 | 26259,828 | 577 | 1 | 1 | 0 | 1 | 0 | 0 | 0 | 0 | 0 | 1 | 1 | 0 | 0 | 0 | 0 | 0 |
| 49.89759 | 23.76368 | LVI_R_2 | lvir2_1 | Ukraine | R | 0 | 0 | 0 | 0 | 0 | 0 | 1 | 320,44 | 1062,762355 | 313 | 50376,166 | 4320 | 1 | 0 | 0 | 1 | 0 | 0 | 0 | 0 | 0 | 0 | 1 | 1 | 0 | 0 | 0 | 0 |
| 49.89742 | 23.75958 | LVI_R_2 | lvir2_2 | Ukraine | R | 0 | 0 | 0 | 0 | 0 | 0 | 0 | 339,683 | 1062,879487 | 313 | 45462,677 | 5056 | 1 | 0 | 0 | 1 | 0 | 0 | 0 | 0 | 1 | 0 | 1 | 0 | 0 | 0 | 0 | 0 |
| 49.89618 | 23.75912 | LVI_R_2 | lvir2_3 | Ukraine | R | 0 | 0 | 0 | 0 | 0 | 0 | 0 | 349,706 | 1062,725391 | 313 | 20544,237 | 9984 | 1 | 1 | 0 | 1 | 0 | 0 | 0 | 0 | 1 | 0 | 1 | 1 | 0 | 0 | 0 | 0 |
| 49.89555 | 23.75631 | LVI_R_2 | lvir2_4 | Ukraine | R | 0 | 0 | 0 | 0 | 0 | 0 | 0 | 323,664 | 1062,796691 | 313 | 43372,062 | 97 | 1 | 0 | 0 | 1 | 0 | 1 | 0 | 0 | 0 | 0 | 1 | 0 | 0 | 0 | 0 | 0 |
| 49.89734 | 23.7537 | LVI_R_2 | lvir2_5 | Ukraine | R | 0 | 0 | 0 | 0 | 0 | 0 | 0 | 299,272 | 1063,045574 | 313 | 34485,406 | 0 | 1 | 0 | 0 | 1 | 0 | 1 | 0 | 0 | 0 | 0 | 0 | 0 | 0 | 0 | 0 | 0 |
| 49.849406 | 24.143656 | LVI_R_3 | lvir3_1 | Ukraine | R | 1 | 1 | 0 | 0 | 0 | 0 | 0 | 238,138 | 1046,618214 | 332 | 6253,663 | 2750 | 0 | 1 | 0 | 1 | 0 | 0 | 0 | 0 | 1 | 0 | 1 | 0 | 0 | 0 | 0 | 0 |
| 49.846761 | 24.143186 | LVI_R_3 | lvir3_2 | Ukraine | R | 1 | 1 | 0 | 0 | 0 | 0 | 0 | 240,841 | 1046,368595 | 332 | 8341,147 | 3623 | 0 | 1 | 0 | 0 | 0 | 0 | 0 | 1 | 1 | 0 | 0 | 1 | 0 | 0 | 1 | 0 |
| 49.845296 | 24.146713 | LVI_R_3 | lvir3_3 | Ukraine | R | 0 | 0 | 0 | 0 | 0 | 0 | 0 | 249,647 | 1046,112931 | 332 | 20063,078 | 5951 | 0 | 1 | 0 | 0 | 0 | 0 | 0 | 0 | 1 | 0 | 0 | 1 | 0 | 0 | 0 | 0 |
| 49.844690 | 24.150760 | LVI_R_3 | lvir3_4 | Ukraine | R | 0 | 0 | 0 | 0 | 0 | 0 | 0 | 248,496 | 1045,937371 | 332 | 11221,3 | 5702 | 0 | 1 | 0 | 0 | 0 | 0 | 0 | 0 | 1 | 0 | 1 | 1 | 0 | 0 | 0 | 0 |
| 49.842276 | 24.152587 | LVI_R_3 | lvir3_5 | Ukraine | R | 1 | 1 | 0 | 0 | 0 | 0 | 0 | 255,626 | 1045,640335 | 332 | 13348,19 | 3363 | 1 | 1 | 0 | 1 | 0 | 1 | 0 | 0 | 1 | 0 | 1 | 1 | 1 | 0 | 0 | 0 |
| 49.776495 | 24.088248 | LVI_R_4 | lvir4_1 | Ukraine | R | 0 | 0 | 0 | 0 | 0 | 0 | 0 | 350,236 | 1039,606502 | 331 | 10557,186 | 4150 | 0 | 1 | 0 | 1 | 0 | 0 | 0 | 0 | 1 | 0 | 1 | 1 | 0 | 0 | 1 | 0 |
| 49.773896 | 24.090038 | LVI_R_4 | lvir4_2 | Ukraine | R | 0 | 0 | 0 | 0 | 0 | 0 | 0 | 348,022 | 1039,904985 | 331 | 11598,658 | 4176 | 0 | 1 | 0 | 1 | 0 | 0 | 0 | 0 | 1 | 0 | 1 | 1 | 0 | 0 | 1 | 0 |
| 49.771233 | 24.091170 | LVI_R_4 | lvir4_3 | Ukraine | R | 1 | 1 | 0 | 0 | 0 | 0 | 0 | 351,532 | 1040,191197 | 331 | 10703,275 | 9423 | 0 | 1 | 0 | 1 | 0 | 0 | 0 | 1 | 1 | 0 | 1 | 1 | 0 | 0 | 0 | 0 |
| 49.768486 | 24.091439 | LVI_R_4 | lvir4_4 | Ukraine | R | 0 | 0 | 0 | 0 | 0 | 0 | 0 | 348,901 | 1040,494834 | 331 | 19504,651 | 6397 | 1 | 1 | 0 | 1 | 0 | 0 | 1 | 0 | 1 | 0 | 1 | 1 | 0 | 0 | 0 | 0 |
| 49.766305 | 24.094001 | LVI_R_4 | lvir4_5 | Ukraine | R | 0 | 0 | 1 | 1 | 0 | 0 | 0 | 352,698 | 1040,812217 | 331 | 19268,269 | 5973 | 1 | 1 | 0 | 1 | 0 | 0 | 0 | 0 | 1 | 0 | 0 | 1 | 0 | 0 | 1 | 0 |
| 49.85153 | 24.04252 | LVI_U_1 | lviu1_1 | Ukraine | U | 1 | 1 | 0 | 0 | 0 | 0 | 0 | 281,15 | 1049,773352 | 320 | 32012,92 | 14138 | 1 | 1 | 0 | 1 | 0 | 0 | 1 | 1 | 0 | 0 | 0 | 1 | 0 | 1 | 1 | 0 |
| 49.85085 | 24.03859 | LVI_U_1 | lviu1_2 | Ukraine | U | 1 | 1 | 0 | 0 | 0 | 0 | 0 | 292,648 | 1049,81746 | 320 | 45340,129 | 7201 | 1 | 1 | 0 | 1 | 0 | 0 | 1 | 1 | 0 | 0 | 1 | 1 | 0 | 0 | 1 | 0 |
| 49.85042 | 24.03452 | LVI_U_1 | lviu1_3 | Ukraine | U | 2 | 1 | 0 | 0 | 0 | 0 | 0 | 301,863 | 1049,89571 | 320 | 52120,449 | 4650 | 1 | 0 | 0 | 1 | 0 | 0 | 0 | 0 | 0 | 0 | 0 | 1 | 1 | 0 | 1 | 0 |
| 49.84934 | 24.03079 | LVI_U_1 | lviu1_4 | Ukraine | U | 1 | 1 | 0 | 0 | 0 | 0 | 0 | 313,476 | 1049,887637 | 320 | 47543 | 5269 | 1 | 1 | 0 | 1 | 0 | 0 | 0 | 0 | 0 | 0 | 0 | 0 | 1 | 0 | 1 | 0 |
| 49.84699 | 24.02999 | LVI_U_1 | lviu1_5 | Ukraine | U | 0 | 0 | 0 | 0 | 0 | 0 | 0 | 297,128 | 1049,671916 | 320 | 16387,213 | 24330 | 1 | 1 | 0 | 0 | 1 | 0 | 0 | 1 | 0 | 0 | 0 | 0 | 0 | 1 | 0 | 0 |
| 49.85152 | 24.04251 | LVI_U_2 | lviu2_1 | Ukraine | U | 0 | 0 | 0 | 0 | 0 | 0 | 1 | 281,15 | 1047,472768 | 332 | 12921,005 | 11925 | 1 | 1 | 0 | 0 | 1 | 0 | 1 | 1 | 1 | 0 | 0 | 1 | 0 | 1 | 1 | 0 |
| 49.85084 | 24.03858 | LVI_U_2 | lviu2_2 | Ukraine | U | 0 | 0 | 0 | 0 | 0 | 0 | 0 | 292,648 | 1047,328676 | 332 | 36789,669 | 7575 | 1 | 1 | 0 | 0 | 1 | 0 | 0 | 1 | 1 | 0 | 0 | 1 | 1 | 0 | 0 | 0 |
| 49.8304 | 24.05945 | LVI_U_2 | lviu2_3 | Ukraine | U | 1 | 1 | 0 | 0 | 0 | 0 | 0 | 323,311 | 1047,144149 | 332 | 30125,969 | 1026 | 1 | 1 | 0 | 1 | 1 | 1 | 0 | 1 | 0 | 0 | 0 | 0 | 1 | 0 | 1 | 0 |
| 49.82835 | 24.05812 | LVI_U_2 | lviu2_4 | Ukraine | U | 0 | 0 | 0 | 0 | 0 | 0 | 1 | 310,232 | 1046,961569 | 332 | 22047,036 | 6635 | 1 | 1 | 0 | 1 | 1 | 1 | 0 | 1 | 0 | 0 | 1 | 0 | 1 | 0 | 0 | 0 |
| 49.82608 | 24.05928 | LVI_U_2 | lviu2_5 | Ukraine | U | 2 | 1 | 0 | 0 | 0 | 0 | 1 | 313,613 | 1046,696257 | 332 | 56820,853 | 5699 | 1 | 0 | 0 | 1 | 0 | 0 | 0 | 0 | 0 | 0 | 0 | 0 | 1 | 0 | 0 | 0 |
| 49.818731 | 24.083768 | LVI_U_3 | lviu3_1 | Ukraine | U | 2 | 1 | 0 | 0 | 0 | 0 | 1 | 350,709 | 1045,237636 | 326 | 58460,526 | 2075 | 1 | 1 | 0 | 1 | 0 | 0 | 0 | 1 | 0 | 0 | 0 | 0 | 1 | 0 | 0 | 0 |
| 49.818112 | 24.087785 | LVI_U_3 | lviu3_2 | Ukraine | U | 2 | 1 | 0 | 0 | 0 | 0 | 0 | 319,891 | 1045,042721 | 326 | 45926,33 | 860 | 1 | 1 | 0 | 1 | 0 | 0 | 0 | 0 | 0 | 0 | 1 | 0 | 0 | 0 | 0 | 0 |
| 49.816862 | 24.091602 | LVI_U_3 | lviu3_3 | Ukraine | U | 0 | 0 | 0 | 0 | 0 | 0 | 0 | 316,678 | 1044,811128 | 326 | 70385,739 | 0 | 1 | 0 | 0 | 1 | 0 | 0 | 0 | 0 | 0 | 0 | 0 | 0 | 0 | 0 | 0 | 0 |
| 49.815159 | 24.094792 | LVI_U_3 | lviu3_4 | Ukraine | U | 0 | 0 | 0 | 0 | 0 | 0 | 0 | 312,214 | 1044,545766 | 326 | 70855 | 0 | 1 | 0 | 0 | 1 | 0 | 0 | 0 | 0 | 0 | 0 | 0 | 0 | 0 | 0 | 0 | 0 |
| 49.814393 | 24.099133 | LVI_U_3 | lviu3_5 | Ukraine | U | 0 | 0 | 0 | 0 | 0 | 0 | 0 | 297,741 | 1044,3491 | 326 | 70855 | 0 | 1 | 0 | 0 | 1 | 0 | 0 | 0 | 0 | 0 | 0 | 0 | 0 | 0 | 0 | 0 | 0 |
| 49.826110 | 24.029908 | LVI_U_4 | lviu4_1 | Ukraine | U | 0 | 0 | 0 | 0 | 0 | 0 | 1 | 308,697 | 1047,563264 | 322 | 55712,048 | 4940 | 1 | 1 | 0 | 1 | 0 | 0 | 0 | 0 | 0 | 0 | 1 | 0 | 1 | 0 | 0 | 1 |
| 49.823997 | 24.027808 | LVI_U_4 | lviu4_2 | Ukraine | U | 1 | 1 | 0 | 0 | 0 | 0 | 0 | 339,72 | 1047,408503 | 322 | 60456,502 | 3515 | 1 | 1 | 0 | 1 | 0 | 0 | 0 | 0 | 0 | 0 | 0 | 0 | 1 | 0 | 0 | 0 |
| 49.822067 | 24.025225 | LVI_U_4 | lviu4_3 | Ukraine | U | 0 | 0 | 0 | 0 | 0 | 0 | 0 | 340,654 | 1047,284479 | 322 | 58242,399 | 3836 | 1 | 1 | 0 | 1 | 0 | 0 | 0 | 0 | 0 | 0 | 1 | 0 | 0 | 0 | 0 | 0 |
| 49.822763 | 24.020994 | LVI_U_4 | lviu4_4 | Ukraine | U | 2 | 1 | 0 | 0 | 0 | 0 | 1 | 345,682 | 1047,478886 | 322 | 45077,715 | 2592 | 1 | 1 | 0 | 1 | 0 | 0 | 0 | 0 | 0 | 0 | 0 | 0 | 0 | 1 | 1 | 0 |
| 49.825509 | 24.021633 | LVI_U_4 | lviu4_5 | Ukraine | U | 0 | 0 | 0 | 0 | 0 | 0 | 1 | 345,946 | 1047,744654 | 322 | 44748,706 | 2996 | 1 | 1 | 0 | 0 | 0 | 0 | 0 | 0 | 0 | 0 | 0 | 0 | 1 | 0 | 1 | 0 |
| 52.16897 | 20.73737 | MAZ_R_37 | mazr37_1 | Poland | R | 0 | 0 | 0 | 0 | 0 | 0 | 1 | 92,708 | 1385,449748 | 39 | 12404,338 | 5393 | 1 | 1 | 0 | 0 | 0 | 0 | 1 | 1 | 1 | 0 | 1 | 0 | 1 | 0 | 1 | 0 |
| 52.16977 | 20.73334 | MAZ_R_37 | mazr37_2 | Poland | R | 0 | 0 | 0 | 0 | 0 | 0 | 1 | 92,664 | 1385,66907 | 39 | 11136,419 | 2958 | 0 | 1 | 0 | 0 | 0 | 0 | 1 | 1 | 1 | 1 | 1 | 0 | 1 | 0 | 0 | 0 |
| 52.17044 | 20.72933 | MAZ_R_37 | mazr37_3 | Poland | R | 0 | 0 | 0 | 0 | 0 | 0 | 1 | 93 | 1385,874486 | 39 | 10818,559 | 3584 | 0 | 1 | 1 | 0 | 0 | 0 | 1 | 1 | 1 | 1 | 1 | 0 | 0 | 0 | 1 | 0 |
| 52.17123 | 20.72506 | MAZ_R_37 | mazr37_4 | Poland | R | 0 | 0 | 0 | 0 | 0 | 0 | 1 | 93,336 | 1386,09487 | 39 | 15998,679 | 1486 | 1 | 1 | 0 | 0 | 0 | 1 | 1 | 1 | 1 | 1 | 1 | 0 | 0 | 0 | 0 | 0 |
| 52.17078 | 20.72149 | MAZ_R_37 | mazr37_5 | Poland | R | 0 | 0 | 0 | 0 | 0 | 0 | 1 | 93 | 1386,177507 | 39 | 9481,785 | 1968 | 1 | 1 | 0 | 1 | 0 | 0 | 1 | 0 | 0 | 1 | 1 | 0 | 0 | 0 | 0 | 0 |
| 52.17094 | 20.71803 | MAZ_R_37 | mazr37_6 | Poland | R | 0 | 0 | 0 | 0 | 0 | 0 | 1 | 93 | 1386,313855 | 39 | 3161,926 | 791 | 0 | 1 | 0 | 0 | 0 | 0 | 1 | 0 | 1 | 0 | 1 | 0 | 0 | 0 | 0 | 0 |
| 52.16878 | 20.71642 | MAZ_R_37 | mazr37_7 | Poland | R | 0 | 0 | 0 | 0 | 0 | 0 | 0 | 92,348 | 1386,165415 | 39 | 4439,224 | 88 | 0 | 1 | 0 | 0 | 0 | 0 | 1 | 0 | 1 | 1 | 1 | 0 | 0 | 0 | 0 | 0 |
| 52.16699 | 20.71262 | MAZ_R_37 | mazr37_8 | Poland | R | 0 | 0 | 0 | 0 | 0 | 0 | 1 | 92,404 | 1386,126598 | 39 | 8123,368 | 38 | 0 | 1 | 0 | 0 | 0 | 0 | 1 | 0 | 0 | 1 | 1 | 0 | 0 | 0 | 0 | 0 |
| 52.2201 | 20.90034 | MAZ_U_1 | mazu1_1 | Poland | U | 0 | 0 | 0 | 0 | 0 | 0 | 1 | 112 | 1384,734292 | 34 | 16679,8 | 12305 | 0 | 1 | 0 | 0 | 0 | 0 | 0 | 1 | 1 | 0 | 0 | 1 | 1 | 0 | 0 | 0 |
| 52.22272 | 20.89994 | MAZ_U_1 | mazu1_2 | Poland | U | 0 | 0 | 0 | 0 | 0 | 0 | 1 | 111,792 | 1384,993811 | 34 | 25683,825 | 16256 | 1 | 1 | 0 | 0 | 0 | 0 | 1 | 1 | 0 | 0 | 0 | 0 | 1 | 0 | 0 | 0 |
| 52.22509 | 20.9003 | MAZ_U_1 | mazu1_3 | Poland | U | 0 | 0 | 0 | 0 | 0 | 0 | 1 | 112 | 1385,214953 | 34 | 24583,398 | 10798 | 0 | 1 | 0 | 0 | 0 | 0 | 1 | 1 | 0 | 0 | 0 | 0 | 1 | 0 | 0 | 0 |
| 52.22728 | 20.90022 | MAZ_U_1 | mazu1_4 | Poland | U | 0 | 0 | 0 | 0 | 0 | 0 | 1 | 111,373 | 1385,43278 | 34 | 18767,529 | 16059 | 0 | 1 | 0 | 0 | 0 | 0 | 0 | 1 | 0 | 0 | 0 | 0 | 1 | 1 | 0 | 0 |
| 52.22993 | 20.90113 | MAZ_U_1 | mazu1_5 | Poland | U | 0 | 0 | 0 | 0 | 0 | 0 | 1 | 110,252 | 1385,652134 | 34 | 15234,913 | 10859 | 0 | 1 | 0 | 0 | 0 | 0 | 1 | 1 | 0 | 0 | 0 | 0 | 1 | 0 | 1 | 0 |
| 52.23256 | 20.90182 | MAZ_U_1 | mazu1_6 | Poland | U | 0 | 0 | 0 | 0 | 0 | 0 | 1 | 110 | 1385,880631 | 34 | 16181,558 | 4288 | 0 | 1 | 0 | 0 | 1 | 0 | 0 | 1 | 0 | 0 | 0 | 0 | 1 | 0 | 0 | 0 |
| 52.23487 | 20.90234 | MAZ_U_1 | mazu1_7 | Poland | U | 0 | 0 | 0 | 0 | 0 | 0 | 1 | 109,468 | 1386,085065 | 34 | 22861,977 | 9919 | 0 | 1 | 0 | 0 | 0 | 0 | 0 | 1 | 0 | 0 | 0 | 0 | 1 | 1 | 0 | 0 |
| 52.23709 | 20.90293 | MAZ_U_1 | mazu1_8 | Poland | U | 0 | 0 | 0 | 0 | 0 | 0 | 1 | 110,763 | 1386,278544 | 34 | 27422,192 | 9449 | 0 | 1 | 0 | 0 | 0 | 0 | 0 | 1 | 0 | 0 | 0 | 0 | 1 | 0 | 0 | 0 |
| 52.28031 | 20.91769 | MAZ_U_10 | mazu10_1 | Poland | U | 0 | 0 | 0 | 0 | 0 | 0 | 0 | 102 | 1389,945353 | 27 | 42260,485 | 8467 | 1 | 1 | 0 | 1 | 0 | 0 | 1 | 0 | 0 | 0 | 0 | 0 | 1 | 0 | 0 | 0 |
| 52.28246 | 20.92022 | MAZ_U_10 | mazu10_2 | Poland | U | 0 | 0 | 0 | 0 | 0 | 0 | 1 | 102 | 1390,066862 | 27 | 33325,057 | 5406 | 1 | 1 | 0 | 1 | 0 | 0 | 0 | 1 | 0 | 0 | 1 | 0 | 0 | 1 | 1 | 0 |
| 52.28458 | 20.92262 | MAZ_U_10 | mazu10_3 | Poland | U | 2 | 1 | 0 | 0 | 0 | 0 | 1 | 101,488 | 1390,190453 | 27 | 39761,831 | 5165 | 1 | 1 | 0 | 1 | 0 | 0 | 0 | 1 | 0 | 0 | 0 | 0 | 1 | 1 | 0 | 0 |
| 52.28681 | 20.92539 | MAZ_U_10 | mazu10_4 | Poland | U | 0 | 0 | 0 | 0 | 0 | 0 | 1 | 100,497 | 1390,311188 | 27 | 37971,244 | 6556 | 1 | 1 | 0 | 1 | 0 | 0 | 0 | 1 | 0 | 0 | 0 | 0 | 1 | 1 | 0 | 0 |
| 52.28908 | 20.92821 | MAZ_U_10 | mazu10_5 | Poland | U | 0 | 0 | 0 | 0 | 0 | 0 | 1 | 99,695 | 1390,435204 | 27 | 36198,184 | 7290 | 1 | 0 | 0 | 1 | 0 | 0 | 0 | 0 | 0 | 0 | 0 | 0 | 0 | 1 | 1 | 0 |
| 52.291 | 20.93077 | MAZ_U_10 | mazu10_6 | Poland | U | 0 | 0 | 0 | 0 | 0 | 0 | 1 | 100 | 1390,533633 | 27 | 9906,996 | 6946 | 0 | 1 | 0 | 0 | 0 | 0 | 0 | 1 | 0 | 0 | 0 | 0 | 1 | 1 | 1 | 0 |
| 52.29339 | 20.93362 | MAZ_U_10 | mazu10_7 | Poland | U | 0 | 0 | 0 | 0 | 0 | 0 | 1 | 100,025 | 1390,667982 | 27 | 22343,825 | 5771 | 1 | 1 | 1 | 1 | 0 | 0 | 0 | 1 | 1 | 0 | 0 | 1 | 1 | 0 | 1 | 0 |
| 52.29572 | 20.93643 | MAZ_U_10 | mazu10_8 | Poland | U | 0 | 0 | 0 | 0 | 0 | 0 | 1 | 101 | 1390,797547 | 27 | 17577,038 | 3870 | 0 | 1 | 1 | 0 | 0 | 0 | 0 | 1 | 1 | 0 | 0 | 1 | 1 | 0 | 0 | 0 |
| 52.2341 | 21.06984 | MAZ_U_11 | mazu11_1 | Poland | U | 0 | 0 | 2 | 1 | 0 | 0 | 1 | 80,678 | 1380,339942 | 34 | 40608,675 | 623 | 1 | 1 | 1 | 0 | 0 | 0 | 0 | 1 | 1 | 1 | 0 | 1 | 0 | 0 | 0 | 0 |
| 52.23593 | 21.0669 | MAZ_U_11 | mazu11_2 | Poland | U | 0 | 0 | 0 | 0 | 0 | 0 | 1 | 81,415 | 1380,616598 | 34 | 39544,448 | 7115 | 0 | 1 | 1 | 0 | 0 | 0 | 1 | 1 | 1 | 1 | 0 | 1 | 1 | 0 | 0 | 0 |
| 52.23832 | 21.0654 | MAZ_U_11 | mazu11_3 | Poland | U | 0 | 0 | 0 | 0 | 0 | 0 | 0 | 81,162 | 1380,897223 | 34 | 34949,685 | 6024 | 0 | 1 | 0 | 0 | 0 | 0 | 1 | 1 | 1 | 1 | 0 | 1 | 1 | 0 | 0 | 0 |
| 52.24026 | 21.06306 | MAZ_U_11 | mazu11_4 | Poland | U | 0 | 0 | 0 | 0 | 0 | 0 | 1 | 82,064 | 1381,163716 | 34 | 30563,179 | 8472 | 1 | 1 | 0 | 0 | 1 | 0 | 1 | 1 | 1 | 1 | 0 | 0 | 1 | 0 | 1 | 0 |
| 52.24229 | 21.06125 | MAZ_U_11 | mazu11_5 | Poland | U | 0 | 0 | 0 | 0 | 0 | 0 | 1 | 81,378 | 1381,42082 | 34 | 41844,96 | 230 | 1 | 1 | 1 | 0 | 1 | 0 | 0 | 0 | 0 | 0 | 1 | 0 | 0 | 0 | 0 | 0 |
| 52.24455 | 21.06024 | MAZ_U_11 | mazu11_6 | Poland | U | 1 | 1 | 0 | 0 | 0 | 0 | 1 | 81,259 | 1381,672899 | 34 | 23297,649 | 134 | 1 | 1 | 0 | 0 | 0 | 0 | 1 | 1 | 0 | 1 | 0 | 0 | 1 | 0 | 0 | 0 |
| 52.24442 | 21.06462 | MAZ_U_11 | mazu11_7 | Poland | U | 1 | 1 | 0 | 0 | 0 | 0 | 1 | 80,576 | 1381,513735 | 34 | 19016,541 | 4757 | 0 | 1 | 0 | 0 | 0 | 0 | 0 | 1 | 0 | 1 | 0 | 0 | 1 | 0 | 0 | 0 |

| | | | | | | | | | | | | | | | | | | | | | | | | | | | | | | | | | |
|---|---|---|---|---|---|---|---|---|---|---|---|---|---|---|---|---|---|---|---|---|---|---|---|---|---|---|---|---|---|---|---|---|---|
| 52.24418 | 21.0693 | MAZ_U_11 | mazu11_8 | Poland | U | 0 | 0 | 0 | 0 | 0 | 0 | 1 | 83 | 1381,332404 | 34 | 15162,44 | 11573 | 0 | 1 | 0 | 0 | 0 | 0 | 0 | 1 | 0 | 1 | 0 | 0 | 1 | 0 | 0 | 0 |
| 52.2263 | 21.10686 | MAZ_U_12 | mazu12_1 | Poland | U | 0 | 0 | 0 | 0 | 0 | 0 | 0 | 81,793 | 1378,340115 | 35 | 9737,654 | 19499 | 0 | 1 | 0 | 0 | 0 | 0 | 1 | 1 | 0 | 0 | 0 | 0 | 0 | 1 | 0 | 0 |
| 52.22772 | 21.10325 | MAZ_U_12 | mazu12_2 | Poland | U | 0 | 0 | 0 | 0 | 0 | 0 | 1 | 81,446 | 1378,598731 | 35 | 10432,363 | 16747 | 0 | 1 | 0 | 0 | 0 | 0 | 1 | 1 | 0 | 0 | 0 | 0 | 1 | 1 | 0 | 0 |
| 52.2283 | 21.09879 | MAZ_U_12 | mazu12_3 | Poland | U | 0 | 0 | 0 | 0 | 0 | 0 | 0 | 82 | 1378,804266 | 35 | 14291,554 | 9438 | 0 | 1 | 0 | 0 | 0 | 0 | 1 | 1 | 0 | 0 | 0 | 0 | 1 | 0 | 0 | 0 |
| 52.22879 | 21.09407 | MAZ_U_12 | mazu12_4 | Poland | U | 0 | 0 | 0 | 0 | 0 | 0 | 1 | 82,908 | 1379,009958 | 35 | 9661,548 | 7442 | 0 | 1 | 0 | 0 | 0 | 0 | 1 | 1 | 0 | 0 | 0 | 0 | 1 | 0 | 0 | 0 |
| 52.22959 | 21.08975 | MAZ_U_12 | mazu12_5 | Poland | U | 0 | 0 | 0 | 0 | 0 | 0 | 0 | 81,524 | 1379,232884 | 35 | 8233,965 | 6226 | 0 | 1 | 0 | 0 | 0 | 0 | 0 | 1 | 0 | 0 | 0 | 0 | 1 | 0 | 0 | 0 |
| 52.22946 | 21.08657 | MAZ_U_12 | mazu12_6 | Poland | U | 0 | 0 | 0 | 0 | 0 | 0 | 1 | 81,901 | 1379,444466 | 35 | 11968,545 | 3603 | 0 | 1 | 0 | 0 | 1 | 0 | 0 | 0 | 0 | 1 | 0 | 0 | 1 | 0 | 0 | 0 |
| 52.22933 | 21.08269 | MAZ_U_12 | mazu12_7 | Poland | U | 0 | 0 | 0 | 0 | 0 | 0 | 1 | 81,728 | 1379,326653 | 35 | 10134,319 | 14066 | 0 | 1 | 0 | 0 | 0 | 0 | 0 | 1 | 0 | 0 | 0 | 0 | 1 | 0 | 0 | 0 |
| 52.2306 | 21.07911 | MAZ_U_12 | mazu12_8 | Poland | U | 0 | 0 | 0 | 0 | 0 | 0 | 1 | 81 | 1379,687342 | 35 | 11928,337 | 9544 | 0 | 1 | 0 | 0 | 0 | 0 | 0 | 1 | 0 | 1 | 0 | 0 | 1 | 0 | 0 | 0 |
| 52.22017 | 21.09933 | MAZ_U_13 | mazu13_1 | Poland | U | 0 | 0 | 0 | 0 | 0 | 0 | 1 | 84 | 1377,998521 | 36 | 19329,627 | 11494 | 0 | 1 | 0 | 0 | 0 | 0 | 1 | 1 | 0 | 0 | 0 | 0 | 1 | 0 | 1 | 0 |
| 52.21917 | 21.096 | MAZ_U_13 | mazu13_2 | Poland | U | 0 | 0 | 0 | 0 | 0 | 0 | 0 | 84,8 | 1378,012565 | 36 | 12801,59 | 10043 | 0 | 1 | 0 | 0 | 0 | 0 | 0 | 1 | 0 | 0 | 0 | 0 | 1 | 0 | 1 | 0 |
| 52.21991 | 21.09088 | MAZ_U_13 | mazu13_3 | Poland | U | 1 | 1 | 0 | 0 | 0 | 0 | 1 | 84 | 1378,256432 | 36 | 29997,519 | 6776 | 1 | 1 | 0 | 1 | 0 | 0 | 0 | 1 | 0 | 0 | 1 | 0 | 1 | 0 | 0 | 0 |
| 52.22071 | 21.08669 | MAZ_U_13 | mazu13_4 | Poland | U | 0 | 0 | 0 | 0 | 0 | 0 | 1 | 85,739 | 1378,474416 | 36 | 25660,373 | 15787 | 1 | 1 | 0 | 1 | 0 | 0 | 0 | 1 | 0 | 0 | 0 | 0 | 1 | 0 | 0 | 0 |
| 52.22166 | 21.08195 | MAZ_U_13 | mazu13_5 | Poland | U | 0 | 0 | 0 | 0 | 0 | 0 | 1 | 84,072 | 1378,72577 | 36 | 22677,051 | 9538 | 1 | 1 | 0 | 1 | 0 | 0 | 0 | 1 | 0 | 0 | 0 | 0 | 1 | 0 | 0 | 0 |
| 52.22351 | 21.07958 | MAZ_U_13 | mazu13_6 | Poland | U | 0 | 0 | 2 | 1 | 0 | 0 | 1 | 83,364 | 1378,984987 | 36 | 29298,621 | 13533 | 1 | 1 | 0 | 0 | 1 | 0 | 0 | 1 | 0 | 0 | 0 | 0 | 1 | 1 | 0 | 0 |
| 52.22514 | 21.07617 | MAZ_U_13 | mazu13_7 | Poland | U | 1 | 1 | 1 | 1 | 0 | 0 | 1 | 83 | 1379,258024 | 36 | 31579,004 | 7906 | 1 | 1 | 0 | 0 | 1 | 0 | 0 | 1 | 0 | 0 | 0 | 1 | 0 | 1 | 0 | 0 |
| 52.22533 | 21.07255 | MAZ_U_13 | mazu13_8 | Poland | U | 0 | 0 | 0 | 0 | 0 | 0 | 1 | 83,188 | 1379,397441 | 36 | 18227,437 | 9786 | 0 | 1 | 0 | 0 | 0 | 0 | 1 | 1 | 0 | 0 | 0 | 0 | 1 | 0 | 0 | 0 |
| 52.23287 | 21.12008 | MAZ_U_14 | mazu14_1 | Poland | U | 0 | 0 | 0 | 0 | 0 | 0 | 0 | 85,576 | 1378,535274 | 34 | 13223,399 | 14722 | 0 | 1 | 0 | 0 | 0 | 0 | 0 | 1 | 0 | 0 | 0 | 0 | 1 | 0 | 0 | 0 |
| 52.23505 | 21.12136 | MAZ_U_14 | mazu14_2 | Poland | U | 0 | 0 | 0 | 0 | 0 | 0 | 0 | 83,019 | 1378,706127 | 34 | 17102,859 | 17958 | 0 | 1 | 0 | 0 | 0 | 0 | 0 | 1 | 0 | 0 | 0 | 0 | 1 | 0 | 1 | 0 |
| 52.23698 | 21.12002 | MAZ_U_14 | mazu14_3 | Poland | U | 0 | 0 | 0 | 0 | 0 | 0 | 1 | 85,191 | 1378,935054 | 34 | 5219,533 | 20330 | 0 | 1 | 0 | 0 | 0 | 0 | 1 | 0 | 0 | 0 | 0 | 0 | 1 | 1 | 0 | 0 |
| 52.24061 | 21.12051 | MAZ_U_14 | mazu14_4 | Poland | U | 0 | 0 | 0 | 0 | 0 | 0 | 0 | 85,328 | 1379,270332 | 34 | 14902,003 | 16455 | 0 | 1 | 0 | 0 | 0 | 0 | 0 | 1 | 0 | 0 | 0 | 0 | 0 | 1 | 0 | 0 |
| 52.2433 | 21.12027 | MAZ_U_14 | mazu14_5 | Poland | U | 0 | 0 | 0 | 0 | 0 | 0 | 0 | 85,975 | 1379,539418 | 34 | 18318,245 | 9634 | 0 | 1 | 0 | 1 | 0 | 0 | 1 | 1 | 0 | 0 | 0 | 0 | 0 | 1 | 0 | 0 |
| 52.24432 | 21.11706 | MAZ_U_14 | mazu14_6 | Poland | U | 0 | 0 | 0 | 0 | 0 | 0 | 0 | 85 | 1379,747372 | 34 | 16769,496 | 12895 | 1 | 1 | 1 | 0 | 0 | 0 | 1 | 1 | 0 | 0 | 0 | 1 | 0 | 1 | 0 | 0 |
| 52.24469 | 21.11243 | MAZ_U_14 | mazu14_7 | Poland | U | 0 | 0 | 0 | 0 | 0 | 0 | 0 | 85 | 1379,938063 | 34 | 8359,499 | 15670 | 0 | 1 | 0 | 0 | 0 | 0 | 0 | 1 | 0 | 0 | 0 | 0 | 0 | 1 | 0 | 0 |
| 52.24221 | 21.11156 | MAZ_U_14 | mazu14_8 | Poland | U | 1 | 1 | 0 | 0 | 0 | 0 | 1 | 84,589 | 1379,726286 | 34 | 39247,03 | 4832 | 1 | 1 | 0 | 0 | 1 | 0 | 0 | 1 | 0 | 0 | 0 | 0 | 1 | 1 | 0 | 0 |
| 52.21729 | 21.10583 | MAZ_U_15 | mazu15_1 | Poland | U | 0 | 0 | 0 | 0 | 0 | 0 | 1 | 84 | 1377,500132 | 37 | 16460,158 | 8360 | 1 | 1 | 0 | 1 | 0 | 0 | 0 | 1 | 1 | 0 | 0 | 1 | 1 | 0 | 0 | 0 |
| 52.21467 | 21.10743 | MAZ_U_15 | mazu15_2 | Poland | U | 0 | 0 | 0 | 0 | 0 | 0 | 1 | 84 | 1377,193217 | 37 | 3851,616 | 11805 | 0 | 1 | 1 | 0 | 0 | 0 | 0 | 1 | 0 | 0 | 0 | 1 | 0 | 1 | 0 | 0 |
| 52.21218 | 21.10882 | MAZ_U_15 | mazu15_3 | Poland | U | 0 | 0 | 0 | 0 | 0 | 0 | 1 | 84 | 1376,90495 | 37 | 2514,428 | 12660 | 0 | 1 | 0 | 0 | 0 | 0 | 0 | 1 | 0 | 0 | 0 | 1 | 0 | 1 | 0 | 0 |
| 52.20954 | 21.11039 | MAZ_U_15 | mazu15_4 | Poland | U | 0 | 0 | 0 | 0 | 0 | 0 | 1 | 85 | 1376,596475 | 37 | 1459,226 | 11074 | 0 | 1 | 0 | 0 | 0 | 0 | 1 | 1 | 0 | 0 | 0 | 1 | 0 | 1 | 0 | 0 |
| 52.20783 | 21.11268 | MAZ_U_15 | mazu15_5 | Poland | U | 0 | 0 | 0 | 0 | 0 | 0 | 0 | 84,934 | 1376,354625 | 37 | 525,881 | 10876 | 0 | 1 | 0 | 0 | 0 | 0 | 0 | 1 | 0 | 0 | 0 | 1 | 0 | 1 | 0 | 0 |
| 52.20866 | 21.11651 | MAZ_U_15 | mazu15_6 | Poland | U | 0 | 0 | 0 | 0 | 0 | 0 | 0 | 84 | 1376,30895 | 37 | 116,978 | 8423 | 0 | 0 | 0 | 0 | 0 | 0 | 0 | 0 | 0 | 0 | 0 | 1 | 0 | 1 | 0 | 0 |
| 52.21113 | 21.11621 | MAZ_U_15 | mazu15_7 | Poland | U | 0 | 0 | 0 | 0 | 0 | 0 | 0 | 83,4 | 1376,555387 | 37 | 959,985 | 7817 | 1 | 1 | 0 | 0 | 1 | 0 | 0 | 1 | 0 | 0 | 0 | 1 | 0 | 1 | 0 | 0 |
| 52.21369 | 21.11467 | MAZ_U_15 | mazu15_8 | Poland | U | 0 | 0 | 0 | 0 | 0 | 0 | 1 | 83,053 | 1376,855334 | 37 | 1293,661 | 9400 | 0 | 1 | 0 | 0 | 0 | 0 | 0 | 1 | 0 | 0 | 0 | 1 | 0 | 1 | 0 | 0 |
| 52.17101 | 21.03845 | MAZ_U_16 | mazu16_1 | Poland | U | 0 | 0 | 0 | 0 | 0 | 0 | 1 | 91,216 | 1375,293703 | 41 | 30075,573 | 5275 | 1 | 1 | 0 | 0 | 1 | 0 | 0 | 1 | 0 | 1 | 0 | 0 | 1 | 0 | 0 | 0 |
| 52.16876 | 21.0349 | MAZ_U_16 | mazu16_2 | Poland | U | 0 | 0 | 0 | 0 | 0 | 0 | 1 | 91 | 1375,195237 | 41 | 24270,925 | 1659 | 1 | 1 | 0 | 0 | 1 | 0 | 0 | 1 | 0 | 1 | 0 | 0 | 1 | 0 | 0 | 0 |
| 52.16796 | 21.03072 | MAZ_U_16 | mazu16_3 | Poland | U | 0 | 0 | 0 | 0 | 0 | 0 | 1 | 91 | 1375,259785 | 41 | 43876,555 | 105 | 1 | 1 | 0 | 0 | 1 | 0 | 0 | 1 | 0 | 1 | 0 | 0 | 0 | 0 | 0 | 0 |
| 52.16785 | 21.02596 | MAZ_U_16 | mazu16_4 | Poland | U | 0 | 0 | 1 | 1 | 0 | 0 | 1 | 93,141 | 1375,41217 | 41 | 23081,204 | 3850 | 1 | 1 | 0 | 0 | 1 | 0 | 0 | 1 | 0 | 1 | 0 | 0 | 0 | 0 | 1 | 0 |
| 52.16797 | 21.02131 | MAZ_U_16 | mazu16_5 | Poland | U | 0 | 0 | 0 | 0 | 0 | 0 | 1 | 94 | 1375,580958 | 41 | 33585,37 | 2437 | 1 | 1 | 0 | 0 | 1 | 0 | 0 | 1 | 0 | 1 | 0 | 0 | 1 | 0 | 0 | 0 |
| 52.16896 | 21.01695 | MAZ_U_16 | mazu16_6 | Poland | U | 0 | 0 | 0 | 0 | 0 | 0 | 1 | 95,256 | 1375,823143 | 41 | 19245,26 | 11230 | 1 | 1 | 0 | 0 | 1 | 0 | 0 | 1 | 0 | 1 | 0 | 0 | 1 | 0 | 1 | 0 |
| 52.16985 | 21.01309 | MAZ_U_16 | mazu16_7 | Poland | U | 0 | 0 | 0 | 0 | 0 | 0 | 1 | 100,279 | 1376,039743 | 41 | 25678,756 | 13060 | 1 | 1 | 0 | 0 | 1 | 0 | 0 | 1 | 0 | 1 | 0 | 0 | 0 | 1 | 0 | 0 |
| 52.17106 | 21.01007 | MAZ_U_16 | mazu16_8 | Poland | U | 0 | 0 | 0 | 0 | 0 | 0 | 0 | 101 | 1376,259235 | 41 | 16320,614 | 11962 | 0 | 1 | 0 | 0 | 0 | 0 | 0 | 1 | 0 | 0 | 0 | 0 | 1 | 0 | 0 | 0 |
| 52.22057 | 20.91028 | MAZ_U_2 | mazu2_1 | Poland | U | 0 | 0 | 0 | 0 | 0 | 0 | 0 | 113 | 1384,434513 | 34 | 13647,408 | 13100 | 0 | 1 | 1 | 0 | 0 | 0 | 0 | 1 | 1 | 0 | 0 | 1 | 1 | 0 | 0 | 0 |
| 52.2232 | 20.91064 | MAZ_U_2 | mazu2_2 | Poland | U | 0 | 0 | 0 | 0 | 0 | 0 | 1 | 113,638 | 1384,676866 | 34 | 12448,967 | 12069 | 0 | 1 | 0 | 0 | 0 | 0 | 0 | 1 | 1 | 0 | 0 | 1 | 1 | 0 | 0 | 0 |
| 52.2248 | 20.9111 | MAZ_U_2 | mazu2_3 | Poland | U | 0 | 0 | 0 | 0 | 0 | 0 | 1 | 113,28 | 1384,81503 | 34 | 21058,825 | 13172 | 0 | 1 | 0 | 0 | 0 | 0 | 1 | 1 | 1 | 0 | 0 | 1 | 1 | 0 | 0 | 0 |
| 52.22744 | 20.91084 | MAZ_U_2 | mazu2_4 | Poland | U | 0 | 0 | 0 | 0 | 0 | 0 | 1 | 112 | 1385,078933 | 34 | 15334,567 | 10048 | 0 | 1 | 0 | 0 | 0 | 0 | 1 | 1 | 0 | 0 | 0 | 0 | 1 | 0 | 0 | 0 |
| 52.23002 | 20.91037 | MAZ_U_2 | mazu2_5 | Poland | U | 0 | 0 | 0 | 0 | 0 | 0 | 1 | 111,668 | 1385,343445 | 34 | 8281,488 | 10157 | 0 | 1 | 0 | 0 | 0 | 0 | 1 | 1 | 0 | 0 | 0 | 1 | 1 | 0 | 0 | 0 |
| 52.23223 | 20.91069 | MAZ_U_2 | mazu2_6 | Poland | U | 0 | 0 | 0 | 0 | 0 | 0 | 1 | 111 | 1385,545361 | 34 | 16994,516 | 10041 | 0 | 1 | 0 | 0 | 1 | 0 | 0 | 1 | 0 | 0 | 1 | 0 | 1 | 0 | 0 | 0 |
| 52.23444 | 20.90976 | MAZ_U_2 | mazu2_7 | Poland | U | 0 | 0 | 0 | 0 | 0 | 0 | 1 | 110 | 1385,790361 | 34 | 14228,412 | 7377 | 0 | 1 | 0 | 0 | 1 | 0 | 1 | 1 | 0 | 0 | 0 | 0 | 1 | 1 | 0 | 0 |
| 52.23692 | 20.90994 | MAZ_U_2 | mazu2_8 | Poland | U | 0 | 0 | 0 | 0 | 0 | 0 | 1 | 110 | 1386,022255 | 34 | 12471,598 | 11930 | 0 | 1 | 0 | 0 | 0 | 0 | 0 | 1 | 0 | 0 | 0 | 0 | 1 | 0 | 1 | 0 |
| 51.39848 | 21.15823 | MAZ_U_20 | mazu20_1 | Poland | U | 0 | 0 | 0 | 0 | 0 | 0 | 1 | 171 | 1296,972577 | 127 | 55227,452 | 2893 | 1 | 0 | 0 | 0 | 1 | 0 | 0 | 1 | 0 | 0 | 0 | 0 | 0 | 1 | 0 | 0 |
| 51.39804 | 21.16133 | MAZ_U_20 | mazu20_2 | Poland | U | 0 | 0 | 0 | 0 | 0 | 0 | 1 | 170 | 1296,819719 | 127 | 25836,98 | 20189 | 1 | 1 | 0 | 0 | 1 | 0 | 0 | 1 | 0 | 0 | 0 | 0 | 0 | 1 | 1 | 0 |
| 51.40033 | 21.16268 | MAZ_U_20 | mazu20_3 | Poland | U | 0 | 0 | 0 | 0 | 0 | 0 | 0 | 172,066 | 1296,989334 | 127 | 8485,804 | 24311 | 0 | 1 | 0 | 0 | 0 | 0 | 0 | 1 | 0 | 0 | 0 | 0 | 1 | 1 | 0 | 0 |
| 51.4025 | 21.16466 | MAZ_U_20 | mazu20_4 | Poland | U | 0 | 0 | 3 | 1 | 0 | 0 | 1 | 169,224 | 1297,12505 | 127 | 24257,207 | 13315 | 1 | 1 | 0 | 0 | 1 | 0 | 1 | 1 | 0 | 0 | 0 | 0 | 1 | 1 | 0 | 0 |
| 51.40317 | 21.16781 | MAZ_U_20 | mazu20_5 | Poland | U | 1 | 1 | 2 | 1 | 0 | 0 | 0 | 164,128 | 1297,075496 | 127 | 40385,102 | 4100 | 1 | 1 | 0 | 0 | 1 | 0 | 0 | 1 | 1 | 1 | 1 | 1 | 1 | 0 | 0 | 0 |
| 51.40307 | 21.17112 | MAZ_U_20 | mazu20_6 | Poland | U | 0 | 0 | 0 | 0 | 0 | 0 | 1 | 163,002 | 1296,94692 | 127 | 22141,047 | 9790 | 1 | 1 | 0 | 0 | 1 | 0 | 0 | 1 | 1 | 1 | 0 | 1 | 1 | 0 | 0 | 0 |
| 51.40393 | 21.17415 | MAZ_U_20 | mazu20_7 | Poland | U | 0 | 0 | 0 | 0 | 0 | 0 | 1 | 167,148 | 1296,920356 | 127 | 13370,973 | 16708 | 1 | 1 | 0 | 0 | 0 | 0 | 0 | 1 | 1 | 0 | 0 | 1 | 1 | 0 | 1 | 0 |
| 51.40327 | 21.17805 | MAZ_U_20 | mazu20_8 | Poland | U | 0 | 0 | 2 | 1 | 0 | 0 | 0 | 164,783 | 1296,717129 | 127 | 32002,426 | 5561 | 1 | 1 | 0 | 1 | 0 | 0 | 1 | 1 | 1 | 1 | 0 | 1 | 1 | 0 | 1 | 0 |
| 51.41025 | 21.15367 | MAZ_U_21 | mazu21_1 | Poland | U | 1 | 1 | 0 | 0 | 0 | 0 | 1 | 159,921 | 1298,258299 | 127 | 22985,274 | 12206 | 0 | 1 | 0 | 0 | 1 | 0 | 0 | 1 | 0 | 0 | 0 | 0 | 1 | 1 | 0 | 0 |
| 51.41136 | 21.15095 | MAZ_U_21 | mazu21_2 | Poland | U | 1 | 1 | 2 | 1 | 0 | 0 | 0 | 161,104 | 1298,462947 | 127 | 20603,216 | 10596 | 0 | 1 | 0 | 0 | 1 | 0 | 0 | 1 | 0 | 0 | 0 | 0 | 1 | 0 | 0 | 0 |
| 51.4113 | 21.14648 | MAZ_U_21 | mazu21_3 | Poland | U | 0 | 0 | 0 | 0 | 0 | 0 | 1 | 152,648 | 1298,61678 | 127 | 13143,692 | 8850 | 0 | 1 | 0 | 0 | 0 | 0 | 0 | 1 | 1 | 0 | 0 | 1 | 1 | 0 | 0 | 0 |
| 51.41035 | 21.1426 | MAZ_U_21 | mazu21_4 | Poland | U | 0 | 0 | 0 | 0 | 0 | 0 | 1 | 150,374 | 1298,665667 | 127 | 15867,16 | 6536 | 0 | 1 | 0 | 0 | 1 | 0 | 1 | 0 | 0 | 0 | 0 | 0 | 0 | 0 | 1 | 0 |
| 51.40816 | 21.14103 | MAZ_U_21 | mazu21_5 | Poland | U | 0 | 0 | 0 | 0 | 0 | 0 | 1 | 150,552 | 1298,514467 | 127 | 28040,697 | 2300 | 0 | 1 | 0 | 0 | 1 | 0 | 1 | 0 | 0 | 0 | 0 | 0 | 0 | 0 | 1 | 0 |
| 51.40587 | 21.13938 | MAZ_U_21 | mazu21_6 | Poland | U | 0 | 0 | 0 | 0 | 0 | 0 | 0 | 151,101 | 1298,354593 | 127 | 29904,383 | 6330 | 1 | 1 | 0 | 0 | 1 | 0 | 1 | 1 | 1 | 0 | 1 | 1 | 0 | 0 | 0 | 0 |
| 51.40484 | 21.13486 | MAZ_U_21 | mazu21_7 | Poland | U | 0 | 0 | 0 | 0 | 0 | 0 | 1 | 152,714 | 1298,421068 | 127 | 17172,132 | 15288 | 0 | 1 | 0 | 0 | 0 | 0 | 1 | 1 | 1 | 0 | 0 | 1 | 0 | 0 | 1 | 0 |
| 51.40259 | 21.13154 | MAZ_U_21 | mazu21_8 | Poland | U | 0 | 0 | 0 | 0 | 0 | 0 | 1 | 151,632 | 1298,328773 | 127 | 32407,762 | 2666 | 1 | 1 | 0 | 1 | 0 | 0 | 1 | 0 | 1 | 0 | 0 | 1 | 0 | 0 | 0 | 0 |
| 52.27344 | 20.97158 | MAZ_U_29 | mazu29_1 | Poland | U | 0 | 0 | 0 | 0 | 0 | 0 | 0 | 91,511 | 1387,455962 | 29 | 42393,471 | 4412 | 1 | 1 | 0 | 0 | 1 | 0 | 1 | 1 | 0 | 0 | 0 | 0 | 1 | 0 | 0 | 0 |
| 52.2758 | 20.9687 | MAZ_U_29 | mazu29_2 | Poland | U | 0 | 0 | 0 | 0 | 0 | 0 | 0 | 98 | 1387,782402 | 29 | 24709,83 | 11538 | 1 | 1 | 0 | 1 | 1 | 0 | 0 | 1 | 0 | 0 | 0 | 0 | 1 | 1 | 0 | 0 |
| 52.27816 | 20.96918 | MAZ_U_29 | mazu29_3 | Poland | U | 0 | 0 | 1 | 1 | 0 | 0 | 0 | 87 | 1387,993198 | 29 | 26007,325 | 8959 | 1 | 1 | 0 | 0 | 1 | 0 | 0 | 1 | 0 | 0 | 0 | 0 | 1 | 1 | 0 | 0 |
| 52.27917 | 20.97289 | MAZ_U_29 | mazu29_4 | Poland | U | 0 | 0 | 0 | 0 | 0 | 0 | 0 | 84,012 | 1387,965871 | 29 | 9096,889 | 13717 | 0 | 1 | 0 | 0 | 0 | 0 | 1 | 1 | 0 | 0 | 0 | 0 | 1 | 0 | 1 | 0 |
| 52.28108 | 20.97291 | MAZ_U_29 | mazu29_5 | Poland | U | 0 | 0 | 0 | 0 | 0 | 0 | 0 | 83,423 | 1388,149813 | 29 | 16063,677 | 10219 | 0 | 1 | 0 | 0 | 0 | 0 | 1 | 1 | 1 | 0 | 0 | 1 | 1 | 0 | 1 | 0 |
| 52.28284 | 20.97518 | MAZ_U_29 | mazu29_6 | Poland | U | 0 | 0 | 0 | 0 | 0 | 0 | 0 | 83 | 1388,243207 | 29 | 18553,382 | 9897 | 1 | 1 | 0 | 0 | 1 | 0 | 1 | 1 | 0 | 0 | 0 | 0 | 1 | 0 | 0 | 0 |
| 52.28512 | 20.97686 | MAZ_U_29 | mazu29_7 | Poland | U | 0 | 0 | 0 | 0 | 0 | 0 | 0 | 81,173 | 1388,406967 | 29 | 29792,055 | 10776 | 1 | 1 | 0 | 0 | 1 | 0 | 0 | 1 | 0 | 0 | 0 | 0 | 1 | 0 | 0 | 0 |
| 52.28775 | 20.97806 | MAZ_U_29 | mazu29_8 | Poland | U | 0 | 0 | 0 | 0 | 0 | 0 | 0 | 79,029 | 1388,621041 | 29 | 35465,306 | 4883 | 1 | 1 | 0 | 0 | 1 | 0 | 1 | 1 | 1 | 1 | 0 | 1 | 1 | 0 | 0 | 0 |
| 52.22367 | 20.92364 | MAZ_U_3 | mazu3_1 | Poland | U | 0 | 0 | 0 | 0 | 0 | 0 | 0 | 113,372 | 1384,277716 | 34 | 30842,914 | 7830 | 1 | 1 | 0 | 1 | 0 | 0 | 1 | 1 | 0 | 0 | 0 | 0 | 0 | 0 | 1 | 0 |
| 52.22612 | 20.92171 | MAZ_U_3 | mazu3_2 | Poland | U | 0 | 0 | 2 | 1 | 0 | 0 | 1 | 114 | 1384,580252 | 34 | 54841,452 | 2300 | 1 | 1 | 0 | 1 | 0 | 0 | 1 | 0 | 0 | 0 | 0 | 0 | 0 | 0 | 0 | 0 |
| 52.22771 | 20.91937 | MAZ_U_3 | mazu3_3 | Poland | U | 0 | 0 | 0 | 0 | 0 | 0 | 1 | 113 | 1384,812653 | 34 | 50312,317 | 5404 | 1 | 1 | 0 | 1 | 0 | 0 | 1 | 1 | 0 | 0 | 0 | 0 | 1 | 1 | 0 | 0 |
| 52.22969 | 20.92028 | MAZ_U_3 | mazu3_4 | Poland | U | 0 | 0 | 0 | 0 | 0 | 0 | 0 | 114 | 1384,973676 | 34 | 24930,791 | 9169 | 1 | 1 | 0 | 1 | 0 | 0 | 1 | 1 | 1 | 0 | 0 | 1 | 1 | 0 | 0 | 0 |
| 52.23228 | 20.92158 | MAZ_U_3 | mazu3_5 | Poland | U | 0 | 0 | 0 | 0 | 0 | 0 | 1 | 112,857 | 1385,177588 | 34 | 7324,052 | 14271 | 0 | 1 | 0 | 0 | 0 | 0 | 1 | 1 | 1 | 0 | 0 | 1 | 0 | 0 | 1 | 0 |
| 52.23504 | 20.92082 | MAZ_U_3 | mazu3_6 | Poland | U | 0 | 0 | 0 | 0 | 0 | 0 | 1 | 111 | 1385,471291 | 34 | 17907,832 | 11063 | 1 | 1 | 0 | 0 | 0 | 0 | 1 | 1 | 1 | 0 | 0 | 1 | 1 | 0 | 0 | 0 |
| 52.23691 | 20.91983 | MAZ_U_3 | mazu3_7 | Poland | U | 0 | 0 | 0 | 0 | 0 | 0 | 1 | 111 | 1385,685139 | 34 | 26862,816 | 14882 | 1 | 1 | 0 | 0 | 0 | 0 | 1 | 1 | 1 | 0 | 0 | 1 | 1 | 0 | 0 | 0 |
| 52.23941 | 20.91955 | MAZ_U_3 | mazu3_8 | Poland | U | 0 | 0 | 0 | 0 | 0 | 0 | 1 | 111,923 | 1385,935965 | 34 | 14422,678 | 17291 | 0 | 1 | 0 | 0 | 0 | 0 | 1 | 1 | 0 | 0 | 0 | 0 | 1 | 0 | 1 | 0 |
| 52.26109 | 20.96445 | MAZ_U_30 | mazu30_1 | Poland | U | 0 | 0 | 0 | 0 | 0 | 0 | 0 | 106 | 1386,504748 | 30 | 16740,812 | 12289 | 1 | 1 | 0 | 0 | 1 | 0 | 1 | 1 | 0 | 0 | 0 | 0 | 1 | 1 | 0 | 0 |
| 52.26287 | 20.96676 | MAZ_U_30 | mazu30_2 | Poland | U | 2 | 1 | 0 | 0 | 0 | 0 | 0 | 105,78 | 1386,597562 | 30 | 23820,901 | 16229 | 1 | 1 | 1 | 0 | 1 | 0 | 1 | 1 | 0 | 0 | 0 | 0 | 1 | 1 | 0 | 0 |
| 52.26471 | 20.96855 | MAZ_U_30 | mazu30_3 | Poland | U | 0 | 0 | 0 | 0 | 0 | 0 | 0 | 105,21 | 1386,715147 | 30 | 16192,746 | 12440 | 0 | 1 | 0 | 0 | 0 | 0 | 1 | 1 | 0 | 0 | 0 | 0 | 1 | 0 | 0 | 0 |
| 52.26688 | 20.97107 | MAZ_U_30 | mazu30_4 | Poland | U | 2 | 1 | 2 | 1 | 0 | 0 | 0 | 102,346 | 1386,839034 | 30 | 41822,932 | 8194 | 1 | 1 | 0 | 0 | 1 | 0 | 0 | 1 | 0 | 0 | 0 | 0 | 1 | 0 | 0 | 0 |
| 52.26843 | 20.97532 | MAZ_U_30 | mazu30_5 | Poland | U | 0 | 0 | 0 | 0 | 0 | 0 | 0 | 99,652 | 1386,845085 | 30 | 33503,689 | 12557 | 1 | 1 | 0 | 0 | 1 | 0 | 1 | 1 | 0 | 0 | 0 | 0 | 1 | 0 | 0 | 0 |
| 52.26982 | 20.97925 | MAZ_U_30 | mazu30_6 | Poland | U | 0 | 0 | 0 | 0 | 0 | 0 | 0 | 97 | 1386,846858 | 30 | 17854,108 | 25296 | 0 | 1 | 0 | 0 | 1 | 0 | 1 | 1 | 0 | 0 | 0 | 0 | 1 | 1 | 0 | 0 |
| 52.27203 | 20.98047 | MAZ_U_30 | mazu30_7 | Poland | U | 0 | 0 | 1 | 1 | 0 | 0 | 0 | 95,171 | 1387,019451 | 30 | 31570,36 | 14506 | 1 | 1 | 1 | 0 | 1 | 0 | 1 | 1 | 0 | 0 | 0 | 0 | 1 | 1 | 0 | 0 |
| 52.27289 | 20.98525 | MAZ_U_30 | mazu30_8 | Poland | U | 0 | 0 | 0 | 0 | 0 | 0 | 0 | 85,636 | 1386,941079 | 30 | 34825,918 | 17429 | 0 | 1 | 0 | 0 | 0 | 0 | 1 | 1 | 1 | 0 | 0 | 0 | 1 | 0 | 0 | 0 |
| 52.2611 | 20.9988 | MAZ_U_31 | mazu31_1 | Poland | U | 0 | 0 | 0 | 0 | 0 | 0 | 0 | 96,694 | 1385,34329 | 30 | 27144,005 | 12094 | 1 | 1 | 0 | 1 | 0 | 0 | 0 | 1 | 0 | 0 | 0 | 0 | 1 | 0 | 0 | 0 |
| 52.26338 | 20.99617 | MAZ_U_31 | mazu31_2 | Poland | U | 1 | 1 | 0 | 0 | 0 | 0 | 0 | 93,62 | 1385,652866 | 30 | 30809,441 | 8403 | 1 | 1 | 0 | 1 | 0 | 0 | 0 | 1 | 0 | 0 | 0 | 0 | 1 | 1 | 0 | 0 |
| 52.26596 | 20.99513 | MAZ_U_31 | mazu31_3 | Poland | U | 0 | 0 | 0 | 0 | 0 | 0 | 0 | 89,225 | 1385,937213 | 30 | 32520,915 | 8389 | 1 | 1 | 0 | 1 | 1 | 0 | 0 | 1 | 0 | 0 | 0 | 0 | 1 | 0 | 0 | 0 |
| 52.2688 | 20.99727 | MAZ_U_31 | mazu31_4 | Poland | U | 0 | 0 | 0 | 0 | 0 | 0 | 0 | 91,383 | 1386,139917 | 30 | 29640,68 | 9782 | 1 | 1 | 1 | 1 | 1 | 0 | 0 | 1 | 0 | 1 | 0 | 0 | 1 | 1 | 1 | 0 |
| 52.27084 | 20.99611 | MAZ_U_31 | mazu31_5 | Poland | U | 0 | 0 | 0 | 0 | 0 | 0 | 0 | 85,98 | 1386,376273 | 30 | 19739,243 | 7601 | 0 | 1 | 0 | 0 | 1 | 0 | 1 | 1 | 0 | 0 | 0 | 0 | 0 | 1 | 1 | 0 |
| 52.27261 | 20.99502 | MAZ_U_31 | mazu31_6 | Poland | U | 0 | 0 | 0 | 0 | 0 | 0 | 0 | 88,971 | 1386,584285 | 30 | 12786,797 | 4183 | 0 | 1 | 0 | 0 | 1 | 0 | 1 | 1 | 0 | 0 | 0 | 0 | 1 | 0 | 1 | 0 |
| 52.27507 | 20.99431 | MAZ_U_31 | mazu31_7 | Poland | U | 0 | 0 | 0 | 0 | 0 | 0 | 0 | 83,174 | 1386,846391 | 30 | 13979,741 | 2161 | 0 | 1 | 0 | 0 | 1 | 0 | 1 | 0 | 0 | 1 | 0 | 0 | 0 | 0 | 1 | 0 |
| 52.27743 | 20.99258 | MAZ_U_31 | mazu31_8 | Poland | U | 0 | 0 | 0 | 0 | 0 | 0 | 0 | 82,251 | 1387,133107 | 30 | 36154,907 | 2100 | 1 | 1 | 0 | 0 | 1 | 0 | 1 | 0 | 0 | 1 | 1 | 0 | 0 | 0 | 1 | 0 |
| 52.19698 | 20.98077 | MAZ_U_32 | mazu32_1 | Poland | U | 0 | 0 | 2 | 1 | 0 | 0 | 0 | 109,128 | 1379,763043 | 37 | 24424,807 | 17385 | 0 | 1 | 0 | 0 | 0 | 0 | 1 | 1 | 0 | 0 | 0 | 0 | 1 | 0 | 0 | 0 |
| 52.19784 | 20.97765 | MAZ_U_32 | mazu32_2 | Poland | U | 0 | 0 | 0 | 0 | 0 | 0 | 0 | 109 | 1379,950859 | 37 | 15261,619 | 19485 | 0 | 1 | 0 | 0 | 0 | 0 | 0 | 1 | 0 | 0 | 0 | 0 | 1 | 1 | 0 | 0 |
| 52.19794 | 20.97371 | MAZ_U_32 | mazu32_3 | Poland | U | 2 | 1 | 0 | 0 | 0 | 0 | 1 | 108,356 | 1380,098737 | 37 | 33626,301 | 7935 | 0 | 1 | 0 | 0 | 0 | 0 | 0 | 1 | 1 | 0 | 1 | 0 | 1 | 0 | 0 | 0 |
| 52.19882 | 20.97002 | MAZ_U_32 | mazu32_4 | Poland | U | 0 | 0 | 0 | 0 | 0 | 0 | 1 | 107 | 1380,301997 | 37 | 50513,231 | 3729 | 0 | 1 | 0 | 0 | 0 | 0 | 0 | 1 | 1 | 0 | 1 | 0 | 1 | 0 | 0 | 0 |
| 52.1999 | 20.96566 | MAZ_U_32 | mazu32_5 | Poland | U | 0 | 0 | 0 | 0 | 0 | 0 | 1 | 108 | 1380,553669 | 37 | 13462,152 | 31067 | 0 | 1 | 0 | 0 | 0 | 0 | 1 | 1 | 1 | 0 | 0 | 0 | 0 | 1 | 0 | 0 |
| 52.20224 | 20.96408 | MAZ_U_32 | mazu32_6 | Poland | U | 1 | 1 | 0 | 0 | 0 | 0 | 0 | 107 | 1380,833456 | 37 | 19437,364 | 15483 | 0 | 1 | 0 | 0 | 1 | 0 | 0 | 1 | 0 | 0 | 0 | 1 | 1 | 1 | 0 | 0 |

| | | | | | | | | | | | | | | | | | | | | | | | | | | | | | | | | |
|---|---|---|---|---|---|---|---|---|---|---|---|---|---|---|---|---|---|---|---|---|---|---|---|---|---|---|---|---|---|---|---|---|
| 52.20505 | 20.96325 | MAZ_U_32 | mazu32_7 | Poland | U | 0 | 0 | 0 | 0 | 0 | 0 | 1 | 109 | 1381,13289 | 37 | 31804,961 | 4239 | 1 | 1 | 0 | 0 | 1 | 0 | 1 | 1 | 0 | 1 | 0 | 0 | 1 | 0 | 0 | 0 |
| 52.20751 | 20.96255 | MAZ_U_32 | mazu32_8 | Poland | U | 0 | 0 | 0 | 0 | 0 | 0 | 0 | 108 | 1381,39379 | 37 | 22124,739 | 3306 | 0 | 1 | 0 | 0 | 1 | 0 | 0 | 1 | 0 | 1 | 0 | 0 | 1 | 0 | 0 | 0 |
| 52.24539 | 20.92624 | MAZ_U_4 | mazu4_1 | Poland | U | 0 | 0 | 0 | 0 | 0 | 0 | 0 | 110,681 | 1386,284991 | 33 | 26595,779 | 2087 | 0 | 1 | 0 | 0 | 0 | 0 | 1 | 1 | 1 | 0 | 0 | 1 | 1 | 0 | 0 | 0 |
| 52.24405 | 20.92989 | MAZ_U_4 | mazu4_2 | Poland | U | 0 | 0 | 0 | 0 | 0 | 0 | 1 | 111,166 | 1386,030954 | 33 | 9526,385 | 8045 | 0 | 1 | 0 | 0 | 0 | 0 | 1 | 0 | 1 | 0 | 0 | 1 | 0 | 0 | 1 | 0 |
| 52.24308 | 20.93377 | MAZ_U_4 | mazu4_3 | Poland | U | 0 | 0 | 0 | 0 | 0 | 0 | 1 | 112 | 1385,805502 | 33 | 11619,916 | 33003 | 0 | 1 | 0 | 0 | 0 | 0 | 1 | 0 | 1 | 0 | 0 | 1 | 0 | 0 | 1 | 0 |
| 52.24215 | 20.93795 | MAZ_U_4 | mazu4_4 | Poland | U | 0 | 0 | 0 | 0 | 0 | 0 | 1 | 112,839 | 1385,574181 | 33 | 10132,888 | 10858 | 0 | 1 | 0 | 0 | 0 | 0 | 1 | 1 | 1 | 0 | 0 | 1 | 1 | 1 | 0 | 0 |
| 52.24295 | 20.94223 | MAZ_U_4 | mazu4_5 | Poland | U | 0 | 0 | 0 | 0 | 0 | 0 | 1 | 112 | 1385,506924 | 33 | 13426,945 | 19407 | 0 | 1 | 0 | 0 | 0 | 0 | 1 | 1 | 0 | 0 | 0 | 1 | 1 | 0 | 0 | 0 |
| 52.24466 | 20.94382 | MAZ_U_4 | mazu4_6 | Poland | U | 0 | 0 | 0 | 0 | 0 | 0 | 1 | 111 | 1385,61475 | 33 | 9322,284 | 15799 | 0 | 0 | 0 | 0 | 0 | 0 | 1 | 1 | 0 | 0 | 0 | 1 | 1 | 0 | 0 | 0 |
| 52.24604 | 20.94519 | MAZ_U_4 | mazu4_7 | Poland | U | 0 | 0 | 0 | 0 | 0 | 0 | 1 | 109,666 | 1385,702088 | 33 | 12878,553 | 17878 | 0 | 0 | 0 | 0 | 0 | 0 | 1 | 1 | 0 | 0 | 0 | 1 | 1 | 0 | 0 | 0 |
| 52.2463 | 20.9497 | MAZ_U_4 | mazu4_8 | Poland | U | 0 | 0 | 0 | 0 | 0 | 0 | 1 | 109,32 | 1385,575299 | 33 | 10184,793 | 17287 | 0 | 0 | 0 | 0 | 0 | 0 | 1 | 1 | 1 | 0 | 0 | 0 | 1 | 0 | 0 | 0 |
| 52.20649 | 20.86154 | MAZ_U_49 | mazu49_1 | Poland | U | 0 | 0 | 0 | 0 | 0 | 0 | 1 | 108 | 1384,749041 | 37 | 12020,775 | 8722 | 0 | 1 | 1 | 0 | 0 | 0 | 0 | 0 | 1 | 0 | 0 | 1 | 0 | 0 | 0 | 0 |
| 52.20445 | 20.86082 | MAZ_U_49 | mazu49_2 | Poland | U | 1 | 1 | 0 | 0 | 0 | 0 | 0 | 107 | 1384,578409 | 37 | 16312,184 | 9229 | 0 | 1 | 1 | 0 | 0 | 0 | 0 | 0 | 1 | 0 | 0 | 1 | 1 | 0 | 0 | 0 |
| 52.20193 | 20.8615 | MAZ_U_49 | mazu49_3 | Poland | U | 0 | 0 | 0 | 0 | 0 | 0 | 1 | 106,948 | 1384,311325 | 37 | 10748,395 | 11345 | 0 | 1 | 1 | 0 | 0 | 0 | 0 | 0 | 1 | 0 | 0 | 1 | 1 | 0 | 0 | 0 |
| 52.19932 | 20.86036 | MAZ_U_49 | mazu49_4 | Poland | U | 0 | 0 | 0 | 0 | 0 | 0 | 1 | 105 | 1384,100203 | 37 | 13644,267 | 9765 | 0 | 1 | 1 | 0 | 0 | 0 | 1 | 0 | 1 | 0 | 0 | 1 | 1 | 0 | 0 | 0 |
| 52.19728 | 20.86094 | MAZ_U_49 | mazu49_5 | Poland | U | 0 | 0 | 0 | 0 | 0 | 0 | 1 | 103,208 | 1383,885248 | 37 | 4311,547 | 22515 | 0 | 1 | 1 | 0 | 0 | 0 | 1 | 0 | 0 | 0 | 0 | 1 | 0 | 1 | 0 | 0 |
| 52.19581 | 20.86205 | MAZ_U_49 | mazu49_6 | Poland | U | 0 | 0 | 0 | 0 | 0 | 0 | 1 | 104,432 | 1383,70608 | 37 | 7289,453 | 24985 | 0 | 1 | 1 | 0 | 0 | 0 | 1 | 1 | 0 | 0 | 0 | 0 | 1 | 0 | 0 | 0 |
| 52.193 | 20.86269 | MAZ_U_49 | mazu49_7 | Poland | U | 1 | 1 | 0 | 0 | 0 | 0 | 1 | 104,937 | 1383,413926 | 37 | 18439,834 | 13510 | 0 | 1 | 1 | 0 | 0 | 0 | 1 | 1 | 0 | 0 | 0 | 0 | 1 | 1 | 0 | 0 |
| 52.19084 | 20.86499 | MAZ_U_49 | mazu49_8 | Poland | U | 0 | 0 | 0 | 0 | 0 | 0 | 1 | 105,035 | 1383,126323 | 37 | 13164,019 | 14060 | 0 | 1 | 1 | 0 | 0 | 0 | 1 | 1 | 1 | 0 | 0 | 1 | 1 | 0 | 1 | 0 |
| 52.24006 | 20.95144 | MAZ_U_5 | mazu5_1 | Poland | U | 0 | 0 | 0 | 0 | 0 | 0 | 1 | 110 | 1384,91418 | 34 | 32580,269 | 230 | 0 | 1 | 0 | 0 | 1 | 0 | 0 | 0 | 0 | 0 | 0 | 0 | 0 | 0 | 0 | 0 |
| 52.2375 | 20.95039 | MAZ_U_5 | mazu5_2 | Poland | U | 2 | 1 | 0 | 0 | 0 | 0 | 1 | 111 | 1384,702079 | 34 | 20254,644 | 9858 | 0 | 1 | 0 | 0 | 0 | 0 | 1 | 1 | 0 | 0 | 0 | 0 | 1 | 0 | 0 | 0 |
| 52.23474 | 20.94996 | MAZ_U_5 | mazu5_3 | Poland | U | 0 | 0 | 0 | 0 | 0 | 0 | 1 | 112,936 | 1384,450382 | 34 | 22752,473 | 2384 | 0 | 1 | 0 | 0 | 1 | 0 | 1 | 1 | 0 | 0 | 0 | 0 | 1 | 0 | 0 | 0 |
| 52.23208 | 20.95041 | MAZ_U_5 | mazu5_4 | Poland | U | 0 | 0 | 0 | 0 | 0 | 0 | 1 | 111,512 | 1384,177882 | 34 | 26017,739 | 3800 | 0 | 1 | 0 | 0 | 1 | 0 | 1 | 0 | 0 | 0 | 0 | 0 | 0 | 0 | 0 | 0 |
| 52.22928 | 20.94955 | MAZ_U_5 | mazu5_5 | Poland | U | 0 | 0 | 0 | 0 | 0 | 0 | 1 | 112,253 | 1383,938636 | 34 | 32357,832 | 15480 | 1 | 1 | 0 | 0 | 1 | 0 | 0 | 1 | 0 | 0 | 0 | 0 | 1 | 0 | 1 | 0 |
| 52.22798 | 20.94674 | MAZ_U_5 | mazu5_6 | Poland | U | 0 | 0 | 0 | 0 | 0 | 0 | 0 | 113,728 | 1383,908579 | 34 | 29151,671 | 9849 | 1 | 1 | 0 | 0 | 0 | 1 | 0 | 1 | 0 | 0 | 0 | 0 | 0 | 0 | 1 | 0 |
| 52.22714 | 20.94316 | MAZ_U_5 | mazu5_7 | Poland | U | 0 | 0 | 0 | 0 | 0 | 0 | 1 | 115,255 | 1383,948958 | 34 | 25254,957 | 12922 | 1 | 1 | 0 | 0 | 0 | 1 | 0 | 1 | 0 | 0 | 0 | 0 | 1 | 0 | 1 | 0 |
| 52.22725 | 20.93814 | MAZ_U_5 | mazu5_8 | Poland | U | 0 | 0 | 0 | 0 | 0 | 0 | 1 | 115 | 1384,129398 | 34 | 38254,906 | 870 | 1 | 1 | 0 | 0 | 1 | 1 | 1 | 1 | 0 | 0 | 0 | 0 | 0 | 0 | 1 | 0 |
| 52.1996 | 20.89224 | MAZ_U_50 | mazu50_1 | Poland | U | 0 | 0 | 0 | 0 | 0 | 0 | 1 | 108,44 | 1383,032934 | 37 | 4767,114 | 14979 | 0 | 1 | 1 | 0 | 0 | 0 | 1 | 1 | 1 | 0 | 0 | 1 | 1 | 0 | 1 | 0 |
| 52.1985 | 20.88887 | MAZ_U_50 | mazu50_2 | Poland | U | 0 | 0 | 0 | 0 | 0 | 0 | 1 | 108,373 | 1383,042884 | 37 | 9759,222 | 17196 | 0 | 1 | 0 | 0 | 0 | 0 | 1 | 0 | 1 | 0 | 0 | 1 | 1 | 0 | 1 | 0 |
| 52.198 | 20.88535 | MAZ_U_50 | mazu50_3 | Poland | U | 0 | 0 | 0 | 0 | 0 | 0 | 1 | 108 | 1383,116433 | 37 | 3565,866 | 14666 | 0 | 1 | 0 | 0 | 0 | 0 | 1 | 0 | 0 | 0 | 0 | 0 | 1 | 1 | 1 | 0 |
| 52.19935 | 20.88258 | MAZ_U_50 | mazu50_4 | Poland | U | 0 | 0 | 0 | 0 | 0 | 0 | 0 | 108,098 | 1383,342382 | 37 | 11147,97 | 7463 | 0 | 1 | 0 | 0 | 0 | 0 | 0 | 1 | 0 | 0 | 0 | 0 | 1 | 1 | 0 | 0 |
| 52.19865 | 20.87934 | MAZ_U_50 | mazu50_5 | Poland | U | 0 | 0 | 0 | 0 | 0 | 0 | 1 | 107,087 | 1383,384801 | 37 | 9456,267 | 8551 | 0 | 1 | 0 | 0 | 0 | 0 | 0 | 1 | 1 | 0 | 0 | 1 | 1 | 0 | 0 | 0 |
| 52.2005 | 20.87582 | MAZ_U_50 | mazu50_6 | Poland | U | 0 | 0 | 1 | 1 | 0 | 0 | 1 | 108 | 1383,683818 | 37 | 15717,461 | 4168 | 0 | 1 | 0 | 0 | 0 | 0 | 1 | 1 | 0 | 0 | 0 | 0 | 1 | 0 | 1 | 0 |
| 52.20255 | 20.8733 | MAZ_U_50 | mazu50_7 | Poland | U | 0 | 0 | 0 | 0 | 0 | 0 | 1 | 108 | 1383,966735 | 37 | 17087,021 | 14425 | 0 | 1 | 0 | 0 | 1 | 0 | 0 | 0 | 0 | 0 | 1 | 0 | 0 | 0 | 1 | 0 |
| 52.20502 | 20.87117 | MAZ_U_50 | mazu50_8 | Poland | U | 0 | 0 | 0 | 0 | 0 | 0 | 1 | 108 | 1384,278218 | 37 | 19754,617 | 28183 | 0 | 1 | 0 | 0 | 1 | 0 | 0 | 1 | 1 | 0 | 0 | 1 | 0 | 0 | 1 | 0 |
| 52.19297 | 20.90607 | MAZ_U_51 | mazu51_1 | Poland | U | 0 | 0 | 0 | 0 | 0 | 0 | 1 | 111 | 1381,751521 | 38 | 17389,574 | 18538 | 1 | 1 | 0 | 0 | 0 | 0 | 1 | 1 | 0 | 0 | 1 | 0 | 0 | 1 | 0 | 0 |
| 52.1941 | 20.90233 | MAZ_U_51 | mazu51_2 | Poland | U | 0 | 0 | 0 | 0 | 0 | 0 | 1 | 110,705 | 1381,920559 | 38 | 28042,457 | 16757 | 1 | 1 | 0 | 0 | 0 | 0 | 1 | 1 | 0 | 0 | 1 | 0 | 1 | 0 | 0 | 0 |
| 52.19521 | 20.89899 | MAZ_U_51 | mazu51_3 | Poland | U | 0 | 0 | 0 | 0 | 0 | 0 | 1 | 110 | 1382,052416 | 38 | 19723,689 | 12680 | 0 | 1 | 0 | 0 | 0 | 0 | 1 | 1 | 0 | 0 | 0 | 1 | 1 | 0 | 0 | 0 |
| 52.19515 | 20.89628 | MAZ_U_51 | mazu51_4 | Poland | U | 0 | 0 | 0 | 0 | 0 | 0 | 1 | 109,788 | 1382,15764 | 38 | 29045,148 | 12272 | 1 | 1 | 0 | 0 | 1 | 0 | 1 | 1 | 0 | 0 | 0 | 0 | 1 | 1 | 0 | 0 |
| 52.19313 | 20.89504 | MAZ_U_51 | mazu51_5 | Poland | U | 0 | 0 | 0 | 0 | 0 | 0 | 1 | 109 | 1382,351082 | 38 | 15088,585 | 14147 | 0 | 1 | 0 | 0 | 0 | 0 | 1 | 1 | 0 | 0 | 0 | 0 | 1 | 1 | 0 | 0 |
| 52.19099 | 20.89309 | MAZ_U_51 | mazu51_6 | Poland | U | 0 | 0 | 1 | 1 | 0 | 0 | 1 | 108 | 1382,377315 | 38 | 20507,068 | 16307 | 0 | 1 | 0 | 0 | 0 | 0 | 1 | 1 | 0 | 0 | 0 | 0 | 1 | 1 | 0 | 0 |
| 52.19013 | 20.88869 | MAZ_U_51 | mazu51_7 | Poland | U | 0 | 0 | 0 | 0 | 0 | 0 | 1 | 108 | 1382,607212 | 38 | 11614,391 | 16270 | 0 | 1 | 0 | 0 | 0 | 0 | 1 | 1 | 1 | 0 | 0 | 1 | 1 | 1 | 0 | 0 |
| 52.19205 | 20.88504 | MAZ_U_51 | mazu51_8 | Poland | U | 1 | 1 | 0 | 0 | 0 | 0 | 1 | 107,144 | 1382,462358 | 38 | 30800,091 | 5621 | 1 | 1 | 0 | 0 | 1 | 0 | 0 | 1 | 0 | 0 | 0 | 1 | 1 | 0 | 0 | 0 |
| 52.23864 | 21.04088 | MAZ_U_55 | mazu55_1 | Poland | U | 2 | 1 | 0 | 0 | 0 | 0 | 1 | 80,168 | 1382,873607 | 33 | 31928,769 | 2100 | 1 | 1 | 0 | 0 | 1 | 0 | 0 | 0 | 0 | 0 | 0 | 0 | 0 | 0 | 0 | 0 |
| 52.24089 | 21.03841 | MAZ_U_55 | mazu55_2 | Poland | U | 0 | 0 | 0 | 0 | 0 | 0 | 0 | 80,424 | 1382,31269 | 33 | 33586,181 | 4300 | 1 | 1 | 0 | 0 | 1 | 0 | 0 | 0 | 0 | 1 | 0 | 0 | 0 | 0 | 1 | 0 |
| 52.24299 | 21.03558 | MAZ_U_55 | mazu55_3 | Poland | U | 0 | 0 | 0 | 0 | 0 | 0 | 0 | 80,764 | 1383,106275 | 33 | 31319,409 | 2000 | 1 | 1 | 0 | 0 | 1 | 0 | 0 | 0 | 0 | 1 | 0 | 0 | 0 | 0 | 1 | 0 |
| 52.24477 | 21.03312 | MAZ_U_55 | mazu55_4 | Poland | U | 0 | 0 | 0 | 0 | 0 | 0 | 0 | 80,364 | 1382,17319 | 33 | 38188,341 | 2000 | 1 | 1 | 0 | 0 | 1 | 0 | 0 | 0 | 0 | 1 | 0 | 0 | 0 | 0 | 0 | 0 |
| 52.24649 | 21.03008 | MAZ_U_55 | mazu55_5 | Poland | U | 0 | 0 | 0 | 0 | 0 | 0 | 0 | 80,094 | 1383,339562 | 33 | 35808,024 | 4100 | 1 | 1 | 0 | 1 | 1 | 0 | 0 | 1 | 0 | 1 | 0 | 0 | 1 | 0 | 0 | 0 |
| 52.24825 | 21.02824 | MAZ_U_55 | mazu55_6 | Poland | U | 0 | 0 | 0 | 0 | 0 | 0 | 0 | 83,999 | 1382,240636 | 33 | 21856,961 | 10532 | 1 | 1 | 0 | 0 | 1 | 0 | 0 | 1 | 0 | 1 | 0 | 0 | 1 | 0 | 0 | 0 |
| 52.24951 | 21.02496 | MAZ_U_55 | mazu55_7 | Poland | U | 0 | 0 | 0 | 0 | 0 | 0 | 1 | 80,092 | 1383,622877 | 33 | 33497,919 | 7087 | 1 | 1 | 0 | 1 | 1 | 0 | 1 | 1 | 0 | 1 | 0 | 0 | 1 | 0 | 1 | 0 |
| 52.25165 | 21.0227 | MAZ_U_55 | mazu55_8 | Poland | U | 0 | 0 | 0 | 0 | 0 | 0 | 1 | 81,643 | 1382,553687 | 33 | 51220,842 | 2000 | 1 | 1 | 0 | 1 | 1 | 0 | 0 | 1 | 0 | 1 | 1 | 0 | 0 | 0 | 0 | 0 |
| 52.2576 | 20.90062 | MAZ_U_6 | mazu6_1 | Poland | U | 0 | 0 | 0 | 0 | 0 | 0 | 1 | 108,508 | 1388,334814 | 31 | 24092,686 | 8252 | 1 | 1 | 0 | 0 | 1 | 0 | 1 | 1 | 1 | 0 | 0 | 0 | 1 | 1 | 0 | 0 |
| 52.25608 | 20.90379 | MAZ_U_6 | mazu6_2 | Poland | U | 0 | 0 | 0 | 0 | 0 | 0 | 1 | 108,072 | 1388,080384 | 31 | 16624,593 | 11409 | 0 | 1 | 0 | 0 | 0 | 0 | 1 | 1 | 0 | 0 | 0 | 0 | 1 | 1 | 0 | 0 |
| 52.2547 | 20.90793 | MAZ_U_6 | mazu6_3 | Poland | U | 0 | 0 | 0 | 0 | 0 | 0 | 1 | 109 | 1387,80421 | 31 | 25594,531 | 7429 | 0 | 1 | 0 | 0 | 0 | 0 | 1 | 1 | 0 | 0 | 0 | 0 | 1 | 0 | 0 | 0 |
| 52.25344 | 20.91172 | MAZ_U_6 | mazu6_4 | Poland | U | 0 | 0 | 0 | 0 | 0 | 0 | 1 | 109 | 1387,555729 | 31 | 12976,775 | 14137 | 0 | 1 | 0 | 0 | 0 | 0 | 1 | 1 | 1 | 0 | 0 | 1 | 1 | 0 | 0 | 0 |
| 52.2551 | 20.91498 | MAZ_U_6 | mazu6_5 | Poland | U | 0 | 0 | 0 | 0 | 0 | 0 | 1 | 108 | 1387,604939 | 31 | 21848,217 | 14598 | 1 | 1 | 0 | 0 | 0 | 0 | 0 | 1 | 0 | 0 | 0 | 0 | 1 | 1 | 0 | 0 |
| 52.25726 | 20.91685 | MAZ_U_6 | mazu6_6 | Poland | U | 0 | 0 | 0 | 0 | 0 | 0 | 1 | 107,954 | 1387,749402 | 31 | 14710,974 | 14301 | 0 | 1 | 0 | 0 | 0 | 0 | 0 | 1 | 1 | 0 | 0 | 0 | 1 | 1 | 0 | 0 |
| 52.25973 | 20.91799 | MAZ_U_6 | mazu6_7 | Poland | U | 0 | 0 | 0 | 0 | 0 | 0 | 1 | 107,257 | 1387,948832 | 31 | 5787,141 | 13517 | 0 | 1 | 0 | 0 | 0 | 0 | 0 | 1 | 1 | 0 | 1 | 0 | 0 | 1 | 0 | 0 |
| 52.26182 | 20.91945 | MAZ_U_6 | mazu6_8 | Poland | U | 0 | 0 | 0 | 0 | 0 | 0 | 1 | 107,552 | 1388,100984 | 31 | 6703,649 | 12282 | 0 | 1 | 0 | 0 | 0 | 0 | 0 | 1 | 0 | 0 | 0 | 0 | 1 | 0 | 0 | 0 |
| 52.25395 | 20.92929 | MAZ_U_7 | mazu7_1 | Poland | U | 0 | 0 | 0 | 0 | 0 | 0 | 1 | 108,566 | 1387,006006 | 31 | 16956,41 | 5701 | 1 | 1 | 1 | 1 | 0 | 0 | 0 | 1 | 0 | 0 | 0 | 0 | 1 | 0 | 0 | 0 |
| 52.25304 | 20.93287 | MAZ_U_7 | mazu7_2 | Poland | U | 1 | 1 | 0 | 0 | 0 | 0 | 1 | 107 | 1386,798595 | 31 | 56817,294 | 1871 | 1 | 0 | 1 | 1 | 1 | 0 | 0 | 1 | 0 | 0 | 1 | 0 | 0 | 1 | 0 | 0 |
| 52.25424 | 20.93523 | MAZ_U_7 | mazu7_3 | Poland | U | 1 | 1 | 0 | 0 | 0 | 0 | 1 | 108 | 1386,832654 | 31 | 31669,019 | 11757 | 1 | 1 | 0 | 0 | 1 | 0 | 0 | 1 | 1 | 0 | 0 | 0 | 0 | 1 | 0 | 0 |
| 52.25606 | 20.93723 | MAZ_U_7 | mazu7_4 | Poland | U | 0 | 0 | 0 | 0 | 0 | 0 | 1 | 107,184 | 1386,940513 | 31 | 48002,98 | 2415 | 1 | 1 | 0 | 0 | 1 | 0 | 0 | 0 | 0 | 0 | 0 | 0 | 0 | 0 | 0 | 0 |
| 52.25749 | 20.94077 | MAZ_U_7 | mazu7_5 | Poland | U | 0 | 0 | 0 | 0 | 0 | 0 | 1 | 104,008 | 1386,958529 | 31 | 40778,004 | 2644 | 1 | 1 | 0 | 0 | 1 | 0 | 0 | 0 | 0 | 0 | 1 | 0 | 0 | 0 | 0 | 1 |
| 52.25767 | 20.94377 | MAZ_U_7 | mazu7_6 | Poland | U | 1 | 1 | 0 | 0 | 0 | 0 | 1 | 105,388 | 1386,873738 | 31 | 40104,325 | 2190 | 1 | 1 | 0 | 1 | 1 | 0 | 0 | 1 | 0 | 0 | 1 | 0 | 0 | 0 | 0 | 0 |
| 52.25936 | 20.94645 | MAZ_U_7 | mazu7_7 | Poland | U | 2 | 1 | 0 | 0 | 0 | 0 | 1 | 104 | 1387,033947 | 31 | 43643,66 | 3405 | 1 | 1 | 0 | 1 | 0 | 0 | 0 | 1 | 0 | 0 | 1 | 0 | 0 | 0 | 0 | 0 |
| 52.26135 | 20.94954 | MAZ_U_7 | mazu7_8 | Poland | U | 0 | 0 | 0 | 0 | 0 | 0 | 1 | 102,564 | 1386,946055 | 31 | 28755,525 | 9978 | 1 | 1 | 0 | 1 | 0 | 0 | 0 | 1 | 0 | 0 | 1 | 0 | 0 | 0 | 1 | 1 |
| 52.26928 | 20.92994 | MAZ_U_8 | mazu8_1 | Poland | U | 0 | 0 | 0 | 0 | 0 | 0 | 0 | 105 | 1388,464882 | 29 | 24300,919 | 6288 | 1 | 1 | 1 | 0 | 0 | 0 | 0 | 1 | 1 | 0 | 0 | 1 | 0 | 1 | 0 | 0 |
| 52.27007 | 20.92576 | MAZ_U_8 | mazu8_2 | Poland | U | 1 | 1 | 0 | 0 | 0 | 0 | 0 | 104,197 | 1388,684433 | 29 | 19107,775 | 3165 | 1 | 1 | 1 | 0 | 0 | 0 | 0 | 1 | 1 | 0 | 0 | 1 | 1 | 1 | 0 | 0 |
| 52.27087 | 20.92132 | MAZ_U_8 | mazu8_3 | Poland | U | 0 | 0 | 0 | 0 | 0 | 0 | 1 | 104,868 | 1388,912307 | 29 | 22989,207 | 5161 | 1 | 1 | 1 | 1 | 0 | 0 | 0 | 1 | 1 | 0 | 0 | 1 | 1 | 0 | 0 | 0 |
| 52.27178 | 20.91741 | MAZ_U_8 | mazu8_4 | Poland | U | 0 | 0 | 0 | 0 | 0 | 0 | 0 | 105 | 1389,135873 | 29 | 18410,811 | 6283 | 1 | 1 | 1 | 0 | 1 | 0 | 0 | 1 | 1 | 0 | 0 | 1 | 1 | 0 | 0 | 0 |
| 52.27268 | 20.91342 | MAZ_U_8 | mazu8_5 | Poland | U | 0 | 0 | 0 | 0 | 0 | 0 | 0 | 105,648 | 1389,356228 | 29 | 9267,34 | 11712 | 1 | 1 | 0 | 1 | 0 | 0 | 1 | 1 | 1 | 0 | 0 | 0 | 1 | 0 | 1 | 0 |
| 52.2736 | 20.90951 | MAZ_U_8 | mazu8_6 | Poland | U | 0 | 0 | 0 | 0 | 0 | 0 | 0 | 105,733 | 1389,576081 | 29 | 6040,091 | 8774 | 0 | 1 | 0 | 1 | 0 | 0 | 0 | 1 | 0 | 0 | 1 | 0 | 0 | 1 | 1 | 0 |
| 52.27451 | 20.90559 | MAZ_U_8 | mazu8_7 | Poland | U | 0 | 0 | 0 | 0 | 0 | 0 | 0 | 105 | 1389,796972 | 29 | 17341,843 | 5625 | 1 | 1 | 0 | 1 | 0 | 0 | 0 | 1 | 1 | 1 | 0 | 1 | 0 | 0 | 1 | 0 |
| 52.27646 | 20.904 | MAZ_U_8 | mazu8_8 | Poland | U | 0 | 0 | 0 | 0 | 1 | 1 | 0 | 104,744 | 1390,038945 | 29 | 32608,197 | 4954 | 1 | 0 | 1 | 1 | 0 | 0 | 0 | 0 | 1 | 1 | 0 | 1 | 0 | 0 | 0 | 1 |
| 52.27195 | 20.95764 | MAZ_U_9 | mazu9_1 | Poland | U | 0 | 0 | 0 | 0 | 0 | 0 | 1 | 97,02 | 1387,783125 | 29 | 24551,462 | 5548 | 0 | 1 | 0 | 0 | 0 | 0 | 0 | 1 | 0 | 0 | 0 | 0 | 1 | 0 | 0 | 0 |
| 52.27379 | 20.95438 | MAZ_U_9 | mazu9_2 | Poland | U | 0 | 0 | 0 | 0 | 0 | 0 | 1 | 98,139 | 1388,071833 | 29 | 21099,897 | 8691 | 0 | 1 | 0 | 0 | 0 | 0 | 0 | 1 | 0 | 0 | 0 | 0 | 1 | 0 | 0 | 0 |
| 52.27426 | 20.95026 | MAZ_U_9 | mazu9_3 | Poland | U | 0 | 0 | 0 | 0 | 0 | 0 | 1 | 99 | 1388,256931 | 29 | 31710,608 | 10879 | 0 | 1 | 0 | 0 | 0 | 0 | 0 | 1 | 0 | 0 | 0 | 0 | 1 | 0 | 0 | 0 |
| 52.2742 | 20.94577 | MAZ_U_9 | mazu9_4 | Poland | U | 0 | 0 | 0 | 0 | 0 | 0 | 0 | 100,201 | 1388,402628 | 29 | 8279,156 | 12683 | 0 | 1 | 0 | 0 | 0 | 0 | 0 | 1 | 0 | 0 | 0 | 0 | 1 | 1 | 0 | 0 |
| 52.2749 | 20.94161 | MAZ_U_9 | mazu9_5 | Poland | U | 0 | 0 | 0 | 0 | 0 | 0 | 1 | 101,36 | 1388,611267 | 29 | 22077,752 | 11221 | 0 | 1 | 0 | 0 | 0 | 0 | 1 | 1 | 0 | 0 | 0 | 0 | 1 | 1 | 0 | 0 |
| 52.27596 | 20.93761 | MAZ_U_9 | mazu9_6 | Poland | U | 0 | 0 | 0 | 0 | 0 | 0 | 1 | 103 | 1388,849387 | 29 | 10878,376 | 12799 | 0 | 1 | 0 | 0 | 0 | 0 | 1 | 1 | 1 | 0 | 0 | 1 | 1 | 1 | 1 | 0 |
| 52.27773 | 20.93506 | MAZ_U_9 | mazu9_7 | Poland | U | 0 | 0 | 0 | 0 | 0 | 0 | 1 | 102,828 | 1389,106356 | 29 | 18203,398 | 6641 | 0 | 1 | 0 | 0 | 0 | 0 | 1 | 1 | 0 | 0 | 0 | 1 | 1 | 0 | 0 | 0 |
| 52.27993 | 20.93208 | MAZ_U_9 | mazu9_8 | Poland | U | 0 | 0 | 0 | 0 | 0 | 0 | 1 | 100,123 | 1389,420199 | 29 | 15800,806 | 8065 | 0 | 1 | 0 | 0 | 0 | 0 | 0 | 1 | 0 | 0 | 0 | 0 | 0 | 1 | 0 | 1 |
| 49.91677 | 20.16472 | MLP_R_1 | mlpr1_1 | Poland | R | 0 | 0 | 0 | 0 | 0 | 0 | 0 | 278,38 | 1201,981726 | 223 | 10259,758 | 2629 | 1 | 1 | 0 | 1 | 0 | 0 | 1 | 1 | 1 | 0 | 0 | 1 | 0 | 0 | 0 | 0 |
| 49.91814 | 20.16185 | MLP_R_1 | mlpr1_2 | Poland | R | 0 | 0 | 0 | 0 | 1 | 1 | 1 | 292,392 | 1202,225033 | 223 | 14143,97 | 1938 | 1 | 1 | 0 | 1 | 0 | 0 | 1 | 1 | 1 | 0 | 0 | 1 | 0 | 0 | 0 | 0 |
| 49.91912 | 20.15838 | MLP_R_1 | mlpr1_3 | Poland | R | 4 | 1 | 0 | 0 | 0 | 0 | 1 | 296,028 | 1202,470338 | 223 | 33469,829 | 639 | 1 | 1 | 0 | 1 | 0 | 0 | 1 | 1 | 1 | 0 | 1 | 0 | 0 | 0 | 0 | 0 |
| 49.91803 | 20.15594 | MLP_R_1 | mlpr1_4 | Poland | R | 0 | 0 | 0 | 0 | 0 | 0 | 1 | 309,292 | 1202,486145 | 223 | 15125,426 | 1562 | 1 | 1 | 0 | 1 | 0 | 0 | 1 | 1 | 1 | 0 | 1 | 1 | 0 | 0 | 0 | 0 |
| 49.91549 | 20.15605 | MLP_R_1 | mlpr1_5 | Poland | R | 0 | 0 | 1 | 1 | 0 | 0 | 1 | 310,528 | 1202,26122 | 223 | 17611,153 | 837 | 1 | 1 | 0 | 1 | 0 | 0 | 1 | 1 | 1 | 0 | 1 | 0 | 0 | 0 | 0 | 0 |
| 49.91317 | 20.15626 | MLP_R_1 | mlpr1_6 | Poland | R | 1 | 1 | 1 | 1 | 0 | 0 | 1 | 302,419 | 1202,050932 | 223 | 25017,132 | 1276 | 1 | 1 | 0 | 1 | 0 | 0 | 1 | 1 | 1 | 0 | 1 | 1 | 0 | 0 | 0 | 0 |
| 49.91236 | 20.1604 | MLP_R_1 | mlpr1_7 | Poland | R | 0 | 0 | 0 | 0 | 0 | 0 | 0 | 293,282 | 1201,793618 | 223 | 19501,301 | 0 | 1 | 1 | 0 | 1 | 0 | 0 | 1 | 0 | 0 | 0 | 0 | 0 | 0 | 0 | 0 | 0 |
| 49.91277 | 20.16438 | MLP_R_1 | mlpr1_8 | Poland | R | 0 | 0 | 0 | 0 | 0 | 0 | 0 | 270,232 | 1201,651234 | 223 | 16842,203 | 1252 | 1 | 1 | 0 | 1 | 0 | 0 | 1 | 0 | 1 | 0 | 1 | 0 | 0 | 0 | 0 | 0 |
| 50.19105 | 20.23196 | MLP_R_10 | mlpr10_1 | Poland | R | 0 | 0 | 0 | 0 | 0 | 0 | 0 | 236,189 | 1222,965939 | 215 | 7152,393 | 0 | 0 | 1 | 0 | 0 | 0 | 0 | 1 | 0 | 0 | 0 | 0 | 0 | 0 | 0 | 0 | 0 |
| 50.19206 | 20.23532 | MLP_R_10 | mlpr10_2 | Poland | R | 0 | 0 | 0 | 0 | 0 | 0 | 0 | 225,521 | 1222,907381 | 215 | 17676,137 | 0 | 0 | 1 | 0 | 0 | 0 | 0 | 1 | 0 | 0 | 0 | 0 | 0 | 0 | 0 | 0 | 0 |
| 50.19429 | 20.23684 | MLP_R_10 | mlpr10_3 | Poland | R | 0 | 0 | 0 | 0 | 0 | 0 | 0 | 218,405 | 1223,037976 | 215 | 21038,153 | 2124 | 0 | 1 | 0 | 1 | 0 | 0 | 1 | 0 | 1 | 0 | 1 | 0 | 0 | 0 | 0 | 1 |
| 50.1959 | 20.23478 | MLP_R_10 | mlpr10_4 | Poland | R | 2 | 1 | 0 | 0 | 0 | 0 | 1 | 219,914 | 1223,269503 | 215 | 36638,201 | 5794 | 1 | 1 | 0 | 1 | 0 | 0 | 1 | 1 | 0 | 0 | 1 | 0 | 1 | 0 | 0 | 1 |
| 50.19805 | 20.23536 | MLP_R_10 | mlpr10_5 | Poland | R | 0 | 0 | 1 | 1 | 0 | 0 | 1 | 210,014 | 1223,432848 | 215 | 27810,163 | 0 | 1 | 1 | 0 | 1 | 0 | 0 | 1 | 0 | 0 | 1 | 1 | 0 | 0 | 0 | 0 | 0 |
| 50.20038 | 20.2344 | MLP_R_10 | mlpr10_6 | Poland | R | 0 | 0 | 0 | 0 | 0 | 0 | 0 | 210,41 | 1223,680042 | 215 | 590,513 | 0 | 0 | 1 | 0 | 0 | 0 | 0 | 1 | 0 | 0 | 0 | 0 | 0 | 0 | 0 | 0 | 0 |
| 50.20288 | 20.23538 | MLP_R_10 | mlpr10_7 | Poland | R | 0 | 0 | 0 | 0 | 0 | 0 | 0 | 210,601 | 1223,857858 | 215 | 3293,461 | 1741 | 0 | 1 | 0 | 0 | 0 | 0 | 1 | 0 | 1 | 0 | 1 | 0 | 0 | 0 | 0 | 0 |
| 50.20388 | 20.23056 | MLP_R_10 | mlpr10_8 | Poland | R | 0 | 0 | 0 | 0 | 0 | 0 | 0 | 212,953 | 1224,155947 | 215 | 12803,465 | 3460 | 1 | 1 | 0 | 1 | 0 | 0 | 1 | 1 | 1 | 0 | 1 | 1 | 0 | 0 | 0 | 0 |
| 50.11096 | 19.80047 | MLP_R_11 | mlpr11_1 | Poland | R | 0 | 0 | 0 | 0 | 0 | 0 | 0 | 254,552 | 1235,206525 | 190 | 9612,667 | 8165 | 0 | 1 | 1 | 0 | 0 | 0 | 0 | 1 | 0 | 0 | 1 | 0 | 0 | 0 | 0 | 0 |
| 50.10871 | 19.79826 | MLP_R_11 | mlpr11_2 | Poland | R | 0 | 0 | 0 | 0 | 0 | 0 | 1 | 269,592 | 1235,115288 | 190 | 6420,104 | 4954 | 0 | 1 | 1 | 0 | 0 | 0 | 0 | 1 | 1 | 0 | 0 | 1 | 0 | 0 | 0 | 0 |
| 50.10701 | 19.79839 | MLP_R_11 | mlpr11_3 | Poland | R | 1 | 1 | 0 | 0 | 0 | 0 | 1 | 274,876 | 1234,965967 | 190 | 9689,404 | 4072 | 1 | 1 | 1 | 0 | 0 | 0 | 0 | 1 | 1 | 0 | 0 | 1 | 0 | 0 | 0 | 0 |
| 50.10711 | 19.80122 | MLP_R_11 | mlpr11_4 | Poland | R | 1 | 1 | 0 | 0 | 0 | 0 | 0 | 269,438 | 1234,843941 | 190 | 9671,034 | 5205 | 1 | 1 | 1 | 0 | 0 | 0 | 0 | 1 | 1 | 0 | 0 | 1 | 0 | 0 | 0 | 0 |
| 50.1054 | 19.80322 | MLP_R_11 | mlpr11_5 | Poland | R | 0 | 0 | 0 | 0 | 0 | 0 | 0 | 254,337 | 1234,606086 | 190 | 12487,466 | 4468 | 0 | 1 | 1 | 0 | 0 | 0 | 0 | 1 | 1 | 0 | 0 | 1 | 0 | 0 | 0 | 0 |

| | | | | | | | | | | | | | | | | | | | | | | | | | | | | | | | | |
|---|---|---|---|---|---|---|---|---|---|---|---|---|---|---|---|---|---|---|---|---|---|---|---|---|---|---|---|---|---|---|---|---|
| 50.10625 | 19.80549 | MLP_R_11 | mlpr11_6 | Poland | R | 1 | 1 | 0 | 0 | 0 | 0 | 0 | 270,264 | 1234,572623 | 190 | 11503,817 | 5460 | 0 | 1 | 1 | 0 | 0 | 0 | 0 | 1 | 1 | 0 | 0 | 1 | 0 | 0 | 0 | 0 |
| 50.10799 | 19.80763 | MLP_R_11 | mlpr11_7 | Poland | R | 0 | 0 | 0 | 0 | 0 | 0 | 1 | 262,878 | 1234,629479 | 190 | 6862,298 | 4284 | 0 | 1 | 1 | 0 | 0 | 0 | 0 | 1 | 0 | 0 | 0 | 1 | 0 | 0 | 0 | 0 |
| 50.11052 | 19.80915 | MLP_R_11 | mlpr11_8 | Poland | R | 0 | 0 | 0 | 0 | 0 | 0 | 0 | 253,744 | 1234,774488 | 190 | 8789,18 | 5086 | 1 | 1 | 1 | 0 | 0 | 0 | 0 | 1 | 0 | 0 | 0 | 1 | 0 | 0 | 0 | 0 |
| 49.88334 | 19.67119 | MLP_R_12 | mlpr12_1 | Poland | R | 1 | 1 | 0 | 0 | 0 | 0 | 1 | 293,384 | 1221,755252 | 194 | 37281,922 | 0 | 1 | 1 | 0 | 1 | 0 | 0 | 1 | 0 | 0 | 0 | 0 | 0 | 0 | 0 | 0 | 0 |
| 49.88189 | 19.67363 | MLP_R_12 | mlpr12_2 | Poland | R | 0 | 0 | 0 | 0 | 0 | 0 | 1 | 290,06 | 1221,518779 | 194 | 7566,255 | 0 | 0 | 1 | 0 | 0 | 0 | 0 | 1 | 0 | 0 | 0 | 0 | 0 | 0 | 0 | 0 | 0 |
| 49.88069 | 19.67659 | MLP_R_12 | mlpr12_3 | Poland | R | 0 | 0 | 0 | 0 | 0 | 0 | 1 | 280,134 | 1221,279639 | 194 | 16671,712 | 3920 | 1 | 1 | 0 | 1 | 0 | 0 | 1 | 0 | 0 | 1 | 1 | 1 | 0 | 0 | 0 | 0 |
| 49.88272 | 19.67803 | MLP_R_12 | mlpr12_4 | Poland | R | 0 | 0 | 0 | 0 | 0 | 0 | 1 | 272 | 1221,384049 | 194 | 13994,617 | 4203 | 1 | 1 | 0 | 1 | 0 | 0 | 1 | 1 | 1 | 0 | 1 | 0 | 1 | 0 | 0 | 0 |
| 49.88453 | 19.67891 | MLP_R_12 | mlpr12_5 | Poland | R | 0 | 0 | 0 | 0 | 0 | 0 | 1 | 273,559 | 1221,496038 | 194 | 17836,266 | 772 | 0 | 1 | 0 | 0 | 0 | 1 | 1 | 0 | 0 | 0 | 1 | 0 | 0 | 0 | 0 | 1 |
| 49.8866 | 19.67785 | MLP_R_12 | mlpr12_6 | Poland | R | 0 | 0 | 0 | 0 | 0 | 0 | 0 | 268,978 | 1221,720136 | 194 | 31237,336 | 1231 | 1 | 1 | 0 | 1 | 0 | 0 | 1 | 0 | 1 | 0 | 0 | 1 | 0 | 0 | 0 | 1 |
| 49.88891 | 19.6761 | MLP_R_12 | mlpr12_7 | Poland | R | 2 | 1 | 0 | 0 | 0 | 0 | 1 | 269,887 | 1221,997508 | 194 | 34496,094 | 1533 | 1 | 1 | 0 | 1 | 0 | 0 | 1 | 0 | 0 | 0 | 0 | 1 | 0 | 0 | 0 | 0 |
| 49.89059 | 19.67382 | MLP_R_12 | mlpr12_8 | Poland | R | 0 | 0 | 0 | 0 | 0 | 0 | 0 | 273,248 | 1222,246017 | 194 | 18917,081 | 5240 | 1 | 1 | 0 | 1 | 0 | 0 | 1 | 0 | 1 | 0 | 1 | 1 | 0 | 0 | 0 | 1 |
| 50.1721 | 19.40267 | MLP_R_13 | mlpr13_1 | Poland | R | 0 | 0 | 0 | 0 | 0 | 0 | 1 | 309,56 | 1258,676415 | 161 | 4877,578 | 5301 | 0 | 1 | 0 | 0 | 0 | 0 | 0 | 1 | 1 | 0 | 0 | 1 | 0 | 0 | 0 | 0 |
| 50.17409 | 19.40359 | MLP_R_13 | mlpr13_2 | Poland | R | 0 | 0 | 0 | 0 | 0 | 0 | 1 | 317,055 | 1258,799584 | 161 | 5366,268 | 9448 | 0 | 1 | 0 | 0 | 0 | 0 | 0 | 1 | 1 | 0 | 0 | 1 | 0 | 0 | 0 | 0 |
| 50.17602 | 19.40516 | MLP_R_13 | mlpr13_3 | Poland | R | 0 | 0 | 0 | 0 | 0 | 0 | 1 | 327,672 | 1258,891643 | 161 | 6468,983 | 3976 | 0 | 1 | 0 | 1 | 0 | 0 | 0 | 0 | 1 | 0 | 1 | 0 | 0 | 0 | 0 | 0 |
| 50.17702 | 19.40134 | MLP_R_13 | mlpr13_4 | Poland | R | 0 | 0 | 0 | 0 | 0 | 0 | 1 | 322,68 | 1259,156738 | 161 | 3448,525 | 4791 | 0 | 1 | 0 | 0 | 0 | 0 | 0 | 1 | 1 | 0 | 1 | 1 | 0 | 0 | 0 | 0 |
| 50.1771 | 19.39735 | MLP_R_13 | mlpr13_5 | Poland | R | 0 | 0 | 0 | 0 | 0 | 0 | 1 | 315,56 | 1259,344283 | 161 | 18895,739 | 3134 | 1 | 1 | 0 | 1 | 0 | 0 | 1 | 0 | 1 | 0 | 0 | 1 | 0 | 0 | 0 | 1 |
| 50.17643 | 19.39359 | MLP_R_13 | mlpr13_6 | Poland | R | 2 | 1 | 0 | 0 | 0 | 0 | 1 | 312,924 | 1259,463297 | 161 | 22376,755 | 1031 | 1 | 1 | 0 | 1 | 0 | 0 | 1 | 1 | 1 | 0 | 1 | 0 | 0 | 0 | 1 | 0 |
| 50.17521 | 19.3958 | MLP_R_13 | mlpr13_7 | Poland | R | 1 | 1 | 0 | 0 | 0 | 0 | 0 | 310,029 | 1259,255588 | 161 | 48667,064 | 340 | 1 | 1 | 0 | 1 | 0 | 0 | 0 | 0 | 0 | 1 | 0 | 0 | 0 | 0 | 1 | 0 |
| 50.17356 | 19.39691 | MLP_R_13 | mlpr13_8 | Poland | R | 0 | 0 | 0 | 0 | 0 | 0 | 0 | 307,147 | 1259,062891 | 161 | 30243,36 | 4918 | 1 | 1 | 0 | 1 | 0 | 0 | 0 | 0 | 0 | 1 | 0 | 0 | 0 | 0 | 0 | 0 |
| 50.22979 | 19.77411 | MLP_R_14 | mlpr14_1 | Poland | R | 1 | 1 | 0 | 0 | 0 | 0 | 1 | 466,796 | 1246,636912 | 182 | 6480,231 | 3517 | 0 | 1 | 1 | 0 | 0 | 1 | 0 | 0 | 1 | 0 | 0 | 1 | 0 | 0 | 0 | 0 |
| 50.22882 | 19.76968 | MLP_R_14 | mlpr14_2 | Poland | R | 0 | 0 | 0 | 0 | 0 | 0 | 1 | 450,714 | 1246,75223 | 182 | 7886,209 | 4037 | 0 | 1 | 1 | 0 | 0 | 0 | 0 | 0 | 1 | 0 | 1 | 1 | 0 | 0 | 0 | 0 |
| 50.22647 | 19.76798 | MLP_R_14 | mlpr14_3 | Poland | R | 1 | 1 | 0 | 0 | 0 | 0 | 1 | 426,292 | 1246,625897 | 182 | 19506,748 | 5519 | 1 | 1 | 1 | 1 | 0 | 0 | 1 | 0 | 1 | 0 | 0 | 1 | 0 | 0 | 0 | 0 |
| 50.22417 | 19.7701 | MLP_R_14 | mlpr14_4 | Poland | R | 4 | 1 | 0 | 0 | 0 | 0 | 1 | 417,092 | 1246,33345 | 182 | 28922,832 | 4086 | 1 | 1 | 1 | 1 | 0 | 0 | 0 | 0 | 1 | 0 | 1 | 1 | 0 | 0 | 0 | 0 |
| 50.22184 | 19.77234 | MLP_R_14 | mlpr14_5 | Poland | R | 1 | 1 | 0 | 0 | 0 | 0 | 1 | 408,793 | 1246,03021 | 182 | 30750,429 | 3145 | 1 | 1 | 1 | 1 | 0 | 0 | 0 | 1 | 1 | 0 | 1 | 0 | 0 | 0 | 0 | 1 |
| 50.22 | 19.77549 | MLP_R_14 | mlpr14_6 | Poland | R | 1 | 1 | 0 | 0 | 0 | 0 | 1 | 402 | 1245,729087 | 182 | 29159,763 | 3121 | 1 | 1 | 1 | 1 | 0 | 0 | 0 | 1 | 1 | 0 | 1 | 1 | 0 | 0 | 0 | 1 |
| 50.2185 | 19.77855 | MLP_R_14 | mlpr14_7 | Poland | R | 1 | 1 | 0 | 0 | 0 | 0 | 1 | 396 | 1245,461006 | 182 | 42572,529 | 1118 | 1 | 1 | 1 | 1 | 0 | 0 | 0 | 1 | 1 | 0 | 1 | 0 | 0 | 0 | 0 | 1 |
| 50.21749 | 19.7822 | MLP_R_14 | mlpr14_8 | Poland | R | 1 | 1 | 0 | 0 | 0 | 0 | 0 | 390,592 | 1245,210886 | 182 | 43453,217 | 0 | 1 | 0 | 1 | 1 | 0 | 0 | 0 | 0 | 0 | 1 | 0 | 0 | 0 | 0 | 0 | 1 |
| 49.74051 | 19.98897 | MLP_R_15 | mlpr15_1 | Poland | R | 0 | 0 | 0 | 0 | 0 | 0 | 0 | 345,556 | 1194,848451 | 221 | 30709,167 | 0 | 1 | 1 | 1 | 1 | 0 | 0 | 0 | 0 | 0 | 1 | 0 | 0 | 0 | 0 | 0 | 0 |
| 49.74049 | 19.98423 | MLP_R_15 | mlpr15_2 | Poland | R | 1 | 1 | 0 | 0 | 0 | 0 | 0 | 337,818 | 1195,071426 | 221 | 16327,81 | 2781 | 1 | 0 | 0 | 1 | 0 | 0 | 0 | 0 | 1 | 1 | 1 | 1 | 0 | 0 | 0 | 0 |
| 49.7404 | 19.98105 | MLP_R_15 | mlpr15_3 | Poland | R | 0 | 0 | 0 | 0 | 0 | 0 | 1 | 332,463 | 1195,211049 | 221 | 29248,946 | 2535 | 1 | 0 | 0 | 1 | 0 | 0 | 0 | 0 | 1 | 1 | 0 | 1 | 0 | 0 | 0 | 0 |
| 49.73796 | 19.98131 | MLP_R_15 | mlpr15_4 | Poland | R | 0 | 0 | 0 | 0 | 0 | 0 | 0 | 332,051 | 1194,989824 | 221 | 34016,324 | 1353 | 1 | 0 | 0 | 1 | 0 | 0 | 0 | 0 | 1 | 1 | 1 | 1 | 0 | 0 | 0 | 0 |
| 49.73558 | 19.98256 | MLP_R_15 | mlpr15_5 | Poland | R | 0 | 0 | 0 | 0 | 0 | 0 | 0 | 335,344 | 1194,726287 | 221 | 25573,062 | 2547 | 1 | 1 | 0 | 1 | 0 | 0 | 0 | 0 | 1 | 0 | 1 | 1 | 0 | 0 | 0 | 0 |
| 49.73373 | 19.97984 | MLP_R_15 | mlpr15_6 | Poland | R | 2 | 1 | 0 | 0 | 0 | 0 | 1 | 334 | 1194,694123 | 221 | 36866,89 | 0 | 1 | 1 | 0 | 1 | 0 | 0 | 0 | 0 | 1 | 1 | 0 | 0 | 0 | 0 | 0 | 0 |
| 49.73238 | 19.9837 | MLP_R_15 | mlpr15_7 | Poland | R | 0 | 0 | 0 | 0 | 0 | 0 | 0 | 339,32 | 1194,401336 | 221 | 20761,143 | 1158 | 1 | 1 | 1 | 1 | 0 | 0 | 0 | 0 | 1 | 0 | 1 | 1 | 0 | 0 | 0 | 0 |
| 49.73158 | 19.98746 | MLP_R_15 | mlpr15_8 | Poland | R | 1 | 1 | 0 | 0 | 0 | 0 | 0 | 356,044 | 1194,155717 | 221 | 38179,007 | 2066 | 1 | 1 | 1 | 1 | 0 | 0 | 0 | 0 | 1 | 0 | 0 | 1 | 0 | 0 | 0 | 0 |
| 49.96767 | 20.51131 | MLP_R_16 | mlpr16_1 | Poland | R | 0 | 0 | 0 | 0 | 0 | 0 | 0 | 226,82 | 1191,219902 | 243 | 28316,883 | 3833 | 0 | 1 | 0 | 1 | 0 | 0 | 1 | 0 | 1 | 0 | 1 | 0 | 0 | 0 | 0 | 1 |
| 49.96781 | 20.50778 | MLP_R_16 | mlpr16_2 | Poland | R | 2 | 1 | 0 | 0 | 0 | 0 | 0 | 214,993 | 1191,236222 | 243 | 15705,488 | 2214 | 1 | 1 | 0 | 1 | 0 | 0 | 1 | 0 | 1 | 1 | 1 | 0 | 0 | 0 | 0 | 0 |
| 49.96768 | 20.50361 | MLP_R_16 | mlpr16_3 | Poland | R | 0 | 0 | 0 | 0 | 0 | 0 | 0 | 219,299 | 1191,395583 | 243 | 14304,096 | 747 | 1 | 1 | 0 | 1 | 0 | 0 | 0 | 0 | 1 | 1 | 1 | 0 | 0 | 0 | 0 | 0 |
| 49.96923 | 20.50575 | MLP_R_16 | mlpr16_4 | Poland | R | 0 | 0 | 0 | 0 | 0 | 0 | 0 | 233,137 | 1191,398155 | 243 | 19889,878 | 2842 | 1 | 1 | 0 | 1 | 0 | 0 | 0 | 0 | 1 | 1 | 1 | 0 | 0 | 0 | 0 | 0 |
| 49.97036 | 20.50948 | MLP_R_16 | mlpr16_5 | Poland | R | 0 | 0 | 0 | 0 | 0 | 0 | 0 | 219,334 | 1191,460943 | 243 | 9815,561 | 3176 | 1 | 1 | 0 | 1 | 0 | 0 | 0 | 0 | 1 | 1 | 1 | 0 | 0 | 0 | 0 | 0 |
| 49.9718 | 20.51243 | MLP_R_16 | mlpr16_6 | Poland | R | 0 | 0 | 1 | 1 | 0 | 0 | 1 | 222,571 | 1191,416317 | 243 | 29571,977 | 3270 | 1 | 1 | 0 | 1 | 0 | 0 | 0 | 0 | 1 | 1 | 1 | 0 | 1 | 0 | 0 | 0 |
| 49.97182 | 20.51614 | MLP_R_16 | mlpr16_7 | Poland | R | 0 | 0 | 0 | 0 | 0 | 0 | 1 | 213,645 | 1191,246326 | 243 | 16998,64 | 3457 | 1 | 1 | 0 | 1 | 0 | 0 | 0 | 0 | 1 | 1 | 1 | 0 | 0 | 0 | 0 | 0 |
| 49.97332 | 20.51955 | MLP_R_16 | mlpr16_8 | Poland | R | 1 | 1 | 0 | 0 | 0 | 0 | 1 | 213 | 1191,078983 | 243 | 18142,48 | 1936 | 1 | 1 | 0 | 1 | 0 | 0 | 0 | 1 | 1 | 1 | 1 | 0 | 0 | 0 | 0 | 0 |
| 49.85757 | 20.20881 | MLP_R_2 | mlpr2_1 | Poland | R | 0 | 0 | 0 | 0 | 0 | 0 | 1 | 334,976 | 1194,865475 | 228 | 7022,127 | 241 | 0 | 1 | 0 | 0 | 0 | 0 | 1 | 1 | 1 | 0 | 1 | 0 | 0 | 0 | 0 | 0 |
| 49.85682 | 20.20514 | MLP_R_2 | mlpr2_2 | Poland | R | 0 | 0 | 2 | 1 | 0 | 0 | 1 | 300,82 | 1194,962529 | 228 | 23646,807 | 0 | 1 | 1 | 0 | 1 | 0 | 0 | 1 | 0 | 1 | 0 | 0 | 0 | 0 | 0 | 0 | 0 |
| 49.85475 | 20.20558 | MLP_R_2 | mlpr2_3 | Poland | R | 0 | 0 | 0 | 0 | 0 | 0 | 1 | 276,388 | 1194,767405 | 228 | 21151,111 | 0 | 1 | 1 | 0 | 1 | 0 | 1 | 1 | 0 | 0 | 0 | 1 | 0 | 0 | 0 | 0 | 0 |
| 49.85322 | 20.20848 | MLP_R_2 | mlpr2_4 | Poland | R | 0 | 0 | 1 | 1 | 0 | 0 | 1 | 272,841 | 1194,503006 | 228 | 22306,309 | 3792 | 1 | 1 | 0 | 1 | 0 | 0 | 1 | 1 | 0 | 0 | 1 | 1 | 1 | 0 | 0 | 0 |
| 49.85129 | 20.21089 | MLP_R_2 | mlpr2_5 | Poland | R | 2 | 1 | 0 | 0 | 0 | 0 | 1 | 271,891 | 1194,225879 | 228 | 32200,665 | 1255 | 1 | 1 | 0 | 1 | 0 | 0 | 1 | 0 | 0 | 0 | 1 | 0 | 0 | 0 | 0 | 1 |
| 49.85189 | 20.21352 | MLP_R_2 | mlpr2_6 | Poland | R | 2 | 1 | 0 | 0 | 0 | 0 | 1 | 298,216 | 1194,162093 | 228 | 30641,952 | 1730 | 1 | 1 | 0 | 1 | 0 | 0 | 1 | 0 | 1 | 0 | 1 | 0 | 0 | 0 | 0 | 0 |
| 49.85405 | 20.21492 | MLP_R_2 | mlpr2_7 | Poland | R | 0 | 0 | 0 | 0 | 0 | 0 | 1 | 331,032 | 1194,286424 | 228 | 21737,231 | 0 | 1 | 1 | 0 | 1 | 0 | 0 | 1 | 0 | 1 | 0 | 0 | 0 | 0 | 0 | 0 | 1 |
| 49.85622 | 20.21657 | MLP_R_2 | mlpr2_8 | Poland | R | 0 | 0 | 0 | 0 | 0 | 0 | 0 | 351,308 | 1194,399234 | 228 | 17894,575 | 461 | 1 | 1 | 0 | 1 | 0 | 0 | 1 | 0 | 0 | 1 | 0 | 0 | 0 | 0 | 0 | 0 |
| 50.10513 | 20.30364 | MLP_R_3 | mlpr3_1 | Poland | R | 0 | 0 | 1 | 1 | 0 | 0 | 1 | 189,572 | 1212,270092 | 223 | 6832,75 | 3193 | 0 | 1 | 0 | 0 | 0 | 0 | 0 | 1 | 1 | 1 | 1 | 1 | 0 | 0 | 0 | 0 |
| 50.10309 | 20.302 | MLP_R_3 | mlpr3_2 | Poland | R | 1 | 1 | 0 | 0 | 0 | 0 | 1 | 189,248 | 1212,163363 | 223 | 4958,333 | 2775 | 0 | 1 | 0 | 0 | 0 | 0 | 0 | 0 | 1 | 1 | 0 | 1 | 0 | 0 | 0 | 0 |
| 50.10103 | 20.30485 | MLP_R_3 | mlpr3_3 | Poland | R | 1 | 1 | 1 | 1 | 1 | 1 | 0 | 187,618 | 1211,861187 | 223 | 9773,955 | 1592 | 0 | 1 | 0 | 0 | 0 | 0 | 0 | 0 | 0 | 1 | 0 | 1 | 0 | 0 | 0 | 0 |
| 50.10006 | 20.3093 | MLP_R_3 | mlpr3_4 | Poland | R | 0 | 0 | 0 | 0 | 0 | 0 | 0 | 187 | 1211,578615 | 223 | 5750,015 | 0 | 0 | 1 | 0 | 1 | 0 | 0 | 0 | 0 | 0 | 1 | 0 | 0 | 0 | 0 | 0 | 0 |
| 50.10002 | 20.31421 | MLP_R_3 | mlpr3_5 | Poland | R | 0 | 0 | 2 | 1 | 0 | 0 | 0 | 186,928 | 1211,355001 | 223 | 3168,961 | 0 | 1 | 1 | 0 | 0 | 0 | 0 | 0 | 0 | 0 | 1 | 0 | 0 | 0 | 0 | 0 | 0 |
| 50.10177 | 20.31618 | MLP_R_3 | mlpr3_6 | Poland | R | 1 | 1 | 0 | 0 | 0 | 0 | 0 | 187,248 | 1211,423529 | 223 | 6164,69 | 2031 | 1 | 1 | 0 | 0 | 0 | 0 | 0 | 0 | 1 | 1 | 0 | 1 | 0 | 0 | 0 | 0 |
| 50.10377 | 20.3172 | MLP_R_3 | mlpr3_7 | Poland | R | 0 | 0 | 0 | 0 | 0 | 0 | 1 | 198,936 | 1211,558405 | 223 | 5450,514 | 4931 | 0 | 1 | 0 | 1 | 0 | 0 | 0 | 0 | 1 | 1 | 0 | 1 | 0 | 0 | 0 | 0 |
| 50.10418 | 20.32084 | MLP_R_3 | mlpr3_8 | Poland | R | 2 | 1 | 0 | 0 | 0 | 0 | 1 | 197,977 | 1211,436978 | 223 | 14024,207 | 4563 | 1 | 1 | 0 | 0 | 0 | 0 | 0 | 1 | 0 | 1 | 0 | 1 | 0 | 0 | 0 | 0 |
| 50.15409 | 20.57799 | MLP_R_4 | mlpr4_1 | Poland | R | 1 | 1 | 1 | 1 | 3 | 1 | 0 | 179 | 1204,757768 | 239 | 14801,97 | 2082 | 1 | 1 | 1 | 1 | 0 | 0 | 1 | 0 | 1 | 0 | 0 | 1 | 0 | 0 | 0 | 0 |
| 50.15147 | 20.57917 | MLP_R_4 | mlpr4_2 | Poland | R | 0 | 0 | 0 | 0 | 2 | 1 | 1 | 183,803 | 1204,475739 | 239 | 7168,297 | 2600 | 1 | 1 | 0 | 1 | 0 | 0 | 0 | 1 | 1 | 0 | 0 | 1 | 0 | 0 | 0 | 1 |
| 50.15081 | 20.5831 | MLP_R_4 | mlpr4_3 | Poland | R | 0 | 0 | 0 | 0 | 0 | 0 | 1 | 181,575 | 1204,250958 | 239 | 32472,699 | 0 | 1 | 1 | 0 | 1 | 0 | 0 | 1 | 0 | 0 | 0 | 0 | 0 | 0 | 0 | 0 | 0 |
| 50.14987 | 20.58653 | MLP_R_4 | mlpr4_4 | Poland | R | 1 | 1 | 0 | 0 | 0 | 0 | 0 | 181,428 | 1204,019702 | 239 | 12215,693 | 0 | 1 | 1 | 0 | 1 | 0 | 0 | 0 | 0 | 0 | 1 | 0 | 0 | 0 | 0 | 0 | 1 |
| 50.14771 | 20.58625 | MLP_R_4 | mlpr4_5 | Poland | R | 0 | 0 | 0 | 0 | 0 | 0 | 1 | 181,39 | 1203,839951 | 239 | 8857,189 | 274 | 1 | 1 | 0 | 1 | 0 | 0 | 1 | 1 | 1 | 1 | 1 | 0 | 0 | 0 | 0 | 0 |
| 50.14831 | 20.58219 | MLP_R_4 | mlpr4_6 | Poland | R | 0 | 0 | 2 | 1 | 0 | 0 | 0 | 181,022 | 1204,070637 | 239 | 26067,149 | 790 | 1 | 0 | 0 | 1 | 0 | 0 | 0 | 0 | 1 | 0 | 1 | 1 | 0 | 0 | 0 | 0 |
| 50.14844 | 20.57796 | MLP_R_4 | mlpr4_7 | Poland | R | 0 | 0 | 0 | 0 | 0 | 0 | 1 | 187,401 | 1204,255912 | 239 | 30907,252 | 320 | 1 | 0 | 0 | 1 | 0 | 0 | 0 | 0 | 1 | 0 | 1 | 0 | 0 | 0 | 0 | 0 |
| 50.14906 | 20.57414 | MLP_R_4 | mlpr4_8 | Poland | R | 2 | 1 | 0 | 0 | 0 | 0 | 1 | 219,386 | 1204,475719 | 239 | 23329,984 | 0 | 1 | 1 | 0 | 1 | 0 | 0 | 0 | 0 | 0 | 1 | 0 | 0 | 0 | 0 | 0 | 0 |
| 50.16575 | 20.0715 | MLP_R_5 | mlpr5_1 | Poland | R | 2 | 1 | 0 | 0 | 0 | 0 | 0 | 280,5 | 1227,801581 | 206 | 23593,542 | 1697 | 1 | 1 | 0 | 1 | 0 | 0 | 1 | 0 | 1 | 0 | 1 | 1 | 0 | 0 | 1 | 0 |
| 50.16471 | 20.07521 | MLP_R_5 | mlpr5_2 | Poland | R | 1 | 1 | 0 | 0 | 0 | 0 | 0 | 266 | 1227,548925 | 206 | 18805,444 | 2582 | 1 | 1 | 0 | 1 | 0 | 0 | 1 | 0 | 1 | 0 | 1 | 1 | 0 | 0 | 1 | 1 |
| 50.16608 | 20.07888 | MLP_R_5 | mlpr5_3 | Poland | R | 0 | 0 | 0 | 0 | 1 | 1 | 0 | 284,076 | 1227,509805 | 206 | 22783,754 | 3323 | 1 | 1 | 0 | 1 | 0 | 0 | 1 | 0 | 1 | 0 | 0 | 1 | 0 | 0 | 0 | 0 |
| 50.16483 | 20.08232 | MLP_R_5 | mlpr5_4 | Poland | R | 0 | 0 | 0 | 0 | 0 | 0 | 0 | 293,863 | 1227,244818 | 206 | 4808,869 | 6413 | 0 | 1 | 0 | 0 | 0 | 0 | 1 | 0 | 1 | 0 | 0 | 1 | 0 | 0 | 0 | 0 |
| 50.16315 | 20.08429 | MLP_R_5 | mlpr5_5 | Poland | R | 2 | 1 | 0 | 0 | 0 | 0 | 1 | 289,479 | 1227,007665 | 206 | 19409,841 | 7287 | 1 | 1 | 0 | 1 | 0 | 0 | 1 | 0 | 1 | 0 | 0 | 1 | 1 | 0 | 1 | 0 |
| 50.16108 | 20.08359 | MLP_R_5 | mlpr5_6 | Poland | R | 0 | 0 | 0 | 0 | 0 | 0 | 1 | 265,597 | 1226,858166 | 206 | 8184,269 | 7200 | 1 | 1 | 0 | 1 | 0 | 0 | 1 | 1 | 1 | 0 | 0 | 1 | 1 | 0 | 1 | 0 |
| 50.15923 | 20.08111 | MLP_R_5 | mlpr5_7 | Poland | R | 0 | 0 | 0 | 0 | 0 | 0 | 1 | 270,463 | 1226,807485 | 206 | 12040,633 | 4652 | 0 | 1 | 0 | 1 | 0 | 0 | 1 | 0 | 1 | 0 | 0 | 1 | 0 | 0 | 1 | 0 |
| 50.16059 | 20.0784 | MLP_R_5 | mlpr5_8 | Poland | R | 0 | 0 | 0 | 0 | 0 | 0 | 1 | 278,027 | 1227,046664 | 206 | 3930,191 | 4289 | 0 | 1 | 0 | 0 | 0 | 0 | 1 | 1 | 1 | 0 | 0 | 1 | 0 | 0 | 1 | 0 |
| 50.29705 | 20.04346 | MLP_R_6 | mlpr6_1 | Poland | R | 1 | 1 | 0 | 0 | 0 | 0 | 0 | 251 | 1240,523674 | 199 | 23317,36 | 1644 | 1 | 1 | 1 | 1 | 0 | 0 | 0 | 0 | 1 | 1 | 0 | 1 | 0 | 0 | 0 | 1 |
| 50.29857 | 20.04692 | MLP_R_6 | mlpr6_2 | Poland | R | 2 | 1 | 0 | 0 | 0 | 0 | 1 | 252,94 | 1240,509056 | 199 | 24804,615 | 1761 | 1 | 1 | 0 | 1 | 0 | 0 | 0 | 1 | 1 | 1 | 1 | 1 | 0 | 0 | 0 | 1 |
| 50.30095 | 20.04911 | MLP_R_6 | mlpr6_3 | Poland | R | 0 | 0 | 0 | 0 | 0 | 0 | 1 | 253 | 1240,619018 | 199 | 19424,442 | 795 | 1 | 1 | 0 | 1 | 0 | 0 | 1 | 1 | 1 | 1 | 0 | 1 | 0 | 0 | 0 | 1 |
| 50.3032 | 20.05128 | MLP_R_6 | mlpr6_4 | Poland | R | 1 | 1 | 0 | 0 | 0 | 0 | 0 | 255,52 | 1240,725357 | 199 | 9481,879 | 1735 | 1 | 1 | 0 | 1 | 0 | 0 | 0 | 1 | 1 | 1 | 1 | 1 | 0 | 0 | 0 | 0 |
| 50.30617 | 20.05123 | MLP_R_6 | mlpr6_5 | Poland | R | 1 | 1 | 0 | 0 | 0 | 0 | 0 | 258,235 | 1240,981557 | 199 | 12398,198 | 1035 | 1 | 1 | 0 | 1 | 0 | 0 | 1 | 1 | 1 | 1 | 1 | 0 | 0 | 0 | 0 | 0 |
| 50.30868 | 20.05124 | MLP_R_6 | mlpr6_6 | Poland | R | 1 | 1 | 2 | 1 | 0 | 0 | 1 | 258,248 | 1241,204134 | 199 | 29542,323 | 1697 | 1 | 1 | 0 | 1 | 0 | 0 | 1 | 0 | 1 | 1 | 0 | 1 | 0 | 0 | 0 | 0 |
| 50.31112 | 20.05263 | MLP_R_6 | mlpr6_7 | Poland | R | 1 | 1 | 0 | 0 | 0 | 0 | 0 | 261,628 | 1241,357426 | 199 | 21728,601 | 1451 | 1 | 1 | 0 | 1 | 0 | 0 | 0 | 0 | 1 | 1 | 0 | 1 | 0 | 0 | 0 | 0 |
| 50.31304 | 20.05426 | MLP_R_6 | mlpr6_8 | Poland | R | 2 | 1 | 0 | 0 | 0 | 0 | 1 | 263 | 1241,453815 | 199 | 8189,322 | 1451 | 1 | 1 | 0 | 1 | 0 | 0 | 0 | 0 | 1 | 1 | 0 | 1 | 0 | 0 | 0 | 1 |
| 49.98146 | 19.63878 | MLP_R_7 | mlpr7_1 | Poland | R | 0 | 0 | 0 | 0 | 0 | 0 | 1 | 214,663 | 1231,576357 | 186 | 24632,269 | 0 | 0 | 1 | 0 | 1 | 0 | 0 | 1 | 0 | 0 | 1 | 0 | 0 | 0 | 0 | 0 | 1 |
| 49.97919 | 19.63659 | MLP_R_7 | mlpr7_2 | Poland | R | 1 | 1 | 0 | 0 | 0 | 0 | 0 | 215,219 | 1231,481922 | 186 | 16207,204 | 176 | 0 | 1 | 0 | 0 | 0 | 0 | 1 | 0 | 0 | 1 | 1 | 0 | 0 | 0 | 0 | 1 |
| 49.9779 | 19.63609 | MLP_R_7 | mlpr7_3 | Poland | R | 0 | 0 | 0 | 0 | 0 | 0 | 0 | 214 | 1231,396109 | 186 | 31804,88 | 1165 | 0 | 1 | 0 | 0 | 0 | 0 | 1 | 0 | 0 | 1 | 1 | 0 | 0 | 0 | 0 | 0 |
| 49.97873 | 19.6401 | MLP_R_7 | mlpr7_4 | Poland | R | 4 | 1 | 0 | 0 | 0 | 0 | 0 | 213,428 | 1231,278847 | 186 | 23828,724 | 384 | 0 | 1 | 0 | 0 | 0 | 0 | 1 | 1 | 1 | 0 | 1 | 0 | 0 | 0 | 0 | 1 |
| 49.97852 | 19.64415 | MLP_R_7 | mlpr7_5 | Poland | R | 1 | 1 | 0 | 0 | 0 | 0 | 0 | 214,344 | 1231,074331 | 186 | 8412,095 | 927 | 0 | 1 | 0 | 0 | 0 | 0 | 1 | 1 | 1 | 0 | 1 | 0 | 0 | 0 | 0 | 0 |
| 49.98039 | 19.64628 | MLP_R_7 | mlpr7_6 | Poland | R | 0 | 0 | 0 | 0 | 0 | 0 | 1 | 211,042 | 1231,131863 | 186 | 24543,328 | 510 | 0 | 1 | 0 | 0 | 0 | 0 | 0 | 1 | 0 | 0 | 1 | 0 | 0 | 0 | 0 | 0 |
| 49.98176 | 19.64948 | MLP_R_7 | mlpr7_7 | Poland | R | 0 | 0 | 0 | 0 | 0 | 0 | 0 | 213 | 1231,100687 | 186 | 16151,586 | 1016 | 1 | 1 | 0 | 1 | 0 | 0 | 0 | 1 | 1 | 1 | 1 | 0 | 0 | 0 | 0 | 1 |
| 49.98453 | 19.64883 | MLP_R_7 | mlpr7_8 | Poland | R | 1 | 1 | 0 | 0 | 0 | 0 | 1 | 214,399 | 1231,365423 | 186 | 31764,633 | 320 | 1 | 1 | 0 | 1 | 0 | 0 | 0 | 1 | 0 | 1 | 1 | 0 | 0 | 0 | 0 | 1 |
| 50.01073 | 19.51424 | MLP_R_8 | mlpr8_1 | Poland | R | 1 | 1 | 0 | 0 | 0 | 0 | 1 | 219 | 1239,852803 | 176 | 23713,896 | 792 | 1 | 1 | 0 | 1 | 0 | 0 | 1 | 0 | 0 | 0 | 1 | 0 | 0 | 0 | 0 | 1 |
| 50.01343 | 19.5128 | MLP_R_8 | mlpr8_2 | Poland | R | 0 | 0 | 0 | 0 | 0 | 0 | 0 | 217,84 | 1240,132555 | 176 | 6270,844 | 0 | 0 | 1 | 0 | 0 | 0 | 0 | 1 | 0 | 0 | 1 | 0 | 0 | 0 | 0 | 0 | 0 |
| 50.0163 | 19.51323 | MLP_R_8 | mlpr8_3 | Poland | R | 1 | 1 | 0 | 0 | 0 | 0 | 0 | 216,625 | 1240,362308 | 176 | 16052,149 | 1722 | 1 | 1 | 0 | 1 | 0 | 0 | 1 | 0 | 1 | 1 | 0 | 1 | 0 | 0 | 0 | 0 |
| 50.01878 | 19.51556 | MLP_R_8 | mlpr8_4 | Poland | R | 0 | 0 | 0 | 0 | 0 | 0 | 1 | 216,608 | 1240,470334 | 176 | 3973,735 | 0 | 1 | 1 | 0 | 1 | 0 | 0 | 1 | 0 | 0 | 1 | 0 | 0 | 0 | 0 | 0 | 1 |
| 50.02027 | 19.51936 | MLP_R_8 | mlpr8_5 | Poland | R | 2 | 1 | 0 | 0 | 0 | 0 | 0 | 217,944 | 1240,413202 | 176 | 7274,484 | 0 | 1 | 1 | 0 | 1 | 0 | 0 | 1 | 0 | 0 | 1 | 0 | 0 | 0 | 0 | 0 | 0 |
| 50.02083 | 19.52366 | MLP_R_8 | mlpr8_6 | Poland | R | 1 | 1 | 0 | 0 | 0 | 0 | 1 | 216,814 | 1240,259455 | 176 | 22404,855 | 0 | 1 | 1 | 0 | 1 | 0 | 0 | 0 | 0 | 0 | 0 | 0 | 0 | 0 | 0 | 0 | 0 |
| 50.02033 | 19.52771 | MLP_R_8 | mlpr8_7 | Poland | R | 0 | 0 | 0 | 0 | 0 | 0 | 0 | 218,092 | 1240,029808 | 176 | 5475,121 | 0 | 1 | 1 | 0 | 1 | 0 | 0 | 1 | 0 | 0 | 1 | 0 | 0 | 0 | 0 | 0 | 0 |
| 50.01835 | 19.52978 | MLP_R_8 | mlpr8_8 | Poland | R | 1 | 1 | 0 | 0 | 0 | 0 | 0 | 216,84 | 1239,765097 | 176 | 12245,499 | 1150 | 0 | 1 | 0 | 1 | 0 | 0 | 1 | 0 | 1 | 1 | 1 | 0 | 0 | 0 | 0 | 1 |
| 49.97527 | 20.26274 | MLP_R_9 | mlpr9_1 | Poland | R | 0 | 0 | 0 | 0 | 0 | 0 | 1 | 262,484 | 1202,690369 | 224 | 16594,19 | 3551 | 0 | 1 | 0 | 0 | 0 | 0 | 1 | 0 | 1 | 0 | 0 | 1 | 0 | 0 | 0 | 0 |
| 49.97665 | 20.2596 | MLP_R_9 | mlpr9_2 | Poland | R | 0 | 0 | 0 | 0 | 0 | 0 | 0 | 273,034 | 1202,953645 | 224 | 5066,791 | 4823 | 0 | 1 | 0 | 0 | 0 | 0 | 1 | 0 | 1 | 0 | 1 | 0 | 1 | 0 | 1 | 0 |
| 49.97375 | 20.26186 | MLP_R_9 | mlpr9_3 | Poland | R | 2 | 1 | 0 | 0 | 0 | 0 | 1 | 261,696 | 1202,593167 | 224 | 24507,46 | 2052 | 1 | 1 | 0 | 0 | 1 | 0 | 1 | 0 | 0 | 0 | 1 | 0 | 0 | 0 | 0 | 0 |
| 49.97243 | 20.26448 | MLP_R_9 | mlpr9_4 | Poland | R | 0 | 0 | 0 | 0 | 0 | 0 | 0 | 246,309 | 1202,36362 | 224 | 4452,937 | 4451 | 0 | 1 | 0 | 0 | 0 | 0 | 1 | 0 | 0 | 0 | 1 | 0 | 0 | 0 | 1 | 0 |

| | | | | | | | | | | | | | | | | | | | | | | | | | | | | | | | | | |
|---|---|---|---|---|---|---|---|---|---|---|---|---|---|---|---|---|---|---|---|---|---|---|---|---|---|---|---|---|---|---|---|---|---|
| 49.97299 | 20.26843 | MLP_R_9 | mlpr9_5 | Poland | R | 0 | 0 | 2 | 1 | 0 | 0 | 1 | 236,652 | 1202,236075 | 224 | 11786,572 | 1981 | 0 | 1 | 0 | 0 | 0 | 0 | 1 | 0 | 1 | 0 | 1 | 0 | 0 | 0 | 0 | 0 |
| 49.97265 | 20.27246 | MLP_R_9 | mlpr9_6 | Poland | R | 0 | 0 | 0 | 0 | 0 | 0 | 0 | 228,066 | 1202,027081 | 224 | 3512,895 | 1783 | 0 | 1 | 0 | 0 | 0 | 0 | 1 | 0 | 1 | 0 | 0 | 1 | 0 | 0 | 0 | 0 |
| 49.96971 | 20.27134 | MLP_R_9 | mlpr9_7 | Poland | R | 0 | 0 | 0 | 0 | 0 | 0 | 0 | 222,956 | 1201,82131 | 224 | 1815,341 | 0 | 0 | 1 | 0 | 0 | 0 | 0 | 1 | 0 | 0 | 0 | 0 | 0 | 0 | 0 | 0 | 1 |
| 49.9686 | 20.2673 | MLP_R_9 | mlpr9_8 | Poland | R | 0 | 0 | 1 | 1 | 0 | 0 | 0 | 225 | 1201,904075 | 224 | 2788,807 | 0 | 0 | 1 | 0 | 0 | 0 | 0 | 1 | 0 | 0 | 0 | 0 | 0 | 0 | 0 | 0 | 0 |
| 50.0585 | 19.93384 | MLP_U_0 | mlpu0_1 | Poland | U | 0 | 0 | 0 | 0 | 0 | 0 | 0 | 205 | 1224,649527 | 201 | 31093 | 26000 | 1 | 0 | 0 | 0 | 1 | 0 | 0 | 1 | 0 | 0 | 0 | 0 | 0 | 1 | 0 | 0 |
| 50.06167 | 19.93249 | MLP_U_0 | mlpu0_2 | Poland | U | 0 | 0 | 0 | 0 | 0 | 0 | 0 | 207,012 | 1224,98411 | 201 | 19550,5 | 44600 | 1 | 0 | 0 | 0 | 1 | 0 | 0 | 0 | 0 | 0 | 0 | 0 | 0 | 1 | 0 | 0 |
| 50.06446 | 19.93483 | MLP_U_0 | mlpu0_3 | Poland | U | 0 | 0 | 0 | 0 | 0 | 0 | 0 | 209,366 | 1225,11942 | 201 | 24242,5 | 39600 | 1 | 0 | 0 | 0 | 1 | 0 | 0 | 0 | 0 | 0 | 0 | 0 | 0 | 1 | 0 | 0 |
| 50.06552 | 19.93966 | MLP_U_0 | mlpu0_4 | Poland | U | 1 | 1 | 0 | 0 | 0 | 0 | 0 | 213,901 | 1224,993601 | 201 | 25547,295 | 36100 | 1 | 0 | 0 | 0 | 1 | 0 | 0 | 0 | 0 | 0 | 0 | 0 | 0 | 1 | 0 | 0 |
| 50.06416 | 19.94367 | MLP_U_0 | mlpu0_5 | Poland | U | 2 | 1 | 0 | 0 | 0 | 0 | 1 | 213,019 | 1224,695116 | 201 | 23336,474 | 29500 | 1 | 0 | 0 | 0 | 1 | 0 | 0 | 0 | 0 | 0 | 0 | 0 | 0 | 1 | 0 | 0 |
| 50.06131 | 19.9438 | MLP_U_0 | mlpu0_6 | Poland | U | 0 | 0 | 0 | 0 | 0 | 0 | 0 | 210,432 | 1224,442702 | 201 | 23397,572 | 38000 | 1 | 0 | 0 | 0 | 1 | 0 | 0 | 1 | 0 | 0 | 0 | 0 | 0 | 1 | 0 | 0 |
| 50.05911 | 19.94064 | MLP_U_0 | mlpu0_7 | Poland | U | 0 | 0 | 0 | 0 | 1 | 1 | 0 | 206,492 | 1224,395178 | 201 | 24654,273 | 38500 | 1 | 0 | 0 | 0 | 1 | 0 | 0 | 1 | 0 | 0 | 0 | 0 | 0 | 1 | 0 | 0 |
| 50.05638 | 19.93984 | MLP_U_0 | mlpu0_8 | Poland | U | 2 | 1 | 0 | 0 | 0 | 0 | 0 | 205,134 | 1224,185566 | 201 | 16498 | 43600 | 1 | 1 | 0 | 0 | 1 | 0 | 0 | 1 | 0 | 0 | 0 | 0 | 0 | 1 | 0 | 0 |
| 50.01866 | 19.99565 | MLP_U_1 | mlpu1_1 | Poland | U | 2 | 1 | 0 | 0 | 2 | 1 | 1 | 213,116 | 1218,418904 | 207 | 46324,456 | 7800 | 1 | 1 | 1 | 0 | 1 | 0 | 0 | 1 | 1 | 1 | 1 | 1 | 1 | 0 | 0 | 0 |
| 50.02052 | 19.99841 | MLP_U_1 | mlpu1_2 | Poland | U | 0 | 0 | 0 | 0 | 2 | 1 | 1 | 208,071 | 1218,455045 | 207 | 21425,895 | 13500 | 1 | 1 | 1 | 0 | 1 | 0 | 0 | 1 | 1 | 1 | 0 | 1 | 1 | 0 | 0 | 1 |
| 50.02111 | 20.00115 | MLP_U_1 | mlpu1_3 | Poland | U | 2 | 1 | 1 | 1 | 0 | 0 | 0 | 207 | 1218,383982 | 207 | 36118,315 | 7300 | 1 | 1 | 1 | 1 | 1 | 0 | 0 | 1 | 1 | 1 | 0 | 1 | 0 | 0 | 0 | 0 |
| 50.02032 | 20.00519 | MLP_U_1 | mlpu1_4 | Poland | U | 2 | 1 | 0 | 0 | 0 | 0 | 1 | 207,58 | 1218,131975 | 207 | 38743,227 | 7500 | 1 | 1 | 1 | 0 | 1 | 0 | 0 | 1 | 0 | 1 | 0 | 0 | 1 | 0 | 0 | 0 |
| 50.01923 | 20.00914 | MLP_U_1 | mlpu1_5 | Poland | U | 1 | 1 | 0 | 0 | 0 | 0 | 0 | 212,984 | 1217,865761 | 207 | 39716,816 | 6600 | 1 | 1 | 0 | 1 | 0 | 0 | 1 | 1 | 0 | 0 | 1 | 0 | 1 | 0 | 0 | 1 |
| 50.01736 | 20.01233 | MLP_U_1 | mlpu1_6 | Poland | U | 1 | 1 | 0 | 0 | 0 | 0 | 0 | 214,572 | 1217,558387 | 207 | 31152,956 | 16800 | 1 | 1 | 1 | 1 | 0 | 0 | 1 | 1 | 0 | 0 | 0 | 0 | 1 | 0 | 1 | 0 |
| 50.01484 | 20.01373 | MLP_U_1 | mlpu1_7 | Poland | U | 2 | 1 | 0 | 0 | 1 | 1 | 1 | 222,428 | 1217,27239 | 207 | 38303,054 | 19400 | 1 | 1 | 1 | 1 | 1 | 0 | 1 | 1 | 0 | 1 | 0 | 0 | 1 | 0 | 0 | 0 |
| 50.01214 | 20.01426 | MLP_U_1 | mlpu1_8 | Poland | U | 2 | 1 | 0 | 0 | 2 | 1 | 1 | 221,336 | 1217,019266 | 207 | 52502,226 | 10400 | 1 | 1 | 1 | 1 | 1 | 0 | 0 | 1 | 1 | 0 | 0 | 0 | 0 | 0 | 1 | 0 |
| 50.08495 | 19.92136 | MLP_U_10 | mlpu10_1 | Poland | U | 2 | 1 | 0 | 0 | 0 | 0 | 1 | 226,546 | 1227,495595 | 198 | 37497,341 | 11200 | 1 | 1 | 1 | 0 | 1 | 0 | 0 | 1 | 1 | 0 | 0 | 1 | 1 | 0 | 0 | 0 |
| 50.08774 | 19.92114 | MLP_U_10 | mlpu10_2 | Poland | U | 0 | 0 | 0 | 0 | 0 | 0 | 1 | 224,136 | 1227,749788 | 198 | 33973,836 | 3500 | 0 | 1 | 0 | 0 | 1 | 0 | 0 | 1 | 1 | 0 | 0 | 1 | 0 | 0 | 0 | 0 |
| 50.08893 | 19.91919 | MLP_U_10 | mlpu10_3 | Poland | U | 0 | 0 | 0 | 0 | 0 | 0 | 0 | 224,852 | 1227,937545 | 198 | 25265,995 | 17300 | 1 | 1 | 0 | 0 | 0 | 0 | 0 | 1 | 0 | 0 | 0 | 0 | 1 | 0 | 0 | 1 |
| 50.08823 | 19.91591 | MLP_U_10 | mlpu10_4 | Poland | U | 0 | 0 | 1 | 1 | 0 | 0 | 0 | 224,827 | 1228,025699 | 198 | 15124,237 | 32300 | 0 | 1 | 0 | 0 | 0 | 0 | 0 | 1 | 0 | 0 | 0 | 1 | 1 | 0 | 0 | 0 |
| 50.08686 | 19.91346 | MLP_U_10 | mlpu10_5 | Poland | U | 2 | 1 | 0 | 0 | 1 | 1 | 1 | 222,696 | 1228,017028 | 198 | 15292,103 | 27400 | 1 | 1 | 0 | 0 | 0 | 0 | 0 | 1 | 0 | 0 | 0 | 1 | 1 | 0 | 0 | 0 |
| 50.08646 | 19.91094 | MLP_U_10 | mlpu10_6 | Poland | U | 0 | 0 | 0 | 0 | 0 | 0 | 0 | 223 | 1228,096805 | 198 | 15710,774 | 29600 | 0 | 1 | 0 | 0 | 0 | 0 | 0 | 1 | 0 | 0 | 0 | 1 | 1 | 0 | 0 | 0 |
| 50.08405 | 19.91134 | MLP_U_10 | mlpu10_7 | Poland | U | 0 | 0 | 0 | 0 | 0 | 0 | 0 | 219,102 | 1227,869819 | 198 | 7320,771 | 31900 | 0 | 1 | 0 | 0 | 0 | 0 | 0 | 1 | 0 | 0 | 0 | 1 | 1 | 0 | 0 | 0 |
| 50.08284 | 19.91478 | MLP_U_10 | mlpu10_8 | Poland | U | 0 | 0 | 0 | 0 | 0 | 0 | 0 | 216 | 1227,611204 | 198 | 5099,241 | 23500 | 0 | 1 | 0 | 0 | 0 | 0 | 0 | 1 | 0 | 0 | 0 | 1 | 1 | 0 | 0 | 0 |
| 50.0665 | 19.99409 | MLP_U_11 | mlpu11_1 | Poland | U | 2 | 1 | 0 | 0 | 1 | 1 | 1 | 199,6 | 1222,625661 | 204 | 20524,7 | 7100 | 0 | 1 | 0 | 0 | 1 | 0 | 1 | 1 | 1 | 0 | 0 | 1 | 1 | 0 | 0 | 0 |
| 50.06789 | 19.99583 | MLP_U_11 | mlpu11_2 | Poland | U | 2 | 1 | 0 | 0 | 0 | 0 | 1 | 200,399 | 1222,668914 | 204 | 16394,869 | 100 | 0 | 1 | 0 | 0 | 1 | 0 | 0 | 0 | 0 | 0 | 0 | 0 | 0 | 0 | 0 | 0 |
| 50.07033 | 19.99529 | MLP_U_11 | mlpu11_3 | Poland | U | 1 | 1 | 0 | 0 | 1 | 1 | 1 | 206 | 1222,906816 | 204 | 55873,168 | 0 | 1 | 1 | 1 | 0 | 1 | 0 | 0 | 0 | 0 | 0 | 0 | 0 | 0 | 0 | 0 | 1 |
| 50.07313 | 19.99534 | MLP_U_11 | mlpu11_4 | Poland | U | 1 | 1 | 0 | 0 | 1 | 1 | 1 | 208 | 1223,146192 | 204 | 36080,784 | 4000 | 1 | 1 | 1 | 0 | 1 | 0 | 1 | 1 | 1 | 0 | 0 | 0 | 1 | 0 | 0 | 0 |
| 50.07471 | 19.99246 | MLP_U_11 | mlpu11_5 | Poland | U | 2 | 1 | 0 | 0 | 0 | 0 | 1 | 213,899 | 1223,410108 | 204 | 34295,929 | 13700 | 1 | 1 | 0 | 1 | 1 | 0 | 0 | 1 | 0 | 0 | 0 | 0 | 1 | 0 | 1 | 0 |
| 50.07442 | 19.98898 | MLP_U_11 | mlpu11_6 | Poland | U | 0 | 0 | 0 | 0 | 0 | 0 | 1 | 211,182 | 1223,542375 | 204 | 38623,714 | 9600 | 1 | 1 | 1 | 0 | 1 | 0 | 1 | 1 | 0 | 0 | 0 | 0 | 1 | 0 | 1 | 0 |
| 50.0723 | 19.99098 | MLP_U_11 | mlpu11_7 | Poland | U | 2 | 1 | 0 | 0 | 0 | 0 | 1 | 208,468 | 1223,271665 | 204 | 61621,18 | 300 | 1 | 0 | 1 | 0 | 1 | 0 | 0 | 0 | 0 | 0 | 0 | 0 | 0 | 0 | 0 | 0 |
| 50.07022 | 19.98961 | MLP_U_11 | mlpu11_8 | Poland | U | 0 | 0 | 0 | 0 | 0 | 0 | 1 | 201 | 1223,14954 | 204 | 69644,239 | 200 | 1 | 1 | 1 | 0 | 1 | 0 | 0 | 0 | 0 | 0 | 1 | 0 | 0 | 0 | 0 | 0 |
| 50.045 | 19.99165 | MLP_U_12 | mlpu12_1 | Poland | U | 2 | 1 | 0 | 0 | 0 | 0 | 1 | 200 | 1220,882351 | 205 | 19417,309 | 18200 | 0 | 1 | 0 | 0 | 1 | 0 | 0 | 1 | 1 | 0 | 1 | 0 | 1 | 0 | 0 | 0 |
| 50.04683 | 19.99336 | MLP_U_12 | mlpu12_2 | Poland | U | 2 | 1 | 0 | 0 | 0 | 0 | 1 | 199,096 | 1220,962233 | 205 | 34815,291 | 10800 | 1 | 1 | 0 | 0 | 1 | 0 | 0 | 1 | 0 | 0 | 1 | 0 | 1 | 0 | 0 | 1 |
| 50.04929 | 19.99173 | MLP_U_12 | mlpu12_3 | Poland | U | 1 | 1 | 0 | 0 | 1 | 1 | 1 | 199,556 | 1221,255118 | 205 | 32730,601 | 9500 | 1 | 1 | 0 | 0 | 1 | 0 | 0 | 1 | 0 | 0 | 1 | 0 | 1 | 0 | 0 | 0 |
| 50.05113 | 19.99395 | MLP_U_12 | mlpu12_4 | Poland | U | 1 | 1 | 0 | 0 | 0 | 0 | 1 | 200 | 1221,302725 | 205 | 20741,187 | 3700 | 1 | 1 | 0 | 1 | 1 | 0 | 0 | 1 | 0 | 0 | 0 | 0 | 1 | 0 | 0 | 0 |
| 50.04987 | 19.99691 | MLP_U_12 | mlpu12_5 | Poland | U | 1 | 1 | 0 | 0 | 0 | 0 | 1 | 200,476 | 1221,039258 | 205 | 16186,296 | 0 | 0 | 1 | 0 | 1 | 1 | 0 | 0 | 1 | 0 | 0 | 0 | 0 | 0 | 0 | 0 | 0 |
| 50.0479 | 20.00006 | MLP_U_12 | mlpu12_6 | Poland | U | 0 | 0 | 0 | 0 | 0 | 0 | 1 | 200,56 | 1220,72926 | 205 | 11862,237 | 0 | 0 | 1 | 0 | 1 | 0 | 0 | 0 | 0 | 0 | 0 | 0 | 0 | 0 | 0 | 0 | 1 |
| 50.0461 | 20.00325 | MLP_U_12 | mlpu12_7 | Poland | U | 2 | 1 | 0 | 0 | 0 | 0 | 1 | 200,632 | 1220,445236 | 205 | 27923,491 | 2500 | 0 | 1 | 0 | 1 | 0 | 0 | 0 | 0 | 1 | 0 | 1 | 0 | 0 | 0 | 0 | 0 |
| 50.04347 | 20.00281 | MLP_U_12 | mlpu12_8 | Poland | U | 1 | 1 | 0 | 0 | 0 | 0 | 1 | 198 | 1220,242543 | 205 | 22455,214 | 4600 | 1 | 1 | 0 | 0 | 1 | 0 | 0 | 1 | 1 | 0 | 1 | 0 | 0 | 0 | 0 | 0 |
| 50.04328 | 19.94062 | MLP_U_14 | mlpu14_1 | Poland | U | 0 | 0 | 0 | 0 | 1 | 1 | 0 | 204,379 | 1223,029658 | 202 | 22445,928 | 14900 | 1 | 1 | 0 | 0 | 1 | 0 | 0 | 1 | 0 | 1 | 0 | 0 | 1 | 1 | 0 | 0 |
| 50.04131 | 19.9426 | MLP_U_14 | mlpu14_2 | Poland | U | 0 | 0 | 0 | 0 | 0 | 0 | 0 | 204 | 1222,770257 | 202 | 9637,287 | 47400 | 0 | 1 | 0 | 0 | 0 | 0 | 0 | 1 | 0 | 0 | 0 | 0 | 1 | 1 | 0 | 0 |
| 50.0393 | 19.94391 | MLP_U_14 | mlpu14_3 | Poland | U | 0 | 0 | 0 | 0 | 0 | 0 | 0 | 220,92 | 1222,537435 | 202 | 32210,416 | 20700 | 1 | 1 | 0 | 0 | 1 | 0 | 0 | 1 | 0 | 0 | 0 | 0 | 0 | 1 | 0 | 0 |
| 50.04172 | 19.9471 | MLP_U_14 | mlpu14_4 | Poland | U | 2 | 1 | 0 | 0 | 0 | 0 | 0 | 221,46 | 1222,602329 | 202 | 42188,666 | 23000 | 1 | 0 | 0 | 0 | 1 | 0 | 0 | 1 | 0 | 0 | 0 | 0 | 0 | 1 | 0 | 0 |
| 50.04181 | 19.95045 | MLP_U_14 | mlpu14_5 | Poland | U | 0 | 0 | 0 | 0 | 0 | 0 | 0 | 222,64 | 1222,458584 | 202 | 43079,696 | 8000 | 1 | 1 | 0 | 0 | 1 | 0 | 0 | 1 | 0 | 0 | 0 | 0 | 0 | 1 | 0 | 0 |
| 50.04255 | 19.95408 | MLP_U_14 | mlpu14_6 | Poland | U | 0 | 0 | 0 | 0 | 0 | 0 | 0 | 229,384 | 1222,358042 | 202 | 28055,228 | 25400 | 1 | 1 | 0 | 1 | 0 | 0 | 0 | 1 | 1 | 0 | 0 | 0 | 1 | 0 | 0 | 0 |
| 50.04372 | 19.95762 | MLP_U_14 | mlpu14_7 | Poland | U | 0 | 0 | 0 | 0 | 0 | 0 | 0 | 210,019 | 1222,301188 | 202 | 26011,648 | 26300 | 1 | 0 | 0 | 1 | 1 | 0 | 0 | 1 | 0 | 0 | 0 | 1 | 0 | 1 | 0 | 0 |
| 50.04471 | 19.96011 | MLP_U_14 | mlpu14_8 | Poland | U | 0 | 0 | 0 | 0 | 0 | 0 | 0 | 201,968 | 1222,272303 | 202 | 7050,556 | 36500 | 0 | 1 | 0 | 0 | 0 | 0 | 0 | 1 | 0 | 0 | 0 | 0 | 0 | 1 | 1 | 0 |
| 50.059 | 19.87046 | MLP_U_15 | mlpu15_1 | Poland | U | 0 | 0 | 0 | 0 | 0 | 0 | 0 | 226,088 | 1227,562396 | 196 | 25801,415 | 11500 | 1 | 0 | 1 | 1 | 0 | 0 | 0 | 0 | 1 | 0 | 0 | 1 | 0 | 0 | 0 | 0 |
| 50.06106 | 19.87081 | MLP_U_15 | mlpu15_2 | Poland | U | 0 | 0 | 0 | 0 | 0 | 0 | 1 | 222,153 | 1227,72381 | 196 | 13584,597 | 15700 | 0 | 1 | 1 | 0 | 0 | 0 | 0 | 0 | 1 | 0 | 0 | 1 | 0 | 0 | 0 | 0 |
| 50.06326 | 19.87245 | MLP_U_15 | mlpu15_3 | Poland | U | 3 | 1 | 0 | 0 | 0 | 0 | 1 | 217,3 | 1227,838255 | 196 | 40116,625 | 11500 | 1 | 1 | 1 | 0 | 1 | 0 | 0 | 0 | 1 | 0 | 0 | 1 | 0 | 0 | 0 | 0 |
| 50.06554 | 19.87322 | MLP_U_15 | mlpu15_4 | Poland | U | 2 | 1 | 0 | 0 | 0 | 0 | 1 | 210,056 | 1227,999247 | 196 | 37833,495 | 6200 | 1 | 1 | 1 | 0 | 1 | 0 | 0 | 0 | 1 | 0 | 0 | 1 | 1 | 0 | 0 | 0 |
| 50.06684 | 19.87117 | MLP_U_15 | mlpu15_5 | Poland | U | 1 | 1 | 0 | 0 | 0 | 0 | 0 | 208,624 | 1228,203266 | 196 | 19481,073 | 17300 | 0 | 1 | 1 | 0 | 1 | 0 | 0 | 1 | 1 | 0 | 0 | 1 | 1 | 0 | 0 | 0 |
| 50.06685 | 19.86703 | MLP_U_15 | mlpu15_6 | Poland | U | 0 | 0 | 0 | 0 | 0 | 0 | 1 | 223,032 | 1228,391559 | 196 | 22178,061 | 18500 | 0 | 1 | 1 | 0 | 0 | 0 | 0 | 1 | 1 | 0 | 0 | 1 | 1 | 0 | 0 | 0 |
| 50.06683 | 19.86276 | MLP_U_15 | mlpu15_7 | Poland | U | 1 | 1 | 0 | 0 | 1 | 1 | 0 | 220,962 | 1228,583418 | 196 | 20668,837 | 17700 | 1 | 1 | 1 | 0 | 0 | 0 | 1 | 1 | 1 | 0 | 0 | 1 | 1 | 1 | 0 | 1 |
| 50.06415 | 19.86143 | MLP_U_15 | mlpu15_8 | Poland | U | 2 | 1 | 0 | 0 | 0 | 0 | 0 | 228,99 | 1228,415623 | 196 | 30936,587 | 9100 | 1 | 1 | 1 | 1 | 0 | 0 | 0 | 1 | 1 | 1 | 0 | 1 | 0 | 0 | 0 | 1 |
| 50.09554 | 20.00582 | MLP_U_16 | mlpu16_1 | Poland | U | 1 | 1 | 0 | 0 | 0 | 0 | 0 | 246,045 | 1225,696355 | 203 | 38316,478 | 10800 | 1 | 1 | 0 | 1 | 1 | 0 | 0 | 1 | 0 | 0 | 0 | 0 | 1 | 0 | 0 | 1 |
| 50.09778 | 20.00646 | MLP_U_16 | mlpu16_2 | Poland | U | 2 | 1 | 0 | 0 | 0 | 0 | 0 | 248,482 | 1225,583346 | 203 | 43660,239 | 14200 | 1 | 1 | 0 | 1 | 0 | 0 | 0 | 1 | 0 | 0 | 0 | 0 | 0 | 1 | 1 | 1 |
| 50.09937 | 20.00428 | MLP_U_16 | mlpu16_3 | Poland | U | 1 | 1 | 1 | 1 | 1 | 1 | 0 | 257,184 | 1225,453845 | 203 | 32184,492 | 16300 | 1 | 1 | 0 | 1 | 0 | 0 | 0 | 1 | 1 | 0 | 0 | 0 | 1 | 1 | 0 | 0 |
| 50.10126 | 20.00167 | MLP_U_16 | mlpu16_4 | Poland | U | 2 | 1 | 0 | 0 | 0 | 0 | 1 | 270,75 | 1225,45145 | 203 | 43948,708 | 4400 | 1 | 1 | 0 | 0 | 1 | 0 | 0 | 1 | 1 | 0 | 0 | 1 | 1 | 0 | 0 | 0 |
| 50.10064 | 19.99708 | MLP_U_16 | mlpu16_5 | Poland | U | 3 | 1 | 1 | 1 | 0 | 0 | 1 | 274,524 | 1225,299354 | 203 | 46983,916 | 600 | 1 | 1 | 0 | 0 | 1 | 0 | 0 | 1 | 1 | 0 | 0 | 1 | 0 | 0 | 0 | 1 |
| 50.09911 | 19.99404 | MLP_U_16 | mlpu16_6 | Poland | U | 1 | 1 | 0 | 0 | 0 | 0 | 0 | 258,655 | 1225,019339 | 203 | 25930,5 | 4800 | 0 | 1 | 0 | 0 | 1 | 0 | 0 | 1 | 0 | 0 | 0 | 0 | 1 | 0 | 1 | 1 |
| 50.09831 | 19.98966 | MLP_U_16 | mlpu16_7 | Poland | U | 1 | 1 | 0 | 0 | 0 | 0 | 1 | 227,084 | 1224,78297 | 203 | 24569,383 | 15900 | 0 | 1 | 0 | 0 | 0 | 0 | 0 | 1 | 0 | 0 | 0 | 0 | 1 | 0 | 0 | 1 |
| 50.09746 | 19.98539 | MLP_U_16 | mlpu16_8 | Poland | U | 1 | 1 | 0 | 0 | 0 | 0 | 1 | 224,202 | 1224,617422 | 203 | 22178,992 | 8300 | 0 | 1 | 0 | 0 | 1 | 0 | 0 | 1 | 0 | 0 | 0 | 0 | 1 | 0 | 0 | 0 |
| 50.0309 | 19.99091 | MLP_U_2 | mlpu2_1 | Poland | U | 1 | 1 | 0 | 0 | 1 | 1 | 0 | 200,07 | 1219,689621 | 206 | 7915,157 | 22200 | 0 | 1 | 0 | 0 | 0 | 0 | 1 | 0 | 0 | 1 | 0 | 0 | 0 | 0 | 1 | 0 |
| 50.03046 | 19.99468 | MLP_U_2 | mlpu2_2 | Poland | U | 0 | 0 | 0 | 0 | 0 | 0 | 1 | 199,084 | 1219,48232 | 206 | 6245,107 | 24600 | 0 | 1 | 0 | 0 | 0 | 0 | 1 | 0 | 0 | 1 | 0 | 0 | 0 | 0 | 1 | 1 |
| 50.03052 | 19.99945 | MLP_U_2 | mlpu2_3 | Poland | U | 0 | 0 | 0 | 0 | 0 | 0 | 0 | 199,145 | 1219,276495 | 206 | 13461,027 | 24500 | 0 | 1 | 0 | 0 | 0 | 0 | 1 | 1 | 0 | 1 | 0 | 0 | 0 | 0 | 1 | 0 |
| 50.0286 | 19.99908 | MLP_U_2 | mlpu2_4 | Poland | U | 0 | 0 | 0 | 0 | 0 | 0 | 0 | 201,012 | 1219,1174 | 206 | 23558,121 | 22300 | 1 | 1 | 0 | 1 | 0 | 0 | 1 | 1 | 1 | 0 | 1 | 0 | 0 | 0 | 1 | 0 |
| 50.02749 | 19.9968 | MLP_U_2 | mlpu2_5 | Poland | U | 0 | 0 | 0 | 0 | 0 | 0 | 0 | 202,889 | 1219,1238 | 206 | 23712,64 | 13500 | 1 | 1 | 0 | 1 | 0 | 0 | 1 | 1 | 1 | 0 | 1 | 1 | 0 | 0 | 1 | 1 |
| 50.02716 | 19.9934 | MLP_U_2 | mlpu2_6 | Poland | U | 0 | 0 | 1 | 1 | 0 | 0 | 0 | 201,822 | 1219,247084 | 206 | 27609,999 | 11000 | 0 | 1 | 0 | 1 | 0 | 0 | 1 | 1 | 1 | 0 | 0 | 1 | 1 | 0 | 1 | 0 |
| 50.02675 | 19.98903 | MLP_U_2 | mlpu2_7 | Poland | U | 0 | 0 | 2 | 1 | 0 | 0 | 0 | 204 | 1219,415493 | 206 | 26529,608 | 17300 | 1 | 1 | 1 | 0 | 1 | 0 | 1 | 1 | 1 | 0 | 0 | 1 | 1 | 0 | 0 | 0 |
| 50.02639 | 19.98542 | MLP_U_2 | mlpu2_8 | Poland | U | 1 | 1 | 0 | 0 | 0 | 0 | 1 | 205,992 | 1219,553542 | 206 | 17062,983 | 27400 | 1 | 1 | 0 | 1 | 0 | 0 | 0 | 1 | 1 | 0 | 0 | 1 | 0 | 1 | 1 | 0 |
| 50.04003 | 19.91735 | MLP_U_3 | mlpu3_1 | Poland | U | 0 | 0 | 0 | 0 | 0 | 0 | 1 | 209,935 | 1223,803614 | 201 | 48515,194 | 1200 | 1 | 1 | 1 | 1 | 0 | 0 | 1 | 0 | 1 | 0 | 1 | 0 | 0 | 0 | 0 | 1 |
| 50.03773 | 19.91535 | MLP_U_3 | mlpu3_2 | Poland | U | 0 | 0 | 0 | 0 | 0 | 0 | 1 | 209,955 | 1223,695009 | 201 | 42641,228 | 0 | 1 | 1 | 1 | 1 | 0 | 0 | 0 | 0 | 0 | 1 | 0 | 0 | 0 | 0 | 0 | 0 |
| 50.03536 | 19.91389 | MLP_U_3 | mlpu3_3 | Poland | U | 0 | 0 | 0 | 0 | 0 | 0 | 0 | 214,288 | 1223,558325 | 201 | 39510,634 | 0 | 1 | 1 | 1 | 1 | 0 | 0 | 0 | 0 | 1 | 0 | 1 | 0 | 0 | 0 | 0 | 1 |
| 50.03424 | 19.91698 | MLP_U_3 | mlpu3_4 | Poland | U | 0 | 0 | 0 | 0 | 0 | 0 | 0 | 206,23 | 1223,319711 | 201 | 14195,549 | 700 | 0 | 1 | 0 | 1 | 0 | 0 | 0 | 0 | 1 | 0 | 1 | 0 | 0 | 0 | 0 | 0 |
| 50.03394 | 19.92057 | MLP_U_3 | mlpu3_5 | Poland | U | 1 | 1 | 0 | 0 | 1 | 1 | 0 | 206,816 | 1223,131163 | 201 | 21984,466 | 14700 | 1 | 1 | 0 | 1 | 0 | 0 | 0 | 1 | 1 | 0 | 0 | 0 | 1 | 0 | 1 | 1 |
| 50.03416 | 19.9239 | MLP_U_3 | mlpu3_6 | Poland | U | 0 | 0 | 0 | 0 | 0 | 0 | 0 | 207,024 | 1223,0008 | 201 | 8707,362 | 33000 | 0 | 1 | 0 | 0 | 0 | 0 | 1 | 1 | 0 | 0 | 0 | 0 | 1 | 0 | 1 | 0 |
| 50.03588 | 19.92733 | MLP_U_3 | mlpu3_7 | Poland | U | 2 | 1 | 0 | 0 | 0 | 0 | 0 | 205,897 | 1222,992184 | 201 | 30459,245 | 10900 | 0 | 1 | 0 | 0 | 1 | 0 | 1 | 1 | 1 | 1 | 0 | 1 | 1 | 0 | 1 | 0 |
| 50.0374 | 19.93091 | MLP_U_3 | mlpu3_8 | Poland | U | 0 | 0 | 0 | 0 | 0 | 0 | 0 | 202,978 | 1222,962816 | 201 | 27869,123 | 2500 | 1 | 1 | 0 | 1 | 0 | 0 | 0 | 1 | 1 | 1 | 0 | 1 | 1 | 0 | 1 | 0 |
| 50.01751 | 19.80224 | MLP_U_4 | mlpu4_1 | Poland | U | 0 | 0 | 0 | 0 | 0 | 0 | 1 | 206,074 | 1227,111614 | 194 | 15900,837 | 0 | 1 | 1 | 1 | 1 | 0 | 0 | 1 | 1 | 0 | 1 | 0 | 1 | 0 | 1 | 0 | 1 |
| 50.01761 | 19.80566 | MLP_U_4 | mlpu4_2 | Poland | U | 0 | 0 | 0 | 0 | 0 | 0 | 0 | 209 | 1226,962495 | 194 | 23807,041 | 0 | 1 | 1 | 1 | 1 | 0 | 1 | 1 | 1 | 0 | 0 | 0 | 1 | 0 | 0 | 0 | 0 |
| 50.01981 | 19.80314 | MLP_U_4 | mlpu4_3 | Poland | U | 2 | 1 | 0 | 0 | 0 | 0 | 1 | 212,876 | 1227,260735 | 194 | 8652,982 | 0 | 1 | 1 | 1 | 1 | 1 | 0 | 1 | 1 | 1 | 1 | 0 | 1 | 0 | 1 | 0 | 1 |
| 50.0216 | 19.80423 | MLP_U_4 | mlpu4_4 | Poland | U | 2 | 1 | 0 | 0 | 0 | 0 | 1 | 207,994 | 1227,372312 | 194 | 11620,918 | 0 | 1 | 1 | 0 | 1 | 0 | 0 | 0 | 1 | 0 | 1 | 0 | 0 | 0 | 0 | 0 | 1 |
| 50.0236 | 19.80693 | MLP_U_4 | mlpu4_5 | Poland | U | 1 | 1 | 0 | 0 | 0 | 0 | 1 | 205,998 | 1227,417588 | 194 | 41886,861 | 1000 | 0 | 1 | 0 | 1 | 0 | 0 | 0 | 0 | 0 | 1 | 0 | 0 | 0 | 0 | 0 | 0 |
| 50.02509 | 19.81044 | MLP_U_4 | mlpu4_6 | Poland | U | 0 | 0 | 0 | 0 | 0 | 0 | 1 | 205,605 | 1227,381775 | 194 | 27826,652 | 10300 | 0 | 1 | 0 | 0 | 0 | 0 | 0 | 1 | 0 | 1 | 0 | 0 | 0 | 0 | 0 | 0 |
| 50.02642 | 19.81396 | MLP_U_4 | mlpu4_7 | Poland | U | 3 | 1 | 0 | 0 | 0 | 0 | 0 | 205,971 | 1227,334208 | 194 | 15604,131 | 7000 | 1 | 1 | 0 | 1 | 0 | 0 | 1 | 0 | 0 | 1 | 0 | 0 | 0 | 0 | 0 | 1 |
| 50.02414 | 19.81599 | MLP_U_4 | mlpu4_8 | Poland | U | 2 | 1 | 0 | 0 | 0 | 0 | 1 | 205,468 | 1227,048562 | 194 | 10935,37 | 16900 | 1 | 1 | 0 | 1 | 0 | 0 | 1 | 0 | 0 | 1 | 0 | 0 | 0 | 0 | 0 | 1 |
| 50.09411 | 19.9418 | MLP_U_5 | mlpu5_1 | Poland | U | 1 | 1 | 0 | 0 | 0 | 0 | 0 | 216,698 | 1227,365509 | 200 | 31785,383 | 7800 | 0 | 1 | 0 | 0 | 1 | 0 | 0 | 1 | 0 | 0 | 1 | 0 | 0 | 0 | 0 | 0 |
| 50.09256 | 19.94524 | MLP_U_5 | mlpu5_2 | Poland | U | 1 | 1 | 0 | 0 | 0 | 0 | 0 | 216 | 1227,077561 | 200 | 21286,224 | 19900 | 0 | 1 | 0 | 0 | 1 | 0 | 1 | 1 | 1 | 0 | 0 | 1 | 1 | 0 | 1 | 0 |
| 50.09169 | 19.94925 | MLP_U_5 | mlpu5_3 | Poland | U | 0 | 0 | 0 | 0 | 0 | 0 | 0 | 215 | 1226,825357 | 200 | 16015,944 | 21900 | 0 | 1 | 0 | 0 | 0 | 0 | 0 | 1 | 1 | 0 | 0 | 1 | 1 | 1 | 0 | 0 |
| 50.08933 | 19.94885 | MLP_U_5 | mlpu5_4 | Poland | U | 0 | 0 | 0 | 0 | 0 | 0 | 0 | 214,082 | 1226,635789 | 200 | 12017,249 | 38300 | 0 | 1 | 0 | 0 | 0 | 0 | 0 | 1 | 0 | 0 | 0 | 1 | 1 | 0 | 1 | 0 |
| 50.08757 | 19.9476 | MLP_U_5 | mlpu5_5 | Poland | U | 1 | 1 | 1 | 1 | 0 | 0 | 1 | 210,852 | 1226,538917 | 200 | 17655,229 | 26500 | 1 | 1 | 0 | 0 | 0 | 0 | 0 | 1 | 0 | 0 | 0 | 1 | 1 | 0 | 1 | 0 |
| 50.08916 | 19.94424 | MLP_U_5 | mlpu5_6 | Poland | U | 0 | 0 | 1 | 1 | 0 | 0 | 1 | 212,287 | 1226,82961 | 200 | 37876,823 | 12300 | 1 | 1 | 0 | 0 | 0 | 0 | 1 | 1 | 1 | 0 | 0 | 1 | 1 | 0 | 1 | 0 |
| 50.09082 | 19.94087 | MLP_U_5 | mlpu5_7 | Poland | U | 0 | 0 | 0 | 0 | 0 | 0 | 0 | 218,79 | 1227,125243 | 200 | 24819,231 | 13600 | 1 | 1 | 0 | 0 | 0 | 0 | 1 | 1 | 0 | 0 | 0 | 1 | 1 | 0 | 1 | 0 |
| 50.08958 | 19.93871 | MLP_U_5 | mlpu5_8 | Poland | U | 1 | 1 | 0 | 0 | 0 | 0 | 1 | 221,818 | 1227,040179 | 200 | 13336,398 | 27400 | 0 | 1 | 0 | 0 | 0 | 0 | 1 | 1 | 0 | 0 | 0 | 0 | 1 | 1 | 0 | 0 |
| 50.07617 | 19.95356 | MLP_U_6 | mlpu6_1 | Poland | U | 4 | 1 | 0 | 0 | 0 | 0 | 0 | 214,827 | 1225,286198 | 201 | 58688,573 | 0 | 0 | 1 | 0 | 0 | 0 | 1 | 0 | 0 | 0 | 0 | 0 | 0 | 0 | 0 | 0 | 1 |
| 50.07619 | 19.95687 | MLP_U_6 | mlpu6_2 | Poland | U | 2 | 1 | 0 | 0 | 0 | 0 | 1 | 212 | 1225,140408 | 201 | 42837,004 | 10800 | 1 | 1 | 0 | 0 | 0 | 1 | 0 | 1 | 0 | 0 | 0 | 0 | 0 | 1 | 0 | 0 |
| 50.07746 | 19.95937 | MLP_U_6 | mlpu6_3 | Poland | U | 0 | 0 | 0 | 0 | 0 | 0 | 0 | 209 | 1225,136019 | 201 | 23044,391 | 16300 | 1 | 1 | 0 | 0 | 0 | 1 | 0 | 1 | 1 | 1 | 0 | 1 | 0 | 1 | 0 | 0 |

| | | | | | | | | | | | | | | | | | | | | | | | | | | | | | | | | | |
|---|---|---|---|---|---|---|---|---|---|---|---|---|---|---|---|---|---|---|---|---|---|---|---|---|---|---|---|---|---|---|---|---|---|
| 50.0798 | 19.96099 | MLP_U_6 | mlpu6_4 | Poland | U | 1 | 1 | 0 | 0 | 0 | 0 | 0 | 208 | 1225,26658 | 201 | 17387,643 | 26500 | 0 | 1 | 1 | 0 | 0 | 0 | 1 | 1 | 1 | 0 | 0 | 1 | 1 | 1 | 1 | 0 |
| 50.07987 | 19.96414 | MLP_U_6 | mlpu6_5 | Poland | U | 0 | 0 | 0 | 0 | 0 | 0 | 0 | 207,577 | 1225,130905 | 201 | 22461,973 | 26200 | 1 | 1 | 0 | 0 | 0 | 0 | 0 | 1 | 1 | 0 | 0 | 1 | 1 | 0 | 0 | 0 |
| 50.0781 | 19.96464 | MLP_U_6 | mlpu6_6 | Poland | U | 1 | 1 | 0 | 0 | 0 | 0 | 0 | 207,84 | 1224,954823 | 201 | 22739,143 | 20500 | 1 | 1 | 0 | 0 | 0 | 0 | 1 | 1 | 1 | 0 | 1 | 1 | 1 | 0 | 1 | 0 |
| 50.07808 | 19.96814 | MLP_U_6 | mlpu6_7 | Poland | U | 0 | 0 | 0 | 0 | 0 | 0 | 0 | 206,912 | 1224,795722 | 201 | 20333,465 | 21900 | 1 | 1 | 0 | 0 | 0 | 0 | 1 | 1 | 1 | 0 | 1 | 0 | 1 | 0 | 1 | 0 |
| 50.07936 | 19.96924 | MLP_U_6 | mlpu6_8 | Poland | U | 0 | 0 | 0 | 0 | 0 | 0 | 0 | 207 | 1224,85794 | 201 | 17375,843 | 27400 | 0 | 1 | 0 | 0 | 0 | 0 | 1 | 1 | 1 | 0 | 0 | 1 | 1 | 1 | 0 | 0 |
| 50.07878 | 20.0551 | MLP_U_7 | mlpu7_1 | Poland | U | 0 | 0 | 0 | 0 | 0 | 0 | 0 | 206,17 | 1220,958437 | 207 | 29830,181 | 6100 | 1 | 1 | 0 | 0 | 1 | 0 | 0 | 1 | 0 | 1 | 0 | 1 | 0 | 0 | 0 | 1 |
| 50.08063 | 20.05194 | MLP_U_7 | mlpu7_2 | Poland | U | 0 | 0 | 0 | 0 | 0 | 0 | 0 | 206,268 | 1221,259139 | 207 | 21679,044 | 2800 | 1 | 1 | 0 | 0 | 1 | 0 | 0 | 0 | 0 | 1 | 1 | 0 | 0 | 0 | 0 | 0 |
| 50.0805 | 20.04892 | MLP_U_7 | mlpu7_3 | Poland | U | 0 | 0 | 0 | 0 | 0 | 0 | 0 | 207 | 1221,383278 | 207 | 22582,884 | 15100 | 1 | 1 | 0 | 0 | 1 | 0 | 1 | 1 | 0 | 1 | 1 | 0 | 1 | 0 | 0 | 0 |
| 50.08233 | 20.04951 | MLP_U_7 | mlpu7_4 | Poland | U | 0 | 0 | 0 | 0 | 0 | 0 | 0 | 208 | 1221,516522 | 207 | 29956,078 | 12300 | 1 | 1 | 0 | 1 | 1 | 0 | 0 | 1 | 0 | 0 | 1 | 0 | 1 | 0 | 0 | 0 |
| 50.08355 | 20.05295 | MLP_U_7 | mlpu7_5 | Poland | U | 1 | 1 | 0 | 0 | 0 | 0 | 1 | 205,44 | 1221,468505 | 207 | 20210,407 | 10200 | 1 | 1 | 0 | 1 | 0 | 0 | 0 | 1 | 1 | 1 | 0 | 1 | 0 | 0 | 0 | 0 |
| 50.08255 | 20.05592 | MLP_U_7 | mlpu7_6 | Poland | U | 0 | 0 | 0 | 0 | 0 | 0 | 0 | 208,056 | 1221,248455 | 207 | 17807,117 | 10800 | 0 | 1 | 0 | 0 | 0 | 0 | 0 | 1 | 1 | 0 | 1 | 1 | 0 | 0 | 0 | 0 |
| 50.08189 | 20.05894 | MLP_U_7 | mlpu7_7 | Poland | U | 0 | 0 | 0 | 0 | 0 | 0 | 0 | 210,988 | 1221,056693 | 207 | 14352,797 | 14200 | 0 | 1 | 0 | 0 | 0 | 0 | 0 | 0 | 1 | 0 | 0 | 1 | 0 | 0 | 0 | 0 |
| 50.07978 | 20.05941 | MLP_U_7 | mlpu7_8 | Poland | U | 1 | 1 | 0 | 0 | 0 | 0 | 1 | 209,876 | 1220,851986 | 207 | 23476,408 | 11000 | 1 | 1 | 1 | 1 | 0 | 0 | 0 | 0 | 1 | 0 | 0 | 1 | 0 | 0 | 0 | 0 |
| 50.05375 | 20.05612 | MLP_U_8 | mlpu8_1 | Poland | U | 2 | 1 | 0 | 0 | 0 | 0 | 0 | 197,984 | 1218,762867 | 209 | 50298,915 | 3000 | 1 | 1 | 0 | 1 | 0 | 0 | 0 | 0 | 1 | 0 | 0 | 1 | 0 | 0 | 0 | 1 |
| 50.05369 | 20.05977 | MLP_U_8 | mlpu8_2 | Poland | U | 2 | 1 | 0 | 0 | 0 | 0 | 0 | 198 | 1218,571609 | 209 | 52430,054 | 2700 | 1 | 1 | 0 | 1 | 0 | 0 | 0 | 0 | 1 | 0 | 0 | 1 | 0 | 0 | 0 | 0 |
| 50.05501 | 20.06131 | MLP_U_8 | mlpu8_3 | Poland | U | 2 | 1 | 0 | 0 | 0 | 0 | 0 | 196,69 | 1218,6568 | 209 | 30390,74 | 3000 | 1 | 1 | 0 | 1 | 0 | 0 | 0 | 0 | 1 | 1 | 1 | 1 | 0 | 0 | 0 | 0 |
| 50.05803 | 20.06032 | MLP_U_8 | mlpu8_4 | Poland | U | 0 | 0 | 0 | 0 | 0 | 0 | 0 | 196,77 | 1218,921506 | 209 | 34466,08 | 1800 | 1 | 1 | 0 | 1 | 0 | 0 | 1 | 0 | 0 | 1 | 1 | 1 | 0 | 0 | 0 | 1 |
| 50.0606 | 20.05952 | MLP_U_8 | mlpu8_5 | Poland | U | 0 | 0 | 0 | 0 | 0 | 0 | 0 | 197,772 | 1219,177563 | 209 | 9527,494 | 9900 | 1 | 1 | 0 | 1 | 0 | 0 | 1 | 1 | 1 | 0 | 1 | 1 | 0 | 0 | 0 | 0 |
| 50.0626 | 20.05739 | MLP_U_8 | mlpu8_6 | Poland | U | 0 | 0 | 0 | 0 | 0 | 0 | 1 | 199 | 1219,449841 | 209 | 10048,059 | 9500 | 0 | 1 | 1 | 0 | 0 | 0 | 1 | 1 | 1 | 0 | 0 | 1 | 0 | 0 | 0 | 0 |
| 50.06414 | 20.05394 | MLP_U_8 | mlpu8_7 | Poland | U | 0 | 0 | 0 | 0 | 0 | 0 | 1 | 202,904 | 1219,738324 | 209 | 16822,048 | 19900 | 1 | 1 | 1 | 1 | 0 | 0 | 1 | 1 | 1 | 0 | 0 | 1 | 0 | 1 | 0 | 0 |
| 50.06568 | 20.05196 | MLP_U_8 | mlpu8_8 | Poland | U | 1 | 1 | 0 | 0 | 0 | 0 | 0 | 205,577 | 1219,960354 | 209 | 18225,001 | 16300 | 1 | 1 | 1 | 0 | 1 | 0 | 1 | 1 | 0 | 0 | 0 | 1 | 1 | 1 | 0 | 0 |
| 50.01648 | 19.92274 | MLP_U_9 | mlpu9_1 | Poland | U | 2 | 1 | 0 | 0 | 0 | 0 | 0 | 222 | 1221,532571 | 202 | 41598,707 | 12900 | 1 | 1 | 1 | 0 | 1 | 0 | 0 | 1 | 1 | 0 | 0 | 1 | 1 | 0 | 0 | 0 |
| 50.01473 | 19.92247 | MLP_U_9 | mlpu9_2 | Poland | U | 1 | 1 | 0 | 0 | 0 | 0 | 0 | 224,972 | 1221,389487 | 202 | 50030,354 | 3400 | 1 | 1 | 0 | 1 | 1 | 0 | 0 | 1 | 0 | 0 | 0 | 0 | 0 | 0 | 0 | 0 |
| 50.01293 | 19.91948 | MLP_U_9 | mlpu9_3 | Poland | U | 2 | 1 | 0 | 0 | 0 | 0 | 0 | 229,968 | 1221,36973 | 202 | 44268,928 | 7700 | 1 | 1 | 1 | 1 | 0 | 0 | 0 | 1 | 0 | 0 | 0 | 0 | 1 | 0 | 0 | 0 |
| 50.01191 | 19.91577 | MLP_U_9 | mlpu9_4 | Poland | U | 2 | 1 | 0 | 0 | 0 | 0 | 0 | 235,076 | 1221,449785 | 202 | 58869,684 | 6000 | 1 | 1 | 1 | 1 | 0 | 1 | 0 | 1 | 0 | 0 | 0 | 0 | 1 | 0 | 0 | 0 |
| 50.01271 | 19.91228 | MLP_U_9 | mlpu9_5 | Poland | U | 2 | 1 | 0 | 0 | 0 | 0 | 0 | 244,599 | 1221,68503 | 202 | 31663,628 | 15000 | 1 | 1 | 1 | 1 | 0 | 0 | 0 | 1 | 0 | 0 | 0 | 0 | 1 | 0 | 0 | 0 |
| 50.01451 | 19.90912 | MLP_U_9 | mlpu9_6 | Poland | U | 0 | 0 | 0 | 0 | 0 | 0 | 0 | 234,764 | 1221,979284 | 202 | 6605,358 | 24000 | 1 | 1 | 0 | 0 | 0 | 0 | 1 | 1 | 1 | 0 | 0 | 1 | 1 | 0 | 0 | 0 |
| 50.01634 | 19.90593 | MLP_U_9 | mlpu9_7 | Poland | U | 0 | 0 | 0 | 0 | 0 | 0 | 1 | 225 | 1222,279423 | 202 | 17207,541 | 19400 | 0 | 1 | 1 | 0 | 0 | 0 | 1 | 1 | 1 | 0 | 0 | 1 | 1 | 0 | 0 | 0 |
| 50.0176 | 19.90211 | MLP_U_9 | mlpu9_8 | Poland | U | 1 | 1 | 0 | 0 | 0 | 0 | 1 | 229,145 | 1222,563522 | 202 | 38318,673 | 6700 | 1 | 1 | 0 | 1 | 1 | 0 | 1 | 0 | 0 | 1 | 0 | 0 | 0 | 0 | 1 | 1 |
| 43.68123 | 24.9061 | NIK_R_1 | nikr1_1 | Bulgaria | R | 1 | 1 | 1 | 1 | 1 | 1 | 0 | 174,331 | 449,5353568 | 472 | 26839,91 | 0 | 1 | 1 | 0 | 1 | 0 | 0 | 1 | 0 | 0 | 0 | 0 | 0 | 0 | 0 | 0 | 0 |
| 43.68348 | 24.90327 | NIK_R_1 | nikr1_2 | Bulgaria | R | 0 | 0 | 0 | 0 | 0 | 0 | 0 | 163,187 | 449,8719282 | 472 | 37174,351 | 0 | 1 | 1 | 0 | 1 | 0 | 0 | 1 | 0 | 0 | 0 | 0 | 0 | 0 | 0 | 0 | 0 |
| 43.68582 | 24.90001 | NIK_R_1 | nikr1_3 | Bulgaria | R | 0 | 0 | 0 | 0 | 0 | 0 | 0 | 148,986 | 450,2361546 | 472 | 36916,198 | 0 | 1 | 1 | 0 | 1 | 0 | 0 | 1 | 0 | 0 | 0 | 0 | 0 | 0 | 0 | 0 | 1 |
| 43.68825 | 24.89823 | NIK_R_1 | nikr1_4 | Bulgaria | R | 0 | 0 | 0 | 0 | 0 | 0 | 0 | 134,091 | 450,5164583 | 472 | 35383,665 | 0 | 1 | 1 | 0 | 1 | 0 | 0 | 1 | 0 | 0 | 0 | 0 | 0 | 0 | 0 | 0 | 0 |
| 43.69014 | 24.89507 | NIK_R_1 | nikr1_5 | Bulgaria | R | 1 | 1 | 2 | 1 | 0 | 0 | 0 | 117,127 | 450,846198 | 472 | 46507,791 | 0 | 1 | 1 | 0 | 1 | 0 | 0 | 1 | 0 | 0 | 0 | 0 | 0 | 0 | 0 | 0 | 0 |
| 43.70032 | 24.88796 | NIK_U_1 | niku1_1 | Bulgaria | U | 2 | 1 | 2 | 1 | 0 | 0 | 0 | 156,384 | 451,5930747 | 472 | 18369,988 | 10056 | 1 | 1 | 1 | 1 | 0 | 0 | 0 | 1 | 1 | 0 | 0 | 1 | 0 | 0 | 0 | 0 |
| 43.70078 | 24.88977 | NIK_U_1 | niku1_2 | Bulgaria | U | 0 | 0 | 0 | 0 | 0 | 0 | 0 | 136,199 | 451,7488291 | 472 | 15699,919 | 12466 | 1 | 1 | 1 | 1 | 0 | 0 | 0 | 0 | 0 | 0 | 1 | 1 | 0 | 0 | 0 | 0 |
| 43.70226 | 24.89076 | NIK_U_1 | niku1_3 | Bulgaria | U | 0 | 0 | 0 | 0 | 0 | 0 | 0 | 112,428 | 451,9561293 | 472 | 15628,661 | 8984 | 1 | 1 | 1 | 1 | 0 | 0 | 0 | 0 | 1 | 0 | 0 | 1 | 0 | 0 | 0 | 0 |
| 43.70153 | 24.89324 | NIK_U_1 | niku1_4 | Bulgaria | U | 0 | 0 | 0 | 0 | 0 | 0 | 0 | 79,275 | 451,9164826 | 472 | 4716,114 | 15031 | 0 | 1 | 0 | 0 | 0 | 0 | 0 | 0 | 1 | 0 | 0 | 1 | 0 | 0 | 0 | 0 |
| 43.69891 | 24.89285 | NIK_U_1 | niku1_5 | Bulgaria | U | 0 | 0 | 0 | 0 | 0 | 0 | 0 | 62,288 | 451,9996068 | 472 | 2161,925 | 9708 | 0 | 1 | 0 | 0 | 0 | 0 | 0 | 0 | 0 | 0 | 1 | 0 | 0 | 0 | 0 | 0 |
| 50.73186 | 17.94088 | OPO_R_2 | opor2_1 | Poland | R | 0 | 0 | 0 | 0 | 0 | 0 | 0 | 154 | 1374,299781 | 41 | 2292,372 | 0 | 0 | 1 | 0 | 0 | 0 | 0 | 1 | 0 | 0 | 0 | 0 | 0 | 0 | 0 | 0 | 0 |
| 50.72988 | 17.94055 | OPO_R_2 | opor2_2 | Poland | R | 0 | 0 | 0 | 0 | 0 | 0 | 0 | 153 | 1374,157726 | 41 | 388,464 | 0 | 1 | 1 | 0 | 1 | 0 | 0 | 0 | 0 | 0 | 0 | 0 | 0 | 0 | 0 | 0 | 0 |
| 50.72729 | 17.94077 | OPO_R_2 | opor2_3 | Poland | R | 2 | 1 | 0 | 0 | 0 | 0 | 1 | 153 | 1373,9352 | 41 | 26628,351 | 420 | 1 | 1 | 0 | 1 | 0 | 0 | 0 | 0 | 0 | 1 | 0 | 0 | 0 | 0 | 0 | 1 |
| 50.72801 | 17.93766 | OPO_R_2 | opor2_4 | Poland | R | 2 | 1 | 0 | 0 | 0 | 0 | 1 | 152,482 | 1374,147602 | 41 | 21275,712 | 0 | 1 | 1 | 0 | 1 | 0 | 0 | 0 | 0 | 0 | 1 | 0 | 0 | 0 | 0 | 0 | 0 |
| 50.72633 | 17.93504 | OPO_R_2 | opor2_5 | Poland | R | 0 | 0 | 0 | 0 | 0 | 0 | 0 | 152,788 | 1374,133223 | 41 | 28262,414 | 0 | 1 | 1 | 0 | 1 | 0 | 0 | 1 | 0 | 0 | 1 | 0 | 0 | 0 | 0 | 0 | 0 |
| 50.7157 | 17.91302 | OPO_R_4 | opor4_1 | Poland | R | 2 | 1 | 0 | 0 | 0 | 0 | 1 | 152,52 | 1374,335309 | 39 | 13296,544 | 2228 | 0 | 1 | 0 | 1 | 0 | 0 | 1 | 0 | 1 | 0 | 1 | 1 | 0 | 0 | 1 | 0 |
| 50.7177 | 17.91005 | OPO_R_4 | opor4_2 | Poland | R | 0 | 0 | 0 | 0 | 0 | 0 | 0 | 151,87 | 1374,645388 | 39 | 13683,153 | 3879 | 0 | 1 | 0 | 0 | 0 | 0 | 1 | 1 | 1 | 1 | 1 | 1 | 0 | 0 | 0 | 0 |
| 50.71972 | 17.90664 | OPO_R_4 | opor4_3 | Poland | R | 2 | 1 | 0 | 0 | 0 | 0 | 0 | 151,095 | 1374,971678 | 39 | 11711,888 | 4054 | 0 | 1 | 0 | 0 | 0 | 0 | 0 | 0 | 1 | 1 | 1 | 1 | 0 | 0 | 0 | 0 |
| 50.72028 | 17.90257 | OPO_R_4 | opor4_4 | Poland | R | 0 | 0 | 0 | 0 | 0 | 0 | 1 | 153 | 1375,214857 | 39 | 12350,572 | 10812 | 0 | 1 | 0 | 0 | 0 | 0 | 0 | 1 | 1 | 0 | 0 | 1 | 1 | 1 | 0 | 0 |
| 50.72258 | 17.90035 | OPO_R_4 | opor4_5 | Poland | R | 0 | 0 | 0 | 0 | 0 | 0 | 0 | 152,452 | 1375,507332 | 39 | 21903,902 | 3053 | 1 | 1 | 0 | 1 | 0 | 0 | 0 | 1 | 1 | 1 | 1 | 0 | 0 | 0 | 0 | 0 |
| 50.68035 | 17.94936 | OPO_U_2 | opou2_1 | Poland | U | 0 | 0 | 0 | 0 | 1 | 1 | 1 | 176 | 1366,419818 | 44 | 14172,943 | 20417 | 0 | 1 | 0 | 0 | 0 | 1 | 1 | 1 | 0 | 0 | 0 | 0 | 1 | 1 | 0 | 0 |
| 50.67767 | 17.95007 | OPO_U_2 | opou2_2 | Poland | U | 0 | 0 | 0 | 0 | 0 | 0 | 0 | 175 | 1369,46315 | 44 | 24440,611 | 687 | 0 | 1 | 0 | 0 | 1 | 0 | 0 | 0 | 0 | 0 | 1 | 0 | 1 | 0 | 0 | 0 |
| 50.67503 | 17.95091 | OPO_U_2 | opou2_3 | Poland | U | 0 | 0 | 0 | 0 | 0 | 0 | 1 | 174 | 1369,209419 | 44 | 2851,677 | 12280 | 0 | 1 | 0 | 0 | 0 | 0 | 0 | 1 | 0 | 0 | 0 | 0 | 1 | 1 | 0 | 0 |
| 50.67246 | 17.94994 | OPO_U_2 | opou2_4 | Poland | U | 0 | 0 | 0 | 0 | 0 | 0 | 0 | 175 | 1369,048418 | 44 | 7656,734 | 11120 | 0 | 1 | 0 | 0 | 0 | 0 | 0 | 1 | 1 | 0 | 0 | 1 | 1 | 1 | 1 | 0 |
| 50.67097 | 17.94782 | OPO_U_2 | opou2_5 | Poland | U | 0 | 0 | 0 | 0 | 0 | 0 | 0 | 174 | 1369,030359 | 44 | 19134,077 | 1500 | 0 | 1 | 0 | 0 | 0 | 0 | 1 | 0 | 1 | 0 | 0 | 1 | 0 | 0 | 1 | 0 |
| 50.67154 | 17.9438 | OPO_U_2 | opou2_6 | Poland | U | 0 | 0 | 0 | 0 | 0 | 0 | 0 | 172,602 | 1369,27291 | 44 | 11253,061 | 6002 | 0 | 1 | 0 | 0 | 0 | 0 | 1 | 0 | 1 | 0 | 0 | 1 | 0 | 0 | 1 | 0 |
| 50.67288 | 17.94003 | OPO_U_2 | opou2_7 | Poland | U | 0 | 0 | 0 | 0 | 0 | 0 | 1 | 170,632 | 1369,562884 | 44 | 20495,029 | 3800 | 0 | 1 | 0 | 0 | 0 | 0 | 1 | 0 | 1 | 0 | 0 | 1 | 0 | 0 | 1 | 0 |
| 50.67421 | 17.9363 | OPO_U_2 | opou2_8 | Poland | U | 0 | 0 | 0 | 0 | 0 | 0 | 0 | 168,114 | 1369,851593 | 44 | 11908,39 | 9539 | 0 | 1 | 0 | 0 | 0 | 0 | 1 | 1 | 1 | 0 | 0 | 1 | 1 | 0 | 1 | 0 |
| 50.65042 | 17.9201 | OPO_U_3 | opou3_1 | Poland | U | 0 | 0 | 0 | 0 | 0 | 0 | 1 | 153 | 1368,714839 | 44 | 55003,653 | 0 | 1 | 0 | 0 | 0 | 1 | 0 | 0 | 0 | 0 | 0 | 0 | 0 | 0 | 0 | 0 | 0 |
| 50.65279 | 17.91983 | OPO_U_3 | opou3_2 | Poland | U | 2 | 1 | 0 | 0 | 0 | 0 | 1 | 152,602 | 1368,920371 | 44 | 54112,033 | 0 | 1 | 0 | 0 | 0 | 1 | 0 | 0 | 0 | 0 | 0 | 0 | 0 | 0 | 0 | 0 | 0 |
| 50.65528 | 17.92088 | OPO_U_3 | opou3_3 | Poland | U | 1 | 1 | 0 | 0 | 0 | 0 | 0 | 151,999 | 1369,069998 | 44 | 51671,64 | 0 | 1 | 1 | 0 | 0 | 1 | 0 | 0 | 0 | 0 | 0 | 0 | 0 | 0 | 0 | 0 | 0 |
| 50.65803 | 17.92171 | OPO_U_3 | opou3_4 | Poland | U | 0 | 0 | 0 | 0 | 0 | 0 | 0 | 149,439 | 1369,252128 | 44 | 17549,452 | 2000 | 1 | 1 | 0 | 0 | 1 | 0 | 1 | 0 | 0 | 0 | 1 | 0 | 0 | 0 | 0 | 0 |
| 50.65973 | 17.91942 | OPO_U_3 | opou3_5 | Poland | U | 2 | 1 | 0 | 0 | 0 | 0 | 0 | 152 | 1369,500617 | 44 | 25836,161 | 4257 | 1 | 1 | 0 | 0 | 1 | 0 | 0 | 1 | 0 | 0 | 0 | 1 | 1 | 0 | 0 | 0 |
| 50.66359 | 17.99073 | OPO_U_4 | opou4_1 | Poland | U | 0 | 0 | 0 | 0 | 0 | 0 | 0 | 160,952 | 1366,353882 | 44 | 3896,016 | 11295 | 0 | 1 | 0 | 0 | 0 | 0 | 0 | 1 | 1 | 0 | 0 | 1 | 0 | 0 | 0 | 0 |
| 50.66541 | 17.98931 | OPO_U_4 | opou4_2 | Poland | U | 0 | 0 | 0 | 0 | 0 | 0 | 1 | 160 | 1366,569187 | 44 | 6871,253 | 11470 | 0 | 1 | 0 | 0 | 0 | 0 | 1 | 1 | 1 | 0 | 0 | 1 | 0 | 0 | 0 | 0 |
| 50.66609 | 17.98536 | OPO_U_4 | opou4_3 | Poland | U | 0 | 0 | 0 | 0 | 1 | 1 | 0 | 163,704 | 1366,81637 | 44 | 10871,554 | 10477 | 0 | 1 | 0 | 0 | 0 | 1 | 0 | 1 | 1 | 0 | 0 | 1 | 1 | 0 | 0 | 0 |
| 50.66545 | 17.98124 | OPO_U_4 | opou4_4 | Poland | U | 0 | 0 | 0 | 0 | 0 | 0 | 1 | 164,62 | 1366,963962 | 44 | 10970,327 | 4007 | 0 | 1 | 0 | 0 | 0 | 1 | 0 | 1 | 1 | 0 | 0 | 1 | 0 | 0 | 1 | 0 |
| 50.66748 | 17.97897 | OPO_U_4 | opou4_5 | Poland | U | 1 | 1 | 1 | 1 | 0 | 0 | 1 | 170 | 1367,238257 | 44 | 2687,052 | 11135 | 0 | 1 | 0 | 0 | 0 | 0 | 0 | 1 | 1 | 0 | 0 | 1 | 1 | 0 | 0 | 0 |
| 49.46319 | 19.81097 | ORA_R_1 | orar1_1 | Poland | R | 1 | 1 | 0 | 0 | 0 | 0 | 0 | 655 | 1179,885092 | 229 | 32604,394 | 0 | 1 | 1 | 1 | 1 | 0 | 0 | 1 | 0 | 0 | 0 | 0 | 0 | 0 | 0 | 0 | 0 |
| 49.461 | 19.80947 | ORA_R_1 | orar1_2 | Poland | R | 2 | 1 | 0 | 0 | 0 | 0 | 0 | 656 | 1179,774955 | 229 | 12104,502 | 0 | 0 | 1 | 1 | 0 | 0 | 0 | 1 | 0 | 0 | 0 | 0 | 0 | 0 | 0 | 0 | 0 |
| 49.45979 | 19.80698 | ORA_R_1 | orar1_3 | Poland | R | 0 | 0 | 0 | 0 | 0 | 0 | 0 | 656,756 | 1179,794912 | 229 | 53999,756 | 0 | 1 | 1 | 1 | 1 | 0 | 0 | 0 | 0 | 0 | 0 | 0 | 0 | 0 | 0 | 0 | 0 |
| 49.46149 | 19.80299 | ORA_R_1 | orar1_4 | Poland | R | 0 | 0 | 0 | 0 | 0 | 0 | 0 | 657 | 1180,1292 | 229 | 43140,021 | 2213 | 1 | 0 | 1 | 1 | 0 | 0 | 0 | 0 | 0 | 0 | 0 | 0 | 0 | 0 | 1 | 0 |
| 49.46311 | 19.7991 | ORA_R_1 | orar1_5 | Poland | R | 1 | 1 | 0 | 0 | 0 | 0 | 0 | 652,611 | 1180,448443 | 229 | 45501,87 | 689 | 1 | 1 | 1 | 1 | 0 | 0 | 0 | 0 | 0 | 0 | 1 | 0 | 0 | 0 | 0 | 0 |
| 49.47001 | 19.63731 | ORA_R_2 | orar2_1 | Poland | R | 0 | 0 | 0 | 0 | 0 | 0 | 1 | 614,695 | 1188,842771 | 220 | 25824,907 | 5047 | 1 | 1 | 1 | 1 | 0 | 0 | 0 | 1 | 1 | 0 | 1 | 0 | 0 | 0 | 0 | 0 |
| 49.46732 | 19.63939 | ORA_R_2 | orar2_2 | Poland | R | 0 | 0 | 0 | 0 | 0 | 0 | 1 | 609,069 | 1188,522079 | 220 | 13789,535 | 3160 | 1 | 1 | 0 | 1 | 0 | 0 | 0 | 0 | 1 | 1 | 1 | 0 | 0 | 0 | 0 | 1 |
| 49.46441 | 19.63899 | ORA_R_2 | orar2_3 | Poland | R | 0 | 0 | 0 | 0 | 0 | 0 | 1 | 607,362 | 1188,298643 | 220 | 18135,781 | 4168 | 1 | 1 | 0 | 1 | 0 | 0 | 0 | 0 | 1 | 1 | 1 | 0 | 0 | 0 | 0 | 0 |
| 49.46142 | 19.64138 | ORA_R_2 | orar2_4 | Poland | R | 0 | 0 | 0 | 0 | 0 | 0 | 0 | 604,892 | 1187,936225 | 220 | 26841,731 | 2321 | 1 | 1 | 0 | 1 | 0 | 0 | 0 | 0 | 0 | 1 | 1 | 0 | 0 | 0 | 0 | 0 |
| 49.4585 | 19.64105 | ORA_R_2 | orar2_5 | Poland | R | 0 | 0 | 0 | 0 | 0 | 0 | 0 | 602,78 | 1187,711687 | 220 | 23764,624 | 0 | 1 | 1 | 0 | 1 | 0 | 0 | 0 | 0 | 0 | 1 | 0 | 0 | 0 | 0 | 0 | 0 |
| 49.4853 | 19.70148 | ORA_U_1 | orau1_1 | Poland | U | 0 | 0 | 0 | 0 | 0 | 0 | 1 | 662,496 | 1187,000764 | 222 | 6652,963 | 3198 | 0 | 1 | 1 | 0 | 0 | 1 | 0 | 1 | 0 | 1 | 1 | 1 | 0 | 0 | 0 | 0 |
| 49.48362 | 19.69963 | ORA_U_1 | orau1_2 | Poland | U | 0 | 0 | 0 | 0 | 0 | 0 | 0 | 642,357 | 1186,950066 | 222 | 4798,05 | 6937 | 0 | 1 | 1 | 0 | 0 | 0 | 0 | 1 | 1 | 1 | 0 | 1 | 0 | 0 | 0 | 0 |
| 49.48136 | 19.69771 | ORA_U_1 | orau1_3 | Poland | U | 0 | 0 | 0 | 0 | 0 | 0 | 1 | 620,756 | 1186,854042 | 222 | 5276,207 | 8812 | 0 | 1 | 1 | 0 | 0 | 0 | 0 | 0 | 1 | 1 | 0 | 1 | 0 | 0 | 0 | 0 |
| 49.48015 | 19.69329 | ORA_U_1 | orau1_4 | Poland | U | 0 | 0 | 0 | 0 | 0 | 0 | 0 | 617,844 | 1186,967815 | 222 | 7477,135 | 8641 | 0 | 1 | 1 | 0 | 0 | 0 | 0 | 1 | 0 | 1 | 0 | 1 | 0 | 1 | 0 | 0 |
| 49.4792 | 19.68927 | ORA_U_1 | orau1_5 | Poland | U | 1 | 1 | 0 | 0 | 0 | 0 | 0 | 618,231 | 1187,083356 | 222 | 16505,011 | 4046 | 0 | 1 | 1 | 0 | 0 | 0 | 0 | 0 | 0 | 1 | 0 | 1 | 0 | 0 | 0 | 0 |
| 49.43929 | 19.85513 | ORA_U_2 | orau2_1 | Poland | U | 0 | 0 | 0 | 0 | 0 | 0 | 1 | 671,296 | 1175,77141 | 234 | 13154,95 | 13446 | 0 | 1 | 0 | 0 | 0 | 0 | 0 | 1 | 0 | 0 | 0 | 1 | 0 | 1 | 0 | 0 |
| 49.43698 | 19.85313 | ORA_U_2 | orau2_2 | Poland | U | 0 | 0 | 0 | 0 | 0 | 0 | 1 | 674,234 | 1175,679848 | 234 | 4589,737 | 17408 | 0 | 1 | 0 | 0 | 0 | 0 | 0 | 1 | 1 | 0 | 0 | 1 | 0 | 1 | 0 | 1 |
| 49.43444 | 19.85429 | ORA_U_2 | orau2_3 | Poland | U | 0 | 0 | 0 | 0 | 0 | 0 | 1 | 676,993 | 1175,408026 | 234 | 20216,777 | 6965 | 1 | 1 | 0 | 1 | 0 | 0 | 0 | 1 | 1 | 1 | 0 | 1 | 1 | 0 | 0 | 0 |
| 49.4319 | 19.85178 | ORA_U_2 | orau2_4 | Poland | U | 0 | 0 | 0 | 0 | 0 | 0 | 1 | 679,527 | 1175,317844 | 234 | 28557,068 | 4420 | 1 | 1 | 0 | 1 | 0 | 0 | 0 | 0 | 1 | 1 | 0 | 1 | 0 | 1 | 0 | 0 |
| 49.42971 | 19.84899 | ORA_U_2 | orau2_5 | Poland | U | 1 | 1 | 0 | 0 | 0 | 0 | 1 | 683,564 | 1175,269746 | 234 | 32520,84 | 368 | 1 | 1 | 0 | 1 | 0 | 0 | 1 | 0 | 0 | 1 | 0 | 0 | 0 | 0 | 1 | 0 |
| 43.73377 | 23.97318 | ORI_R_1 | orir1_1 | Bulgaria | R | 0 | 0 | 0 | 0 | 0 | 0 | 0 | 104,329 | 514,1478107 | 402 | 49211,151 | 851 | 1 | 1 | 0 | 1 | 0 | 0 | 0 | 0 | 0 | 0 | 1 | 0 | 0 | 0 | 0 | 0 |
| 43.73203 | 23.97622 | ORI_R_1 | orir1_2 | Bulgaria | R | 0 | 0 | 1 | 1 | 0 | 0 | 0 | 112,6 | 513,8428706 | 402 | 58899,665 | 2106 | 1 | 0 | 1 | 1 | 0 | 0 | 0 | 0 | 0 | 0 | 0 | 0 | 1 | 0 | 0 | 0 |
| 43.73164 | 23.98058 | ORI_R_1 | orir1_3 | Bulgaria | R | 0 | 0 | 0 | 0 | 0 | 0 | 0 | 102,104 | 513,5227958 | 402 | 45442,835 | 0 | 1 | 1 | 1 | 1 | 0 | 0 | 1 | 0 | 0 | 0 | 0 | 0 | 0 | 0 | 0 | 0 |
| 43.73116 | 23.98435 | ORI_R_1 | orir1_4 | Bulgaria | R | 0 | 0 | 0 | 0 | 0 | 0 | 0 | 112,308 | 513,2364659 | 402 | 11853,52 | 7946 | 1 | 0 | 0 | 1 | 0 | 0 | 0 | 0 | 0 | 0 | 0 | 0 | 0 | 0 | 1 | 0 |
| 43.73032 | 23.9883 | ORI_R_1 | orir1_5 | Bulgaria | R | 0 | 0 | 1 | 1 | 0 | 0 | 0 | 117,614 | 512,9180096 | 402 | 34318,55 | 0 | 1 | 1 | 0 | 1 | 0 | 0 | 1 | 0 | 1 | 0 | 1 | 0 | 0 | 0 | 0 | 0 |
| 43.83452 | 23.84544 | ORI_R_2 | orir2_1 | Romania | R | 2 | 1 | 0 | 0 | 0 | 0 | 0 | 32,697 | 528,8790175 | 399 | 5293,537 | 0 | 0 | 1 | 0 | 0 | 0 | 0 | 1 | 0 | 0 | 0 | 0 | 0 | 0 | 0 | 0 | 0 |
| 43.83706 | 23.84682 | ORI_R_2 | orir2_2 | Romania | R | 2 | 1 | 0 | 0 | 0 | 0 | 0 | 29,398 | 528,9408885 | 399 | 5938,251 | 0 | 0 | 1 | 0 | 0 | 0 | 0 | 1 | 0 | 1 | 1 | 1 | 0 | 0 | 0 | 0 | 0 |
| 43.83981 | 23.84856 | ORI_R_2 | orir2_3 | Romania | R | 1 | 1 | 0 | 0 | 0 | 0 | 0 | 41,316 | 528,9922688 | 399 | 49464,28 | 0 | 1 | 0 | 0 | 1 | 0 | 0 | 0 | 0 | 0 | 0 | 1 | 0 | 0 | 0 | 0 | 0 |
| 43.84252 | 23.85046 | ORI_R_2 | orir2_4 | Romania | R | 0 | 0 | 1 | 1 | 0 | 0 | 0 | 38,982 | 529,029565 | 399 | 64169,945 | 0 | 1 | 0 | 0 | 1 | 0 | 0 | 0 | 0 | 0 | 0 | 0 | 0 | 0 | 0 | 0 | 0 |
| 43.84557 | 23.85239 | ORI_R_2 | orir2_5 | Romania | R | 0 | 0 | 1 | 1 | 0 | 0 | 0 | 33,021 | 529,0866009 | 399 | 30372,831 | 0 | 1 | 1 | 0 | 1 | 0 | 0 | 1 | 0 | 0 | 0 | 0 | 0 | 0 | 0 | 0 | 1 |
| 43.74016 | 23.96221 | ORI_U_1 | oriu1_1 | Bulgaria | U | 0 | 0 | 0 | 0 | 0 | 0 | 0 | 40,644 | 515,281201 | 402 | 30747,654 | 6948 | 1 | 0 | 0 | 1 | 0 | 0 | 0 | 0 | 0 | 0 | 1 | 0 | 0 | 0 | 0 | 0 |
| 43.73992 | 23.95963 | ORI_U_1 | oriu1_2 | Bulgaria | U | 0 | 0 | 0 | 0 | 0 | 0 | 0 | 67,392 | 515,4391656 | 402 | 47566,505 | 4652 | 1 | 1 | 0 | 1 | 0 | 0 | 0 | 0 | 1 | 0 | 0 | 1 | 0 | 0 | 0 | 0 |
| 43.73995 | 23.955 | ORI_U_1 | oriu1_3 | Bulgaria | U | 0 | 0 | 0 | 0 | 0 | 0 | 1 | 90,9 | 515,7559979 | 402 | 34592,76 | 6174 | 1 | 1 | 0 | 1 | 0 | 0 | 1 | 1 | 1 | 0 | 0 | 1 | 0 | 0 | 0 | 0 |
| 43.73791 | 23.95899 | ORI_U_1 | oriu1_4 | Bulgaria | U | 0 | 0 | 0 | 0 | 0 | 0 | 0 | 100,289 | 515,3625897 | 402 | 23660,637 | 10194 | 0 | 1 | 0 | 1 | 1 | 0 | 0 | 1 | 1 | 0 | 0 | 1 | 0 | 1 | 0 | 0 |
| 43.73696 | 23.96137 | ORI_U_1 | oriu1_5 | Bulgaria | U | 0 | 0 | 0 | 0 | 0 | 0 | 1 | 94,04 | 515,1459936 | 402 | 33983,299 | 9934 | 1 | 1 | 0 | 0 | 1 | 0 | 0 | 0 | 1 | 0 | 0 | 1 | 0 | 1 | 0 | 0 |
| 43.77953 | 23.9604 | ORI_U_2 | oriu2_1 | Romania | U | 0 | 0 | 5 | 1 | 0 | 0 | 1 | 35,92 | 517,7941014 | 400 | 14044,201 | 2203 | 1 | 1 | 0 | 1 | 0 | 0 | 0 | 1 | 1 | 1 | 0 | 1 | 0 | 0 | 0 | 0 |

| | | | | | | | | | | | | | | | | | | | | | | | | | | | | | | | | | |
|---|---|---|---|---|---|---|---|---|---|---|---|---|---|---|---|---|---|---|---|---|---|---|---|---|---|---|---|---|---|---|---|---|---|
| 43.78003 | 23.95621 | ORI_U_2 | oriu2_2 | Romania | U | 0 | 0 | 1 | 1 | 0 | 0 | 1 | 34,216 | 518,1030067 | 400 | 21994,709 | 3204 | 0 | 1 | 0 | 1 | 0 | 0 | 0 | 0 | 1 | 1 | 0 | 1 | 0 | 0 | 0 | 0 |
| 43.78124 | 23.95246 | ORI_U_2 | oriu2_3 | Romania | U | 0 | 0 | 2 | 1 | 0 | 0 | 1 | 35 | 518,42933 | 400 | 15752,829 | 3791 | 0 | 1 | 0 | 0 | 0 | 0 | 0 | 0 | 1 | 1 | 1 | 1 | 0 | 0 | 0 | 0 |
| 43.78387 | 23.95323 | ORI_U_2 | oriu2_4 | Romania | U | 0 | 0 | 0 | 0 | 0 | 0 | 1 | 38,068 | 518,5406945 | 400 | 24614,992 | 8171 | 0 | 1 | 0 | 0 | 0 | 0 | 0 | 0 | 1 | 0 | 0 | 1 | 0 | 0 | 0 | 0 |
| 43.78595 | 23.95465 | ORI_U_2 | oriu2_5 | Romania | U | 0 | 0 | 3 | 1 | 0 | 0 | 1 | 40,009 | 518,5705688 | 400 | 18807,77 | 7585 | 0 | 1 | 0 | 0 | 1 | 0 | 0 | 0 | 1 | 0 | 0 | 1 | 0 | 0 | 0 | 0 |
| 50.05492 | 15.81161 | PAR_R_1 | parr1_1 | Czech | R | 2 | 1 | 0 | 0 | 0 | 0 | 1 | 223,855 | 1434,162799 | 1 | 24002,233 | 769 | 1 | 1 | 0 | 1 | 0 | 0 | 0 | 0 | 1 | 0 | 1 | 0 | 0 | 0 | 0 | 1 |
| 50.05472 | 15.80755 | PAR_R_1 | parr1_2 | Czech | R | 1 | 1 | 0 | 0 | 0 | 0 | 1 | 220,001 | 1434,374673 | 1 | 27837,719 | 533 | 1 | 1 | 0 | 1 | 0 | 0 | 0 | 0 | 1 | 0 | 1 | 0 | 0 | 0 | 0 | 0 |
| 50.0539 | 15.80363 | PAR_R_1 | parr1_3 | Czech | R | 0 | 0 | 0 | 0 | 0 | 0 | 1 | 219 | 1434,536825 | 1 | 11393,558 | 95 | 1 | 1 | 0 | 1 | 0 | 0 | 0 | 0 | 1 | 0 | 1 | 0 | 0 | 0 | 0 | 0 |
| 50.05232 | 15.80035 | PAR_R_1 | parr1_4 | Czech | R | 3 | 1 | 0 | 0 | 0 | 0 | 0 | 219 | 1434,610214 | 1 | 17655,142 | 100 | 1 | 1 | 0 | 1 | 0 | 0 | 0 | 0 | 1 | 1 | 1 | 0 | 0 | 0 | 0 | 0 |
| 50.05092 | 15.79823 | PAR_R_1 | parr1_5 | Czech | R | 0 | 0 | 0 | 0 | 0 | 0 | 0 | 219 | 1434,631393 | 1 | 21156,998 | 615 | 1 | 1 | 0 | 1 | 0 | 0 | 0 | 0 | 1 | 1 | 1 | 0 | 0 | 0 | 0 | 0 |
| 50.03945 | 15.77629 | PAR_U_1 | paru1_1 | Czech | U | 0 | 0 | 0 | 0 | 0 | 0 | 0 | 216,98 | 1435,060328 | 1 | 18685,863 | 10621 | 0 | 1 | 0 | 0 | 1 | 0 | 0 | 0 | 0 | 0 | 0 | 0 | 0 | 1 | 1 | 0 |
| 50.04109 | 15.77395 | PAR_U_1 | paru1_2 | Czech | U | 1 | 1 | 0 | 0 | 0 | 0 | 1 | 215 | 1435,301298 | 1 | 30451,991 | 3528 | 0 | 1 | 0 | 0 | 1 | 0 | 0 | 0 | 0 | 0 | 0 | 0 | 1 | 0 | 0 | 0 |
| 50.04361 | 15.77546 | PAR_U_1 | paru1_3 | Czech | U | 2 | 1 | 0 | 0 | 0 | 0 | 1 | 219,335 | 1435,392478 | 1 | 18885,394 | 2205 | 0 | 1 | 0 | 0 | 1 | 0 | 0 | 0 | 1 | 0 | 0 | 1 | 0 | 0 | 0 | 1 |
| 50.04447 | 15.77896 | PAR_U_1 | paru1_4 | Czech | U | 2 | 1 | 0 | 0 | 0 | 0 | 1 | 218,068 | 1435,256875 | 1 | 23067,731 | 1073 | 1 | 1 | 0 | 0 | 1 | 0 | 0 | 0 | 1 | 1 | 1 | 0 | 1 | 0 | 0 | 1 |
| 50.0447 | 15.7831 | PAR_U_1 | paru1_5 | Czech | U | 4 | 1 | 0 | 0 | 0 | 0 | 1 | 216 | 1435,041646 | 1 | 38893,04 | 6045 | 1 | 1 | 0 | 0 | 1 | 0 | 0 | 0 | 0 | 1 | 0 | 1 | 1 | 0 | 0 | 0 |
| 50.07525 | 14.25502 | PRA_R_1 | prar1_1 | Czech | R | 1 | 1 | 0 | 0 | 0 | 0 | 1 | 355,935 | 1524,523549 | -96 | 16762,316 | 10218 | 1 | 1 | 0 | 1 | 0 | 0 | 0 | 1 | 1 | 0 | 0 | 1 | 1 | 0 | 0 | 1 |
| 50.07321 | 14.25165 | PRA_R_1 | prar1_2 | Czech | R | 5 | 1 | 0 | 0 | 0 | 0 | 1 | 347,967 | 1524,586826 | -96 | 59662,877 | 187 | 1 | 0 | 0 | 1 | 0 | 0 | 0 | 0 | 0 | 1 | 1 | 0 | 0 | 0 | 0 | 1 |
| 50.07136 | 14.24851 | PRA_R_1 | prar1_3 | Czech | R | 4 | 1 | 0 | 0 | 0 | 0 | 0 | 351,638 | 1524,653879 | -96 | 66585,443 | 369 | 1 | 0 | 0 | 1 | 0 | 0 | 0 | 0 | 0 | 1 | 1 | 0 | 0 | 0 | 0 | 1 |
| 50.06927 | 14.24705 | PRA_R_1 | prar1_4 | Czech | R | 2 | 1 | 0 | 0 | 0 | 0 | 0 | 352,019 | 1524,591558 | -96 | 21737,853 | 6742 | 1 | 0 | 0 | 1 | 0 | 0 | 0 | 0 | 0 | 1 | 0 | 1 | 1 | 0 | 0 | 1 |
| 50.06612 | 14.24727 | PRA_R_1 | prar1_5 | Czech | R | 5 | 1 | 0 | 0 | 0 | 0 | 1 | 353,802 | 1524,387512 | -96 | 53759,632 | 713 | 1 | 0 | 0 | 1 | 0 | 0 | 0 | 0 | 0 | 1 | 1 | 0 | 0 | 0 | 0 | 1 |
| 50.06686 | 14.35437 | PRA_R_2 | prar2_1 | Czech | R | 2 | 1 | 0 | 0 | 0 | 0 | 1 | 271,764 | 1518,176286 | -89 | 50870,478 | 820 | 1 | 1 | 0 | 0 | 1 | 0 | 0 | 0 | 1 | 1 | 1 | 0 | 0 | 0 | 0 | 1 |
| 50.06733 | 14.35883 | PRA_R_2 | prar2_2 | Czech | R | 3 | 1 | 0 | 0 | 0 | 0 | 1 | 292,553 | 1517,957725 | -89 | 42881,118 | 6512 | 1 | 1 | 0 | 0 | 1 | 0 | 0 | 1 | 1 | 0 | 1 | 0 | 1 | 0 | 0 | 1 |
| 50.06549 | 14.36097 | PRA_R_2 | prar2_3 | Czech | R | 0 | 0 | 0 | 0 | 0 | 0 | 1 | 299,588 | 1517,693707 | -89 | 29017,271 | 9141 | 1 | 1 | 0 | 0 | 0 | 0 | 1 | 1 | 1 | 0 | 1 | 0 | 0 | 0 | 1 | 1 |
| 50.06478 | 14.36542 | PRA_R_2 | prar2_4 | Czech | R | 1 | 1 | 0 | 0 | 0 | 0 | 0 | 298,616 | 1517,409998 | -89 | 27974,916 | 7097 | 1 | 1 | 0 | 0 | 0 | 0 | 1 | 1 | 1 | 0 | 1 | 1 | 0 | 0 | 1 | 1 |
| 50.06474 | 14.36927 | PRA_R_2 | prar2_5 | Czech | R | 0 | 0 | 0 | 0 | 0 | 0 | 0 | 305,308 | 1517,156629 | -89 | 33803,485 | 5336 | 1 | 1 | 1 | 1 | 0 | 0 | 1 | 1 | 1 | 0 | 0 | 0 | 1 | 0 | 1 | 0 |
| 50.08214 | 14.40397 | PRA_U_1 | prau1_1 | Czech | U | 1 | 1 | 0 | 0 | 0 | 0 | 0 | 199,42 | 1516,270877 | -87 | 27790,253 | 15231 | 1 | 1 | 0 | 0 | 1 | 0 | 0 | 1 | 1 | 0 | 0 | 0 | 0 | 1 | 0 | 1 |
| 50.0833 | 14.40086 | PRA_U_1 | prau1_2 | Czech | U | 2 | 1 | 0 | 0 | 0 | 0 | 1 | 232,784 | 1516,527604 | -87 | 43757,792 | 0 | 1 | 1 | 0 | 0 | 1 | 0 | 0 | 1 | 1 | 0 | 0 | 0 | 0 | 0 | 0 | 1 |
| 50.08491 | 14.39757 | PRA_U_1 | prau1_3 | Czech | U | 1 | 1 | 1 | 1 | 0 | 0 | 1 | 271,447 | 1516,824139 | -87 | 56757,109 | 455 | 1 | 1 | 0 | 0 | 1 | 0 | 0 | 0 | 1 | 0 | 0 | 1 | 0 | 0 | 0 | 1 |
| 50.08494 | 14.39351 | PRA_U_1 | prau1_4 | Czech | U | 2 | 1 | 0 | 0 | 0 | 0 | 1 | 272,443 | 1517,06286 | -87 | 58536,957 | 996 | 1 | 1 | 0 | 0 | 1 | 0 | 0 | 0 | 1 | 0 | 1 | 0 | 0 | 0 | 0 | 0 |
| 50.08715 | 14.39285 | PRA_U_1 | prau1_5 | Czech | U | 3 | 1 | 0 | 0 | 0 | 0 | 1 | 241,288 | 1517,243352 | -87 | 27392,123 | 5930 | 0 | 1 | 0 | 0 | 0 | 0 | 1 | 1 | 1 | 0 | 1 | 0 | 0 | 1 | 0 | 1 |
| 50.07689 | 14.44608 | PRA_U_2 | prau2_1 | Czech | U | 2 | 1 | 0 | 0 | 0 | 0 | 0 | 255,566 | 1513,484711 | -84 | 21561,76 | 22582 | 0 | 1 | 0 | 0 | 1 | 0 | 0 | 1 | 1 | 0 | 0 | 0 | 0 | 1 | 0 | 0 |
| 50.07449 | 14.44774 | PRA_U_2 | prau2_2 | Czech | U | 0 | 0 | 0 | 0 | 0 | 0 | 1 | 269 | 1513,229113 | -84 | 20687,341 | 20872 | 0 | 1 | 0 | 0 | 1 | 0 | 0 | 1 | 1 | 0 | 0 | 0 | 1 | 1 | 0 | 1 |
| 50.07342 | 14.45268 | PRA_U_2 | prau2_3 | Czech | U | 0 | 0 | 0 | 0 | 0 | 0 | 1 | 248,848 | 1512,882804 | -84 | 26815,27 | 12806 | 0 | 1 | 1 | 0 | 0 | 0 | 0 | 1 | 1 | 0 | 1 | 0 | 1 | 0 | 0 | 0 |
| 50.0738 | 14.45719 | PRA_U_2 | prau2_4 | Czech | U | 0 | 0 | 0 | 0 | 0 | 0 | 1 | 264,165 | 1512,646999 | -84 | 20923,839 | 17332 | 0 | 1 | 1 | 0 | 0 | 0 | 0 | 1 | 1 | 0 | 0 | 0 | 1 | 1 | 0 | 0 |
| 50.07408 | 14.46091 | PRA_U_2 | prau2_5 | Czech | U | 0 | 0 | 0 | 0 | 0 | 0 | 0 | 267,688 | 1512,436145 | -84 | 17988,484 | 14670 | 0 | 1 | 1 | 0 | 0 | 0 | 0 | 1 | 1 | 0 | 1 | 0 | 1 | 1 | 0 | 0 |
| 51.34456 | 21.05643 | RAD_R_1 | radr1_1 | Poland | R | 2 | 1 | 0 | 0 | 0 | 0 | 1 | 189,584 | 1295,542306 | 133 | 47185,934 | 768 | 1 | 1 | 1 | 1 | 0 | 0 | 0 | 0 | 1 | 0 | 1 | 0 | 0 | 0 | 0 | 0 |
| 51.34172 | 21.05668 | RAD_R_1 | radr1_2 | Poland | R | 0 | 0 | 0 | 0 | 0 | 0 | 1 | 190 | 1295,263845 | 133 | 24830,794 | 3349 | 1 | 1 | 1 | 1 | 0 | 0 | 0 | 0 | 1 | 0 | 1 | 0 | 0 | 0 | 0 | 0 |
| 51.34067 | 21.06008 | RAD_R_1 | radr1_3 | Poland | R | 0 | 0 | 0 | 0 | 0 | 0 | 1 | 190 | 1295,041067 | 133 | 25247,029 | 7153 | 1 | 1 | 0 | 1 | 0 | 0 | 1 | 0 | 0 | 0 | 1 | 0 | 0 | 0 | 1 | 0 |
| 51.34101 | 21.06506 | RAD_R_1 | radr1_4 | Poland | R | 1 | 1 | 0 | 0 | 0 | 0 | 1 | 195,636 | 1294,89035 | 133 | 15526,298 | 4655 | 0 | 1 | 1 | 1 | 0 | 0 | 0 | 0 | 1 | 0 | 0 | 1 | 0 | 0 | 0 | 0 |
| 51.34302 | 21.06641 | RAD_R_1 | radr1_5 | Poland | R | 0 | 0 | 0 | 0 | 0 | 0 | 1 | 195,934 | 1295,032122 | 133 | 20817,356 | 4618 | 1 | 1 | 1 | 1 | 0 | 0 | 1 | 0 | 1 | 0 | 1 | 0 | 0 | 0 | 1 | 0 |
| 51.31472 | 20.98458 | RAD_R_2 | radr2_1 | Poland | R | 0 | 0 | 0 | 0 | 0 | 0 | 0 | 183 | 1295,356675 | 136 | 8948,031 | 5667 | 1 | 1 | 0 | 0 | 1 | 1 | 1 | 1 | 1 | 0 | 0 | 1 | 1 | 0 | 0 | 0 |
| 51.31571 | 20.98068 | RAD_R_2 | radr2_2 | Poland | R | 0 | 0 | 0 | 0 | 0 | 0 | 0 | 187,556 | 1295,595058 | 136 | 9704,477 | 3634 | 0 | 1 | 0 | 0 | 0 | 1 | 1 | 0 | 1 | 0 | 0 | 1 | 0 | 0 | 0 | 0 |
| 51.31341 | 20.97849 | RAD_R_2 | radr2_3 | Poland | R | 0 | 0 | 0 | 0 | 0 | 0 | 1 | 182 | 1295,45691 | 136 | 17951,868 | 6054 | 1 | 1 | 0 | 0 | 1 | 0 | 1 | 1 | 1 | 0 | 0 | 1 | 1 | 1 | 0 | 0 |
| 51.31178 | 20.9807 | RAD_R_2 | radr2_4 | Poland | R | 1 | 1 | 0 | 0 | 0 | 0 | 1 | 181 | 1295,223771 | 136 | 51563,452 | 4748 | 1 | 0 | 0 | 0 | 1 | 0 | 0 | 0 | 0 | 0 | 0 | 0 | 0 | 1 | 0 | 0 |
| 51.31079 | 20.98556 | RAD_R_2 | radr2_5 | Poland | R | 3 | 1 | 0 | 0 | 0 | 0 | 1 | 180,016 | 1294,949515 | 136 | 32268,719 | 0 | 1 | 1 | 0 | 0 | 1 | 0 | 0 | 0 | 0 | 0 | 0 | 0 | 0 | 0 | 0 | 0 |
| 52.18354 | 20.66518 | ROK_R_1 | rokr1_1 | Poland | R | 0 | 0 | 0 | 0 | 0 | 0 | 1 | 90,834 | 1389,373457 | 36 | 11839 | 71 | 0 | 1 | 0 | 0 | 0 | 1 | 1 | 0 | 0 | 1 | 0 | 0 | 0 | 0 | 0 | 0 |
| 52.18593 | 20.66346 | ROK_R_1 | rokr1_2 | Poland | R | 0 | 0 | 0 | 0 | 0 | 0 | 1 | 89,841 | 1389,665449 | 36 | 6416 | 0 | 0 | 1 | 0 | 0 | 0 | 0 | 1 | 0 | 0 | 0 | 0 | 0 | 0 | 0 | 0 | 0 |
| 52.18829 | 20.66111 | ROK_R_1 | rokr1_3 | Poland | R | 0 | 0 | 0 | 0 | 0 | 0 | 0 | 90 | 1389,961574 | 36 | 7683 | 0 | 0 | 1 | 0 | 0 | 0 | 0 | 1 | 0 | 0 | 0 | 0 | 0 | 0 | 0 | 0 | 0 |
| 52.19073 | 20.6588 | ROK_R_1 | rokr1_4 | Poland | R | 0 | 0 | 0 | 0 | 0 | 0 | 0 | 90 | 1390,277151 | 36 | 7639 | 159 | 1 | 1 | 0 | 1 | 0 | 0 | 1 | 0 | 1 | 0 | 1 | 0 | 0 | 0 | 0 | 0 |
| 52.19314 | 20.65642 | ROK_R_1 | rokr1_5 | Poland | R | 0 | 0 | 0 | 0 | 0 | 0 | 1 | 89 | 1390,592005 | 36 | 10053 | 2412 | 0 | 1 | 0 | 0 | 0 | 0 | 1 | 1 | 1 | 0 | 0 | 1 | 0 | 0 | 1 | 0 |
| 48.65418 | 20.56025 | ROZ_R_1 | rozr1_1 | Slovakia | R | 0 | 0 | 1 | 1 | 1 | 1 | 0 | 388,001 | 1076,287892 | 367 | 27041,214 | 3597 | 1 | 1 | 0 | 1 | 0 | 0 | 0 | 0 | 0 | 0 | 0 | 1 | 0 | 0 | 0 | 0 |
| 48.65554 | 20.55625 | ROZ_R_1 | rozr1_2 | Slovakia | R | 0 | 0 | 2 | 1 | 0 | 0 | 0 | 389,972 | 1076,596373 | 367 | 44537,7 | 40 | 1 | 1 | 0 | 1 | 0 | 0 | 1 | 0 | 0 | 0 | 0 | 0 | 0 | 0 | 0 | 0 |
| 48.65466 | 20.55205 | ROZ_R_1 | rozr1_3 | Slovakia | R | 0 | 0 | 0 | 0 | 0 | 0 | 0 | 363,017 | 1076,731855 | 367 | 9268,407 | 4739 | 0 | 1 | 0 | 1 | 0 | 0 | 1 | 0 | 1 | 0 | 1 | 0 | 0 | 0 | 0 | 0 |
| 48.65749 | 20.54924 | ROZ_R_1 | rozr1_4 | Slovakia | R | 0 | 0 | 0 | 0 | 0 | 0 | 0 | 360,037 | 1077,103596 | 367 | 6953,012 | 1488 | 1 | 1 | 0 | 1 | 0 | 0 | 1 | 1 | 0 | 0 | 1 | 0 | 0 | 0 | 1 | 0 |
| 48.65752 | 20.55261 | ROZ_R_1 | rozr1_5 | Slovakia | R | 0 | 0 | 0 | 0 | 0 | 0 | 0 | 381,936 | 1076,940579 | 367 | 23149,801 | 836 | 0 | 1 | 0 | 0 | 0 | 0 | 1 | 0 | 1 | 0 | 1 | 0 | 1 | 0 | 0 | 0 |
| 48.59913 | 20.47425 | ROZ_R_2 | rozr2_1 | Slovakia | R | 0 | 0 | 0 | 0 | 0 | 0 | 0 | 244,06 | 1076,009947 | 356 | 40283,008 | 1500 | 1 | 1 | 0 | 1 | 0 | 0 | 1 | 0 | 0 | 0 | 0 | 0 | 0 | 0 | 1 | 0 |
| 48.60133 | 20.47509 | ROZ_R_2 | rozr2_2 | Slovakia | R | 0 | 0 | 0 | 0 | 0 | 0 | 0 | 245,533 | 1076,149506 | 356 | 43307,391 | 2000 | 1 | 1 | 0 | 1 | 0 | 0 | 1 | 0 | 0 | 0 | 1 | 0 | 0 | 0 | 1 | 0 |
| 48.60389 | 20.47692 | ROZ_R_2 | rozr2_3 | Slovakia | R | 0 | 0 | 0 | 0 | 0 | 0 | 0 | 247,848 | 1076,267212 | 356 | 25423,363 | 1000 | 1 | 1 | 0 | 1 | 0 | 0 | 1 | 0 | 0 | 0 | 0 | 0 | 0 | 0 | 1 | 0 |
| 48.6056 | 20.4818 | ROZ_R_2 | rozr2_4 | Slovakia | R | 0 | 0 | 0 | 0 | 0 | 0 | 0 | 248,403 | 1076,165167 | 356 | 5342,284 | 0 | 1 | 1 | 0 | 1 | 0 | 0 | 1 | 0 | 0 | 0 | 0 | 0 | 0 | 0 | 0 | 0 |
| 48.60388 | 20.47691 | ROZ_R_2 | rozr2_5 | Slovakia | R | 1 | 1 | 0 | 0 | 0 | 0 | 0 | 247,848 | 1076,280395 | 356 | 18870,695 | 0 | 1 | 1 | 0 | 1 | 0 | 0 | 1 | 0 | 0 | 0 | 0 | 0 | 0 | 0 | 0 | 0 |
| 48.66044 | 20.52754 | ROZ_U_1 | rozu1_1 | Slovakia | U | 2 | 1 | 0 | 0 | 0 | 0 | 0 | 300,644 | 1078,417379 | 367 | 13183,271 | 15080 | 0 | 1 | 0 | 0 | 0 | 0 | 1 | 1 | 0 | 0 | 0 | 1 | 0 | 1 | 0 | 0 |
| 48.66192 | 20.52561 | ROZ_U_1 | rozu1_2 | Slovakia | U | 1 | 1 | 0 | 0 | 0 | 0 | 0 | 302,588 | 1078,63442 | 367 | 22894,343 | 9322 | 1 | 1 | 0 | 1 | 0 | 0 | 1 | 0 | 1 | 0 | 0 | 1 | 0 | 0 | 0 | 0 |
| 48.6646 | 20.52721 | ROZ_U_1 | rozu1_3 | Slovakia | U | 4 | 1 | 0 | 0 | 0 | 0 | 0 | 306,03 | 1078,77629 | 367 | 38683,98 | 1689 | 1 | 1 | 0 | 1 | 0 | 1 | 0 | 1 | 0 | 1 | 1 | 0 | 0 | 0 | 0 | 0 |
| 48.66553 | 20.52999 | ROZ_U_1 | rozu1_4 | Slovakia | U | 1 | 1 | 1 | 1 | 0 | 0 | 0 | 315,872 | 1078,717453 | 367 | 11812,685 | 4149 | 1 | 1 | 0 | 1 | 0 | 1 | 0 | 1 | 0 | 0 | 0 | 1 | 0 | 0 | 1 | 0 |
| 48.66342 | 20.53202 | ROZ_U_1 | rozu1_5 | Slovakia | U | 0 | 0 | 0 | 0 | 0 | 0 | 0 | 315,04 | 1078,442249 | 367 | 3834,291 | 18987 | 0 | 1 | 0 | 0 | 0 | 0 | 0 | 1 | 1 | 0 | 0 | 1 | 0 | 1 | 0 | 0 |
| 48.65252 | 20.54119 | ROZ_U_2 | rozu2_1 | Slovakia | U | 1 | 1 | 0 | 0 | 0 | 0 | 0 | 318,212 | 1077,088631 | 367 | 23348,146 | 5498 | 1 | 1 | 0 | 1 | 0 | 0 | 1 | 1 | 0 | 1 | 0 | 0 | 1 | 0 | 0 | 0 |
| 48.65081 | 20.53842 | ROZ_U_2 | rozu2_2 | Slovakia | U | 0 | 0 | 1 | 1 | 0 | 0 | 0 | 308,312 | 1077,085087 | 367 | 9461,584 | 10517 | 0 | 1 | 1 | 0 | 0 | 0 | 0 | 1 | 0 | 0 | 0 | 0 | 1 | 0 | 0 | 0 |
| 48.64883 | 20.53716 | ROZ_U_2 | rozu2_3 | Slovakia | U | 0 | 0 | 0 | 0 | 0 | 0 | 0 | 305,988 | 1076,982479 | 367 | 18995,471 | 12964 | 0 | 1 | 1 | 0 | 0 | 0 | 0 | 1 | 0 | 0 | 0 | 0 | 1 | 0 | 1 | 0 |
| 48.64714 | 20.53233 | ROZ_U_2 | rozu2_4 | Slovakia | U | 0 | 0 | 0 | 0 | 0 | 0 | 0 | 290,431 | 1077,08223 | 367 | 10491,467 | 6846 | 1 | 1 | 1 | 1 | 0 | 0 | 1 | 0 | 0 | 0 | 0 | 0 | 1 | 0 | 0 | 0 |
| 48.64869 | 20.52896 | ROZ_U_2 | rozu2_5 | Slovakia | U | 0 | 0 | 0 | 0 | 0 | 0 | 0 | 291,467 | 1077,376362 | 367 | 3505,06 | 10400 | 0 | 1 | 0 | 0 | 0 | 0 | 0 | 0 | 1 | 0 | 0 | 1 | 0 | 0 | 0 | 0 |
| 50.08756 | 21.87551 | RZE_R_1 | rzer1_1 | Poland | R | 0 | 0 | 2 | 1 | 0 | 0 | 0 | 210,035 | 1146,242743 | 278 | 21461,935 | 0 | 1 | 0 | 0 | 1 | 0 | 0 | 0 | 0 | 0 | 0 | 0 | 0 | 0 | 0 | 0 | 1 |
| 50.08758 | 21.87182 | RZE_R_1 | rzer1_2 | Poland | R | 1 | 1 | 0 | 0 | 1 | 1 | 0 | 210,447 | 1146,38687 | 278 | 8818,156 | 0 | 1 | 1 | 0 | 1 | 0 | 0 | 1 | 0 | 0 | 0 | 0 | 0 | 0 | 0 | 0 | 0 |
| 50.0863 | 21.86905 | RZE_R_1 | rzer1_3 | Poland | R | 0 | 0 | 0 | 0 | 0 | 0 | 0 | 210,606 | 1146,369129 | 278 | 0 | 0 | 0 | 0 | 0 | 0 | 0 | 0 | 0 | 0 | 0 | 0 | 0 | 0 | 0 | 0 | 0 | 0 |
| 50.0838 | 21.86948 | RZE_R_1 | rzer1_4 | Poland | R | 0 | 0 | 1 | 1 | 1 | 1 | 0 | 212,593 | 1146,113845 | 278 | 2029,252 | 0 | 0 | 1 | 0 | 0 | 0 | 0 | 1 | 0 | 0 | 0 | 0 | 0 | 0 | 0 | 0 | 0 |
| 50.08299 | 21.86508 | RZE_R_1 | rzer1_5 | Poland | R | 0 | 0 | 0 | 0 | 0 | 0 | 0 | 213,78 | 1146,206117 | 278 | 3542,561 | 0 | 0 | 1 | 0 | 0 | 0 | 0 | 1 | 0 | 0 | 0 | 0 | 0 | 0 | 0 | 0 | 0 |
| 50.0722 | 21.83677 | RZE_R_2 | rzer2_1 | Poland | R | 0 | 0 | 0 | 0 | 0 | 0 | 0 | 233,05 | 1146,27482 | 278 | 13689,379 | 8237 | 1 | 1 | 1 | 0 | 0 | 0 | 0 | 1 | 1 | 0 | 0 | 1 | 0 | 0 | 0 | 0 |
| 50.07264 | 21.83967 | RZE_R_2 | rzer2_2 | Poland | R | 0 | 0 | 0 | 0 | 0 | 0 | 1 | 232,221 | 1146,204359 | 278 | 11615,627 | 7646 | 1 | 1 | 1 | 0 | 0 | 0 | 0 | 1 | 0 | 0 | 0 | 1 | 0 | 0 | 0 | 0 |
| 50.07506 | 21.83931 | RZE_R_2 | rzer2_3 | Poland | R | 2 | 1 | 2 | 1 | 0 | 0 | 1 | 229,679 | 1146,448705 | 278 | 34944,054 | 9088 | 1 | 0 | 0 | 1 | 1 | 0 | 0 | 1 | 0 | 0 | 0 | 0 | 1 | 1 | 0 | 1 |
| 50.0774 | 21.83835 | RZE_R_2 | rzer2_4 | Poland | R | 2 | 1 | 2 | 1 | 0 | 0 | 1 | 220,338 | 1146,70319 | 278 | 12653,622 | 5243 | 1 | 1 | 0 | 0 | 1 | 0 | 0 | 1 | 1 | 0 | 0 | 1 | 1 | 0 | 1 | 0 |
| 50.07929 | 21.83665 | RZE_R_2 | rzer2_5 | Poland | R | 0 | 0 | 0 | 0 | 0 | 0 | 0 | 218,523 | 1146,946445 | 278 | 6970,505 | 7303 | 0 | 1 | 1 | 0 | 0 | 0 | 0 | 1 | 1 | 0 | 0 | 1 | 0 | 0 | 1 | 0 |
| 50.07943 | 22.03171 | RZE_R_3 | rzer3_1 | Poland | R | 0 | 0 | 0 | 0 | 0 | 0 | 1 | 203 | 1139,588807 | 280 | 26602,027 | 10602 | 0 | 1 | 0 | 0 | 0 | 0 | 0 | 0 | 1 | 0 | 0 | 1 | 0 | 0 | 0 | 0 |
| 50.07805 | 22.03497 | RZE_R_3 | rzer3_2 | Poland | R | 1 | 1 | 0 | 0 | 0 | 0 | 1 | 201,126 | 1139,321283 | 280 | 13596,876 | 13890 | 1 | 1 | 0 | 1 | 0 | 0 | 0 | 0 | 1 | 0 | 0 | 1 | 0 | 0 | 0 | 0 |
| 50.07854 | 22.03973 | RZE_R_3 | rzer3_3 | Poland | R | 1 | 1 | 0 | 0 | 0 | 0 | 0 | 198,979 | 1139,199288 | 280 | 8260,087 | 10506 | 0 | 1 | 0 | 0 | 0 | 0 | 0 | 0 | 1 | 0 | 1 | 1 | 1 | 0 | 0 | 0 |
| 50.0813 | 22.04024 | RZE_R_3 | rzer3_4 | Poland | R | 0 | 0 | 0 | 0 | 0 | 0 | 0 | 198 | 1139,432518 | 280 | 22333,287 | 5811 | 0 | 1 | 0 | 0 | 0 | 0 | 0 | 1 | 1 | 0 | 1 | 0 | 0 | 0 | 0 | 0 |
| 50.08362 | 22.04235 | RZE_R_3 | rzer3_5 | Poland | R | 1 | 1 | 0 | 0 | 0 | 0 | 0 | 198 | 1139,579561 | 280 | 30508,431 | 5269 | 0 | 1 | 0 | 0 | 0 | 0 | 1 | 0 | 1 | 0 | 1 | 0 | 0 | 0 | 0 | 0 |
| 50.00826 | 21.98867 | RZE_R_4 | rzer4_1 | Poland | R | 0 | 0 | 0 | 0 | 0 | 0 | 0 | 200,528 | 1134,472208 | 288 | 30256,148 | 1689 | 1 | 1 | 0 | 1 | 0 | 0 | 1 | 0 | 0 | 1 | 1 | 0 | 0 | 0 | 0 | 1 |
| 50.00658 | 21.9849 | RZE_R_4 | rzer4_2 | Poland | R | 0 | 0 | 0 | 0 | 0 | 0 | 0 | 201,048 | 1134,455117 | 288 | 24837,824 | 255 | 1 | 1 | 0 | 1 | 0 | 0 | 1 | 0 | 0 | 1 | 0 | 0 | 0 | 0 | 0 | 0 |
| 50.00409 | 21.98121 | RZE_R_4 | rzer4_3 | Poland | R | 1 | 1 | 1 | 1 | 0 | 0 | 0 | 200,626 | 1134,338192 | 288 | 20405,461 | 0 | 1 | 1 | 0 | 1 | 0 | 0 | 1 | 0 | 0 | 1 | 0 | 0 | 0 | 0 | 0 | 0 |
| 50.00414 | 21.97712 | RZE_R_4 | rzer4_4 | Poland | R | 0 | 0 | 0 | 0 | 0 | 0 | 0 | 203,551 | 1134,484449 | 288 | 47099,417 | 1442 | 1 | 1 | 0 | 1 | 0 | 0 | 0 | 0 | 1 | 0 | 1 | 0 | 0 | 0 | 0 | 0 |
| 50.00621 | 21.97927 | RZE_R_4 | rzer4_5 | Poland | R | 1 | 1 | 0 | 0 | 0 | 0 | 0 | 215,116 | 1134,625243 | 288 | 26887,716 | 3265 | 1 | 1 | 0 | 1 | 0 | 0 | 1 | 0 | 1 | 0 | 1 | 0 | 0 | 0 | 1 | 0 |
| 50.03976 | 21.98108 | RZE_U_1 | rzeu1_1 | Poland | U | 0 | 0 | 0 | 0 | 0 | 0 | 0 | 210,864 | 1137,727606 | 285 | 8996,037 | 5835 | 0 | 1 | 0 | 0 | 1 | 0 | 1 | 1 | 0 | 0 | 0 | 0 | 1 | 0 | 0 | 0 |
| 50.03803 | 21.97925 | RZE_U_1 | rzeu1_2 | Poland | U | 1 | 1 | 0 | 0 | 0 | 0 | 1 | 211 | 1137,633059 | 285 | 19641,441 | 8601 | 0 | 1 | 1 | 0 | 1 | 0 | 0 | 1 | 0 | 0 | 0 | 1 | 1 | 0 | 0 | 0 |
| 50.03547 | 21.97797 | RZE_U_1 | rzeu1_3 | Poland | U | 0 | 0 | 0 | 0 | 0 | 0 | 0 | 213,213 | 1137,4413 | 285 | 7425,661 | 11609 | 0 | 1 | 1 | 0 | 0 | 0 | 1 | 1 | 0 | 0 | 1 | 0 | 1 | 0 | 0 | 1 |
| 50.0342 | 21.98082 | RZE_U_1 | rzeu1_4 | Poland | U | 1 | 1 | 0 | 0 | 1 | 1 | 0 | 210,162 | 1137,211009 | 285 | 10613,654 | 12546 | 0 | 1 | 1 | 0 | 0 | 0 | 0 | 1 | 1 | 0 | 0 | 1 | 1 | 1 | 0 | 0 |
| 50.03573 | 21.98327 | RZE_U_1 | rzeu1_5 | Poland | U | 0 | 0 | 0 | 0 | 0 | 0 | 0 | 209 | 1137,264493 | 285 | 10674,687 | 14153 | 0 | 1 | 1 | 0 | 0 | 0 | 0 | 1 | 1 | 0 | 0 | 1 | 1 | 0 | 0 | 0 |
| 50.02662 | 22.04317 | RZE_U_2 | rzeu2_1 | Poland | U | 0 | 0 | 0 | 0 | 0 | 0 | 0 | 208 | 1134,142122 | 286 | 13648,64 | 8893 | 1 | 1 | 1 | 0 | 1 | 0 | 0 | 1 | 1 | 0 | 1 | 1 | 1 | 1 | 0 | 0 |
| 50.0255 | 22.04057 | RZE_U_2 | rzeu2_2 | Poland | U | 1 | 1 | 0 | 0 | 0 | 0 | 1 | 209,2 | 1134,134288 | 286 | 45333,32 | 4407 | 1 | 0 | 0 | 0 | 1 | 0 | 0 | 1 | 0 | 0 | 0 | 0 | 0 | 1 | 0 | 0 |
| 50.02377 | 22.04011 | RZE_U_2 | rzeu2_3 | Poland | U | 1 | 1 | 1 | 1 | 1 | 1 | 1 | 215,428 | 1133,988802 | 286 | 21700,601 | 7853 | 1 | 1 | 1 | 0 | 1 | 0 | 0 | 1 | 0 | 0 | 0 | 1 | 0 | 1 | 0 | 1 |
| 50.02469 | 22.03698 | RZE_U_2 | rzeu2_4 | Poland | U | 0 | 0 | 0 | 0 | 0 | 0 | 1 | 211,116 | 1134,192754 | 286 | 10675,419 | 10439 | 0 | 1 | 1 | 0 | 0 | 0 | 0 | 0 | 1 | 0 | 0 | 1 | 1 | 0 | 0 | 0 |
| 50.02715 | 22.0355 | RZE_U_2 | rzeu2_5 | Poland | U | 0 | 0 | 0 | 0 | 0 | 0 | 0 | 204,74 | 1134,481929 | 286 | 10914,251 | 9703 | 0 | 1 | 1 | 0 | 0 | 0 | 1 | 1 | 1 | 0 | 0 | 1 | 0 | 0 | 1 | 0 |
| 50.04805 | 22.00252 | RZE_U_3 | rzeu3_1 | Poland | U | 2 | 1 | 0 | 0 | 0 | 0 | 1 | 207 | 1137,690674 | 282 | 26602,027 | 11434 | 1 | 1 | 0 | 0 | 1 | 0 | 1 | 1 | 0 | 0 | 0 | 0 | 1 | 0 | 0 | 0 |
| 50.04892 | 22.00666 | RZE_U_3 | rzeu3_2 | Poland | U | 0 | 0 | 0 | 0 | 0 | 0 | 1 | 201,003 | 1137,66426 | 282 | 13596,876 | 8456 | 0 | 1 | 1 | 0 | 0 | 0 | 0 | 1 | 1 | 0 | 0 | 1 | 1 | 0 | 0 | 0 |
| 50.04893 | 22.01097 | RZE_U_3 | rzeu3_3 | Poland | U | 0 | 0 | 0 | 0 | 0 | 0 | 0 | 199,162 | 1137,493937 | 282 | 8260,087 | 14449 | 0 | 1 | 0 | 0 | 0 | 0 | 0 | 1 | 1 | 0 | 0 | 1 | 1 | 0 | 1 | 0 |
| 50.04896 | 22.01503 | RZE_U_3 | rzeu3_4 | Poland | U | 0 | 0 | 0 | 0 | 0 | 0 | 1 | 199,232 | 1137,348274 | 282 | 22333,287 | 0 | 1 | 1 | 0 | 1 | 0 | 0 | 0 | 1 | 1 | 1 | 1 | 0 | 0 | 0 | 0 | 1 |

| | | | | | | | | | | | | | | | | | | | | | | | | | | | | | | | | | |
|---|---|---|---|---|---|---|---|---|---|---|---|---|---|---|---|---|---|---|---|---|---|---|---|---|---|---|---|---|---|---|---|---|---|
| 50.05009 | 22.01795 | RZE_U_3 | rzeu3_5 | Poland | U | 0 | 0 | 1 | 1 | 0 | 0 | 1 | 199,62 | 1137,314406 | 282 | 30508,431 | 100 | 1 | 1 | 0 | 1 | 0 | 0 | 1 | 1 | 1 | 1 | 1 | 0 | 0 | 0 | 0 | 0 |
| 50.03638 | 22.01257 | RZE_U_4 | rzeu4_1 | Poland | U | 2 | 1 | 0 | 0 | 0 | 0 | 1 | 200,032 | 1136,225459 | 285 | 28366,057 | 8802 | 1 | 1 | 0 | 0 | 1 | 0 | 0 | 1 | 0 | 0 | 0 | 0 | 0 | 1 | 0 | 0 |
| 50.03365 | 22.01096 | RZE_U_4 | rzeu4_2 | Poland | U | 0 | 0 | 0 | 0 | 0 | 0 | 0 | 198,888 | 1136,044791 | 285 | 7540,29 | 8205 | 0 | 1 | 0 | 0 | 0 | 0 | 0 | 1 | 0 | 0 | 1 | 0 | 1 | 0 | 1 | 0 |
| 50.03207 | 22.00686 | RZE_U_4 | rzeu4_3 | Poland | U | 0 | 0 | 2 | 1 | 0 | 0 | 1 | 198,167 | 1136,021493 | 285 | 23111,132 | 7183 | 1 | 1 | 0 | 0 | 1 | 0 | 1 | 1 | 0 | 0 | 0 | 1 | 0 | 1 | 1 | 0 |
| 50.02975 | 22.0039 | RZE_U_4 | rzeu4_4 | Poland | U | 0 | 0 | 0 | 0 | 0 | 0 | 0 | 197,928 | 1135,935785 | 285 | 7478,825 | 4588 | 0 | 1 | 0 | 1 | 0 | 0 | 0 | 1 | 1 | 1 | 0 | 1 | 1 | 0 | 1 | 0 |
| 50.02723 | 22.00177 | RZE_U_4 | rzeu4_5 | Poland | U | 1 | 1 | 0 | 0 | 0 | 0 | 1 | 199,39 | 1135,790071 | 285 | 17160,394 | 6822 | 0 | 1 | 0 | 1 | 0 | 0 | 0 | 1 | 0 | 1 | 0 | 0 | 1 | 0 | 0 | 1 |
| 48.0486 | 19.83049 | SAL_R_1 | salr1_1 | Hungary | R | 0 | 0 | 0 | 0 | 0 | 0 | 1 | 229,657 | 1066,117366 | 293 | 5012,282 | 1736 | 0 | 1 | 0 | 0 | 0 | 0 | 1 | 0 | 1 | 0 | 1 | 1 | 0 | 0 | 0 | 0 |
| 48.04872 | 19.8254 | SAL_R_1 | salr1_2 | Hungary | R | 0 | 0 | 0 | 0 | 0 | 0 | 0 | 221,508 | 1066,404266 | 293 | 14597,69 | 6941 | 0 | 1 | 0 | 0 | 0 | 0 | 0 | 1 | 1 | 0 | 0 | 1 | 0 | 0 | 0 | 0 |
| 48.04875 | 19.82046 | SAL_R_1 | salr1_3 | Hungary | R | 0 | 0 | 1 | 1 | 0 | 0 | 0 | 221,672 | 1066,675882 | 293 | 10685,795 | 7567 | 0 | 1 | 0 | 0 | 0 | 0 | 0 | 1 | 1 | 1 | 0 | 1 | 0 | 0 | 0 | 0 |
| 48.05036 | 19.81729 | SAL_R_1 | salr1_4 | Hungary | R | 0 | 0 | 1 | 1 | 0 | 0 | 1 | 227,356 | 1066,970907 | 293 | 13354,318 | 6922 | 1 | 1 | 1 | 1 | 0 | 0 | 0 | 1 | 1 | 0 | 0 | 1 | 0 | 0 | 0 | 0 |
| 48.05129 | 19.81384 | SAL_R_1 | salr1_5 | Hungary | R | 0 | 0 | 0 | 0 | 0 | 0 | 0 | 225,356 | 1067,229537 | 293 | 18744,829 | 7048 | 1 | 1 | 1 | 1 | 0 | 0 | 0 | 0 | 1 | 0 | 0 | 1 | 0 | 0 | 0 | 0 |
| 47.99437 | 19.86542 | SAL_R_2 | salr2_1 | Hungary | R | 0 | 0 | 0 | 0 | 0 | 0 | 0 | 197,244 | 1060,102643 | 293 | 18249,362 | 0 | 1 | 1 | 0 | 1 | 0 | 0 | 1 | 0 | 0 | 1 | 0 | 0 | 0 | 0 | 0 | 0 |
| 47.99278 | 19.86205 | SAL_R_2 | salr2_2 | Hungary | R | 0 | 0 | 0 | 0 | 0 | 0 | 0 | 196,989 | 1060,166929 | 293 | 2637,211 | 0 | 0 | 1 | 0 | 0 | 0 | 0 | 0 | 0 | 0 | 1 | 0 | 0 | 0 | 0 | 0 | 0 |
| 47.99352 | 19.85866 | SAL_R_2 | salr2_3 | Hungary | R | 0 | 0 | 0 | 0 | 0 | 0 | 0 | 194,832 | 1060,408283 | 293 | 14639,604 | 0 | 0 | 1 | 0 | 1 | 0 | 0 | 0 | 0 | 0 | 1 | 0 | 0 | 0 | 0 | 0 | 0 |
| 47.99305 | 19.85507 | SAL_R_2 | salr2_4 | Hungary | R | 0 | 0 | 0 | 0 | 0 | 0 | 0 | 193,02 | 1060,569247 | 293 | 5325,492 | 0 | 0 | 1 | 0 | 1 | 0 | 0 | 0 | 0 | 0 | 1 | 0 | 0 | 0 | 0 | 0 | 0 |
| 47.99348 | 19.85059 | SAL_R_2 | salr2_5 | Hungary | R | 0 | 0 | 2 | 1 | 0 | 0 | 0 | 192,065 | 1060,847001 | 293 | 5406,295 | 0 | 0 | 1 | 0 | 0 | 0 | 0 | 0 | 0 | 0 | 1 | 0 | 0 | 0 | 0 | 0 | 0 |
| 48.10329 | 19.8077 | SAL_U_1 | salu1_1 | Hungary | U | 0 | 0 | 0 | 0 | 0 | 0 | 0 | 254,764 | 1071,510045 | 293 | 7021,095 | 6387 | 0 | 1 | 0 | 0 | 0 | 0 | 1 | 1 | 0 | 0 | 0 | 0 | 0 | 1 | 0 | 0 |
| 48.10425 | 19.8103 | SAL_U_1 | salu1_2 | Hungary | U | 1 | 1 | 0 | 0 | 0 | 0 | 0 | 275,824 | 1071,44205 | 293 | 32634,141 | 16255 | 1 | 1 | 1 | 1 | 0 | 0 | 0 | 1 | 0 | 0 | 0 | 0 | 0 | 1 | 0 | 0 |
| 48.10563 | 19.81215 | SAL_U_1 | salu1_3 | Hungary | U | 1 | 1 | 0 | 0 | 0 | 0 | 0 | 287,575 | 1071,44641 | 293 | 19404,572 | 10398 | 1 | 1 | 1 | 1 | 0 | 0 | 0 | 1 | 0 | 0 | 0 | 0 | 1 | 0 | 0 | 0 |
| 48.1071 | 19.81116 | SAL_U_1 | salu1_4 | Hungary | U | 1 | 1 | 0 | 0 | 0 | 0 | 0 | 260,109 | 1071,612085 | 293 | 10220,499 | 16630 | 0 | 1 | 1 | 0 | 0 | 1 | 0 | 1 | 0 | 0 | 0 | 0 | 1 | 0 | 0 | 0 |
| 48.10826 | 19.81521 | SAL_U_1 | salu1_5 | Hungary | U | 0 | 0 | 0 | 0 | 0 | 0 | 0 | 268,699 | 1071,480293 | 293 | 26889,145 | 19848 | 0 | 1 | 1 | 0 | 0 | 1 | 0 | 1 | 0 | 1 | 1 | 0 | 1 | 0 | 1 | 0 |
| 48.01489 | 19.83609 | SAL_U_2 | salu2_1 | Hungary | U | 0 | 0 | 0 | 0 | 0 | 0 | 0 | 188,046 | 1063,258922 | 293 | 10466,23 | 1277 | 1 | 1 | 1 | 1 | 0 | 0 | 1 | 1 | 1 | 0 | 1 | 0 | 0 | 0 | 0 | 0 |
| 48.01173 | 19.83656 | SAL_U_2 | salu2_2 | Hungary | U | 5 | 1 | 0 | 0 | 0 | 0 | 0 | 195,772 | 1062,994167 | 293 | 28043,945 | 3140 | 1 | 1 | 1 | 1 | 0 | 0 | 0 | 1 | 0 | 1 | 1 | 1 | 0 | 0 | 0 | 0 |
| 48.0092 | 19.83612 | SAL_U_2 | salu2_3 | Hungary | U | 2 | 1 | 0 | 0 | 0 | 0 | 0 | 200,032 | 1062,826694 | 293 | 56336,88 | 4111 | 1 | 1 | 1 | 1 | 1 | 0 | 0 | 0 | 0 | 0 | 1 | 1 | 0 | 0 | 0 | 0 |
| 48.00599 | 19.83601 | SAL_U_2 | salu2_4 | Hungary | U | 0 | 0 | 0 | 0 | 1 | 1 | 0 | 195,795 | 1062,590482 | 293 | 11656,079 | 12669 | 1 | 1 | 0 | 1 | 0 | 0 | 0 | 1 | 1 | 0 | 0 | 1 | 0 | 0 | 0 | 0 |
| 48.00309 | 19.83494 | SAL_U_2 | salu2_5 | Hungary | U | 0 | 0 | 0 | 0 | 0 | 0 | 0 | 190,059 | 1062,429947 | 293 | 4008,922 | 5324 | 0 | 1 | 0 | 0 | 0 | 0 | 0 | 1 | 1 | 0 | 1 | 1 | 0 | 0 | 0 | 0 |
| 46.28525 | 20.23487 | SEG_R_1 | segr1_1 | Hungary | R | 2 | 1 | 0 | 0 | 0 | 0 | 1 | 75 | 917,795916 | 267 | 30180,406 | 135 | 1 | 1 | 0 | 1 | 0 | 0 | 1 | 0 | 0 | 0 | 0 | 0 | 0 | 0 | 0 | 1 |
| 46.28314 | 20.23182 | SEG_R_1 | segr1_2 | Hungary | R | 3 | 1 | 0 | 0 | 0 | 0 | 1 | 74,552 | 917,8502418 | 267 | 23089,855 | 0 | 1 | 0 | 0 | 1 | 0 | 0 | 0 | 0 | 0 | 0 | 0 | 0 | 0 | 0 | 0 | 1 |
| 46.2811 | 20.22873 | SEG_R_1 | segr1_3 | Hungary | R | 5 | 1 | 0 | 0 | 0 | 0 | 1 | 75,04 | 917,9108563 | 267 | 20345,104 | 0 | 1 | 1 | 0 | 1 | 0 | 0 | 1 | 0 | 0 | 0 | 0 | 0 | 0 | 0 | 0 | 0 |
| 46.27896 | 20.22543 | SEG_R_1 | segr1_4 | Hungary | R | 2 | 1 | 0 | 0 | 0 | 0 | 1 | 74,548 | 917,9786944 | 267 | 27049,07 | 0 | 1 | 1 | 0 | 1 | 0 | 0 | 1 | 0 | 0 | 0 | 0 | 0 | 0 | 0 | 0 | 1 |
| 46.27668 | 20.22272 | SEG_R_1 | segr1_5 | Hungary | R | 3 | 1 | 0 | 0 | 0 | 0 | 1 | 77,942 | 918,0010231 | 267 | 19798,564 | 0 | 1 | 1 | 0 | 1 | 0 | 0 | 1 | 0 | 0 | 1 | 0 | 0 | 0 | 0 | 0 | 1 |
| 46.20692 | 20.11188 | SEG_R_2 | segr2_1 | Hungary | R | 0 | 0 | 3 | 1 | 0 | 0 | 1 | 74,088 | 920,5115353 | 255 | 10548,431 | 10205 | 0 | 1 | 0 | 0 | 0 | 0 | 0 | 0 | 1 | 0 | 0 | 1 | 0 | 0 | 0 | 0 |
| 46.20553 | 20.11428 | SEG_R_2 | segr2_2 | Hungary | R | 1 | 1 | 0 | 0 | 0 | 0 | 1 | 76,038 | 920,2711593 | 255 | 23445,558 | 4790 | 1 | 1 | 0 | 1 | 0 | 0 | 1 | 0 | 1 | 1 | 1 | 1 | 0 | 0 | 0 | 0 |
| 46.20281 | 20.11396 | SEG_R_2 | segr2_3 | Hungary | R | 2 | 1 | 2 | 1 | 0 | 0 | 1 | 79,024 | 920,1196083 | 255 | 34528,144 | 2713 | 1 | 1 | 0 | 1 | 0 | 0 | 1 | 1 | 1 | 1 | 0 | 1 | 0 | 0 | 0 | 1 |
| 46.20038 | 20.11156 | SEG_R_2 | segr2_4 | Hungary | R | 2 | 1 | 2 | 1 | 0 | 0 | 1 | 76,248 | 920,1176003 | 255 | 30484,515 | 1722 | 1 | 1 | 1 | 1 | 0 | 0 | 0 | 0 | 1 | 0 | 0 | 1 | 0 | 0 | 0 | 1 |
| 46.19803 | 20.10927 | SEG_R_2 | segr2_5 | Hungary | R | 4 | 1 | 2 | 1 | 0 | 0 | 1 | 75,372 | 920,1180529 | 255 | 31347,027 | 1805 | 1 | 0 | 0 | 1 | 0 | 0 | 0 | 0 | 1 | 0 | 0 | 1 | 0 | 0 | 0 | 1 |
| 46.24299 | 20.15686 | SEG_U_1 | segu1_1 | Hungary | U | 2 | 1 | 0 | 0 | 0 | 0 | 0 | 76,532 | 919,9701858 | 261 | 42503,628 | 14414 | 1 | 1 | 0 | 1 | 0 | 0 | 1 | 1 | 0 | 0 | 1 | 0 | 1 | 0 | 1 | 1 |
| 46.24457 | 20.1553 | SEG_U_1 | segu1_2 | Hungary | U | 3 | 1 | 0 | 0 | 0 | 0 | 1 | 84,124 | 920,1699877 | 261 | 43013,654 | 7294 | 1 | 1 | 0 | 1 | 0 | 0 | 1 | 1 | 0 | 1 | 0 | 0 | 1 | 0 | 0 | 0 |
| 46.2472 | 20.15613 | SEG_U_1 | segu1_3 | Hungary | U | 3 | 1 | 0 | 0 | 0 | 0 | 1 | 83,023 | 920,2862804 | 261 | 42889,866 | 7370 | 1 | 1 | 0 | 1 | 0 | 0 | 0 | 1 | 0 | 1 | 0 | 0 | 1 | 1 | 0 | 1 |
| 46.24875 | 20.15769 | SEG_U_1 | segu1_4 | Hungary | U | 3 | 1 | 0 | 0 | 0 | 0 | 1 | 81,316 | 920,2873812 | 261 | 20426,564 | 9652 | 1 | 1 | 0 | 1 | 1 | 0 | 0 | 1 | 0 | 0 | 0 | 0 | 1 | 1 | 0 | 1 |
| 46.24835 | 20.16123 | SEG_U_1 | segu1_5 | Hungary | U | 0 | 0 | 0 | 0 | 0 | 0 | 0 | 80,692 | 920,0387735 | 261 | 62781,951 | 2945 | 1 | 1 | 0 | 0 | 1 | 0 | 0 | 0 | 0 | 0 | 0 | 0 | 0 | 0 | 0 | 0 |
| 46.24822 | 20.18599 | SEG_U_2 | segu2_1 | Hungary | U | 3 | 1 | 2 | 1 | 0 | 0 | 1 | 78,822 | 918,471689 | 261 | 26788,219 | 8115 | 1 | 1 | 1 | 1 | 0 | 0 | 0 | 0 | 1 | 1 | 0 | 1 | 0 | 0 | 0 | 1 |
| 46.2504 | 20.18732 | SEG_U_2 | segu2_2 | Hungary | U | 2 | 1 | 0 | 0 | 0 | 0 | 1 | 77,69 | 918,5282155 | 261 | 38595,656 | 1862 | 1 | 1 | 0 | 1 | 0 | 0 | 0 | 0 | 1 | 1 | 1 | 1 | 0 | 0 | 0 | 1 |
| 46.2519 | 20.18455 | SEG_U_2 | segu2_3 | Hungary | U | 6 | 1 | 0 | 0 | 0 | 0 | 1 | 83,581 | 918,798863 | 261 | 50569,753 | 1066 | 1 | 0 | 0 | 1 | 0 | 0 | 0 | 0 | 0 | 1 | 1 | 0 | 0 | 0 | 0 | 1 |
| 46.25272 | 20.1808 | SEG_U_2 | segu2_4 | Hungary | U | 1 | 1 | 0 | 0 | 0 | 0 | 0 | 81,136 | 919,087345 | 261 | 48435,636 | 1897 | 1 | 0 | 0 | 1 | 0 | 0 | 0 | 0 | 0 | 1 | 1 | 1 | 0 | 0 | 0 | 0 |
| 46.25355 | 20.17702 | SEG_U_2 | segu2_5 | Hungary | U | 1 | 1 | 0 | 0 | 0 | 0 | 0 | 83,4 | 919,3785434 | 261 | 35859,105 | 3676 | 1 | 1 | 0 | 1 | 0 | 0 | 0 | 0 | 1 | 1 | 0 | 1 | 0 | 0 | 0 | 1 |
| 42.59243 | 25.41669 | STA_R_1 | star1_1 | Bulgaria | R | 0 | 0 | 0 | 0 | 0 | 0 | 0 | 336 | 348,7675971 | 515 | 26758,671 | 0 | 0 | 1 | 0 | 1 | 0 | 0 | 1 | 0 | 0 | 0 | 0 | 0 | 0 | 0 | 0 | 0 |
| 42.59019 | 25.41939 | STA_R_1 | star1_2 | Bulgaria | R | 0 | 0 | 0 | 0 | 0 | 0 | 1 | 335,938 | 348,4569323 | 515 | 8028,549 | 2131 | 0 | 1 | 0 | 0 | 0 | 0 | 1 | 0 | 0 | 0 | 0 | 0 | 1 | 0 | 0 | 0 |
| 42.58758 | 25.42188 | STA_R_1 | star1_3 | Bulgaria | R | 0 | 0 | 0 | 0 | 0 | 0 | 0 | 332,877 | 348,143781 | 515 | 6112,314 | 0 | 0 | 1 | 0 | 0 | 0 | 0 | 1 | 0 | 0 | 0 | 0 | 0 | 0 | 0 | 0 | 0 |
| 42.58496 | 25.42309 | STA_R_1 | star1_4 | Bulgaria | R | 0 | 0 | 0 | 0 | 0 | 0 | 0 | 332,73 | 347,9221765 | 515 | 5992,972 | 0 | 0 | 1 | 0 | 0 | 0 | 0 | 1 | 0 | 0 | 0 | 0 | 0 | 0 | 0 | 0 | 1 |
| 42.58259 | 25.42388 | STA_R_1 | star1_5 | Bulgaria | R | 0 | 0 | 0 | 0 | 0 | 0 | 0 | 334,676 | 347,7455667 | 515 | 26935,26 | 3259 | 0 | 1 | 0 | 0 | 1 | 0 | 1 | 1 | 1 | 0 | 1 | 0 | 0 | 0 | 0 | 0 |
| 42.57974 | 25.39947 | STA_R_2 | star2_1 | Bulgaria | R | 0 | 0 | 2 | 1 | 0 | 0 | 0 | 330 | 349,3884549 | 515 | 18758,581 | 0 | 0 | 1 | 0 | 0 | 0 | 0 | 1 | 0 | 0 | 1 | 0 | 0 | 0 | 0 | 0 | 0 |
| 42.57855 | 25.40251 | STA_R_2 | star2_2 | Bulgaria | R | 0 | 0 | 0 | 0 | 0 | 0 | 0 | 330,256 | 349,1065086 | 515 | 7041,774 | 100 | 0 | 1 | 0 | 0 | 0 | 0 | 0 | 0 | 0 | 1 | 0 | 0 | 0 | 0 | 0 | 0 |
| 42.57757 | 25.40618 | STA_R_2 | star2_3 | Bulgaria | R | 0 | 0 | 1 | 1 | 0 | 0 | 0 | 330,19 | 348,7881263 | 515 | 10023,19 | 69 | 0 | 1 | 0 | 0 | 0 | 0 | 1 | 0 | 0 | 1 | 1 | 0 | 0 | 0 | 0 | 0 |
| 42.57686 | 25.40992 | STA_R_2 | star2_4 | Bulgaria | R | 0 | 0 | 1 | 1 | 1 | 1 | 0 | 331,896 | 348,4775785 | 515 | 3707,797 | 0 | 0 | 1 | 0 | 0 | 0 | 0 | 1 | 0 | 0 | 1 | 0 | 0 | 0 | 0 | 0 | 0 |
| 42.57638 | 25.41373 | STA_R_2 | star2_5 | Bulgaria | R | 1 | 1 | 0 | 0 | 0 | 0 | 0 | 329,982 | 348,174625 | 515 | 7612,167 | 0 | 0 | 1 | 0 | 0 | 0 | 0 | 1 | 0 | 0 | 1 | 0 | 0 | 0 | 0 | 0 | 0 |
| 41.81562 | 26.13619 | SVI_R_1 | svir1_1 | Bulgaria | R | 0 | 0 | 2 | 1 | 0 | 0 | 0 | 58,616 | 261,0020353 | 472 | 6997,941 | 0 | 1 | 1 | 0 | 1 | 0 | 0 | 1 | 0 | 0 | 0 | 0 | 0 | 0 | 0 | 0 | 0 |
| 41.81631 | 26.13387 | SVI_R_1 | svir1_2 | Bulgaria | R | 1 | 1 | 2 | 1 | 0 | 0 | 0 | 57 | 261,14576 | 472 | 37960,841 | 0 | 1 | 1 | 0 | 1 | 0 | 0 | 0 | 0 | 0 | 1 | 0 | 0 | 0 | 0 | 0 | 0 |
| 41.81127 | 26.13606 | SVI_R_1 | svir1_3 | Bulgaria | R | 0 | 0 | 1 | 1 | 0 | 0 | 0 | 54,039 | 260,8735579 | 472 | 37843,11 | 0 | 1 | 1 | 0 | 1 | 0 | 0 | 0 | 0 | 0 | 1 | 0 | 0 | 0 | 0 | 0 | 1 |
| 41.81146 | 26.1404 | SVI_R_1 | svir1_4 | Bulgaria | R | 0 | 0 | 0 | 0 | 0 | 0 | 0 | 54,256 | 260,5349407 | 472 | 20534,824 | 0 | 0 | 1 | 0 | 1 | 0 | 0 | 0 | 0 | 0 | 1 | 0 | 0 | 0 | 0 | 0 | 0 |
| 41.81263 | 26.1431 | SVI_R_1 | svir1_5 | Bulgaria | R | 0 | 0 | 2 | 1 | 0 | 0 | 0 | 56 | 260,3577042 | 472 | 27205,613 | 0 | 1 | 1 | 0 | 1 | 0 | 0 | 0 | 0 | 0 | 1 | 0 | 0 | 0 | 0 | 0 | 0 |
| 41.81779 | 26.15813 | SVI_R_2 | svir2_1 | Bulgaria | R | 0 | 0 | 0 | 0 | 1 | 1 | 0 | 72,256 | 259,5604336 | 472 | 9231,082 | 0 | 1 | 1 | 0 | 1 | 0 | 0 | 1 | 0 | 0 | 0 | 0 | 0 | 0 | 0 | 0 | 1 |
| 41.81502 | 26.1597 | SVI_R_2 | svir2_2 | Bulgaria | R | 0 | 0 | 0 | 0 | 0 | 0 | 0 | 67,922 | 259,3507089 | 472 | 31115,099 | 120 | 1 | 1 | 0 | 1 | 0 | 0 | 1 | 0 | 0 | 1 | 0 | 0 | 0 | 0 | 0 | 0 |
| 41.81173 | 26.16146 | SVI_R_2 | svir2_3 | Bulgaria | R | 0 | 0 | 1 | 1 | 0 | 0 | 0 | 69,942 | 259,0903053 | 472 | 28383,548 | 0 | 1 | 1 | 0 | 1 | 0 | 0 | 1 | 0 | 0 | 0 | 0 | 0 | 0 | 0 | 0 | 0 |
| 41.80899 | 26.16298 | SVI_R_2 | svir2_4 | Bulgaria | R | 0 | 0 | 0 | 0 | 0 | 0 | 0 | 66,092 | 258,8167571 | 472 | 22731,751 | 0 | 1 | 1 | 0 | 1 | 0 | 0 | 1 | 0 | 0 | 0 | 0 | 0 | 0 | 0 | 0 | 0 |
| 41.80674 | 26.16498 | SVI_R_2 | svir2_5 | Bulgaria | R | 0 | 0 | 2 | 1 | 0 | 0 | 0 | 65,664 | 258,5390673 | 472 | 7468,179 | 0 | 0 | 1 | 0 | 0 | 0 | 0 | 1 | 0 | 0 | 0 | 0 | 0 | 0 | 0 | 0 | 0 |
| 41.7647 | 26.20737 | SVI_U_1 | sviu1_1 | Bulgaria | U | 0 | 0 | 0 | 0 | 0 | 0 | 0 | 58,431 | 253,7327034 | 474 | 14039,404 | 15081 | 0 | 1 | 0 | 0 | 1 | 0 | 0 | 1 | 0 | 0 | 0 | 1 | 0 | 1 | 0 | 0 |
| 41.76382 | 26.20955 | SVI_U_1 | sviu1_2 | Bulgaria | U | 0 | 0 | 0 | 0 | 0 | 0 | 0 | 60,188 | 253,5311357 | 474 | 25595,699 | 7994 | 0 | 1 | 0 | 0 | 1 | 0 | 0 | 1 | 0 | 0 | 0 | 1 | 0 | 1 | 0 | 0 |
| 41.7629 | 26.21204 | SVI_U_1 | sviu1_3 | Bulgaria | U | 0 | 0 | 0 | 0 | 0 | 0 | 0 | 53,904 | 253,3031425 | 474 | 9905,414 | 12227 | 0 | 1 | 0 | 0 | 1 | 0 | 1 | 0 | 0 | 0 | 0 | 1 | 0 | 0 | 1 | 0 |
| 41.76426 | 26.21442 | SVI_U_1 | sviu1_4 | Bulgaria | U | 0 | 0 | 0 | 0 | 0 | 0 | 1 | 57,066 | 253,1538116 | 474 | 14705,806 | 4941 | 1 | 1 | 0 | 0 | 0 | 0 | 0 | 1 | 0 | 0 | 0 | 1 | 0 | 0 | 1 | 0 |
| 41.76652 | 26.21431 | SVI_U_1 | sviu1_5 | Bulgaria | U | 0 | 0 | 0 | 0 | 0 | 0 | 1 | 58,788 | 253,2331915 | 474 | 6686,129 | 2006 | 0 | 1 | 1 | 0 | 0 | 0 | 1 | 0 | 1 | 0 | 0 | 1 | 0 | 0 | 0 | 0 |
| 41.76825 | 26.20258 | SVI_U_2 | sviu2_1 | Bulgaria | U | 0 | 0 | 0 | 0 | 0 | 0 | 0 | 56,4 | 254,2239471 | 474 | 7727,919 | 20651 | 0 | 1 | 0 | 0 | 0 | 0 | 0 | 1 | 0 | 0 | 0 | 1 | 0 | 0 | 1 | 0 |
| 41.77067 | 26.20252 | SVI_U_2 | sviu2_2 | Bulgaria | U | 0 | 0 | 1 | 1 | 0 | 0 | 0 | 57,03 | 254,3003025 | 474 | 10612,83 | 20131 | 0 | 1 | 0 | 0 | 1 | 0 | 0 | 1 | 0 | 0 | 0 | 1 | 0 | 0 | 0 | 0 |
| 41.77139 | 26.19976 | SVI_U_2 | sviu2_3 | Bulgaria | U | 0 | 0 | 0 | 0 | 0 | 0 | 0 | 54 | 254,5422853 | 474 | 10113,734 | 20012 | 0 | 1 | 0 | 0 | 1 | 0 | 0 | 1 | 1 | 0 | 1 | 1 | 1 | 0 | 1 | 0 |
| 41.77188 | 26.19601 | SVI_U_2 | sviu2_4 | Bulgaria | U | 0 | 0 | 1 | 1 | 0 | 0 | 0 | 56,94 | 254,8593005 | 474 | 26066,069 | 3454 | 1 | 1 | 1 | 0 | 1 | 0 | 0 | 1 | 0 | 0 | 0 | 0 | 1 | 0 | 1 | 0 |
| 41.77427 | 26.19721 | SVI_U_2 | sviu2_5 | Bulgaria | U | 0 | 0 | 0 | 0 | 0 | 0 | 0 | 51,372 | 254,8392151 | 474 | 25279,8 | 2000 | 1 | 1 | 1 | 0 | 1 | 0 | 1 | 0 | 0 | 0 | 0 | 0 | 0 | 0 | 1 | 0 |
| 50.97425 | 21.09598 | SWK_R_1 | swkr1_1 | Poland | R | 0 | 0 | 0 | 0 | 0 | 0 | 0 | 218,97 | 1259,148275 | 174 | 14674,171 | 0 | 0 | 1 | 0 | 0 | 0 | 0 | 1 | 0 | 0 | 1 | 0 | 0 | 0 | 0 | 0 | 0 |
| 50.97586 | 21.09848 | SWK_R_1 | swkr1_2 | Poland | R | 0 | 0 | 0 | 0 | 0 | 0 | 1 | 220,149 | 1259,20554 | 174 | 11389,355 | 0 | 0 | 1 | 0 | 0 | 0 | 0 | 0 | 0 | 0 | 1 | 0 | 0 | 0 | 0 | 0 | 0 |
| 50.97546 | 21.1018 | SWK_R_1 | swkr1_3 | Poland | R | 0 | 0 | 0 | 0 | 0 | 0 | 1 | 217,957 | 1259,038079 | 174 | 19145,577 | 0 | 0 | 1 | 0 | 0 | 0 | 0 | 1 | 0 | 0 | 1 | 0 | 0 | 0 | 0 | 0 | 1 |
| 50.97675 | 21.10494 | SWK_R_1 | swkr1_4 | Poland | R | 0 | 0 | 0 | 0 | 0 | 0 | 1 | 217,451 | 1259,045229 | 174 | 9479,553 | 0 | 0 | 1 | 0 | 0 | 0 | 0 | 0 | 0 | 0 | 1 | 0 | 0 | 0 | 0 | 0 | 1 |
| 50.97757 | 21.10917 | SWK_R_1 | swkr1_5 | Poland | R | 1 | 1 | 0 | 0 | 0 | 0 | 1 | 217,243 | 1258,964463 | 174 | 11741,594 | 200 | 0 | 1 | 0 | 0 | 0 | 0 | 0 | 0 | 0 | 1 | 0 | 0 | 0 | 0 | 0 | 1 |
| 50.97729 | 21.11284 | SWK_R_1 | swkr1_6 | Poland | R | 0 | 0 | 0 | 0 | 0 | 0 | 0 | 215,956 | 1258,79607 | 174 | 12918,443 | 0 | 0 | 1 | 0 | 0 | 0 | 0 | 0 | 0 | 0 | 1 | 0 | 0 | 0 | 0 | 0 | 0 |
| 50.97646 | 21.11667 | SWK_R_1 | swkr1_7 | Poland | R | 1 | 1 | 0 | 0 | 0 | 0 | 1 | 213,777 | 1258,575494 | 174 | 11691,924 | 0 | 0 | 1 | 0 | 0 | 0 | 0 | 0 | 0 | 0 | 1 | 0 | 0 | 0 | 0 | 0 | 0 |
| 50.97526 | 21.11999 | SWK_R_1 | swkr1_8 | Poland | R | 0 | 0 | 0 | 0 | 0 | 0 | 1 | 213,907 | 1258,340477 | 174 | 13795,874 | 0 | 0 | 1 | 0 | 0 | 0 | 0 | 0 | 0 | 0 | 1 | 0 | 0 | 0 | 0 | 0 | 0 |
| 50.97498 | 21.06288 | SWK_R_2 | swkr2_1 | Poland | R | 0 | 0 | 0 | 0 | 0 | 0 | 0 | 227,71 | 1260,467605 | 174 | 10304,093 | 1234 | 0 | 1 | 0 | 0 | 0 | 0 | 0 | 0 | 0 | 1 | 0 | 1 | 0 | 0 | 0 | 0 |
| 50.97377 | 21.06637 | SWK_R_2 | swkr2_2 | Poland | R | 0 | 0 | 0 | 0 | 0 | 0 | 0 | 226,971 | 1260,219861 | 174 | 13271,869 | 1032 | 0 | 1 | 0 | 0 | 0 | 0 | 1 | 0 | 1 | 1 | 0 | 1 | 0 | 0 | 0 | 0 |
| 50.97273 | 21.06999 | SWK_R_2 | swkr2_3 | Poland | R | 0 | 0 | 0 | 0 | 0 | 0 | 1 | 225,928 | 1259,986366 | 174 | 17832,712 | 745 | 0 | 1 | 0 | 0 | 0 | 0 | 1 | 0 | 1 | 1 | 0 | 1 | 0 | 0 | 0 | 0 |
| 50.97094 | 21.07267 | SWK_R_2 | swkr2_4 | Poland | R | 0 | 0 | 0 | 0 | 0 | 0 | 0 | 224,384 | 1259,716031 | 174 | 11551,545 | 617 | 0 | 1 | 0 | 0 | 0 | 0 | 1 | 0 | 0 | 1 | 1 | 0 | 0 | 0 | 0 | 0 |
| 50.97163 | 21.07636 | SWK_R_2 | swkr2_5 | Poland | R | 0 | 0 | 0 | 0 | 0 | 0 | 1 | 228,59 | 1259,640435 | 174 | 8037,611 | 2553 | 0 | 1 | 0 | 0 | 0 | 0 | 0 | 0 | 1 | 1 | 0 | 1 | 0 | 0 | 0 | 0 |
| 50.97015 | 21.07959 | SWK_R_2 | swkr2_6 | Poland | R | 2 | 1 | 0 | 0 | 0 | 0 | 1 | 223,54 | 1259,379922 | 174 | 8027,876 | 483 | 0 | 1 | 0 | 0 | 0 | 0 | 0 | 0 | 0 | 1 | 1 | 0 | 0 | 0 | 0 | 0 |
| 50.97106 | 21.08364 | SWK_R_2 | swkr2_7 | Poland | R | 0 | 0 | 0 | 0 | 0 | 0 | 0 | 223,019 | 1259,314573 | 174 | 15208,736 | 2045 | 0 | 1 | 0 | 0 | 0 | 0 | 1 | 0 | 1 | 1 | 0 | 1 | 0 | 0 | 0 | 0 |
| 50.9725 | 21.08611 | SWK_R_2 | swkr2_8 | Poland | R | 0 | 0 | 0 | 0 | 0 | 0 | 1 | 224 | 1259,355824 | 174 | 13235,383 | 1640 | 0 | 1 | 0 | 0 | 0 | 0 | 1 | 0 | 1 | 1 | 0 | 1 | 0 | 0 | 0 | 1 |
| 51.04892 | 21.06956 | SWK_U_1 | swku1_1 | Poland | U | 0 | 0 | 0 | 0 | 0 | 0 | 0 | 219,224 | 1267,151931 | 166 | 33821,155 | 8499 | 1 | 1 | 0 | 0 | 0 | 0 | 0 | 1 | 0 | 0 | 0 | 1 | 1 | 0 | 0 | 0 |
| 51.05055 | 21.07152 | SWK_U_1 | swku1_2 | Poland | U | 0 | 0 | 0 | 0 | 0 | 0 | 0 | 229,472 | 1267,23099 | 166 | 32305,029 | 5981 | 1 | 1 | 0 | 0 | 1 | 0 | 1 | 1 | 0 | 0 | 0 | 0 | 1 | 0 | 1 | 0 |
| 51.05172 | 21.0752 | SWK_U_1 | swku1_3 | Poland | U | 0 | 0 | 1 | 1 | 0 | 0 | 1 | 257,384 | 1267,207624 | 166 | 22655,313 | 8338 | 1 | 1 | 0 | 0 | 1 | 0 | 0 | 1 | 0 | 0 | 0 | 0 | 1 | 1 | 0 | 0 |
| 51.05234 | 21.07936 | SWK_U_1 | swku1_4 | Poland | U | 0 | 0 | 0 | 0 | 0 | 0 | 0 | 264,871 | 1267,111616 | 166 | 14643,59 | 13895 | 0 | 1 | 0 | 0 | 0 | 0 | 0 | 1 | 0 | 0 | 0 | 0 | 1 | 1 | 0 | 0 |
| 51.05069 | 21.08116 | SWK_U_1 | swku1_5 | Poland | U | 0 | 0 | 0 | 0 | 0 | 0 | 0 | 266 | 1266,887943 | 166 | 11980,454 | 9797 | 0 | 1 | 0 | 0 | 0 | 0 | 0 | 1 | 0 | 0 | 0 | 0 | 1 | 1 | 0 | 0 |
| 51.04815 | 21.08114 | SWK_U_1 | swku1_6 | Poland | U | 0 | 0 | 0 | 0 | 0 | 0 | 1 | 248,853 | 1266,650102 | 166 | 13003,454 | 8297 | 0 | 1 | 0 | 0 | 0 | 0 | 1 | 1 | 0 | 0 | 0 | 0 | 1 | 0 | 0 | 0 |
| 51.04565 | 21.0814 | SWK_U_1 | swku1_7 | Poland | U | 0 | 0 | 0 | 0 | 0 | 0 | 1 | 234,72 | 1266,404403 | 166 | 18447,173 | 7491 | 0 | 1 | 0 | 0 | 0 | 0 | 0 | 1 | 1 | 0 | 0 | 1 | 1 | 0 | 0 | 0 |
| 51.04373 | 21.07838 | SWK_U_1 | swku1_8 | Poland | U | 0 | 0 | 0 | 0 | 0 | 0 | 0 | 209,596 | 1266,337121 | 166 | 12415,151 | 13736 | 1 | 1 | 0 | 0 | 0 | 0 | 1 | 1 | 1 | 0 | 0 | 1 | 0 | 0 | 1 | 0 |
| 49.99293 | 20.89937 | TAR_R_1 | tarr1_1 | Poland | R | 0 | 0 | 1 | 1 | 0 | 0 | 0 | 231,816 | 1176,692475 | 267 | 21923,112 | 5891 | 1 | 1 | 1 | 1 | 0 | 0 | 0 | 1 | 0 | 0 | 1 | 1 | 0 | 0 | 0 | 0 |
| 49.99497 | 20.90055 | TAR_R_1 | tarr1_2 | Poland | R | 0 | 0 | 0 | 0 | 0 | 0 | 0 | 207,306 | 1176,826276 | 267 | 7574,082 | 7014 | 1 | 1 | 0 | 1 | 0 | 0 | 0 | 1 | 1 | 0 | 1 | 1 | 0 | 1 | 0 | 0 |
| 49.99498 | 20.89546 | TAR_R_1 | tarr1_3 | Poland | R | 0 | 0 | 0 | 0 | 0 | 0 | 0 | 191,344 | 1177,043217 | 267 | 6927,25 | 0 | 1 | 1 | 0 | 1 | 0 | 0 | 1 | 0 | 0 | 0 | 0 | 0 | 0 | 0 | 0 | 0 |

| | | | | | | | | | | | | | | | | | | | | | | | | | | | | | | | | | |
|---|---|---|---|---|---|---|---|---|---|---|---|---|---|---|---|---|---|---|---|---|---|---|---|---|---|---|---|---|---|---|---|---|---|
| 49.99244 | 20.89361 | TAR_R_1 | tarr1_4 | Poland | R | 1 | 1 | 0 | 0 | 0 | 0 | 1 | 193,57 | 1176,890725 | 267 | 8652,434 | 100 | 1 | 1 | 0 | 1 | 0 | 0 | 1 | 0 | 0 | 1 | 0 | 0 | 0 | 0 | 0 | 0 |
| 49.99034 | 20.89365 | TAR_R_1 | tarr1_5 | Poland | R | 2 | 1 | 0 | 0 | 0 | 0 | 1 | 192,109 | 1176,70211 | 267 | 20051,016 | 69 | 1 | 1 | 0 | 1 | 0 | 0 | 1 | 0 | 0 | 1 | 0 | 0 | 0 | 0 | 0 | 0 |
| 49.98546 | 20.89981 | TAR_R_2 | tarr2_1 | Poland | R | 2 | 1 | 0 | 0 | 0 | 0 | 0 | 227,934 | 1176,001195 | 267 | 41810,96 | 2435 | 1 | 0 | 1 | 1 | 0 | 0 | 0 | 0 | 1 | 0 | 0 | 1 | 0 | 0 | 0 | 0 |
| 49.98527 | 20.89569 | TAR_R_2 | tarr2_2 | Poland | R | 0 | 0 | 0 | 0 | 0 | 0 | 1 | 205,494 | 1176,158197 | 267 | 60689,761 | 548 | 1 | 0 | 0 | 1 | 0 | 0 | 0 | 0 | 0 | 0 | 0 | 0 | 0 | 0 | 0 | 0 |
| 49.98596 | 20.89181 | TAR_R_2 | tarr2_3 | Poland | R | 1 | 1 | 0 | 0 | 0 | 0 | 0 | 193,737 | 1176,384038 | 267 | 8462,934 | 0 | 1 | 0 | 0 | 1 | 0 | 0 | 0 | 0 | 0 | 0 | 0 | 0 | 0 | 0 | 0 | 0 |
| 49.98327 | 20.88921 | TAR_R_2 | tarr2_4 | Poland | R | 1 | 1 | 0 | 0 | 0 | 0 | 0 | 194 | 1176,252782 | 267 | 14766,243 | 0 | 1 | 0 | 0 | 1 | 0 | 0 | 0 | 0 | 0 | 1 | 0 | 0 | 0 | 0 | 0 | 0 |
| 49.981 | 20.88698 | TAR_R_2 | tarr2_5 | Poland | R | 0 | 0 | 0 | 0 | 0 | 0 | 0 | 194,077 | 1176,144632 | 267 | 2315,999 | 0 | 0 | 1 | 0 | 0 | 0 | 0 | 1 | 0 | 0 | 1 | 0 | 0 | 0 | 0 | 0 | 1 |
| 50.01927 | 20.98497 | TAR_U_1 | taru1_1 | Poland | U | 2 | 1 | 0 | 0 | 0 | 0 | 1 | 214,776 | 1175,480266 | 272 | 49539,861 | 1687 | 1 | 1 | 0 | 0 | 1 | 0 | 0 | 1 | 0 | 0 | 0 | 0 | 0 | 1 | 0 | 0 |
| 50.02098 | 20.98699 | TAR_U_1 | taru1_2 | Poland | U | 0 | 0 | 0 | 0 | 0 | 0 | 1 | 205,605 | 1175,550637 | 272 | 23094,267 | 5951 | 1 | 1 | 1 | 0 | 1 | 0 | 0 | 1 | 1 | 0 | 0 | 1 | 0 | 0 | 0 | 0 |
| 50.02376 | 20.98835 | TAR_U_1 | taru1_3 | Poland | U | 0 | 0 | 0 | 0 | 0 | 0 | 0 | 211 | 1175,746205 | 272 | 5350,523 | 8377 | 0 | 1 | 1 | 0 | 0 | 0 | 0 | 1 | 1 | 0 | 0 | 1 | 1 | 0 | 0 | 1 |
| 50.02692 | 20.98934 | TAR_U_1 | taru1_4 | Poland | U | 1 | 1 | 1 | 1 | 0 | 0 | 1 | 207 | 1175,98982 | 272 | 16333,608 | 500 | 1 | 1 | 0 | 0 | 0 | 0 | 0 | 1 | 1 | 1 | 0 | 1 | 0 | 0 | 0 | 0 |
| 50.03016 | 20.98843 | TAR_U_1 | taru1_5 | Poland | U | 1 | 1 | 1 | 1 | 0 | 0 | 1 | 210,924 | 1176,322065 | 272 | 32253,085 | 500 | 0 | 1 | 0 | 1 | 0 | 0 | 0 | 0 | 1 | 0 | 0 | 1 | 0 | 0 | 0 | 1 |
| 50.00529 | 20.99669 | TAR_U_2 | taru2_1 | Poland | U | 0 | 0 | 2 | 1 | 0 | 0 | 1 | 209,077 | 1173,727126 | 272 | 8172,674 | 9859 | 1 | 1 | 0 | 0 | 0 | 0 | 1 | 1 | 0 | 0 | 0 | 1 | 1 | 0 | 1 | 0 |
| 50.00559 | 21.00094 | TAR_U_2 | taru2_2 | Poland | U | 0 | 0 | 0 | 0 | 0 | 0 | 1 | 210,876 | 1173,577728 | 272 | 13645,97 | 4890 | 1 | 1 | 0 | 0 | 0 | 0 | 1 | 1 | 0 | 0 | 0 | 0 | 0 | 1 | 1 | 0 |
| 50.00453 | 21.00613 | TAR_U_2 | taru2_3 | Poland | U | 1 | 1 | 0 | 0 | 0 | 0 | 1 | 214,519 | 1173,265613 | 272 | 37974,132 | 4203 | 1 | 1 | 1 | 0 | 1 | 0 | 1 | 1 | 0 | 0 | 0 | 0 | 1 | 1 | 1 | 0 |
| 50.00603 | 21.0081 | TAR_U_2 | taru2_4 | Poland | U | 0 | 0 | 0 | 0 | 0 | 0 | 1 | 213,292 | 1173,317841 | 272 | 22269,966 | 12586 | 1 | 0 | 0 | 0 | 1 | 0 | 0 | 0 | 0 | 0 | 0 | 0 | 1 | 1 | 1 | 0 |
| 50.00751 | 21.00739 | TAR_U_2 | taru2_5 | Poland | U | 1 | 1 | 1 | 1 | 0 | 0 | 1 | 209,964 | 1173,481761 | 272 | 30082,895 | 4469 | 1 | 1 | 1 | 0 | 1 | 0 | 0 | 1 | 1 | 0 | 0 | 1 | 1 | 1 | 0 | 0 |
| 49.490025 | 25.605769 | TER_R_1 | terr1_1 | Ukraine | R | 0 | 0 | 0 | 0 | 0 | 0 | 0 | 314,804 | 985,0685263 | 350 | 27235,784 | 2900 | 1 | 1 | 0 | 1 | 0 | 0 | 0 | 0 | 0 | 1 | 0 | 0 | 0 | 0 | 1 | 0 |
| 49.492708 | 25.606678 | TER_R_1 | terr1_2 | Ukraine | R | 1 | 1 | 0 | 0 | 0 | 0 | 0 | 304,968 | 984,8905482 | 350 | 20382,472 | 4476 | 1 | 1 | 0 | 1 | 0 | 0 | 0 | 0 | 0 | 1 | 0 | 0 | 0 | 0 | 1 | 0 |
| 49.495293 | 25.607295 | TER_R_1 | terr1_3 | Ukraine | R | 1 | 1 | 0 | 0 | 0 | 0 | 1 | 301,262 | 984,9731026 | 350 | 43436,152 | 1507 | 1 | 1 | 1 | 1 | 0 | 1 | 0 | 0 | 1 | 1 | 0 | 1 | 0 | 0 | 0 | 0 |
| 49.498071 | 25.607219 | TER_R_1 | terr1_4 | Ukraine | R | 0 | 0 | 0 | 0 | 0 | 0 | 1 | 300,933 | 985,2702313 | 350 | 11330,502 | 5908 | 0 | 1 | 0 | 1 | 0 | 0 | 0 | 0 | 1 | 1 | 1 | 1 | 0 | 0 | 0 | 0 |
| 49.499420 | 25.603545 | TER_R_1 | terr1_5 | Ukraine | R | 0 | 0 | 1 | 1 | 0 | 0 | 1 | 297,088 | 985,5548916 | 350 | 12917,775 | 4565 | 0 | 1 | 0 | 1 | 0 | 0 | 0 | 0 | 1 | 1 | 0 | 1 | 0 | 0 | 0 | 0 |
| 49.602581 | 25.553699 | TER_R_2 | terr2_1 | Ukraine | R | 1 | 1 | 0 | 0 | 0 | 0 | 1 | 305,193 | 972,9956838 | 362 | 20493,482 | 2153 | 1 | 1 | 0 | 1 | 0 | 0 | 0 | 0 | 1 | 1 | 1 | 0 | 0 | 0 | 0 | 0 |
| 49.602093 | 25.557791 | TER_R_2 | terr2_2 | Ukraine | R | 0 | 0 | 0 | 0 | 0 | 0 | 0 | 306,978 | 972,7599824 | 362 | 19243,827 | 4640 | 1 | 1 | 0 | 1 | 0 | 0 | 0 | 0 | 1 | 1 | 1 | 1 | 0 | 0 | 1 | 0 |
| 49.603685 | 25.560401 | TER_R_2 | terr2_3 | Ukraine | R | 1 | 1 | 0 | 0 | 0 | 0 | 0 | 308,153 | 972,4626022 | 362 | 31379,765 | 7755 | 1 | 1 | 0 | 1 | 0 | 0 | 0 | 0 | 1 | 1 | 1 | 0 | 0 | 0 | 1 | 0 |
| 49.606175 | 25.559914 | TER_R_2 | terr2_4 | Ukraine | R | 0 | 0 | 0 | 0 | 0 | 0 | 0 | 315,063 | 972,2056703 | 362 | 35497,338 | 12880 | 1 | 1 | 0 | 1 | 0 | 0 | 0 | 0 | 0 | 1 | 0 | 0 | 0 | 0 | 1 | 0 |
| 49.608926 | 25.560118 | TER_R_2 | terr2_5 | Ukraine | R | 0 | 0 | 0 | 0 | 0 | 0 | 1 | 312,408 | 971,9364135 | 362 | 42173,183 | 8906 | 1 | 1 | 0 | 1 | 0 | 0 | 0 | 0 | 0 | 0 | 1 | 0 | 0 | 0 | 1 | 0 |
| 49.557623 | 25.586040 | TER_U_1 | teru1_1 | Ukraine | U | 2 | 1 | 0 | 0 | 0 | 0 | 1 | 309,63 | 978,6501702 | 355 | 50340,063 | 1117 | 1 | 1 | 0 | 0 | 1 | 0 | 0 | 1 | 0 | 0 | 1 | 0 | 0 | 0 | 0 | 1 |
| 49.554741 | 25.586921 | TER_U_1 | teru1_2 | Ukraine | U | 0 | 0 | 0 | 0 | 0 | 0 | 0 | 306,983 | 978,8457757 | 355 | 30939,234 | 7862 | 1 | 1 | 0 | 0 | 1 | 0 | 0 | 1 | 0 | 0 | 0 | 0 | 1 | 0 | 1 | 0 |
| 49.552184 | 25.586081 | TER_U_1 | teru1_3 | Ukraine | U | 0 | 0 | 0 | 0 | 0 | 0 | 0 | 305,058 | 978,938835 | 355 | 9890,914 | 7794 | 0 | 1 | 0 | 0 | 0 | 0 | 0 | 1 | 0 | 0 | 0 | 0 | 0 | 1 | 0 | 0 |
| 49.550295 | 25.582254 | TER_U_1 | teru1_4 | Ukraine | U | 0 | 0 | 0 | 0 | 0 | 0 | 0 | 306 | 979,1901726 | 355 | 10404,022 | 6494 | 0 | 1 | 0 | 0 | 1 | 0 | 0 | 1 | 0 | 1 | 0 | 0 | 1 | 0 | 0 | 0 |
| 49.548884 | 25.580454 | TER_U_1 | teru1_5 | Ukraine | U | 0 | 0 | 0 | 0 | 0 | 0 | 0 | 305,634 | 979,4999564 | 355 | 34897,734 | 923 | 1 | 1 | 0 | 0 | 1 | 0 | 0 | 1 | 0 | 1 | 0 | 0 | 0 | 0 | 1 | 1 |
| 49.557851 | 25.629836 | TER_U_2 | teru2_1 | Ukraine | U | 0 | 0 | 0 | 0 | 0 | 0 | 0 | 333,419 | 978,7321476 | 355 | 43925,303 | 5189 | 1 | 1 | 0 | 0 | 1 | 0 | 0 | 1 | 0 | 0 | 0 | 0 | 1 | 0 | 0 | 0 |
| 49.556159 | 25.633110 | TER_U_2 | teru2_2 | Ukraine | U | 0 | 0 | 0 | 0 | 0 | 0 | 0 | 350,345 | 978,4495905 | 355 | 45862,649 | 3990 | 1 | 1 | 0 | 0 | 1 | 0 | 0 | 0 | 0 | 0 | 0 | 0 | 0 | 1 | 0 | 1 |
| 49.553742 | 25.634324 | TER_U_2 | teru2_3 | Ukraine | U | 0 | 0 | 0 | 0 | 0 | 0 | 0 | 355,796 | 978,090954 | 355 | 64938,097 | 3058 | 1 | 1 | 0 | 0 | 1 | 0 | 0 | 0 | 0 | 0 | 1 | 0 | 0 | 0 | 1 | 0 |
| 49.551269 | 25.632388 | TER_U_2 | teru2_4 | Ukraine | U | 0 | 0 | 0 | 0 | 0 | 0 | 1 | 340,835 | 977,8694279 | 355 | 42535,108 | 0 | 1 | 1 | 0 | 0 | 1 | 0 | 0 | 0 | 0 | 0 | 0 | 0 | 0 | 0 | 0 | 0 |
| 49.549486 | 25.629116 | TER_U_2 | teru2_5 | Ukraine | U | 0 | 0 | 0 | 0 | 0 | 0 | 1 | 362,856 | 977,7380599 | 355 | 27209,347 | 6357 | 0 | 1 | 0 | 0 | 1 | 0 | 0 | 1 | 0 | 0 | 0 | 0 | 0 | 1 | 1 | 0 |
| 45.65192 | 21.27266 | TIM_R_1 | timr1_1 | Romania | R | 1 | 1 | 1 | 1 | 1 | 1 | 0 | 88 | 811,3878311 | 313 | 19564,386 | 345 | 0 | 1 | 0 | 0 | 0 | 0 | 1 | 0 | 1 | 0 | 1 | 0 | 0 | 0 | 0 | 0 |
| 45.65432 | 21.27476 | TIM_R_1 | timr1_2 | Romania | R | 1 | 1 | 0 | 0 | 0 | 0 | 0 | 88 | 811,4113838 | 313 | 22150,139 | 0 | 0 | 1 | 0 | 0 | 0 | 0 | 1 | 0 | 0 | 1 | 0 | 0 | 0 | 0 | 0 | 0 |
| 45.65714 | 21.27622 | TIM_R_1 | timr1_3 | Romania | R | 2 | 1 | 0 | 0 | 0 | 0 | 0 | 88 | 811,503327 | 313 | 30697,801 | 0 | 0 | 1 | 0 | 0 | 0 | 0 | 1 | 0 | 0 | 1 | 0 | 0 | 0 | 0 | 0 | 0 |
| 45.65945 | 21.27789 | TIM_R_1 | timr1_4 | Romania | R | 1 | 1 | 0 | 0 | 0 | 0 | 0 | 88 | 811,5480953 | 313 | 25567,616 | 0 | 0 | 1 | 0 | 0 | 0 | 0 | 1 | 0 | 0 | 1 | 0 | 0 | 0 | 0 | 0 | 0 |
| 45.66189 | 21.28004 | TIM_R_1 | timr1_5 | Romania | R | 0 | 0 | 0 | 0 | 0 | 0 | 0 | 88 | 811,5704754 | 313 | 13305,09 | 0 | 0 | 1 | 0 | 0 | 0 | 0 | 1 | 0 | 0 | 0 | 0 | 0 | 0 | 0 | 0 | 0 |
| 45.65328 | 21.26449 | TIM_R_2 | timr2_1 | Romania | R | 0 | 0 | 0 | 0 | 1 | 1 | 0 | 88 | 811,9946272 | 313 | 60023,595 | 0 | 1 | 1 | 0 | 1 | 0 | 0 | 0 | 0 | 0 | 0 | 0 | 0 | 0 | 0 | 0 | 0 |
| 45.65588 | 21.26495 | TIM_R_2 | timr2_2 | Romania | R | 0 | 0 | 0 | 0 | 0 | 0 | 0 | 88 | 812,1327675 | 313 | 66885,166 | 0 | 1 | 0 | 0 | 1 | 0 | 0 | 0 | 0 | 0 | 0 | 0 | 0 | 0 | 0 | 0 | 0 |
| 45.6584 | 21.26531 | TIM_R_2 | timr2_3 | Romania | R | 0 | 0 | 0 | 0 | 0 | 0 | 1 | 88 | 812,2737155 | 313 | 38557,381 | 567 | 1 | 1 | 0 | 1 | 0 | 0 | 1 | 0 | 0 | 0 | 1 | 0 | 0 | 0 | 0 | 0 |
| 45.66091 | 21.26302 | TIM_R_2 | timr2_4 | Romania | R | 0 | 0 | 0 | 0 | 0 | 0 | 0 | 88 | 812,5824173 | 313 | 39382,794 | 346 | 1 | 1 | 0 | 1 | 0 | 0 | 1 | 0 | 0 | 0 | 1 | 0 | 0 | 0 | 0 | 0 |
| 45.66201 | 21.25985 | TIM_R_2 | timr2_5 | Romania | R | 0 | 0 | 2 | 1 | 0 | 0 | 0 | 88 | 812,8543967 | 313 | 36259,982 | 0 | 1 | 1 | 0 | 1 | 0 | 0 | 1 | 0 | 0 | 0 | 0 | 0 | 0 | 0 | 0 | 0 |
| 45.7508 | 21.22258 | TIM_U_1 | timu1_1 | Romania | U | 2 | 1 | 0 | 0 | 0 | 0 | 0 | 87 | 821,0028646 | 317 | 38363,244 | 1422 | 1 | 1 | 0 | 0 | 1 | 0 | 0 | 0 | 0 | 0 | 0 | 0 | 0 | 1 | 0 | 0 |
| 45.74974 | 21.22625 | TIM_U_1 | timu1_2 | Romania | U | 1 | 1 | 0 | 0 | 0 | 0 | 0 | 87 | 820,7000873 | 317 | 41505,683 | 6485 | 1 | 1 | 0 | 0 | 1 | 0 | 1 | 0 | 0 | 0 | 0 | 0 | 1 | 0 | 0 | 0 |
| 45.75012 | 21.23014 | TIM_U_1 | timu1_3 | Romania | U | 0 | 0 | 0 | 0 | 0 | 0 | 0 | 87 | 820,4819155 | 317 | 25279,77 | 11975 | 0 | 1 | 0 | 0 | 1 | 0 | 1 | 1 | 0 | 0 | 0 | 1 | 0 | 0 | 0 | 0 |
| 45.75135 | 21.23421 | TIM_U_1 | timu1_4 | Romania | U | 0 | 0 | 3 | 1 | 0 | 0 | 0 | 87 | 820,3062497 | 317 | 24843,766 | 8197 | 0 | 1 | 0 | 0 | 1 | 0 | 1 | 1 | 0 | 0 | 0 | 1 | 0 | 0 | 0 | 0 |
| 45.75294 | 21.23786 | TIM_U_1 | timu1_5 | Romania | U | 0 | 0 | 0 | 0 | 0 | 0 | 0 | 87,411 | 820,183966 | 317 | 43767,235 | 3692 | 0 | 1 | 0 | 0 | 1 | 0 | 0 | 1 | 0 | 0 | 0 | 0 | 1 | 0 | 0 | 0 |
| 45.76006 | 21.22497 | TIM_U_2 | timu2_1 | Romania | U | 0 | 0 | 0 | 0 | 0 | 0 | 0 | 88 | 821,4611081 | 317 | 56709,466 | 3841 | 1 | 1 | 0 | 0 | 1 | 0 | 0 | 0 | 0 | 0 | 0 | 0 | 1 | 0 | 0 | 0 |
| 45.76126 | 21.227 | TIM_U_2 | timu2_2 | Romania | U | 0 | 0 | 0 | 0 | 0 | 0 | 0 | 88 | 821,412743 | 317 | 46749,055 | 6004 | 1 | 1 | 0 | 0 | 1 | 0 | 1 | 1 | 0 | 0 | 0 | 0 | 1 | 0 | 0 | 0 |
| 45.76367 | 21.22678 | TIM_U_2 | timu2_3 | Romania | U | 0 | 0 | 0 | 0 | 0 | 0 | 0 | 89 | 821,5851442 | 317 | 18249,109 | 4395 | 0 | 1 | 0 | 0 | 0 | 0 | 1 | 1 | 0 | 0 | 0 | 0 | 1 | 0 | 0 | 0 |
| 45.76596 | 21.22665 | TIM_U_2 | timu2_4 | Romania | U | 0 | 0 | 0 | 0 | 0 | 0 | 0 | 89 | 821,7439734 | 317 | 2775,082 | 37252 | 0 | 0 | 0 | 0 | 0 | 0 | 0 | 0 | 0 | 0 | 0 | 0 | 0 | 1 | 1 | 0 |
| 45.76839 | 21.2261 | TIM_U_2 | timu2_5 | Romania | U | 0 | 0 | 2 | 1 | 0 | 0 | 0 | 89 | 821,9386912 | 317 | 10103,07 | 10475 | 0 | 1 | 0 | 0 | 0 | 1 | 0 | 1 | 0 | 0 | 1 | 0 | 0 | 1 | 1 | 0 |
| 48.89853 | 17.94221 | TRE_R_1 | trer1_1 | Slovak | R | 0 | 0 | 1 | 1 | 0 | 0 | 1 | 326,948 | 1232,493505 | 199 | 22175,221 | 4453 | 1 | 1 | 0 | 1 | 0 | 0 | 0 | 1 | 1 | 0 | 1 | 1 | 1 | 0 | 1 | 0 |
| 48.89728 | 17.9457 | TRE_R_1 | trer1_2 | Slovak | R | 0 | 0 | 0 | 0 | 0 | 0 | 1 | 297,44 | 1232,76754 | 199 | 9701,103 | 1945 | 0 | 1 | 0 | 1 | 0 | 0 | 1 | 0 | 1 | 0 | 1 | 1 | 0 | 0 | 0 | 0 |
| 48.89778 | 17.9495 | TRE_R_1 | trer1_3 | Slovak | R | 0 | 0 | 0 | 0 | 0 | 0 | 0 | 258,402 | 1232,938775 | 199 | 14325,817 | 129 | 1 | 1 | 0 | 1 | 0 | 0 | 1 | 0 | 1 | 0 | 1 | 0 | 0 | 0 | 0 | 1 |
| 48.89607 | 17.9521 | TRE_R_1 | trer1_4 | Slovak | R | 1 | 1 | 0 | 0 | 0 | 0 | 0 | 237,625 | 1233,223769 | 199 | 10563,001 | 0 | 1 | 1 | 0 | 1 | 0 | 0 | 1 | 0 | 0 | 0 | 0 | 0 | 0 | 0 | 0 | 0 |
| 48.90062 | 17.94022 | TRE_R_1 | trer1_5 | Slovak | R | 2 | 1 | 0 | 0 | 0 | 0 | 0 | 336,768 | 1233,485029 | 199 | 13960,231 | 0 | 1 | 1 | 1 | 1 | 0 | 0 | 1 | 0 | 0 | 0 | 0 | 0 | 0 | 0 | 0 | 1 |
| 48.89212 | 17.9531 | TRE_R_2 | trer2_1 | Slovak | R | 0 | 0 | 0 | 0 | 0 | 0 | 1 | 233,531 | 1232,150476 | 201 | 17496,77 | 5429 | 1 | 1 | 0 | 1 | 0 | 0 | 0 | 1 | 1 | 0 | 1 | 1 | 0 | 0 | 0 | 0 |
| 48.88952 | 17.95533 | TRE_R_2 | trer2_2 | Slovak | R | 0 | 0 | 0 | 0 | 0 | 0 | 1 | 233,54 | 1231,866489 | 201 | 7597,995 | 8553 | 0 | 1 | 0 | 0 | 0 | 0 | 1 | 0 | 1 | 0 | 1 | 1 | 1 | 0 | 0 | 0 |
| 48.8888 | 17.95198 | TRE_R_2 | trer2_3 | Slovak | R | 1 | 1 | 3 | 1 | 1 | 1 | 1 | 255,342 | 1231,98631 | 201 | 12510,246 | 0 | 1 | 1 | 0 | 1 | 0 | 0 | 1 | 0 | 0 | 0 | 0 | 0 | 0 | 0 | 0 | 1 |
| 48.8885 | 17.94765 | TRE_R_2 | trer2_4 | Slovak | R | 1 | 1 | 0 | 0 | 0 | 0 | 0 | 291,272 | 1232,212958 | 201 | 24934,096 | 0 | 1 | 1 | 1 | 1 | 0 | 0 | 0 | 0 | 0 | 0 | 0 | 0 | 0 | 0 | 0 | 0 |
| 48.89009 | 17.94281 | TRE_R_2 | trer2_5 | Slovak | R | 0 | 0 | 0 | 0 | 0 | 0 | 0 | 340,198 | 1232,583552 | 201 | 4477,029 | 0 | 0 | 1 | 0 | 1 | 0 | 0 | 0 | 0 | 0 | 0 | 0 | 0 | 0 | 0 | 0 | 1 |
| 48.90163 | 18.04288 | TRE_U_1 | treu1_1 | Slovak | U | 0 | 0 | 2 | 1 | 0 | 0 | 1 | 212,403 | 1227,846786 | 162 | 8278,215 | 4469 | 0 | 1 | 0 | 1 | 0 | 0 | 0 | 0 | 1 | 1 | 1 | 0 | 0 | 0 | 1 | 0 |
| 48.90299 | 18.04586 | TRE_U_1 | treu1_2 | Slovak | U | 0 | 0 | 2 | 1 | 0 | 0 | 1 | 213,764 | 1227,782624 | 162 | 19048,884 | 4203 | 0 | 1 | 0 | 1 | 0 | 0 | 0 | 0 | 0 | 1 | 1 | 0 | 0 | 0 | 0 | 0 |
| 48.90419 | 18.04992 | TRE_U_1 | treu1_3 | Slovak | U | 0 | 0 | 4 | 1 | 0 | 0 | 1 | 210 | 1227,641148 | 162 | 6939,884 | 1076 | 0 | 1 | 0 | 1 | 0 | 0 | 0 | 0 | 0 | 1 | 1 | 0 | 0 | 0 | 0 | 1 |
| 48.90512 | 18.05359 | TRE_U_1 | treu1_4 | Slovak | U | 1 | 1 | 0 | 0 | 0 | 0 | 1 | 209,634 | 1227,504417 | 162 | 17887,088 | 0 | 1 | 1 | 0 | 1 | 0 | 0 | 0 | 0 | 0 | 1 | 0 | 0 | 0 | 0 | 0 | 1 |
| 48.90312 | 18.05548 | TRE_U_1 | treu1_5 | Slovak | U | 2 | 1 | 0 | 0 | 0 | 0 | 1 | 215,791 | 1227,249433 | 162 | 23878,801 | 5750 | 1 | 1 | 0 | 1 | 0 | 0 | 0 | 0 | 0 | 1 | 1 | 0 | 0 | 0 | 1 | 1 |
| 48.8964 | 18.04623 | TRE_U_2 | treu2_1 | Slovak | U | 0 | 0 | 0 | 0 | 0 | 0 | 0 | 210 | 1227,232792 | 161 | 31690,835 | 6714 | 0 | 1 | 0 | 0 | 1 | 0 | 0 | 1 | 0 | 0 | 1 | 0 | 1 | 0 | 1 | 1 |
| 48.89434 | 18.04341 | TRE_U_2 | treu2_2 | Slovak | U | 0 | 0 | 0 | 0 | 0 | 0 | 0 | 243,212 | 1227,291753 | 161 | 10323,488 | 22702 | 1 | 1 | 0 | 1 | 0 | 0 | 0 | 1 | 0 | 0 | 0 | 0 | 0 | 1 | 0 | 0 |
| 48.89294 | 18.04161 | TRE_U_2 | treu2_3 | Slovak | U | 2 | 1 | 0 | 0 | 0 | 0 | 1 | 247,442 | 1227,296243 | 161 | 48046,326 | 12255 | 1 | 0 | 0 | 1 | 0 | 0 | 0 | 1 | 0 | 0 | 0 | 0 | 0 | 1 | 0 | 1 |
| 48.89157 | 18.04401 | TRE_U_2 | treu2_4 | Slovak | U | 1 | 1 | 0 | 0 | 1 | 1 | 0 | 300,348 | 1227,056641 | 161 | 67207,131 | 1793 | 1 | 1 | 0 | 1 | 1 | 0 | 0 | 0 | 1 | 0 | 0 | 1 | 0 | 0 | 0 | 0 |
| 48.88935 | 18.04184 | TRE_U_2 | treu2_5 | Slovak | U | 2 | 1 | 2 | 1 | 0 | 0 | 1 | 251,584 | 1227,014951 | 161 | 29788,416 | 10586 | 1 | 1 | 0 | 1 | 1 | 1 | 0 | 1 | 1 | 0 | 0 | 1 | 0 | 0 | 0 | 0 |
| 49.23406 | 15.81937 | TRZ_R_1 | trzr1_1 | Czech | R | 3 | 1 | 0 | 0 | 0 | 0 | 0 | 405,451 | 1378,220034 | 40 | 46358,732 | 0 | 1 | 1 | 0 | 1 | 0 | 0 | 1 | 0 | 1 | 1 | 1 | 0 | 0 | 0 | 0 | 0 |
| 49.23161 | 15.82053 | TRZ_R_1 | trzr1_2 | Czech | R | 1 | 1 | 0 | 0 | 0 | 0 | 0 | 405,446 | 1377,988218 | 40 | 53005,6 | 231 | 1 | 1 | 1 | 1 | 0 | 0 | 1 | 0 | 1 | 1 | 1 | 0 | 0 | 0 | 0 | 0 |
| 49.229 | 15.82223 | TRZ_R_1 | trzr1_3 | Czech | R | 2 | 1 | 0 | 0 | 0 | 0 | 0 | 407,83 | 1377,720903 | 40 | 41440,82 | 1195 | 1 | 1 | 1 | 1 | 0 | 0 | 1 | 0 | 0 | 1 | 1 | 0 | 0 | 0 | 0 | 0 |
| 49.22705 | 15.82527 | TRZ_R_1 | trzr1_4 | Czech | R | 0 | 0 | 0 | 0 | 0 | 0 | 0 | 406,98 | 1377,414004 | 40 | 37696,983 | 29 | 1 | 1 | 0 | 1 | 0 | 0 | 1 | 0 | 0 | 1 | 1 | 0 | 0 | 0 | 0 | 0 |
| 49.22688 | 15.82933 | TRZ_R_1 | trzr1_5 | Czech | R | 0 | 0 | 0 | 0 | 0 | 0 | 0 | 409,716 | 1377,163458 | 40 | 14943,279 | 6988 | 1 | 1 | 0 | 1 | 0 | 0 | 1 | 1 | 1 | 1 | 0 | 1 | 0 | 0 | 0 | 0 |
| 49.19782 | 15.83515 | TRZ_R_2 | trzr2_1 | Czech | R | 1 | 1 | 0 | 0 | 0 | 0 | 0 | 445,916 | 1374,924516 | 42 | 15043,151 | 617 | 1 | 1 | 0 | 1 | 0 | 0 | 1 | 0 | 1 | 1 | 0 | 1 | 0 | 0 | 0 | 1 |
| 49.19568 | 15.83286 | TRZ_R_2 | trzr2_2 | Czech | R | 1 | 1 | 0 | 0 | 0 | 0 | 0 | 444,867 | 1374,919989 | 42 | 15932,582 | 1892 | 1 | 1 | 1 | 1 | 0 | 0 | 1 | 0 | 1 | 1 | 0 | 1 | 0 | 0 | 0 | 0 |
| 49.19436 | 15.83601 | TRZ_R_2 | trzr2_3 | Czech | R | 3 | 1 | 0 | 0 | 0 | 0 | 0 | 458,404 | 1374,647909 | 42 | 25117,072 | 465 | 1 | 1 | 0 | 1 | 0 | 0 | 1 | 0 | 0 | 0 | 1 | 0 | 0 | 0 | 0 | 0 |
| 49.19317 | 15.83949 | TRZ_R_2 | trzr2_4 | Czech | R | 0 | 0 | 0 | 0 | 0 | 0 | 0 | 466,882 | 1374,364807 | 42 | 21398,142 | 4643 | 0 | 1 | 0 | 1 | 0 | 0 | 1 | 0 | 1 | 1 | 1 | 1 | 0 | 0 | 0 | 0 |
| 49.19464 | 15.84228 | TRZ_R_2 | trzr2_5 | Czech | R | 0 | 0 | 0 | 0 | 0 | 0 | 0 | 482,885 | 1374,295884 | 42 | 16667,21 | 6217 | 0 | 1 | 1 | 0 | 0 | 0 | 1 | 0 | 1 | 0 | 0 | 1 | 0 | 0 | 0 | 0 |
| 49.21807 | 15.88489 | TRZ_U_1 | trzu1_1 | Czech | U | 1 | 1 | 0 | 0 | 0 | 0 | 0 | 406,294 | 1373,31664 | 44 | 37903,189 | 9909 | 1 | 1 | 1 | 1 | 1 | 0 | 0 | 0 | 0 | 0 | 0 | 1 | 1 | 0 | 0 | 0 |
| 49.21927 | 15.88117 | TRZ_U_1 | trzu1_2 | Czech | U | 1 | 1 | 0 | 0 | 0 | 0 | 0 | 429,379 | 1373,614462 | 44 | 30738,727 | 539 | 0 | 1 | 1 | 0 | 1 | 0 | 1 | 0 | 1 | 0 | 1 | 0 | 0 | 0 | 0 | 0 |
| 49.2195 | 15.87776 | TRZ_U_1 | trzu1_3 | Czech | U | 0 | 0 | 0 | 0 | 0 | 0 | 0 | 439,013 | 1373,82775 | 44 | 13403,479 | 7081 | 0 | 1 | 0 | 0 | 0 | 1 | 1 | 1 | 1 | 0 | 0 | 1 | 1 | 0 | 0 | 0 |
| 49.21906 | 15.87384 | TRZ_U_1 | trzu1_4 | Czech | U | 0 | 0 | 0 | 0 | 0 | 0 | 0 | 435,464 | 1374,030528 | 44 | 9971,946 | 12759 | 0 | 1 | 0 | 0 | 0 | 0 | 0 | 1 | 1 | 0 | 0 | 1 | 0 | 1 | 0 | 0 |
| 49.22048 | 15.87129 | TRZ_U_1 | trzu1_5 | Czech | U | 0 | 0 | 0 | 0 | 0 | 0 | 0 | 448 | 1374,274728 | 44 | 5819,432 | 13548 | 0 | 1 | 0 | 0 | 1 | 0 | 1 | 0 | 1 | 1 | 0 | 1 | 0 | 0 | 0 | 0 |
| 49.20815 | 15.85472 | TRZ_U_2 | trzu2_1 | Czech | U | 1 | 1 | 0 | 0 | 0 | 0 | 0 | 451,68 | 1374,443437 | 44 | 15043,151 | 8985 | 1 | 1 | 0 | 0 | 0 | 0 | 1 | 1 | 1 | 0 | 0 | 1 | 1 | 0 | 0 | 0 |
| 49.20858 | 15.85855 | TRZ_U_2 | trzu2_2 | Czech | U | 0 | 0 | 0 | 0 | 0 | 0 | 0 | 454,577 | 1374,245989 | 44 | 15932,582 | 10773 | 1 | 1 | 1 | 1 | 0 | 0 | 1 | 1 | 1 | 0 | 0 | 1 | 1 | 1 | 0 | 0 |
| 49.20873 | 15.86259 | TRZ_U_2 | trzu2_3 | Czech | U | 0 | 0 | 0 | 0 | 0 | 0 | 0 | 449,099 | 1374,017224 | 44 | 25117,072 | 9023 | 1 | 1 | 1 | 1 | 0 | 0 | 1 | 1 | 0 | 0 | 0 | 0 | 1 | 0 | 1 | 0 |
| 49.20896 | 15.86679 | TRZ_U_2 | trzu2_4 | Czech | U | 2 | 1 | 0 | 0 | 0 | 0 | 0 | 450,324 | 1373,78477 | 44 | 21398,142 | 9560 | 1 | 1 | 1 | 1 | 0 | 0 | 1 | 1 | 0 | 0 | 0 | 0 | 1 | 0 | 0 | 1 |
| 49.21148 | 15.86791 | TRZ_U_2 | trzu2_5 | Czech | U | 0 | 0 | 0 | 0 | 0 | 0 | 0 | 446,344 | 1373,88336 | 44 | 16667,21 | 12068 | 1 | 1 | 1 | 1 | 0 | 0 | 1 | 0 | 1 | 0 | 0 | 1 | 0 | 0 | 1 | 0 |
| 48.3494 | 16.04684 | TUL_R_1 | tulr1_1 | Austria | R | 3 | 1 | 0 | 0 | 0 | 0 | 0 | 176,06 | 1308,624348 | 21 | 21944,586 | 0 | 1 | 1 | 1 | 1 | 0 | 0 | 0 | 0 | 1 | 0 | 0 | 0 | 0 | 0 | 0 | 0 |
| 48.34964 | 16.05238 | TUL_R_1 | tulr1_2 | Austria | R | 0 | 0 | 1 | 1 | 0 | 0 | 0 | 175,432 | 1308,294919 | 21 | 18263,07 | 0 | 1 | 1 | 1 | 1 | 0 | 0 | 0 | 0 | 0 | 0 | 0 | 0 | 0 | 0 | 0 | 0 |
| 48.35219 | 16.05389 | TUL_R_1 | tulr1_3 | Austria | R | 1 | 1 | 0 | 0 | 0 | 0 | 0 | 176,884 | 1308,354389 | 21 | 44439,199 | 0 | 1 | 1 | 1 | 1 | 0 | 0 | 0 | 0 | 0 | 0 | 0 | 0 | 0 | 0 | 0 | 1 |
| 48.35489 | 16.05438 | TUL_R_1 | tulr1_4 | Austria | R | 0 | 0 | 0 | 0 | 0 | 0 | 1 | 177,768 | 1308,497707 | 21 | 35861,384 | 1496 | 1 | 1 | 1 | 1 | 0 | 0 | 0 | 1 | 0 | 0 | 1 | 0 | 0 | 0 | 0 | 1 |
| 48.35791 | 16.05511 | TUL_R_1 | tulr1_5 | Austria | R | 0 | 0 | 0 | 0 | 0 | 0 | 0 | 176,584 | 1308,628203 | 21 | 19693,082 | 0 | 1 | 1 | 0 | 1 | 0 | 0 | 0 | 0 | 0 | 0 | 0 | 0 | 0 | 0 | 0 | 0 |
| 48.39884 | 16.09531 | TUL_R_2 | tulr2_1 | Austria | R | 1 | 1 | 0 | 0 | 0 | 0 | 1 | 193,467 | 1308,646999 | 26 | 11576,494 | 207 | 0 | 1 | 0 | 1 | 0 | 0 | 0 | 0 | 0 | 0 | 1 | 0 | 0 | 0 | 0 | 1 |

| | | | | | | | | | | | | | | | | | | | | | | | | | | | | | | | | | |
|---|---|---|---|---|---|---|---|---|---|---|---|---|---|---|---|---|---|---|---|---|---|---|---|---|---|---|---|---|---|---|---|---|---|
| 48.39661 | 16.09479 | TUL_R_2 | tulr2_2 | Austria | R | 0 | 0 | 1 | 1 | 0 | 0 | 1 | 203,736 | 1308,546875 | 26 | 11014,64 | 3488 | 1 | 1 | 0 | 1 | 0 | 1 | 0 | 0 | 0 | 0 | 0 | 0 | 1 | 0 | 0 | 0 |
| 48.39652 | 16.0901 | TUL_R_2 | tulr2_3 | Austria | R | 0 | 0 | 1 | 1 | 0 | 0 | 1 | 178,888 | 1308,924187 | 26 | 17907,54 | 0 | 1 | 0 | 0 | 1 | 0 | 0 | 0 | 0 | 0 | 0 | 0 | 0 | 0 | 0 | 0 | 0 |
| 48.3971 | 16.08473 | TUL_R_2 | tulr2_4 | Austria | R | 0 | 0 | 1 | 1 | 0 | 0 | 0 | 180,828 | 1309,218481 | 26 | 13261,432 | 0 | 1 | 1 | 0 | 1 | 0 | 0 | 0 | 0 | 0 | 0 | 0 | 0 | 0 | 0 | 0 | 0 |
| 48.39732 | 16.08003 | TUL_R_2 | tulr2_5 | Austria | R | 0 | 0 | 0 | 0 | 0 | 0 | 0 | 180,112 | 1309,485503 | 26 | 15731,368 | 0 | 1 | 1 | 0 | 1 | 0 | 0 | 0 | 0 | 0 | 0 | 0 | 0 | 0 | 0 | 0 | 0 |
| 48.32965 | 16.06932 | TUL_U_1 | tulu1_1 | Austria | U | 0 | 0 | 0 | 0 | 0 | 0 | 1 | 177,856 | 1306,024608 | 19 | 15228,954 | 16090 | 1 | 0 | 0 | 1 | 0 | 0 | 0 | 0 | 1 | 0 | 0 | 1 | 0 | 1 | 0 | 0 |
| 48.32986 | 16.07344 | TUL_U_1 | tulu1_2 | Austria | U | 0 | 0 | 1 | 1 | 0 | 0 | 1 | 175,31 | 1305,768463 | 19 | 34535,562 | 9798 | 1 | 1 | 0 | 1 | 0 | 0 | 0 | 1 | 1 | 0 | 0 | 1 | 0 | 1 | 0 | 1 |
| 48.33201 | 16.07351 | TUL_U_1 | tulu1_3 | Austria | U | 0 | 0 | 0 | 0 | 0 | 0 | 0 | 175 | 1305,909663 | 19 | 19909,106 | 1293 | 1 | 1 | 0 | 1 | 1 | 0 | 0 | 0 | 0 | 1 | 1 | 0 | 0 | 0 | 0 | 0 |
| 48.33313 | 16.06991 | TUL_U_1 | tulu1_4 | Austria | U | 2 | 1 | 0 | 0 | 0 | 0 | 1 | 176,536 | 1306,202369 | 19 | 20003,484 | 2300 | 0 | 1 | 0 | 0 | 1 | 0 | 0 | 0 | 0 | 0 | 1 | 0 | 0 | 0 | 0 | 0 |
| 48.33295 | 16.06658 | TUL_U_1 | tulu1_5 | Austria | U | 0 | 0 | 0 | 0 | 0 | 0 | 0 | 176,76 | 1306,39768 | 19 | 4192,083 | 11850 | 0 | 1 | 0 | 0 | 0 | 0 | 1 | 1 | 0 | 0 | 0 | 0 | 0 | 1 | 0 | 0 |
| 48.33042 | 16.04926 | TUL_U_2 | tulu2_1 | Austria | U | 0 | 0 | 0 | 0 | 0 | 0 | 1 | 179,996 | 1307,318522 | 18 | 9383,939 | 27438 | 0 | 1 | 0 | 0 | 0 | 0 | 0 | 1 | 1 | 0 | 0 | 1 | 0 | 1 | 0 | 0 |
| 48.33257 | 16.04735 | TUL_U_2 | tulu2_2 | Austria | U | 2 | 1 | 0 | 0 | 0 | 0 | 1 | 174,388 | 1307,570943 | 18 | 17458,507 | 9842 | 0 | 1 | 0 | 1 | 1 | 0 | 0 | 1 | 0 | 1 | 0 | 0 | 1 | 1 | 0 | 0 |
| 48.33183 | 16.0435 | TUL_U_2 | tulu2_3 | Austria | U | 3 | 1 | 0 | 0 | 0 | 0 | 1 | 175,753 | 1307,763875 | 18 | 43583,919 | 2157 | 1 | 0 | 0 | 1 | 0 | 0 | 0 | 0 | 1 | 1 | 0 | 1 | 0 | 0 | 0 | 1 |
| 48.32988 | 16.04199 | TUL_U_2 | tulu2_4 | Austria | U | 1 | 1 | 0 | 0 | 1 | 1 | 1 | 179 | 1307,738302 | 18 | 21558,636 | 7891 | 1 | 1 | 0 | 0 | 0 | 0 | 1 | 0 | 1 | 1 | 0 | 1 | 0 | 0 | 0 | 1 |
| 48.32777 | 16.04359 | TUL_U_2 | tulu2_5 | Austria | U | 0 | 0 | 0 | 0 | 0 | 0 | 0 | 178,148 | 1307,511209 | 18 | 10274,461 | 7689 | 0 | 1 | 0 | 0 | 0 | 0 | 1 | 0 | 1 | 0 | 0 | 1 | 0 | 0 | 0 | 0 |
| 43.7425 | 24.82151 | TUR_R_1 | turr1_1 | Romania | R | 0 | 0 | 0 | 0 | 0 | 0 | 0 | 28,564 | 459,1057846 | 479 | 23682,875 | 0 | 0 | 1 | 0 | 0 | 0 | 0 | 1 | 0 | 1 | 0 | 0 | 0 | 0 | 0 | 0 | 0 |
| 43.74377 | 24.82528 | TUR_R_1 | turr1_2 | Romania | R | 0 | 0 | 1 | 1 | 0 | 0 | 0 | 27 | 458,956147 | 479 | 15306,566 | 0 | 1 | 1 | 0 | 1 | 0 | 0 | 1 | 0 | 0 | 0 | 0 | 0 | 0 | 0 | 0 | 0 |
| 43.74552 | 24.82831 | TUR_R_1 | turr1_3 | Romania | R | 0 | 0 | 0 | 0 | 0 | 0 | 0 | 26,956 | 458,8920606 | 479 | 18608,304 | 0 | 1 | 1 | 0 | 1 | 0 | 0 | 1 | 0 | 0 | 1 | 0 | 0 | 0 | 0 | 0 | 0 |
| 43.74679 | 24.83211 | TUR_R_1 | turr1_4 | Romania | R | 0 | 0 | 1 | 1 | 0 | 0 | 1 | 25,377 | 458,7262447 | 479 | 4408,322 | 0 | 0 | 1 | 0 | 0 | 0 | 0 | 1 | 0 | 0 | 0 | 0 | 0 | 0 | 0 | 0 | 1 |
| 43.7471 | 24.8358 | TUR_R_1 | turr1_5 | Romania | R | 0 | 0 | 0 | 0 | 0 | 0 | 0 | 27,12 | 458,5129378 | 479 | 2188,837 | 0 | 0 | 1 | 0 | 0 | 0 | 0 | 1 | 1 | 0 | 0 | 1 | 0 | 0 | 0 | 0 | 0 |
| 43.74783 | 24.86961 | TUR_U_1 | turu1_1 | Romania | U | 0 | 0 | 0 | 0 | 0 | 0 | 0 | 40,356 | 456,4403706 | 481 | 18784,336 | 16325 | 0 | 1 | 0 | 0 | 1 | 0 | 1 | 1 | 0 | 0 | 0 | 0 | 1 | 0 | 0 | 0 |
| 43.74892 | 24.87249 | TUR_U_1 | turu1_2 | Romania | U | 0 | 0 | 0 | 0 | 0 | 0 | 0 | 34,812 | 456,3402478 | 481 | 16225,053 | 23572 | 0 | 1 | 0 | 0 | 0 | 0 | 1 | 1 | 0 | 0 | 0 | 1 | 1 | 1 | 0 | 0 |
| 43.74636 | 24.87397 | TUR_U_1 | turu1_3 | Romania | U | 0 | 0 | 0 | 0 | 0 | 0 | 1 | 34,366 | 456,0685571 | 481 | 3411,711 | 42248 | 0 | 1 | 0 | 0 | 0 | 0 | 0 | 0 | 1 | 0 | 0 | 1 | 0 | 0 | 0 | 0 |
| 43.74471 | 24.87219 | TUR_U_1 | turu1_4 | Romania | U | 0 | 0 | 2 | 1 | 0 | 0 | 1 | 32,956 | 456,0672096 | 481 | 5779,492 | 43723 | 0 | 1 | 0 | 0 | 0 | 0 | 1 | 1 | 1 | 0 | 0 | 1 | 1 | 0 | 1 | 0 |
| 43.74403 | 24.86861 | TUR_U_1 | turu1_5 | Romania | U | 0 | 0 | 0 | 0 | 0 | 0 | 0 | 38,512 | 456,2432625 | 481 | 22392,738 | 32663 | 0 | 1 | 0 | 0 | 0 | 0 | 1 | 1 | 1 | 0 | 0 | 1 | 1 | 0 | 1 | 0 |
| 52.11662 | 20.54234 | WAR_R_1 | warr1_1 | Poland | R | 1 | 1 | 0 | 0 | 0 | 0 | 0 | 97 | 1387,346265 | 43 | 49163 | 0 | 1 | 1 | 1 | 1 | 0 | 0 | 0 | 0 | 0 | 0 | 0 | 0 | 0 | 0 | 0 | 0 |
| 52.11939 | 20.54152 | WAR_R_1 | warr1_2 | Poland | R | 0 | 0 | 0 | 0 | 0 | 0 | 0 | 96 | 1387,639259 | 43 | 52996 | 0 | 1 | 0 | 1 | 1 | 0 | 0 | 0 | 0 | 0 | 0 | 0 | 0 | 0 | 0 | 0 | 0 |
| 52.12002 | 20.54603 | WAR_R_1 | warr1_3 | Poland | R | 1 | 1 | 0 | 0 | 0 | 0 | 0 | 97 | 1387,534893 | 43 | 41384 | 0 | 1 | 0 | 1 | 1 | 0 | 0 | 0 | 0 | 0 | 0 | 0 | 0 | 0 | 0 | 0 | 0 |
| 52.11894 | 20.54978 | WAR_R_1 | warr1_4 | Poland | R | 1 | 1 | 0 | 0 | 0 | 0 | 0 | 97 | 1387,293816 | 43 | 47969 | 0 | 1 | 0 | 1 | 1 | 0 | 0 | 0 | 0 | 0 | 0 | 0 | 0 | 0 | 0 | 0 | 0 |
| 52.11852 | 20.55426 | WAR_R_1 | warr1_5 | Poland | R | 0 | 0 | 0 | 0 | 0 | 0 | 0 | 97 | 1387,100339 | 43 | 52949 | 0 | 1 | 0 | 1 | 1 | 0 | 0 | 0 | 0 | 0 | 1 | 0 | 0 | 0 | 0 | 0 | 0 |
| 52.0603 | 20.65543 | WAR_R_2 | warr2_1 | Poland | R | 0 | 0 | 0 | 0 | 0 | 0 | 0 | 138 | 1377,959759 | 51 | 27759,202 | 1649 | 1 | 0 | 1 | 1 | 0 | 0 | 0 | 0 | 1 | 0 | 1 | 0 | 0 | 0 | 0 | 0 |
| 52.05844 | 20.6579 | WAR_R_2 | warr2_2 | Poland | R | 0 | 0 | 0 | 0 | 0 | 0 | 1 | 140 | 1377,69651 | 51 | 31997 | 2213 | 1 | 0 | 1 | 1 | 0 | 0 | 0 | 0 | 0 | 0 | 1 | 0 | 0 | 0 | 0 | 0 |
| 52.0566 | 20.66149 | WAR_R_2 | warr2_3 | Poland | R | 0 | 0 | 0 | 0 | 0 | 0 | 0 | 140,847 | 1377,394039 | 51 | 18297 | 1716 | 1 | 0 | 1 | 1 | 0 | 0 | 0 | 0 | 1 | 0 | 1 | 0 | 0 | 0 | 0 | 1 |
| 52.05439 | 20.6626 | WAR_R_2 | warr2_4 | Poland | R | 0 | 0 | 0 | 0 | 0 | 0 | 0 | 138 | 1377,143162 | 51 | 11630 | 704 | 1 | 1 | 1 | 1 | 0 | 0 | 1 | 0 | 1 | 0 | 1 | 0 | 0 | 0 | 0 | 0 |
| 52.05245 | 20.66028 | WAR_R_2 | warr2_5 | Poland | R | 2 | 1 | 0 | 0 | 0 | 0 | 0 | 135,18 | 1377,041325 | 51 | 29501 | 989 | 1 | 1 | 1 | 1 | 0 | 0 | 1 | 0 | 1 | 1 | 1 | 0 | 0 | 0 | 0 | 0 |
| 52.1278 | 20.71107 | WAR_R_3 | warr3_1 | Poland | R | 1 | 1 | 0 | 0 | 0 | 0 | 0 | 99 | 1382,43848 | 44 | 51625 | 3718 | 1 | 0 | 1 | 1 | 0 | 0 | 0 | 0 | 0 | 0 | 0 | 1 | 0 | 0 | 0 | 0 |
| 52.12735 | 20.71549 | WAR_R_3 | warr3_2 | Poland | R | 1 | 1 | 0 | 0 | 0 | 0 | 0 | 98 | 1382,242539 | 44 | 47015 | 3282 | 1 | 0 | 1 | 1 | 0 | 0 | 0 | 0 | 1 | 0 | 0 | 1 | 0 | 0 | 0 | 0 |
| 52.12758 | 20.71957 | WAR_R_3 | warr3_3 | Poland | R | 0 | 0 | 0 | 0 | 0 | 0 | 0 | 98,26 | 1382,114326 | 44 | 41022 | 3322 | 1 | 1 | 1 | 1 | 0 | 0 | 0 | 1 | 0 | 0 | 0 | 1 | 0 | 0 | 0 | 0 |
| 52.12532 | 20.72124 | WAR_R_3 | warr3_4 | Poland | R | 2 | 1 | 0 | 0 | 0 | 0 | 0 | 101,848 | 1381,83303 | 44 | 36188 | 7439 | 0 | 1 | 1 | 1 | 0 | 0 | 0 | 0 | 0 | 0 | 0 | 1 | 0 | 0 | 0 | 0 |
| 52.12295 | 20.7194 | WAR_R_3 | warr3_5 | Poland | R | 2 | 1 | 0 | 0 | 0 | 0 | 0 | 103 | 1381,680003 | 44 | 51137 | 3965 | 1 | 0 | 1 | 1 | 0 | 0 | 0 | 0 | 0 | 0 | 1 | 0 | 0 | 0 | 0 | 0 |
| 52.1507 | 20.91538 | WAR_R_4 | warr4_1 | Poland | R | 0 | 0 | 2 | 1 | 0 | 0 | 1 | 103 | 1377,534193 | 43 | 18121,917 | 6593 | 0 | 1 | 0 | 0 | 0 | 0 | 1 | 1 | 0 | 0 | 0 | 1 | 0 | 1 | 0 | 0 |
| 52.14793 | 20.91563 | WAR_R_4 | warr4_2 | Poland | R | 0 | 0 | 0 | 0 | 0 | 0 | 0 | 102,331 | 1377,256726 | 43 | 9794,397 | 0 | 0 | 1 | 0 | 0 | 0 | 0 | 0 | 0 | 0 | 1 | 0 | 0 | 0 | 0 | 0 | 0 |
| 52.14522 | 20.91696 | WAR_R_4 | warr4_3 | Poland | R | 2 | 1 | 0 | 0 | 0 | 0 | 0 | 104 | 1376,950518 | 43 | 39999,231 | 0 | 1 | 1 | 0 | 1 | 0 | 0 | 0 | 0 | 0 | 1 | 0 | 0 | 0 | 0 | 0 | 1 |
| 52.14327 | 20.92057 | WAR_R_4 | warr4_4 | Poland | R | 1 | 1 | 0 | 0 | 0 | 0 | 0 | 103,988 | 1376,638631 | 43 | 18456,327 | 0 | 0 | 1 | 0 | 1 | 0 | 0 | 0 | 0 | 0 | 1 | 0 | 0 | 0 | 0 | 0 | 0 |
| 52.14146 | 20.92276 | WAR_R_4 | warr4_5 | Poland | R | 0 | 0 | 0 | 0 | 0 | 0 | 0 | 104 | 1376,389438 | 43 | 44543,883 | 2791 | 1 | 1 | 0 | 1 | 0 | 0 | 1 | 0 | 0 | 1 | 1 | 0 | 0 | 0 | 0 | 0 |
| 52.02427 | 20.78944 | WAR_R_9 | warr9_1 | Poland | R | 0 | 0 | 0 | 0 | 0 | 0 | 0 | 140,628 | 1369,789307 | 56 | 19879,329 | 1192 | 1 | 1 | 1 | 1 | 0 | 0 | 1 | 1 | 1 | 0 | 1 | 0 | 0 | 0 | 0 | 0 |
| 52.0228 | 20.78571 | WAR_R_9 | warr9_2 | Poland | R | 0 | 0 | 0 | 0 | 0 | 0 | 0 | 142,92 | 1369,7796 | 56 | 11922,074 | 3031 | 0 | 1 | 1 | 0 | 0 | 0 | 0 | 1 | 1 | 0 | 0 | 1 | 1 | 0 | 0 | 0 |
| 52.02157 | 20.78184 | WAR_R_9 | warr9_3 | Poland | R | 1 | 1 | 0 | 0 | 0 | 0 | 0 | 143,348 | 1369,798836 | 56 | 19452,875 | 2237 | 0 | 1 | 1 | 1 | 0 | 0 | 0 | 1 | 1 | 1 | 1 | 1 | 0 | 0 | 0 | 0 |
| 52.02043 | 20.77785 | WAR_R_9 | warr9_4 | Poland | R | 0 | 0 | 0 | 0 | 0 | 0 | 0 | 145,164 | 1369,83057 | 56 | 34508,585 | 1822 | 1 | 1 | 1 | 1 | 0 | 0 | 0 | 1 | 1 | 1 | 1 | 1 | 0 | 0 | 0 | 0 |
| 52.01842 | 20.77542 | WAR_R_9 | warr9_5 | Poland | R | 0 | 0 | 0 | 0 | 0 | 0 | 0 | 147,2 | 1369,724786 | 56 | 40084,689 | 2090 | 1 | 1 | 1 | 1 | 0 | 0 | 0 | 1 | 1 | 1 | 1 | 1 | 0 | 0 | 0 | 0 |
| 43.10101 | 25.62934 | WIE_R_1 | wier1_1 | Bulgaria | R | 0 | 0 | 0 | 0 | 0 | 0 | 0 | 164,419 | 363,8321492 | 529 | 59038,037 | 0 | 1 | 0 | 0 | 1 | 0 | 0 | 0 | 0 | 0 | 1 | 0 | 0 | 0 | 0 | 0 | 0 |
| 43.10016 | 25.63303 | WIE_R_1 | wier1_2 | Bulgaria | R | 0 | 0 | 0 | 0 | 0 | 0 | 0 | 155,576 | 363,5373559 | 529 | 62078,184 | 0 | 1 | 0 | 0 | 1 | 0 | 0 | 0 | 0 | 0 | 1 | 0 | 0 | 0 | 0 | 0 | 0 |
| 43.09852 | 25.63607 | WIE_R_1 | wier1_3 | Bulgaria | R | 0 | 0 | 0 | 0 | 0 | 0 | 0 | 156,867 | 363,2277898 | 529 | 58372,601 | 190 | 1 | 0 | 0 | 1 | 0 | 0 | 0 | 0 | 0 | 1 | 1 | 0 | 0 | 0 | 0 | 1 |
| 43.09626 | 25.63735 | WIE_R_1 | wier1_4 | Bulgaria | R | 0 | 0 | 0 | 0 | 0 | 0 | 0 | 140,452 | 362,9931895 | 529 | 48899,449 | 4783 | 1 | 1 | 0 | 1 | 0 | 0 | 0 | 1 | 1 | 1 | 0 | 1 | 0 | 0 | 0 | 0 |
| 43.09423 | 25.63977 | WIE_R_1 | wier1_5 | Bulgaria | R | 0 | 0 | 0 | 0 | 0 | 0 | 0 | 144,034 | 362,7049633 | 529 | 29951,898 | 15603 | 1 | 1 | 0 | 1 | 0 | 0 | 0 | 1 | 1 | 1 | 0 | 1 | 0 | 0 | 1 | 0 |
| 43.10502 | 25.66325 | WIE_R_2 | wier2_1 | Bulgaria | R | 0 | 0 | 0 | 0 | 0 | 0 | 0 | 360,657 | 361,8873049 | 529 | 70537,921 | 0 | 1 | 1 | 0 | 1 | 0 | 0 | 0 | 0 | 0 | 0 | 0 | 0 | 0 | 0 | 0 | 0 |
| 43.10556 | 25.6596 | WIE_R_2 | wier2_2 | Bulgaria | R | 1 | 1 | 0 | 0 | 0 | 0 | 0 | 365,321 | 362,1578938 | 529 | 69737,146 | 0 | 1 | 1 | 0 | 1 | 0 | 0 | 0 | 0 | 0 | 0 | 0 | 0 | 0 | 0 | 0 | 0 |
| 43.10582 | 25.65596 | WIE_R_2 | wier2_3 | Bulgaria | R | 0 | 0 | 0 | 0 | 0 | 0 | 0 | 372,978 | 362,4101549 | 529 | 64703,264 | 70 | 1 | 1 | 0 | 1 | 0 | 0 | 0 | 0 | 0 | 0 | 0 | 0 | 0 | 0 | 0 | 0 |
| 43.10455 | 25.65335 | WIE_R_2 | wier2_4 | Bulgaria | R | 0 | 0 | 0 | 0 | 0 | 0 | 0 | 380,997 | 362,5000323 | 529 | 63303,351 | 176 | 1 | 1 | 0 | 1 | 0 | 0 | 0 | 0 | 0 | 0 | 0 | 0 | 0 | 0 | 0 | 0 |
| 43.1028 | 25.65124 | WIE_R_2 | wier2_5 | Bulgaria | R | 0 | 0 | 1 | 1 | 0 | 0 | 0 | 383,602 | 362,5210443 | 529 | 35679,317 | 0 | 1 | 1 | 0 | 1 | 0 | 0 | 0 | 0 | 0 | 0 | 0 | 0 | 0 | 0 | 0 | 0 |
| 43.07556 | 25.62659 | WIE_U_1 | wieu1_1 | Bulgaria | U | 0 | 0 | 1 | 1 | 0 | 0 | 0 | 188,564 | 362,3396871 | 529 | 15875,291 | 20552 | 0 | 1 | 0 | 0 | 1 | 0 | 1 | 1 | 0 | 0 | 0 | 1 | 1 | 0 | 0 | 0 |
| 43.07444 | 25.62957 | WIE_U_1 | wieu1_2 | Bulgaria | U | 0 | 0 | 0 | 0 | 0 | 0 | 0 | 185 | 362,0705545 | 529 | 27766,154 | 15918 | 1 | 1 | 0 | 1 | 1 | 0 | 0 | 0 | 0 | 0 | 0 | 1 | 0 | 0 | 0 | 0 |
| 43.07648 | 25.63151 | WIE_U_1 | wieu1_3 | Bulgaria | U | 0 | 0 | 0 | 0 | 0 | 0 | 0 | 171,513 | 362,0746133 | 529 | 12790,223 | 25023 | 0 | 1 | 0 | 0 | 0 | 0 | 0 | 1 | 0 | 0 | 0 | 0 | 1 | 1 | 0 | 0 |
| 43.07864 | 25.63112 | WIE_U_1 | wieu1_4 | Bulgaria | U | 0 | 0 | 0 | 0 | 0 | 0 | 0 | 178,104 | 362,2436055 | 529 | 21923,079 | 21244 | 0 | 1 | 0 | 0 | 1 | 0 | 1 | 1 | 0 | 0 | 0 | 0 | 1 | 1 | 0 | 0 |
| 43.08029 | 25.63104 | WIE_U_1 | wieu1_5 | Bulgaria | U | 0 | 0 | 0 | 0 | 0 | 0 | 0 | 192,921 | 362,3562359 | 529 | 21074,241 | 23674 | 0 | 1 | 0 | 0 | 1 | 0 | 0 | 1 | 0 | 0 | 0 | 0 | 0 | 1 | 0 | 0 |
| 43.07321 | 25.64247 | WIE_U_2 | wieu2_1 | Bulgaria | U | 2 | 1 | 0 | 0 | 0 | 0 | 0 | 171,952 | 361,1431875 | 529 | 57354,765 | 2799 | 1 | 0 | 1 | 1 | 0 | 0 | 0 | 0 | 0 | 0 | 0 | 0 | 1 | 0 | 1 | 0 |
| 43.07452 | 25.6395 | WIE_U_2 | wieu2_2 | Bulgaria | U | 2 | 1 | 0 | 0 | 0 | 0 | 0 | 155,141 | 361,4220253 | 529 | 49843,486 | 2876 | 1 | 1 | 1 | 1 | 0 | 0 | 1 | 1 | 0 | 1 | 0 | 0 | 1 | 0 | 1 | 0 |
| 43.07727 | 25.64017 | WIE_U_2 | wieu2_3 | Bulgaria | U | 0 | 0 | 0 | 0 | 0 | 0 | 0 | 164,015 | 361,5594839 | 529 | 59786,735 | 0 | 1 | 0 | 1 | 1 | 1 | 0 | 1 | 0 | 0 | 0 | 0 | 0 | 0 | 0 | 0 | 1 |
| 43.08048 | 25.63923 | WIE_U_2 | wieu2_4 | Bulgaria | U | 2 | 1 | 0 | 0 | 0 | 0 | 0 | 164,062 | 361,8302045 | 529 | 57746,077 | 400 | 1 | 1 | 1 | 1 | 1 | 0 | 1 | 1 | 0 | 1 | 0 | 0 | 1 | 0 | 0 | 0 |
| 43.07824 | 25.64173 | WIE_U_2 | wieu2_5 | Bulgaria | U | 0 | 0 | 0 | 0 | 0 | 0 | 0 | 198,776 | 361,5224275 | 529 | 49662,408 | 200 | 1 | 0 | 1 | 1 | 1 | 0 | 0 | 1 | 0 | 1 | 0 | 0 | 1 | 0 | 0 | 0 |
| 52.65196 | 19.15226 | WLO_R_1 | wlor1_1 | Poland | R | 0 | 0 | 0 | 0 | 0 | 0 | 1 | 58,944 | 1488,762009 | -27 | 12979,435 | 111 | 1 | 1 | 0 | 1 | 0 | 0 | 0 | 0 | 0 | 1 | 0 | 0 | 0 | 0 | 0 | 1 |
| 52.65035 | 19.15408 | WLO_R_1 | wlor1_2 | Poland | R | 0 | 0 | 0 | 0 | 0 | 0 | 1 | 54 | 1488,545254 | -27 | 9512,935 | 32 | 1 | 1 | 0 | 1 | 0 | 0 | 0 | 1 | 0 | 0 | 0 | 0 | 0 | 0 | 0 | 0 |
| 52.64922 | 19.15644 | WLO_R_1 | wlor1_3 | Poland | R | 0 | 0 | 0 | 0 | 0 | 0 | 0 | 54,808 | 1488,352847 | -27 | 30240,606 | 2032 | 1 | 1 | 0 | 1 | 0 | 0 | 0 | 0 | 0 | 0 | 0 | 1 | 0 | 0 | 0 | 1 |
| 52.64887 | 19.16083 | WLO_R_1 | wlor1_4 | Poland | R | 0 | 0 | 0 | 0 | 0 | 0 | 1 | 55 | 1488,151989 | -27 | 18199,637 | 2543 | 1 | 1 | 0 | 1 | 0 | 0 | 0 | 1 | 0 | 0 | 1 | 1 | 0 | 0 | 0 | 0 |
| 52.6464 | 19.16266 | WLO_R_1 | wlor1_5 | Poland | R | 0 | 0 | 0 | 0 | 0 | 0 | 1 | 56 | 1487,850236 | -27 | 11346,654 | 617 | 1 | 1 | 0 | 1 | 0 | 0 | 0 | 1 | 0 | 0 | 1 | 0 | 0 | 0 | 0 | 1 |
| 52.65305 | 19.12642 | WLO_R_2 | wlor2_1 | Poland | R | 0 | 0 | 0 | 0 | 0 | 0 | 1 | 56 | 1489,846975 | -28 | 7411,051 | 8119 | 0 | 1 | 0 | 0 | 0 | 0 | 0 | 1 | 0 | 0 | 0 | 0 | 1 | 0 | 0 | 0 |
| 52.65425 | 19.12292 | WLO_R_2 | wlor2_2 | Poland | R | 0 | 0 | 0 | 0 | 0 | 0 | 1 | 56 | 1490,091363 | -28 | 9147,801 | 6183 | 0 | 1 | 0 | 0 | 0 | 0 | 0 | 1 | 1 | 0 | 0 | 1 | 1 | 0 | 0 | 0 |
| 52.65378 | 19.11925 | WLO_R_2 | wlor2_3 | Poland | R | 0 | 0 | 0 | 0 | 0 | 0 | 1 | 52,458 | 1490,188383 | -28 | 17378,482 | 1858 | 0 | 1 | 0 | 0 | 0 | 0 | 1 | 1 | 1 | 0 | 1 | 1 | 0 | 0 | 1 | 1 |
| 52.65588 | 19.11663 | WLO_R_2 | wlor2_4 | Poland | R | 0 | 0 | 0 | 0 | 0 | 0 | 1 | 48,35 | 1490,481042 | -28 | 20013,987 | 42 | 0 | 1 | 0 | 1 | 0 | 0 | 0 | 0 | 1 | 1 | 0 | 0 | 0 | 0 | 0 | 0 |
| 52.65632 | 19.11217 | WLO_R_2 | wlor2_5 | Poland | R | 0 | 0 | 0 | 0 | 0 | 0 | 1 | 47,248 | 1490,692938 | -28 | 23263,986 | 0 | 0 | 1 | 0 | 1 | 0 | 0 | 0 | 0 | 0 | 1 | 0 | 0 | 0 | 0 | 0 | 0 |
| 52.65727 | 19.04815 | WLO_U_1 | wlou1_1 | Poland | U | 0 | 0 | 0 | 0 | 0 | 0 | 1 | 48,718 | 1493,235773 | -30 | 35451,874 | 869 | 1 | 1 | 0 | 0 | 1 | 0 | 1 | 0 | 0 | 0 | 0 | 0 | 0 | 0 | 0 | 1 |
| 52.65871 | 19.05137 | WLO_U_1 | wlou1_2 | Poland | U | 0 | 0 | 0 | 0 | 0 | 0 | 1 | 48 | 1493,244897 | -30 | 23750,8 | 463 | 1 | 1 | 0 | 0 | 1 | 0 | 0 | 0 | 0 | 0 | 0 | 0 | 0 | 0 | 0 | 0 |
| 52.6596 | 19.05585 | WLO_U_1 | wlou1_3 | Poland | U | 0 | 0 | 0 | 0 | 0 | 0 | 1 | 48,673 | 1493,152816 | -30 | 37002,468 | 4554 | 1 | 1 | 0 | 0 | 1 | 0 | 0 | 1 | 0 | 0 | 0 | 0 | 0 | 1 | 1 | 0 |
| 52.66069 | 19.05959 | WLO_U_1 | wlou1_4 | Poland | U | 0 | 0 | 0 | 0 | 0 | 0 | 1 | 47,968 | 1493,10802 | -30 | 27525,16 | 7404 | 0 | 1 | 0 | 0 | 1 | 0 | 0 | 1 | 0 | 0 | 0 | 0 | 0 | 1 | 0 | 0 |
| 52.66114 | 19.06377 | WLO_U_1 | wlou1_5 | Poland | U | 1 | 1 | 0 | 0 | 0 | 0 | 1 | 51,168 | 1492,992268 | -30 | 37368,546 | 4438 | 1 | 1 | 0 | 0 | 1 | 0 | 0 | 1 | 0 | 0 | 0 | 0 | 0 | 1 | 0 | 1 |
| 52.64873 | 19.04591 | WLO_U_2 | wlou2_1 | Poland | U | 0 | 0 | 0 | 0 | 0 | 0 | 1 | 57,572 | 1492,508632 | -30 | 11959,801 | 7746 | 1 | 1 | 0 | 0 | 0 | 0 | 0 | 1 | 1 | 0 | 0 | 1 | 0 | 0 | 0 | 0 |
| 52.6474 | 19.04363 | WLO_U_2 | wlou2_2 | Poland | U | 0 | 0 | 0 | 0 | 0 | 0 | 1 | 58,956 | 1492,634529 | -30 | 7949,229 | 12905 | 1 | 1 | 0 | 0 | 0 | 0 | 0 | 1 | 1 | 0 | 0 | 1 | 0 | 0 | 0 | 0 |
| 52.64705 | 19.03951 | WLO_U_2 | wlou2_3 | Poland | U | 0 | 0 | 0 | 0 | 0 | 0 | 1 | 59 | 1492,812396 | -30 | 4996,401 | 13575 | 0 | 1 | 0 | 0 | 0 | 0 | 0 | 0 | 1 | 0 | 0 | 1 | 1 | 0 | 0 | 0 |
| 52.64766 | 19.03634 | WLO_U_2 | wlou2_4 | Poland | U | 0 | 0 | 0 | 0 | 0 | 0 | 1 | 59,899 | 1493,069592 | -30 | 22291,029 | 6762 | 1 | 1 | 0 | 1 | 0 | 0 | 1 | 1 | 1 | 0 | 0 | 1 | 0 | 0 | 0 | 0 |
| 52.64994 | 19.03507 | WLO_U_2 | wlou2_5 | Poland | U | 0 | 0 | 0 | 0 | 0 | 0 | 1 | 52,054 | 1492,54356 | -30 | 17955,873 | 3580 | 0 | 1 | 1 | 0 | 0 | 0 | 0 | 0 | 1 | 0 | 0 | 1 | 0 | 0 | 0 | 0 |
| 43.15758 | 23.50627 | WRA_R_1 | wrar1_1 | Bulgaria | R | 1 | 1 | 0 | 0 | 0 | 0 | 0 | 786,344 | 516,6457986 | 365 | 53158,892 | 0 | 1 | 0 | 1 | 1 | 0 | 0 | 0 | 0 | 0 | 0 | 0 | 0 | 0 | 0 | 0 | 0 |
| 43.16008 | 23.50645 | WRA_R_1 | wrar1_2 | Bulgaria | R | 0 | 0 | 0 | 0 | 0 | 0 | 0 | 708,801 | 516,7477257 | 365 | 66623,663 | 0 | 1 | 0 | 1 | 1 | 0 | 0 | 0 | 0 | 0 | 0 | 0 | 0 | 0 | 0 | 0 | 0 |
| 43.16257 | 23.50448 | WRA_R_1 | wrar1_3 | Bulgaria | R | 0 | 0 | 0 | 0 | 0 | 0 | 1 | 663,583 | 517,0073479 | 365 | 48406,389 | 0 | 1 | 1 | 1 | 1 | 0 | 0 | 1 | 0 | 0 | 0 | 0 | 0 | 0 | 0 | 0 | 0 |
| 43.16553 | 23.50353 | WRA_R_1 | wrar1_4 | Bulgaria | R | 0 | 0 | 0 | 0 | 0 | 0 | 0 | 602,882 | 517,2128429 | 365 | 32019,353 | 243 | 1 | 1 | 1 | 1 | 0 | 0 | 1 | 0 | 0 | 1 | 1 | 0 | 0 | 0 | 0 | 0 |
| 43.16816 | 23.50507 | WRA_R_1 | wrar1_5 | Bulgaria | R | 0 | 0 | 0 | 0 | 0 | 0 | 1 | 567,6 | 517,219905 | 365 | 39548,301 | 194 | 1 | 1 | 1 | 1 | 0 | 0 | 1 | 1 | 1 | 1 | 1 | 0 | 0 | 0 | 0 | 0 |
| 43.18933 | 23.52362 | WRA_R_2 | wrar2_1 | Bulgaria | R | 1 | 1 | 0 | 0 | 0 | 0 | 0 | 450,907 | 516,8291472 | 368 | 43550,514 | 969 | 1 | 1 | 1 | 1 | 0 | 0 | 1 | 0 | 0 | 1 | 1 | 0 | 0 | 0 | 1 | 0 |
| 43.1903 | 23.52699 | WRA_R_2 | wrar2_2 | Bulgaria | R | 0 | 0 | 0 | 0 | 0 | 0 | 0 | 452,656 | 516,6249854 | 368 | 61223,931 | 0 | 1 | 1 | 1 | 1 | 0 | 0 | 1 | 0 | 0 | 1 | 0 | 0 | 0 | 0 | 0 | 0 |
| 43.19093 | 23.53058 | WRA_R_2 | wrar2_3 | Bulgaria | R | 0 | 0 | 0 | 0 | 0 | 0 | 0 | 451,056 | 516,3897144 | 368 | 48985,513 | 0 | 1 | 1 | 1 | 1 | 0 | 0 | 1 | 0 | 0 | 1 | 0 | 0 | 0 | 0 | 0 | 0 |
| 43.19144 | 23.53459 | WRA_R_2 | wrar2_4 | Bulgaria | R | 0 | 0 | 0 | 0 | 0 | 0 | 0 | 441,683 | 516,1172791 | 368 | 56430,075 | 262 | 1 | 1 | 1 | 1 | 0 | 0 | 1 | 0 | 0 | 1 | 1 | 0 | 0 | 0 | 0 | 0 |
| 43.19166 | 23.53952 | WRA_R_2 | wrar2_5 | Bulgaria | R | 0 | 0 | 1 | 1 | 0 | 0 | 0 | 414,036 | 515,7649073 | 368 | 65248,036 | 0 | 1 | 1 | 1 | 1 | 0 | 0 | 1 | 0 | 0 | 1 | 1 | 0 | 0 | 0 | 0 | 0 |
| 43.19909 | 23.57053 | WRA_U_1 | wrau1_1 | Bulgaria | U | 1 | 1 | 0 | 0 | 0 | 0 | 1 | 373,619 | 513,8295661 | 370 | 14536,725 | 1339 | 0 | 1 | 1 | 0 | 0 | 0 | 1 | 1 | 0 | 0 | 0 | 0 | 1 | 0 | 1 | 0 |
| 43.19981 | 23.56663 | WRA_U_1 | wrau1_2 | Bulgaria | U | 0 | 0 | 0 | 0 | 0 | 0 | 0 | 375,278 | 514,1500767 | 370 | 26368,312 | 5163 | 1 | 1 | 0 | 0 | 1 | 0 | 1 | 1 | 0 | 0 | 1 | 0 | 0 | 0 | 1 | 0 |
| 43.20192 | 23.56441 | WRA_U_1 | wrau1_3 | Bulgaria | U | 0 | 0 | 0 | 0 | 0 | 0 | 0 | 373,336 | 514,4119095 | 370 | 19093,741 | 10374 | 1 | 1 | 0 | 0 | 1 | 0 | 1 | 1 | 1 | 0 | 0 | 0 | 1 | 0 | 1 | 0 |
| 43.20357 | 23.56119 | WRA_U_1 | wrau1_4 | Bulgaria | U | 0 | 0 | 2 | 1 | 0 | 0 | 0 | 377,58 | 514,7266456 | 370 | 23920,736 | 12986 | 1 | 1 | 0 | 0 | 0 | 0 | 1 | 1 | 0 | 0 | 0 | 0 | 1 | 0 | 0 | 0 |

| | | | | | | | | | | | | | | | | | | | | | | | | | | | | | | | | | |
|---|---|---|---|---|---|---|---|---|---|---|---|---|---|---|---|---|---|---|---|---|---|---|---|---|---|---|---|---|---|---|---|---|---|
| 43.20547 | 23.55875 | WRA_U_1 | wrau1_5 | Bulgaria | U | 0 | 0 | 0 | 0 | 0 | 0 | 0 | 370,692 | 514,9966286 | 370 | 21226,91 | 18362 | 0 | 1 | 0 | 0 | 0 | 0 | 1 | 1 | 0 | 0 | 0 | 0 | 1 | 0 | 0 | 0 |
| 43.19847 | 23.54943 | WRA_U_2 | wrau2_1 | Bulgaria | U | 0 | 0 | 0 | 0 | 0 | 0 | 0 | 407,462 | 515,3530716 | 370 | 45953,599 | 5444 | 1 | 1 | 1 | 1 | 0 | 0 | 0 | 1 | 0 | 0 | 0 | 1 | 1 | 0 | 0 | 0 |
| 43.20063 | 23.54896 | WRA_U_2 | wrau2_2 | Bulgaria | U | 0 | 0 | 0 | 0 | 0 | 0 | 0 | 383,187 | 515,4893472 | 370 | 13173,732 | 10239 | 0 | 1 | 1 | 0 | 1 | 0 | 1 | 1 | 0 | 0 | 0 | 1 | 1 | 1 | 0 | 1 |
| 43.20313 | 23.55018 | WRA_U_2 | wrau2_3 | Bulgaria | U | 0 | 0 | 0 | 0 | 0 | 0 | 0 | 378,826 | 515,5171021 | 370 | 9726,221 | 26661 | 0 | 1 | 0 | 0 | 1 | 0 | 0 | 1 | 0 | 0 | 0 | 0 | 1 | 1 | 0 | 0 |
| 43.20522 | 23.54845 | WRA_U_2 | wrau2_4 | Bulgaria | U | 0 | 0 | 0 | 0 | 0 | 0 | 0 | 375,175 | 515,7425694 | 370 | 7395,969 | 24864 | 0 | 1 | 1 | 0 | 1 | 0 | 0 | 1 | 0 | 0 | 0 | 0 | 1 | 1 | 0 | 0 |
| 43.20773 | 23.54763 | WRA_U_2 | wrau2_5 | Bulgaria | U | 0 | 0 | 1 | 1 | 0 | 0 | 0 | 366,252 | 515,9215505 | 370 | 15836,204 | 17402 | 0 | 1 | 0 | 0 | 1 | 0 | 0 | 1 | 0 | 0 | 0 | 0 | 1 | 1 | 0 | 0 |
| 51.07322 | 17.14337 | WRO_R_1 | wror1_1 | Poland | R | 0 | 0 | 0 | 0 | 0 | 0 | 1 | 120 | 1440,524815 | 1 | 6987,729 | 0 | 0 | 1 | 0 | 0 | 0 | 0 | 1 | 0 | 0 | 0 | 0 | 0 | 0 | 0 | 0 | 0 |
| 51.07076 | 17.14101 | WRO_R_1 | wror1_2 | Poland | R | 3 | 1 | 0 | 0 | 0 | 0 | 1 | 119,168 | 1440,441086 | 1 | 13496,978 | 0 | 1 | 1 | 0 | 1 | 0 | 0 | 1 | 0 | 0 | 0 | 0 | 0 | 0 | 0 | 0 | 0 |
| 51.07036 | 17.13721 | WRO_R_1 | wror1_3 | Poland | R | 2 | 1 | 0 | 0 | 0 | 0 | 1 | 119 | 1440,599972 | 1 | 25325,967 | 0 | 1 | 1 | 0 | 1 | 0 | 0 | 1 | 0 | 0 | 1 | 0 | 0 | 0 | 0 | 0 | 0 |
| 51.07218 | 17.13455 | WRO_R_1 | wror1_4 | Poland | R | 2 | 1 | 0 | 0 | 0 | 0 | 1 | 119 | 1440,87439 | 1 | 30390,388 | 0 | 1 | 1 | 0 | 1 | 0 | 0 | 0 | 0 | 0 | 0 | 0 | 0 | 0 | 0 | 0 | 0 |
| 51.07373 | 17.13101 | WRO_R_1 | wror1_5 | Poland | R | 1 | 1 | 0 | 0 | 0 | 0 | 1 | 118,636 | 1441,170637 | 1 | 27663,307 | 0 | 1 | 1 | 0 | 1 | 0 | 0 | 1 | 0 | 0 | 1 | 0 | 0 | 0 | 0 | 0 | 0 |
| 51.08315 | 17.12038 | WRO_R_2 | wror2_1 | Poland | R | 2 | 1 | 0 | 0 | 0 | 0 | 1 | 118,798 | 1442,441193 | 1 | 9887,327 | 0 | 0 | 1 | 0 | 1 | 0 | 0 | 1 | 0 | 0 | 0 | 0 | 0 | 0 | 0 | 0 | 0 |
| 51.08501 | 17.11752 | WRO_R_2 | wror2_2 | Poland | R | 1 | 1 | 0 | 0 | 0 | 0 | 1 | 120,856 | 1442,728814 | 1 | 44517,4 | 0 | 1 | 0 | 0 | 1 | 0 | 0 | 0 | 0 | 1 | 0 | 0 | 0 | 0 | 0 | 0 | 0 |
| 51.08774 | 17.11639 | WRO_R_2 | wror2_3 | Poland | R | 2 | 1 | 0 | 0 | 0 | 0 | 1 | 118,862 | 1443,001498 | 1 | 43529,146 | 0 | 1 | 1 | 0 | 1 | 0 | 0 | 1 | 0 | 1 | 0 | 0 | 0 | 0 | 0 | 0 | 0 |
| 51.09038 | 17.11554 | WRO_R_2 | wror2_4 | Poland | R | 2 | 1 | 0 | 0 | 0 | 0 | 1 | 119,291 | 1443,253143 | 1 | 45525,597 | 0 | 1 | 1 | 0 | 1 | 0 | 0 | 0 | 0 | 0 | 0 | 0 | 0 | 0 | 0 | 0 | 0 |
| 51.09312 | 17.11446 | WRO_R_2 | wror2_5 | Poland | R | 1 | 1 | 0 | 0 | 0 | 0 | 1 | 118,112 | 1443,524005 | 1 | 33027,772 | 497 | 1 | 1 | 0 | 1 | 0 | 0 | 1 | 0 | 0 | 0 | 1 | 0 | 0 | 0 | 0 | 1 |
| 51.07358 | 17.00974 | WRO_U_1 | wrou1_1 | Poland | U | 2 | 1 | 0 | 0 | 0 | 0 | 1 | 123 | 1447,119539 | 1 | 57502,84 | 1000 | 1 | 1 | 0 | 0 | 1 | 0 | 1 | 0 | 0 | 0 | 0 | 0 | 0 | 0 | 1 | 1 |
| 51.0759 | 17.01003 | WRO_U_1 | wrou1_2 | Poland | U | 1 | 1 | 0 | 0 | 0 | 0 | 1 | 122,214 | 1447,288553 | 1 | 44943,832 | 2342 | 1 | 1 | 0 | 0 | 1 | 0 | 0 | 1 | 0 | 0 | 0 | 0 | 1 | 0 | 0 | 0 |
| 51.07872 | 17.01049 | WRO_U_1 | wrou1_3 | Poland | U | 2 | 1 | 0 | 0 | 0 | 0 | 1 | 122,857 | 1447,488874 | 1 | 49446,582 | 4897 | 1 | 1 | 0 | 0 | 1 | 0 | 1 | 1 | 0 | 0 | 0 | 0 | 1 | 0 | 0 | 0 |
| 51.08187 | 17.01195 | WRO_U_1 | wrou1_4 | Poland | U | 0 | 0 | 0 | 0 | 0 | 0 | 1 | 124 | 1447,666956 | 1 | 27241,147 | 13605 | 1 | 1 | 0 | 0 | 0 | 0 | 1 | 1 | 0 | 0 | 0 | 0 | 1 | 0 | 0 | 0 |
| 51.08505 | 17.01285 | WRO_U_1 | wrou1_5 | Poland | U | 0 | 0 | 0 | 0 | 0 | 0 | 1 | 124,18 | 1447,87394 | 1 | 29189,178 | 14541 | 1 | 1 | 0 | 0 | 0 | 0 | 1 | 1 | 0 | 0 | 0 | 0 | 1 | 0 | 0 | 0 |
| 51.07334 | 17.02589 | WRO_U_2 | wrou2_1 | Poland | U | 0 | 0 | 0 | 0 | 0 | 0 | 1 | 123 | 1446,304859 | 1 | 38374,043 | 2636 | 0 | 1 | 0 | 0 | 0 | 0 | 0 | 1 | 1 | 0 | 0 | 1 | 1 | 0 | 0 | 0 |
| 51.07561 | 17.02706 | WRO_U_2 | wrou2_2 | Poland | U | 0 | 0 | 0 | 0 | 0 | 0 | 1 | 126,085 | 1446,426852 | 1 | 25178,136 | 4391 | 0 | 1 | 0 | 0 | 1 | 0 | 0 | 1 | 1 | 0 | 0 | 1 | 1 | 0 | 0 | 0 |
| 51.07833 | 17.02683 | WRO_U_2 | wrou2_3 | Poland | U | 2 | 1 | 0 | 0 | 0 | 0 | 1 | 124 | 1446,652955 | 1 | 63347,484 | 0 | 1 | 1 | 0 | 0 | 1 | 0 | 0 | 0 | 1 | 0 | 0 | 1 | 0 | 0 | 0 | 0 |
| 51.08092 | 17.02785 | WRO_U_2 | wrou2_4 | Poland | U | 2 | 1 | 0 | 0 | 0 | 0 | 1 | 125,179 | 1446,80849 | 1 | 65411,822 | 520 | 1 | 1 | 0 | 0 | 1 | 0 | 0 | 0 | 1 | 0 | 0 | 1 | 0 | 0 | 0 | 0 |
| 51.08219 | 17.0303 | WRO_U_2 | wrou2_5 | Poland | U | 1 | 1 | 0 | 0 | 0 | 0 | 1 | 125,929 | 1446,788294 | 1 | 55597,886 | 0 | 1 | 1 | 0 | 0 | 1 | 0 | 1 | 0 | 1 | 0 | 0 | 1 | 0 | 0 | 0 | 1 |
| 48.21767 | 15.24447 | YBB_R_1 | ybbr1_1 | Austria | R | 3 | 1 | 0 | 0 | 0 | 0 | 1 | 214,412 | 1351,642327 | 3 | 30356,686 | 7455 | 1 | 0 | 0 | 1 | 0 | 0 | 0 | 0 | 1 | 1 | 0 | 0 | 0 | 0 | 0 | 0 |
| 48.21626 | 15.23974 | YBB_R_1 | ybbr1_2 | Austria | R | 2 | 1 | 0 | 0 | 0 | 0 | 0 | 214,608 | 1352,010779 | 3 | 36688,915 | 0 | 1 | 1 | 0 | 1 | 0 | 0 | 0 | 0 | 0 | 1 | 0 | 0 | 0 | 0 | 0 | 0 |
| 48.21496 | 15.23619 | YBB_R_1 | ybbr1_3 | Austria | R | 2 | 1 | 0 | 0 | 0 | 0 | 1 | 216,037 | 1351,809941 | 3 | 35320,055 | 0 | 1 | 0 | 0 | 1 | 0 | 0 | 0 | 0 | 0 | 0 | 0 | 0 | 0 | 0 | 0 | 1 |
| 48.21316 | 15.23146 | YBB_R_1 | ybbr1_4 | Austria | R | 1 | 1 | 0 | 0 | 0 | 0 | 0 | 211,752 | 1351,634101 | 3 | 22285,871 | 0 | 1 | 0 | 0 | 1 | 0 | 0 | 0 | 0 | 0 | 0 | 0 | 0 | 0 | 0 | 0 | 0 |
| 48.21177 | 15.23603 | YBB_R_1 | ybbr1_5 | Austria | R | 1 | 1 | 0 | 0 | 0 | 0 | 1 | 212,256 | 1351,434867 | 3 | 30344,331 | 0 | 1 | 0 | 0 | 1 | 0 | 0 | 0 | 0 | 0 | 0 | 0 | 0 | 0 | 0 | 0 | 0 |
| 48.19704 | 15.10402 | YBB_R_2 | ybbr2_1 | Austria | R | 1 | 1 | 0 | 0 | 0 | 0 | 1 | 234,825 | 1359,261634 | 2 | 28495,253 | 4116 | 1 | 1 | 0 | 1 | 0 | 0 | 0 | 0 | 0 | 0 | 0 | 1 | 0 | 0 | 0 | 0 |
| 48.19899 | 15.10579 | YBB_R_2 | ybbr2_2 | Austria | R | 2 | 1 | 0 | 0 | 0 | 0 | 0 | 249,118 | 1359,263797 | 2 | 26454,413 | 5316 | 1 | 1 | 0 | 1 | 0 | 0 | 0 | 0 | 1 | 0 | 0 | 1 | 0 | 0 | 0 | 0 |
| 48.2015 | 15.10426 | YBB_R_2 | ybbr2_3 | Austria | R | 4 | 1 | 0 | 0 | 0 | 0 | 0 | 283,528 | 1359,502757 | 2 | 30564,021 | 1339 | 1 | 1 | 0 | 1 | 0 | 0 | 0 | 0 | 1 | 0 | 1 | 0 | 0 | 0 | 0 | 1 |
| 48.20168 | 15.10063 | YBB_R_2 | ybbr2_4 | Austria | R | 0 | 0 | 0 | 0 | 0 | 0 | 1 | 314,449 | 1359,744788 | 2 | 30886,876 | 0 | 1 | 1 | 0 | 1 | 0 | 0 | 0 | 0 | 0 | 0 | 0 | 0 | 0 | 0 | 0 | 1 |
| 48.20022 | 15.09724 | YBB_R_2 | ybbr2_5 | Austria | R | 1 | 1 | 0 | 0 | 0 | 0 | 1 | 304,781 | 1359,88222 | 2 | 22673,63 | 1792 | 1 | 1 | 0 | 1 | 0 | 0 | 0 | 1 | 0 | 0 | 1 | 0 | 0 | 0 | 0 | 0 |
| 48.21213 | 15.21741 | YBB_U_1 | ybbu1_1 | Austria | U | 0 | 0 | 0 | 0 | 0 | 0 | 1 | 213,452 | 1352,851559 | 3 | 4323,9 | 13974 | 0 | 1 | 0 | 0 | 0 | 0 | 0 | 0 | 1 | 0 | 0 | 1 | 0 | 1 | 0 | 0 |
| 48.2123 | 15.2121 | YBB_U_1 | ybbu1_2 | Austria | U | 0 | 0 | 0 | 0 | 0 | 0 | 1 | 215,317 | 1353,187262 | 3 | 4462,18 | 18151 | 0 | 1 | 0 | 0 | 1 | 0 | 0 | 1 | 0 | 0 | 0 | 0 | 0 | 1 | 0 | 0 |
| 48.21184 | 15.20769 | YBB_U_1 | ybbu1_3 | Austria | U | 0 | 0 | 0 | 0 | 0 | 0 | 1 | 215 | 1353,435297 | 3 | 5908,403 | 13185 | 0 | 1 | 0 | 0 | 0 | 0 | 0 | 1 | 0 | 1 | 0 | 0 | 0 | 1 | 0 | 0 |
| 48.2118 | 15.20373 | YBB_U_1 | ybbu1_4 | Austria | U | 1 | 1 | 0 | 0 | 0 | 0 | 1 | 210,765 | 1353,70782 | 3 | 7993,773 | 7751 | 0 | 1 | 0 | 0 | 0 | 0 | 1 | 1 | 1 | 1 | 0 | 1 | 1 | 0 | 0 | 0 |
| 48.212 | 15.1996 | YBB_U_1 | ybbu1_5 | Austria | U | 1 | 1 | 0 | 0 | 0 | 0 | 0 | 213,8 | 1353,990061 | 3 | 8073,653 | 6983 | 0 | 1 | 0 | 0 | 0 | 0 | 0 | 0 | 1 | 1 | 0 | 1 | 0 | 0 | 0 | 1 |
| 48.18807 | 15.08309 | YBB_U_2 | ybbu2_1 | Austria | U | 0 | 0 | 0 | 0 | 0 | 0 | 0 | 228,928 | 1360,098499 | 1 | 11659,318 | 8068 | 0 | 1 | 0 | 1 | 0 | 1 | 0 | 0 | 1 | 0 | 1 | 1 | 0 | 1 | 0 | 0 |
| 48.18589 | 15.08333 | YBB_U_2 | ybbu2_2 | Austria | U | 1 | 1 | 0 | 0 | 0 | 0 | 1 | 224,981 | 1359,958881 | 1 | 18462,243 | 10731 | 0 | 1 | 1 | 0 | 1 | 0 | 0 | 0 | 1 | 0 | 0 | 1 | 0 | 1 | 0 | 1 |
| 48.18471 | 15.08693 | YBB_U_2 | ybbu2_3 | Austria | U | 0 | 0 | 0 | 0 | 0 | 0 | 1 | 223,862 | 1359,669357 | 1 | 6412,452 | 14631 | 0 | 1 | 1 | 0 | 0 | 0 | 0 | 0 | 1 | 0 | 0 | 1 | 0 | 0 | 0 | 0 |
| 48.18624 | 15.0888 | YBB_U_2 | ybbu2_4 | Austria | U | 0 | 0 | 0 | 0 | 0 | 0 | 0 | 225 | 1359,64231 | 1 | 6727,079 | 11973 | 0 | 1 | 0 | 0 | 0 | 0 | 0 | 0 | 1 | 0 | 0 | 1 | 1 | 0 | 0 | 1 |
| 48.18762 | 15.09113 | YBB_U_2 | ybbu2_5 | Austria | U | 0 | 0 | 0 | 0 | 0 | 0 | 0 | 224 | 1359,568093 | 1 | 5365,463 | 10192 | 0 | 1 | 0 | 0 | 0 | 0 | 0 | 0 | 1 | 0 | 0 | 1 | 1 | 0 | 0 | 0 |
| 42.4374 | 25.63887 | STA_U_1 | stau1_1 | Bulgaria | U | 0 | 0 | 0 | 0 | 0 | 0 | 0 | 213,708 | 324,7278328 | 537 | 65871,068 | 0 | 1 | 1 | 1 | 1 | 0 | 0 | 0 | 0 | 0 | 0 | 0 | 0 | 0 | 0 | 0 | 0 |
| 42.44002 | 25.63759 | STA_U_1 | stau1_2 | Bulgaria | U | 0 | 0 | 1 | 1 | 0 | 0 | 0 | 222,373 | 324,9526456 | 537 | 40805,405 | 7358 | 1 | 1 | 1 | 1 | 1 | 0 | 0 | 1 | 1 | 0 | 0 | 0 | 0 | 0 | 1 | 0 |
| 42.44296 | 25.63649 | STA_U_1 | stau1_3 | Bulgaria | U | 0 | 0 | 0 | 0 | 0 | 0 | 0 | 231,656 | 325,1756498 | 537 | 55156,633 | 313 | 1 | 1 | 1 | 1 | 1 | 0 | 0 | 1 | 0 | 1 | 0 | 0 | 0 | 0 | 0 | 1 |
| 42.444 | 25.63891 | STA_U_1 | stau1_4 | Bulgaria | U | 0 | 0 | 0 | 0 | 0 | 0 | 0 | 229,77 | 325,0488208 | 537 | 49360,197 | 499 | 1 | 1 | 1 | 1 | 0 | 0 | 1 | 0 | 0 | 1 | 1 | 0 | 0 | 0 | 0 | 0 |
| 42.44244 | 25.64107 | STA_U_1 | stau1_5 | Bulgaria | U | 0 | 0 | 0 | 0 | 0 | 0 | 0 | 220,032 | 324,8134571 | 537 | 13428,504 | 159 | 1 | 1 | 0 | 1 | 0 | 0 | 1 | 0 | 1 | 0 | 1 | 0 | 0 | 0 | 0 | 0 |
| 42.43517 | 25.65407 | STA_U_2 | stau2_1 | Bulgaria | U | 0 | 0 | 0 | 0 | 0 | 0 | 0 | 223,175 | 323,4969862 | 537 | 35344,402 | 1243 | 1 | 1 | 1 | 1 | 0 | 0 | 0 | 0 | 0 | 0 | 1 | 0 | 0 | 0 | 0 | 0 |
| 42.43331 | 25.65132 | STA_U_2 | stau2_2 | Bulgaria | U | 0 | 0 | 0 | 0 | 0 | 0 | 0 | 213,319 | 323,608292 | 537 | 21288,592 | 5292 | 1 | 1 | 1 | 0 | 1 | 0 | 1 | 1 | 0 | 0 | 0 | 1 | 1 | 0 | 0 | 0 |
| 42.43177 | 25.65319 | STA_U_2 | stau2_3 | Bulgaria | U | 0 | 0 | 1 | 1 | 1 | 1 | 0 | 212,936 | 323,395588 | 537 | 42880,514 | 5168 | 1 | 1 | 1 | 1 | 1 | 0 | 1 | 1 | 1 | 0 | 0 | 1 | 0 | 0 | 0 | 0 |
| 42.43043 | 25.65638 | STA_U_2 | stau2_4 | Bulgaria | U | 0 | 0 | 0 | 0 | 0 | 0 | 0 | 219,562 | 323,0933596 | 537 | 38537,658 | 6973 | 1 | 1 | 1 | 1 | 0 | 0 | 1 | 1 | 1 | 0 | 0 | 1 | 1 | 0 | 0 | 0 |
| 42.43116 | 25.65962 | STA_U_2 | stau2_5 | Bulgaria | U | 0 | 0 | 1 | 1 | 1 | 1 | 0 | 215,111 | 322,8902582 | 537 | 59322,267 | 0 | 1 | 1 | 1 | 1 | 0 | 0 | 0 | 0 | 0 | 0 | 0 | 0 | 0 | 0 | 0 | 0 |
| 49.19945 | 18.75622 | ZIL_R_1 | zilr1_1 | Slovak | R | 1 | 1 | 0 | 0 | 0 | 0 | 0 | 393,608 | 1211,697363 | 202 | 59343,766 | 903 | 1 | 1 | 1 | 1 | 0 | 0 | 0 | 0 | 1 | 0 | 1 | 0 | 0 | 0 | 0 | 0 |
| 49.19681 | 18.75633 | ZIL_R_1 | zilr1_2 | Slovak | R | 2 | 1 | 0 | 0 | 0 | 0 | 0 | 388,227 | 1211,490992 | 202 | 56746,121 | 381 | 1 | 0 | 1 | 1 | 0 | 0 | 0 | 1 | 0 | 0 | 1 | 0 | 0 | 0 | 1 | 0 |
| 49.19313 | 18.75743 | ZIL_R_1 | zilr1_3 | Slovak | R | 0 | 0 | 0 | 0 | 0 | 0 | 0 | 380,196 | 1211,145445 | 202 | 3424,034 | 0 | 1 | 1 | 1 | 1 | 0 | 0 | 0 | 0 | 1 | 0 | 1 | 0 | 0 | 0 | 0 | 0 |
| 49.19 | 18.75773 | ZIL_R_1 | zilr1_4 | Slovak | R | 0 | 0 | 0 | 0 | 0 | 0 | 0 | 413 | 1210,888902 | 202 | 4167,735 | 0 | 0 | 1 | 0 | 1 | 0 | 0 | 0 | 0 | 1 | 0 | 1 | 0 | 0 | 0 | 0 | 0 |
| 49.18709 | 18.75806 | ZIL_R_1 | zilr1_5 | Slovak | R | 0 | 0 | 1 | 1 | 0 | 0 | 0 | 422,484 | 1210,64677 | 202 | 3589,63 | 0 | 0 | 1 | 0 | 1 | 0 | 0 | 1 | 0 | 0 | 0 | 1 | 0 | 0 | 0 | 0 | 0 |
| 49.1919 | 18.78691 | ZIL_R_2 | zilr2_1 | Slovak | R | 0 | 0 | 0 | 0 | 0 | 0 | 1 | 373,82 | 1209,508001 | 203 | 10378,657 | 6641 | 0 | 1 | 0 | 0 | 0 | 0 | 1 | 0 | 0 | 0 | 0 | 0 | 0 | 0 | 0 | 0 |
| 49.19455 | 18.78604 | ZIL_R_2 | zilr2_2 | Slovak | R | 1 | 1 | 0 | 0 | 0 | 0 | 0 | 364,841 | 1209,75816 | 203 | 29985,612 | 2975 | 1 | 1 | 0 | 1 | 1 | 1 | 0 | 0 | 1 | 0 | 0 | 1 | 0 | 0 | 0 | 1 |
| 49.19633 | 18.78831 | ZIL_R_2 | zilr2_3 | Slovak | R | 2 | 1 | 0 | 0 | 0 | 0 | 0 | 408,368 | 1209,776973 | 203 | 61804,549 | 0 | 1 | 0 | 0 | 1 | 0 | 0 | 0 | 0 | 0 | 0 | 0 | 0 | 0 | 0 | 0 | 0 |
| 49.19367 | 18.79078 | ZIL_R_2 | zilr2_4 | Slovak | R | 2 | 1 | 0 | 0 | 0 | 0 | 1 | 401,231 | 1209,440738 | 203 | 50591,194 | 2311 | 1 | 1 | 1 | 1 | 0 | 0 | 0 | 0 | 1 | 0 | 1 | 1 | 0 | 0 | 0 | 1 |
| 49.19193 | 18.79371 | ZIL_R_2 | zilr2_5 | Slovak | R | 0 | 0 | 0 | 0 | 0 | 0 | 0 | 376,844 | 1209,152846 | 203 | 25798,381 | 6373 | 1 | 1 | 1 | 1 | 0 | 0 | 0 | 0 | 1 | 0 | 0 | 1 | 0 | 0 | 0 | 0 |
| 49.21152 | 18.77075 | ZIL_U_1 | zilu1_1 | Slovak | U | 1 | 1 | 0 | 0 | 0 | 0 | 1 | 360 | 1211,870746 | 200 | 21971,601 | 7585 | 0 | 1 | 1 | 0 | 0 | 1 | 1 | 1 | 0 | 0 | 0 | 1 | 1 | 1 | 0 | 0 |
| 49.21394 | 18.77001 | ZIL_U_1 | zilu1_2 | Slovak | U | 2 | 1 | 0 | 0 | 0 | 0 | 1 | 357,583 | 1212,098771 | 200 | 11721,199 | 10567 | 0 | 1 | 1 | 0 | 0 | 1 | 0 | 1 | 1 | 0 | 1 | 1 | 1 | 1 | 0 | 0 |
| 49.21484 | 18.76645 | ZIL_U_1 | zilu1_3 | Slovak | U | 1 | 1 | 0 | 0 | 0 | 0 | 0 | 356,093 | 1212,355499 | 200 | 16032,39 | 12571 | 0 | 1 | 1 | 0 | 0 | 0 | 0 | 1 | 0 | 0 | 0 | 0 | 1 | 1 | 0 | 0 |
| 49.21375 | 18.76276 | ZIL_U_1 | zilu1_4 | Slovak | U | 1 | 1 | 0 | 0 | 0 | 0 | 1 | 355,936 | 1212,461318 | 200 | 30932,997 | 9325 | 0 | 1 | 1 | 0 | 0 | 0 | 0 | 1 | 0 | 0 | 0 | 0 | 1 | 1 | 0 | 0 |
| 49.21496 | 18.75972 | ZIL_U_1 | zilu1_5 | Slovak | U | 1 | 1 | 0 | 0 | 0 | 0 | 1 | 349,13 | 1212,714731 | 200 | 31719,845 | 9512 | 0 | 1 | 1 | 0 | 1 | 0 | 0 | 0 | 0 | 0 | 0 | 0 | 0 | 1 | 0 | 0 |
| 49.2219 | 18.73419 | ZIL_U_2 | zilu2_1 | Slovak | U | 0 | 0 | 0 | 0 | 0 | 0 | 0 | 347,16 | 1214,587375 | 199 | 9219,762 | 17200 | 0 | 1 | 1 | 0 | 0 | 0 | 1 | 1 | 0 | 0 | 0 | 0 | 1 | 0 | 0 | 0 |
| 49.22401 | 18.73383 | ZIL_U_2 | zilu2_2 | Slovak | U | 0 | 0 | 0 | 0 | 0 | 0 | 0 | 344,212 | 1214,775416 | 199 | 9333,797 | 18113 | 0 | 1 | 0 | 0 | 0 | 0 | 1 | 1 | 1 | 0 | 0 | 1 | 1 | 0 | 0 | 0 |
| 49.22605 | 18.73476 | ZIL_U_2 | zilu2_3 | Slovak | U | 1 | 1 | 0 | 0 | 0 | 0 | 0 | 338,274 | 1214,879453 | 199 | 8431,281 | 24366 | 1 | 1 | 1 | 1 | 0 | 0 | 0 | 1 | 0 | 0 | 0 | 0 | 1 | 0 | 0 | 0 |
| 49.22663 | 18.73863 | ZIL_U_2 | zilu2_4 | Slovak | U | 0 | 0 | 0 | 0 | 0 | 0 | 0 | 332,282 | 1214,721962 | 199 | 8409,089 | 25164 | 0 | 1 | 1 | 0 | 1 | 0 | 1 | 1 | 0 | 0 | 0 | 0 | 1 | 0 | 0 | 0 |
| 49.22591 | 18.74209 | ZIL_U_2 | zilu2_5 | Slovak | U | 0 | 0 | 0 | 0 | 0 | 0 | 0 | 331,869 | 1214,488871 | 199 | 6250,187 | 26847 | 0 | 1 | 0 | 0 | 1 | 0 | 1 | 1 | 0 | 0 | 0 | 0 | 1 | 0 | 0 | 0 |
| 51.11416 | 15.88817 | ZLO_R_1 | zlor1_1 | Poland | R | 2 | 1 | 0 | 0 | 0 | 0 | 0 | 230,676 | 1506,871433 | -62 | 19012,112 | 3860 | 1 | 1 | 0 | 1 | 0 | 0 | 0 | 0 | 1 | 1 | 1 | 0 | 0 | 0 | 0 | 0 |
| 51.11593 | 15.88994 | ZLO_R_1 | zlor1_2 | Poland | R | 2 | 1 | 0 | 0 | 0 | 0 | 0 | 238,06 | 1506,922888 | -62 | 14229,507 | 3167 | 1 | 1 | 0 | 0 | 0 | 0 | 1 | 0 | 0 | 1 | 1 | 0 | 0 | 0 | 1 | 0 |
| 51.11886 | 15.8894 | ZLO_R_1 | zlor1_3 | Poland | R | 2 | 1 | 0 | 0 | 0 | 0 | 0 | 249,744 | 1507,169916 | -62 | 22555,689 | 1000 | 1 | 1 | 0 | 1 | 0 | 0 | 0 | 0 | 0 | 1 | 0 | 0 | 0 | 0 | 1 | 0 |
| 51.12125 | 15.88968 | ZLO_R_1 | zlor1_4 | Poland | R | 1 | 1 | 0 | 0 | 0 | 0 | 0 | 265,44 | 1507,337305 | -62 | 23238,861 | 1000 | 1 | 1 | 0 | 1 | 0 | 0 | 0 | 0 | 0 | 1 | 0 | 0 | 0 | 0 | 0 | 0 |
| 51.12117 | 15.89483 | ZLO_R_1 | zlor1_5 | Poland | R | 2 | 1 | 0 | 0 | 0 | 0 | 0 | 268,572 | 1507,063749 | -62 | 38008,899 | 1000 | 1 | 1 | 0 | 1 | 0 | 0 | 0 | 0 | 0 | 1 | 0 | 0 | 0 | 0 | 1 | 0 |
| 51.13561 | 15.88253 | ZLO_R_2 | zlor2_1 | Poland | R | 1 | 0 | 0 | 0 | 0 | 0 | 0 | 198,024 | 1507,71491 | -62 | 11574,867 | 0 | 1 | 1 | 0 | 1 | 0 | 0 | 1 | 0 | 0 | 0 | 0 | 0 | 0 | 0 | 0 | 0 |
| 51.13496 | 15.8871 | ZLO_R_2 | zlor2_2 | Poland | R | 2 | 0 | 0 | 0 | 0 | 0 | 0 | 197,022 | 1507,980521 | -62 | 6073,396 | 0 | 0 | 1 | 0 | 0 | 0 | 0 | 1 | 0 | 0 | 0 | 0 | 0 | 0 | 0 | 0 | 0 |
| 51.13504 | 15.89142 | ZLO_R_2 | zlor2_3 | Poland | R | 2 | 1 | 0 | 0 | 0 | 0 | 1 | 195,769 | 1508,289009 | -62 | 22066,924 | 39 | 1 | 1 | 0 | 1 | 0 | 0 | 1 | 0 | 1 | 0 | 1 | 0 | 0 | 0 | 0 | 0 |
| 51.13411 | 15.89603 | ZLO_R_2 | zlor2_4 | Poland | R | 0 | 1 | 0 | 0 | 0 | 0 | 1 | 195,076 | 1508,508778 | -62 | 10360,986 | 63 | 0 | 1 | 0 | 0 | 0 | 0 | 1 | 0 | 0 | 0 | 1 | 0 | 0 | 0 | 0 | 0 |
| 51.1335 | 15.90015 | ZLO_R_2 | zlor2_5 | Poland | R | 0 | 1 | 0 | 0 | 0 | 0 | 1 | 193,975 | 1508,786406 | -62 | 23210,618 | 691 | 1 | 1 | 0 | 1 | 0 | 0 | 1 | 0 | 1 | 0 | 1 | 0 | 0 | 0 | 0 | 0 |
| 51.12746 | 15.91028 | ZLO_U_1 | zlou1_1 | Poland | U | 1 | 1 | 0 | 0 | 0 | 0 | 1 | 189,005 | 1506,755552 | -62 | 23006,041 | 5357 | 1 | 1 | 0 | 1 | 0 | 0 | 0 | 1 | 0 | 1 | 1 | 0 | 0 | 1 | 1 | 1 |
| 51.12556 | 15.91078 | ZLO_U_1 | zlou1_2 | Poland | U | 1 | 1 | 0 | 0 | 0 | 0 | 1 | 200,026 | 1506,531842 | -62 | 44668,177 | 7526 | 1 | 1 | 0 | 1 | 0 | 0 | 0 | 0 | 1 | 0 | 0 | 1 | 0 | 1 | 0 | 0 |
| 51.12275 | 15.91185 | ZLO_U_1 | zlou1_3 | Poland | U | 0 | 0 | 0 | 0 | 0 | 0 | 1 | 220,638 | 1506,272496 | -62 | 12417,643 | 11089 | 0 | 1 | 0 | 1 | 1 | 0 | 0 | 0 | 1 | 0 | 0 | 1 | 0 | 1 | 0 | 0 |
| 51.12132 | 15.91569 | ZLO_U_1 | zlou1_4 | Poland | U | 0 | 0 | 0 | 0 | 0 | 0 | 1 | 231 | 1505,988835 | -62 | 7471,22 | 18444 | 0 | 1 | 0 | 0 | 0 | 0 | 0 | 1 | 0 | 0 | 0 | 1 | 1 | 1 | 0 | 0 |
| 51.12288 | 15.91921 | ZLO_U_1 | zlou1_5 | Poland | U | 0 | 0 | 0 | 0 | 0 | 0 | 1 | 223,818 | 1505,929237 | -62 | 24349,085 | 8016 | 1 | 1 | 0 | 1 | 0 | 0 | 0 | 1 | 1 | 0 | 0 | 1 | 0 | 1 | 0 | 0 |
| 51.12767 | 15.92036 | ZLO_U_2 | zlou2_1 | Poland | U | 1 | 1 | 0 | 0 | 0 | 0 | 1 | 211,937 | 1506,237849 | -62 | 23403,268 | 9901 | 1 | 1 | 0 | 0 | 1 | 1 | 0 | 1 | 0 | 0 | 0 | 1 | 1 | 1 | 0 | 0 |
| 51.1304 | 15.92077 | ZLO_U_2 | zlou2_2 | Poland | U | 0 | 0 | 0 | 0 | 0 | 0 | 1 | 185 | 1506,415164 | -62 | 34290,065 | 984 | 1 | 1 | 0 | 1 | 0 | 1 | 0 | 1 | 0 | 1 | 1 | 0 | 0 | 0 | 1 | 0 |
| 51.13187 | 15.9245 | ZLO_U_2 | zlou2_3 | Poland | U | 1 | 1 | 0 | 0 | 0 | 0 | 1 | 185,322 | 1506,346857 | -62 | 28689,427 | 1245 | 1 | 1 | 0 | 1 | 0 | 1 | 0 | 0 | 1 | 1 | 1 | 0 | 0 | 0 | 0 | 1 |
| 51.13076 | 15.93062 | ZLO_U_2 | zlou2_4 | Poland | U | 2 | 1 | 0 | 0 | 0 | 0 | 1 | 196,288 | 1505,944843 | -62 | 45009,845 | 6442 | 1 | 0 | 0 | 1 | 0 | 0 | 0 | 1 | 0 | 0 | 0 | 1 | 0 | 0 | 0 | 0 |
| 51.12796 | 15.92821 | ZLO_U_2 | zlou2_5 | Poland | U | 0 | 0 | 0 | 0 | 0 | 0 | 1 | 207,277 | 1505,851069 | -62 | 16437,388 | 8206 | 1 | 0 | 1 | 1 | 0 | 0 | 0 | 1 | 1 | 0 | 0 | 1 | 1 | 0 | 0 | 0 |
| 48.55361 | 19.28751 | ZVO_R_1 | zvor1_1 | Slovakia | R | 0 | 0 | 1 | 1 | 0 | 0 | 1 | 342,968 | 1133,908786 | 238 | 9041,569 | 6799 | 0 | 1 | 0 | 0 | 0 | 0 | 1 | 0 | 1 | 1 | 0 | 1 | 0 | 0 | 1 | 1 |
| 48.55551 | 19.28856 | ZVO_R_1 | zvor1_2 | Slovakia | R | 1 | 1 | 0 | 0 | 0 | 0 | 1 | 350,806 | 1133,996405 | 238 | 2793,017 | 11781 | 0 | 1 | 0 | 0 | 0 | 0 | 0 | 0 | 1 | 1 | 0 | 1 | 0 | 0 | 0 | 0 |
| 48.55706 | 19.28783 | ZVO_R_1 | zvor1_3 | Slovakia | R | 1 | 1 | 0 | 0 | 0 | 0 | 1 | 349,423 | 1134,153758 | 238 | 17962,484 | 7110 | 1 | 1 | 0 | 1 | 0 | 0 | 0 | 0 | 1 | 0 | 0 | 1 | 0 | 0 | 1 | 0 |
| 48.55962 | 19.28657 | ZVO_R_1 | zvor1_4 | Slovakia | R | 1 | 1 | 0 | 0 | 0 | 0 | 1 | 350,412 | 1134,415653 | 238 | 22408,428 | 3251 | 1 | 1 | 1 | 1 | 0 | 0 | 1 | 0 | 0 | 0 | 0 | 1 | 0 | 0 | 0 | 0 |
| 48.56224 | 19.28539 | ZVO_R_1 | zvor1_5 | Slovakia | R | 0 | 0 | 0 | 0 | 0 | 0 | 1 | 353,706 | 1134,679541 | 238 | 4446,388 | 0 | 1 | 1 | 0 | 1 | 0 | 0 | 0 | 0 | 0 | 0 | 0 | 0 | 0 | 0 | 0 | 0 |
| 48.5493 | 19.31247 | ZVO_R_2 | zvor2_1 | Slovakia | R | 0 | 0 | 0 | 0 | 0 | 0 | 1 | 359 | 1132,237466 | 240 | 1530,786 | 7964 | 0 | 1 | 0 | 0 | 0 | 0 | 0 | 0 | 1 | 0 | 0 | 1 | 1 | 0 | 0 | 0 |
| 48.55189 | 19.31366 | ZVO_R_2 | zvor2_2 | Slovakia | R | 0 | 0 | 0 | 0 | 0 | 0 | 1 | 352,913 | 1132,371195 | 240 | 4261,851 | 4037 | 1 | 1 | 0 | 1 | 0 | 0 | 1 | 0 | 1 | 0 | 0 | 1 | 0 | 0 | 1 | 0 |

| | | | | | | | | | | | | | | | | | | | | | | | | | | | | | | | | | |
|---|---|---|---|---|---|---|---|---|---|---|---|---|---|---|---|---|---|---|---|---|---|---|---|---|---|---|---|---|---|---|---|---|---|
| 48.55328 | 19.31076 | ZVO_R_2 | zvor2_3 | Slovakia | R | 2 | 1 | 0 | 0 | 0 | 0 | 1 | 349,96 | 1132,633358 | 240 | 10371,817 | 3072 | 0 | 1 | 0 | 0 | 0 | 0 | 1 | 0 | 0 | 0 | 0 | 0 | 0 | 0 | 1 | 0 |
| 48.55482 | 19.31179 | ZVO_R_2 | zvor2_4 | Slovakia | R | 1 | 1 | 0 | 0 | 0 | 0 | 1 | 344,36 | 1132,695551 | 240 | 18820,66 | 1700 | 0 | 1 | 0 | 1 | 0 | 0 | 1 | 0 | 0 | 1 | 0 | 0 | 0 | 0 | 0 | 0 |
| 48.55705 | 19.31342 | ZVO_R_2 | zvor2_5 | Slovakia | R | 0 | 0 | 0 | 0 | 0 | 0 | 1 | 345 | 1132,778776 | 240 | 11092,032 | 0 | 0 | 1 | 0 | 1 | 0 | 0 | 1 | 0 | 0 | 1 | 0 | 0 | 0 | 0 | 0 | 0 |
| 48.57102 | 19.11748 | ZVO_U_1 | zvou1_1 | Slovakia | U | 1 | 1 | 0 | 0 | 0 | 0 | 1 | 287,624 | 1144,4029 | 228 | 19420,513 | 20543 | 1 | 1 | 0 | 1 | 0 | 1 | 0 | 1 | 0 | 1 | 1 | 0 | 0 | 1 | 1 | 1 |
| 48.57187 | 19.12095 | ZVO_U_1 | zvou1_2 | Slovakia | U | 1 | 1 | 0 | 0 | 0 | 0 | 1 | 284,887 | 1144,280808 | 228 | 27381,122 | 6415 | 0 | 1 | 1 | 0 | 1 | 0 | 0 | 1 | 0 | 0 | 0 | 0 | 1 | 0 | 0 | 0 |
| 48.57192 | 19.12449 | ZVO_U_1 | zvou1_3 | Slovakia | U | 1 | 1 | 0 | 0 | 0 | 0 | 0 | 283,238 | 1144,090732 | 228 | 18750,272 | 24954 | 0 | 1 | 1 | 0 | 1 | 0 | 1 | 1 | 0 | 0 | 0 | 0 | 0 | 1 | 1 | 1 |
| 48.57265 | 19.12856 | ZVO_U_1 | zvou1_4 | Slovakia | U | 0 | 0 | 0 | 0 | 0 | 0 | 1 | 285,037 | 1143,926601 | 228 | 30730,421 | 17119 | 1 | 1 | 0 | 1 | 0 | 0 | 0 | 1 | 1 | 0 | 0 | 1 | 0 | 1 | 1 | 0 |
| 48.57385 | 19.13218 | ZVO_U_1 | zvou1_5 | Slovakia | U | 0 | 0 | 0 | 0 | 0 | 0 | 1 | 285,611 | 1143,821185 | 228 | 29701,704 | 12313 | 1 | 1 | 0 | 1 | 0 | 0 | 1 | 1 | 0 | 0 | 1 | 0 | 0 | 0 | 1 | 0 |
| 48.57639 | 19.10985 | ZVO_U_2 | zvou2_1 | Slovakia | U | 0 | 0 | 0 | 0 | 0 | 0 | 1 | 284,458 | 1145,221721 | 228 | 22446,916 | 4898 | 1 | 1 | 0 | 1 | 0 | 0 | 1 | 0 | 0 | 0 | 0 | 0 | 0 | 1 | 0 | 0 |
| 48.57557 | 19.11369 | ZVO_U_2 | zvou2_2 | Slovakia | U | 0 | 0 | 0 | 0 | 0 | 0 | 0 | 286,113 | 1144,952219 | 228 | 31280,601 | 9695 | 1 | 1 | 0 | 1 | 0 | 0 | 1 | 1 | 0 | 1 | 1 | 0 | 1 | 0 | 0 | 1 |
| 48.57775 | 19.11368 | ZVO_U_2 | zvou2_3 | Slovakia | U | 2 | 1 | 0 | 0 | 0 | 0 | 0 | 284 | 1145,116612 | 228 | 42065,589 | 2637 | 1 | 1 | 0 | 0 | 1 | 0 | 1 | 0 | 0 | 1 | 0 | 0 | 1 | 0 | 0 | 0 |
| 48.58045 | 19.11406 | ZVO_U_2 | zvou2_4 | Slovakia | U | 0 | 0 | 0 | 0 | 0 | 0 | 0 | 284 | 1145,300655 | 228 | 19533,063 | 321 | 0 | 1 | 0 | 0 | 1 | 0 | 0 | 1 | 1 | 1 | 0 | 1 | 0 | 0 | 0 | 1 |
| 48.5807 | 19.11796 | ZVO_U_2 | zvou2_5 | Slovakia | U | 0 | 0 | 0 | 0 | 0 | 0 | 1 | 285 | 1145,108391 | 228 | 17834,374 | 2848 | 0 | 1 | 0 | 1 | 0 | 0 | 1 | 0 | 1 | 1 | 0 | 1 | 0 | 0 | 0 | 0 |

Table S1. Database of examined transects with records of woodpeckers and environmental variables collected across Lesser Poland.

| NS | EW | T_name | Altitude | Distance_DE | Country | Region | Type | Location | Point_ID | GW | GWpr | SW | SWpr | H | Hpr | Syntopy_all | Syntopy_GW | Syntopy_SW | Allotopy_GW | Allotopy_SW | CO | CF | OF | SMHD | C_C_area | Infstr_area | Green_C | Green_O | Green_F | Dense_canopy | Loose_canopy | Con_trees | Forest_patch | Urban_Park | Graveyard | Alleys | Midestate_wood | Orchard | Riparian_wood | B_S_Sporadic | B_S_Dense | B_T_Sporadic | B_T_Dense | Other_Infrastructure | GrW | St |
|---|---|---|---|---|---|---|---|---|---|---|---|---|---|---|---|---|---|---|---|---|---|---|---|---|---|---|---|---|---|---|---|---|---|---|---|---|---|---|---|---|---|---|---|---|---|---|
| 50.0585 | 19.93384 | MLP_U_0 | 205 | 201 | Poland | Northern | U | MLP | mlpu0_1 | 0 | 0 | 0 | 0 | 0 | 0 | 0 | | | 0 | 0 | NA | NA | NA | | 31093 | 26000 | 31093 | 0 | 0 | 1 | 0 | 0 | 0 | 1 | 0 | 0 | 1 | 0 | 0 | 0 | 0 | 0 | 1 | 0 | 0 | 0 |
| 50.06167 | 19.93249 | MLP_U_0 | 207 | 201 | Poland | Northern | U | MLP | mlpu0_2 | 0 | 0 | 0 | 0 | 0 | 0 | 0 | | | 0 | 0 | NA | NA | NA | | 19551 | 44600 | 19551 | 0 | 0 | 1 | 0 | 0 | 0 | 1 | 0 | 0 | 0 | 0 | 0 | 0 | 0 | 0 | 1 | 0 | 0 | 0 |
| 50.06446 | 19.93483 | MLP_U_0 | 209 | 201 | Poland | Northern | U | MLP | mlpu0_3 | 0 | 0 | 0 | 0 | 0 | 0 | 0 | | | 0 | 0 | NA | NA | NA | | 24243 | 39600 | 24243 | 0 | 0 | 1 | 0 | 0 | 0 | 1 | 0 | 0 | 0 | 0 | 0 | 0 | 0 | 0 | 1 | 0 | 0 | 0 |
| 50.06552 | 19.93966 | MLP_U_0 | 214 | 201 | Poland | Northern | U | MLP | mlpu0_4 | 1 | 1 | 0 | 0 | 0 | 0 | 0 | 0 | | 1 | 0 | NA | NA | NA | | 25547 | 36100 | 25547 | 0 | 0 | 1 | 0 | 0 | 0 | 1 | 0 | 0 | 0 | 0 | 0 | 0 | 0 | 0 | 1 | 0 | 0 | 0 |
| 50.06416 | 19.94367 | MLP_U_0 | 213 | 201 | Poland | Northern | U | MLP | mlpu0_5 | 2 | 1 | 0 | 0 | 0 | 0 | 0 | 0 | | 1 | 0 | NA | NA | NA | | 23336 | 29500 | 23336 | 0 | 0 | 1 | 0 | 0 | 0 | 1 | 0 | 0 | 0 | 0 | 0 | 0 | 0 | 0 | 1 | 0 | 0 | 1 |
| 50.06131 | 19.9438 | MLP_U_0 | 210 | 201 | Poland | Northern | U | MLP | mlpu0_6 | 0 | 0 | 0 | 0 | 0 | 0 | 0 | | | 0 | 0 | NA | NA | NA | | 23398 | 38000 | 23398 | 0 | 0 | 1 | 0 | 0 | 0 | 1 | 0 | 0 | 1 | 0 | 0 | 0 | 0 | 0 | 1 | 0 | 0 | 0 |
| 50.05911 | 19.94064 | MLP_U_0 | 206 | 201 | Poland | Northern | U | MLP | mlpu0_7 | 0 | 0 | 0 | 0 | 1 | 1 | 0 | | | 0 | 0 | NA | NA | NA | | 24654 | 38500 | 24654 | 0 | 0 | 1 | 0 | 0 | 0 | 1 | 0 | 0 | 1 | 0 | 0 | 0 | 0 | 0 | 1 | 0 | 0 | 0 |
| 50.05638 | 19.93984 | MLP_U_0 | 205 | 201 | Poland | Northern | U | MLP | mlpu0_8 | 2 | 1 | 0 | 0 | 0 | 0 | 0 | 0 | | 1 | 0 | NA | NA | NA | | 16498 | 43600 | 16498 | 0 | 0 | 1 | 1 | 0 | 0 | 1 | 0 | 0 | 1 | 0 | 0 | 0 | 0 | 0 | 1 | 0 | 0 | 0 |
| 50.08494 | 19.92136 | MLP_U_10 | 227 | 198 | Poland | Northern | U | MLP | mlpu10_1 | 2 | 1 | 0 | 0 | 0 | 0 | 0 | 0 | | 1 | 0 | 1 | NA | NA | 1 | 37497 | 11200 | 19393 | 18104 | 0 | 1 | 1 | 1 | 0 | 1 | 0 | 0 | 1 | 1 | 0 | 0 | 1 | 1 | 0 | 0 | 0 | 1 |
| 50.08774 | 19.92114 | MLP_U_10 | 224 | 198 | Poland | Northern | U | MLP | mlpu10_2 | 0 | 0 | 0 | 0 | 0 | 0 | 0 | | | 0 | 0 | 1 | NA | NA | 1 | 33974 | 3500 | 19799 | 14175 | 0 | 0 | 1 | 0 | 0 | 1 | 0 | 0 | 1 | 1 | 0 | 0 | 1 | 0 | 0 | 0 | 0 | 1 |
| 50.08893 | 19.91919 | MLP_U_10 | 225 | 198 | Poland | Northern | U | MLP | mlpu10_3 | 0 | 0 | 0 | 0 | 0 | 0 | 0 | | | 0 | 0 | 14 | NA | NA | 14 | 25266 | 17300 | 25266 | 0 | 0 | 1 | 1 | 0 | 0 | 0 | 0 | 0 | 1 | 0 | 0 | 0 | 0 | 1 | 0 | 0 | 1 | 0 |
| 50.08823 | 19.91591 | MLP_U_10 | 225 | 198 | Poland | Northern | U | MLP | mlpu10_4 | 0 | 0 | 1 | 1 | 0 | 0 | 0 | | 0 | 0 | 1 | 189 | NA | NA | 189 | 15124 | 32300 | 15124 | 0 | 0 | 0 | 1 | 0 | 0 | 0 | 0 | 0 | 1 | 0 | 0 | 0 | 1 | 1 | 0 | 0 | 0 | 0 |
| 50.08686 | 19.91346 | MLP_U_10 | 223 | 198 | Poland | Northern | U | MLP | mlpu10_5 | 2 | 1 | 0 | 0 | 1 | 1 | 0 | 0 | | 1 | 0 | NA | NA | NA | | 15292 | 27400 | 15292 | 0 | 0 | 1 | 1 | 0 | 0 | 0 | 0 | 0 | 1 | 0 | 0 | 0 | 1 | 1 | 0 | 0 | 0 | 1 |
| 50.08646 | 19.91094 | MLP_U_10 | 223 | 198 | Poland | Northern | U | MLP | mlpu10_6 | 0 | 0 | 0 | 0 | 0 | 0 | 0 | | | 0 | 0 | NA | NA | NA | | 15711 | 29600 | 15711 | 0 | 0 | 0 | 1 | 0 | 0 | 0 | 0 | 0 | 1 | 0 | 0 | 0 | 1 | 1 | 0 | 0 | 0 | 0 |
| 50.08405 | 19.91134 | MLP_U_10 | 219 | 198 | Poland | Northern | U | MLP | mlpu10_7 | 0 | 0 | 0 | 0 | 0 | 0 | 0 | | | 0 | 0 | NA | NA | NA | | 7321 | 31900 | 7321 | 0 | 0 | 0 | 1 | 0 | 0 | 0 | 0 | 0 | 1 | 0 | 0 | 0 | 1 | 1 | 0 | 0 | 0 | 0 |
| 50.08284 | 19.91478 | MLP_U_10 | 216 | 198 | Poland | Northern | U | MLP | mlpu10_8 | 0 | 0 | 0 | 0 | 0 | 0 | 0 | | | 0 | 0 | NA | NA | NA | | 5099 | 23500 | 5099 | 0 | 0 | 0 | 1 | 0 | 0 | 0 | 0 | 0 | 1 | 0 | 0 | 0 | 1 | 1 | 0 | 0 | 0 | 0 |
| 50.04328 | 19.94062 | MLP_U_14 | 204 | 202 | Poland | Northern | U | MLP | mlpu14_1 | 0 | 0 | 0 | 0 | 1 | 1 | 0 | | | 0 | 0 | NA | 210 | NA | 210 | 22446 | 14900 | 22446 | 0 | 0 | 1 | 1 | 0 | 0 | 1 | 0 | 0 | 1 | 0 | 1 | 0 | 0 | 1 | 1 | 0 | 0 | 0 |
| 50.04131 | 19.9426 | MLP_U_14 | 204 | 202 | Poland | Northern | U | MLP | mlpu14_2 | 0 | 0 | 0 | 0 | 0 | 0 | 0 | | | 0 | 0 | 46 | 1 | 111 | 1 | 9637 | 47400 | 9637 | 0 | 0 | 0 | 1 | 0 | 0 | 0 | 0 | 0 | 1 | 0 | 0 | 0 | 0 | 1 | 1 | 0 | 0 | 0 |
| 50.0393 | 19.94391 | MLP_U_14 | 221 | 202 | Poland | Northern | U | MLP | mlpu14_3 | 0 | 0 | 0 | 0 | 0 | 0 | 0 | | | 0 | 0 | 150 | 1 | 255 | 1 | 32210 | 20700 | 13355 | 0 | 18856 | 1 | 1 | 0 | 0 | 1 | 0 | 0 | 1 | 0 | 0 | 0 | 0 | 0 | 1 | 0 | 0 | 0 |
| 50.04172 | 19.9471 | MLP_U_14 | 221 | 202 | Poland | Northern | U | MLP | mlpu14_4 | 2 | 1 | 0 | 0 | 0 | 0 | 0 | 0 | | 1 | 0 | 35 | 45 | NA | 35 | 42189 | 23000 | 34730 | 0 | 7459 | 1 | 0 | 0 | 0 | 1 | 0 | 0 | 1 | 0 | 0 | 0 | 0 | 0 | 1 | 0 | 0 | 0 |
| 50.04181 | 19.95045 | MLP_U_14 | 223 | 202 | Poland | Northern | U | MLP | mlpu14_5 | 0 | 0 | 0 | 0 | 0 | 0 | 0 | | | 0 | 0 | 1 | 1 | 30 | 1 | 43080 | 8000 | 41966 | 1113 | 0 | 1 | 1 | 0 | 0 | 1 | 0 | 0 | 1 | 0 | 0 | 0 | 0 | 0 | 1 | 0 | 0 | 0 |
| 50.04255 | 19.95408 | MLP_U_14 | 229 | 202 | Poland | Northern | U | MLP | mlpu14_6 | 0 | 0 | 0 | 0 | 0 | 0 | 0 | | | 0 | 0 | 10 | 20 | 21 | 10 | 28055 | 25400 | 16069 | 4327 | 7658 | 1 | 1 | 0 | 1 | 0 | 0 | 0 | 1 | 1 | 0 | 0 | 0 | 1 | 0 | 0 | 0 | 0 |
| 50.04372 | 19.95762 | MLP_U_14 | 210 | 202 | Poland | Northern | U | MLP | mlpu14_7 | 0 | 0 | 0 | 0 | 0 | 0 | 0 | | | 0 | 0 | 5 | 20 | 30 | 5 | 26012 | 26300 | 3688 | 0 | 22323 | 1 | 0 | 0 | 1 | 1 | 0 | 0 | 1 | 0 | 0 | 0 | 1 | 0 | 1 | 0 | 0 | 0 |
| 50.0447 | 19.96011 | MLP_U_14 | 202 | 202 | Poland | Northern | U | MLP | mlpu14_8 | 0 | 0 | 0 | 0 | 0 | 0 | 0 | | | 0 | 0 | 200 | 100 | NA | 100 | 7051 | 36500 | 7051 | 0 | 0 | 0 | 1 | 0 | 0 | 0 | 0 | 0 | 1 | 0 | 0 | 0 | 0 | 0 | 1 | 1 | 0 | 0 |
| 50.13706 | 19.62114 | MLP_S_11 | 289 | 177 | Poland | Northern | S | MLP | mlps11_1 | 0 | 0 | 0 | 0 | 0 | 0 | 0 | | | 0 | 0 | 13 | 11 | 7 | 7 | 24614 | 5496 | 11066 | 4614 | 8934 | 1 | 1 | 0 | 0 | 1 | 0 | 0 | 1 | 1 | 0 | 0 | 1 | 0 | 0 | 0 | 0 | 0 |
| 50.13515 | 19.61977 | MLP_S_11 | 289 | 177 | Poland | Northern | S | MLP | mlps11_2 | 2 | 1 | 0 | 0 | 0 | 0 | 0 | 0 | | 1 | 0 | 24 | 25 | 50 | 24 | 15290 | 7398 | 7979 | 2513 | 4798 | 1 | 1 | 1 | 0 | 1 | 0 | 0 | 1 | 1 | 0 | 0 | 1 | 0 | 0 | 0 | 0 | 1 |
| 50.13263 | 19.62091 | MLP_S_11 | 272 | 177 | Poland | Northern | S | MLP | mlps11_3 | 1 | 1 | 0 | 0 | 0 | 0 | 0 | 0 | | 1 | 0 | 106 | 37 | 41 | 37 | 16514 | 7202 | 3365 | 2058 | 11091 | 1 | 1 | 1 | 0 | 1 | 0 | 0 | 1 | 0 | 0 | 0 | 1 | 0 | 0 | 0 | 1 | 1 |
| 50.13273 | 19.62546 | MLP_S_11 | 267 | 177 | Poland | Northern | S | MLP | mlps11_4 | 0 | 0 | 0 | 0 | 0 | 0 | 0 | | | 0 | 0 | 31 | 1 | 107 | 1 | 24649 | 7144 | 24000 | 649 | 0 | 1 | 1 | 1 | 0 | 1 | 0 | 0 | 1 | 0 | 0 | 0 | 1 | 0 | 0 | 0 | 0 | 1 |
| 50.13427 | 19.6284 | MLP_S_11 | 262 | 177 | Poland | Northern | S | MLP | mlps11_5 | 1 | 1 | 0 | 0 | 0 | 0 | 0 | 0 | | 1 | 0 | 150 | 1 | 315 | 1 | 43001 | 5450 | 43001 | 0 | 0 | 1 | 1 | 0 | 0 | 1 | 0 | 0 | 1 | 0 | 0 | 0 | 0 | 0 | 1 | 0 | 0 | 1 |
| 50.13637 | 19.63057 | MLP_S_11 | 262 | 177 | Poland | Northern | S | MLP | mlps11_6 | 2 | 1 | 0 | 0 | 0 | 0 | 0 | 0 | | 1 | 0 | 224 | 173 | NA | 173 | 37056 | 11973 | 37056 | 0 | 0 | 1 | 0 | 0 | 0 | 1 | 0 | 0 | 1 | 0 | 0 | 0 | 1 | 0 | 1 | 0 | 0 | 0 |
| 50.1387 | 19.63187 | MLP_S_11 | 264 | 177 | Poland | Northern | S | MLP | mlps11_7 | 1 | 1 | 0 | 0 | 0 | 0 | 0 | 0 | | 1 | 0 | 67 | NA | NA | 67 | 26401 | 10060 | 25306 | 1095 | 0 | 1 | 0 | 0 | 0 | 1 | 0 | 0 | 1 | 0 | 0 | 0 | 1 | 0 | 0 | 0 | 0 | 0 |
| 50.14078 | 19.633 | MLP_S_11 | 266 | 177 | Poland | Northern | S | MLP | mlps11_8 | 2 | 1 | 0 | 0 | 0 | 0 | 0 | 0 | | 1 | 0 | 73 | NA | NA | 73 | 17116 | 6336 | 15506 | 1609 | 0 | 1 | 1 | 1 | 0 | 1 | 0 | 0 | 1 | 0 | 0 | 0 | 1 | 0 | 0 | 0 | 0 | 0 |
| 50.11096 | 19.80047 | MLP_R_11 | 255 | 190 | Poland | Northern | R | MLP | mlpr11_1 | 0 | 0 | 0 | 0 | 0 | 0 | 0 | | | 0 | 0 | 151 | 178 | NA | 151 | 9613 | 8165 | 9613 | 0 | 0 | 0 | 1 | 1 | 0 | 0 | 0 | 0 | 1 | 0 | 0 | 0 | 1 | 0 | 0 | 0 | 0 | 0 |
| 50.10871 | 19.79826 | MLP_R_11 | 270 | 190 | Poland | Northern | R | MLP | mlpr11_2 | 0 | 0 | 0 | 0 | 0 | 0 | 0 | | | 0 | 0 | 23 | 148 | 205 | 23 | 6420 | 4954 | 5790 | 630 | 0 | 0 | 1 | 1 | 0 | 0 | 0 | 0 | 1 | 1 | 0 | 0 | 1 | 0 | 0 | 0 | 0 | 1 |
| 50.10701 | 19.79839 | MLP_R_11 | 275 | 190 | Poland | Northern | R | MLP | mlpr11_3 | 1 | 1 | 0 | 0 | 0 | 0 | 0 | 0 | | 1 | 0 | 18 | 165 | 165 | 18 | 9689 | 4072 | 7784 | 1905 | 0 | 1 | 1 | 1 | 0 | 0 | 0 | 0 | 1 | 1 | 0 | 0 | 1 | 0 | 0 | 0 | 0 | 1 |
| 50.10711 | 19.80122 | MLP_R_11 | 269 | 190 | Poland | Northern | R | MLP | mlpr11_4 | 1 | 1 | 0 | 0 | 0 | 0 | 0 | 0 | | 1 | 0 | 41 | 142 | 258 | 41 | 9671 | 5205 | 8230 | 1441 | 0 | 1 | 1 | 1 | 0 | 0 | 0 | 0 | 1 | 1 | 0 | 0 | 1 | 0 | 0 | 0 | 0 | 0 |
| 50.1054 | 19.80322 | MLP_R_11 | 254 | 190 | Poland | Northern | R | MLP | mlpr11_5 | 0 | 0 | 0 | 0 | 0 | 0 | 0 | | | 0 | 0 | 1 | 143 | 63 | 1 | 12487 | 4468 | 3788 | 8699 | 0 | 0 | 1 | 1 | 0 | 0 | 0 | 0 | 1 | 1 | 0 | 0 | 1 | 0 | 0 | 0 | 0 | 0 |
| 50.10625 | 19.80549 | MLP_R_11 | 270 | 190 | Poland | Northern | R | MLP | mlpr11_6 | 1 | 1 | 0 | 0 | 0 | 0 | 0 | 0 | | 1 | 0 | 104 | 263 | 296 | 104 | 11504 | 5460 | 10778 | 726 | 0 | 0 | 1 | 1 | 0 | 0 | 0 | 0 | 1 | 1 | 0 | 0 | 1 | 0 | 0 | 0 | 0 | 0 |
| 50.10799 | 19.80763 | MLP_R_11 | 263 | 190 | Poland | Northern | R | MLP | mlpr11_7 | 0 | 0 | 0 | 0 | 0 | 0 | 0 | | | 0 | 0 | 54 | 270 | 282 | 54 | 6862 | 4284 | 5984 | 878 | 0 | 0 | 1 | 1 | 0 | 0 | 0 | 0 | 1 | 0 | 0 | 0 | 1 | 0 | 0 | 0 | 0 | 1 |
| 50.11052 | 19.80915 | MLP_R_11 | 254 | 190 | Poland | Northern | R | MLP | mlpr11_8 | 0 | 0 | 0 | 0 | 0 | 0 | 0 | | | 0 | 0 | 87 | NA | NA | 87 | 8789 | 5086 | 8789 | 0 | 0 | 1 | 1 | 1 | 0 | 0 | 0 | 0 | 1 | 0 | 0 | 0 | 1 | 0 | 0 | 0 | 0 | 0 |
| 50.01648 | 19.92274 | MLP_U_9 | 222 | 202 | Poland | Northern | U | MLP | mlpu9_1 | 2 | 1 | 0 | 0 | 0 | 0 | 0 | 0 | | 1 | 0 | 14 | 55 | 238 | 14 | 41599 | 12900 | 40138 | 1461 | 0 | 1 | 1 | 1 | 0 | 1 | 0 | 0 | 1 | 1 | 0 | 0 | 1 | 1 | 0 | 0 | 0 | 0 |
| 50.01473 | 19.92247 | MLP_U_9 | 225 | 202 | Poland | Northern | U | MLP | mlpu9_2 | 1 | 1 | 0 | 0 | 0 | 0 | 0 | 0 | | 1 | 0 | 114 | 10 | NA | 10 | 50030 | 3400 | 50030 | 0 | 0 | 1 | 1 | 0 | 1 | 1 | 0 | 0 | 1 | 0 | 0 | 0 | 0 | 0 | 0 | 0 | 0 | 0 |
| 50.01293 | 19.91948 | MLP_U_9 | 230 | 202 | Poland | Northern | U | MLP | mlpu9_3 | 2 | 1 | 0 | 0 | 0 | 0 | 0 | 0 | | 1 | 0 | NA | 19 | NA | 19 | 44269 | 7700 | 7456 | 0 | 36812 | 1 | 1 | 1 | 1 | 0 | 0 | 0 | 1 | 0 | 0 | 0 | 0 | 1 | 0 | 0 | 0 | 0 |
| 50.01191 | 19.91577 | MLP_U_9 | 235 | 202 | Poland | Northern | U | MLP | mlpu9_4 | 2 | 1 | 0 | 0 | 0 | 0 | 0 | 0 | | 1 | 0 | NA | 26 | NA | 26 | 58870 | 6000 | 1651 | 0 | 57219 | 1 | 1 | 1 | 1 | 0 | 1 | 0 | 1 | 0 | 0 | 0 | 0 | 1 | 0 | 0 | 0 | 0 |
| 50.01271 | 19.91228 | MLP_U_9 | 245 | 202 | Poland | Northern | U | MLP | mlpu9_5 | 2 | 1 | 0 | 0 | 0 | 0 | 0 | 0 | | 1 | 0 | NA | 113 | NA | 113 | 31664 | 15000 | 2068 | 0 | 29596 | 1 | 1 | 1 | 1 | 0 | 0 | 0 | 1 | 0 | 0 | 0 | 0 | 1 | 0 | 0 | 0 | 0 |
| 50.01451 | 19.90912 | MLP_U_9 | 235 | 202 | Poland | Northern | U | MLP | mlpu9_6 | 0 | 0 | 0 | 0 | 0 | 0 | 0 | | | 0 | 0 | NA | NA | NA | 0 | 6605 | 24000 | 6605 | 0 | 0 | 1 | 1 | 0 | 0 | 0 | 0 | 1 | 1 | 1 | 0 | 0 | 1 | 1 | 0 | 0 | 0 | 0 |
| 50.01634 | 19.90593 | MLP_U_9 | 225 | 202 | Poland | Northern | U | MLP | mlpu9_7 | 0 | 0 | 0 | 0 | 0 | 0 | 0 | | | 0 | 0 | NA | 193 | NA | 193 | 17208 | 19400 | 17208 | 0 | 0 | 0 | 1 | 1 | 0 | 0 | 0 | 1 | 1 | 1 | 0 | 0 | 1 | 1 | 0 | 0 | 0 | 1 |
| 50.0176 | 19.90211 | MLP_U_9 | 229 | 202 | Poland | Northern | U | MLP | mlpu9_8 | 1 | 1 | 0 | 0 | 0 | 0 | 0 | 0 | | 1 | 0 | NA | 16 | NA | 16 | 38319 | 6700 | 38319 | 0 | 0 | 1 | 1 | 0 | 1 | 1 | 0 | 1 | 0 | 0 | 1 | 0 | 0 | 0 | 0 | 1 | 1 | 1 |
| 50.0665 | 19.99409 | MLP_U_11 | 200 | 204 | Poland | Northern | U | MLP | mlpu11_1 | 2 | 1 | 0 | 0 | 1 | 1 | 0 | 0 | | 1 | 0 | 10 | NA | NA | 10 | 20525 | 7100 | 19482 | 1042 | 0 | 0 | 1 | 0 | 0 | 1 | 0 | 1 | 1 | 1 | 0 | 0 | 1 | 1 | 0 | 0 | 0 | 1 |
| 50.06789 | 19.99583 | MLP_U_11 | 200 | 204 | Poland | Northern | U | MLP | mlpu11_2 | 2 | 1 | 0 | 0 | 0 | 0 | 0 | 0 | | 1 | 0 | 140 | NA | NA | 140 | 16395 | 100 | 16395 | 0 | 0 | 0 | 1 | 0 | 0 | 1 | 0 | 0 | 0 | 0 | 0 | 0 | 0 | 0 | 0 | 0 | 0 | 1 |
| 50.07033 | 19.99529 | MLP_U_11 | 206 | 204 | Poland | Northern | U | MLP | mlpu11_3 | 1 | 1 | 0 | 0 | 1 | 1 | 0 | 0 | | 1 | 0 | NA | NA | | 0 | 55873 | 0 | 55873 | 0 | 0 | 1 | 1 | 1 | 0 | 1 | 0 | 0 | 0 | 0 | 0 | 0 | 0 | 0 | 0 | 0 | 1 | 1 |
| 50.07313 | 19.99534 | MLP_U_11 | 208 | 204 | Poland | Northern | U | MLP | mlpu11_4 | 1 | 1 | 0 | 0 | 1 | 1 | 0 | 0 | | 1 | 0 | NA | NA | | 0 | 36081 | 4000 | 36081 | 0 | 0 | 1 | 1 | 1 | 0 | 1 | 0 | 1 | 1 | 1 | 0 | 0 | 0 | 1 | 0 | 0 | 0 | 1 |
| 50.07471 | 19.99246 | MLP_U_11 | 214 | 204 | Poland | Northern | U | MLP | mlpu11_5 | 2 | 1 | 0 | 0 | 0 | 0 | 0 | 0 | | 1 | 0 | NA | NA | NA | 0 | 34296 | 13700 | 34296 | 0 | 0 | 1 | 1 | 0 | 1 | 1 | 0 | 0 | 1 | 0 | 0 | 0 | 0 | 1 | 0 | 1 | 0 | 1 |
| 50.07442 | 19.98898 | MLP_U_11 | 211 | 204 | Poland | Northern | U | MLP | mlpu11_6 | 0 | 0 | 0 | 0 | 0 | 0 | 0 | | | 0 | 0 | NA | NA | NA | 0 | 38624 | 9600 | 38624 | 0 | 0 | 1 | 1 | 1 | 0 | 1 | 0 | 1 | 1 | 0 | 0 | 0 | 0 | 1 | 0 | 1 | 0 | 1 |
| 50.0723 | 19.99098 | MLP_U_11 | 208 | 204 | Poland | Northern | U | MLP | mlpu11_7 | 2 | 1 | 0 | 0 | 0 | 0 | 0 | 0 | | 1 | 0 | 150 | 69 | NA | 69 | 61621 | 300 | 61621 | 0 | 0 | 1 | 0 | 1 | 0 | 1 | 0 | 0 | 0 | 0 | 0 | 0 | 0 | 0 | 0 | 0 | 0 | 1 |
| 50.07022 | 19.98961 | MLP_U_11 | 201 | 204 | Poland | Northern | U | MLP | mlpu11_8 | 0 | 0 | 0 | 0 | 0 | 0 | 0 | | | 0 | 0 | 82 | 1 | 73 | 1 | 69644 | 200 | 60005 | 0 | 9639 | 1 | 1 | 1 | 0 | 1 | 0 | 0 | 0 | 0 | 0 | 1 | 0 | 0 | 0 | 0 | 0 | 1 |
| 50.045 | 19.99165 | MLP_U_12 | 200 | 205 | Poland | Northern | U | MLP | mlpu12_1 | 2 | 1 | 0 | 0 | 0 | 0 | 0 | 0 | | 1 | 0 | 10 | 46 | 85 | 10 | 19417 | 18200 | 11440 | 7111 | 866 | 0 | 1 | 0 | 0 | 1 | 0 | 0 | 1 | 1 | 0 | 1 | 0 | 1 | 0 | 0 | 0 | 1 |
| 50.04683 | 19.99336 | MLP_U_12 | 199 | 205 | Poland | Northern | U | MLP | mlpu12_2 | 2 | 1 | 0 | 0 | 0 | 0 | 0 | 0 | | 1 | 0 | 62 | NA | NA | 62 | 34815 | 10800 | 34815 | 0 | 0 | 1 | 1 | 0 | 0 | 1 | 0 | 0 | 1 | 0 | 0 | 1 | 0 | 1 | 0 | 0 | 1 | 1 |
| 50.04929 | 19.99173 | MLP_U_12 | 200 | 205 | Poland | Northern | U | MLP | mlpu12_3 | 1 | 1 | 0 | 0 | 1 | 1 | 0 | 0 | | 1 | 0 | NA | 250 | | 250 | 32731 | 9500 | 32731 | 0 | 0 | 1 | 1 | 0 | 0 | 1 | 0 | 0 | 1 | 0 | 0 | 1 | 0 | 1 | 0 | 0 | 0 | 1 |
| 50.05113 | 19.99395 | MLP_U_12 | 200 | 205 | Poland | Northern | U | MLP | mlpu12_4 | 1 | 1 | 0 | 0 | 0 | 0 | 0 | 0 | | 1 | 0 | NA | 42 | NA | 42 | 20741 | 3700 | 9766 | 0 | 10975 | 1 | 1 | 0 | 1 | 1 | 0 | 0 | 1 | 0 | 0 | 0 | 0 | 1 | 0 | 0 | 0 | 1 |
| 50.04987 | 19.99691 | MLP_U_12 | 200 | 205 | Poland | Northern | U | MLP | mlpu12_5 | 1 | 1 | 0 | 0 | 0 | 0 | 0 | 0 | | 1 | 0 | NA | 88 | NA | 88 | 16186 | 0 | 3241 | 0 | 12945 | 0 | 1 | 0 | 1 | 1 | 0 | 0 | 1 | 0 | 0 | 0 | 0 | 0 | 0 | 0 | 0 | 1 |
| 50.0479 | 20.00006 | MLP_U_12 | 201 | 205 | Poland | Northern | U | MLP | mlpu12_6 | 0 | 0 | 0 | 0 | 0 | 0 | 0 | | | 0 | 0 | NA | NA | 83 | 83 | 11862 | 0 | 0 | 0 | 11862 | 0 | 1 | 0 | 1 | 0 | 0 | 0 | 0 | 0 | 0 | 0 | 0 | 0 | 0 | 0 | 1 | 1 |
| 50.0461 | 20.00325 | MLP_U_12 | 201 | 205 | Poland | Northern | U | MLP | mlpu12_7 | 2 | 1 | 0 | 0 | 0 | 0 | 0 | 0 | | 1 | 0 | 150 | 126 | 20 | 20 | 27923 | 2500 | 0 | 16571 | 11353 | 0 | 1 | 0 | 1 | 0 | 0 | 0 | 0 | 1 | 0 | 1 | 0 | 0 | 0 | 0 | 0 | 1 |
| 50.04347 | 20.00281 | MLP_U_12 | 198 | 205 | Poland | Northern | U | MLP | mlpu12_8 | 1 | 1 | 0 | 0 | 0 | 0 | 0 | 0 | | 1 | 0 | 96 | 59 | 93 | 59 | 22455 | 4600 | 10169 | 12287 | 0 | 1 | 1 | 0 | 0 | 1 | 0 | 0 | 1 | 1 | 0 | 1 | 0 | 0 | 0 | 0 | 0 | 1 |
| 49.86879 | 19.68298 | MLP_S_12 | 321 | 195 | Poland | Northern | S | MLP | mlps12_1 | 2 | 1 | 0 | 0 | 0 | 0 | 0 | 0 | | 1 | 0 | NA | 120 | NA | 120 | 20550 | 8718 | 0 | 3469 | 17080 | 1 | 1 | 0 | 1 | 0 | 0 | 0 | 0 | 1 | 0 | 1 | 1 | 1 | 0 | 0 | 1 | 1 |
| 49.8685 | 19.68633 | MLP_S_12 | 317 | 195 | Poland | Northern | S | MLP | mlps12_2 | 1 | 1 | 0 | 0 | 0 | 0 | 0 | 0 | | 1 | 0 | NA | 48 | NA | 48 | 13578 | 9751 | 1414 | 2361 | 9803 | 1 | 1 | 0 | 1 | 0 | 0 | 0 | 1 | 1 | 0 | 0 | 1 | 1 | 0 | 0 | 0 | 1 |
| 49.86656 | 19.68921 | MLP_S_12 | 306 | 195 | Poland | Northern | S | MLP | mlps12_3 | 0 | 0 | 0 | 0 | 0 | 0 | 0 | | | 0 | 0 | NA | 125 | NA | 125 | 11190 | 7476 | 6493 | 1429 | 3268 | 0 | 1 | 0 | 0 | 0 | 0 | 1 | 0 | 1 | 0 | 0 | 1 | 0 | 0 | 0 | 0 | 0 |
| 49.86765 | 19.69116 | MLP_S_12 | 300 | 195 | Poland | Northern | S | MLP | mlps12_4 | 1 | 1 | 0 | 0 | 0 | 0 | 0 | 0 | | 1 | 0 | NA | 121 | NA | 121 | 33175 | 5043 | 4595 | 0 | 28580 | 1 | 1 | 0 | 1 | 0 | 0 | 1 | 0 | 1 | 0 | 0 | 1 | 0 | 0 | 0 | 0 | 1 |
| 49.86978 | 19.69031 | MLP_S_12 | 293 | 195 | Poland | Northern | S | MLP | mlps12_5 | 0 | 0 | 0 | 0 | 0 | 0 | 0 | | | 0 | 0 | 107 | 25 | 135 | 25 | 39860 | 6956 | 845 | 0 | 39015 | 1 | 0 | 0 | 1 | 0 | 0 | 0 | 0 | 1 | 0 | 0 | 1 | 0 | 0 | 1 | 0 | 0 |
| 49.87189 | 19.68893 | MLP_S_12 | 290 | 195 | Poland | Northern | S | MLP | mlps12_6 | 1 | 1 | 0 | 0 | 0 | 0 | 0 | 0 | | 1 | 0 | 107 | 25 | 135 | 25 | 14159 | 13846 | 5157 | 0 | 9002 | 1 | 1 | 0 | 1 | 0 | 0 | 0 | 1 | 1 | 0 | 0 | 1 | 0 | 0 | 1 | 0 | 0 |
| 49.87392 | 19.68736 | MLP_S_12 | 283 | 195 | Poland | Northern | S | MLP | mlps12_7 | 3 | 1 | 0 | 0 | 1 | 1 | 0 | 0 | | 1 | 0 | 92 | 53 | 80 | 53 | 22872 | 15774 | 3980 | 0 | 18892 | 1 | 0 | 0 | 1 | 0 | 0 | 0 | 1 | 0 | 0 | 1 | 1 | 0 | 0 | 1 | 0 | 1 |
| 49.87503 | 19.68396 | MLP_S_12 | 295 | 195 | Poland | Northern | S | MLP | mlps12_8 | 0 | 0 | 0 | 0 | 0 | 0 | 0 | | | 0 | 0 | 189 | 95 | 54 | 54 | 12415 | 12627 | 12415 | 0 | 0 | 0 | 1 | 0 | 0 | 0 | 0 | 0 | 1 | 1 | 0 | 0 | 1 | 0 | 0 | 0 | 0 | 1 |
| 50.1432 | 19.40027 | MLP_S_13 | 295 | 162 | Poland | Northern | S | MLP | mlps13_1 | 1 | 1 | 0 | 0 | 0 | 0 | 0 | 0 | | 1 | 0 | NA | 154 | NA | 154 | 38484 | 9100 | 38484 | 0 | 0 | 1 | 1 | 0 | 0 | 1 | 0 | 0 | 1 | 0 | 0 | 0 | 0 | 1 | 1 | 0 | 1 | 0 |
| 50.14232 | 19.39692 | MLP_S_13 | 286 | 162 | Poland | Northern | S | MLP | mlps13_2 | 0 | 0 | 0 | 0 | 0 | 0 | 0 | | | 0 | 0 | 132 | 131 | NA | 131 | 4590 | 11817 | 4590 | 0 | 0 | 1 | 1 | 0 | 0 | 0 | 0 | 0 | 1 | 0 | 0 | 0 | 0 | 1 | 1 | 0 | 0 | 0 |
| 50.14378 | 19.39304 | MLP_S_13 | 284 | 162 | Poland | Northern | S | MLP | mlps13_3 | 0 | 0 | 0 | 0 | 0 | 0 | 0 | | | 0 | 0 | 44 | 112 | 126 | 44 | 5505 | 11238 | 4139 | 1365 | 0 | 0 | 1 | 0 | 0 | 0 | 0 | 0 | 1 | 1 | 0 | 0 | 1 | 1 | 0 | 0 | 0 | 0 |
| 50.14481 | 19.39658 | MLP_S_13 | 292 | 162 | Poland | Northern | S | MLP | mlps13_4 | 0 | 0 | 0 | 0 | 0 | 0 | 0 | | | 0 | 0 | 35 | 104 | 155 | 35 | 8317 | 9197 | 4428 | 1445 | 2444 | 0 | 1 | 0 | 0 | 0 | 0 | 0 | 1 | 1 | 0 | 0 | 1 | 1 | 0 | 1 | 0 | 1 |
| 50.14624 | 19.3999 | MLP_S_13 | 303 | 162 | Poland | Northern | S | MLP | mlps13_5 | 0 | 0 | 0 | 0 | 0 | 0 | 0 | | | 0 | 0 | 143 | 118 | 109 | 109 | 7965 | 6020 | 4547 | 0 | 3417 | 1 | 1 | 0 | 1 | 0 | 1 | 1 | 1 | 1 | 0 | 0 | 1 | 0 | 0 | 1 | 0 | 0 |
| 50.14729 | 19.40261 | MLP_S_13 | 306 | 162 | Poland | Northern | S | MLP | mlps13_6 | 2 | 1 | 0 | 0 | 0 | 0 | 0 | 0 | | 1 | 0 | 60 | 80 | NA | 60 | 15365 | 2002 | 15365 | 0 | 0 | 0 | 1 | 0 | 0 | 0 | 1 | 0 | 1 | 0 | 0 | 0 | 1 | 0 | 0 | 0 | 0 | 1 |
| 50.14696 | 19.40561 | MLP_S_13 | 303 | 162 | Poland | Northern | S | MLP | mlps13_7 | 0 | 0 | 0 | 0 | 1 | 1 | 0 | | | 0 | 0 | 77 | 85 | | 77 | 10321 | 9986 | 10321 | 0 | 0 | 0 | 1 | 0 | 0 | 0 | 1 | 1 | 1 | 0 | 0 | 0 | 1 | 1 | 0 | 0 | 0 | 1 |
| 50.14581 | 19.40783 | MLP_S_13 | 299 | 162 | Poland | Northern | S | MLP | mlps13_8 | 1 | 1 | 0 | 0 | 0 | 0 | 0 | 0 | | 1 | 0 | NA | NA | NA | 0 | 8474 | 13031 | 8474 | 0 | 0 | 0 | 1 | 0 | 0 | 0 | 0 | 0 | 1 | 0 | 0 | 0 | 0 | 1 | 0 | 0 | 0 | 0 |
| 49.88334 | 19.67119 | MLP_R_12 | 293 | 194 | Poland | Northern | R | MLP | mlpr12_1 | 1 | 1 | 0 | 0 | 0 | 0 | 0 | 0 | | 1 | 0 | NA | 9 | NA | 9 | 37282 | 0 | 0 | 0 | 37282 | 1 | 1 | 0 | 1 | 0 | 0 | 1 | 0 | 0 | 0 | 0 | 0 | 0 | 0 | 0 | 0 | 1 |
| 49.88189 | 19.67363 | MLP_R_12 | 290 | 194 | Poland | Northern | R | MLP | mlpr12_2 | 0 | 0 | 0 | 0 | 0 | 0 | 0 | | | 0 | 0 | 274 | 106 | 185 | 106 | 7566 | 0 | 4133 | 0 | 3433 | 0 | 1 | 0 | 0 | 0 | 0 | 1 | 0 | 0 | 0 | 0 | 0 | 0 | 0 | 0 | 0 | 1 |
| 49.88069 | 19.67659 | MLP_R_12 | 280 | 194 | Poland | Northern | R | MLP | mlpr12_3 | 0 | 0 | 0 | 0 | 0 | 0 | 0 | | | 0 | 0 | 238 | 1 | 135 | 1 | 16672 | 3920 | 6338 | 0 | 10334 | 1 | 1 | 0 | 1 | 0 | 0 | 1 | 0 | 0 | 1 | 1 | 1 | 0 | 0 | 0 | 0 | 1 |
| 49.88272 | 19.67803 | MLP_R_12 | 272 | 194 | Poland | Northern | R | MLP | mlpr12_4 | 0 | 0 | 0 | 0 | 0 | 0 | 0 | | | 0 | 0 | 24 | 66 | 63 | 24 | 13995 | 4203 | 1043 | 1351 | 11600 | 1 | 1 | 0 | 1 | 0 | 0 | 1 | 1 | 1 | 0 | 1 | 0 | 1 | 0 | 0 | 0 | 1 |
| 49.88453 | 19.67891 | MLP_R_12 | 274 | 194 | Poland | Northern | R | MLP | mlpr12_5 | 0 | 0 | 0 | 0 | 0 | 0 | 0 | | | 0 | 0 | 96 | 28 | 56 | 28 | 17836 | 772 | 1135 | 700 | 16001 | 0 | 1 | 0 | 0 | 0 | 1 | 1 | 0 | 0 | 0 | 1 | 0 | 0 | 0 | 0 | 1 | 1 |
| 49.8866 | 19.67785 | MLP_R_12 | 269 | 194 | Poland | Northern | R | MLP | mlpr12_6 | 0 | 0 | 0 | 0 | 0 | 0 | 0 | | | 0 | 0 | 208 | 50 | 33 | 33 | 31237 | 1231 | 445 | 0 | 30793 | 1 | 1 | 0 | 1 | 0 | 0 | 1 | 0 | 1 | 0 | 0 | 1 | 0 | 0 | 0 | 1 | 0 |
| 49.88891 | 19.6761 | MLP_R_12 | 270 | 194 | Poland | Northern | R | MLP | mlpr12_7 | 2 | 1 | 0 | 0 | 0 | 0 | 0 | 0 | | 1 | 0 | 106 | 106 | 55 | 55 | 34496 | 1533 | 0 | 925 | 33571 | 1 | 1 | 0 | 1 | 0 | 0 | 1 | 0 | 0 | 0 | 0 | 1 | 0 | 0 | 0 | 0 | 1 |
| 49.89059 | 19.67382 | MLP_R_12 | 273 | 194 | Poland | Northern | R | MLP | mlpr12_8 | 0 | 0 | 0 | 0 | 0 | 0 | 0 | | | 0 | 0 | 137 | 53 | 131 | 53 | 18917 | 5240 | 680 | 0 | 18237 | 1 | 1 | 0 | 1 | 0 | 0 | 1 | 0 | 1 | 0 | 1 | 1 | 0 | 0 | 0 | 1 | 0 |
| 50.1721 | 19.40267 | MLP_R_13 | 310 | 161 | Poland | Northern | R | MLP | mlpr13_1 | 0 | 0 | 0 | 0 | 0 | 0 | 0 | | | 0 | 0 | 97 | 225 | 116 | 97 | 4878 | 5301 | 0 | 3270 | 1607 | 0 | 1 | 0 | 0 | 0 | 0 | 0 | 1 | 1 | 0 | 0 | 1 | 0 | 0 | 0 | 0 | 1 |
| 50.17409 | 19.40359 | MLP_R_13 | 317 | 161 | Poland | Northern | R | MLP | mlpr13_2 | 0 | 0 | 0 | 0 | 0 | 0 | 0 | | | 0 | 0 | 66 | 42 | 163 | 42 | 5366 | 9448 | 4880 | 486 | 0 | 0 | 1 | 0 | 0 | 0 | 0 | 0 | 1 | 1 | 0 | 0 | 1 | 0 | 0 | 0 | 0 | 1 |
| 50.17602 | 19.40516 | MLP_R_13 | 328 | 161 | Poland | Northern | R | MLP | mlpr13_3 | 0 | 0 | 0 | 0 | 0 | 0 | 0 | | | 0 | 0 | 59 | 37 | 17 | 17 | 6469 | 3976 | 0 | 1168 | 5301 | 0 | 1 | 0 | 1 | 0 | 0 | 0 | 0 | 1 | 0 | 1 | 0 | 0 | 0 | 0 | 0 | 1 |
| 50.17702 | 19.40134 | MLP_R_13 | 323 | 161 | Poland | Northern | R | MLP | mlpr13_4 | 0 | 0 | 0 | 0 | 0 | 0 | 0 | | | 0 | 0 | 168 | 134 | 45 | 45 | 3449 | 4791 | 0 | 2162 | 1286 | 0 | 1 | 0 | 0 | 0 | 0 | 0 | 1 | 1 | 0 | 1 | 1 | 0 | 0 | 0 | 0 | 1 |
| 50.1771 | 19.39735 | MLP_R_13 | 316 | 161 | Poland | Northern | R | MLP | mlpr13_5 | 0 | 0 | 0 | 0 | 0 | 0 | 0 | | | 0 | 0 | 53 | 40 | 66 | 40 | 18896 | 3134 | 4600 | 1053 | 13242 | 1 | 1 | 0 | 1 | 0 | 0 | 1 | 0 | 1 | 0 | 0 | 1 | 0 | 0 | 0 | 1 | 1 |
| 50.17643 | 19.39359 | MLP_R_13 | 313 | 161 | Poland | Northern | R | MLP | mlpr13_6 | 2 | 1 | 0 | 0 | 0 | 0 | 0 | 0 | | 1 | 0 | 38 | 18 | 106 | 18 | 22377 | 1031 | 8183 | 291 | 13904 | 1 | 1 | 0 | 1 | 0 | 0 | 1 | 1 | 1 | 0 | 1 | 0 | 0 | 0 | 1 | 0 | 1 |
| 50.17521 | 19.3958 | MLP_R_13 | 310 | 161 | Poland | Northern | R | MLP | mlpr13_7 | 1 | 1 | 0 | 0 | 0 | 0 | 0 | 0 | | 1 | 0 | 338 | 1 | 102 | 1 | 48667 | 340 | 0 | 0 | 48667 | 1 | 1 | 0 | 1 | 0 | 0 | 0 | 0 | 0 | 1 | 0 | 0 | 0 | 0 | 1 | 0 | 0 |
| 50.17356 | 19.39691 | MLP_R_13 | 307 | 161 | Poland | Northern | R | MLP | mlpr13_8 | 0 | 0 | 0 | 0 | 0 | 0 | 0 | | | 0 | 0 | 102 | 163 | 16 | 16 | 30243 | 4918 | 0 | 652 | 29591 | 1 | 1 | 0 | 1 | 0 | 0 | 0 | 0 | 0 | 1 | 0 | 0 | 0 | 0 | 0 | 0 | 0 |
| 49.9664 | 20.43476 | MLP_S_9 | 243 | 242 | Poland | Northern | S | MLP | mlps9_1 | 1 | 1 | 0 | 0 | 0 | 0 | 0 | 0 | | 1 | 0 | 30 | NA | NA | 30 | 15197 | 8936 | 9192 | 6005 | 0 | 0 | 1 | 0 | 0 | 0 | 0 | 0 | 1 | 1 | 0 | 0 | 1 | 0 | 1 | 0 | 0 | 1 |
| 49.96849 | 20.43394 | MLP_S_9 | 212 | 242 | Poland | Northern | S | MLP | mlps9_2 | 0 | 0 | 2 | 1 | 0 | 0 | 0 | | 0 | 0 | 1 | 55 | NA | NA | 55 | 14723 | 14423 | 12601 | 2122 | 0 | 0 | 1 | 1 | 0 | 0 | 0 | 0 | 1 | 0 | 0 | 0 | 0 | 1 | 1 | 0 | 0 | 0 |
| 49.96775 | 20.43093 | MLP_S_9 | 211 | 242 | Poland | Northern | S | MLP | mlps9_3 | 0 | 0 | 0 | 0 | 1 | 1 | 0 | | | 0 | 0 | 71 | NA | | 71 | 19896 | 20670 | 19896 | 0 | 0 | 1 | 1 | 0 | 0 | 1 | 0 | 0 | 1 | 0 | 0 | 0 | 0 | 0 | 1 | 0 | 1 | 0 |
| 49.96755 | 20.42792 | MLP_S_9 | 220 | 242 | Poland | Northern | S | MLP | mlps9_4 | 0 | 0 | 0 | 0 | 0 | 0 | 0 | | | 0 | 0 | 134 | NA | NA | 134 | 15008 | 21824 | 15008 | 0 | 0 | 1 | 1 | 0 | 0 | 1 | 0 | 0 | 1 | 0 | 0 | 0 | 0 | 1 | 1 | 0 | 0 | 0 |
| 49.96531 | 20.42694 | MLP_S_9 | 223 | 242 | Poland | Northern | S | MLP | mlps9_5 | 2 | 1 | 0 | 0 | 0 | 0 | 0 | 0 | | 1 | 0 | 51 | NA | NA | 51 | 42442 | 4189 | 42442 | 0 | 0 | 1 | 0 | 0 | 0 | 1 | 0 | 0 | 0 | 0 | 0 | 0 | 0 | 1 | 1 | 0 | 0 | 1 |
| 49.9637 | 20.42584 | MLP_S_9 | 246 | 242 | Poland | Northern | S | MLP | mlps9_6 | 2 | 1 | 0 | 0 | 0 | 0 | 0 | 0 | | 1 | 0 | 70 | NA | NA | 70 | 69836 | 10 | 69836 | 0 | 0 | 1 | 0 | 0 | 0 | 1 | 0 | 0 | 0 | 0 | 0 | 0 | 0 | 0 | 0 | 0 | 0 | 1 |
| 49.96236 | 20.42739 | MLP_S_9 | 267 | 242 | Poland | Northern | S | MLP | mlps9_7 | 2 | 1 | 0 | 0 | 0 | 0 | 0 | 0 | | 1 | 0 | 30 | NA | NA | 30 | 45941 | 3855 | 42893 | 3048 | 0 | 1 | 0 | 0 | 0 | 1 | 0 | 0 | 0 | 1 | 0 | 0 | 1 | 0 | 0 | 0 | 0 | 1 |
| 49.96204 | 20.43041 | MLP_S_9 | 265 | 242 | Poland | Northern | S | MLP | mlps9_8 | 1 | 1 | 0 | 0 | 0 | 0 | 0 | 0 | | 1 | 0 | 90 | NA | NA | 90 | 16740 | 6337 | 7403 | 9338 | 0 | 1 | 1 | 1 | 0 | 1 | 0 | 0 | 0 | 1 | 0 | 0 | 1 | 0 | 0 | 0 | 0 | 1 |
| 50.19764 | 20.28674 | MLP_S_10 | 204 | 219 | Poland | Northern | S | MLP | mlps10_1 | 0 | 0 | 0 | 0 | 0 | 0 | 0 | | | 0 | 0 | 131 | NA | NA | 131 | 3558 | 7278 | 0 | 3048 | 510 | 0 | 1 | 0 | 0 | 0 | 0 | 1 | 0 | 1 | 0 | 0 | 1 | 0 | 0 | 1 | 0 | 0 |
| 50.19552 | 20.28889 | MLP_S_10 | 204 | 219 | Poland | Northern | S | MLP | mlps10_2 | 2 | 1 | 2 | 1 | 2 | 1 | 1 | 1 | 1 | 0 | 0 | 83 | NA | | 83 | 25057 | 4388 | 25057 | 0 | 0 | 1 | 1 | 0 | 0 | 1 | 0 | 1 | 1 | 0 | 1 | 1 | 0 | 0 | 0 | 1 | 1 | 1 |
| 50.19523 | 20.29318 | MLP_S_10 | 204 | 219 | Poland | Northern | S | MLP | mlps10_3 | 2 | 1 | 0 | 0 | 0 | 0 | 0 | 0 | | 1 | 0 | 106 | 40 | 1 | 1 | 31193 | 487 | 15361 | 1076 | 14756 | 0 | 1 | 0 | 1 | 1 | 0 | 1 | 0 | 1 | 1 | 1 | 0 | 0 | 0 | 0 | 0 | 1 |
| 50.19297 | 20.29397 | MLP_S_10 | 210 | 219 | Poland | Northern | S | MLP | mlps10_4 | 2 | 1 | 0 | 0 | 0 | 0 | 0 | 0 | | 1 | 0 | 126 | 50 | 66 | 50 | 12131 | 15167 | 2795 | 1121 | 8215 | 1 | 1 | 0 | 1 | 0 | 0 | 0 | 1 | 1 | 1 | 0 | 1 | 1 | 1 | 0 | 0 | 0 |
| 50.19242 | 20.29788 | MLP_S_10 | 207 | 219 | Poland | Northern | S | MLP | mlps10_5 | 0 | 0 | 0 | 0 | 0 | 0 | 0 | | | 0 | 0 | 159 | 86 | 243 | 86 | 14070 | 8455 | 1853 | 0 | 12218 | 1 | 1 | 1 | 1 | 0 | 0 | 0 | 1 | 0 | 0 | 0 | 0 | 1 | 0 | 1 | 1 | 0 |
| 50.19438 | 20.29887 | MLP_S_10 | 203 | 219 | Poland | Northern | S | MLP | mlps10_6 | 0 | 0 | 1 | 1 | 0 | 0 | 0 | | 0 | 0 | 1 | 415 | 280 | 36 | 36 | 11017 | 1593 | 0 | 896 | 10121 | 0 | 1 | 0 | 1 | 1 | 0 | 0 | 1 | 1 | 1 | 0 | 1 | 0 | 0 | 1 | 0 | 1 |
| 50.1968 | 20.30085 | MLP_S_10 | 203 | 219 | Poland | Northern | S | MLP | mlps10_7 | 0 | 0 | 0 | 0 | 0 | 0 | 0 | | | 0 | 0 | 183 | 282 | NA | 183 | 3376 | 0 | 2651 | 0 | 725 | 0 | 1 | 0 | 0 | 0 | 0 | 1 | 0 | 0 | 0 | 0 | 0 | 0 | 0 | 1 | 0 | 0 |
| 50.19851 | 20.30337 | MLP_S_10 | 203 | 219 | Poland | Northern | S | MLP | mlps10_8 | 0 | 0 | 0 | 0 | 0 | 0 | 0 | | | 0 | 0 | 174 | NA | NA | 174 | 4159 | 4229 | 0 | 0 | 4159 | 0 | 1 | 0 | 0 | 0 | 0 | 1 | 0 | 0 | 0 | 0 | 0 | 0 | 0 | 0 | 0 | 0 |
| 49.97527 | 20.26274 | MLP_R_9 | 262 | 224 | Poland | Northern | R | MLP | mlpr9_1 | 0 | 0 | 0 | 0 | 0 | 0 | 0 | | | 0 | 0 | 59 | 110 | 284 | 59 | 16594 | 3551 | 14993 | 1601 | 0 | 0 | 1 | 0 | 0 | 0 | 0 | 1 | 0 | 1 | 0 | 0 | 1 | 0 | 0 | 0 | 0 | 1 |
| 49.97665 | 20.2596 | MLP_R_9 | 273 | 224 | Poland | Northern | R | MLP | mlpr9_2 | 0 | 0 | 0 | 0 | 0 | 0 | 0 | | | 0 | 0 | 39 | 131 | 103 | 39 | 5067 | 4823 | 4166 | 901 | 0 | 0 | 1 | 0 | 0 | 0 | 0 | 1 | 0 | 1 | 0 | 1 | 0 | 1 | 0 | 1 | 0 | 0 |
| 49.97375 | 20.26186 | MLP_R_9 | 262 | 224 | Poland | Northern | R | MLP | mlpr9_3 | 2 | 1 | 0 | 0 | 0 | 0 | 0 | 0 | | 1 | 0 | 64 | 103 | NA | 64 | 24507 | 2052 | 24507 | 0 | 0 | 1 | 1 | 0 | 0 | 1 | 0 | 1 | 0 | 0 | 0 | 1 | 0 | 0 | 0 | 0 | 0 | 1 |
| 49.97243 | 20.26448 | MLP_R_9 | 246 | 224 | Poland | Northern | R | MLP | mlpr9_4 | 0 | 0 | 0 | 0 | 0 | 0 | 0 | | | 0 | 0 | 155 | 204 | NA | 155 | 4453 | 4451 | 4453 | 0 | 0 | 0 | 1 | 0 | 0 | 0 | 0 | 1 | 0 | 0 | 0 | 1 | 0 | 0 | 0 | 1 | 0 | 0 |

| | | | | | | | | | | | | | | | | | | | | | | | | | | | | | | | | | | | | | | | | | | | | | | |
|---|---|---|---|---|---|---|---|---|---|---|---|---|---|---|---|---|---|---|---|---|---|---|---|---|---|---|---|---|---|---|---|---|---|---|---|---|---|---|---|---|---|---|---|---|---|---|
| 49.97299 | 20.26843 | MLP_R_9 | 237 | 224 | Poland | Northern | R | MLP | mlpr9_5 | 0 | 0 | 2 | 1 | 0 | 0 | 0 | | 0 | 0 | 1 | 32 | NA | NA | 32 | 11787 | 1981 | 11787 | 0 | 0 | 0 | 1 | 0 | 0 | 0 | 0 | 1 | 0 | 1 | 0 | 1 | 0 | 0 | 0 | 0 | 0 | 1 |
| 49.97265 | 20.27246 | MLP_R_9 | 228 | 224 | Poland | Northern | R | MLP | mlpr9_6 | 0 | 0 | 0 | 0 | 0 | 0 | 0 | | | 0 | 0 | 169 | NA | NA | 169 | 3513 | 1783 | 3513 | 0 | 0 | 0 | 1 | 0 | 0 | 0 | 0 | 1 | 0 | 1 | 0 | 0 | 1 | 0 | 0 | 0 | 0 | 0 |
| 49.96971 | 20.27134 | MLP_R_9 | 223 | 224 | Poland | Northern | R | MLP | mlpr9_7 | 0 | 0 | 0 | 0 | 0 | 0 | 0 | | | 0 | 0 | NA | NA | NA | 0 | 1815 | 0 | 1815 | 0 | 0 | 0 | 1 | 0 | 0 | 0 | 0 | 1 | 0 | 0 | 0 | 0 | 0 | 0 | 0 | 0 | 1 | 0 |
| 49.9686 | 20.2673 | MLP_R_9 | 225 | 224 | Poland | Northern | R | MLP | mlpr9_8 | 0 | 0 | 1 | 1 | 0 | 0 | 0 | | 0 | 0 | 1 | NA | 159 | NA | 159 | 2789 | 0 | 2789 | 0 | 0 | 0 | 1 | 0 | 0 | 0 | 0 | 1 | 0 | 0 | 0 | 0 | 0 | 0 | 0 | 0 | 0 | 0 |
| 50.19105 | 20.23196 | MLP_R_10 | 236 | 215 | Poland | Northern | R | MLP | mlpr10_1 | 0 | 0 | 0 | 0 | 0 | 0 | 0 | | | 0 | 0 | NA | NA | NA | 0 | 7152 | 0 | 0 | 0 | 7152 | 0 | 1 | 0 | 0 | 0 | 0 | 1 | 0 | 0 | 0 | 0 | 0 | 0 | 0 | 0 | 0 | 0 |
| 50.19206 | 20.23532 | MLP_R_10 | 226 | 215 | Poland | Northern | R | MLP | mlpr10_2 | 0 | 0 | 0 | 0 | 0 | 0 | 0 | | | 0 | 0 | NA | 204 | NA | 204 | 17676 | 0 | 0 | 0 | 17676 | 0 | 1 | 0 | 0 | 0 | 0 | 1 | 0 | 0 | 0 | 0 | 0 | 0 | 0 | 0 | 0 | 0 |
| 50.19429 | 20.23684 | MLP_R_10 | 218 | 215 | Poland | Northern | R | MLP | mlpr10_3 | 0 | 0 | 0 | 0 | 0 | 0 | 0 | | | 0 | 0 | 216 | 46 | 79 | 46 | 21038 | 2124 | 1822 | 564 | 18651 | 0 | 1 | 0 | 1 | 0 | 0 | 1 | 0 | 1 | 0 | 1 | 0 | 0 | 0 | 0 | 1 | 0 |
| 50.1959 | 20.23478 | MLP_R_10 | 220 | 215 | Poland | Northern | R | MLP | mlpr10_4 | 2 | 1 | 0 | 0 | 0 | 0 | 0 | 0 | | 1 | 0 | 124 | 13 | 116 | 13 | 36638 | 5794 | 13483 | 0 | 23155 | 1 | 1 | 0 | 1 | 0 | 0 | 1 | 1 | 0 | 0 | 1 | 0 | 1 | 0 | 0 | 1 | 1 |
| 50.19805 | 20.23536 | MLP_R_10 | 210 | 215 | Poland | Northern | R | MLP | mlpr10_5 | 0 | 0 | 1 | 1 | 0 | 0 | 0 | | 0 | 0 | 1 | 152 | 2 | 92 | 2 | 27810 | 0 | 0 | 0 | 27810 | 1 | 1 | 0 | 1 | 0 | 0 | 1 | 0 | 0 | 1 | 1 | 0 | 0 | 0 | 0 | 0 | 1 |
| 50.20038 | 20.2344 | MLP_R_10 | 210 | 215 | Poland | Northern | R | MLP | mlpr10_6 | 0 | 0 | 0 | 0 | 0 | 0 | 0 | | | 0 | 0 | NA | NA | NA | 0 | 591 | 0 | 0 | 0 | 591 | 0 | 1 | 0 | 0 | 0 | 0 | 1 | 0 | 0 | 0 | 0 | 0 | 0 | 0 | 0 | 0 | 0 |
| 50.20288 | 20.23538 | MLP_R_10 | 211 | 215 | Poland | Northern | R | MLP | mlpr10_7 | 0 | 0 | 0 | 0 | 0 | 0 | 0 | | | 0 | 0 | 143 | 83 | NA | 83 | 3293 | 1741 | 0 | 2322 | 971 | 0 | 1 | 0 | 0 | 0 | 0 | 1 | 0 | 1 | 0 | 1 | 0 | 0 | 0 | 0 | 0 | 0 |
| 50.20388 | 20.23056 | MLP_R_10 | 213 | 215 | Poland | Northern | R | MLP | mlpr10_8 | 0 | 0 | 0 | 0 | 0 | 0 | 0 | | | 0 | 0 | 91 | 107 | 182 | 91 | 12803 | 3460 | 2131 | 1891 | 8782 | 1 | 1 | 0 | 1 | 0 | 0 | 1 | 1 | 1 | 0 | 1 | 1 | 0 | 0 | 0 | 0 | 0 |
| 50.01866 | 19.99565 | MLP_U_1 | 213 | 207 | Poland | Northern | U | MLP | mlpu1_1 | 2 | 1 | 0 | 0 | 2 | 1 | 0 | 0 | | 1 | 0 | 5 | 95 | | 5 | 46324 | 7800 | 46324 | 0 | 0 | 1 | 1 | 1 | 0 | 1 | 0 | 0 | 1 | 1 | 1 | 1 | 1 | 1 | 0 | 0 | 0 | 1 |
| 50.02052 | 19.99841 | MLP_U_1 | 208 | 207 | Poland | Northern | U | MLP | mlpu1_2 | 0 | 0 | 0 | 0 | 2 | 1 | 0 | | | 0 | 0 | 50 | NA | NA | 50 | 21426 | 13500 | 19204 | 2221 | 0 | 1 | 1 | 1 | 0 | 1 | 0 | 0 | 1 | 1 | 1 | 0 | 1 | 1 | 0 | 0 | 1 | 1 |
| 50.02111 | 20.00115 | MLP_U_1 | 207 | 207 | Poland | Northern | U | MLP | mlpu1_3 | 2 | 1 | 1 | 1 | 0 | 0 | 1 | 1 | 1 | 0 | 0 | 20 | NA | NA | 20 | 36118 | 7300 | 34869 | 1250 | 0 | 1 | 1 | 1 | 1 | 1 | 0 | 0 | 1 | 1 | 1 | 0 | 1 | 0 | 0 | 0 | 0 | 0 |
| 50.02032 | 20.00519 | MLP_U_1 | 208 | 207 | Poland | Northern | U | MLP | mlpu1_4 | 2 | 1 | 0 | 0 | 0 | 0 | 0 | 0 | | 1 | 0 | 115 | NA | NA | 115 | 38743 | 7500 | 38743 | 0 | 0 | 1 | 1 | 1 | 0 | 1 | 0 | 0 | 1 | 0 | 1 | 0 | 0 | 1 | 0 | 0 | 0 | 1 |
| 50.01923 | 20.00914 | MLP_U_1 | 213 | 207 | Poland | Northern | U | MLP | mlpu1_5 | 1 | 1 | 0 | 0 | 0 | 0 | 0 | 0 | | 1 | 0 | 180 | 50 | 200 | 50 | 39717 | 6600 | 9817 | 0 | 29900 | 1 | 1 | 0 | 1 | 0 | 0 | 1 | 1 | 0 | 0 | 1 | 0 | 1 | 0 | 0 | 1 | 0 |
| 50.01736 | 20.01233 | MLP_U_1 | 215 | 207 | Poland | Northern | U | MLP | mlpu1_6 | 1 | 1 | 0 | 0 | 0 | 0 | 0 | 0 | | 1 | 0 | NA | 80 | NA | 80 | 31153 | 16800 | 5403 | 0 | 25750 | 1 | 1 | 1 | 1 | 0 | 0 | 1 | 1 | 0 | 0 | 0 | 0 | 1 | 0 | 1 | 0 | 0 |
| 50.01484 | 20.01373 | MLP_U_1 | 222 | 207 | Poland | Northern | U | MLP | mlpu1_7 | 2 | 1 | 0 | 0 | 1 | 1 | 0 | 0 | | 1 | 0 | NA | 1 | NA | 1 | 38303 | 19400 | 28407 | 0 | 9896 | 1 | 1 | 1 | 1 | 1 | 0 | 1 | 1 | 0 | 1 | 0 | 0 | 1 | 0 | 0 | 0 | 1 |
| 50.01214 | 20.01426 | MLP_U_1 | 221 | 207 | Poland | Northern | U | MLP | mlpu1_8 | 2 | 1 | 0 | 0 | 2 | 1 | 0 | 0 | | 1 | 0 | NA | 1 | NA | 1 | 52502 | 10400 | 43330 | 0 | 9172 | 1 | 1 | 1 | 1 | 1 | 0 | 0 | 1 | 1 | 0 | 0 | 0 | 0 | 0 | 1 | 0 | 1 |
| 50.0309 | 19.99091 | MLP_U_2 | 200 | 206 | Poland | Northern | U | MLP | mlpu2_1 | 1 | 1 | 0 | 0 | 1 | 1 | 0 | 0 | | 1 | 0 | 170 | 70 | | 70 | 7915 | 22200 | 7915 | 0 | 0 | 0 | 1 | 0 | 0 | 0 | 0 | 1 | 0 | 0 | 1 | 0 | 0 | 0 | 0 | 1 | 0 | 0 |
| 50.03046 | 19.99468 | MLP_U_2 | 199 | 206 | Poland | Northern | U | MLP | mlpu2_2 | 0 | 0 | 0 | 0 | 0 | 0 | 0 | | | 0 | 0 | 214 | 70 | NA | 70 | 6245 | 24600 | 6245 | 0 | 0 | 0 | 1 | 0 | 0 | 0 | 0 | 1 | 0 | 0 | 1 | 0 | 0 | 0 | 0 | 1 | 1 | 1 |
| 50.03052 | 19.99945 | MLP_U_2 | 199 | 206 | Poland | Northern | U | MLP | mlpu2_3 | 0 | 0 | 0 | 0 | 0 | 0 | 0 | | | 0 | 0 | 133 | 155 | NA | 133 | 13461 | 24500 | 13461 | 0 | 0 | 0 | 1 | 0 | 0 | 0 | 0 | 1 | 1 | 0 | 1 | 0 | 0 | 0 | 0 | 1 | 0 | 0 |
| 50.0286 | 19.99908 | MLP_U_2 | 201 | 206 | Poland | Northern | U | MLP | mlpu2_4 | 0 | 0 | 0 | 0 | 0 | 0 | 0 | | | 0 | 0 | 100 | NA | 25 | 25 | 23558 | 22300 | 1110 | 0 | 22448 | 1 | 1 | 0 | 1 | 0 | 0 | 1 | 1 | 1 | 0 | 1 | 0 | 0 | 0 | 1 | 0 | 0 |
| 50.02749 | 19.9968 | MLP_U_2 | 203 | 206 | Poland | Northern | U | MLP | mlpu2_5 | 0 | 0 | 0 | 0 | 0 | 0 | 0 | | | 0 | 0 | 25 | 30 | 25 | 25 | 23713 | 13500 | 3930 | 3921 | 15861 | 1 | 1 | 0 | 1 | 0 | 0 | 1 | 1 | 1 | 0 | 1 | 1 | 0 | 0 | 1 | 1 | 0 |
| 50.02715 | 19.9934 | MLP_U_2 | 202 | 206 | Poland | Northern | U | MLP | mlpu2_6 | 0 | 0 | 1 | 1 | 0 | 0 | 0 | | 0 | 0 | 1 | 20 | 30 | 50 | 20 | 27610 | 11000 | 14129 | 1729 | 11752 | 0 | 1 | 0 | 1 | 0 | 0 | 1 | 1 | 1 | 0 | 0 | 1 | 1 | 0 | 1 | 0 | 0 |
| 50.02675 | 19.98903 | MLP_U_2 | 204 | 206 | Poland | Northern | U | MLP | mlpu2_7 | 0 | 0 | 2 | 1 | 0 | 0 | 0 | | 0 | 0 | 1 | 20 | 38 | 90 | 20 | 26530 | 17300 | 20050 | 6480 | 0 | 1 | 1 | 1 | 0 | 1 | 0 | 1 | 1 | 1 | 0 | 0 | 1 | 1 | 0 | 0 | 0 | 0 |
| 50.02639 | 19.98542 | MLP_U_2 | 206 | 206 | Poland | Northern | U | MLP | mlpu2_8 | 1 | 1 | 0 | 0 | 0 | 0 | 0 | 0 | | 1 | 0 | 20 | 10 | 96 | 10 | 17063 | 27400 | 10016 | 648 | 6399 | 1 | 1 | 0 | 1 | 0 | 0 | 0 | 1 | 1 | 0 | 0 | 1 | 0 | 1 | 1 | 0 | 1 |
| 49.98335 | 20.05949 | MLP_S_1 | 249 | 212 | Poland | Northern | S | MLP | mlps1_1 | 2 | 1 | 0 | 0 | 0 | 0 | 0 | 0 | | 1 | 0 | 227 | NA | NA | 227 | 19432 | 16371 | 19432 | 0 | 0 | 1 | 1 | 0 | 0 | 1 | 0 | 0 | 1 | 0 | 0 | 0 | 0 | 1 | 1 | 0 | 0 | 1 |
| 49.98242 | 20.05697 | MLP_S_1 | 249 | 212 | Poland | Northern | S | MLP | mlps1_2 | 0 | 0 | 0 | 0 | 0 | 0 | 0 | | | 0 | 0 | 88 | NA | NA | 88 | 14265 | 10010 | 13749 | 516 | 0 | 0 | 1 | 0 | 0 | 1 | 0 | 1 | 1 | 1 | 0 | 0 | 0 | 1 | 0 | 0 | 0 | 0 |
| 49.9798 | 20.05686 | MLP_S_1 | 278 | 212 | Poland | Northern | S | MLP | mlps1_3 | 0 | 0 | 2 | 1 | 0 | 0 | 0 | | 0 | 0 | 1 | 88 | NA | 139 | 88 | 36400 | 5997 | 0 | 36400 | 0 | 1 | 1 | 0 | 0 | 0 | 0 | 0 | 1 | 1 | 0 | 1 | 1 | 1 | 0 | 0 | 1 | 1 |
| 49.97719 | 20.05694 | MLP_S_1 | 310 | 212 | Poland | Northern | S | MLP | mlps1_4 | 1 | 1 | 0 | 0 | 0 | 0 | 0 | 0 | | 1 | 0 | NA | NA | 30 | 30 | 39008 | 3283 | 0 | 21430 | 17578 | 1 | 1 | 1 | 1 | 0 | 0 | 0 | 1 | 1 | 0 | 0 | 1 | 0 | 0 | 0 | 0 | 1 |
| 49.97621 | 20.05988 | MLP_S_1 | 314 | 212 | Poland | Northern | S | MLP | mlps1_5 | 2 | 1 | 0 | 0 | 1 | 1 | 0 | 0 | | 1 | 0 | 147 | 1 | 1 | 1 | 46752 | 2108 | 0 | 3830 | 42922 | 1 | 1 | 1 | 1 | 0 | 0 | 0 | 1 | 1 | 0 | 1 | 1 | 0 | 0 | 0 | 0 | 1 |
| 49.97669 | 20.06369 | MLP_S_1 | 312 | 212 | Poland | Northern | S | MLP | mlps1_6 | 2 | 1 | 0 | 0 | 0 | 0 | 0 | 0 | | 1 | 0 | 6 | 1 | 230 | 1 | 64194 | 39 | 61928 | 2266 | 0 | 1 | 0 | 1 | 1 | 1 | 0 | 0 | 0 | 1 | 0 | 0 | 1 | 0 | 0 | 0 | 0 | 1 |
| 49.97849 | 20.06655 | MLP_S_1 | 290 | 212 | Poland | Northern | S | MLP | mlps1_7 | 0 | 0 | 0 | 0 | 0 | 0 | 0 | | | 0 | 0 | 23 | NA | NA | 23 | 29390 | 4603 | 0 | 29390 | 0 | 1 | 1 | 0 | 0 | 1 | 0 | 0 | 1 | 1 | 0 | 1 | 1 | 0 | 0 | 0 | 0 | 1 |
| 49.97991 | 20.06374 | MLP_S_1 | 256 | 212 | Poland | Northern | S | MLP | mlps1_8 | 1 | 1 | 1 | 1 | 1 | 1 | 1 | 1 | 1 | 0 | 0 | 20 | 166 | 210 | 20 | 32758 | 7205 | 20485 | 12272 | 0 | 1 | 1 | 0 | 0 | 1 | 0 | 0 | 1 | 1 | 0 | 0 | 1 | 1 | 0 | 1 | 0 | 1 |
| 49.8847 | 20.09445 | MLP_S_2 | 243 | 220 | Poland | Northern | S | MLP | mlps2_1 | 0 | 0 | 0 | 0 | 0 | 0 | 0 | | | 0 | 0 | 200 | 10 | NA | 10 | 4648 | 11093 | 0 | 3595 | 1054 | 0 | 1 | 0 | 1 | 0 | 0 | 0 | 0 | 1 | 1 | 0 | 1 | 0 | 0 | 1 | 0 | 0 |
| 49.88479 | 20.09132 | MLP_S_2 | 242 | 220 | Poland | Northern | S | MLP | mlps2_2 | 2 | 1 | 0 | 0 | 0 | 0 | 0 | 0 | | 1 | 0 | 112 | 116 | 247 | 112 | 22467 | 5472 | 0 | 2507 | 19960 | 1 | 1 | 1 | 1 | 0 | 0 | 1 | 1 | 1 | 1 | 0 | 1 | 0 | 0 | 1 | 1 | 1 |
| 49.88321 | 20.0878 | MLP_S_2 | 244 | 220 | Poland | Northern | S | MLP | mlps2_3 | 0 | 0 | 0 | 0 | 0 | 0 | 0 | | | 0 | 0 | 135 | 28 | 257 | 28 | 9147 | 4725 | 0 | 3746 | 5401 | 0 | 1 | 1 | 0 | 1 | 0 | 1 | 1 | 1 | 1 | 0 | 1 | 1 | 0 | 0 | 0 | 1 |
| 49.88111 | 20.08596 | MLP_S_2 | 245 | 220 | Poland | Northern | S | MLP | mlps2_4 | 0 | 0 | 0 | 0 | 0 | 0 | 0 | | | 0 | 0 | NA | 1 | NA | 1 | 17985 | 1970 | 8842 | 0 | 9143 | 1 | 1 | 1 | 0 | 1 | 0 | 1 | 1 | 0 | 0 | 0 | 1 | 0 | 0 | 1 | 0 | 1 |
| 49.87883 | 20.08428 | MLP_S_2 | 246 | 220 | Poland | Northern | S | MLP | mlps2_5 | 1 | 1 | 0 | 0 | 0 | 0 | 0 | 0 | | 1 | 0 | NA | 1 | NA | 1 | 24543 | 778 | 4762 | 0 | 19781 | 1 | 1 | 1 | 1 | 0 | 0 | 1 | 1 | 0 | 0 | 0 | 0 | 1 | 0 | 0 | 0 | 1 |
| 49.87734 | 20.08679 | MLP_S_2 | 299 | 220 | Poland | Northern | S | MLP | mlps2_6 | 2 | 1 | 0 | 0 | 0 | 0 | 0 | 0 | | 1 | 0 | 300 | 77 | 11 | 11 | 35596 | 2160 | 116 | 5249 | 30231 | 1 | 1 | 1 | 1 | 0 | 0 | 0 | 1 | 1 | 1 | 1 | 0 | 1 | 0 | 0 | 1 | 1 |
| 49.87691 | 20.08939 | MLP_S_2 | 284 | 220 | Poland | Northern | S | MLP | mlps2_7 | 1 | 1 | 0 | 0 | 0 | 0 | 0 | 0 | | 1 | 0 | 120 | 68 | 11 | 11 | 29660 | 2612 | 0 | 5589 | 24072 | 1 | 1 | 1 | 1 | 0 | 1 | 0 | 1 | 1 | 1 | 0 | 1 | 0 | 0 | 0 | 0 | 1 |
| 49.87675 | 20.0934 | MLP_S_2 | 271 | 220 | Poland | Northern | S | MLP | mlps2_8 | 1 | 1 | 0 | 0 | 0 | 0 | 0 | 0 | | 1 | 0 | 164 | 36 | 132 | 36 | 23013 | 4312 | 0 | 4081 | 18932 | 1 | 1 | 1 | 1 | 0 | 0 | 0 | 1 | 1 | 0 | 0 | 1 | 0 | 0 | 0 | 0 | 1 |
| 50.03595 | 20.20838 | MLP_S_3 | 198 | 220 | Poland | Northern | S | MLP | mlps3_1 | 2 | 1 | 0 | 0 | 0 | 0 | 0 | 0 | | 1 | 0 | NA | NA | 1 | 1 | 11369 | 9262 | 0 | 4782 | 6587 | 1 | 1 | 1 | 1 | 0 | 0 | 0 | 0 | 1 | 1 | 0 | 1 | 1 | 0 | 0 | 1 | 1 |
| 50.03801 | 20.21066 | MLP_S_3 | 195 | 220 | Poland | Northern | S | MLP | mlps3_2 | 1 | 1 | 0 | 0 | 0 | 0 | 0 | 0 | | 1 | 0 | NA | NA | 1 | 1 | 17012 | 7230 | 0 | 7054 | 9958 | 0 | 1 | 1 | 1 | 0 | 0 | 0 | 1 | 1 | 1 | 0 | 1 | 1 | 0 | 0 | 0 | 0 |
| 50.03906 | 20.21362 | MLP_S_3 | 190 | 220 | Poland | Northern | S | MLP | mlps3_3 | 0 | 0 | 0 | 0 | 0 | 0 | 0 | | | 0 | 0 | NA | NA | 52 | 52 | 9320 | 7296 | 0 | 1106 | 8214 | 0 | 1 | 1 | 0 | 0 | 0 | 1 | 1 | 1 | 1 | 1 | 1 | 1 | 1 | 1 | 0 | 1 |
| 50.0401 | 20.21437 | MLP_S_3 | 193 | 220 | Poland | Northern | S | MLP | mlps3_4 | 0 | 0 | 0 | 0 | 0 | 0 | 0 | | | 0 | 0 | NA | NA | 18 | 18 | 10266 | 5551 | 0 | 1581 | 8685 | 1 | 1 | 1 | 1 | 0 | 0 | 1 | 0 | 1 | 1 | 1 | 1 | 0 | 0 | 1 | 0 | 1 |
| 50.0398 | 20.218 | MLP_S_3 | 192 | 220 | Poland | Northern | S | MLP | mlps3_5 | 0 | 0 | 0 | 0 | 0 | 0 | 0 | | | 0 | 0 | NA | NA | 34 | 34 | 9850 | 10348 | 0 | 3175 | 6676 | 1 | 1 | 1 | 0 | 0 | 0 | 0 | 0 | 1 | 0 | 0 | 1 | 1 | 0 | 1 | 0 | 0 |
| 50.03831 | 20.2184 | MLP_S_3 | 192 | 220 | Poland | Northern | S | MLP | mlps3_6 | 0 | 0 | 0 | 0 | 0 | 0 | 0 | | | 0 | 0 | NA | NA | 207 | 207 | 521 | 9540 | 0 | 521 | 0 | 0 | 1 | 0 | 0 | 0 | 0 | 0 | 0 | 1 | 0 | 0 | 1 | 1 | 0 | 1 | 0 | 0 |
| 50.03632 | 20.21915 | MLP_S_3 | 192 | 220 | Poland | Northern | S | MLP | mlps3_7 | 0 | 0 | 0 | 0 | 0 | 0 | 0 | | | 0 | 0 | 211 | 334 | NA | 211 | 463 | 10557 | 463 | 0 | 0 | 0 | 1 | 0 | 0 | 0 | 0 | 0 | 1 | 0 | 0 | 1 | 0 | 1 | 0 | 1 | 0 | 0 |
| 50.03442 | 20.21702 | MLP_S_3 | 200 | 220 | Poland | Northern | S | MLP | mlps3_8 | 0 | 0 | 0 | 0 | 0 | 0 | 0 | | | 0 | 0 | 62 | NA | NA | 62 | 10732 | 11425 | 10460 | 272 | 0 | 1 | 1 | 0 | 0 | 1 | 0 | 1 | 1 | 0 | 0 | 0 | 1 | 0 | 1 | 0 | 0 | 1 |
| 50.12779 | 20.37366 | MLP_S_4 | 193 | 228 | Poland | Northern | S | MLP | mlps4_1 | 0 | 0 | 1 | 1 | 0 | 0 | 0 | | 0 | 0 | 1 | 74 | 142 | 47 | 47 | 11784 | 7237 | 2833 | 5330 | 3621 | 0 | 1 | 0 | 0 | 0 | 1 | 0 | 1 | 1 | 0 | 0 | 1 | 0 | 0 | 0 | 0 | 1 |
| 50.12978 | 20.37572 | MLP_S_4 | 198 | 228 | Poland | Northern | S | MLP | mlps4_2 | 0 | 0 | 0 | 0 | 0 | 0 | 0 | | | 0 | 0 | 139 | 153 | 1 | 1 | 23843 | 8061 | 3898 | 2204 | 17742 | 1 | 1 | 1 | 1 | 0 | 0 | 0 | 1 | 1 | 0 | 0 | 1 | 1 | 0 | 0 | 0 | 1 |
| 50.13172 | 20.37869 | MLP_S_4 | 196 | 228 | Poland | Northern | S | MLP | mlps4_3 | 2 | 1 | 0 | 0 | 0 | 0 | 0 | 0 | | 1 | 0 | 160 | 200 | 112 | 112 | 21544 | 10542 | 0 | 2077 | 19467 | 1 | 1 | 0 | 1 | 0 | 0 | 0 | 0 | 1 | 0 | 0 | 1 | 0 | 0 | 0 | 0 | 1 |
| 50.13311 | 20.38255 | MLP_S_4 | 195 | 228 | Poland | Northern | S | MLP | mlps4_4 | 0 | 0 | 2 | 1 | 0 | 0 | 0 | | 0 | 0 | 1 | NA | NA | 20 | 20 | 14323 | 4910 | 0 | 3029 | 11294 | 1 | 1 | 0 | 1 | 0 | 0 | 0 | 1 | 1 | 0 | 0 | 1 | 1 | 0 | 0 | 1 | 0 |
| 50.13423 | 20.38637 | MLP_S_4 | 195 | 228 | Poland | Northern | S | MLP | mlps4_5 | 1 | 1 | 0 | 0 | 1 | 1 | 0 | 0 | | 1 | 0 | NA | NA | 45 | 45 | 13317 | 1845 | 0 | 3046 | 10271 | 1 | 1 | 0 | 1 | 0 | 0 | 1 | 0 | 1 | 0 | 1 | 1 | 1 | 0 | 0 | 1 | 1 |
| 50.1343 | 20.39053 | MLP_S_4 | 196 | 228 | Poland | Northern | S | MLP | mlps4_6 | 0 | 0 | 0 | 0 | 0 | 0 | 0 | | | 0 | 0 | NA | NA | 51 | 51 | 7353 | 956 | 0 | 649 | 6704 | 0 | 1 | 0 | 0 | 0 | 0 | 1 | 1 | 0 | 0 | 1 | 0 | 0 | 0 | 0 | 0 | 1 |
| 50.13306 | 20.3941 | MLP_S_4 | 195 | 228 | Poland | Northern | S | MLP | mlps4_7 | 0 | 0 | 0 | 0 | 0 | 0 | 0 | | | 0 | 0 | NA | NA | 154 | 154 | 1933 | 2469 | 0 | 1933 | 0 | 0 | 1 | 0 | 0 | 0 | 0 | 0 | 0 | 1 | 0 | 0 | 1 | 0 | 0 | 0 | 0 | 0 |
| 50.13084 | 20.39601 | MLP_S_4 | 195 | 228 | Poland | Northern | S | MLP | mlps4_8 | 1 | 1 | 0 | 0 | 0 | 0 | 0 | 0 | | 1 | 0 | NA | NA | 32 | 32 | 11238 | 4268 | 0 | 579 | 10658 | 1 | 1 | 0 | 1 | 0 | 0 | 1 | 1 | 1 | 0 | 0 | 1 | 0 | 1 | 0 | 1 | 1 |
| 50.24579 | 20.07224 | MLP_S_5 | 241 | 201 | Poland | Northern | S | MLP | mlps5_1 | 0 | 0 | 0 | 0 | 0 | 0 | 0 | | | 0 | 0 | 218 | 153 | 1 | 1 | 17094 | 3995 | 0 | 1186 | 15909 | 1 | 1 | 0 | 1 | 0 | 0 | 0 | 0 | 1 | 0 | 0 | 1 | 0 | 0 | 0 | 0 | 0 |
| 50.24571 | 20.06785 | MLP_S_5 | 245 | 201 | Poland | Northern | S | MLP | mlps5_2 | 0 | 0 | 0 | 0 | 0 | 0 | 0 | | | 0 | 0 | NA | NA | 23 | 23 | 10298 | 5016 | 0 | 1280 | 9018 | 0 | 1 | 0 | 1 | 0 | 0 | 0 | 0 | 1 | 0 | 0 | 1 | 0 | 0 | 0 | 0 | 1 |
| 50.24549 | 20.06396 | MLP_S_5 | 236 | 201 | Poland | Northern | S | MLP | mlps5_3 | 0 | 0 | 0 | 0 | 0 | 0 | 0 | | | 0 | 0 | NA | NA | 81 | 81 | 16583 | 380 | 0 | 0 | 16583 | 1 | 1 | 0 | 1 | 0 | 0 | 0 | 0 | 0 | 1 | 1 | 0 | 0 | 0 | 0 | 0 | 1 |
| 50.24304 | 20.06517 | MLP_S_5 | 235 | 201 | Poland | Northern | S | MLP | mlps5_4 | 0 | 0 | 0 | 0 | 0 | 0 | 0 | | | 0 | 0 | NA | NA | 22 | 22 | 23838 | 1215 | 0 | 942 | 22896 | 0 | 1 | 0 | 1 | 0 | 0 | 1 | 0 | 1 | 1 | 1 | 0 | 0 | 0 | 0 | 0 | 1 |
| 50.24132 | 20.06847 | MLP_S_5 | 237 | 201 | Poland | Northern | S | MLP | mlps5_5 | 2 | 1 | 0 | 0 | 0 | 0 | 0 | 0 | | 1 | 0 | NA | NA | 18 | 18 | 22597 | 2492 | 0 | 1569 | 21028 | 0 | 1 | 0 | 1 | 0 | 0 | 1 | 0 | 1 | 1 | 1 | 1 | 0 | 0 | 0 | 0 | 1 |
| 50.24082 | 20.07274 | MLP_S_5 | 234 | 201 | Poland | Northern | S | MLP | mlps5_6 | 1 | 1 | 0 | 0 | 0 | 0 | 0 | 0 | | 1 | 0 | NA | NA | 19 | 19 | 14121 | 3979 | 0 | 1393 | 12729 | 1 | 1 | 0 | 1 | 0 | 0 | 0 | 0 | 1 | 0 | 1 | 1 | 0 | 0 | 0 | 0 | 1 |
| 50.23996 | 20.07652 | MLP_S_5 | 233 | 201 | Poland | Northern | S | MLP | mlps5_7 | 0 | 0 | 0 | 0 | 0 | 0 | 0 | | | 0 | 0 | 35 | 84 | 15 | 15 | 14191 | 7302 | 0 | 6774 | 7417 | 0 | 1 | 0 | 1 | 0 | 0 | 0 | 0 | 0 | 0 | 1 | 1 | 0 | 0 | 0 | 0 | 1 |
| 50.24157 | 20.07773 | MLP_S_5 | 236 | 201 | Poland | Northern | S | MLP | mlps5_8 | 0 | 0 | 0 | 0 | 0 | 0 | 0 | | | 0 | 0 | 93 | 156 | 48 | 48 | 19847 | 7292 | 0 | 3179 | 16669 | 0 | 1 | 0 | 1 | 0 | 0 | 0 | 0 | 0 | 0 | 0 | 1 | 0 | 0 | 0 | 1 | 1 |
| 50.35291 | 20.03217 | MLP_S_6 | 315 | 195 | Poland | Northern | S | MLP | mlps6_1 | 0 | 0 | 0 | 0 | 0 | 0 | 0 | | | 0 | 0 | 33 | NA | NA | 33 | 5433 | 10047 | 1378 | 4055 | 0 | 0 | 1 | 0 | 0 | 0 | 0 | 0 | 1 | 1 | 0 | 0 | 1 | 1 | 0 | 0 | 0 | 0 |
| 50.35407 | 20.02848 | MLP_S_6 | 312 | 195 | Poland | Northern | S | MLP | mlps6_2 | 0 | 0 | 0 | 0 | 0 | 0 | 0 | | | 0 | 0 | 81 | 76 | 291 | 76 | 9203 | 13243 | 4164 | 0 | 5039 | 1 | 1 | 0 | 1 | 0 | 0 | 1 | 1 | 1 | 0 | 0 | 1 | 1 | 1 | 0 | 0 | 0 |
| 50.35614 | 20.02745 | MLP_S_6 | 300 | 195 | Poland | Northern | S | MLP | mlps6_3 | 0 | 0 | 0 | 0 | 0 | 0 | 0 | | | 0 | 0 | NA | 71 | NA | 71 | 4291 | 25636 | 4291 | 0 | 0 | 0 | 1 | 0 | 0 | 0 | 0 | 0 | 1 | 0 | 0 | 0 | 0 | 0 | 1 | 0 | 0 | 0 |
| 50.35812 | 20.0257 | MLP_S_6 | 291 | 195 | Poland | Northern | S | MLP | mlps6_4 | 0 | 0 | 0 | 0 | 0 | 0 | 0 | | | 0 | 0 | 51 | 1 | 2 | 1 | 17314 | 12087 | 8643 | 1438 | 7232 | 1 | 1 | 0 | 1 | 1 | 0 | 1 | 1 | 0 | 0 | 0 | 1 | 0 | 1 | 0 | 0 | 1 |
| 50.35957 | 20.02805 | MLP_S_6 | 286 | 195 | Poland | Northern | S | MLP | mlps6_5 | 0 | 0 | 0 | 0 | 0 | 0 | 0 | | | 0 | 0 | 24 | 45 | 1 | 1 | 20504 | 6526 | 0 | 4646 | 15858 | 1 | 1 | 1 | 1 | 0 | 0 | 1 | 1 | 0 | 0 | 0 | 1 | 0 | 0 | 1 | 1 | 1 |
| 50.35953 | 20.03187 | MLP_S_6 | 287 | 195 | Poland | Northern | S | MLP | mlps6_6 | 1 | 1 | 0 | 0 | 0 | 0 | 0 | 0 | | 1 | 0 | 32 | 106 | 72 | 32 | 15696 | 8211 | 811 | 9738 | 5147 | 1 | 1 | 1 | 0 | 0 | 0 | 1 | 1 | 1 | 0 | 0 | 1 | 0 | 0 | 1 | 0 | 1 |
| 50.35864 | 20.03493 | MLP_S_6 | 285 | 195 | Poland | Northern | S | MLP | mlps6_7 | 2 | 1 | 2 | 1 | 0 | 0 | 1 | 1 | 1 | 0 | 0 | 41 | 143 | NA | 41 | 14748 | 7087 | 14748 | 0 | 0 | 0 | 1 | 0 | 0 | 1 | 0 | 1 | 1 | 0 | 0 | 1 | 0 | 1 | 1 | 0 | 0 | 1 |
| 50.35744 | 20.03667 | MLP_S_6 | 282 | 195 | Poland | Northern | S | MLP | mlps6_8 | 2 | 1 | 0 | 0 | 0 | 0 | 0 | 0 | | 1 | 0 | 87 | 61 | NA | 61 | 20857 | 5075 | 20857 | 0 | 0 | 0 | 1 | 0 | 0 | 1 | 0 | 0 | 1 | 1 | 1 | 0 | 0 | 1 | 0 | 0 | 1 | 1 |
| 49.97128 | 19.83445 | MLP_S_7 | 236 | 198 | Poland | Northern | S | MLP | mlps7_1 | 2 | 1 | 0 | 0 | 0 | 0 | 0 | 0 | | 1 | 0 | 153 | NA | NA | 153 | 22269 | 18178 | 22269 | 0 | 0 | 0 | 1 | 0 | 0 | 0 | 0 | 1 | 1 | 0 | 0 | 0 | 0 | 1 | 0 | 0 | 0 | 0 |
| 49.97218 | 19.83126 | MLP_S_7 | 229 | 198 | Poland | Northern | S | MLP | mlps7_2 | 0 | 0 | 0 | 0 | 0 | 0 | 0 | | | 0 | 0 | NA | NA | NA | | 9741 | 17121 | 9741 | 0 | 0 | 0 | 1 | 0 | 0 | 0 | 0 | 1 | 1 | 0 | 0 | 0 | 1 | 1 | 0 | 0 | 0 | 0 |
| 49.97224 | 19.82772 | MLP_S_7 | 221 | 198 | Poland | Northern | S | MLP | mlps7_3 | 0 | 0 | 0 | 0 | 0 | 0 | 0 | | | 0 | 0 | NA | NA | NA | | 14589 | 15094 | 14589 | 0 | 0 | 1 | 1 | 0 | 0 | 0 | 0 | 0 | 1 | 0 | 0 | 0 | 1 | 1 | 0 | 1 | 0 | 0 |
| 49.97282 | 19.82478 | MLP_S_7 | 216 | 198 | Poland | Northern | S | MLP | mlps7_4 | 0 | 0 | 0 | 0 | 0 | 0 | 0 | | | 0 | 0 | NA | 105 | NA | 105 | 19918 | 12016 | 19918 | 0 | 0 | 1 | 1 | 1 | 0 | 1 | 0 | 1 | 1 | 0 | 0 | 0 | 1 | 1 | 0 | 0 | 0 | 0 |
| 49.97451 | 19.82259 | MLP_S_7 | 212 | 198 | Poland | Northern | S | MLP | mlps7_5 | 1 | 1 | 0 | 0 | 0 | 0 | 0 | 0 | | 1 | 0 | 38 | 60 | 235 | 38 | 38328 | 5901 | 38328 | 0 | 0 | 1 | 1 | 0 | 0 | 1 | 0 | 0 | 1 | 0 | 1 | 1 | 0 | 1 | 0 | 0 | 0 | 1 |
| 49.97516 | 19.81888 | MLP_S_7 | 210 | 198 | Poland | Northern | S | MLP | mlps7_6 | 0 | 0 | 0 | 0 | 0 | 0 | 0 | | | 0 | 0 | NA | 1 | NA | 1 | 21521 | 8595 | 18276 | 3245 | 0 | 1 | 1 | 0 | 0 | 1 | 0 | 1 | 1 | 0 | 1 | 0 | 1 | 1 | 0 | 1 | 1 | 1 |
| 49.97302 | 19.81916 | MLP_S_7 | 212 | 198 | Poland | Northern | S | MLP | mlps7_7 | 2 | 1 | 0 | 0 | 0 | 0 | 0 | 0 | | 1 | 0 | NA | 1 | NA | 1 | 43072 | 5 | 0 | 0 | 43072 | 1 | 1 | 0 | 1 | 1 | 0 | 0 | 0 | 0 | 1 | 0 | 0 | 0 | 0 | 0 | 0 | 1 |
| 49.97088 | 19.81729 | MLP_S_7 | 212 | 198 | Poland | Northern | S | MLP | mlps7_8 | 1 | 1 | 0 | 0 | 0 | 0 | 0 | 0 | | 1 | 0 | NA | 95 | 158 | 95 | 39148 | 0 | 0 | 0 | 39148 | 1 | 1 | 0 | 1 | 1 | 0 | 0 | 1 | 0 | 1 | 0 | 0 | 0 | 0 | 0 | 1 | 0 |
| 49.91677 | 20.16472 | MLP_R_1 | 278 | 223 | Poland | Northern | R | MLP | mlpr1_1 | 0 | 0 | 0 | 0 | 0 | 0 | 0 | | | 0 | 0 | 169 | 197 | 154 | 154 | 10260 | 2629 | 0 | 3034 | 7225 | 1 | 1 | 0 | 1 | 0 | 0 | 1 | 1 | 1 | 0 | 0 | 1 | 0 | 0 | 0 | 0 | 0 |
| 49.91814 | 20.16185 | MLP_R_1 | 292 | 223 | Poland | Northern | R | MLP | mlpr1_2 | 0 | 0 | 0 | 0 | 1 | 1 | 0 | | | 0 | 0 | 138 | 97 | 25 | 25 | 14144 | 1938 | 0 | 5398 | 8746 | 1 | 1 | 0 | 1 | 0 | 0 | 1 | 1 | 1 | 0 | 0 | 1 | 0 | 0 | 0 | 0 | 1 |
| 49.91912 | 20.15838 | MLP_R_1 | 296 | 223 | Poland | Northern | R | MLP | mlpr1_3 | 4 | 1 | 0 | 0 | 0 | 0 | 0 | 0 | | 1 | 0 | NA | NA | 81 | 81 | 33470 | 639 | 0 | 2494 | 30976 | 1 | 1 | 0 | 1 | 0 | 0 | 1 | 1 | 1 | 0 | 1 | 0 | 0 | 0 | 0 | 0 | 1 |
| 49.91803 | 20.15594 | MLP_R_1 | 309 | 223 | Poland | Northern | R | MLP | mlpr1_4 | 0 | 0 | 0 | 0 | 0 | 0 | 0 | | | 0 | 0 | NA | NA | 23 | 23 | 15125 | 1562 | 0 | 3334 | 11792 | 1 | 1 | 0 | 1 | 0 | 0 | 1 | 1 | 1 | 0 | 1 | 1 | 0 | 0 | 0 | 0 | 1 |
| 49.91549 | 20.15605 | MLP_R_1 | 311 | 223 | Poland | Northern | R | MLP | mlpr1_5 | 0 | 0 | 1 | 1 | 0 | 0 | 0 | | 0 | 0 | 1 | NA | NA | 162 | 162 | 17611 | 837 | 0 | 0 | 17611 | 1 | 1 | 0 | 1 | 0 | 0 | 1 | 1 | 1 | 0 | 1 | 0 | 0 | 0 | 0 | 0 | 1 |
| 49.91317 | 20.15626 | MLP_R_1 | 302 | 223 | Poland | Northern | R | MLP | mlpr1_6 | 1 | 1 | 1 | 1 | 0 | 0 | 1 | 1 | 1 | 0 | 0 | NA | NA | 57 | 57 | 25017 | 1276 | 0 | 2298 | 22720 | 1 | 1 | 0 | 1 | 0 | 0 | 1 | 1 | 1 | 0 | 1 | 1 | 0 | 0 | 0 | 0 | 1 |
| 49.91236 | 20.1604 | MLP_R_1 | 293 | 223 | Poland | Northern | R | MLP | mlpr1_7 | 0 | 0 | 0 | 0 | 0 | 0 | 0 | | | 0 | 0 | NA | NA | 162 | 162 | 19501 | 0 | 0 | 0 | 19501 | 1 | 1 | 0 | 1 | 0 | 0 | 1 | 0 | 0 | 0 | 0 | 0 | 0 | 0 | 0 | 0 | 0 |
| 49.91277 | 20.16438 | MLP_R_1 | 270 | 223 | Poland | Northern | R | MLP | mlpr1_8 | 0 | 0 | 0 | 0 | 0 | 0 | 0 | | | 0 | 0 | NA | NA | 32 | 32 | 16842 | 1252 | 0 | 7064 | 9779 | 1 | 1 | 0 | 1 | 0 | 0 | 1 | 0 | 1 | 0 | 1 | 0 | 0 | 0 | 0 | 0 | 0 |
| 49.85757 | 20.20881 | MLP_R_2 | 335 | 228 | Poland | Northern | R | MLP | mlpr2_1 | 0 | 0 | 0 | 0 | 0 | 0 | 0 | | | 0 | 0 | 370 | 186 | 44 | 44 | 7022 | 241 | 0 | 683 | 6340 | 0 | 1 | 0 | 0 | 0 | 0 | 1 | 1 | 1 | 0 | 1 | 0 | 0 | 0 | 0 | 0 | 1 |
| 49.85682 | 20.20514 | MLP_R_2 | 301 | 228 | Poland | Northern | R | MLP | mlpr2_2 | 0 | 0 | 2 | 1 | 0 | 0 | 0 | | 0 | 0 | 1 | NA | 75 | 93 | 75 | 23647 | 0 | 0 | 200 | 23447 | 1 | 1 | 0 | 1 | 0 | 0 | 1 | 0 | 1 | 0 | 0 | 0 | 0 | 0 | 0 | 0 | 1 |
| 49.85475 | 20.20558 | MLP_R_2 | 276 | 228 | Poland | Northern | R | MLP | mlpr2_3 | 0 | 0 | 0 | 0 | 0 | 0 | 0 | | | 0 | 0 | NA | 23 | NA | 23 | 21151 | 0 | 2066 | 508 | 18577 | 1 | 1 | 0 | 1 | 0 | 1 | 1 | 0 | 0 | 0 | 1 | 0 | 0 | 0 | 0 | 0 | 1 |
| 49.85322 | 20.20848 | MLP_R_2 | 273 | 228 | Poland | Northern | R | MLP | mlpr2_4 | 0 | 0 | 1 | 1 | 0 | 0 | 0 | | 0 | 0 | 1 | 279 | 3 | 118 | 3 | 22306 | 3792 | 1195 | 406 | 20705 | 1 | 1 | 0 | 1 | 0 | 0 | 1 | 1 | 0 | 0 | 1 | 1 | 1 | 0 | 0 | 0 | 1 |
| 49.85129 | 20.21089 | MLP_R_2 | 272 | 228 | Poland | Northern | R | MLP | mlpr2_5 | 2 | 1 | 0 | 0 | 0 | 0 | 0 | 0 | | 1 | 0 | 279 | 143 | 12 | 12 | 32201 | 1255 | 0 | 1948 | 30253 | 1 | 1 | 0 | 1 | 0 | 0 | 1 | 0 | 0 | 0 | 1 | 0 | 0 | 0 | 0 | 1 | 1 |
| 49.85189 | 20.21352 | MLP_R_2 | 298 | 228 | Poland | Northern | R | MLP | mlpr2_6 | 2 | 1 | 0 | 0 | 0 | 0 | 0 | 0 | | 1 | 0 | NA | NA | 5 | 5 | 30642 | 1730 | 0 | 7322 | 23320 | 1 | 1 | 0 | 1 | 0 | 0 | 1 | 0 | 1 | 0 | 1 | 0 | 0 | 0 | 0 | 0 | 1 |
| 49.85405 | 20.21492 | MLP_R_2 | 331 | 228 | Poland | Northern | R | MLP | mlpr2_7 | 0 | 0 | 0 | 0 | 0 | 0 | 0 | | | 0 | 0 | NA | NA | 17 | 17 | 21737 | 0 | 0 | 1998 | 19739 | 1 | 1 | 0 | 1 | 0 | 0 | 1 | 0 | 1 | 0 | 0 | 0 | 0 | 0 | 0 | 1 | 1 |
| 49.85622 | 20.21657 | MLP_R_2 | 351 | 228 | Poland | Northern | R | MLP | mlpr2_8 | 0 | 0 | 0 | 0 | 0 | 0 | 0 | | | 0 | 0 | NA | NA | 47 | 47 | 17895 | 461 | 0 | 327 | 17567 | 1 | 1 | 0 | 1 | 0 | 0 | 1 | 0 | 0 | 1 | 0 | 0 | 0 | 0 | 0 | 0 | 0 |
| 50.10513 | 20.30364 | MLP_R_3 | 190 | 223 | Poland | Northern | R | MLP | mlpr3_1 | 0 | 0 | 1 | 1 | 0 | 0 | 0 | | 0 | 0 | 1 | 71 | 162 | 96 | 71 | 6833 | 3193 | 0 | 936 | 5897 | 0 | 1 | 0 | 0 | 0 | 0 | 0 | 1 | 1 | 1 | 1 | 1 | 0 | 0 | 0 | 0 | 1 |
| 50.10309 | 20.302 | MLP_R_3 | 189 | 223 | Poland | Northern | R | MLP | mlpr3_2 | 1 | 1 | 0 | 0 | 0 | 0 | 0 | 0 | | 1 | 0 | NA | NA | 28 | 28 | 4958 | 2775 | 0 | 2068 | 2890 | 0 | 1 | 0 | 0 | 0 | 0 | 0 | 0 | 1 | 1 | 0 | 1 | 0 | 0 | 0 | 0 | 1 |
| 50.10103 | 20.30485 | MLP_R_3 | 188 | 223 | Poland | Northern | R | MLP | mlpr3_3 | 1 | 1 | 1 | 1 | 1 | 1 | 1 | 1 | 1 | 0 | 0 | NA | NA | 83 | 83 | 9774 | 1592 | 0 | 236 | 9538 | 0 | 1 | 0 | 0 | 0 | 0 | 0 | 0 | 0 | 1 | 0 | 1 | 0 | 0 | 0 | 0 | 0 |
| 50.10006 | 20.3093 | MLP_R_3 | 187 | 223 | Poland | Northern | R | MLP | mlpr3_4 | 0 | 0 | 0 | 0 | 0 | 0 | 0 | | | 0 | 0 | NA | NA | 71 | 71 | 5750 | 0 | 0 | 0 | 5750 | 0 | 1 | 0 | 1 | 0 | 0 | 0 | 0 | 0 | 1 | 0 | 0 | 0 | 0 | 0 | 0 | 0 |
| 50.10002 | 20.31421 | MLP_R_3 | 187 | 223 | Poland | Northern | R | MLP | mlpr3_5 | 0 | 0 | 2 | 1 | 0 | 0 | 0 | | 0 | 0 | 1 | NA | NA | 195 | 195 | 3169 | 0 | 0 | 0 | 3169 | 1 | 1 | 0 | 0 | 0 | 0 | 0 | 0 | 0 | 1 | 0 | 0 | 0 | 0 | 0 | 0 | 0 |
| 50.10177 | 20.31618 | MLP_R_3 | 187 | 223 | Poland | Northern | R | MLP | mlpr3_6 | 1 | 1 | 0 | 0 | 0 | 0 | 0 | 0 | | 1 | 0 | NA | NA | 152 | 152 | 6165 | 2031 | 0 | 757 | 5408 | 1 | 1 | 0 | 0 | 0 | 0 | 0 | 0 | 1 | 1 | 0 | 1 | 0 | 0 | 0 | 0 | 0 |
| 50.10377 | 20.3172 | MLP_R_3 | 199 | 223 | Poland | Northern | R | MLP | mlpr3_7 | 0 | 0 | 0 | 0 | 0 | 0 | 0 | | | 0 | 0 | NA | NA | 40 | 40 | 5451 | 4931 | 0 | 545 | 4905 | 0 | 1 | 0 | 1 | 0 | 0 | 0 | 0 | 1 | 1 | 0 | 1 | 0 | 0 | 0 | 0 | 1 |
| 50.10418 | 20.32084 | MLP_R_3 | 198 | 223 | Poland | Northern | R | MLP | mlpr3_8 | 2 | 1 | 0 | 0 | 0 | 0 | 0 | 0 | | 1 | 0 | NA | NA | 78 | 78 | 14024 | 4563 | 0 | 0 | 14024 | 1 | 1 | 0 | 0 | 0 | 0 | 0 | 1 | 0 | 1 | 0 | 1 | 0 | 0 | 0 | 0 | 1 |
| 50.15409 | 20.57799 | MLP_R_4 | 179 | 239 | Poland | Northern | R | MLP | mlpr4_1 | 1 | 1 | 1 | 1 | 3 | 1 | 1 | 1 | 1 | 0 | 0 | 312 | 161 | 62 | 62 | 14802 | 2082 | 0 | 671 | 14131 | 1 | 1 | 1 | 1 | 0 | 0 | 1 | 0 | 1 | 0 | 0 | 1 | 0 | 0 | 0 | 0 | 0 |
| 50.15147 | 20.57917 | MLP_R_4 | 184 | 239 | Poland | Northern | R | MLP | mlpr4_2 | 0 | 0 | 0 | 0 | 2 | 1 | 0 | | | 0 | 0 | 35 | 56 | 33 | 33 | 7168 | 2600 | 937 | 2133 | 4098 | 1 | 1 | 0 | 1 | 0 | 0 | 0 | 1 | 1 | 0 | 0 | 1 | 0 | 0 | 0 | 1 | 1 |
| 50.15081 | 20.5831 | MLP_R_4 | 182 | 239 | Poland | Northern | R | MLP | mlpr4_3 | 0 | 0 | 0 | 0 | 0 | 0 | 0 | | | 0 | 0 | NA | 179 | 20 | 20 | 32473 | 0 | 0 | 0 | 32473 | 1 | 1 | 0 | 1 | 0 | 0 | 1 | 0 | 0 | 0 | 0 | 0 | 0 | 0 | 0 | 0 | 1 |
| 50.14987 | 20.58653 | MLP_R_4 | 181 | 239 | Poland | Northern | R | MLP | mlpr4_4 | 1 | 1 | 0 | 0 | 0 | 0 | 0 | 0 | | 1 | 0 | NA | NA | 132 | 132 | 12216 | 0 | 0 | 0 | 12216 | 1 | 1 | 0 | 1 | 0 | 0 | 0 | 0 | 0 | 1 | 0 | 0 | 0 | 0 | 0 | 1 | 0 |
| 50.14771 | 20.58625 | MLP_R_4 | 181 | 239 | Poland | Northern | R | MLP | mlpr4_5 | 0 | 0 | 0 | 0 | 0 | 0 | 0 | | | 0 | 0 | NA | NA | 24 | 24 | 8857 | 274 | 0 | 469 | 8388 | 1 | 1 | 0 | 1 | 0 | 0 | 1 | 1 | 1 | 1 | 1 | 0 | 0 | 0 | 0 | 0 | 1 |
| 50.14831 | 20.58219 | MLP_R_4 | 181 | 239 | Poland | Northern | R | MLP | mlpr4_6 | 0 | 0 | 2 | 1 | 0 | 0 | 0 | | 0 | 0 | 1 | NA | NA | 19 | 19 | 26067 | 790 | 0 | 604 | 25463 | 1 | 0 | 0 | 1 | 0 | 0 | 0 | 0 | 1 | 0 | 1 | 1 | 0 | 0 | 0 | 0 | 0 |
| 50.14844 | 20.57796 | MLP_R_4 | 187 | 239 | Poland | Northern | R | MLP | mlpr4_7 | 0 | 0 | 0 | 0 | 0 | 0 | 0 | | | 0 | 0 | NA | NA | 44 | 44 | 30907 | 320 | 0 | 0 | 30907 | 1 | 0 | 0 | 1 | 0 | 0 | 0 | 0 | 1 | 0 | 1 | 0 | 0 | 0 | 0 | 0 | 1 |
| 50.14906 | 20.57414 | MLP_R_4 | 219 | 239 | Poland | Northern | R | MLP | mlpr4_8 | 2 | 1 | 0 | 0 | 0 | 0 | 0 | 0 | | 1 | 0 | NA | NA | NA | | 23330 | 0 | 0 | 0 | 23330 | 1 | 1 | 0 | 1 | 0 | 0 | 0 | 0 | 0 | 1 | 0 | 0 | 0 | 0 | 0 | 0 | 1 |
| 50.16575 | 20.0715 | MLP_R_5 | 281 | 206 | Poland | Northern | R | MLP | mlpr5_1 | 2 | 1 | 0 | 0 | 0 | 0 | 0 | 0 | | 1 | 0 | NA | NA | 195 | 195 | 23594 | 1697 | 0 | 0 | 23594 | 1 | 1 | 0 | 1 | 0 | 0 | 1 | 0 | 1 | 0 | 1 | 1 | 0 | 0 | 1 | 0 | 0 |
| 50.16471 | 20.07521 | MLP_R_5 | 266 | 206 | Poland | Northern | R | MLP | mlpr5_2 | 1 | 1 | 0 | 0 | 0 | 0 | 0 | 0 | | 1 | 0 | NA | NA | 39 | 39 | 18805 | 2582 | 0 | 409 | 18396 | 1 | 1 | 0 | 1 | 0 | 0 | 1 | 0 | 1 | 0 | 1 | 1 | 0 | 0 | 1 | 1 | 0 |
| 50.16608 | 20.07888 | MLP_R_5 | 284 | 206 | Poland | Northern | R | MLP | mlpr5_3 | 0 | 0 | 0 | 0 | 1 | 1 | 0 | | | 0 | 0 | NA | NA | 32 | 32 | 22784 | 3323 | 0 | 452 | 22332 | 1 | 1 | 0 | 1 | 0 | 0 | 1 | 0 | 1 | 0 | 0 | 1 | 0 | 0 | 0 | 0 | 0 |
| 50.16483 | 20.08232 | MLP_R_5 | 294 | 206 | Poland | Northern | R | MLP | mlpr5_4 | 0 | 0 | 0 | 0 | 0 | 0 | 0 | | | 0 | 0 | 60 | 179 | 91 | 60 | 4809 | 6413 | 720 | 2274 | 1815 | 0 | 1 | 0 | 0 | 0 | 0 | 1 | 0 | 1 | 0 | 0 | 1 | 0 | 0 | 0 | 0 | 0 |
| 50.16315 | 20.08429 | MLP_R_5 | 289 | 206 | Poland | Northern | R | MLP | mlpr5_5 | 2 | 1 | 0 | 0 | 0 | 0 | 0 | 0 | | 1 | 0 | 18 | 18 | 20 | 18 | 19410 | 7287 | 0 | 717 | 18693 | 1 | 1 | 0 | 1 | 0 | 0 | 1 | 0 | 1 | 0 | 0 | 1 | 1 | 0 | 1 | 0 | 1 |
| 50.16108 | 20.08359 | MLP_R_5 | 266 | 206 | Poland | Northern | R | MLP | mlpr5_6 | 0 | 0 | 0 | 0 | 0 | 0 | 0 | | | 0 | 0 | NA | NA | 48 | 48 | 8184 | 7200 | 0 | 1102 | 7083 | 1 | 1 | 0 | 1 | 0 | 0 | 1 | 1 | 1 | 0 | 0 | 1 | 1 | 0 | 1 | 0 | 1 |
| 50.15923 | 20.08111 | MLP_R_5 | 270 | 206 | Poland | Northern | R | MLP | mlpr5_7 | 0 | 0 | 0 | 0 | 0 | 0 | 0 | | | 0 | 0 | 123 | 180 | 40 | 40 | 12041 | 4652 | 0 | 0 | 12041 | 0 | 1 | 0 | 1 | 0 | 0 | 1 | 0 | 1 | 0 | 0 | 1 | 0 | 0 | 1 | 0 | 1 |
| 50.16059 | 20.0784 | MLP_R_5 | 278 | 206 | Poland | Northern | R | MLP | mlpr5_8 | 0 | 0 | 0 | 0 | 0 | 0 | 0 | | | 0 | 0 | 141 | 249 | 141 | 141 | 3930 | 4289 | 3930 | 0 | 0 | 0 | 1 | 0 | 0 | 0 | 0 | 1 | 1 | 1 | 0 | 0 | 1 | 0 | 0 | 1 | 0 | 1 |

| | | | | | | | | | | | | | | | | | | | | | | | | | | | | | | | | | | | | | | | | | | | | | | |
|---|---|---|---|---|---|---|---|---|---|---|---|---|---|---|---|---|---|---|---|---|---|---|---|---|---|---|---|---|---|---|---|---|---|---|---|---|---|---|---|---|---|---|---|---|---|---|
| 50.29705 | 20.04346 | MLP_R_6 | 251 | 199 | Poland | Northern | R | MLP | mlpr6_1 | 1 | 1 | 0 | 0 | 0 | 0 | 0 | 0 | | 1 | 0 | NA | NA | 126 | 126 | 23317 | 1644 | 0 | 316 | 23002 | 1 | 1 | 1 | 1 | 0 | 0 | 0 | 0 | 1 | 1 | 0 | 1 | 0 | 0 | 0 | 1 | 0 |
| 50.29857 | 20.04692 | MLP_R_6 | 253 | 199 | Poland | Northern | R | MLP | mlpr6_2 | 2 | 1 | 0 | 0 | 0 | 0 | 0 | 0 | | 1 | 0 | 1 | 27 | 73 | 1 | 24805 | 1761 | 2532 | 525 | 21748 | 1 | 1 | 0 | 1 | 0 | 0 | 0 | 1 | 1 | 1 | 1 | 1 | 0 | 0 | 0 | 1 | 1 |
| 50.30095 | 20.04911 | MLP_R_6 | 253 | 199 | Poland | Northern | R | MLP | mlpr6_3 | 0 | 0 | 0 | 0 | 0 | 0 | 0 | | | 0 | 0 | 202 | 64 | 118 | 64 | 19424 | 795 | 0 | 1507 | 17917 | 1 | 1 | 0 | 1 | 0 | 0 | 1 | 1 | 1 | 1 | 0 | 1 | 0 | 0 | 0 | 1 | 1 |
| 50.3032 | 20.05128 | MLP_R_6 | 256 | 199 | Poland | Northern | R | MLP | mlpr6_4 | 1 | 1 | 0 | 0 | 0 | 0 | 0 | 0 | | 1 | 0 | NA | NA | 63 | 63 | 9482 | 1735 | 0 | 804 | 8677 | 1 | 1 | 0 | 1 | 0 | 0 | 0 | 1 | 1 | 1 | 1 | 1 | 0 | 0 | 0 | 0 | 0 |
| 50.30617 | 20.05123 | MLP_R_6 | 258 | 199 | Poland | Northern | R | MLP | mlpr6_5 | 1 | 1 | 0 | 0 | 0 | 0 | 0 | 0 | | 1 | 0 | NA | NA | 34 | 34 | 12398 | 1035 | 0 | 693 | 11705 | 1 | 1 | 0 | 1 | 0 | 0 | 1 | 1 | 1 | 1 | 1 | 0 | 0 | 0 | 0 | 0 | 0 |
| 50.30868 | 20.05124 | MLP_R_6 | 258 | 199 | Poland | Northern | R | MLP | mlpr6_6 | 1 | 1 | 2 | 1 | 0 | 0 | 1 | 1 | 1 | 0 | 0 | NA | NA | 10 | 10 | 29542 | 1697 | 0 | 7857 | 21685 | 1 | 1 | 0 | 1 | 0 | 0 | 1 | 0 | 1 | 1 | 0 | 1 | 0 | 0 | 0 | 0 | 1 |
| 50.31112 | 20.05263 | MLP_R_6 | 262 | 199 | Poland | Northern | R | MLP | mlpr6_7 | 1 | 1 | 0 | 0 | 0 | 0 | 0 | 0 | | 1 | 0 | NA | NA | 7 | 7 | 21729 | 1451 | 0 | 2041 | 19688 | 1 | 1 | 0 | 1 | 0 | 0 | 0 | 0 | 1 | 1 | 0 | 1 | 0 | 0 | 0 | 0 | 0 |
| 50.31304 | 20.05426 | MLP_R_6 | 263 | 199 | Poland | Northern | R | MLP | mlpr6_8 | 2 | 1 | 0 | 0 | 0 | 0 | 0 | 0 | | 1 | 0 | NA | NA | 18 | 18 | 8189 | 1451 | 0 | 1333 | 6856 | 1 | 1 | 0 | 1 | 0 | 0 | 0 | 0 | 1 | 1 | 0 | 1 | 0 | 0 | 0 | 1 | 1 |
| 50.09411 | 19.9418 | MLP_U_5 | 217 | 200 | Poland | Northern | U | MLP | mlpu5_1 | 1 | 1 | 0 | 0 | 0 | 0 | 0 | 0 | | 1 | 0 | 141 | NA | NA | 141 | 31785 | 7800 | 31785 | 0 | 0 | 0 | 1 | 0 | 0 | 1 | 0 | 0 | 1 | 0 | 0 | 1 | 0 | 0 | 0 | 0 | 0 | 0 |
| 50.09256 | 19.94524 | MLP_U_5 | 216 | 200 | Poland | Northern | U | MLP | mlpu5_2 | 1 | 1 | 0 | 0 | 0 | 0 | 0 | 0 | | 1 | 0 | 45 | NA | NA | 45 | 21286 | 19900 | 15889 | 5398 | 0 | 0 | 1 | 0 | 0 | 1 | 0 | 1 | 1 | 1 | 0 | 0 | 1 | 1 | 0 | 1 | 0 | 0 |
| 50.09169 | 19.94925 | MLP_U_5 | 215 | 200 | Poland | Northern | U | MLP | mlpu5_3 | 0 | 0 | 0 | 0 | 0 | 0 | 0 | | | 0 | 0 | 10 | NA | NA | 10 | 16016 | 21900 | 14403 | 1613 | 0 | 0 | 1 | 0 | 0 | 0 | 0 | 0 | 1 | 1 | 0 | 0 | 1 | 1 | 1 | 0 | 0 | 0 |
| 50.08933 | 19.94885 | MLP_U_5 | 214 | 200 | Poland | Northern | U | MLP | mlpu5_4 | 0 | 0 | 0 | 0 | 0 | 0 | 0 | | | 0 | 0 | 60 | NA | NA | 60 | 12017 | 38300 | 11346 | 671 | 0 | 0 | 1 | 0 | 0 | 0 | 0 | 0 | 1 | 0 | 0 | 0 | 1 | 1 | 0 | 1 | 0 | 0 |
| 50.08756 | 19.9476 | MLP_U_5 | 211 | 200 | Poland | Northern | U | MLP | mlpu5_5 | 1 | 1 | 1 | 1 | 0 | 0 | 1 | 1 | 1 | 0 | 0 | 68 | NA | NA | 68 | 17655 | 26500 | 17655 | 0 | 0 | 1 | 1 | 0 | 0 | 0 | 0 | 0 | 1 | 0 | 0 | 0 | 1 | 1 | 0 | 1 | 0 | 1 |
| 50.08916 | 19.94424 | MLP_U_5 | 212 | 200 | Poland | Northern | U | MLP | mlpu5_6 | 0 | 0 | 1 | 1 | 0 | 0 | 0 | | 0 | 0 | 1 | 1 | NA | NA | 1 | 37877 | 12300 | 36572 | 1305 | 0 | 1 | 1 | 0 | 0 | 0 | 0 | 1 | 1 | 1 | 0 | 0 | 1 | 1 | 0 | 1 | 0 | 1 |
| 50.09082 | 19.94087 | MLP_U_5 | 219 | 200 | Poland | Northern | U | MLP | mlpu5_7 | 0 | 0 | 0 | 0 | 0 | 0 | 0 | | | 0 | 0 | 31 | NA | NA | 31 | 24819 | 13600 | 23323 | 1496 | 0 | 1 | 1 | 0 | 0 | 0 | 0 | 1 | 1 | 0 | 0 | 0 | 1 | 1 | 0 | 1 | 0 | 0 |
| 50.08958 | 19.93871 | MLP_U_5 | 222 | 200 | Poland | Northern | U | MLP | mlpu5_8 | 1 | 1 | 0 | 0 | 0 | 0 | 0 | 0 | | 1 | 0 | 25 | NA | NA | 25 | 13336 | 27400 | 12381 | 955 | 0 | 0 | 1 | 0 | 0 | 0 | 0 | 1 | 1 | 0 | 0 | 0 | 0 | 1 | 1 | 0 | 0 | 1 |
| 50.07617 | 19.95356 | MLP_U_6 | 215 | 201 | Poland | Northern | U | MLP | mlpu6_1 | 4 | 1 | 0 | 0 | 0 | 0 | 0 | 0 | | 1 | 0 | NA | NA | NA | | 58689 | 0 | 58689 | 0 | 0 | 0 | 1 | 0 | 0 | 0 | 1 | 0 | 0 | 0 | 0 | 0 | 0 | 0 | 0 | 0 | 1 | 0 |
| 50.07619 | 19.95687 | MLP_U_6 | 212 | 201 | Poland | Northern | U | MLP | mlpu6_2 | 2 | 1 | 0 | 0 | 0 | 0 | 0 | 0 | | 1 | 0 | 22 | NA | NA | 22 | 42837 | 10800 | 42044 | 793 | 0 | 1 | 1 | 0 | 0 | 0 | 1 | 0 | 1 | 0 | 0 | 0 | 0 | 0 | 1 | 0 | 0 | 1 |
| 50.07746 | 19.95937 | MLP_U_6 | 209 | 201 | Poland | Northern | U | MLP | mlpu6_3 | 0 | 0 | 0 | 0 | 0 | 0 | 0 | | | 0 | 0 | 10 | 1 | 5 | 1 | 23044 | 16300 | 4528 | 6322 | 12194 | 1 | 1 | 0 | 0 | 0 | 1 | 0 | 1 | 1 | 1 | 0 | 1 | 0 | 1 | 0 | 0 | 0 |
| 50.0798 | 19.96099 | MLP_U_6 | 208 | 201 | Poland | Northern | U | MLP | mlpu6_4 | 1 | 1 | 0 | 0 | 0 | 0 | 0 | 0 | | 1 | 0 | 11 | 1 | 109 | 1 | 17388 | 26500 | 12798 | 4590 | 0 | 0 | 1 | 1 | 0 | 0 | 0 | 1 | 1 | 1 | 0 | 0 | 1 | 1 | 1 | 1 | 0 | 0 |
| 50.07987 | 19.96414 | MLP_U_6 | 208 | 201 | Poland | Northern | U | MLP | mlpu6_5 | 0 | 0 | 0 | 0 | 0 | 0 | 0 | | | 0 | 0 | 2 | 209 | 140 | 2 | 22462 | 26200 | 11666 | 10796 | 0 | 1 | 1 | 0 | 0 | 0 | 0 | 0 | 1 | 1 | 0 | 0 | 1 | 1 | 0 | 0 | 0 | 0 |
| 50.0781 | 19.96464 | MLP_U_6 | 208 | 201 | Poland | Northern | U | MLP | mlpu6_6 | 1 | 1 | 0 | 0 | 0 | 0 | 0 | 0 | | 1 | 0 | 2 | 151 | 125 | 2 | 22739 | 20500 | 19035 | 3704 | 0 | 1 | 1 | 0 | 0 | 0 | 0 | 1 | 1 | 1 | 0 | 1 | 1 | 1 | 0 | 1 | 0 | 0 |
| 50.07808 | 19.96814 | MLP_U_6 | 207 | 201 | Poland | Northern | U | MLP | mlpu6_7 | 0 | 0 | 0 | 0 | 0 | 0 | 0 | | | 0 | 0 | 37 | NA | NA | 37 | 20333 | 21900 | 20333 | 0 | 0 | 1 | 1 | 0 | 0 | 0 | 0 | 1 | 1 | 1 | 0 | 1 | 0 | 1 | 0 | 1 | 0 | 0 |
| 50.07936 | 19.96924 | MLP_U_6 | 207 | 201 | Poland | Northern | U | MLP | mlpu6_8 | 0 | 0 | 0 | 0 | 0 | 0 | 0 | | | 0 | 0 | 15 | 142 | 71 | 15 | 17376 | 27400 | 8191 | 9185 | 0 | 0 | 1 | 0 | 0 | 0 | 0 | 1 | 1 | 1 | 0 | 0 | 1 | 1 | 1 | 0 | 0 | 0 |
| 50.07878 | 20.0551 | MLP_U_7 | 206 | 207 | Poland | Northern | U | MLP | mlpu7_1 | 0 | 0 | 0 | 0 | 0 | 0 | 0 | | | 0 | 0 | 193 | 133 | 371 | 133 | 29830 | 6100 | 29222 | 608 | 0 | 1 | 1 | 0 | 0 | 1 | 0 | 0 | 1 | 0 | 1 | 0 | 1 | 0 | 0 | 0 | 1 | 0 |
| 50.08063 | 20.05194 | MLP_U_7 | 206 | 207 | Poland | Northern | U | MLP | mlpu7_2 | 0 | 0 | 0 | 0 | 0 | 0 | 0 | | | 0 | 0 | 163 | 148 | NA | 148 | 21679 | 2800 | 21679 | 0 | 0 | 1 | 1 | 0 | 0 | 1 | 0 | 0 | 0 | 0 | 1 | 1 | 0 | 0 | 0 | 0 | 0 | 0 |
| 50.0805 | 20.04892 | MLP_U_7 | 207 | 207 | Poland | Northern | U | MLP | mlpu7_3 | 0 | 0 | 0 | 0 | 0 | 0 | 0 | | | 0 | 0 | NA | 156 | NA | 156 | 22583 | 15100 | 22583 | 0 | 0 | 1 | 1 | 0 | 0 | 1 | 0 | 1 | 1 | 0 | 1 | 1 | 0 | 1 | 0 | 0 | 0 | 0 |
| 50.08233 | 20.04951 | MLP_U_7 | 208 | 207 | Poland | Northern | U | MLP | mlpu7_4 | 0 | 0 | 0 | 0 | 0 | 0 | 0 | | | 0 | 0 | 127 | 1 | 110 | 1 | 29956 | 12300 | 13139 | 264 | 16553 | 1 | 1 | 0 | 1 | 1 | 0 | 0 | 1 | 0 | 0 | 1 | 0 | 1 | 0 | 0 | 0 | 0 |
| 50.08355 | 20.05295 | MLP_U_7 | 205 | 207 | Poland | Northern | U | MLP | mlpu7_5 | 1 | 1 | 0 | 0 | 0 | 0 | 0 | 0 | | 1 | 0 | 15 | 12 | 78 | 12 | 20210 | 10200 | 13239 | 3899 | 3072 | 1 | 1 | 0 | 1 | 0 | 0 | 0 | 1 | 1 | 1 | 0 | 1 | 0 | 0 | 0 | 0 | 1 |
| 50.08255 | 20.05592 | MLP_U_7 | 208 | 207 | Poland | Northern | U | MLP | mlpu7_6 | 0 | 0 | 0 | 0 | 0 | 0 | 0 | | | 0 | 0 | 18 | 140 | 160 | 18 | 17807 | 10800 | 11940 | 5867 | 0 | 0 | 1 | 0 | 0 | 0 | 0 | 0 | 1 | 1 | 0 | 1 | 1 | 0 | 0 | 0 | 0 | 0 |
| 50.08189 | 20.05894 | MLP_U_7 | 211 | 207 | Poland | Northern | U | MLP | mlpu7_7 | 0 | 0 | 0 | 0 | 0 | 0 | 0 | | | 0 | 0 | 47 | NA | 114 | 47 | 14353 | 14200 | 0 | 14353 | 0 | 0 | 1 | 0 | 0 | 0 | 0 | 0 | 0 | 1 | 0 | 0 | 1 | 0 | 0 | 0 | 0 | 0 |
| 50.07978 | 20.05941 | MLP_U_7 | 210 | 207 | Poland | Northern | U | MLP | mlpu7_8 | 1 | 1 | 0 | 0 | 0 | 0 | 0 | 0 | | 1 | 0 | 47 | 155 | 301 | 47 | 23476 | 11000 | 21599 | 1877 | 0 | 1 | 1 | 1 | 1 | 0 | 0 | 0 | 0 | 1 | 0 | 0 | 1 | 0 | 0 | 0 | 0 | 1 |
| 50.05375 | 20.05612 | MLP_U_8 | 198 | 209 | Poland | Northern | U | MLP | mlpu8_1 | 2 | 1 | 0 | 0 | 0 | 0 | 0 | 0 | | 1 | 0 | NA | NA | 32 | 32 | 50299 | 3000 | 0 | 2077 | 48222 | 1 | 1 | 0 | 1 | 0 | 0 | 0 | 0 | 1 | 0 | 0 | 1 | 0 | 0 | 0 | 1 | 0 |
| 50.05369 | 20.05977 | MLP_U_8 | 198 | 209 | Poland | Northern | U | MLP | mlpu8_2 | 2 | 1 | 0 | 0 | 0 | 0 | 0 | 0 | | 1 | 0 | 63 | 16 | 27 | 16 | 52430 | 2700 | 436 | 3383 | 48611 | 1 | 1 | 0 | 1 | 0 | 0 | 0 | 0 | 1 | 0 | 0 | 1 | 0 | 0 | 0 | 0 | 0 |
| 50.05501 | 20.06131 | MLP_U_8 | 197 | 209 | Poland | Northern | U | MLP | mlpu8_3 | 2 | 1 | 0 | 0 | 0 | 0 | 0 | 0 | | 1 | 0 | 64 | 90 | 67 | 64 | 30391 | 3000 | 615 | 1239 | 28537 | 1 | 1 | 0 | 1 | 0 | 0 | 0 | 0 | 1 | 1 | 1 | 1 | 0 | 0 | 0 | 0 | 0 |
| 50.05803 | 20.06032 | MLP_U_8 | 197 | 209 | Poland | Northern | U | MLP | mlpu8_4 | 0 | 0 | 0 | 0 | 0 | 0 | 0 | | | 0 | 0 | 55 | 7 | 108 | 7 | 34466 | 1800 | 1428 | 0 | 33038 | 1 | 1 | 0 | 1 | 0 | 0 | 1 | 0 | 0 | 1 | 1 | 1 | 0 | 0 | 0 | 1 | 0 |
| 50.0606 | 20.05952 | MLP_U_8 | 198 | 209 | Poland | Northern | U | MLP | mlpu8_5 | 0 | 0 | 0 | 0 | 0 | 0 | 0 | | | 0 | 0 | 34 | 20 | 45 | 20 | 9527 | 9900 | 5609 | 1299 | 2620 | 1 | 1 | 0 | 1 | 0 | 0 | 1 | 1 | 1 | 0 | 1 | 1 | 0 | 0 | 0 | 0 | 0 |
| 50.0626 | 20.05739 | MLP_U_8 | 199 | 209 | Poland | Northern | U | MLP | mlpu8_6 | 0 | 0 | 0 | 0 | 0 | 0 | 0 | | | 0 | 0 | 2 | 85 | 40 | 2 | 10048 | 9500 | 3407 | 5842 | 799 | 0 | 1 | 1 | 0 | 0 | 0 | 1 | 1 | 1 | 0 | 0 | 1 | 0 | 0 | 0 | 0 | 1 |
| 50.06414 | 20.05394 | MLP_U_8 | 203 | 209 | Poland | Northern | U | MLP | mlpu8_7 | 0 | 0 | 0 | 0 | 0 | 0 | 0 | | | 0 | 0 | 22 | 19 | 2 | 2 | 16822 | 19900 | 5980 | 3031 | 7811 | 1 | 1 | 1 | 1 | 0 | 0 | 1 | 1 | 1 | 0 | 0 | 1 | 0 | 1 | 0 | 0 | 1 |
| 50.06568 | 20.05196 | MLP_U_8 | 206 | 209 | Poland | Northern | U | MLP | mlpu8_8 | 1 | 1 | 0 | 0 | 0 | 0 | 0 | 0 | | 1 | 0 | 10 | 62 | 80 | 10 | 18225 | 16300 | 17424 | 801 | 0 | 1 | 1 | 1 | 0 | 1 | 0 | 1 | 1 | 0 | 0 | 0 | 1 | 1 | 1 | 0 | 0 | 0 |
| 49.99929 | 19.43311 | MLP_S_8 | 233 | 173 | Poland | Northern | S | MLP | mlps8_1 | 0 | 0 | 0 | 0 | 0 | 0 | 0 | | | 0 | 0 | 27 | 1 | 47 | 1 | 17156 | 1620 | 11245 | 444 | 5467 | 1 | 1 | 0 | 1 | 0 | 1 | 1 | 1 | 1 | 0 | 1 | 0 | 1 | 0 | 0 | 0 | 1 |
| 49.99826 | 19.43616 | MLP_S_8 | 228 | 173 | Poland | Northern | S | MLP | mlps8_2 | 0 | 0 | 0 | 0 | 0 | 0 | 0 | | | 0 | 0 | 1 | 4 | 22 | 1 | 23896 | 5234 | 15181 | 5893 | 2822 | 1 | 1 | 0 | 1 | 0 | 0 | 1 | 1 | 1 | 0 | 1 | 1 | 0 | 0 | 0 | 0 | 0 |
| 49.99875 | 19.44004 | MLP_S_8 | 226 | 173 | Poland | Northern | S | MLP | mlps8_3 | 0 | 0 | 0 | 0 | 0 | 0 | 0 | | | 0 | 0 | 67 | 20 | 6 | 6 | 12086 | 3199 | 841 | 3530 | 7714 | 1 | 1 | 0 | 1 | 0 | 0 | 1 | 1 | 1 | 0 | 1 | 1 | 0 | 0 | 0 | 0 | 0 |
| 49.99761 | 19.44295 | MLP_S_8 | 227 | 173 | Poland | Northern | S | MLP | mlps8_4 | 1 | 1 | 0 | 0 | 0 | 0 | 0 | 0 | | 1 | 0 | 67 | 21 | 50 | 21 | 34326 | 738 | 4699 | 0 | 29627 | 1 | 1 | 0 | 1 | 0 | 0 | 1 | 1 | 0 | 1 | 1 | 0 | 0 | 0 | 0 | 0 | 0 |
| 49.99492 | 19.44249 | MLP_S_8 | 227 | 173 | Poland | Northern | S | MLP | mlps8_5 | 1 | 1 | 0 | 0 | 0 | 0 | 0 | 0 | | 1 | 0 | NA | 1 | 116 | 1 | 47586 | 456 | 0 | 0 | 47586 | 1 | 1 | 0 | 1 | 0 | 0 | 0 | 0 | 0 | 1 | 0 | 0 | 0 | 0 | 1 | 0 | 1 |
| 49.99383 | 19.44014 | MLP_S_8 | 227 | 173 | Poland | Northern | S | MLP | mlps8_6 | 0 | 0 | 0 | 0 | 0 | 0 | 0 | | | 0 | 0 | 18 | 1 | 24 | 1 | 29571 | 4477 | 8159 | 2787 | 18625 | 1 | 1 | 0 | 1 | 0 | 0 | 0 | 1 | 1 | 0 | 1 | 1 | 0 | 0 | 1 | 0 | 1 |
| 49.99259 | 19.43713 | MLP_S_8 | 242 | 173 | Poland | Northern | S | MLP | mlps8_7 | 0 | 0 | 0 | 0 | 0 | 0 | 0 | | | 0 | 0 | 42 | 144 | 128 | 42 | 11033 | 9636 | 5842 | 5191 | 0 | 1 | 1 | 0 | 1 | 0 | 0 | 1 | 1 | 1 | 0 | 1 | 1 | 0 | 1 | 0 | 0 | 0 |
| 49.9928 | 19.43279 | MLP_S_8 | 240 | 173 | Poland | Northern | S | MLP | mlps8_8 | 0 | 0 | 0 | 0 | 0 | 0 | 0 | | | 0 | 0 | 32 | 248 | 157 | 32 | 4252 | 10352 | 1963 | 2289 | 0 | 0 | 1 | 0 | 0 | 0 | 0 | 1 | 1 | 1 | 0 | 1 | 1 | 1 | 0 | 0 | 0 | 0 |
| 49.98146 | 19.63878 | MLP_R_7 | 215 | 186 | Poland | Northern | R | MLP | mlpr7_1 | 0 | 0 | 0 | 0 | 0 | 0 | 0 | | | 0 | 0 | NA | NA | 234 | 234 | 24632 | 0 | 0 | 0 | 24632 | 0 | 1 | 0 | 1 | 0 | 0 | 1 | 0 | 0 | 1 | 0 | 0 | 0 | 0 | 0 | 1 | 1 |
| 49.97919 | 19.63659 | MLP_R_7 | 215 | 186 | Poland | Northern | R | MLP | mlpr7_2 | 1 | 1 | 0 | 0 | 0 | 0 | 0 | 0 | | 1 | 0 | NA | NA | 47 | 47 | 16207 | 176 | 0 | 1356 | 14851 | 0 | 1 | 0 | 0 | 0 | 0 | 1 | 0 | 0 | 1 | 1 | 0 | 0 | 0 | 0 | 1 | 0 |
| 49.9779 | 19.63609 | MLP_R_7 | 214 | 186 | Poland | Northern | R | MLP | mlpr7_3 | 0 | 0 | 0 | 0 | 0 | 0 | 0 | | | 0 | 0 | NA | NA | 43 | 43 | 31805 | 1165 | 0 | 2376 | 29429 | 0 | 1 | 0 | 0 | 0 | 0 | 1 | 0 | 0 | 1 | 1 | 0 | 0 | 0 | 0 | 0 | 0 |
| 49.97873 | 19.6401 | MLP_R_7 | 213 | 186 | Poland | Northern | R | MLP | mlpr7_4 | 4 | 1 | 0 | 0 | 0 | 0 | 0 | 0 | | 1 | 0 | NA | NA | 21 | 21 | 23829 | 384 | 0 | 1746 | 22083 | 0 | 1 | 0 | 0 | 0 | 0 | 1 | 1 | 1 | 0 | 1 | 0 | 0 | 0 | 0 | 1 | 0 |
| 49.97852 | 19.64415 | MLP_R_7 | 214 | 186 | Poland | Northern | R | MLP | mlpr7_5 | 1 | 1 | 0 | 0 | 0 | 0 | 0 | 0 | | 1 | 0 | NA | NA | 33 | 33 | 8412 | 927 | 0 | 2895 | 5517 | 0 | 1 | 0 | 0 | 0 | 0 | 1 | 1 | 1 | 0 | 1 | 0 | 0 | 0 | 0 | 0 | 0 |
| 49.98039 | 19.64628 | MLP_R_7 | 211 | 186 | Poland | Northern | R | MLP | mlpr7_6 | 0 | 0 | 0 | 0 | 0 | 0 | 0 | | | 0 | 0 | NA | NA | 57 | 57 | 24543 | 510 | 0 | 642 | 23902 | 0 | 1 | 0 | 0 | 0 | 0 | 0 | 1 | 0 | 0 | 1 | 0 | 0 | 0 | 0 | 0 | 1 |
| 49.98176 | 19.64948 | MLP_R_7 | 213 | 186 | Poland | Northern | R | MLP | mlpr7_7 | 0 | 0 | 0 | 0 | 0 | 0 | 0 | | | 0 | 0 | NA | NA | 43 | 43 | 16152 | 1016 | 0 | 3451 | 12701 | 1 | 1 | 0 | 1 | 0 | 0 | 0 | 1 | 1 | 1 | 1 | 0 | 0 | 0 | 0 | 1 | 0 |
| 49.98453 | 19.64883 | MLP_R_7 | 214 | 186 | Poland | Northern | R | MLP | mlpr7_8 | 1 | 1 | 0 | 0 | 0 | 0 | 0 | 0 | | 1 | 0 | NA | NA | 23 | 23 | 31765 | 320 | 0 | 2356 | 29408 | 1 | 1 | 0 | 1 | 0 | 0 | 0 | 1 | 0 | 1 | 1 | 0 | 0 | 0 | 0 | 1 | 1 |
| 50.01073 | 19.51424 | MLP_R_8 | 219 | 176 | Poland | Northern | R | MLP | mlpr8_1 | 1 | 1 | 0 | 0 | 0 | 0 | 0 | 0 | | 1 | 0 | NA | NA | NA | | 23714 | 792 | 0 | 0 | 23714 | 1 | 1 | 0 | 1 | 0 | 0 | 1 | 0 | 0 | 0 | 1 | 0 | 0 | 0 | 0 | 1 | 1 |
| 50.01343 | 19.5128 | MLP_R_8 | 218 | 176 | Poland | Northern | R | MLP | mlpr8_2 | 0 | 0 | 0 | 0 | 0 | 0 | 0 | | | 0 | 0 | NA | NA | 104 | 104 | 6271 | 0 | 0 | 0 | 6271 | 0 | 1 | 0 | 0 | 0 | 0 | 1 | 0 | 0 | 1 | 0 | 0 | 0 | 0 | 0 | 0 | 0 |
| 50.0163 | 19.51323 | MLP_R_8 | 217 | 176 | Poland | Northern | R | MLP | mlpr8_3 | 1 | 1 | 0 | 0 | 0 | 0 | 0 | 0 | | 1 | 0 | NA | NA | 21 | 21 | 16052 | 1722 | 0 | 1710 | 14342 | 1 | 1 | 0 | 1 | 0 | 0 | 1 | 0 | 1 | 1 | 0 | 1 | 0 | 0 | 0 | 0 | 0 |
| 50.01878 | 19.51556 | MLP_R_8 | 217 | 176 | Poland | Northern | R | MLP | mlpr8_4 | 0 | 0 | 0 | 0 | 0 | 0 | 0 | | | 0 | 0 | NA | NA | 78 | 78 | 3974 | 0 | 0 | 0 | 3974 | 1 | 1 | 0 | 1 | 0 | 0 | 1 | 0 | 0 | 1 | 0 | 0 | 0 | 0 | 0 | 1 | 1 |
| 50.02027 | 19.51936 | MLP_R_8 | 218 | 176 | Poland | Northern | R | MLP | mlpr8_5 | 2 | 1 | 0 | 0 | 0 | 0 | 0 | 0 | | 1 | 0 | NA | NA | NA | | 7274 | 0 | 0 | 0 | 7274 | 1 | 1 | 0 | 1 | 0 | 0 | 1 | 0 | 0 | 1 | 0 | 0 | 0 | 0 | 0 | 0 | 0 |
| 50.02083 | 19.52366 | MLP_R_8 | 217 | 176 | Poland | Northern | R | MLP | mlpr8_6 | 1 | 1 | 0 | 0 | 0 | 0 | 0 | 0 | | 1 | 0 | NA | NA | NA | | 22405 | 0 | 0 | 0 | 22405 | 1 | 1 | 0 | 1 | 0 | 0 | 0 | 0 | 0 | 0 | 0 | 0 | 0 | 0 | 0 | 0 | 1 |
| 50.02033 | 19.52771 | MLP_R_8 | 218 | 176 | Poland | Northern | R | MLP | mlpr8_7 | 0 | 0 | 0 | 0 | 0 | 0 | 0 | | | 0 | 0 | NA | NA | 145 | 145 | 5475 | 0 | 0 | 0 | 5475 | 1 | 1 | 0 | 1 | 0 | 0 | 1 | 0 | 0 | 1 | 0 | 0 | 0 | 0 | 0 | 0 | 0 |
| 50.01835 | 19.52978 | MLP_R_8 | 217 | 176 | Poland | Northern | R | MLP | mlpr8_8 | 1 | 1 | 0 | 0 | 0 | 0 | 0 | 0 | | 1 | 0 | NA | NA | 50 | 50 | 12245 | 1150 | 0 | 1340 | 10906 | 0 | 1 | 0 | 1 | 0 | 0 | 1 | 0 | 1 | 1 | 1 | 0 | 0 | 0 | 0 | 1 | 0 |
| 50.04003 | 19.91735 | MLP_U_3 | 210 | 201 | Poland | Northern | U | MLP | mlpu3_1 | 0 | 0 | 0 | 0 | 0 | 0 | 0 | | | 0 | 0 | 130 | 64 | 97 | 64 | 48515 | 1200 | 0 | 790 | 47725 | 1 | 1 | 1 | 1 | 0 | 0 | 1 | 0 | 1 | 0 | 1 | 0 | 0 | 0 | 0 | 1 | 1 |
| 50.03773 | 19.91535 | MLP_U_3 | 210 | 201 | Poland | Northern | U | MLP | mlpu3_2 | 0 | 0 | 0 | 0 | 0 | 0 | 0 | | | 0 | 0 | NA | NA | NA | | 42641 | 0 | 0 | 0 | 42641 | 1 | 1 | 1 | 1 | 0 | 0 | 0 | 0 | 0 | 1 | 0 | 0 | 0 | 0 | 0 | 0 | 1 |
| 50.03536 | 19.91389 | MLP_U_3 | 214 | 201 | Poland | Northern | U | MLP | mlpu3_3 | 0 | 0 | 0 | 0 | 0 | 0 | 0 | | | 0 | 0 | 39 | NA | NA | 39 | 39511 | 0 | 0 | 0 | 39511 | 1 | 1 | 1 | 1 | 0 | 0 | 0 | 0 | 1 | 0 | 1 | 0 | 0 | 0 | 0 | 1 | 0 |
| 50.03424 | 19.91698 | MLP_U_3 | 206 | 201 | Poland | Northern | U | MLP | mlpu3_4 | 0 | 0 | 0 | 0 | 0 | 0 | 0 | | | 0 | 0 | NA | 143 | 48 | 48 | 14196 | 700 | 0 | 0 | 14196 | 0 | 1 | 0 | 1 | 0 | 0 | 0 | 0 | 1 | 0 | 1 | 0 | 0 | 0 | 0 | 0 | 0 |
| 50.03394 | 19.92057 | MLP_U_3 | 207 | 201 | Poland | Northern | U | MLP | mlpu3_5 | 1 | 1 | 0 | 0 | 1 | 1 | 0 | 0 | | 1 | 0 | 210 | 145 | 65 | 65 | 21984 | 14700 | 4428 | 0 | 17556 | 1 | 1 | 0 | 1 | 0 | 0 | 0 | 1 | 1 | 0 | 0 | 0 | 1 | 0 | 1 | 1 | 0 |
| 50.03416 | 19.9239 | MLP_U_3 | 207 | 201 | Poland | Northern | U | MLP | mlpu3_6 | 0 | 0 | 0 | 0 | 0 | 0 | 0 | | | 0 | 0 | 212 | 91 | NA | 91 | 8707 | 33000 | 8162 | 0 | 545 | 0 | 1 | 0 | 0 | 0 | 0 | 1 | 1 | 0 | 0 | 0 | 0 | 1 | 0 | 1 | 0 | 0 |
| 50.03588 | 19.92733 | MLP_U_3 | 206 | 201 | Poland | Northern | U | MLP | mlpu3_7 | 2 | 1 | 0 | 0 | 0 | 0 | 0 | 0 | | 1 | 0 | 119 | 132 | 4 | 4 | 30459 | 10900 | 7496 | 16913 | 6050 | 0 | 1 | 0 | 0 | 1 | 0 | 1 | 1 | 1 | 1 | 0 | 1 | 1 | 0 | 1 | 0 | 0 |
| 50.0374 | 19.93091 | MLP_U_3 | 203 | 201 | Poland | Northern | U | MLP | mlpu3_8 | 0 | 0 | 0 | 0 | 0 | 0 | 0 | | | 0 | 0 | 110 | 20 | 1 | 1 | 27869 | 2500 | 2301 | 10524 | 15044 | 1 | 1 | 0 | 1 | 0 | 0 | 0 | 1 | 1 | 1 | 0 | 1 | 1 | 0 | 1 | 0 | 0 |
| 50.01751 | 19.80224 | MLP_U_4 | 206 | 194 | Poland | Northern | U | MLP | mlpu4_1 | 0 | 0 | 0 | 0 | 0 | 0 | 0 | | | 0 | 0 | NA | NA | NA | | 15901 | 0 | 0 | 0 | 15901 | 1 | 1 | 1 | 1 | 0 | 0 | 1 | 1 | 0 | 1 | 0 | 1 | 0 | 1 | 0 | 1 | 1 |
| 50.01761 | 19.80566 | MLP_U_4 | 209 | 194 | Poland | Northern | U | MLP | mlpu4_2 | 0 | 0 | 0 | 0 | 0 | 0 | 0 | | | 0 | 0 | NA | NA | NA | | 23807 | 0 | 0 | 0 | 23807 | 1 | 1 | 1 | 1 | 0 | 1 | 1 | 1 | 0 | 0 | 0 | 1 | 0 | 0 | 0 | 0 | 0 |
| 50.01981 | 19.80314 | MLP_U_4 | 213 | 194 | Poland | Northern | U | MLP | mlpu4_3 | 2 | 1 | 0 | 0 | 0 | 0 | 0 | 0 | | 1 | 0 | NA | NA | NA | | 8653 | 0 | 0 | 0 | 8653 | 1 | 1 | 1 | 1 | 1 | 0 | 1 | 1 | 1 | 1 | 0 | 1 | 0 | 1 | 0 | 1 | 1 |
| 50.0216 | 19.80423 | MLP_U_4 | 208 | 194 | Poland | Northern | U | MLP | mlpu4_4 | 2 | 1 | 0 | 0 | 0 | 0 | 0 | 0 | | 1 | 0 | 31 | 86 | 105 | 31 | 11621 | 0 | 0 | 0 | 11621 | 1 | 1 | 0 | 1 | 0 | 0 | 0 | 1 | 0 | 1 | 0 | 0 | 0 | 0 | 0 | 1 | 1 |
| 50.0236 | 19.80693 | MLP_U_4 | 206 | 194 | Poland | Northern | U | MLP | mlpu4_5 | 1 | 1 | 0 | 0 | 0 | 0 | 0 | 0 | | 1 | 0 | 42 | 1 | 42 | 1 | 41887 | 1000 | 3916 | 0 | 37971 | 0 | 1 | 0 | 1 | 0 | 0 | 0 | 0 | 0 | 1 | 0 | 0 | 0 | 0 | 0 | 0 | 1 |
| 50.02509 | 19.81044 | MLP_U_4 | 206 | 194 | Poland | Northern | U | MLP | mlpu4_6 | 0 | 0 | 0 | 0 | 0 | 0 | 0 | | | 0 | 0 | 13 | 1 | 1 | 1 | 27827 | 10300 | 10586 | 1818 | 15423 | 0 | 1 | 0 | 0 | 0 | 0 | 0 | 1 | 0 | 1 | 0 | 0 | 0 | 0 | 0 | 0 | 1 |
| 50.02642 | 19.81396 | MLP_U_4 | 206 | 194 | Poland | Northern | U | MLP | mlpu4_7 | 3 | 1 | 0 | 0 | 0 | 0 | 0 | 0 | | 1 | 0 | 19 | 140 | 48 | 19 | 15604 | 7000 | 3055 | 97 | 12452 | 1 | 1 | 0 | 1 | 0 | 0 | 1 | 0 | 0 | 1 | 0 | 0 | 0 | 0 | 0 | 1 | 0 |
| 50.02414 | 19.81599 | MLP_U_4 | 205 | 194 | Poland | Northern | U | MLP | mlpu4_8 | 2 | 1 | 0 | 0 | 0 | 0 | 0 | 0 | | 1 | 0 | 50 | 13 | 183 | 13 | 10935 | 16900 | 2452 | 4503 | 3981 | 1 | 1 | 0 | 1 | 0 | 0 | 1 | 0 | 0 | 1 | 0 | 0 | 0 | 0 | 0 | 1 | 1 |
| 50.059 | 19.87046 | MLP_U_15 | 226 | 196 | Poland | Northern | U | MLP | mlpu15_1 | 0 | 0 | 0 | 0 | 0 | 0 | 0 | | | 0 | 0 | 272 | 192 | 54 | 54 | 25801 | 11500 | 0 | 6266 | 19536 | 1 | 0 | 1 | 1 | 0 | 0 | 0 | 0 | 1 | 0 | 0 | 1 | 0 | 0 | 0 | 0 | 0 |
| 50.06106 | 19.87081 | MLP_U_15 | 222 | 196 | Poland | Northern | U | MLP | mlpu15_2 | 0 | 0 | 0 | 0 | 0 | 0 | 0 | | | 0 | 0 | 78 | 101 | 104 | 78 | 13585 | 15700 | 3578 | 9407 | 600 | 0 | 1 | 1 | 0 | 0 | 0 | 0 | 1 | 0 | 0 | 1 | 0 | 0 | 0 | 0 | 0 | 1 |
| 50.06326 | 19.87245 | MLP_U_15 | 217 | 196 | Poland | Northern | U | MLP | mlpu15_3 | 3 | 1 | 0 | 0 | 0 | 0 | 0 | 0 | | 1 | 0 | 21 | NA | NA | 21 | 40117 | 11500 | 37872 | 2244 | 0 | 1 | 1 | 1 | 0 | 1 | 0 | 0 | 0 | 1 | 0 | 0 | 1 | 0 | 0 | 0 | 0 | 1 |
| 50.06554 | 19.87322 | MLP_U_15 | 210 | 196 | Poland | Northern | U | MLP | mlpu15_4 | 2 | 1 | 0 | 0 | 0 | 0 | 0 | 0 | | 1 | 0 | 20 | NA | NA | 20 | 37833 | 6200 | 36850 | 983 | 0 | 1 | 1 | 1 | 0 | 1 | 0 | 0 | 0 | 1 | 0 | 0 | 1 | 1 | 0 | 0 | 0 | 1 |
| 50.06683 | 19.87117 | MLP_U_15 | 209 | 196 | Poland | Northern | U | MLP | mlpu15_5 | 1 | 1 | 0 | 0 | 0 | 0 | 0 | 0 | | 1 | 0 | 38 | NA | NA | 38 | 19481 | 17300 | 14677 | 4804 | 0 | 0 | 1 | 1 | 0 | 1 | 0 | 0 | 1 | 1 | 0 | 0 | 1 | 1 | 0 | 0 | 0 | 0 |
| 50.06685 | 19.86703 | MLP_U_15 | 223 | 196 | Poland | Northern | U | MLP | mlpu15_6 | 0 | 0 | 0 | 0 | 0 | 0 | 0 | | | 0 | 0 | 2 | NA | NA | 2 | 22178 | 18500 | 12748 | 9430 | 0 | 0 | 1 | 1 | 0 | 0 | 0 | 0 | 1 | 1 | 0 | 0 | 1 | 1 | 0 | 0 | 0 | 1 |
| 50.06683 | 19.86276 | MLP_U_15 | 221 | 196 | Poland | Northern | U | MLP | mlpu15_7 | 1 | 1 | 0 | 0 | 1 | 1 | 0 | 0 | | 1 | 0 | 10 | 114 | 256 | 10 | 20669 | 17700 | 18361 | 2308 | 0 | 1 | 1 | 1 | 0 | 0 | 0 | 1 | 1 | 1 | 0 | 0 | 1 | 1 | 1 | 0 | 1 | 0 |
| 50.06415 | 19.86143 | MLP_U_15 | 229 | 196 | Poland | Northern | U | MLP | mlpu15_8 | 2 | 1 | 0 | 0 | 0 | 0 | 0 | 0 | | 1 | 0 | 90 | 120 | 87 | 87 | 30937 | 9100 | 11993 | 0 | 18943 | 1 | 1 | 1 | 1 | 0 | 0 | 0 | 1 | 1 | 1 | 0 | 1 | 0 | 0 | 0 | 1 | 0 |
| 50.09554 | 20.00582 | MLP_U_16 | 246 | 203 | Poland | Northern | U | MLP | mlpu16_1 | 1 | 1 | 0 | 0 | 0 | 0 | 0 | 0 | | 1 | 0 | NA | 1 | NA | 1 | 38316 | 10800 | 0 | 0 | 38316 | 1 | 1 | 0 | 1 | 1 | 0 | 0 | 1 | 0 | 0 | 0 | 0 | 1 | 0 | 0 | 1 | 0 |
| 50.09778 | 20.00646 | MLP_U_16 | 248 | 203 | Poland | Northern | U | MLP | mlpu16_2 | 2 | 1 | 0 | 0 | 0 | 0 | 0 | 0 | | 1 | 0 | 208 | 11 | 2 | 2 | 43660 | 14200 | 2621 | 2825 | 38215 | 1 | 1 | 0 | 1 | 0 | 0 | 0 | 1 | 0 | 0 | 0 | 0 | 0 | 1 | 1 | 1 | 0 |
| 50.09937 | 20.00428 | MLP_U_16 | 257 | 203 | Poland | Northern | U | MLP | mlpu16_3 | 1 | 1 | 1 | 1 | 1 | 1 | 1 | 1 | 1 | 0 | 0 | 10 | 100 | 1 | 1 | 32184 | 16300 | 4599 | 2039 | 25546 | 1 | 1 | 0 | 1 | 0 | 0 | 0 | 1 | 1 | 0 | 0 | 0 | 1 | 1 | 0 | 0 | 0 |
| 50.10126 | 20.00167 | MLP_U_16 | 271 | 203 | Poland | Northern | U | MLP | mlpu16_4 | 2 | 1 | 0 | 0 | 0 | 0 | 0 | 0 | | 1 | 0 | 7 | 1 | 148 | 1 | 43949 | 4400 | 29625 | 3590 | 10734 | 1 | 1 | 0 | 0 | 1 | 0 | 0 | 1 | 1 | 0 | 0 | 1 | 1 | 0 | 0 | 0 | 1 |
| 50.10064 | 19.99708 | MLP_U_16 | 275 | 203 | Poland | Northern | U | MLP | mlpu16_5 | 3 | 1 | 1 | 1 | 0 | 0 | 1 | 1 | 1 | 0 | 0 | 8 | 58 | 216 | 8 | 46984 | 600 | 40270 | 6714 | 0 | 1 | 1 | 0 | 0 | 1 | 0 | 0 | 1 | 1 | 0 | 0 | 1 | 0 | 0 | 0 | 1 | 1 |
| 50.09911 | 19.99404 | MLP_U_16 | 259 | 203 | Poland | Northern | U | MLP | mlpu16_6 | 1 | 1 | 0 | 0 | 0 | 0 | 0 | 0 | | 1 | 0 | 1 | 150 | NA | 1 | 25931 | 4800 | 25470 | 461 | 0 | 0 | 1 | 0 | 0 | 1 | 0 | 0 | 1 | 0 | 0 | 0 | 0 | 1 | 0 | 1 | 1 | 0 |
| 50.09831 | 19.98966 | MLP_U_16 | 227 | 203 | Poland | Northern | U | MLP | mlpu16_7 | 1 | 1 | 0 | 0 | 0 | 0 | 0 | 0 | | 1 | 0 | NA | NA | NA | | 24569 | 15900 | 24569 | 0 | 0 | 0 | 1 | 0 | 0 | 0 | 0 | 0 | 1 | 0 | 0 | 0 | 0 | 1 | 0 | 0 | 1 | 1 |
| 50.09746 | 19.98539 | MLP_U_16 | 224 | 203 | Poland | Northern | U | MLP | mlpu16_8 | 1 | 1 | 0 | 0 | 0 | 0 | 0 | 0 | | 1 | 0 | NA | NA | NA | | 22179 | 8300 | 22179 | 0 | 0 | 0 | 1 | 0 | 0 | 1 | 0 | 0 | 1 | 0 | 0 | 0 | 0 | 1 | 0 | 0 | 0 | 1 |
| 50.2749 | 19.56687 | MLP_S_14 | 356 | 167 | Poland | Northern | S | MLP | mlps14_1 | 0 | 0 | 0 | 0 | 0 | 0 | 0 | | | 0 | 0 | 33 | NA | NA | 33 | 18119 | 16090 | 16242 | 1877 | 0 | 1 | 1 | 1 | 0 | 1 | 0 | 0 | 1 | 1 | 1 | 1 | 1 | 0 | 0 | 1 | 0 | 0 |
| 50.27645 | 19.56909 | MLP_S_14 | 366 | 167 | Poland | Northern | S | MLP | mlps14_2 | 1 | 1 | 0 | 0 | 0 | 0 | 0 | 0 | | 1 | 0 | 191 | NA | NA | 191 | 21152 | 14328 | 21152 | 0 | 0 | 1 | 1 | 0 | 0 | 1 | 1 | 0 | 1 | 0 | 0 | 0 | 1 | 1 | 1 | 0 | 0 | 1 |
| 50.27833 | 19.56716 | MLP_S_14 | 368 | 167 | Poland | Northern | S | MLP | mlps14_3 | 0 | 0 | 0 | 0 | 0 | 0 | 0 | | | 0 | 0 | NA | NA | NA | | 11859 | 19091 | 11859 | 0 | 0 | 0 | 1 | 1 | 0 | 0 | 0 | 0 | 1 | 0 | 0 | 0 | 0 | 1 | 1 | 0 | 1 | 0 |
| 50.27965 | 19.56384 | MLP_S_14 | 367 | 167 | Poland | Northern | S | MLP | mlps14_4 | 0 | 0 | 0 | 0 | 0 | 0 | 0 | | | 0 | 0 | 121 | NA | NA | 121 | 5713 | 21969 | 5713 | 0 | 0 | 0 | 1 | 1 | 0 | 0 | 0 | 0 | 1 | 0 | 0 | 0 | 0 | 0 | 1 | 1 | 0 | 0 |
| 50.28039 | 19.55976 | MLP_S_14 | 366 | 167 | Poland | Northern | S | MLP | mlps14_5 | 0 | 0 | 0 | 0 | 0 | 0 | 0 | | | 0 | 0 | 126 | NA | NA | 126 | 9209 | 21367 | 9209 | 0 | 0 | 1 | 1 | 1 | 0 | 1 | 0 | 0 | 1 | 0 | 0 | 0 | 0 | 0 | 1 | 1 | 0 | 0 |
| 50.27917 | 19.55739 | MLP_S_14 | 363 | 167 | Poland | Northern | S | MLP | mlps14_6 | 0 | 0 | 0 | 0 | 0 | 0 | 0 | | | 0 | 0 | 35 | 207 | 201 | 35 | 20601 | 20282 | 19388 | 1213 | 0 | 1 | 1 | 1 | 0 | 1 | 0 | 0 | 1 | 1 | 0 | 0 | 1 | 1 | 1 | 0 | 1 | 1 |
| 50.27746 | 19.55929 | MLP_S_14 | 361 | 167 | Poland | Northern | S | MLP | mlps14_7 | 0 | 0 | 0 | 0 | 0 | 0 | 0 | | | 0 | 0 | 47 | 280 | 156 | 47 | 7320 | 19434 | 4138 | 3182 | 0 | 1 | 1 | 0 | 0 | 0 | 0 | 0 | 1 | 1 | 0 | 0 | 1 | 0 | 1 | 0 | 0 | 0 |
| 50.275 | 19.55797 | MLP_S_14 | 351 | 167 | Poland | Northern | S | MLP | mlps14_8 | 0 | 0 | 0 | 0 | 0 | 0 | 0 | | | 0 | 0 | 25 | 50 | 172 | 25 | 25686 | 3184 | 23348 | 2337 | 0 | 1 | 1 | 1 | 0 | 1 | 0 | 1 | 1 | 1 | 1 | 1 | 1 | 0 | 0 | 1 | 0 | 1 |
| 50.22979 | 19.77411 | MLP_R_14 | 467 | 182 | Poland | Northern | R | MLP | mlpr14_1 | 1 | 1 | 0 | 0 | 0 | 0 | 0 | 0 | | 1 | 0 | 113 | 42 | 52 | 42 | 6480 | 3517 | 1704 | 2381 | 2396 | 0 | 1 | 1 | 0 | 0 | 1 | 0 | 0 | 1 | 0 | 0 | 1 | 0 | 0 | 0 | 0 | 1 |
| 50.22882 | 19.76968 | MLP_R_14 | 451 | 182 | Poland | Northern | R | MLP | mlpr14_2 | 0 | 0 | 0 | 0 | 0 | 0 | 0 | | | 0 | 0 | 17 | 73 | 41 | 17 | 7886 | 4037 | 4218 | 3668 | 0 | 0 | 1 | 1 | 0 | 0 | 0 | 0 | 0 | 1 | 0 | 1 | 1 | 0 | 0 | 0 | 0 | 1 |
| 50.22647 | 19.76798 | MLP_R_14 | 426 | 182 | Poland | Northern | R | MLP | mlpr14_3 | 1 | 1 | 0 | 0 | 0 | 0 | 0 | 0 | | 1 | 0 | 17 | 3 | 1 | 1 | 19507 | 5519 | 7472 | 8514 | 3520 | 1 | 1 | 1 | 1 | 0 | 0 | 1 | 0 | 1 | 0 | 0 | 1 | 0 | 0 | 0 | 0 | 1 |
| 50.22417 | 19.7701 | MLP_R_14 | 417 | 182 | Poland | Northern | R | MLP | mlpr14_4 | 4 | 1 | 0 | 0 | 0 | 0 | 0 | 0 | | 1 | 0 | 162 | 82 | 1 | 1 | 28923 | 4086 | 0 | 3408 | 25515 | 1 | 1 | 1 | 1 | 0 | 0 | 0 | 0 | 1 | 0 | 1 | 1 | 0 | 0 | 0 | 0 | 1 |
| 50.22184 | 19.77234 | MLP_R_14 | 409 | 182 | Poland | Northern | R | MLP | mlpr14_5 | 1 | 1 | 0 | 0 | 0 | 0 | 0 | 0 | | 1 | 0 | 16 | 20 | 28 | 16 | 30750 | 3145 | 1682 | 1737 | 27331 | 1 | 1 | 1 | 1 | 0 | 0 | 0 | 1 | 1 | 0 | 1 | 0 | 0 | 0 | 0 | 1 | 1 |
| 50.22 | 19.77549 | MLP_R_14 | 402 | 182 | Poland | Northern | R | MLP | mlpr14_6 | 1 | 1 | 0 | 0 | 0 | 0 | 0 | 0 | | 1 | 0 | 1 | 1 | 33 | 1 | 29160 | 3121 | 2159 | 1244 | 25757 | 1 | 1 | 1 | 1 | 0 | 0 | 0 | 1 | 1 | 0 | 1 | 1 | 0 | 0 | 0 | 1 | 1 |
| 50.2185 | 19.77855 | MLP_R_14 | 396 | 182 | Poland | Northern | R | MLP | mlpr14_7 | 1 | 1 | 0 | 0 | 0 | 0 | 0 | 0 | | 1 | 0 | 17 | 1 | 62 | 1 | 42573 | 1118 | 1331 | 911 | 40330 | 1 | 1 | 1 | 1 | 0 | 0 | 0 | 1 | 1 | 0 | 1 | 0 | 0 | 0 | 0 | 1 | 1 |
| 50.21749 | 19.7822 | MLP_R_14 | 391 | 182 | Poland | Northern | R | MLP | mlpr14_8 | 1 | 1 | 0 | 0 | 0 | 0 | 0 | 0 | | 1 | 0 | NA | 104 | 66 | 66 | 43453 | 0 | 0 | 0 | 43453 | 1 | 0 | 1 | 1 | 0 | 0 | 0 | 0 | 0 | 1 | 0 | 0 | 0 | 0 | 0 | 1 | 0 |
| 49.6757 | 20.07198 | MLP_S_15 | 394 | 231 | Poland | Northern | S | MLP | mlps15_1 | 0 | 0 | 0 | 0 | 0 | 0 | 0 | | | 0 | 0 | 17 | 100 | 84 | 17 | 8241 | 13659 | 0 | 4808 | 3433 | 1 | 1 | 0 | 0 | 0 | 0 | 0 | 1 | 1 | 0 | 0 | 1 | 0 | 0 | 1 | 0 | 0 |
| 49.67398 | 20.07191 | MLP_S_15 | 401 | 231 | Poland | Northern | S | MLP | mlps15_2 | 1 | 1 | 0 | 0 | 0 | 0 | 0 | 0 | | 1 | 0 | 90 | 42 | 50 | 42 | 17353 | 9095 | 0 | 3358 | 13995 | 1 | 1 | 1 | 1 | 0 | 0 | 1 | 0 | 1 | 0 | 0 | 1 | 0 | 0 | 1 | 0 | 0 |
| 49.67228 | 20.07293 | MLP_S_15 | 430 | 231 | Poland | Northern | S | MLP | mlps15_3 | 0 | 0 | 0 | 0 | 0 | 0 | 0 | | | 0 | 0 | 124 | 182 | 5 | 5 | 8467 | 3890 | 0 | 7376 | 1091 | 1 | 1 | 0 | 1 | 0 | 0 | 0 | 0 | 1 | 0 | 1 | 1 | 0 | 0 | 0 | 0 | 0 |
| 49.67055 | 20.07224 | MLP_S_15 | 444 | 231 | Poland | Northern | S | MLP | mlps15_4 | 0 | 0 | 0 | 0 | 0 | 0 | 0 | | | 0 | 0 | 204 | 232 | 115 | 115 | 2842 | 4868 | 0 | 2842 | 0 | 0 | 1 | 0 | 0 | 0 | 0 | 0 | 0 | 1 | 0 | 1 | 0 | 0 | 0 | 0 | 0 | 0 |
| 49.66909 | 20.07439 | MLP_S_15 | 432 | 231 | Poland | Northern | S | MLP | mlps15_5 | 0 | 0 | 0 | 0 | 0 | 0 | 0 | | | 0 | 0 | 65 | 224 | 61 | 61 | 10158 | 3763 | 0 | 5279 | 4879 | 0 | 1 | 0 | 0 | 0 | 0 | 1 | 0 | 1 | 1 | 1 | 1 | 0 | 0 | 0 | 0 | 0 |
| 49.6688 | 20.0778 | MLP_S_15 | 410 | 231 | Poland | Northern | S | MLP | mlps15_6 | 0 | 0 | 0 | 0 | 0 | 0 | 0 | | | 0 | 0 | 90 | 148 | 291 | 90 | 14644 | 6773 | 4078 | 8050 | 2516 | 0 | 1 | 0 | 0 | 0 | 0 | 0 | 0 | 1 | 1 | 0 | 0 | 1 | 0 | 0 | 0 | 0 |
| 49.66949 | 20.08126 | MLP_S_15 | 405 | 231 | Poland | Northern | S | MLP | mlps15_7 | 0 | 0 | 0 | 0 | 0 | 0 | 0 | | | 0 | 0 | 31 | 193 | 286 | 31 | 20300 | 14701 | 20032 | 268 | 0 | 1 | 1 | 0 | 0 | 1 | 0 | 0 | 1 | 1 | 0 | 0 | 1 | 0 | 1 | 0 | 0 | 0 |
| 49.67187 | 20.08147 | MLP_S_15 | 398 | 231 | Poland | Northern | S | MLP | mlps15_8 | 0 | 0 | 0 | 0 | 0 | 0 | 0 | | | 0 | 0 | 59 | 162 | NA | 59 | 21546 | 2820 | 21546 | 0 | 0 | 0 | 1 | 0 | 0 | 1 | 0 | 0 | 0 | 0 | 0 | 1 | 0 | 0 | 0 | 1 | 0 | 0 |
| 49.74051 | 19.98897 | MLP_R_15 | 346 | 221 | Poland | Northern | R | MLP | mlpr15_1 | 0 | 0 | 0 | 0 | 0 | 0 | 0 | | | 0 | 0 | NA | NA | 154 | 154 | 30709 | 0 | 0 | 0 | 30709 | 1 | 1 | 1 | 1 | 0 | 0 | 0 | 0 | 0 | 1 | 0 | 0 | 0 | 0 | 0 | 0 | 0 |
| 49.74049 | 19.98423 | MLP_R_15 | 338 | 221 | Poland | Northern | R | MLP | mlpr15_2 | 1 | 1 | 0 | 0 | 0 | 0 | 0 | 0 | | 1 | 0 | NA | NA | 58 | 58 | 16328 | 2781 | 0 | 833 | 15494 | 1 | 0 | 0 | 1 | 0 | 0 | 0 | 0 | 1 | 1 | 1 | 1 | 0 | 0 | 0 | 0 | 0 |
| 49.7404 | 19.98105 | MLP_R_15 | 332 | 221 | Poland | Northern | R | MLP | mlpr15_3 | 0 | 0 | 0 | 0 | 0 | 0 | 0 | | | 0 | 0 | NA | NA | 35 | 35 | 29249 | 2535 | 0 | 1513 | 27736 | 1 | 0 | 0 | 1 | 0 | 0 | 0 | 0 | 1 | 1 | 0 | 1 | 0 | 0 | 0 | 0 | 1 |
| 49.73796 | 19.98131 | MLP_R_15 | 332 | 221 | Poland | Northern | R | MLP | mlpr15_4 | 0 | 0 | 0 | 0 | 0 | 0 | 0 | | | 0 | 0 | NA | NA | 17 | 17 | 34016 | 1353 | 0 | 1269 | 32748 | 1 | 0 | 0 | 1 | 0 | 0 | 0 | 0 | 1 | 1 | 1 | 1 | 0 | 0 | 0 | 0 | 0 |

| | | | | | | | | | | | | | | | | | | | | | | | | | | | | | | | | | | | | | | | | | | | | | | |
|---|---|---|---|---|---|---|---|---|---|---|---|---|---|---|---|---|---|---|---|---|---|---|---|---|---|---|---|---|---|---|---|---|---|---|---|---|---|---|---|---|---|---|---|---|---|---|
| 49.73558 | 19.98256 | MLP_R_15 | 335 | 221 | Poland | Northern | R | MLP | mlpr15_5 | 0 | 0 | 0 | 0 | 0 | 0 | 0 | | | 0 | 0 | NA | NA | 61 | 61 | 25573 | 2547 | 0 | 3747 | 21826 | 1 | 1 | 0 | 1 | 0 | 0 | 0 | 0 | 1 | 0 | 1 | 1 | 0 | 0 | 0 | 0 | 0 |
| 49.73373 | 19.97984 | MLP_R_15 | 334 | 221 | Poland | Northern | R | MLP | mlpr15_6 | 2 | 1 | 0 | 0 | 0 | 0 | 0 | 0 | | 1 | 0 | NA | NA | 144 | 144 | 36867 | 0 | 0 | 0 | 36867 | 1 | 1 | 0 | 1 | 0 | 0 | 0 | 0 | 1 | 1 | 0 | 0 | 0 | 0 | 0 | 0 | 1 |
| 49.73238 | 19.9837 | MLP_R_15 | 339 | 221 | Poland | Northern | R | MLP | mlpr15_7 | 0 | 0 | 0 | 0 | 0 | 0 | 0 | | | 0 | 0 | NA | NA | 45 | 45 | 20761 | 1158 | 0 | 3505 | 17256 | 1 | 1 | 1 | 1 | 0 | 0 | 0 | 0 | 1 | 0 | 1 | 1 | 0 | 0 | 0 | 0 | 0 |
| 49.73158 | 19.98746 | MLP_R_15 | 356 | 221 | Poland | Northern | R | MLP | mlpr15_8 | 1 | 1 | 0 | 0 | 0 | 0 | 0 | 0 | | 1 | 0 | NA | NA | 51 | 51 | 38179 | 2066 | 0 | 841 | 37338 | 1 | 1 | 1 | 1 | 0 | 0 | 0 | 0 | 1 | 0 | 0 | 1 | 0 | 0 | 0 | 0 | 0 |
| 49.97209 | 20.60693 | MLP_S_16 | 228 | 216 | Poland | Northern | S | MLP | mlps16_1 | 1 | 1 | 1 | 1 | 1 | 1 | 1 | 1 | 1 | 0 | 0 | 100 | NA | NA | 100 | 30531 | 7690 | 28236 | 2295 | 0 | 0 | 1 | 0 | 0 | 0 | 1 | 0 | 1 | 1 | 0 | 1 | 0 | 1 | 0 | 1 | 0 | 1 |
| 49.97222 | 20.61051 | MLP_S_16 | 225 | 216 | Poland | Northern | S | MLP | mlps16_2 | 0 | 0 | 2 | 1 | 0 | 0 | 0 | | 0 | 0 | 1 | 116 | NA | NA | 116 | 10627 | 14180 | 8369 | 2258 | 0 | 0 | 1 | 1 | 0 | 0 | 0 | 0 | 1 | 1 | 0 | 0 | 1 | 1 | 0 | 0 | 0 | 1 |
| 49.97011 | 20.61026 | MLP_S_16 | 221 | 216 | Poland | Northern | S | MLP | mlps16_3 | 0 | 0 | 1 | 1 | 0 | 0 | 0 | | 0 | 0 | 1 | 76 | 239 | 411 | 76 | 17911 | 12605 | 5246 | 2065 | 10599 | 0 | 1 | 0 | 0 | 1 | 0 | 0 | 1 | 1 | 0 | 0 | 1 | 1 | 1 | 0 | 0 | 0 |
| 49.9681 | 20.60866 | MLP_S_16 | 225 | 216 | Poland | Northern | S | MLP | mlps16_4 | 1 | 1 | 0 | 0 | 0 | 0 | 0 | 0 | | 1 | 0 | 311 | 61 | 181 | 61 | 21702 | 21205 | 1481 | 629 | 19592 | 1 | 0 | 0 | 1 | 0 | 0 | 0 | 1 | 1 | 1 | 0 | 0 | 0 | 1 | 0 | 0 | 0 |
| 49.96607 | 20.60645 | MLP_S_16 | 223 | 216 | Poland | Northern | S | MLP | mlps16_5 | 0 | 0 | 0 | 0 | 0 | 0 | 0 | | | 0 | 0 | 168 | 64 | 46 | 46 | 7549 | 18609 | 1382 | 2142 | 4026 | 1 | 1 | 0 | 1 | 0 | 0 | 0 | 1 | 1 | 1 | 0 | 0 | 0 | 1 | 0 | 0 | 0 |
| 49.96458 | 20.60293 | MLP_S_16 | 223 | 216 | Poland | Northern | S | MLP | mlps16_6 | 0 | 0 | 0 | 0 | 0 | 0 | 0 | | | 0 | 0 | 80 | 157 | 108 | 80 | 13041 | 11471 | 11203 | 1838 | 0 | 1 | 1 | 0 | 1 | 0 | 0 | 0 | 1 | 1 | 0 | 0 | 1 | 1 | 0 | 1 | 0 | 0 |
| 49.96228 | 20.60117 | MLP_S_16 | 224 | 216 | Poland | Northern | S | MLP | mlps16_7 | 1 | 1 | 0 | 0 | 0 | 0 | 0 | 0 | | 1 | 0 | 37 | 94 | 107 | 37 | 27110 | 8438 | 25444 | 1666 | 0 | 1 | 1 | 0 | 1 | 0 | 0 | 0 | 1 | 1 | 0 | 1 | 0 | 1 | 0 | 0 | 0 | 1 |
| 49.95977 | 20.60172 | MLP_S_16 | 227 | 216 | Poland | Northern | S | MLP | mlps16_8 | 2 | 1 | 0 | 0 | 0 | 0 | 0 | 0 | | 1 | 0 | 133 | 20 | 78 | 20 | 32744 | 21331 | 32744 | 0 | 0 | 1 | 0 | 0 | 0 | 1 | 0 | 0 | 0 | 0 | 1 | 0 | 0 | 0 | 1 | 1 | 0 | 1 |
| 49.96767 | 20.51131 | MLP_R_16 | 227 | 243 | Poland | Northern | R | MLP | mlpr16_1 | 0 | 0 | 0 | 0 | 0 | 0 | 0 | | | 0 | 0 | 30 | 138 | 82 | 30 | 28317 | 3833 | 1949 | 2073 | 24294 | 0 | 1 | 0 | 1 | 0 | 0 | 1 | 0 | 1 | 0 | 1 | 0 | 0 | 0 | 0 | 1 | 0 |
| 49.96781 | 20.50778 | MLP_R_16 | 215 | 243 | Poland | Northern | R | MLP | mlpr16_2 | 2 | 1 | 0 | 0 | 0 | 0 | 0 | 0 | | 1 | 0 | 200 | 198 | 50 | 50 | 15705 | 2214 | 0 | 1397 | 14308 | 1 | 1 | 0 | 1 | 0 | 0 | 1 | 0 | 1 | 1 | 1 | 0 | 0 | 0 | 0 | 0 | 0 |
| 49.96768 | 20.50361 | MLP_R_16 | 219 | 243 | Poland | Northern | R | MLP | mlpr16_3 | 0 | 0 | 0 | 0 | 0 | 0 | 0 | | | 0 | 0 | NA | NA | 60 | 60 | 14304 | 747 | 0 | 278 | 14026 | 1 | 1 | 0 | 1 | 0 | 0 | 0 | 0 | 1 | 1 | 1 | 0 | 0 | 0 | 0 | 0 | 0 |
| 49.96923 | 20.50575 | MLP_R_16 | 233 | 243 | Poland | Northern | R | MLP | mlpr16_4 | 0 | 0 | 0 | 0 | 0 | 0 | 0 | | | 0 | 0 | NA | NA | 8 | 8 | 19890 | 2842 | 0 | 2797 | 17093 | 1 | 1 | 0 | 1 | 0 | 0 | 0 | 0 | 1 | 1 | 1 | 0 | 0 | 0 | 0 | 0 | 0 |
| 49.97036 | 20.50948 | MLP_R_16 | 219 | 243 | Poland | Northern | R | MLP | mlpr16_5 | 0 | 0 | 0 | 0 | 0 | 0 | 0 | | | 0 | 0 | NA | NA | 25 | 25 | 9816 | 3176 | 0 | 2914 | 6902 | 1 | 1 | 0 | 1 | 0 | 0 | 0 | 0 | 1 | 1 | 1 | 0 | 0 | 0 | 0 | 0 | 0 |
| 49.9718 | 20.51243 | MLP_R_16 | 223 | 243 | Poland | Northern | R | MLP | mlpr16_6 | 0 | 0 | 1 | 1 | 0 | 0 | 0 | | 0 | 0 | 1 | NA | NA | 16 | 16 | 29572 | 3270 | 0 | 2514 | 27058 | 1 | 1 | 0 | 1 | 0 | 0 | 0 | 0 | 1 | 1 | 1 | 0 | 1 | 0 | 0 | 0 | 1 |
| 49.97182 | 20.51614 | MLP_R_16 | 214 | 243 | Poland | Northern | R | MLP | mlpr16_7 | 0 | 0 | 0 | 0 | 0 | 0 | 0 | | | 0 | 0 | NA | NA | 36 | 36 | 16999 | 3457 | 0 | 726 | 16273 | 1 | 1 | 0 | 1 | 0 | 0 | 0 | 0 | 1 | 1 | 1 | 0 | 0 | 0 | 0 | 0 | 1 |
| 49.97332 | 20.51955 | MLP_R_16 | 213 | 243 | Poland | Northern | R | MLP | mlpr16_8 | 1 | 1 | 0 | 0 | 0 | 0 | 0 | 0 | | 1 | 0 | NA | NA | 40 | 40 | 18142 | 1936 | 0 | 3821 | 14322 | 1 | 1 | 0 | 1 | 0 | 0 | 0 | 1 | 1 | 1 | 1 | 0 | 0 | 0 | 0 | 0 | 1 |

**Table S5**. Generalised Linear Mixed Models demonstrating the abundance relationships of Great Spotted Woodpecker, Syrian Woodpecker and their hybrids to a mixture of particular habitat characteristic elements: Canopy cover area, Infrastructure area, Dense canopy, Loose canopy, Coniferous trees, Forest patch, Urban park, Graveyard, Alleys, Mid-estate wood, Orchard, Riparian wood, Buildings small sporadic, Buildings small dense, Buildings tall sporadic, Buildings tall dense and Other infrastructure. European scale. Significant p values (<0.05) highlighted in bold.

| Predictors | Estimate | Std. Error | z-value | p |
|---|---|---|---|---|
| Great Spotted Woodpecker | | | | |
| Intercept | -1.598e+00 | 2.382e-01 | -6.709 | **1.96e-11** |
| Canopy cover area | 1.756e-05 | 3.864e-06 | 4.545 | **5.50e-06** |
| Infrastructure area | -1.405e-05 | 1.293e-05 | -1.087 | 0.277 |
| Dense canopy | 2.914e-01 | 1.463e-01 | 1.991 | **0.046** |
| Loose canopy | -7.548e-02 | 1.448e-01 | -0.521 | 0.602 |
| Coniferous trees | 2.100e-01 | 1.226e-01 | 1.713 | 0.087 |
| Forest patch | 2.232e-01 | 1.429e-01 | 1.562 | 0.118 |
| Urban Park | 1.788e-01 | 1.388e-01 | 1.288 | 0.198 |
| Graveyard | 2.126e-01 | 2.181e-01 | 0.975 | 0.330 |
| Alleys | -9.528e-02 | 1.038e-01 | -0.918 | 0.359 |
| Mid-estate wood | 3.122e-02 | 1.202e-01 | 0.260 | 0.795 |
| Orchard | 4.629e-02 | 1.185e-01 | 0.391 | 0.696 |
| Riparian wood | 3.135e-01 | 1.094e-01 | 2.865 | **0.004** |
| Buildings small sporadic | 8.274e-02 | 1.061e-01 | 0.780 | 0.435 |
| Buildings small dense | -1.952e-01 | 1.262e-01 | -1.546 | 0.122 |
| Buildings tall sporadic | 6.249e-02 | 1.282e-01 | 0.488 | 0.626 |
| Buildings tall dense | 2.011e-01 | 1.692e-01 | 1.189 | 0.235 |
| Other infrastructure | -5.513e-02 | 1.365e-01 | -0.404 | 0.686 |
| Syrian Woodpecker | | | | |
| Intercept | -2.003e+00 | 4.177e-01 | -4.795 | **1.63e-06** |
| Canopy cover area | -5.171e-06 | 7.105e-06 | -0.728 | 0.467 |
| Infrastructure area | -1.939e-05 | 2.030e-05 | -0.955 | 0.339 |
| Dense canopy | 6.973e-02 | 2.027e-01 | 0.344 | 0.731 |
| Loose canopy | -1.442e-01 | 3.070e-01 | -0.470 | 0.639 |
| Coniferous trees | -4.166e-01 | 2.353e-01 | -1.771 | 0.077 |
| Forest patch | 1.082e-01 | 2.188e-01 | 0.495 | 0.621 |
| Urban Park | 1.538e-01 | 2.301e-01 | 0.668 | 0.504 |
| Graveyard | 4.428e-01 | 3.799e-01 | 1.166 | 0.244 |
| Alleys | 1.716e-01 | 1.686e-01 | 1.018 | 0.309 |
| Mid-estate wood | 1.264e-01 | 1.941e-01 | 0.651 | 0.515 |
| Orchard | 8.135e-02 | 2.086e-01 | 0.390 | 0.697 |
| Riparian wood | -1.221e-01 | 1.867e-01 | -0.654 | 0.513 |
| Buildings small sporadic | -2.020e-01 | 1.903e-01 | -1.062 | 0.288 |
| Buildings small dense | 4.136e-01 | 2.072e-01 | 1.997 | **0.046** |
| Buildings tall sporadic | -3.081e-02 | 2.226e-01 | -0.138 | 0.890 |
| Buildings tall dense | -1.246e-01 | 2.974e-01 | -0.419 | 0.675 |
| Other infrastructure | -3.066e-01 | 2.378e-01 | -1.290 | 0.197 |

| Hybrid | | | | |
|---|---|---|---|---|
| Intercept | -4.905e+00 | 9.547e-01 | -5.138 | 2.78e-07 |
| Canopy cover area | -8.292e-07 | 1.651e-05 | -0.050 | 0.960 |
| Infrastructure area | 5.247e-05 | 3.616e-05 | 1.451 | 0.147 |
| Dense canopy | 1.072e+00 | 5.961e-01 | 1.799 | 0.072 |
| Loose canopy | -4.855e-02 | 6.566e-01 | -0.074 | 0.941 |
| Coniferous trees | 1.847e-01 | 4.609e-01 | 0.401 | 0.689 |
| Forest patch | -2.745e-01 | 5.396e-01 | -0.509 | 0.611 |
| Urban Park | 9.085e-02 | 5.655e-01 | 0.161 | 0.872 |
| Graveyard | 3.158e-01 | 9.994e-01 | 0.316 | 0.752 |
| Alleys | 5.601e-01 | 3.769e-01 | 1.486 | 0.137 |
| Mid-estate wood | 1.558e-01 | 4.789e-01 | 0.325 | 0.745 |
| Orchard | 3.672e-01 | 4.792e-01 | 0.766 | 0.444 |
| Riparian wood | -3.453e-01 | 4.713e-01 | -0.733 | 0.464 |
| Buildings small sporadic | -6.015e-01 | 4.855e-01 | -1.239 | 0.215 |
| Buildings small dense | -6.300e-02 | 4.884e-01 | -0.129 | 0.897 |
| Buildings tall sporadic | -5.579e-01 | 5.057e-01 | -1.103 | 0.270 |
| Buildings tall dense | -2.758e-01 | 6.893e-01 | -0.400 | 0.689 |
| Other infrastructure | -1.113e+00 | 7.240e-01 | -1.537 | 0.124 |
